# HOOK3 is a scaffold for the opposite-polarity microtubule-based motors cytoplasmic dynein and KIF1C

**DOI:** 10.1101/508887

**Authors:** Agnieszka A. Kendrick, William B. Redwine, Phuoc Tien Tran, Laura Pontano Vaites, Monika Dzieciatkowska, J. Wade Harper, Samara L. Reck-Peterson

**Affiliations:** Department of Cellular and Molecular Medicine, University of California San Diego, La Jolla, CA, 92093.; Department of Cell Biology, Harvard Medical School, Boston, MA 02115.; Section of Cell and Developmental Biology, Division of Biological Sciences, University of California San Diego, La Jolla, CA 92093.; Department of Biochemistry and Molecular Genetics, University of Colorado Denver, Aurora, CO 80045.; Howard Hughes Medical Institute.

**Author notes:** Present address: Stowers Institute for Medical Research, Kansas City, MO 64110. Present address: Department of Molecular and Cellular Biology, Harvard University, Cambridge, MA 02138. Correspondence to: Samara Reck-Peterson, 9500 Gilman Drive, Leichtag 482, La Jolla CA, 92093,; https://orcid.org/0000-0002-1553-465X.

## Abstract

The unidirectional and opposite-polarity microtubule-based motors, dynein and kinesin, drive long-distance intracellular cargo transport. Cellular observations support the existence of mechanisms to couple opposite polarity motors: in cells some cargos rapidly switch directions and kinesin motors can be used to localize dynein. We recently identified an interaction between the cytoplasmic dynein-1 activating adaptor HOOK3 and the kinesin-3 KIF1C. Here we show that KIF1C and dynein/dynactin can exist in a single complex scaffolded by HOOK3. Full-length HOOK3 binds to and activates dynein/dynactin motility. HOOK3 also binds to a short region in the “tail” of KIF1C, but unlike dynein/dynactin, this interaction does not affect the processive motility of KIF1C. HOOK3 scaffolding allows dynein to transport KIF1C towards the microtubule minus end, and KIF1C to transport dynein towards the microtubule plus end. We propose that linking dynein and kinesin motors by dynein activating adaptors may be a general mechanism to regulate bidirectional motility.

## Introduction

In many eukaryotic organisms microtubules and the motors that move on them, kinesins and dynein, power the long distance transport of intracellular cargos. Microtubules are polar structures with their “minus ends” typically located near microtubule organizing centers. Cytoplasmic dynein-1 (“dynein” here) moves cargos towards the microtubule minus end, while kinesins that transport cargos over long distances, such as those in the kinesin-1, −2 and −3 families, move cargos towards the microtubule plus end (Vale, 2003). The cargos of these motors include organelles, other membrane-bound compartments, and large RNA and protein complexes (Reck-Peterson et al., 2018).

In many cases, these cargos can be observed rapidly switching directions. For example, in filamentous fungi endosomes move bidirectionally along microtubules (Abenza et al., 2009; Egan et al., 2012; Wedlich-Söldner et al., 2002) and also drive the bidirectional motility of hitchhiking cargos such as peroxisomes, lipid droplets, endoplasmic reticulum, and ribonucleoprotein complexes (Baumann et al., 2012; Guimaraes et al., 2015; Salogiannis et al., 2016). In human cells, examples of cargos that move bidirectionally on microtubules include lysosomes (Hendricks et al., 2010), secretory vesicles (Barkus et al., 2008; Schlager et al., 2010), autophagosomes (Maday et al., 2012), and protein aggregates (Encalada et al., 2011; Kamal et al., 2000). Purified cargos, such as pigment granules (Rogers et al., 1997) and neuronal transport vesicles (Hendricks et al., 2010) exhibit bidirectional motility along microtubules in vitro. Together, these data suggest that opposite polarity motors are present on the same cargos in many organisms and for many cargo types. There is also evidence that kinesin localizes dynein to microtubule plus ends (Carvalho et al., 2004; Twelvetrees et al., 2016; Zhang et al., 2003), suggesting that these motors could be directly coupled. Given these data, how are teams of opposite polarity motors scaffolded and what controls their directionality?

To answer these questions, we and others have taken a “bottom-up” approach to design artificial scaffolds bearing opposite-polarity motors. For example, dynein and kinesin motors can be scaffolded by DNA origami (Derr et al., 2012) or short DNA oligomers (Belyy et al., 2016). In these cases, the presence of an opposite polarity motor negatively impacts the forward velocity of the other motor type. Such approaches allow the basic biophysical properties of motor teams to be dissected. However, few studies have been done with both physiological motor pairs and physiological scaffolds, primarily because these scaffolds have not been identified or well characterized. One exception is our recent reconstitution of dynein transport to microtubule plus ends by a kinesin (Roberts et al., 2014), a process that occurs in vivo in yeast cells (Moore et al., 2009). Our reconstitution showed that cytoplasmic dynein-1 and the kinesin Kip2 required two additional proteins for scaffolding and that both motors were regulated so that Kip2-driven plus-end-directed motility of dynein to the “start” of its track (microtubule plus ends) is achieved (DeSantis et al., 2017; Roberts et al., 2014).

How are opposite polarity motors scaffolded in mammalian cells? A group of proteins called “dynein activating adaptors” are emerging as candidate scaffold proteins (Fu and Holzbaur, 2014; Reck-Peterson et al., 2018). Processive dynein motility requires an activating adaptor as well as the dynactin complex (McKenney et al., 2014; Schlager et al., 2014). Examples of activating adaptors are the HOOK (HOOK1, HOOK2, and HOOK3), BICD (BICD1, BICD2, BICDL1, and BICDL2), and ninein (NIN and NINL) families of proteins (Reck-Peterson et al., 2018). Recently, we identified an interaction between HOOK3 and the kinesin KIF1C using a proteomics approach (Redwine et al., 2017). The dynein activating adaptors BICD2 and BICDL1 may also interact with kinesin motors (Novarino et al., 2014; Schlager et al., 2010; Splinter et al., 2010). However, for these interactions between dynein activating adaptors and kinesins, it is not known if the interactions are direct, if dynein and kinesin binding is achieved simultaneously, and if the dynein activating adaptor can scaffold bidirectional motility. Here we set out to understand these fundamental points for the dynein activating adaptor HOOK3, cytoplasmic dynein-1, and KIF1C.

## Results

### Endogenous KIF1C and HOOK3 interact specifically

We originally identified the HOOK3-KIF1C interaction using a proximity-dependent biotinylation technique (BioID) (Redwine et al., 2017; Roux et al., 2012). KIF1C is a kinesin-3 family member that is closely related to KIF1A and KIF1B (Dorner et al., 1998). It contains an amino-terminal motor domain and carboxy-terminal “tail” domain with several regions of predicted coiled-coil, a forkhead-associated domain (FHA), and a proline-rich region that is predicted to be unstructured and is distinct from other kinesin-3 family members (Miki et al., 2005). KIF1C interacts with the carboxy-terminus of HOOK3 (Redwine et al., 2017), while dynein and dynactin interact with its amino-terminus (McKenney et al., 2014) (Fig. 1A).

**Figure 1.**
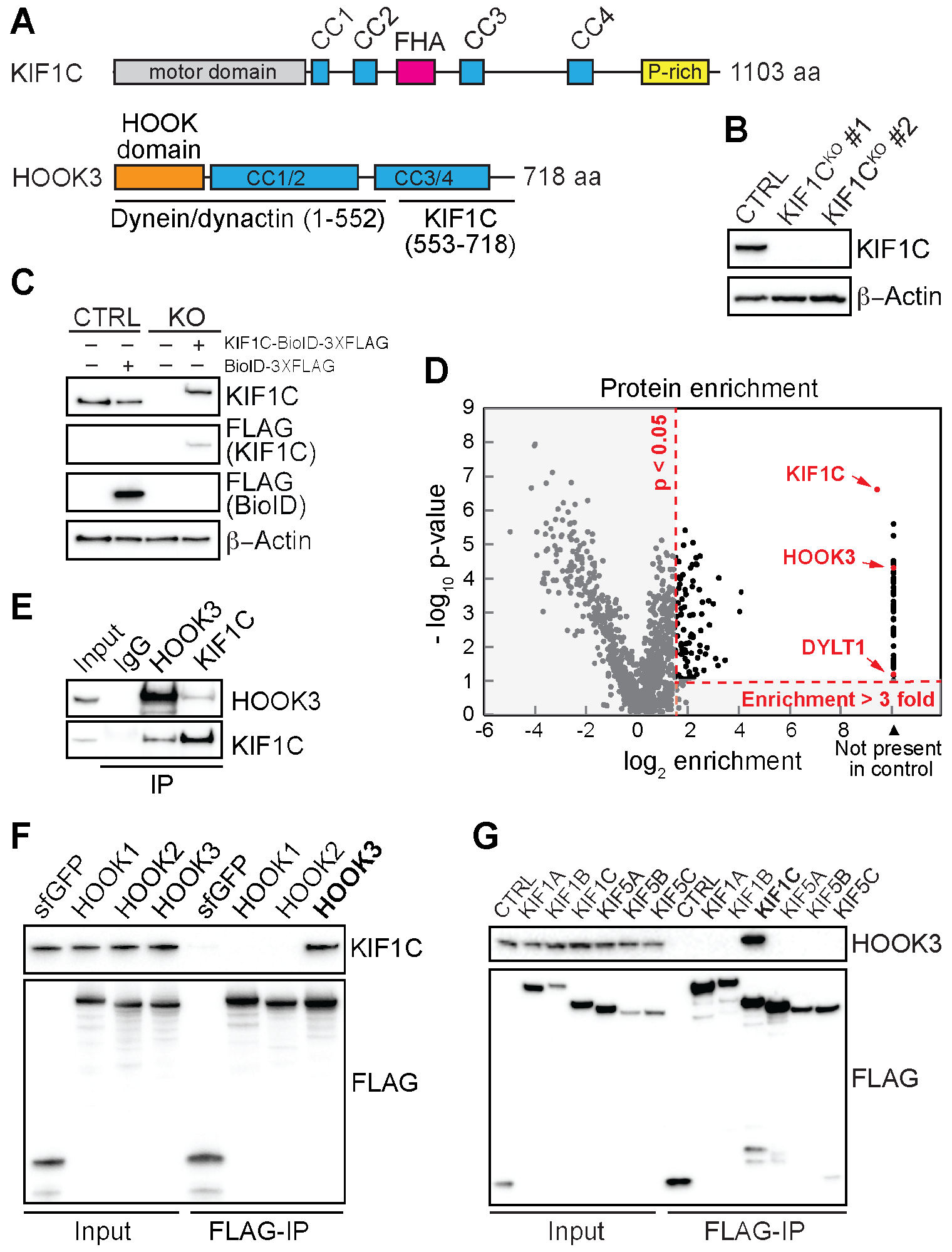
Endogenous HOOK3 and KIF1C interact specifically. **(A)** Domain organization of KIF1C and HOOK3. KIF1C contains an amino-terminal kinesin motor domain and regions of predicted coiled coil (CC), a forkhead-associated domain (FHA) and proline-rich (P-rich) region in its carboxy-terminal “tail”. HOOK3 is largely made up of regions of predicted coiled coil and contains dynein/dynactin and KIF1C binding regions (McKenney et al., 2014; Redwine et al., 2017). The HOOK domain, which is also involved in dynein/dynactin binding (Schroeder and Vale, 2016) is indicated. **(B)** 293T cells were transfected with control CRISPR-Cas9 (CTRL) or with CRISPR-Cas9-gRNA specific for KIF1C. KIF1C knockout (KIF1C^KO^) was confirmed in two different clones by immunoblotting with a KIF1C antibody. Clone #1 was selected for further assays. ß-Actin provided a loading control. **(C)** KIF1C^KO^ cells were infected with viral particles encoding murine stem cell virus (MSCV)-driven KIF1C-BioID-3XFLAG (BioID, promiscuous biotin ligase) plasmid to obtain near-endogenous KIF1C-BioID protein expression levels. Immunoblots were performed using the indicated antibodies. ß-Actin provided a loading control. **(D)** A volcano plot showing enrichment versus significance of proteins identified in KIF1C-BioID experiments relative to control (BioID alone) experiments. A quadrant (dashed red line) bounded by a p-value of 0.05 and 3-fold enrichment contained KIF1C, HOOK3 and DYLT1 (dynein light chain TcTex-1). **(E)** Immunoprecipitation (IP) of endogenous HOOK3 and KIF1C with the indicated antibodies from 293T cells. Immunoblots were performed with HOOK3 or KIF1C antibodies. **(F)** Human HOOK1, HOOK2, and HOOK3 tagged with the HaloTag on the amino-terminus and 3XFLAG at the carboxy-terminus were transiently transfected into 293T cells and immunoprecipitated (IP) with FLAG affinity resin. Immunoblots were performed with KIF1C and FLAG antibodies. 3XFLAG-sfGFP (super folder GFP) provided a control. **(G)** Human KIF1A, KIF1B, KIF1C, KIF5A, KIF5B, and KIF5C were each tagged with BioID-3XFLAG on the carboxy-terminus and stably expressed in 293T cells. Tagged proteins were immunoprecipitated (IP) with FLAG affinity resin and immunoblots were performed with HOOK3 and FLAG antibodies. BioID-3XFLAG provided a control (CTRL).

We began by performing a BioID experiment to identify the protein interactome of KIF1C. Because we wanted to perform this experiment with near-endogenous expression levels of KIF1C (to avoid any artifacts of protein overexpression), we generated KIF1C knockout 293T human cell lines using CRISPR/Cas9-based gene editing. Co-transfection of 293T cells with Cas9 and guides specific for exon 3 of the KIF1C transcript or empty vector, followed by clonal selection yielded two clones with full depletion of KIF1C and a control cell line (Fig. 1B). We then infected one of these cell lines (KIF1C^KO^, clone #1) and the control cell line with retroviral KIF1C-BioID-3XFLAG or BiOID-3XFLAG plasmids driven by the murine stem cell virus (MSCV) promoter to generate stable cells expressing ectopic KIF1C-BioID or BioID alone (Behrends et al., 2010). The KIF1C protein expression levels in these cells were similar to endogenous KIF1C expression levels in 293T cells (Fig. 1C). To perform BioID experiments we lysed cells after growth in biotin-containing medium and isolated biotinylated proteins using streptavidin beads. Biotinylated proteins were then identified by mass spectrometry and significant “hits” were determined using a label-free proteomics approach by comparison to a BioID alone control (Redwine et al., 2017; Zhang et al., 2010). Proteins with an enrichment ratio greater than 3-fold and a p-value above 0.05 relative to the control were considered significant hits. One of the top hits from this experiment was HOOK3 (Fig. 1D, Table S1). We also detected a dynein complex component, the dynein light chain Tctex-Type 1 (DYNLT1). We did not detect other dynein activating adaptors, with the exception of HOOK1, which was a significant hit, but had a relatively low peptide count.

To further characterize the interaction between KIF1C and HOOK3 we immunoprecipitated endogenous KIF1C and HOOK3 from 293T cells. Immunoblots with antibodies against HOOK3 and KIF1C demonstrated that these proteins co-precipitate (Fig. 1E). Because there are three different HOOK homologs in the human genome (HOOK1, HOOK2 and HOOK3) and because we detected both HOOK3 and HOOK1 in our KIF1C BioID data, we next used immunoprecipitation experiments to confirm these interactions and to determine if KIF1C interacted with the third HOOK homolog, HOOK2. We expressed each HOOK homolog with an amino-terminal HaloTag and a carboxy-terminal 3XFLAG tag in 293T cells. We then immunoprecipitated each tagged HOOK protein using FLAG affinity resin and immunoblotted for endogenous KIF1C. This revealed that endogenous KIF1C co-precipitates with HOOK3, but not HOOK1 or HOOK2 (Fig. 1F). The presence of HOOK1 in our KIF1C BioID dataset was likely due to heterodimerization of HOOK1 with HOOK3, rather than HOOK1 interacting with KIF1C. In our previous BioID experiments with HOOK1 and HOOK3, we detected HOOK1 in HOOK3 datasets and vice versa (Redwine et al., 2017), leading us to speculate that these proteins may heterodimerize under some conditions. This is also consistent with a previous study that suggested possible heterodimerization between HOOK family members (Xu et al., 2008).

We next asked if HOOK3 specifically interacts with KIF1C. The two most closely related kinesin-3 family members to KIF1C are KIF1A and KIF1B (Siddiqui and Straube, 2017). In addition, the kinesin-1s, KIF5A, KIF5B, and KIF5C, are well-characterized cargo-transporting plus-end-directed motors. We expressed each of these kinesins with a carboxy-terminal BioID-3XFLAG tag in 293T cells. We then immunoprecipitated each tagged kinesin using FLAG affinity resin and immunoblotted for endogenous HOOK3. This revealed that endogenous HOOK3 co-precipitates with KIF1C, but not KIF1A, KIF1B, KIF5A, KIF5B or KIF5C (Fig. 1G). We conclude that endogenous HOOK3 and KIF1C interact in a specific manner.

### KIF1C is a processive plus-end-directed motor, whose motility is unaffected by HOOK3

To further explore the interaction between KIF1C and HOOK3, we purified full-length KIF1C tagged with SNAP- and 3XFLAG- tags at its carboxy-terminus and full-length HOOK3 tagged with a HaloTag at its amino-terminus and 3XFLAG tag at its carboxy-terminus from 293T cells. Each protein migrated as a single band when analyzed by SDS-PAGE (Fig. S1A). Using total internal reflection fluorescence (TIRF) microscopy with TMR-labeled KIF1C-SNAP-3XFLAG and Alexa-405-labeled microtubules, we visualized the motile properties of KIF1C on microtubules. Single full-length KIF1C molecules moved processively towards the microtubule plus end (Fig. 2A, Fig. S1B-D) with an average velocity of 0.734 ± 0.223 μm/s (Fig. 2B) and run length of 15.83 μm (Fig. 2C). We next monitored the interaction of full-length KIF1C-TMR with full-length HOOK3 labeled with Alexa-488 via its amino-terminal HaloTag using near-simultaneous two-color TIRF microscopy. HOOK3 alone did not interact with microtubules (Fig. 2D, left panel). In contrast, in the presence of KIF1C, HOOK3 moved robustly towards microtubule plus ends and colocalized with KIF1C (Fig. 2D, right panels). HOOK3 did not appear to affect KIF1C’s motile properties as both the velocity and run length of KIF1C were not significantly different in the presence of HOOK3 (Figs. 2E, F and Figs. S1E, F). These data define the single-molecule motile properties of KIF1C and show that HOOK3 co-migrates with processive KIF1C molecules without affecting KIF1C’s motility.

**Figure 2.**
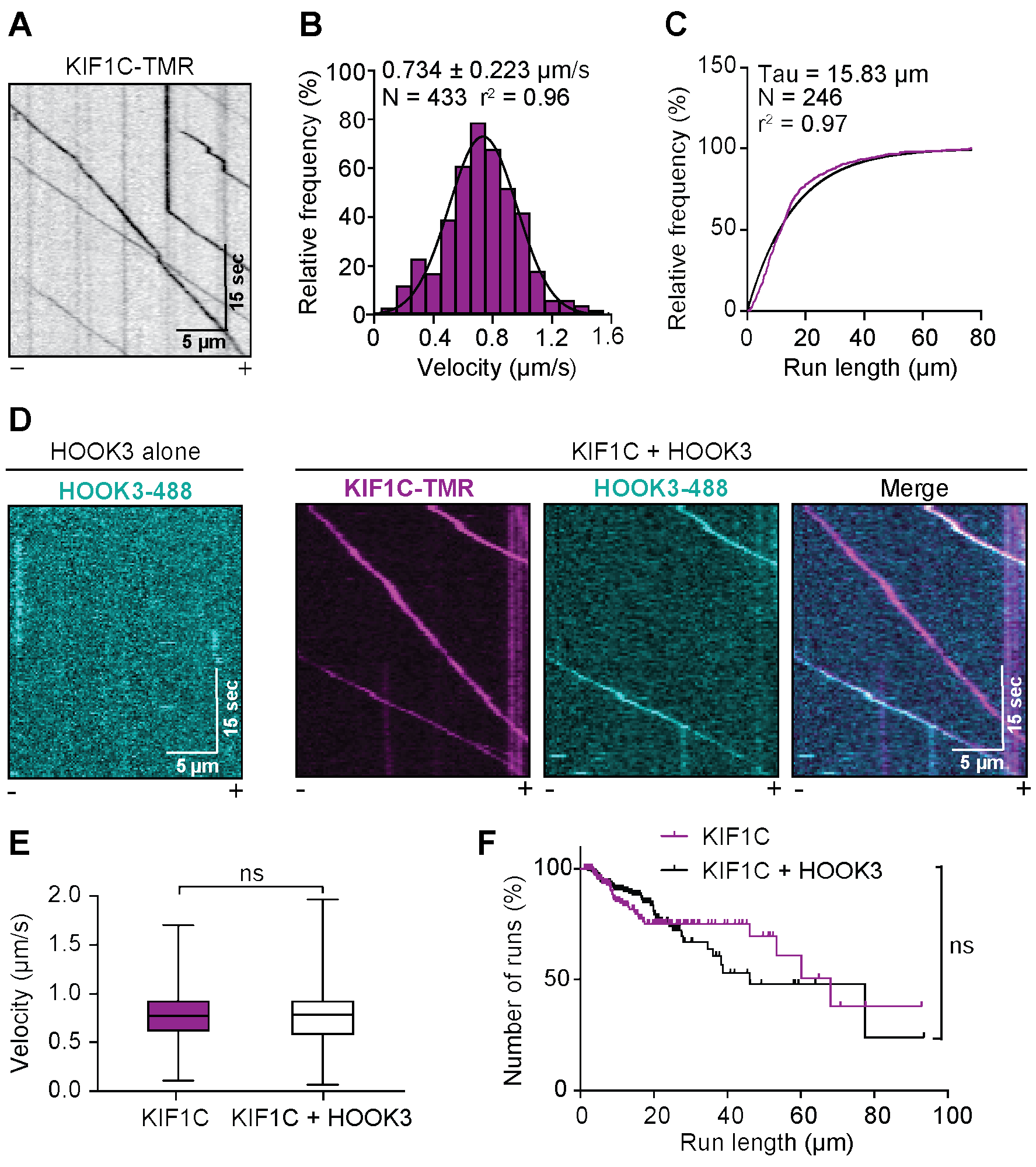
KIF1C is a highly processive kinesin-3 motor. **(A)** Representative kymograph from single-molecule motility assays with full-length KIF1C tagged with SNAP and 3XFLAG, and labeled with TMR via the SNAP-tag. Microtubule polarity is marked with minus (-) and plus (+). **(B)** Single-molecule velocity ± SD of KIF1C-TMR. A Gaussian fit to the data from three independent experiments is shown. **(C)** Run length analysis of KIF1C-TMR. The cumulative frequency distribution was fit to a one phase exponential decay. Mean decay constant is reported from three independent experiments. **(D)** Representative kymograph from single-molecule motility assays with full-length HOOK3 tagged at the amino-terminus with the HaloTag and carboxy-terminus with 3XFLAG, and labeled with Alexa-488 via the HaloTag (HOOK3-488; left panel). KIF1C-TMR in the presence of HOOK3-488 (right panels). Colocalized runs can be seen in the merge in white. Microtubule polarity is marked with minus (-) and plus (+). **(E)** Quantification of velocity ± SD (N = 310-418) from colocalized KIF1C-TMR and HOOK3 runs compared to KIF1C-TMR-only runs. Statistical significance was calculated using an unpaired t-test; ns, no significance. **(F)** Quantification of run length (N = 310-418) from colocalized KIF1C-TMR and HOOK3 runs compared to KIF1C-TMR-only runs. The Kaplan-Meler estimator of run length survivor function (N = 212-301) was used to compare colocalized KIF1C-TMR and HOOK3 runs with KIF1C-TMR-only runs. Statistical significance was calculated with Gehan-Breslow-Wilcoxon test; ns, no significance.

### Fourteen amino acids in the tail of KIF1C are required for HOOK3 binding

We next sought to identify the regions in both KIF1C and HOOK3 responsible for their interaction. We began with KIF1C, generating a series of carboxy-terminal KIF1C truncation constructs, all of which contained a carboxy-terminal 3XFLAG tag (Fig. 3A). We made constructs lacking the carboxy-terminus including the proline-rich region (KIF1C^1-820^), lacking the fourth coiled-coil and the proline-rich region (KIF1C^1-785^), and a deletion mutant that contained a stretch of charged residues as well as two tryptophans (KIF1C^Δ794-807^-3XFLAG) (Fig. 3A). We speculated that the amino acid content in this region might form a protein-protein interaction interface in the KIF1C tail sequence that is otherwise predicted to be mainly unstructured or coiled-coil. Overexpression of these constructs in 293T cells followed by FLAG immunoprecipitations showed that HOOK3 binding to KIF1C is lost when the 14 amino acids (794-807) between coiled-coils 3 and 4 are deleted (Fig. 3B). To confirm the requirement of this region for the KIF1C/ HOOK3 interaction, we purified KIF1C^Δ794-807^ with carboxy-terminal SNAP- and 3XFLAG-tags from 293T cells (Fig. S2A) and labeled it with TMR via the SNAP-tag. Purified KIF1C^Δ794-807^ showed similar velocity and run lengths to wild type KIF1C in a singlemolecule assay (Fig. S2B-D). However, when KIF1C^Δ794-807^-TMR was incubated with HOOK3-Alexa-488 and imaged using near-simultaneous two-color TIRF microscopy, we observed no colocalized events, further demonstrating the importance of this region for HOOK3 binding (Fig. 3C). Taken together our domain mapping identified a 14 amino acid region in the KIF1C tail that is necessary for HOOK3 binding.

**Figure 3.**
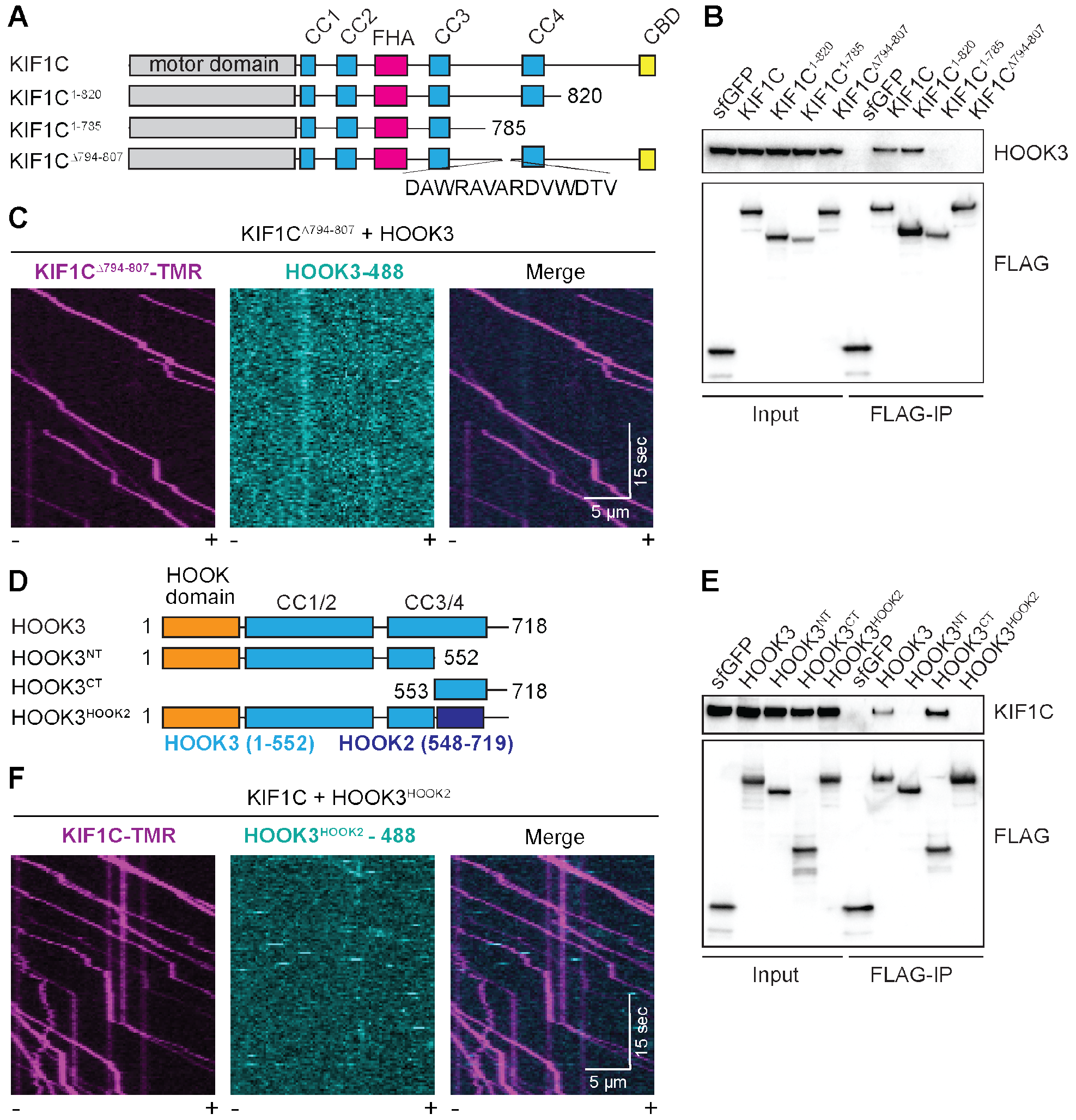
Fourteen amino acids in KIF1C mediate its interaction with HOOK3. **(A)** Schematic of constructs used to map the region of KIF1C that is responsible for binding to HOOK3. **(B)** KIF1C-SNAP-3XFLAG constructs were transiently expressed in 293T cells and immunoprecipitated (FLAG-IP) with FLAG affinity resin. Immunoblots were performed with HOOK3 and FLAG antibodies. 3XFLAG-sfGFP provided a control. **(C)** Representative kymographs from single-molecule motility assays with purified KIF1C^Δ794-807^-TMR in the presence of HOOK3-488. Microtubule polarity is marked with minus (-) and plus (+). **(D)** Schematic of constructs used to map the region of HOOK3 that is responsible for binding to KIF1C. HOOK3^NT^ (AA 1-552), HOOK3^CT^ (AA 553-718), HOOK3^HOOK2^ (a HOOK3 [AA 1-552] and HOOK2 [AA 548-719] chimera). **(E)** HaloTag-HOOK3-3XFLAG constructs were transiently expressed in 293T cells and immunoprecipitated (FLAG-IP) with FLAG affinity resin. Immunoblots were performed with KIF1C and FLAG antibodies. 3XFLAG-sfGFP provided a control. **(F)** Representative kymographs from single-molecule motility assays of KIF1C-TMR in the presence of 488-HOOK3 ^HOOK2^. Microtubule polarity is marked with minus (-) and plus (+).

Next, we set out to map KIF1C’s binding site on HOOK3. We previously showed that the carboxy-terminal region of HOOK3 (amino acids 553-718) is required for the KIF1C interaction (Redwine et al., 2017). To attempt to map this binding site more precisely, we generated a series of constructs lacking various regions in the carboxy-terminal tail of HOOK3. However, these constructs failed to identify a single linear binding site (Fig. S2E), perhaps because the KIF1C binding site on HOOK3 requires a folded domain. As an alternative approach to generate a HOOK3 construct that could no longer bind KIF1C, we designed a chimeric construct in which we replaced amino acids 553-718 of the Halo-HOOK3-3XFLAG construct with the homologous region of HOOK2 (amino acids 548-719; Fig. 3D, Fig S2F), which we showed could not bind KIF1C (Fig. 1F). We then transfected this chimeric construct (HOOK3^HOOK2^, Halo-HOOK3 ^HOOK2^-3XFLAG), full-length HOOK3 (HOOK3, Halo-HOOK3-3XFLAG), HOOK3 lacking the carboxy-terminal region (HOOK3^NT^, Halo-HOOK3^1-552^-3XFLAG), or HOOK3 lacking the amino-terminal region (HOOK3^CT^, Halo-HOOK3^553-718^-3XFLAG) into 293T cells and performed FLAG immunoprecipitations. Only full-length HOOK3 or HOOK3 lacking the amino-terminal region co-immunoprecipitated with endogenous KIF1C (Fig. 3E). To verify that this chimeric HOOK3^HOOK2^ construct does not directly interact with KIF1C, we purified it from insect cells (Fig. S2G) and labeled it with Alexa-488 via its HaloTag. Using near-simultaneous two-color TIRF microscopy we showed that the HOOK3 ^HOOK2^ chimera does not colocalize with KIF1C-TMR in a single-molecule motility assay (Fig. 3F).

### Full-length HOOK3 is a robust dynein activating adaptor

HOOK3 is a well-established dynein activating adaptor (McKenney et al., 2014). However, in vitro assays with pure proteins have previously been performed with a truncated version of HOOK3 (amino acids 1-552, HOOK3^NT^). Here, we aimed to characterize the motile properties of dynein in the presence of full-length HOOK3. Human dynein requires the dynactin complex and an activating adaptor, such as HOOK3, to achieve processive motility (McKenney et al., 2014; Schlager et al., 2014). We purified human dynein and dynactin complexes separately from stable 293T cell lines expressing the dynein intermediate chain (IC2) or the dynactin subunit p62 tagged with the SNAP-tag or HaloTag, respectively, and a 3XFLAG tag (Redwine et al., 2017). We labeled IC2 with Alexa-647, and used non-fluorescently labeled dynactin for these experiments. In the absence of HOOK3, the dynein/dynactin complex is largely stationary in single-molecule motility assays, exhibiting occasional diffusive events and very rare motile events (Fig. S3A). In the absence of dynein, HOOK3 and dynactin are not motile (Fig. S3B). However, the combination of dynein, dynactin and full-length HOOK3 led to robust activation of dynein motility towards microtubule minus ends (Fig. 4A). The velocity (0.730 ± 0.322 μm/s, Fig. 4B) and run length (15.79 μm, Fig. 4C) were comparable to the values we obtained with truncated HOOK3^NT^ (Fig. 4D-F) and were also consistent with previously reported values for truncated HOOK3 (McKenney et al., 2014; Redwine et al., 2017; Schroeder and Vale, 2016) or HOOK3 immunoprecipitated from mammalian cells (Olenick et al., 2016). In addition, the chimeric HOOK3 ^HOOK2^ construct that does not bind KIF1C (Fig. 3E, F) still activated dynein to a similar extent as wild type full-length HOOK3 (Fig. 4G-I).

**Figure 4.**
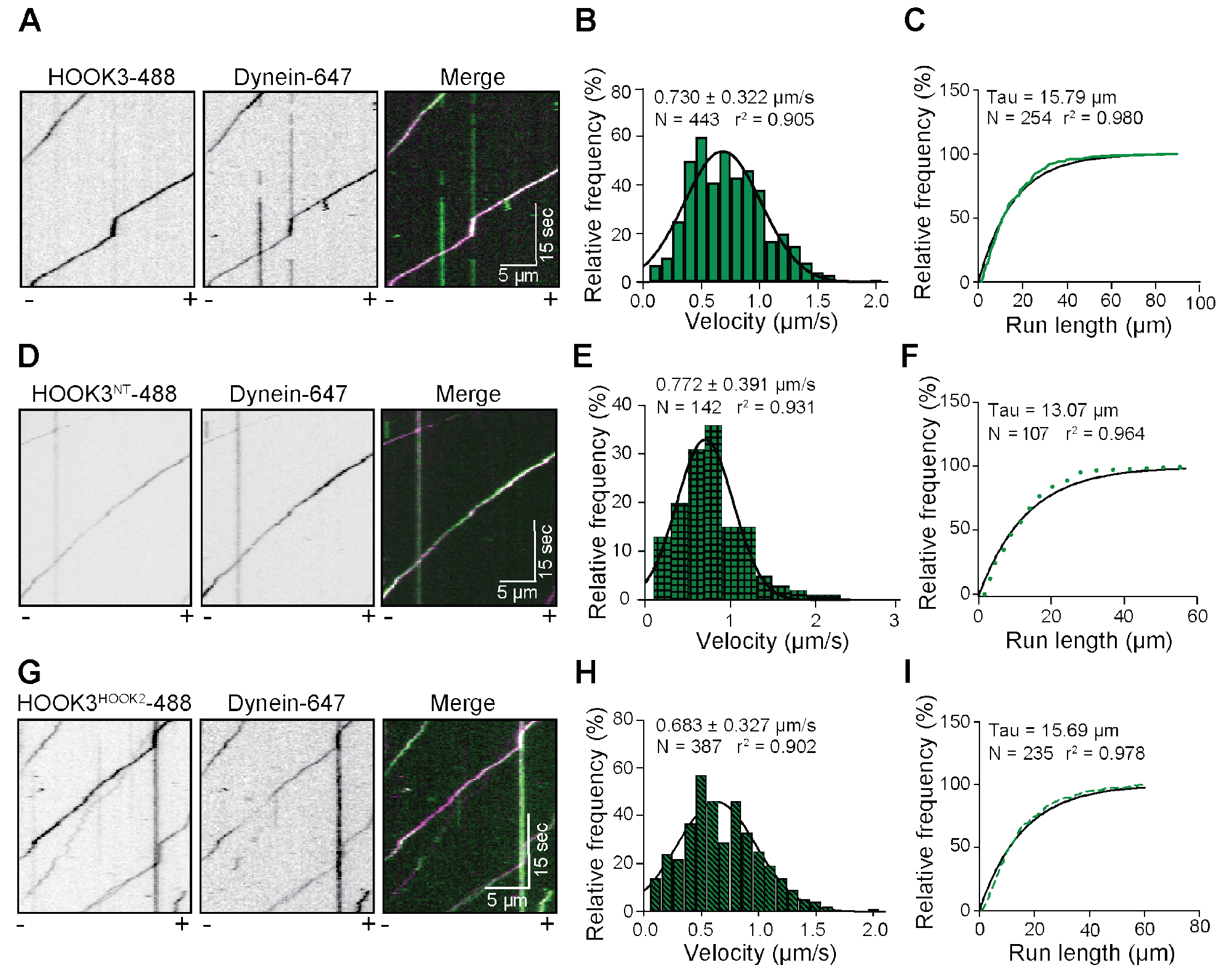
Full-length HOOK3 activates dynein motility. **(A)** Representative kymographs from single-molecule motility assays with purified dynein-647, unlabeled dynactin and full-length HOOK3-488. Microtubule polarity is marked with minus (-) and plus (+). **(B)** Velocity ± SD of dynein/dynactin activated by full-length HOOK3. A Gaussian fit to the data from two independent experiments is shown. **(C)** Run length analysis of dynein/dynactin activated by full-length HOOK3. The cumulative frequency distribution was fit to a one phase exponential decay. Mean decay constant is reported from two independent experiments. **(D)** Representative kymographs from singlemolecule motility assays with purified dynein-647, unlabeled dynactin and HOOK3^1-552^-488. Microtubule polarity is marked with minus (-) and plus (+). **(E)** Velocity ± SD of dynein/dynactin activated by HOOK3^1-552^. A Gaussian fit to the data from two independent experiments is shown. **(F)** Run length analysis of dynein/dynactin activated by HOOK3^1-552^. The cumulative frequency distribution was fit to a one phase exponential decay. Mean decay constant is reported from two independent experiments. **(G)** Representative kymographs from single-molecule motility assays with purified dynein-647, unlabeled dynactin and the HOOK3^HOOK2^-488 chimera. Microtubule polarity is marked with minus (-) and plus (+). **(H)** Velocity ± SD of dynein/dynactin activated by the HOOK3^HOOK2^-488 chimera. A Gaussian fit to the data from two independent experiments is shown. **(I)** Run length analysis of dynein/dynactin activated by the HOOK3^HOOK2^-488chimera. The cumulative frequency distribution was fit to a one phase exponential decay. Mean decay constant is reported from two independent experiments.

### HOOK3 is a scaffold for dynein and KIF1C

Thus far, our experiments indicate that full-length HOOK3 directly associates with KIF1C, and binds to and activates dynein/dynactin complexes (Fig. 5A). We next sought to determine if HOOK3 could bind KIF1C and the dynein/dynactin complex simultaneously. To determine if such a complex exists, we first generated HOOK3 knockout 293T cell lines using CRISPR/Cas9-based gene editing, allowing us to add a tagged version of HOOK3 back to cells lacking HOOK3. This approach attempts to avoid artifacts due to protein over-expression. Co-transfection of 293T cells with Cas9 and guides specific for exon 1 of the HOOK3 transcript followed by clonal selection yielded two clones with HOOK3 knocked out (Fig. 5B). We detected no change in the protein expression levels of KIF1C, a dynein subunit (IC1/2), or a dynactin subunit (p150) in the KIF1C knockout cells (Fig. 5B). We then infected these knockout cells (HOOK3^KO^, clone #1) with a MSCV-driven retroviral plasmid encoding HA-FLAG-HOOK3 to generate cells expressing near-endogenous HOOK3 (Fig. 5C). The protein expression levels of KIF1C, IC1/2, and p150 did not change upon introduction of exogenous HOOK3 (Fig. 5C). We used this cell line to perform immunoprecipitations of HA-FLAG-HOOK3 with FLAG affinity resin under mild conditions to maintain protein-protein interactions. We then analyzed the immunoprecipitate by size exclusion chromatography. Several high molecular weight fractions (centered around A11 to B1) contained dynein and dynactin subunits, as well as HOOK3 and KIF1C (Fig. 5D).

**Figure 5.**
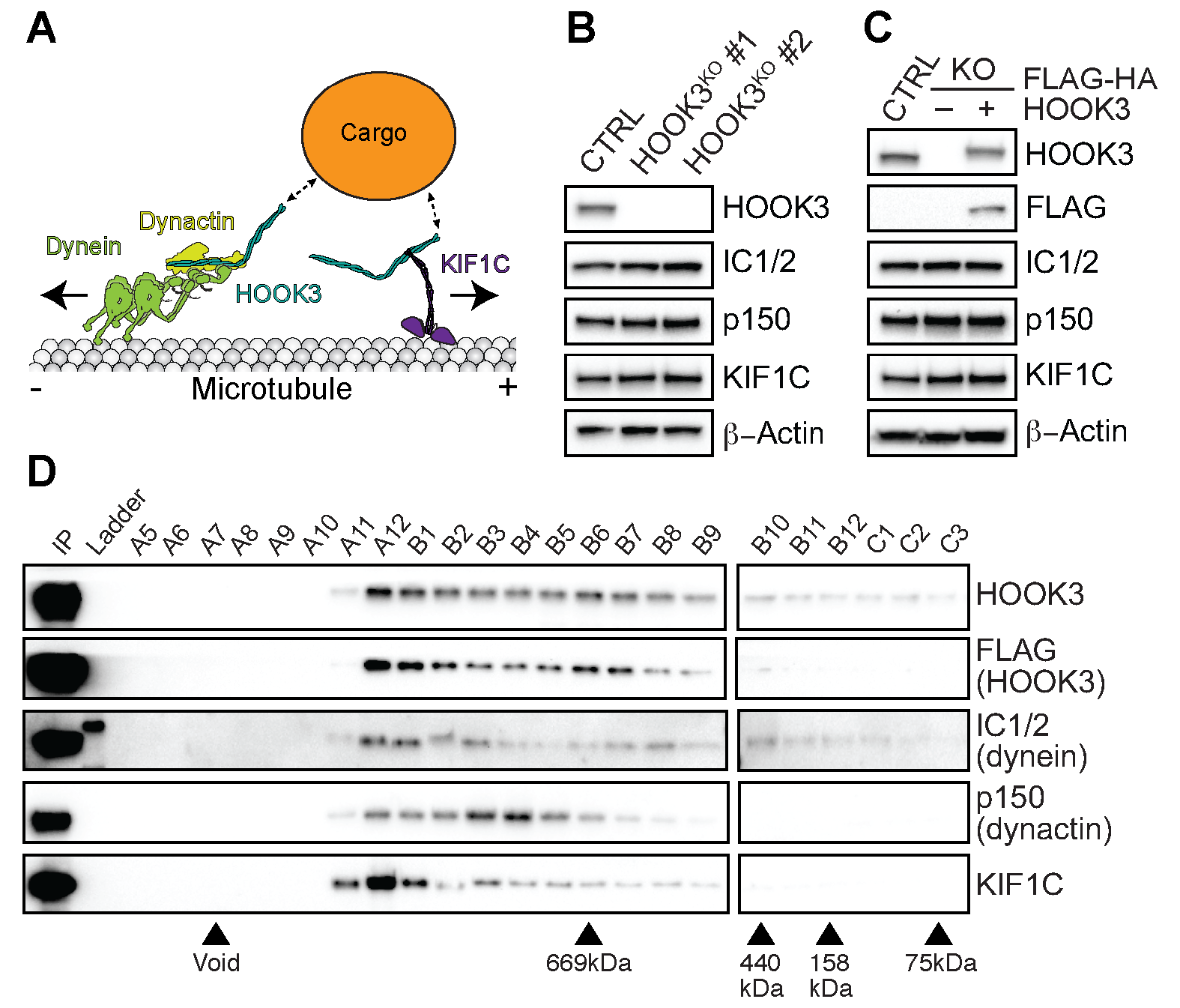
KIF1C is in a complex that contains HOOK3, dynein, and dynactin. **(A)** Schematic of HOOK3 interactions with dynein, dynactin, and KIF1C. **(B)** 293T cells were transfected with control CRISPR-Cas9 (CTRL) or with CRISPR-Cas9-gRNA specific for HOOK3. HOOK3 knockout (HOOK3^KO^) was confirmed in two different clones by immunoblotting with HOOK3 antibody. Clone #1 was selected for further assays. The expression of KIF1C, dynein (intermediate chain 1 and 2, IC1/2) and dynactin (p150 subunit) in the knockout cells lines was examined by immunoblotting with antibodies to each protein. ß-Actin provided a loading control. **(C)** HOOK3^KO^ cells (clone #1) were infected with viral particles encoding murine stem cell virus (MSCV)-driven FLAG-HA-HOOK3 plasmid to obtain near-endogenous HOOK3 expression. Protein expression levels were determined using the indicated antibodies. ß-Actin provided a loading control. **(D)** Size exclusion chromatography analysis of FLAG-HA-HOOK3 complexes purified from HOOK3^KO^ 293T cells expressing near-endogenous HOOK3. After purification on FLAG resin, the extracts were applied to a TSKgel size exclusion column and fractions were analyzed by immunoblotting with the indicated antibodies. Molecular weight standards are indicated (triangles). The data is representative of two independent experiments.

To further examine whether KIF1C and dynein exist in the same HOOK3-scaffolded complexes, we preformed near-simultaneous three-color TIRF microscopy experiments with purified components. For these experiments dynein IC2 was labeled with Alexa-647, HOOK3 with Alexa-488, and KIF1C with TMR. This experimental set up allowed us to detect moving events corresponding to 1) KIF1C alone, 2) KIF1C with HOOK3, 3) dynein with HOOK3, and 4) KIF1C with dynein and HOOK3 (Fig. 6A). Complexes that contained all three labeled components (dynein-467, HOOK3-488 and KIF1C-TMR) moved in either the minus-(Fig. 6B) or plus-(Fig. 6C) end directions. The presence of these three-color colocalized events implies that HOOK3 scaffolds dynein/dynactin and KIF1C to form a complex capable of moving towards the microtubule plus- or minus-end. As expected, we did not observe three-color colocalized runs when TMR-labeled KIF1C^Δ794-807^ (Fig. S4A) or Alexa-488 labeled HOOK3^HOOK2^ (Fig. S4B) were used as opposed to their full-length counterparts. We also did not observe any runs in which the moving molecules changed direction. Next, we quantified the velocity of each detectable species. KIF1C with HOOK3, and dynein/dynactin with HOOK3 both behaved as reported above (Fig. 2D-F, Fig. 4A-C, and Fig. 6D). However, complexes containing KIF1C and dynein/dynactin scaffolded by HOOK3 had slower velocities in the minus-end direction compared to complexes where KIF1C was absent (Fig. 6D). The slowing of the scaffolded complex in the minus-end direction suggests that KIF1C may engage the microtubule when dynein is the primary driver of motility.

**Figure 6.**
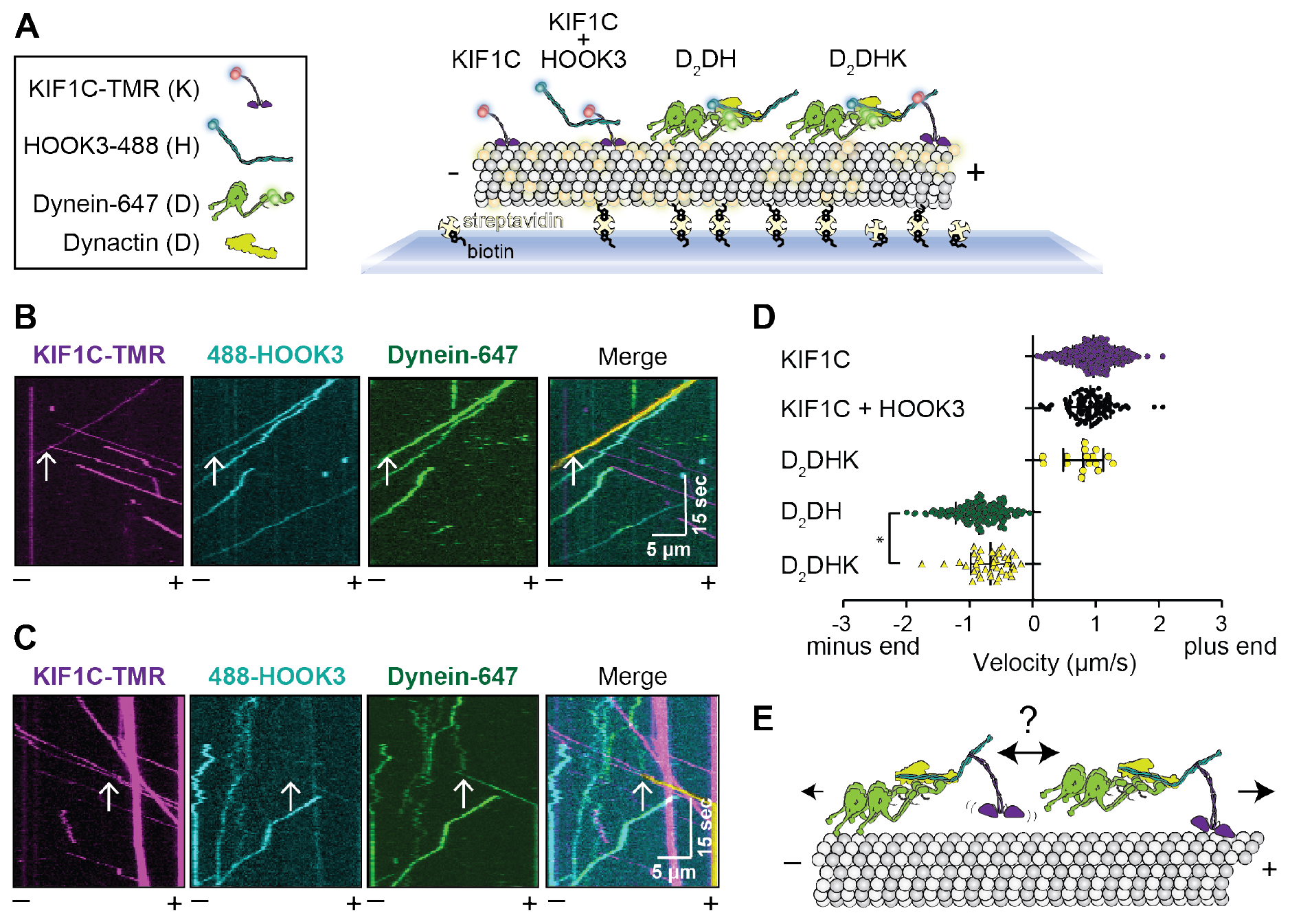
HOOK3 is a scaffold for opposite polarity motors. **(A)** Schematic of experimental set up. Four different species are detectable using 3-color imaging: 1) KIF1C-TMR, 2) KIF1C-TMR with HOOK3-488, 3) Dynein-647 with dynactin and HOOK3-488, and 4) Dynein-647 with dynactin, HOOK3-488, and KIF1C-TMR. **(B)** Representative kymographs from single-molecule motility assays with purified dynein-647, unlabeled dynactin, KIF1C-TMR and HOOK3-488. A three-color colocalized plus-end-directed run is marked with white arrows on each single-channel image and in the merge. The yellow signal in the merge highlights the colocalized run. Microtubule polarity is marked with minus (-) and plus (+). **(C)** Representative kymographs from single-molecule motility assays with purified dynein-647, unlabeled dynactin, KIF1C-TMR and HOOK3-488. A three-color colocalized minus-end-directed run is marked with arrows on each single-channel image and in the merge. The yellow signal in the merge highlights the colocalized run. Microtubule polarity is marked with minus (-) and plus (+). **(D)** Velocity analysis of the indicated complexes (N=16-328). Statistical significance was calculated with a One-Way Anova with Turkey post-test; *, <0.05. Combined data from at least three independent experiments is shown. **(E)** Model of HOOK3 acting as a scaffold for opposite polarity movement on microtubules. The arrows indicate plus- or minus-end velocity.

## Discussion

Here we have shown that the dynein activating adaptor HOOK3 directly scaffolds the opposite-polarity motors cytoplasmic dynein-1/dynactin and the kinesin KIF1C (Fig. 6E). In doing so, we have reported the single-molecule motile properties of KIF1C and full-length HOOK3 in a complex with dynein and dynactin. We mapped the HOOK3 interaction region on KIF1C to 14 amino acids in its tail and showed that while full-length HOOK3 activates dynein/dynactin motility it does not affect the motile properties of KIF1C. Finally, we have reconstituted the HOOK3 scaffold system, showing that dynein/dynactin can transport KIF1C and KIF1C can transport dynein.

### KIF1C is a highly processive motor whose activity is unaffected by HOOK3

Under our analysis conditions full-length KIF1C is a highly processive motor with a characteristic run length that is greater than 15 microns. Kinesin-1s are regulated by autoinhibition via interactions between their motor and tail domains (Friedman and Vale, 1999; Hackney and Stock, 2000; Stock et al., 1999). Several lines of evidence suggest that dimeric kinesin-3 family members are also autoinhibited (Farkhondeh et al., 2015; Hammond et al., 2009; Yamada et al., 2007). The fact that our purified full-length KIF1C is a robust processive motor may indicate that it is not autoinhibited. It is also possible that KIF1C as purified from human 293T cells contains post-translational modifications that could relieve autoinhibition. High-throughput proteomics approaches suggest that KIF1C is phosphorylated, as well as mono-methylated, in multiple regions in its tail (Hornbeck et al., 2015). One of these phosphorylation sites, serine 1092, has been proposed to affect KIF1C’s interaction with 14-3-3 proteins (Dorner et al., 1999). However, how phosphorylation at this site or other candidate posttranslational modifications affects KIF1C’s motor activity has not been investigated. In our experiments the addition of HOOK3 did not modify the motile properties of KIF1C, either in terms of its velocity or its run length. This suggests that HOOK3 is not required to activate KIF1C motility, but rather that HOOK3 serves to link KIF1C to other proteins or cargos.

While we were preparing this manuscript for publication another group reported the motile properties of KIF1C on a preprint server (Siddiqui. 2018 Preprint). In this study, KIF1C purified from insect cells appeared to be largely inactive as observed by single-molecule motility experiments. The addition of HOOK3 or deletion of a region of KIF1C’s third predicted coiled-coil domain increased the landing rate of KIF1C. This study mapped the KIF1C/HOOK3 interaction site to a region (amino acids 714-809) that contained the-14 amino acid (amino acids 794-807) binding site that we identified. We speculate that the differences between our results and this work could be due to the source of the protein, posttranslational state of the protein, purification methods, or motility assay conditions. Further work will be required to differentiate among these possibilities.

### HOOK3 is a scaffold for bidirectional motility

Our size exclusion chromatography and three-color single-molecule experiments show that HOOK3, KIF1C, and dynein/dynactin exist in a complex together. Furthermore, complex formation requires the KIF1C binding site we identified, and the complex does not form when the carboxy-terminus of HOOK3 is replaced by the carboxy-terminus of HOOK2 (the HOOK family member we showed did not bind KIF1C). The analysis of three-color motility experiments indicates that KIF1C can transport dynein/dynactin towards microtubule plus ends and that dynein/dynactin can transport KIF1C towards microtubule minus ends. We do not observe any switches in direction, but we do observe that the presence of KIF1C can slow the velocity of dynein/dynactin/HOOK3 in the minus end direction. This is consistent with a model in which KIF1C engages microtubules while being pulled by dynein towards microtubule minus ends, thus slowing dynein’s velocity (Fig. 6E). Similar velocity decreases have been observed with other opposite polarity motor teams (Belyy et al., 2016; Derr et al., 2012; Roberts et al., 2014). Our data suggest that if dynein and KIF1C share a common cargo that moves bidirectionally, the activity of each motor type may be regulated to achieve changes in direction (Fig. 6E). Our reconstituted system is likely missing these regulatory factors as indicated by the lack of directional switching of dynein/dynactin/HOOK3/KIF1C complexes.

### What is the physiological function of kinesin and dynein complexes scaffolded by HOOK3?

It is possible that HOOK3 scaffolds KIF1C and cytoplasmic dynein for the bidirectional motility of a shared cargo(s). Consistent with this, KIF1C moves bidirectionally in human epithelial cells (Theisen et al., 2012). Multiple cargos for KIF1C have been proposed. KIF1C is implicated in the transport of a5ß1-integrins, and focal adhesion and podosome formation (Efimova et al., 2014; Kopp et al., 2006; Theisen et al., 2012). KIF1C has also been shown to bind to Rab6 (Lee et al., 2015) and KIF1C depletion leads to defects in synaptic vesicle transport (Lipka et al., 2016; Schlager et al., 2014). In contrast, the most likely cargos for HOOK3 are endo-lysosomal compartments (Guo et al., 2016). HOOK3 is part of the FHF complex, named after its components, Fused-Toes homolog (FTS), HOOK-related protein, and FTS and HOOK-interacting protein (FHIP) (Bielska et al., 2014; Xu et al., 2008; Yao et al., 2014). FHF is thought to link the dynein/dynactin complex to Rab5-marked early endosomes, which move bidirectionally in both neurons and filamentous fungi (Bielska et al., 2014; Guo et al., 2016; Yao et al., 2014; Zhang et al., 2014). Thus, at this point no shared cargo for KIF1C and HOOK3 have been reported and this will be an important area for future investigation.

The fact that the KIF1C and HOOK3 cargos that have been described so far are distinct, raises the possibility that the functional role of HOOK3 in scaffolding dynein and KIF1C could be to recycle one or both motors. This could be achieved by KIF1C transporting dynein to the microtubule plus end or dynein/dynactin transporting KIF1C to microtubule minus-ends. Such recycling of dynein by kinesin has been observed in S. *cerevisiae*, where dynein is transported to microtubule plus ends by a kinesin and a set of accessory proteins (Moore et al., 2009). This process is through direct protein-protein interactions, as it has been reconstituted in vitro (Roberts et al., 2014). In addition, in filamentous fungi and neurons, kinesin-1 family members are required for dynein’s plus end localization (Twelvetrees et al., 2016; Zhang et al., 2003). If and how kinesins are recycled is less clear; one study of mammalian kinesin-1 suggests that diffusion is sufficient for its recycling (Blasius et al., 2013). An exciting direction for future studies will be to determine if HOOK3 plays a role in recycling either dynein/dynactin or KIF1C.

### Is scaffolding of dynein/dynactin and kinesin by dynein activating adaptors a general principle?

We have directly demonstrated that the dynein activating adaptor HOOK3 scaffolds KIF1C and dynein/dynactin and that these complexes can move either towards the plus or minus end of microtubules. Do other dynein activating adaptors perform similar functions for dynein/dynactin and other kinesins? There are hints in the literature that this may be the case. For example, interactions between KIF1C and both BICD2 and BICDL1 have also been suggested. In the case of BICD2, network analysis of genes mutated in hereditary spastic paraplegias (HSPs), a disease associated with KIF1C mutations (Dor et al., 2014), identified BICD2 as a possible KIF1C interactor (Novarino et al., 2014). This interaction was confirmed by co-immunoprecipitation with overexpressed proteins (Novarino et al., 2014). In the case of BICDL1, BICDL1 was shown to interact with KIF1C via two-hybrid experiments and co-immunoprecipitation of endogenous KIF1C with overexpressed BICDL1 (Schlager et al., 2010). Other proteins that share a similar general domain structure to the *bona fide* dynein activating adaptors, such as TRAK1, TRAK2, and HAP1 are candidate dynein activating adaptors (Reck-Peterson et al., 2018). Interestingly, TRAK1, TRAK2, and HAP1 have all been shown to interact with kinesin-1 family members and dynein/dynactin subunits (Engelender et al., 1997; Li et al., 1998; Twelvetrees et al., 2010; van Spronsen et al., 2013). Cell biological and in vitro reconstitution experiments will be required to determine if these candidate dynein activating adaptors and other known dynein activating adaptors scaffold dynein/dynactin to kinesin family members for bidirectional motility.

## Acknowledgements

We thank Jenna Chistensen, Zaw Htet, John Salogiannis, and Andres Leschziner for critically reading the manuscript, and members of the Reck-Peterson lab for many lively discussions. S.L.R.-P. is funded by the Howard Hughes Medical Institute and the National Institutes of Health (NIH; R01GM121772). A.A.K. was funded by the NIH (F32GM125224) and currently by the American Cancer Society (PF-18-190-01-CCG). J.W.H. is funded by the NIH (R37NS083524 and RO1NS110395) and a generous gift from Ned Goodnow. J.W.H. is a consultant and founder of Rheostat Therapeutics and a consultant for X-Chem Inc. The other authors have no competing financial interests to declare.

## Author Contributions

A.A.K, W.B.R. and S.L.R.-P. designed the experiments. A.A.K, W.B.R., P.T.T., L.P.V., and M.D. performed the experiments. A.A.K, W.B.R., L.P.V., J.W.H., and S.L.R.-P. interpreted the data. A.A.K. and S.L.R.-P. wrote the original draft of the manuscript. A.A.K, W.B.R., P.T.T., L.P.V., M.D., J.W.H., and S.L.R.-P. reviewed and edited the manuscript.

## Materials and Methods

### Molecular cloning

All plasmids used in this study, unless otherwise stated were constructed by PCR and Gibson isothermal assembly. BioID G2 (Kim et al., 2014) was a kind gift of Kyle Roux (Sanford School of Medicine, University of South Dakota). P62 (isoform 1, 460 aa) was amplified from a human RPE1 cell cDNA library (generated in the Reck-Peterson lab). HOOK3 (clone ID: 5106726), KIF1A (clone ID: 40037561), KIF1B (clone ID: 319918), KIF5A (clone ID: 40148192), KIF5B (clone ID: 8991995) and KIF5C (clone ID: 516562) cDNAs were obtained from Dharmacon. HOOK1 (clone ID: HsCD00044030), HOOK2 isoform 2 (clone ID: HsCD00326811) and KIF1C (clone ID: HsCD00336693) cDNAs were from PlasmidID (Harvard Medical School). HOOK2 isoform 2 clone was mutagenized in the Reck-Peterson lab to generate HOOK2 isoform 1. The HOOK3^ho^°^K2^ chimera construct was generated by replacing HOOK3 amino acids 553-718 with HOOK2 amino acids 548-719 using Gibson isothermal assembly and cloned into pLIB vector containing an amino-terminal His_6_-ZZ-TEV-HaloTag for expression in Sf9 cells. pMSCV-FLAG-HA-HOOK3 was generated using Gateway cloning as previously described (Sowa et al., 2009). The pSpCas9(BB)-2A-Puro (PX459) V2.0 vector was a gift from Feng Zhang (Addgene plasmid #62988).

### Cell lines and transfections

Hum an 293T cells were obtained from ATCC and maintained at 37°C with 5% CO2 in Dulbecco’s Modified Eagle Medium (DMEM, Corning) supplemented with 10% fetal bovine serum (FBS, Gibco) and 1% Penicillin Streptomycin (PenStrep, Corning). Cells were routinely tested for mycoplasma contamination.

#### CRISPR/Cas9-mediated genome editing

Gene editing for creation of KIF1C and HOOK3-depleted 293T cells was performed as described previously (Ran et al., 2013). Briefly, a 20-nucleotide gRNA for HOOK3 was cloned into pX459 CRISPR/Cas9 vectors. The HOOK3 exon 1 gRNA sequence was 5’-GATGTTCAGCGTAGAGTCGC-3’. For KIF1C *in vitro* transcribed 20-nucleotide Alt-R crRNA (Hs.Cas9.KIF1C.1.AD) along with Alt-R CRISPR/Cas9 tracrRNA were purchased from Integrated DNA technologies (IDT). The KIF1C exon 3 crRNA sequence was 5’-TCTCACTAACGCGAGAGAAG −3’. To prepare the Alt-R crRNA and Alt-R tracrRNA duplex, reconstituted oligos (100 μM) were mixed at equimolar concentrations in a sterile PCR water and annealed at 95°C for 5 mins, following slow cooling to room temperature. To generate HOOK3 knockout cells, 200 ng of pX459 vector containing HOOK3 gRNA was diluted in OptiMEM (Gibco) and combined with 1μg/μL polyethylenimine (PEI; Polysciences Inc.) in a 4:1 ratio of PEI:DNA for transfection into 293T cells. KIF1C knockout cell lines were generated by cotransfecting, as described for HOOK3 293T cells with KIF1C crRNA-tracrRNA duplex (10 nM) and Cas9-expressing pX459 vector. 48 hours post-transfection, the cells were pulsed with 1ug/mL puromycin for 24 hours to allow selection of pX459-transfected cells. Following puromycin selection and recovery in DMEM without puromycin, single cell clones were plated in 96-well format by limiting dilution and cultured to allow single colonies to grow out. Clones were expanded to 12-well plates, and samples of resulting clones were screened via immunoblotting with two independent gene-specific antibodies (HOOK3, mouse monoclonal, SCBT cat. No. sc-398924, immunogen from aa202-253 and rabbit polyclonal, Thermo Fisher cat. No. PA5-55172, immunogen from aa214-314) and KIF1C (KIF1C, rabbit polyclonal Novus cat. No. NBP1-85978, immunogen from aa996-1096, and rabbit polyclonal, Thermo Fisher cat. No. PA5-27657 immunogen from aa452-758). A SURVEYOR mutation detection kit (IDT, #706020) was used to detect KIF1C and HOOK3 edited clones.

#### Stable cell lines with near-endogenous protein expression generation

HOOK3^KO^ or KIF1C^KO^ clones were reconstituted with near-endogenous FLAG-HA-HOOK3 or KIF1C-BioID-3XFLAG, respectively, using a retroviral infection/MSCV-driven expression system as described previously (Sowa et al., 2009). Briefly, plasmid DNA (retroviral pMSCV with 3XFLAG-HA-HOOK3, KIF1C-3XFLAG-BioID or BioID-3XFLAG genes inserted) along with viral helper constructs (retroviral MSCV-vsvg, MSCV-gag/pol) were diluted in OptiMEM (Gibco) and combined with 1μg/μL polyethylenimine (PEI; Polysciences Inc.) in a 3:1 ratio of PEI:DNA concentration. The transfection mixture was added to 293T cells, followed by incubation for 12-16 hours. Fresh DMEM was added to the cells, followed by a 24-hour incubation to allow virus production. Viral supernatant was collected, filtered, and added to recipient 293T cells along with 1μg/mL polybrene for infection. Stable cell lines were established by puromycin selection (0.75 μg/mL) for 48-72h. Expression of ectopic proteins was confirmed via immunoblotting with HOOK3 and KIF1C-specific antibodies as well as anti-FLAG-HRP antibody.

#### FLP/FRT stable cell line generation

Dynein (IC2-SNAPf-3XFLAG), dynactin (p62-Halo-3XFLAG), kinesin (KIF1A, KIF1B, KIF1C, KIF5A, KIF5B, and KIF5C) carboxy-terminal BIoID-3XFLAG, and BioID-3XFLAG stable cell lines were created with the FLP/FRT system and T-Rex 293T cells (Invitrogen). These lines were generated as previously described (Redwine et al., 2017). Briefly, one day before transfection cells we plated onto 10 cm dishes. Cells were transfected with 30 μL of Lipofectamine 2000 and a combination of the appropriate pcDNA5/FRT/GOI construct and Flipase expressing pOG44 plasmid (5 μg of total DNA: 9 parts pOG44 + 1 part pcDNA5/FRT/GOI). Following a 24 hour recovery, cells were grown in DMEM containing 10% FBS, 1% PenStrep, and 50 μg/mL Hygromycin B. Colonies were isolated, expanded, and screened for expression of the fusion proteins by immunoblotting with an anti-FLAG M2-HRP antibody.

#### Transient transfections

For small-scale immunoprecipitations from transiently transfected 293T cells, 1.5 x 10^6^ cells were plated onto one 10 cm dish one day before transfection. Transfections were performed with PEI and 2 μg of transfection grade DNA (Purelink midi prep kit, Invitrogen) per dish, with the exception of Halo-HOOK3^1-552^-3xFLAG, where 1 μg of DNA was used due to high protein expression if higher amounts of DNA were used. After 24 hours the media was exchanged for fresh DMEM containing 10% FBS and 1% PenStrep. Cells were then grown for an additional 24 hours before lysate preparation. For large scale protein purifications, 293T cells were plated onto 30 x 15 cm dishes and grown to ~50% confluence. Cells were transiently transfected with PEI and 7.5 μg DNA per plate. The PEI /DNA mixture was added to plates containing fresh DMEM + 10% FBS (no antibiotics) and incubated overnight. The following day the cells were split 1:3 into 90 x 15 cm plates and incubated an additional 24 hours. Cells were collected by pipetting with ice cold 1X phosphate-buffered saline (PBS), centrifuged, and washed twice with 1X PBS. The cells were flash frozen in liquid nitrogen prior to lysis.

### Immunoprecipitations

#### Immunoprecipitation from transiently transfected cells

Transiently transfected cells were collected by decanting the media and washing the cells off the dish with ice cold 1X PBS. Cells were collected by centrifugation at 1000 x g for 3 minutes, washed again with 1X PBS, and then transferred with 1X PBS to Eppendorf tubes for lysis. After spinning 2000 rpm in a microcentrifuge for 4 min and removing the 1X PBS, cells were flash frozen for storage or immediately lysed in 500 μL of dynein lysis buffer (DLB, 30 mM HEPES, pH 7.4; 50 mM KOAc; 2 mM MgOAc; 1 mM EGTA, pH 7.5; 10% glycerol) supplemented with 1 mM DTT, 0.2% Triton X-100, 1X protease inhibitor cocktail (cOmplete Protease Inhibitor Cocktail, Roche) with gentle mixing at 4°C for 20 minutes. Lysates were then centrifuged at maximum speed in a 4°C microcentrifuge for 15 min. For each immunoprecipitation, 420 μL clarified lysate was retrieved and added to 50 μL packed volume of anti-FLAG M2 agarose slurry (Sigma) and incubated for 2 hours at 4°C. Cells were washed four times with 1 mL of DLB, and elutions were carried out with 50 μL of DLB supplemented with 1 mg/mL 3XFLAG peptide.

#### Immunoprecipitation of endogenous proteins

Wild type 293T cells were grown to ~75% confluence and collected by pipetting with cold 1X PBS on ice. For each immunoprecipitation a single 15 cm plate was collected, washed, and resuspended in 1 mL of DLB supplemented with 1 mM DTT, 0.2% Triton X-100, 1X protease inhibitor cocktail (cOmplete Protease Inhibitor Cocktail, Roche). Resuspended cells were gently mixed at 4°C for 15 min, and then centrifuged at maximum speed in a 4°C microcentrifuge. The beads were prepared by incubating appropriate antibodies with Dynabeads Protein G (Fisher Scientific). For each immunoprecipitation sample, 100 μL of bead slurry was washed 3X with 500 μL of 1X PBS and then resuspended in 100 μL of 1X PBS. To this mixture, 4 μg of the appropriate antibody was added (HOOK3, ProteinTech cat. No. 15457-1-AP; KIF1C, Bethyl cat. No. A301-070A; Normal Rabbit IgG, Cell Signaling cat. No. 2729) and incubated for 30 min at room temperature. The resin was washed twice with 1X PBS and then once with DLB. After removing the final wash, 1 mL of cell lysate was added to the prepared resin and incubated for 4 hours at 4°C. The beads were then washed 3 times with 1 mL DLB. To elute proteins, the resin was resuspended in 60 μL of 4X sample buffer and heated at 95°C for 5 minutes. Eluted proteins were analyzed by SDS-PAGE followed by immunoblotting.

### Immunoblotting and antibodies

Lysates and eluates were run on 4-15% polyacrylamide gels (NuPage, Invitrogen), with the exception of gel filtration elution fractions (see below for details), which were separated on 3-8% Tris-Acetate gels (NuPage, Invitrogen). Protein gels were transferred to PVDF membranes for 1.5 hours at 110 V (constant voltage) at 4°C. The membranes were blocked with PBS + 0.05% Tween-20 (v/v, PBST) + 5% dry milk (w/v) and immunoblotted with the appropriate antibodies. All antibodies were diluted in PBST + 1% milk (w/v). Primary antibodies were incubated overnight at 4°C, while secondary antibodies were incubated for 1 hour at room temperature. Immunoblots were visualized with Supersignal West Pico Chemiluminescent reagent (Thermo Fisher) or Supersignal West Femto Chemiluminescent reagent (Thermo Fisher) on a VersaDoc imaging system (BioRad). Image intensity histograms were adjusted in Image lab Version 6.0.1 (BioRad) and then imported into Adobe Illustrator to make figures.

Antibodies used for immunoblots were as follows: anti-FLAG conjugated HRP (Sigma cat. No. A8592, 1:5000 dilution), anti-KIF1C (Novus Biotechnologies cat. No. NBP1-85978, 1:500 dilution), anti-actin (Thermo Fisher cat. No. MAP-15739, 1:4000 dilution), anti-HOOK3 (ProteinTech cat. No. 15457-1-A, 1:1000 dilution), anti-p150 (BD Biosciences cat. No. 610473, 1:1000 dilution) anti-IC1/2 (Santa Cruz Biotechnology cat. No. sc-13524, 1:200 dilution) goat anti-rabbit HRP (sc-2030, Santa Cruz Biotechnology, 1:4000 dilution) and goat anti-mouse HRP (sc-2031, Santa Cruz Biotechnology, 1:4000 dilution).

### BioID sample preparation and mass spectrometry

#### Cell growth and streptavidin purification

Growth of cells and sample preparation for BioID experiments were performed as previously described with slight modifications (Redwine et al., 2017). Briefly, BioID-3XFLAG or KIF1C-BioID-3XFLAG cells were plated at ~20% confluence in 15 cm dishes as four replicates, with each replicate consisting of 8 x 15 cm plates. After 24 hours, biotin was added to the media to a final concentration of 50 μM, and the cells were allowed to grow for another 16 hours. After decanting the media, cells were dislodged from each plate by pipetting with ice-cold 1X PBS. Cells were centrifuged at 1000 x g for 2 min following two washes with ice cold 1X PBS and the cell pellets were resuspended and lysed in 16 mL RIPA buffer (50 mM Tris-HCl, pH 8.0; 150 mM NaCl; 1% (v/v) NP-40, 0.5% (w/v) sodium deoxycholate; 0.1% (w/v) SDS; 1 mM DTT; and protease inhibitors (cOmplete Protease Inhibitor Cocktail, Roche) by gentle rocking for 15 mins at 4°C. The cell lysate was clarified via centrifugation at 66,000 x g for 30 min in a Type 70 Ti rotor (Beckman Coulter; Brea, CA) at 4°C. The clarified lysate was retrieved and combined with pre-washed 0.8 mL streptavidin-conjugated beads (Pierce Strepavidin magnetic beads) and incubated overnight at 4°C with gentle rocking. Bead/lysate mixtures were collected on a magnetic stand into a single 2 mL round-bottom microcentrifuge tube. The beads were then washed 3 times with 2 mL RIPA buffer and once with 1X PBS with immobilization and solution removal performed on a magnetic stand.

#### On-bead digestion

Samples were prepared for mass spectrometry (MS) as follows. After the final wash the beads were resuspended in 100 μL of 50 mM ammonium bicarbonate and the proteins on the beads were reduced with 10 mM DTT for 30 min at room temperature and alkylated with 55 mM iodoacetamide for 30 min in the dark. Protein digestion was carried out with sequencing grade modified Trypsin (Promega) at 1/50 protease/protein (wt/wt) at 37°C overnight. After trypsin digestion, the beads were washed twice with 100 μL of 80% acetonitrile in 1% formic acid and the supernatants were collected. Samples were dried in Speed-Vac (Thermo Fisher) and desalted and concentrated on Thermo Fisher Pierce C18 Tip.

#### MS data acquisition

On bead digested samples were analyzed on an Orbitrap Fusion mass spectrometer (Thermo Fisher) coupled to an Easy-nLC 1200 system (Thermo Fisher) through a nanoelectrospray ion source. Peptides were separated on a self-made C18 analytical column (100 μm internal diameter, x 20 cm length) packed with 2.7 μm Cortecs particles. After equilibration with 3 μL 5% acetonitrile and 0.1% formic acid mixture, the peptides were separated by a 120 min linear gradient from 6% to 42% acetonitrile with 0.1% formic acid at 400nL/min. LC (Optima™ LC/MS, Fisher Scientific) mobile phase solvents and sample dilutions were all made in 0.1% formic acid diluted in water (Buffer A) and 0.1% formic acid in 80 % acetonitrile (Buffer B). Data acquisition was performed using the instrument supplied Xcalibur™ (version 4.1) software. Survey scans covering the mass range of 350–1800 were performed in the Orbitrap by scanning from *m/z* 300-1800 with a resolution of 120,000 (at *m/z* 200), an S-Lens RF Level of 30%, a maximum injection time of 50 milliseconds, and an automatic gain control (AGC) target value of 4e5. For MS2 scan triggering, monoisotopic precursor selection was enabled, charge state filtering was limited to 2-7, an intensity threshold of 2e4 was employed, and dynamic exclusion of previously selected masses was enabled for 45 seconds with a tolerance of 10 ppm. MS2 scans were acquired in the Orbitrap mode with a maximum injection time of 35 ms, quadrupole isolation, an isolation window of 1.6 m/z, HCD collision energy of 30%, and an AGC target value of 5e4.

#### MS data analysis

MS/MS spectra were extracted from raw data files and converted into .mgf files using a Proteome Discoverer Software (ver. 2.1.0.62). These .mgf files were then independently searched against human database using an in-house Mascot server (Version 2.6, Matrix Science). Mass tolerances were +/- 10 ppm for MS peaks, and +/- 25 ppm for MS/MS fragment ions. Trypsin specificity was used allowing for 1 missed cleavage. Met oxidation, protein N-terminal acetylation, N-terminal biotinylation, lysine biotinylation, and peptide N-terminal pyroglutamic acid formation were allowed as variable modifications while carbamidomethyl of Cys was set as a fixed modification. Scaffold (version 4.8, Proteome Software, Portland, OR, USA) was used to validate MS/MS based peptide and protein identifications. Peptide identifications were accepted if they could be established at greater than 95.0% probability as specified by the Peptide Prophet algorithm. Protein identifications were accepted if they could be established at greater than 99.0% probability and contained at least two identified unique peptides.

To estimate relative protein levels, Normalized Spectral Abundance Factor dNSAFs were calculated for each non-redundant protein, as described (Zhang et al., 2010). Average dNSAFs were calculated for each protein using replicates with non-zero dNSAF values. Enrichment of proteins in streptavidin affinity purifications from KIF1C-BioID-3XFLAG tagged stable cell line relative to a control BioID stable cell line was calculated for all replicates as the ratio of average dNSAF (ratio = avg. dNSAF_KIF1C-BioID_: avg. dNSAF_BioID_). The volcano plot (Fig. 1D) was generated by plotting the log2(fold enrichment) against the -log10(p-value), where the p-value (2-tailed Student’s T-test) was computed by comparing the replicate dNSAF values of KIF1C-BioID to the BioID control. Potential KIF1C interactions were included as statistically significant if they were >3-fold enriched in the KIF1C-BioID-3XFLAG dataset and had p-values < 0.05.

### Protein purification

#### KIF1C

Different KIF1C constructs were purified from 293T cells transiently transfected with KIF1C-SNAPf-3XFLAG or KIF1C^Δ794-807^-SNAPf-3XFLAG. Frozen cell pellets from 45 plates were resuspended in 60 mL of BRB80 lysis buffer (80 mM PIPES, pH 6.8; 1 mM MgCL2; 1 mM EGTA; 10% glycerol; 50 mM KOAc) supplemented with 1 mM DTT, 0.5 mM ATP, 0.2% Triton X-100, 1X protease inhibitor cocktail (cOmplete Protease Inhibitor Cocktail, Roche) and gently mixed at 4°C for 15 min. The lysed cells were then centrifuged at 30k x rpm in a Ti70 rotor (Beckman) at 4°C for 30 min. The clarified lysate was retrieved and added to 0.7 mL packed anti-FLAG M2 agarose resin (Sigma) and incubated with gentle mixing at 4°C for 16 hours. After incubation, the lysate/resin mixture was centrifuged at 1000 rpm for 2 min at 4°C to pellet the resin, the supernatant was decanted, and the resin was transferred to a column at 4°C. The column was washed with 50 mL low salt wash buffer (80 mM PIPES, pH 6.8; 1 mM MgCL2; 1 mM EGTA; 10% glycerol; 50 mM KOAc; 1 mM DTT; 0.02% Triton X-100; 0.5 mM Pefabloc), 100 mL high salt wash buffer (80 mM PIPES, pH 6.8; 1 mM MgCL2; 1 mM EGTA,;10% glycerol; 250 mM KOAc; 1 mM DTT; 0.02% Triton X-100; 0.5 mM Pefabloc), and finally with 150 mL of low salt wash buffer. After the final wash the resin was resuspended in an equal volume of low salt wash buffer (700 μL), moved to room temperature and 7 μL of 1 mM SNAP-JF646 (Tocris Bioscience) or SNAP-TMR (Promega) was added and mixed. The mixture was incubated in the dark at room temperature for 10 min. The column was returned to 4°C and washed with 100 mL of low salt wash buffer. For unlabeled KIF1C constructs the labelling steps were omitted. The resin was resuspended in 700 μL of low salt wash buffer containing 2 mg/mL 3X-FLAG peptide and incubated for 30 min at 4°C. The mixture was retrieved and centrifuged through a small filter column to remove the resin. Aliquots were snap frozen in liquid N2 and stored at −80°C. Protein purity was determined on a Sypro (Thermo Fisher) stained SDS-PAGE gels. The labeling efficiency of KIF1C-SNAPf-JF646 was 97%, KIF1C-SNAPf-AlexaTMR was 86%, and KIF1C^Δ794-807^-SNAPf-AlexaTMR was 99%.

#### Full-length HOOK3

Full-length wild type HOOK3 (Halo-HOOK3(1-718)-3XFLAG) was purified from transiently transfected 293T cells. Frozen cells (90 x 15 cm plates) were resuspended in 80 mL of supplemented with 1 mM DTT, 0.5 mM ATP, 0.2% Triton X-100, 1X protease inhibitor cocktail (cOmplete Protease Inhibitor Cocktail, Roche) and gently mixed at 4°C for 15 min. The lysed cells were then centrifuged at 30k x rpm in a Ti70 rotor (Beckman) at 4°C for 30 min. The clarified lysate was retrieved and added to 1.5 mL packed anti-FLAG M2 agarose (Sigma) and incubated with gentle mixing at 4°C for 16 hours. After incubation, the lysate/resin mixture was centrifuged at 1000 rpm for 2 min at 4°C to pellet the resin, the supernatant was decanted, and the resin was transferred to a column at 4°C. The column was washed with 100 mL low salt wash buffer (30 mM HEPES, pH 7.4; 50 mM KOAc; 2 mM MgOAc; 1 mM EGTA, pH 7.5; 10% glycerol; 1 mM DTT; 0.5 mM ATP; 0.5 mM Pefabloc; 0.02% Triton X-100), 100 mL high salt wash buffer (30 mM HEPES, pH 7.4; 250 mM KOAc; 2 mM MgOAc; 1 mM EGTA, pH 7.5; 10% glycerol; 1 mM DTT; 0.5 mM ATP; 0.5 mM Pefabloc; 0.02% Triton X-100), and finally with 100 mL of low salt wash buffer. The resin was then resuspended in an equal volume of low salt wash buffer (1.5 mL) and 20 μL of 1 mM Halo-Alexa488 was added and mixed. The mixture was incubated in the dark at room temperature for 10 min. The column was returned to 4°C and washed with 100 mL of low salt wash buffer. The resin was resuspended in 1000 μL of low salt wash buffer containing 2 mg/mL 3X-FLAG peptide and incubated for 30 min at 4°C. The mixture was retrieved and centrifuged through a small filter column to remove the resin. The eluate was retrieved and 500 μL was loaded onto a Superose 6 Increase 10/300 GL Column connected to an AKTA FPLC (GE) and run in GF150 buffer. Peak fractions containing Alexa-488 labeled Halo-HOOK3-3X FLAG were pooled and concentrated using a 100 kDa MWCO centrifugal filter (Amicon Ultra, Millipore). Aliquots were snap frozen in liquid N2 and stored at - 80°C. Protein purity was checked on a Sypro (Thermo Fisher) stained SDS-PAGE gel. The labeling efficiency was 91%.

#### HOOK3^HOOK2^ chimera

*HOOK3^HOOK2^ chimera* was purified from Baculovirus-infected SF9 insect cells. Cell pellets from 800 mL culture were resuspended in DLB supplemented with 0.5 mM ATP, 0.2% Triton X-100, 300 mM KOAc and 1X protease inhibitor cocktail (cOmplete Protease Inhibitor Cocktail, Roche) and lysed using a Dounce homogenizer (15 strokes with loose plunger and 10 strokes with tight plunger). The lysate was clarified by centrifuging at 183,960 x g for 30 min. The clarified lysate was retrieved and added to 1.5 mL of IgG Sepharose 6 fast Flow affinity resin (GE Healthcare), pre-equilibrated in lysis buffer and incubated with gentle mixing at 4°C for 2 hours. After incubation, the lysate/resin mixture was centrifuged at 1000 rpm for 2 min at 4°C to pellet the resin, the supernatant was decanted, and the resin was transferred to a column at 4°C. The column was washed with 100 mL low salt TEV buffer (10 mM Tris-HCl, pH 8; 2 mM MgOAc; 1 mM EGTA, pH 7.5; 10% glycerol; 1 mM DTT; 250 mM KOAc), 100 mL high salt TEV buffer (10 mM Tris-HCl, pH 8; 2 mM MgOAc; 1 mM EGTA, pH 7.5; 10% glycerol; 1 mM DTT; 500 mM KOAc), and finally with 100 mL of low salt TEV buffer. The resin was then resuspended in an equal volume of low salt TEV buffer supplemented with 0.02% NP40 and TEV protease and incubated ~16 hours following labeling with 20 μL of 1 mM Halo-Alexa488. The mixture was incubated at 4°C in the dark for 2 hours. Following labeling, the mixture was retrieved and centrifuged through a small filter column to remove the resin. The eluate was retrieved and 500 μL was loaded onto a Superose 6 Increase 10/300 GL Column connected to an AKTA FPLC (GE Healthcare) and run in GF150 buffer. Peak fractions containing Alexa-488 labeled Halo-HOOK3^HOOK2^-3X flag were pooled and concentrated using a 100 kDa MWCO centrifugal filter (Amicon Ultra, Millipore). Aliquots were snap frozen in LN2 and stored at −80°C. Protein purity was checked on a Sypro (Thermo Fisher) stained SDS-PAGE gel. The labeling efficiency was 94%.

#### HOOK3^1-552^

The Strep-Halo-HOOK3^1-552^ construct was transformed into BL21-CodonPlus (DE3)-RIPL cells (Agilent). 2L of cells were grown at 37°C in LB media to a 600 nm optical density of 0.4-0.8 before the temperature was reduced to 18°C and expression was induced with 0.5 mM IPTG. After 16-18 hours, the cells were harvested via centrifugation for 6 min at 4°C at 6,000 rpm in a Beckman-Coulter JLA 8.1000 fixed angle rotor. Pellets were resuspended in 40 mL of DLB supplemented with 0.5 mM Pefabloc SC (Sigma-Aldrich) and 1mg/mL lysozyme and incubated at 4°C for 30 min. Cells were lysed via sonication (Branson Digital Sonifier) and clarified via centrifugation at 66,000 x g for 30 min in a Type 70 Ti rotor (Beckman) at 4°C. Supernatant was loaded onto a 5 mL StrepTrap column (GE Healthcare) and washed with 50-100 mL of lysis buffer. Strep-Halo-HOOK3^1-552^ was eluted with 25-50 mL of elution buffer (DLB with 3 mM d-Desthiobiotin). Elution was then applied to a size exclusion chromatography Superose 6 Increase 10/300 GL column (GE Healthcare) that had been equilibrated with GF150 buffer. Peak fractions containing Alexa-488 Strep-Halo-HOOK3^1-552^ were pooled and concentrated using a 100 kDa MWCO centrifugal filter (Amicon Ultra, Millipore). Aliquots were snap frozen in liquid N2 and stored at −80°C. Protein purity was assayed by SDS-PGAE and Sypro (Thermo Fisher) staining. The labeling efficiency was 75%.

#### Dynein and Dynactin

Dynein (IC2-SNAPf-3XFLAG) and dynactin (p62-Halo-3XFLAG) were purified from stable cell line as previously described (Redwine et al., 2017). Briefly, frozen pellets from 293T cells (80 x 15 cm plates, dynein and 160 x 15 cm plates, dynactin) were resuspended in DLB supplemented with 0.5 mM ATP, 0.2% Triton X-100 and 1X protease inhibitor cocktail (cOmplete Protease Inhibitor Cocktail, Roche) and gently mixed at 4°C for 15 min. The lysed cells were then centrifuged at 30,000 rpm in a Ti70 rotor (Beckman) at 4°C for 30 min. The clarified lysate was retrieved and added to 1.5 mL (dynein) or 3 mL (dynactin) packed anti-FLAG M2 agarose resin (Sigma) and incubated with gentle mixing at 4°C for 16 hours. After incubation, the lysate/resin mixture was centrifuged at 1000 rpm for 2 minutes at 4°C to pellet the resin and the supernatant was decanted. The resin was transferred to a column at 4°C and the column was washed with 100 mL low salt wash buffer (30 mM HEPES, pH 7.4; 50 mM KOAc; 2 mM MgOAc; 1 mM EGTA, pH 7.5; 10% glycerol; 1 mM DTT; 0.5 mM ATP; 0.5 mM Pefabloc; 0.02% Triton X-100), 100 mL high salt wash buffer (30 mM HEPES, pH 7.4; 250 mM KOAc; 2 mM MgOAc; 1 mM EGTA, pH 7.5; 10% glycerol; 1 mM DTT; 0.5 mM ATP; 0.5 mM Pefabloc; 0.02% Triton X-100), and finally with 50 mL of low salt wash buffer. After the final wash the resin was resuspended in an equal volume of low salt wash buffer, moved to room temperature and 15 μL of 1 mM SNAP-Alexa-647 was added and mixed. The mixture was incubated in the dark at room temperature for 10 min. The column was returned to 4°C and washed with 100 mL of low salt wash buffer. The labeling steps were omitted when unlabeled protein was desired. The resin was resuspended in 800 μL of low salt wash buffer containing 2 mg/mL 3X-FLAG peptide and incubated for 30 min at 4°C. The mixture was retrieved and centrifuged through a small filter column to remove the resin. The eluate was next loaded onto a Mono Q 5/50 GL 1 mL column on an AKTA FPLC (GE Healthcare). The column was washed with 5 mL of Buffer A (50 mM Tris-HCl, pH 8.0; 2 mM MgOAc; 1 mM EGTA; 1 mM DTT) and then subjected to a 26 mL linear gradient from 35-100% Buffer B mixed with Buffer A (Buffer B = 50 mM Tris-HCl, pH 8.0; 1 M KOAc; 2 mM MgOAc; 1 mM EGTA; 1 mM DTT), followed by 5 mL additional 100% Buffer B. Fractions containing pure dynein (~60-70% Buffer B) or pure dynactin (~75-80% Buffer B) were pooled and buffer exchanged through iterative rounds of dilution and concentration on a 100 kDa MWCO centrifugal filter (Amicon Ultra, Millipore) using “GF150” buffer as a diluent (GF150 = 25 mM HEPES, pH 7.4; 150 mM KCl; 1 mM MgCL2; 1 mM DTT; 10% glycerol). Purity was evaluated on SDS-PAGE gels and protein aliquots were snap frozen in liquid N2 and stored at −80°C. The labeling efficiency of dynein-Alexa647 was 97%.

### Size-exclusion chromatography

For size-exclusion chromatography, FLAG-HA-HOOK3 was immunoprecipitated from 293T HOOK3^KO^ stable cell lines expressing ectopic FLAG-HA-HOOK3 as described for Halo-HOOK3-3XFLAG purification. Briefly, 50 x 15 cm plates were grown to ~ 75% confluency. Following harvesting and lysis in DLB HOOK3 was immunoprecipitated (~16 hours, 4°C) with FLAG-agarose resin. After washing with low salt wash buffer, the protein was eluted off of the beads and elute was applied to a TSKgel G4000SWxL size exclusion column (Tosoh) pre-equilibrated in GF150 without 10% glycerol and supplemented with 1 mM DTT. Samples were fractionated using an ÄKTA Pure system (GE Healthcare) and analyzed by SDS-PAGE and immunoblotting.

### TIRF microscopy

Imaging was performed with an inverted microscope (Nikon, Ti-E Eclipse) equipped with a 100x 1.49 N.A. oil immersion objective (Nikon, Plano Apo). The xy position of the stage was controlled by ProScan linear motor stage controller (Prior). The microscope was equipped with an MLC400B laser launch (Agilent) equipped with 405 nm (30 mW), 488 nm (90 mW), 561 nm (90 mW), and 640 nm (170 mW) laser lines. The excitation and emission paths were filtered using appropriate single bandpass filter cubes (Chroma). The emitted signals were detected with an electron multiplying CCD camera (Andor Technology, iXon Ultra 888). Illumination and image acquisition was controlled by NIS Elements Advanced Research software (Nikon).

### Single-molecule motility assays

Single-molecule motility assays were performed in flow chambers assembled as described previously (Case et al., 1997) using the TIRF microscopy set up described above. Briefly, biotinylated and PEGylated coverslips (Microsurfaces) were used to reduce non-specific binding. Microtubules contained ~10% biotin-tubulin for attachment to streptavidin-coated cover slip and ~10% Alexa Fluor 405 (Thermo Fisher) tubulin for visualization. Imaging buffer was DLB supplemented with 20 μM taxol, 1 mg/mL casein, 1 mM Mg-ATP, 71.5 mM βME (beta mercaptoethanol) and an oxygen scavenger system (0.4% glucose, 45 μg/ml glucose catalase (Sigma-Aldrich), and 1.15 mg/ml glucose oxidase (Sigma-Aldrich). Images were recorded every 0.3-0.4 sec for 3 min. Movies showing significant drift were not analyzed.

For two-color motility assays of KIF1C with HOOK3, 1.125 nM KIF1C-SNAPf-AlexaTMR was mixed with 2.25 nM HOOK3-Alexa488 or 2.25 nM HOOK3^HOOK2^-Alexa488. The two-color motility measurements of dynein, dynactin and different HOOK3 constructs were all performed with 450 pM dynein-Alexa647, 900 pM unlabeled dynactin and 3.25 pM HOOK3 (HOOK3^1-552^-Alexa488, HOOK3-Alexa488 or HOOK3 ^HOOK2^-Alexa488). The three-color single-molecule motility experiments were performed with 450 pM dynein-Alexa647, 900 pM unlabeled dynactin, 130 nM HOOK3 (HOOK3-Alexa488 or HOOK3 ^HOOK2^-Alexa488) and 0.45 - 1.125 nM KIF1C (KIF1C-SNAPf-TMR or KIF1C^Δ794-807^-SNAPf-TMR). Each protein mixture was incubated on ice for 10 min prior to TIRF imaging.

### Polarity marked microtubules

Polarity marked microtubules were prepared according to previously described protocol with slight modifications (Roberts et al., 2014). Brightly-labeled, biotinylated microtubule seeds were polymerized by mixing Alexa405-tubulin (10 μM), biotin-tubulin (10 μM) and unlabeled tubulin (10 μM) with 0.5 mM GMP-CPP (Jena Bioscience) in BRB80 supplemented with 1 mM DTT and incubating for 30 min at 37°C. Following the addition 10X volume of BRB80, polymerized seeds were pelleted in a benchtop centrifuge (15 min at 16,100 x g) and resuspended in a volume of BRB80 equal to the original polymerization volume. GMP-CPP seeds were then mixed with 1:5 diluted dim mix containing 12 μM 405-tubulin, 15 μM unlababled tubulin, 10 μM biotin tubulin and 15 μM NEM-tubulin and incubated for 30 min at 37°C. After incubation equal volume of BRB80 supplemented with 1 mM DTT and 20 μM taxol was added to the mixture and microtubules were incubated for additional 30 min at 37°C to generate polarity-marked microtubules. 1:25 diluted polarity marked microtubules were flown into flow chambers and single molecule-motility analysis were performed as described above.

### Data analysis

The velocity of moving particles was calculated form kymographs generated in ImageJ as described previously (Roberts et al., 2014). Velocities were only calculated from molecules that moved processively for greater than 5 frames. Non-motile or diffusive events were not considered in velocity calculation. Processive events were defined as events that move uni-directionally and do not exhibit directional changes greater than 600 nm. Diffusive events were defined as events that exhibit at least one bi-directional movement greater than 600 nm in each direction. Single-molecule movements that change apparent behavior (e.g. shift from non-motile to processive) were counted as multiple events.

### Bleaching analysis

Bleaching step analysis was performed in a flow chamber, as described above with biotin-488-microtubules immobilized to the coverslips. 450 pM KIF1C-TMR was flown into the chamber in the presence of DLB supplemented with 1mM ATP, 100 uM Taxol and 0.1 mg/ml casein. Images were acquired every 100 ms for 180 s using 562 nm laser at 70% power. Images were analyzed in Image J with Plot Profile function. Steps were manually counted from individual spot profiles.

### Sequence alignment

Protein sequences of different HOOK isoforms were obtained from UniProt. Sequence alignments were performed with Clustal Omega web services (McWilliam et al., 2013) and annotated using Jalview (Biasini et al., 2014).

### Data analysis

Data visualization and statistical analyses were performed using GraphPad Prism (8.0d; GraphPad Software), Excel (version 16.20; Microsoft), and ImageJ (2.0). Brightness and contrast were adjusted in Image J for all kymographs. In addition, images in Fig. 6B and C were manually colored (yellow) in Photoshop (Photoshop CC version 20) to highlight the three-color colocalized runs. For run length analysis data was plotted as a cumulative probability distribution and fit to a cumulative distribution function (one phase decay, least squares fit). The Kaplan-Meler estimator of run length survivor function was used to compare colocalized two-color events. Statistical significance of run length comparison was calculated with Gehan-Breslow-Wilcoxon test. Statistical analyses for velocities of KIF1C and KIF1C + HOOK3 were done using unpaired t-test. Statistical comparisons of the plus-end and minus-end moving events was performed using oneway Anova with Turkey post-test. Exact value of n and evaluation of statistical significance are described in the corresponding figure legends. All experiments were analyzed from at least three independent replicates, except for D_2_DH^NT^ experiments (Fig. 4D), which was analyzed from two independent replicates.

## Supplemental Figures and Figure Legends

**Figure S1.**
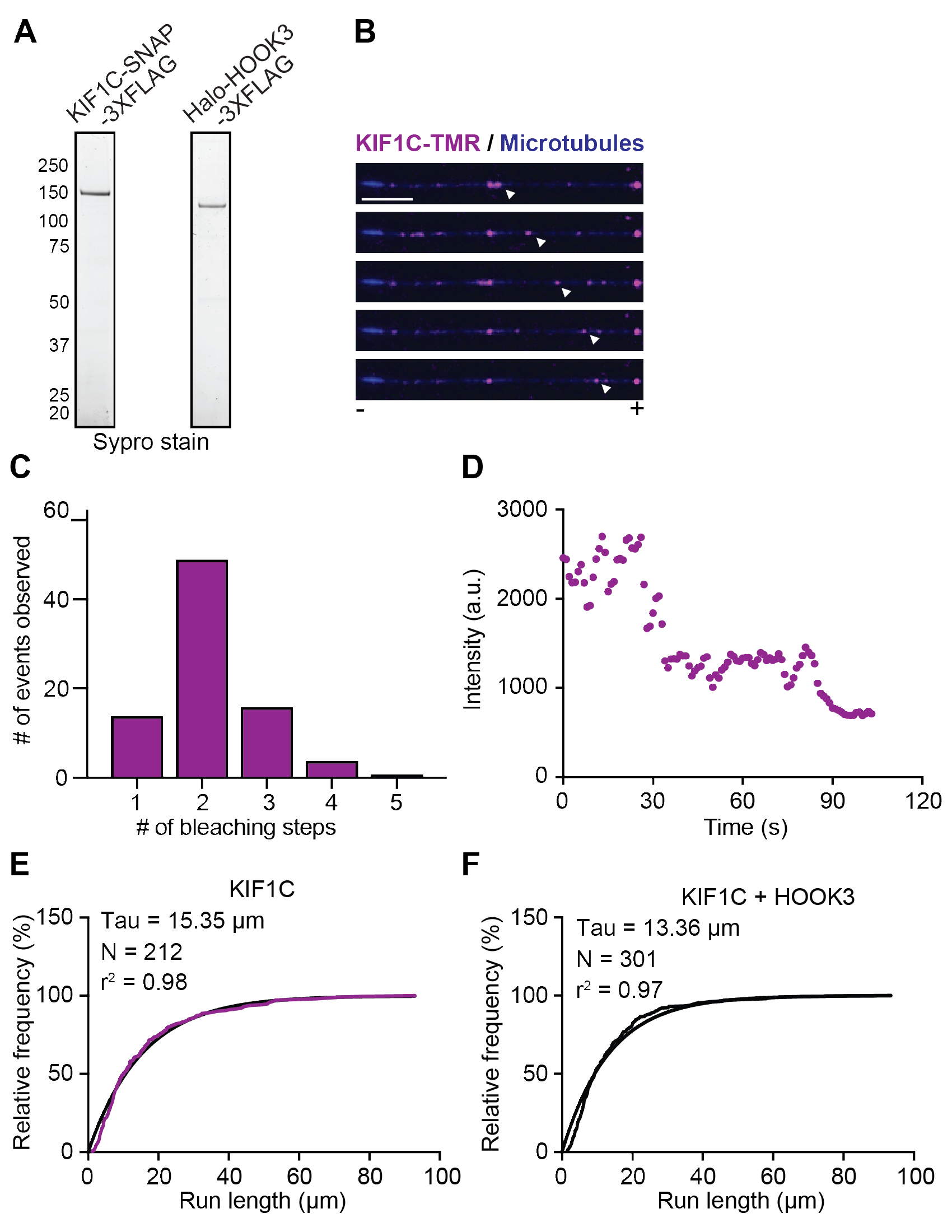
**(A)** SDS-PAGE of purified KIF1C-SNAP-3XFLAG and Halo-HOOK3-3XFLAG used for motility assays. **(B)** KIF1C-TMR (magenta) motility on polarity marked microtubules. The blue seed made with GMPCPP tubulin marks the microtubule minus end. Scale bar is 2 μm. **(C)** The number of photobleaching steps was determined for moving KIF1C molecules (N = 105). **(D)** An example of a two-step photobleaching event. **(E)** Run length analysis of KIF1C-TMR-only runs and **(F)** KIF1C-TMR plus HOOK3-488 colocalized runs from Figure 2F. Cumulative frequency distribution was fit to a one phase exponential decay. Mean decay constants are reported from three independent trails.

**Figure S2.**
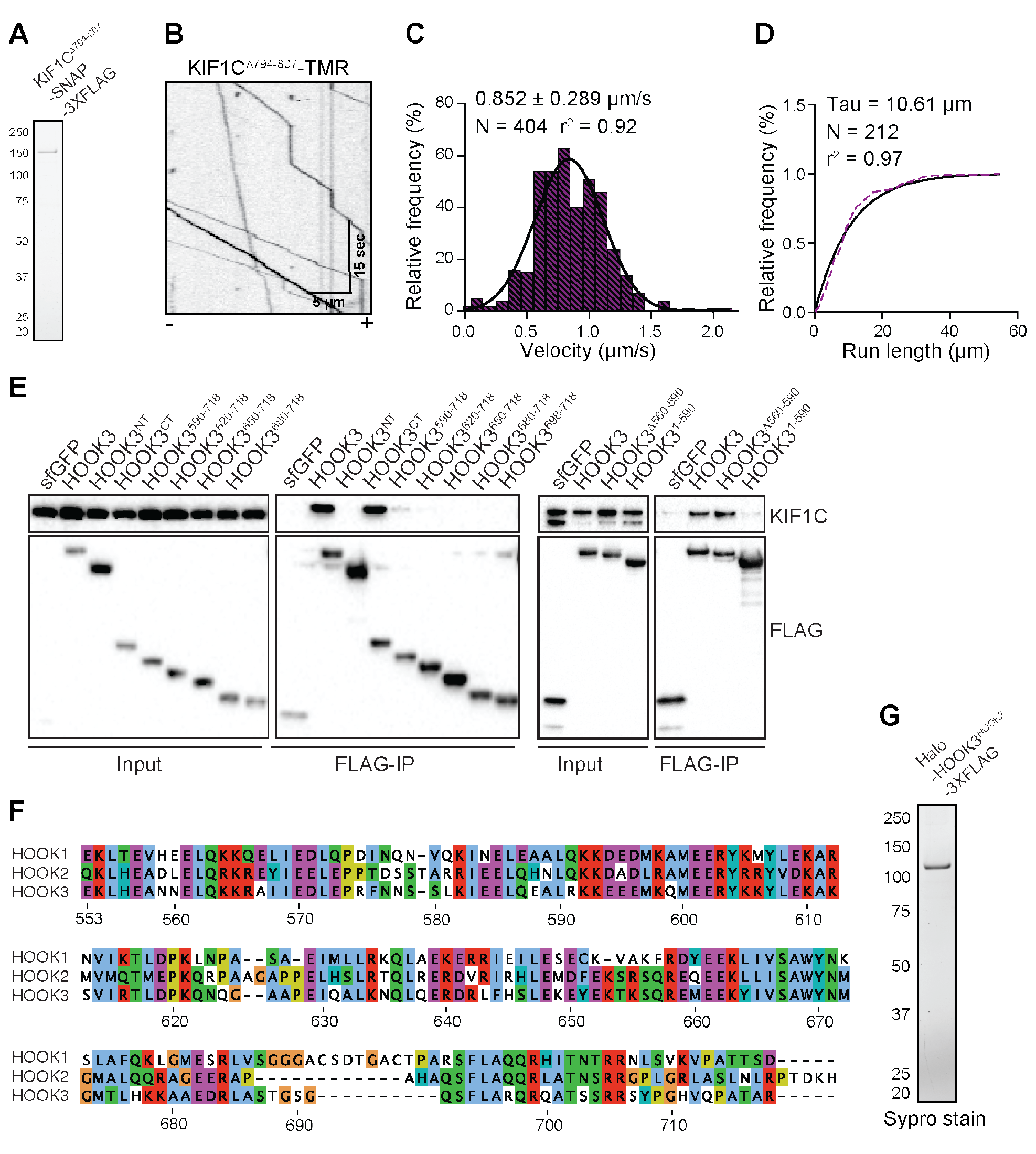
**(A)** SDS-PAGE of KIF1C^Δ794-807^-SNAP-3XFLAG used for motility assays. **(B)** Representative kymograph from single-molecule motility assays with purified full-length Kif1qA7^9^4-^8^07-snap-3XFLAG labeled with TMR. Microtubule polarity is marked with minus (-) and plus (+). **(C)** Single-molecule velocity ± SD of KIF1C^Δ794-807^-TMR. Combined data from three independent experiments is shown. A Gaussian fit to the data from three independent experiments is shown. **(D)** Run length analysis of KIF1CΔ^794-807^-TMR. The cumulative frequency distribution was fit to a one phase exponential decay. Mean decay constant is reported from three independent experiments. **(E)** Indicated HaloTag-HOOK3-3XFLAG constructs were transiently expressed in 293T cells and immunoprecipitated (FLAG-IP) with FLAG affinity resin. Immunoblots were performed with KIF1C and FLAG antibodies. 3XFLAG-sfGFP provided a control. **(F)** Sequence alignment of the carboxy-terminal regions of the three human hook homologs (HOOK3, AA 552-718, HOOK2, AA 548-719, and HOOK1 AA 556-728) made using Clustal Omega. **(G)** SDS-PAGE of Halo-tagged HOOK3^HOOK2^-3XFLAG used for motility assays.

**Figure S3.**
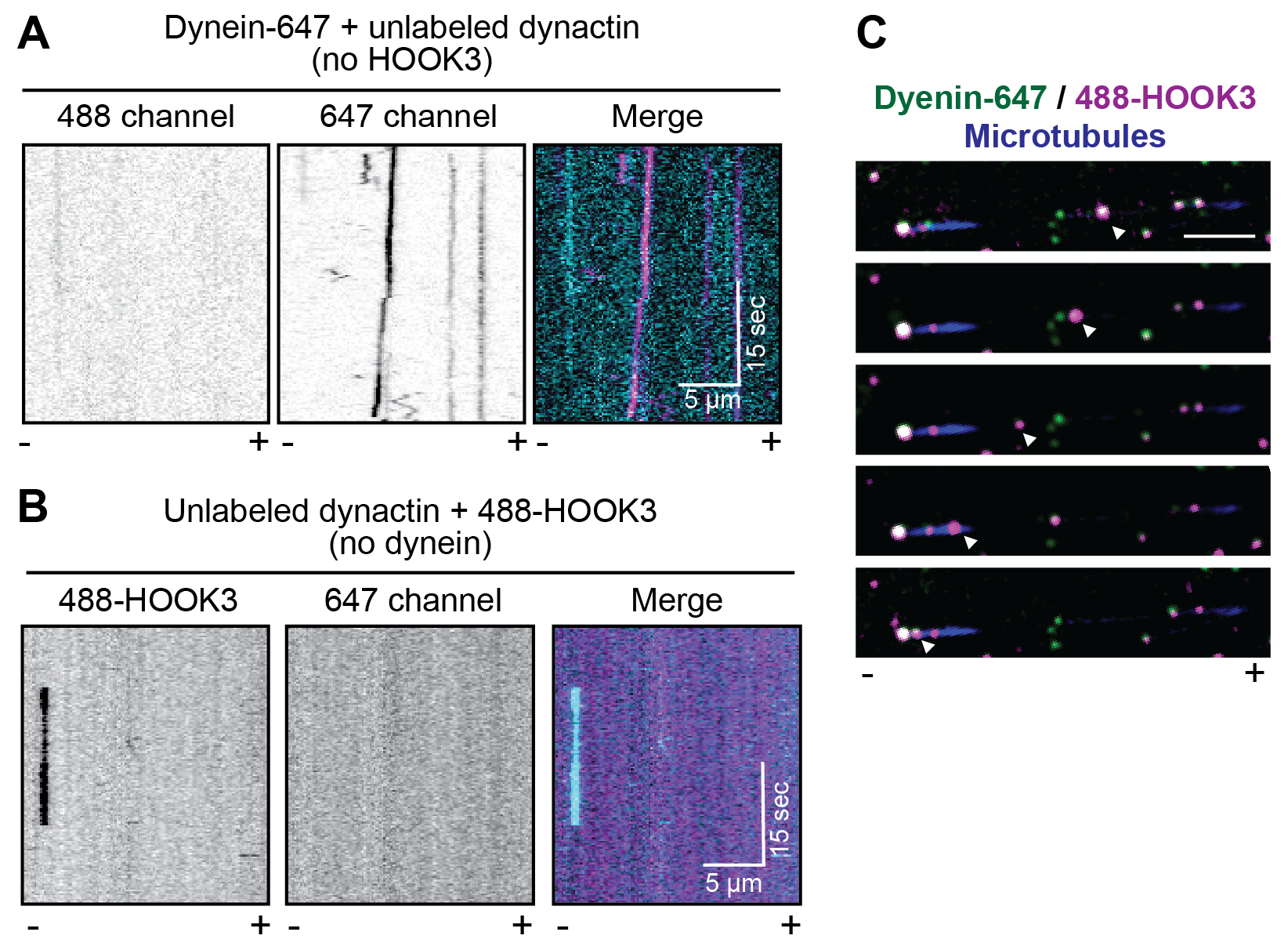
**(A)** Representative kymographs from single-molecule motility assays with dynein-647 and unlabeled dynactin in the absence of HOOK3. Microtubule polarity is marked with minus (-) and plus (+). **(B)** Representative kymographs from singlemolecule motility assays with unlabeled dynactin and HOOK3-488 in the absence of dynein. Microtubule polarity is marked with minus (-) and plus (+). **(C)** Dynein-647 (green) motility in the presence of full-length HOOK3-488 (magenta) and dynactin on polarity marked microtubules. The blue seed made with GMPCPP tubulin marks the microtubule minus end. Scale bar is 2 μm.

**Figure S4.**
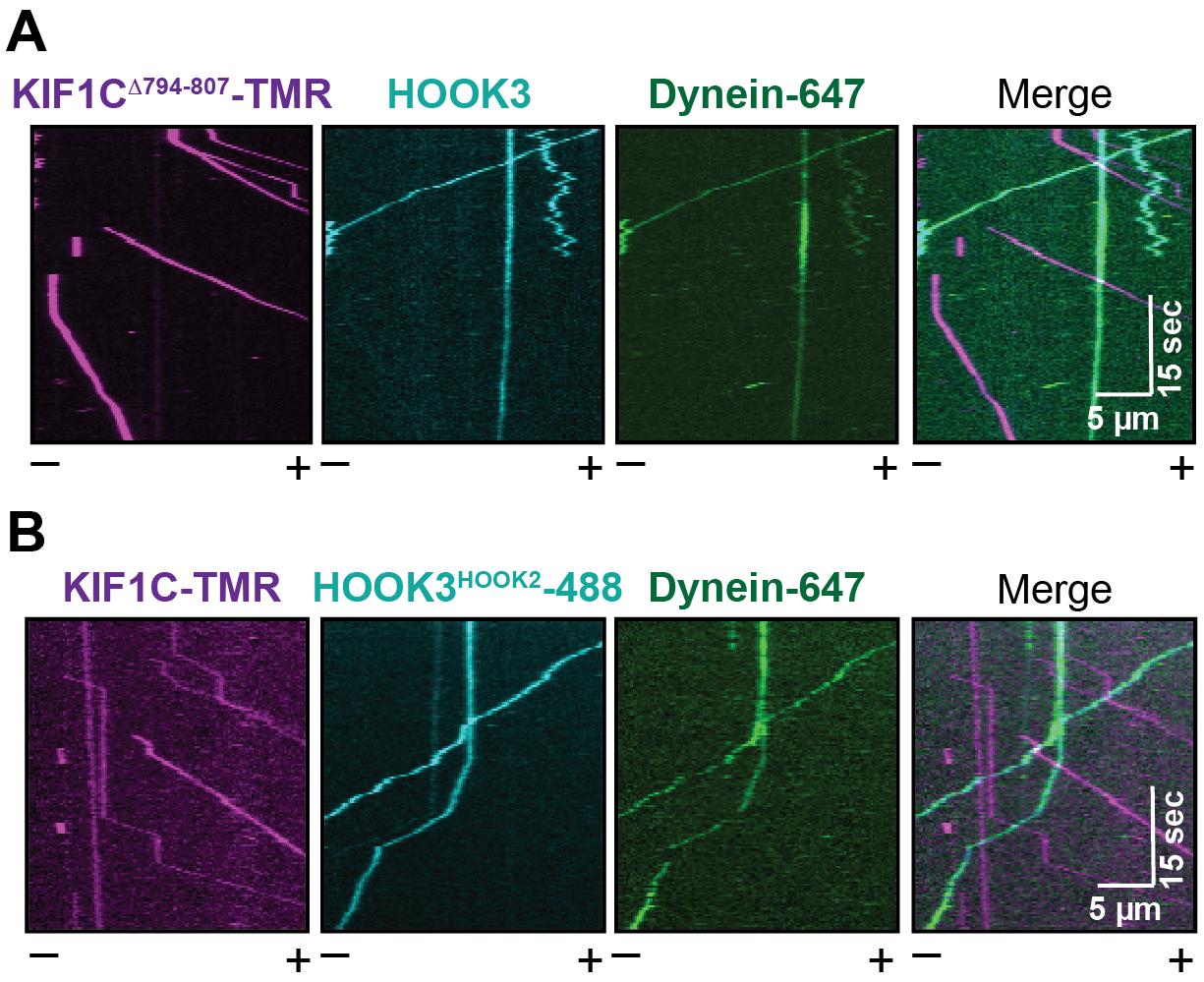
**(A)** Representative kymographs from single-molecule motility assays with purified dynein-647, unlabeled dynactin, KIF1C^Δ794-807^-TMR, and 488-HOOK3. Microtubule polarity is marked with minus (-) and plus (+). **(B)** Representative kymographs from single-molecule motility assays with purified dynein-647, unlabeled dynactin, KIF1C-TMR, and HOOK3^HOOK2^-488 chimera. Microtubule polarity is marked with minus (-) and plus (+).

