## Supplementary material for "HOOK3 is a scaffold for the opposite-polarity microtubule-based motors cytoplasmic dynein and KIF1C": Table S1

| | | | | distrubuted normalized spectral abundance factor (dNSAF) | | | | | | | | | | Spectral Count | | | | | | | | | |
|---|---|---|---|---|---|---|---|---|---|---|---|---|---|---|---|---|---|---|---|---|---|---|---|
| | | | | BioID-3XFLAG | | | | | KIF1C-BioID-3XFLAG | | | | | BioID-3XFLAG | | | | | KIF1C-BioID-3XFLAG | | | | |
| Accession Number | Description | Enrichement | p-value | dNSF BioID S1 | dNSF BioID S2 | dNSF BioID S3 | dNSFBioID S4 | Average dNSF BioID | dNSF KIF1C S1 | dNSF KIF1C S2 | dNSF KIF1C S3 | dNSF KIF1C S4 | dNSF Average KIF1C | SC BioID S1 | SC BioID S2 | SC BioID S3 | SC BioID S4 | SC Average BioID | SC KIF1C S1 | SC KIF1C S2 | SC KIF1C S3 | SC KIF1C S4 | SC Average KIF1C |
| HOOK3 | Protein Hook | 0 | 4.6067E-05 | 0 | 0 | 0 | | 0 | 0.0029553 | 0.0029741 | 0.0042708 | 0.0030702 | 0.0033176 | 0 | 0 | 0 | 0 | 0 | 63 | 63 | 73 | 65 | 66 |
| TDRD3 | Tudor domai | 0 | 0.00024455 | 0 | 0 | 0 | 0 | 0 | 0.00072434 | 0.00072894 | 0.00045168 | 0.00088563 | 0.00069765 | 0 | 0 | 0 | 0 | 0 | 14 | 14 | 7 | 17 | 13 |
| TUT7 | Terminal uric | 0 | 2.8876E-05 | 0 | 0 | 0 | 0 | 0 | 0.000383 | 0.00029474 | 0.00025288 | 0.00034028 | 0.00031773 | 0 | 0 | 0 | 0 | 0 | 17 | 13 | 9 | 15 | 13.5 |
| ANR26 | Ankyrin repe | 0 | 0.00076288 | 0 | 0 | 0 | 0 | 0 | 0.00037446 | 0.000238 | 0.00017206 | 0.00027782 | 0.00026559 | 0 | 0 | 0 | 0 | 0 | 19 | 12 | 7 | 14 | 13 |
| CASC3 | Protein CASC | 0 | 0.00047097 | 0 | 0 | 0 | 0 | 0 | 0.00071867 | 0.0011572 | 0.00059753 | 0.00096485 | 0.00085956 | 0 | 0 | 0 | 0 | 0 | 15 | 24 | 10 | 20 | 17.25 |
| CENPE | Centromere- | 0 | 3.7384E-05 | 0 | 0 | 0 | 0 | 0 | 0.00009976 | 0.00013804 | 9.3312E-05 | 0.00012556 | 0.00011417 | 0 | 0 | 0 | 0 | 0 | 8 | 11 | 6 | 10 | 8.75 |
| YTHD3 | YTH domain- | 0 | 0.002903 | 0 | 0 | 0 | 0 | 0 | 0.00086363 | 0.00057941 | 0.00028722 | 0.00081163 | 0.00063547 | 0 | 0 | 0 | 0 | 0 | 21 | 17 | 9 | 20 | 16.75 |
| RL17 | 60S ribosoma | 0 | 0.00154114 | 0 | 0 | 0 | 0 | 0 | 0.00073221 | 0.00036843 | 0.00091318 | 0.00055295 | 0.00064169 | 0 | 0 | 0 | 0 | 0 | 4 | 2 | 4 | 3 | 3.25 |
| TDRD7 | Tudor domai | 0 | 0.00058416 | 0 | 0 | 0 | 0 | 0 | 0.00033743 | 0.00024696 | 0.00015303 | 0.00027799 | 0.00025385 | 0 | 0 | 0 | 0 | 0 | 11 | 8 | 4 | 9 | 8 |
| AKTIP | AKT-interacti | 0 | 0.00012672 | 0 | 0 | 0 | 0 | 0 | 0.0018456 | 0.0010447 | 0.0015824 | 0.0013937 | 0.0014666 | 0 | 0 | 0 | 0 | 0 | 16 | 9 | 11 | 12 | 12 |
| CCDC9 | Coiled-coil dc | 0 | 0.00029564 | 0 | 0 | 0 | 0 | 0 | 0.00050744 | 0.000383 | 0.00039554 | 0.00025548 | 0.00038537 | 0 | 0 | 0 | 0 | 0 | 8 | 6 | 5 | 4 | 5.75 |
| MEX3A | RNA-binding | 0 | 0.00010073 | 0 | 0 | 0 | 0 | 0 | 0.00064772 | 0.0003911 | 0.00064625 | 0.0006522 | 0.00058432 | 0 | 0 | 0 | 0 | 0 | 10 | 6 | 8 | 10 | 8.5 |
| CTIF | CBP80/20-de | 0 | 0.00417286 | 0 | 0 | 0 | 0 | 0 | 0.00084486 | 0.00073686 | 0.00028098 | 0.0004537 | 0.0005791 | 0 | 0 | 0 | 0 | 0 | 15 | 13 | 4 | 8 | 10 |
| 1433Z | 14-3-3 protei | 0 | 0.00023323 | 0 | 0 | 0 | 0 | 0 | 0.00068738 | 0.0008301 | 0.0010287 | 0.0012458 | 0.000948 | 0 | 0 | 0 | 0 | 0 | 7 | 8 | 8 | 13 | 9 |
| IF4E2 | Eukaryotic tr | 0 | 4.9116E-06 | 0 | 0 | 0 | 0 | 0 | 0.0010998 | 0.00096845 | 0.00085727 | 0.00083056 | 0.00093902 | 0 | 0 | 0 | 0 | 0 | 8 | 7 | 5 | 6 | 6.5 |
| HORN | Hornerin OS= | 0 | 0.00352874 | 0 | 0 | 0 | 0 | 0 | 0.00010636 | 0.00003568 | 8.8434E-05 | 5.9499E-05 | 7.2493E-05 | 0 | 0 | 0 | 0 | 0 | 9 | 3 | 6 | 5 | 5.75 |
| PURB | Transcription | 0 | 0.01934282 | 0 | 0 | 0 | 0 | 0 | 0.00021591 | 0.00032592 | 0.00080781 | 0.0003261 | 0.00041894 | 0 | 0 | 0 | 0 | 0 | 2 | 3 | 6 | 3 | 3.5 |
| E2AK2 | Interferon-in | 0 | 0.00010893 | 0 | 0 | 0 | 0 | 0 | 0.00024451 | 0.00018455 | 0.00015247 | 0.0002462 | 0.00020693 | 0 | 0 | 0 | 0 | 0 | 4 | 3 | 2 | 4 | 3.25 |
| RHEB | GTP-binding | 0 | 0.01012919 | 0 | 0 | 0 | 0 | 0 | 0.0003661 | 0.00036843 | 0.00091318 | 0.00036864 | 0.00050409 | 0 | 0 | 0 | 0 | 0 | 2 | 2 | 4 | 2 | 2.5 |
| NAA10 | N-alpha-acet | 0 | 5.8888E-05 | 0 | 0 | 0 | 0 | 0 | 0.0005733 | 0.00043271 | 0.0003575 | 0.00043295 | 0.00044912 | 0 | 0 | 0 | 0 | 0 | 4 | 3 | 2 | 3 | 3 |
| F16A2 | FTS and Hool | 0 | 9.2734E-05 | 0 | 0 | 0 | 0 | 0 | 0.00024256 | 0.00020923 | 0.00034573 | 0.00027913 | 0.00026916 | 0 | 0 | 0 | 0 | 0 | 7 | 6 | 8 | 8 | 7.25 |
| BIEA | Biliverdin red | 0 | 0.00070432 | 0 | 0 | 0 | 0 | 0 | 0.00022758 | 0.00034354 | 0.00042574 | 0.00022915 | 0.0003065 | 0 | 0 | 0 | 0 | 0 | 2 | 3 | 3 | 2 | 2.5 |
| RL29 | 60S ribosoma | 0 | 0.00475937 | 0 | 0 | 0 | 0 | 0 | 0.0010592 | 0.00042636 | 0.0015851 | 0.0010665 | 0.00103429 | 0 | 0 | 0 | 0 | 0 | 5 | 2 | 6 | 5 | 4.5 |
| ABCB7 | ATP-binding | 0 | 0.00094627 | 0 | 0 | 0 | 0 | 0 | 0.00013437 | 0.00013522 | 0.00011172 | 0.00022549 | 0.0001517 | 0 | 0 | 0 | 0 | 0 | 3 | 3 | 2 | 5 | 3.25 |
| G3P | Glyceraldehy | 0 | 5.8926E-05 | 0 | 0 | 0 | 0 | 0 | 0.00040217 | 0.00030354 | 0.00050157 | 0.00040495 | 0.00040306 | 0 | 0 | 0 | 0 | 0 | 4 | 3 | 4 | 4 | 3.75 |
| GPBP1 | Vasculin OS= | 0 | 0.00082466 | 0 | 0 | 0 | 0 | 0 | 0.00042725 | 0.0003583 | 0.00017762 | 0.0003585 | 0.00033042 | 0 | 0 | 0 | 0 | 0 | 6 | 5 | 2 | 5 | 4.5 |
| RT11 | 28S ribosoma | 0 | 0.00017434 | 0 | 0 | 0 | 0 | 0 | 0.00034723 | 0.00052416 | 0.00064958 | 0.00052445 | 0.00051136 | 0 | 0 | 0 | 0 | 0 | 2 | 3 | 3 | 3 | 2.75 |
| RAF1 | RAF proto-or | 0 | 6.5958E-05 | 0 | 0 | 0 | 0 | 0 | 0.00015593 | 0.00020923 | 0.00012965 | 0.00015701 | 0.00016296 | 0 | 0 | 0 | 0 | 0 | 3 | 4 | 2 | 3 | 3 |
| 1433G | 14-3-3 protei | 0 | 2.199E-06 | 0 | 0 | 0 | 0 | 0 | 0.00040909 | 0.00041169 | 0.0005102 | 0.00041192 | 0.00043573 | 0 | 0 | 0 | 0 | 0 | 5 | 5 | 5 | 7 | 5.5 |
| ERLN1 | Erlin-1 OS=H | 0 | 0.00164118 | 0 | 0 | 0 | 0 | 0 | 0.00029204 | 0.00019593 | 0.00036422 | 0.00049009 | 0.00033557 | 0 | 0 | 0 | 0 | 0 | 5 | 3 | 4 | 7 | 4.75 |
| ARAF | Serine/threo | 0 | 0.00070625 | 0 | 0 | 0 | 0 | 0 | 0.00011116 | 0.00011187 | 0.00020795 | 0.00016789 | 0.00014972 | 0 | 0 | 0 | 0 | 0 | 2 | 2 | 3 | 3 | 2.5 |
| AAKG1 | 5'-AMP-activ | 0 | 3.3835E-05 | 0 | 0 | 0 | 0 | 0 | 0.00030527 | 0.00020481 | 0.00025381 | 0.00030738 | 0.00026782 | 0 | 0 | 0 | 0 | 0 | 3 | 2 | 2 | 3 | 2.5 |
| CE164 | Centrosomal | 0 | 0.0250707 | 0 | 0 | 0 | 0 | 0 | 0.00055367 | 0.00046432 | 0 | 0.00048781 | 0.00050193 | 0 | 0 | 0 | 0 | 0 | 24 | 20 | 0 | 21 | 21.6666667 |
| SMG7 | Protein SMG | 0 | 0.04144503 | 0 | 0 | 0 | 0 | 0 | 0.00032586 | 0.00038755 | 0 | 0.00017897 | 0.00029746 | 0 | 0 | 0 | 0 | 0 | 11 | 13 | 0 | 6 | 10 |
| SMG8 | Protein SMG | 0 | 0.03127906 | 0 | 0 | 0 | 0 | 0 | 0.0002719 | 0.00017102 | 0 | 0.00020533 | 0.00021608 | 0 | 0 | 0 | 0 | 0 | 8 | 5 | 0 | 6 | 6.33333333 |
| CKAP2 | Cytoskeleton | 0 | 0.07211 | 0 | 0 | 0 | 0 | 0 | 9.8628E-05 | 0.00024814 | 0 | 0.00009931 | 0.00014869 | 0 | 0 | 0 | 0 | 0 | 2 | 5 | 0 | 2 | 3 |
| RL32 | 60S ribosoma | 0 | 0.02612185 | 0 | 0 | 0 | 0 | 0 | 0.00049899 | 0 | 0.00062231 | 0.00050244 | 0.00054125 | 0 | 0 | 0 | 0 | 0 | 2 | 0 | 2 | 2 | 2 |
| DYR | Dihydrofolat | 0 | 0.02612741 | 0 | 0 | 0 | 0 | 0 | 0.00036023 | 0.00036252 | 0.00044926 | 0 | 0.00039067 | 0 | 0 | 0 | 0 | 0 | 2 | 2 | 2 | 0 | 2 |
| RAB14 | Ras-related p | 0 | 0.03182713 | 0 | 0 | 0 | 0 | 0 | 0.00046998 | 0.00063062 | 0.00039076 | 0 | 0.00049712 | 0 | 0 | 0 | 0 | 0 | 7 | 6 | 5 | 0 | 6 |
| TIPRL | TIP41-like pr | 0 | 0.02401 | 0 | 0 | 0 | 0 | 0 | 0.00049532 | 0.00049846 | 0 | 0.00049874 | 0.00049751 | 0 | 0 | 0 | 0 | 0 | 4 | 4 | 0 | 4 | 4 |
| LAR4B | La-related pr | 0 | 0.02928142 | 0 | 0 | 0 | 0 | 0 | 0.00018256 | 0.00022964 | 0 | 0.00027573 | 0.00022931 | 0 | 0 | 0 | 0 | 0 | 4 | 5 | 0 | 6 | 5 |
| AGK | Acylglycerol l | 0 | 0.02994198 | 0 | 0 | 0 | 0 | 0 | 0.00023944 | 0.00016064 | 0 | 0.0002411 | 0.00021373 | 0 | 0 | 0 | 0 | 0 | 3 | 2 | 0 | 3 | 2.66666667 |
| PATL1 | Protein PAT1 | 0 | 0.04593411 | 0 | 0 | 0 | 0 | 0 | 0.00021871 | 0.00013206 | 0 | 0.00030831 | 0.00021969 | 0 | 0 | 0 | 0 | 0 | 5 | 3 | 0 | 7 | 5 |
| COPG2 | Coatomer su | 0 | 0.03216146 | 0 | 0 | 0 | 0 | 0 | 0.00015468 | 0.00011675 | 0 | 0.00019469 | 0.00015537 | 0 | 0 | 0 | 0 | 0 | 7 | 3 | 0 | 8 | 6 |
| HAUS2 | HAUS augmir | 0 | 0.07839483 | 0 | 0 | 0 | 0 | 0 | 0 | 0.00028847 | 0.0010725 | 0.00057727 | 0.00064608 | 0 | 0 | 0 | 0 | 0 | 0 | 2 | 6 | 4 | 4 |
| MCRI1 | Mapk-regula | 0 | 0.02727894 | 0 | 0 | 0 | 0 | 0 | 0.0010417 | 0.0013978 | 0 | 0.0013985 | 0.00127933 | 0 | 0 | 0 | 0 | 0 | 3 | 4 | 0 | 4 | 3.66666667 |
| RC3H1 | Roquin-1 OS= | 0 | 0.03491281 | 0 | 0 | 0 | 0 | 0 | 0.00014864 | 0.00026925 | 0 | 0.00020953 | 0.00020914 | 0 | 0 | 0 | 0 | 0 | 5 | 9 | 0 | 7 | 7 |
| ODPB | Pyruvate deh | 0 | 0.02762863 | 0 | 0 | 0 | 0 | 0 | 0.00037528 | 0.00028325 | 0 | 0.00028341 | 0.00031398 | 0 | 0 | 0 | 0 | 0 | 4 | 3 | 0 | 3 | 3.33333333 |
| DYLT1 | Dynein light | 0 | 0.07388278 | 0 | 0 | 0 | 0 | 0 | 0.00059613 | 0 | 0.0018587 | 0.00090038 | 0.0011184 | 0 | 0 | 0 | 0 | 0 | 2 | 0 | 5 | 3 | 3.33333333 |
| CDC20 | Cell division | 0 | 0.03332668 | 0 | 0 | 0 | 0 | 0 | 0.00020249 | 0.00033964 | 0.00033672 | 0 | 0.00029295 | 0 | 0 | 0 | 0 | 0 | 3 | 5 | 4 | 0 | 4 |
| EST1A | Telomerase- | 0 | 0.04444109 | 0 | 0 | 0 | 0 | 0 | 0.00016615 | 7.1661E-05 | 0 | 0.0001434 | 0.00012707 | 0 | 0 | 0 | 0 | 0 | 7 | 3 | 0 | 6 | 5.33333333 |
| QTRT2 | Queuine tRN | 0 | 0.03188172 | 0 | 0 | 0 | 0 | 0 | 0.0004058 | 0.00024503 | 0 | 0.00032689 | 0.00032591 | 0 | 0 | 0 | 0 | 0 | 5 | 3 | 0 | 4 | 4 |
| ABC3C | DNA dC->dU- | 0 | 0.03663395 | 0 | 0 | 0 | 0 | 0 | 0 | 0.00053519 | 0.00088434 | 0.00053549 | 0.00065167 | 0 | 0 | 0 | 0 | 0 | 0 | 3 | 4 | 3 | 3.33333333 |
| KLC2 | Kinesin light | 0 | 0.03562683 | 0 | 0 | 0 | 0 | 0 | 0.00016245 | 0.00010899 | 0.0002026 | 0 | 0.00015801 | 0 | 0 | 0 | 0 | 0 | 3 | 2 | 3 | 0 | 2.66666667 |
| MCRI2 | MAPK regula | 0 | 0.02508594 | 0 | 0 | 0 | 0 | 0 | 0.0012631 | 0.0014829 | 0 | 0.0012718 | 0.00133927 | 0 | 0 | 0 | 0 | 0 | 6 | 7 | 0 | 6 | 6.33333333 |
| SNX30 | Sorting nexin | 0 | 0.03165577 | 0 | 0 | 0 | 0 | 0 | 0.00023122 | 0.00015513 | 0 | 0.00015521 | 0.00018052 | 0 | 0 | 0 | 0 | 0 | 3 | 2 | 0 | 2 | 2.33333333 |
| LS14B | Protein LSM1 | 0 | 0.0270638 | 0 | 0 | 0 | 0 | 0 | 0.00034994 | 0.00035216 | 0 | 0.00026427 | 0.00032212 | 0 | 0 | 0 | 0 | 0 | 4 | 4 | 0 | 3 | 3.66666667 |
| TUT4 | Terminal uric | 0 | 0.03849121 | 0 | 0 | 0 | 0 | 0 | 0.00008195 | 4.1236E-05 | 0 | 6.1888E-05 | 6.1691E-05 | 0 | 0 | 0 | 0 | 0 | 4 | 2 | 0 | 3 | 3 |
| DHX40 | Probable ATF | 0 | 0.02401001 | 0 | 0 | 0 | 0 | 0 | 8.6474E-05 | 8.7023E-05 | 0 | 8.7072E-05 | 8.6856E-05 | 0 | 0 | 0 | 0 | 0 | 2 | 2 | 0 | 2 | 2 |
| PAN2 | PAN2-PAN3 c | 0 | 0.03228981 | 0 | 0 | 0 | 0 | 0 | 8.4064E-05 | 0.000141 | 0 | 0.00011286 | 0.00011264 | 0 | 0 | 0 | 0 | 0 | 3 | 5 | 0 | 4 | 4 |
| ARF | Tumor suppr | 0 | 0.03198939 | 0 | 0 | 0 | 0 | 0 | 0.00076549 | 0 | 0.0012729 | 0.0010277 | 0.00102203 | 0 | 0 | 0 | 0 | 0 | 3 | 0 | 4 | 4 | 3.66666667 |
| RB6I2 | ELKS/Rab6-in | 0 | 0.07174999 | 0 | 0 | 0 | 0 | 0 | 0.00021126 | 6.0745E-05 | 0 | 0.00012156 | 0.00013119 | 0 | 0 | 0 | 0 | 0 | 7 | 2 | 0 | 4 | 4.33333333 |
| PCID2 | PCI domain-c | 0 | 0.03870833 | 0 | 0 | 0 | 0 | 0 | 0.00033766 | 0.0001699 | 0.00031584 | 0 | 0.00027447 | 0 | 0 | 0 | 0 | 0 | 4 | 2 | 3 | 0 | 3 |
| KCC2G | Calcium/calm | 0 | 0.02913339 | 0 | 0 | 0 | 0 | 0 | 0 | 0.00018223 | 0.00015056 | 0.00012156 | 0.00015145 | 0 | 0 | 0 | 0 | 0 | 0 | 3 | 3 | 2 | 2.66666667 |
| KIF7 | Kinesin-like p | 0 | 0.02401 | 0 | 0 | 0 | 0 | 0 | 5.0159E-05 | 5.0477E-05 | 0 | 5.0505E-05 | 5.038E-05 | 0 | 0 | 0 | 0 | 0 | 2 | 2 | 0 | 2 | 2 |
| OFD1 | Oral-facial-di | 0 | 0.04290405 | 0 | 0 | 0 | 0 | 0 | 9.9847E-05 | 0.00013397 | 0.00020754 | 0 | 0.00014712 | 0 | 0 | 0 | 0 | 0 | 3 | 4 | 5 | 0 | 4 |
| CDIPT | CDP-diacylgly | 0 | 0.02612173 | 0 | 0 | 0 | 0 | 0 | 0.00031626 | 0 | 0.00039442 | 0.00031845 | 0.00034304 | 0 | 0 | 0 | 0 | 0 | 2 | 0 | 2 | 2 | 2 |
| HYOU1 | Hypoxia up-r | 0 | 0.07113491 | 0 | 0 | 0 | 0 | 0 | 6.7431E-05 | 0 | 0.00016819 | 6.7897E-05 | 0.00010117 | 0 | 0 | 0 | 0 | 0 | 2 | 0 | 4 | 2 | 2.66666667 |
| K1C17 | Keratin, type | 0 | 0.11283996 | 0 | 0 | 0 | 0 | 0 | 0.00031187 | 0 | 0.00077789 | 0.00015701 | 0.00041559 | 0 | 0 | 0 | 0 | 0 | 10 | 0 | 15 | 8 | 11 |
| CNBP | Cellular nucle | 0 | 0.02991826 | 0 | 0 | 0 | 0 | 0 | 0.00057087 | 0.0005745 | 0 | 0.00038321 | 0.00050953 | 0 | 0 | 0 | 0 | 0 | 3 | 3 | 0 | 2 | 2.66666667 |
| TBA1A | Tubulin alpha | 0 | 0.0443631 | 0 | 0 | 0 | 0 | 0 | 0.00014936 | 0 | 0.00027942 | 0.0001504 | 0.00019306 | 0 | 0 | 0 | 0 | 0 | 201 | 0 | 145 | 178 | 174.666667 |
| 1433B | 14-3-3 protei | 0 | 0.04438408 | 0 | 0 | 0 | 0 | 0 | 0.00027383 | 0.00027557 | 0.00051227 | 0 | 0.00035389 | 0 | 0 | 0 | 0 | 0 | 5 | 4 | 7 | 0 | 5.33333333 |
| GSH0 | Glutamate--c | 0 | 0.02789358 | 0 | 0 | 0 | 0 | 0 | 0.00036878 | 0.00049483 | 0 | 0.00037133 | 0.00041165 | 0 | 0 | 0 | 0 | 0 | 3 | 4 | 0 | 3 | 3.33333333 |
| NUB1 | NEDD8 ultim | 0 | 0.03199234 | 0 | 0 | 0 | 0 | 0 | 0.0001643 | 0.00022046 | 0.00027321 | 0 | 0.00021932 | 0 | 0 | 0 | 0 | 0 | 3 | 4 | 4 | 0 | 3.66666667 |
| AGO3 | Protein argor | 0 | 0.03205652 | 0 | 0 | 0 | 0 | 0 | 7.8329E-05 | 0.00011824 | 0 | 7.8871E-05 | 9.1813E-05 | 0 | 0 | 0 | 0 | 0 | 8 | 10 | 0 | 7 | 8.33333333 |
| IPO5 | Importin-5 O | 0 | 0.18899663 | 0 | 0 | 0 | 0 | 0 | 0 | 0.00015449 | 0 | 6.1831E-05 | 0.00010816 | 0 | 0 | 0 | 0 | 0 | 0 | 5 | 0 | 2 | 3.5 |
| NSA2 | Ribosome bio | 0 | 0.1339784 | 0 | 0 | 0 | 0 | 0 | 0.00025909 | 0 | 0 | 0.00026088 | 0.00025999 | 0 | 0 | 0 | 0 | 0 | 2 | 0 | 0 | 2 | 2 |
| GSK3B | Glycogen syn | 0 | 0.14045077 | 0 | 0 | 0 | 0 | 0 | 0 | 0.00032282 | 0 | 0.00024225 | 0.00028254 | 0 | 0 | 0 | 0 | 0 | 0 | 4 | 0 | 3 | 3.5 |
| BZW1 | Basic leucine | 0 | 0.18821415 | 0 | 0 | 0 | 0 | 0 | 0 | 0.00016179 | 0.00040101 | 0 | 0.0002814 | 0 | 0 | 0 | 0 | 0 | 0 | 2 | 4 | 0 | 3 |
| PAPD1 | Poly(A) RNA | 0 | 0.16254263 | 0 | 0 | 0 | 0 | 0 | 0.00011574 | 0 | 0.00021653 | 0 | 0.00016614 | 0 | 0 | 0 | 0 | 0 | 2 | 0 | 3 | 0 | 2.5 |
| WASL | Neural Wiskc | 0 | 0.16827638 | 0 | 0 | 0 | 0 | 0 | 0 | 0.00013424 | 0 | 0.00026863 | 0.00020144 | 0 | 0 | 0 | 0 | 0 | 0 | 2 | 0 | 4 | 3 |
| RL38 | 60S ribosoma | 0 | 0.14862043 | 0 | 0 | 0 | 0 | 0 | 0 | 0.0019369 | 0.0030004 | 0 | 0.00246865 | 0 | 0 | 0 | 0 | 0 | 0 | 4 | 5 | 0 | 4.5 |
| INT7 | Integrator co | 0 | 0.1339784 | 0 | 0 | 0 | 0 | 0 | 7.0024E-05 | 0 | 0 | 7.0508E-05 | 7.0266E-05 | 0 | 0 | 0 | 0 | 0 | 2 | 0 | 0 | 2 | 2 |
| MP2K1 | Dual specifici | 0 | 0.14666198 | 0 | 0 | 0 | 0 | 0 | 0 | 0.0001725 | 0 | 0.00025889 | 0.0002157 | 0 | 0 | 0 | 0 | 0 | 0 | 2 | 0 | 3 | 2.5 |
| RS15 | 40S ribosoma | 0 | 0.13783908 | 0 | 0 | 0 | 0 | 0 | 0.00092915 | 0 | 0.0011588 | 0 | 0.00104398 | 0 | 0 | 0 | 0 | 0 | 4 | 0 | 4 | 0 | 4 |
| CNOT8 | CCR4-NOT tr | 0 | 0.18882856 | 0 | 0 | 0 | 0 | 0 | 0.0002307 | 0 | 0.00057543 | 0 | 0.00040307 | 0 | 0 | 0 | 0 | 0 | 2 | 0 | 4 | 0 | 3 |
| PJA2 | E3 ubiquitin- | 0 | 0.14017206 | 0 | 0 | 0 | 0 | 0 | 0.00019029 | 0 | 0 | 0.00014371 | 0.000167 | 0 | 0 | 0 | 0 | 0 | 4 | 0 | 0 | 3 | 3.5 |
| NDC80 | Kinetochore | 0 | 0.15089671 | 0 | 0 | 0 | 0 | 0 | 0.00020985 | 0 | 0.00013086 | 0 | 0.00017036 | 0 | 0 | 0 | 0 | 0 | 4 | 0 | 2 | 0 | 3 |
| GTSE1 | G2 and S pha | 0 | 0.14017388 | 0 | 0 | 0 | 0 | 0 | 0.00018712 | 0 | 0 | 0.00014131 | 0.00016422 | 0 | 0 | 0 | 0 | 0 | 4 | 0 | 0 | 3 | 3.5 |
| ROAA | Heterogeneo | 0 | 0.13783921 | 0 | 0 | 0 | 0 | 0 | 0.0002029 | 0 | 0.00025305 | 0 | 0.00022798 | 0 | 0 | 0 | 0 | 0 | 6 | 0 | 4 | 0 | 5 |
| ARF5 | ADP-ribosyla | 0 | 0.13397841 | 0 | 0 | 0 | 0 | 0 | 0.00037424 | 0 | 0 | 0.00037683 | 0.00037554 | 0 | 0 | 0 | 0 | 0 | 16 | 0 | 0 | 20 | 18 |
| EI2BA | Translation ir | 0 | 0.13687149 | 0 | 0 | 0 | 0 | 0 | 0 | 0.0003334 | 0.00027545 | 0 | 0.00030443 | 0 | 0 | 0 | 0 | 0 | 0 | 3 | 2 | 0 | 2.5 |
| LAR1B | La-related pr | 0 | 0.14625137 | 0 | 0 | 0 | 0 | 0 | 0.00011055 | 0.00007417 | 0 | 0 | 0.00009236 | 0 | 0 | 0 | 0 | 0 | 4 | 4 | 0 | 0 | 4 |
| HERC1 | Probable E3 | 0 | 0.15309601 | 0 | 0 | 0 | 0 | 0 | 3.4645E-05 | 0 | 0 | 2.0931E-05 | 2.7788E-05 | 0 | 0 | 0 | 0 | 0 | 5 | 0 | 0 | 3 | 4 |
| QCR1 | Cytochrome | 0 | 0.13397462 | 0 | 0 | 0 | 0 | 0 | 0 | 0.00021185 | 0 | 0.00021197 | 0.00021191 | 0 | 0 | 0 | 0 | 0 | 0 | 3 | 0 | 3 | 3 |
| FBX7 | F-box only pr | 0 | 0.16883069 | 0 | 0 | 0 | 0 | 0 | 0.00012905 | 0 | 0 | 0.00025988 | 0.00019447 | 0 | 0 | 0 | 0 | 0 | 2 | 0 | 0 | 4 | 3 |
| NDUS7 | NADH dehyd | 0 | 0.14625264 | 0 | 0 | 0 | 0 | 0 | 0.00047439 | 0.00031827 | 0 | 0 | 0.00039633 | 0 | 0 | 0 | 0 | 0 | 3 | 2 | 0 | 0 | 2.5 |
| F16B1 | Protein FAM | 0 | 0.13397462 | 0 | 0 | 0 | 0 | 0 | 0 | 8.8616E-05 | 0 | 8.8665E-05 | 8.8641E-05 | 0 | 0 | 0 | 0 | 0 | 0 | 2 | 0 | 2 | 2 |
| YBOX3 | Y-box-bindin | 0 | 0.13397845 | 0 | 0 | 0 | 0 | 0 | 0.00018108 | 0 | 0 | 0.00018234 | 0.00018171 | 0 | 0 | 0 | 0 | 0 | 5 | 0 | 0 | 3 | 4 |
| RC3H2 | Roquin-2 OS= | 0 | 0.18899875 | 0 | 0 | 0 | 0 | 0 | 0 | 0.0001423 | 0 | 5.6951E-05 | 9.9626E-05 | 0 | 0 | 0 | 0 | 0 | 0 | 5 | 0 | 2 | 3.5 |
| MCA3 | Eukaryotic tr | 0 | 0.14701241 | 0 | 0 | 0 | 0 | 0 | 0.00038714 | 0.00058441 | 0 | 0 | 0.00048578 | 0 | 0 | 0 | 0 | 0 | 2 | 3 | 0 | 0 | 2.5 |
| RM16 | 39S ribosoma | 0 | 0.16762771 | 0 | 0 | 0 | 0 | 0 | 0.00053676 | 0 | 0 | 0.00027023 | 0.0004035 | 0 | 0 | 0 | 0 | 0 | 4 | 0 | 0 | 2 | 3 |
| HAUS1 | HAUS augmir | 0 | 0.17546705 | 0 | 0 | 0 | 0 | 0 | 0 | 0 | 0.0010577 | 0.00048798 | 0.00077284 | 0 | 0 | 0 | 0 | 0 | 0 | 0 | 7 | 4 | 5.5 |
| CP062 | UPF0505 pro | 0 | 0.14017397 | 0 | 0 | 0 | 0 | 0 | 0.0001399 | 0 | 0 | 0.00010565 | 0.00012278 | 0 | 0 | 0 | 0 | 0 | 4 | 0 | 0 | 3 | 3.5 |
| ORNT1 | Mitochondria | 0 | 0.13760348 | 0 | 0 | 0 | 0 | 0 | 0 | 0 | 0.00027911 | 0.00022535 | 0.00025223 | 0 | 0 | 0 | 0 | 0 | 0 | 0 | 2 | 2 | 2 |
| RS18 | 40S ribosoma | 0 | 0.16197173 | 0 | 0 | 0 | 0 | 0 | 0 | 0 | 0.00082907 | 0.00044624 | 0.00063766 | 0 | 0 | 0 | 0 | 0 | 0 | 0 | 3 | 2 | 2.5 |
| TNG6 | Transport an | 0 | 0.13760431 | 0 | 0 | 0 | 0 | 0 | 0 | 0 | 7.6794E-05 | 6.2001E-05 | 6.9398E-05 | 0 | 0 | 0 | 0 | 0 | 0 | 0 | 2 | 2 | 2 |
| CC138 | Coiled-coil dc | 0 | 0.13688966 | 0 | 0 | 0 | 0 | 0 | 0 | 0 | 0.00012633 | 0.000153 | 0.00013967 | 0 | 0 | 0 | 0 | 0 | 0 | 0 | 2 | 3 | 2.5 |
| RL26L | 60S ribosoma | 0 | 0.13397842 | 0 | 0 | 0 | 0 | 0 | 0.00046457 | 0 | 0 | 0.00046779 | 0.00046618 | 0 | 0 | 0 | 0 | 0 | 9 | 0 | 0 | 9 | 9 |
| PDIA6 | Protein disul | 0 | 0.13760405 | 0 | 0 | 0 | 0 | 0 | 0 | 0 | 0.00019094 | 0.00015416 | 0.00017255 | 0 | 0 | 0 | 0 | 0 | 0 | 0 | 2 | 2 | 2 |
| PSA | Puromycin-se | 0 | 0.13668383 | 0 | 0 | 0 | 0 | 0 | 0.00010995 | 0 | 9.1417E-05 | 0 | 0.00010068 | 0 | 0 | 0 | 0 | 0 | 3 | 0 | 2 | 0 | 2.5 |
| TT30B | Tetratricope | 0 | 0.13762234 | 0 | 0 | 0 | 0 | 0 | 0 | 0.00010194 | 0.00012633 | 0 | 0.00011414 | 0 | 0 | 0 | 0 | 0 | 0 | 10 | 11 | 0 | 10.5 |
| FACE2 | CAAX prenyl | 0 | 0.14701205 | 0 | 0 | 0 | 0 | 0 | 0.00020475 | 0.00030908 | 0 | 0 | 0.00025692 | 0 | 0 | 0 | 0 | 0 | 2 | 3 | 0 | 0 | 2.5 |
| ACTS | Actin, alpha s | 0 | 0.35591768 | 0 | 0 | 0 | 0 | 0 | 0.00044671 | 0 | 0 | 0 | 0.00044671 | 0 | 0 | 0 | 0 | 0 | 11 | 0 | 0 | 0 | 11 |
| SPICE | Spindle and c | 0 | 0.35591768 | 0 | 0 | 0 | 0 | 0 | 0 | 0.00015858 | 0 | 0 | 0.00015858 | 0 | 0 | 0 | 0 | 0 | 0 | 4 | 0 | 0 | 4 |
| KIF1A | Kinesin-like p | 0 | 0.35591768 | 0 | 0 | 0 | 0 | 0 | 0 | 0 | 9.9423E-05 | 0 | 9.9423E-05 | 0 | 0 | 0 | 0 | 0 | 0 | 0 | 98 | 0 | 98 |
| RBMS1 | RNA-binding | 0 | 0.35591768 | 0 | 0 | 0 | 0 | 0 | 0.00016592 | 0 | 0 | 0 | 0.00016592 | 0 | 0 | 0 | 0 | 0 | 2 | 0 | 0 | 0 | 2 |
| PEX5 | Peroxisomal | 0 | 0.35591768 | 0 | 0 | 0 | 0 | 0 | 0.00010542 | 0 | 0 | 0 | 0.00010542 | 0 | 0 | 0 | 0 | 0 | 2 | 0 | 0 | 0 | 2 |
| BPL1 | Biotin--prote | 0 | 0.35591768 | 0 | 0 | 0 | 0 | 0 | 9.2787E-05 | 0 | 0 | 0 | 9.2787E-05 | 0 | 0 | 0 | 0 | 0 | 2 | 0 | 0 | 0 | 2 |
| RAB1B | Ras-related p | 0 | 0.35591768 | 0 | 0 | 0 | 0 | 0 | 0.00050271 | 0 | 0 | 0 | 0.00050271 | 0 | 0 | 0 | 0 | 0 | 9 | 0 | 0 | 0 | 9 |
| DNJB6 | DnaJ homolo | 0 | 0.35591768 | 0 | 0 | 0 | 0 | 0 | 0 | 0.0004159 | 0 | 0 | 0.0004159 | 0 | 0 | 0 | 0 | 0 | 0 | 4 | 0 | 0 | 4 |
| AGO1 | Protein argor | 0 | 0.35591768 | 0 | 0 | 0 | 0 | 0 | 0 | 0 | 0 | 0.00011872 | 0.00011872 | 0 | 0 | 0 | 0 | 0 | 0 | 0 | 0 | 7 | 7 |
| SRP14 | Signal recogr | 0 | 0.35591768 | 0 | 0 | 0 | 0 | 0 | 0.00074298 | 0 | 0 | 0 | 0.00074298 | 0 | 0 | 0 | 0 | 0 | 3 | 0 | 0 | 0 | 3 |
| ALG13 | Putative bifu | 0 | 0.35591768 | 0 | 0 | 0 | 0 | 0 | 0 | 0 | 0.00011083 | 0 | 0.00011083 | 0 | 0 | 0 | 0 | 0 | 0 | 0 | 3 | 0 | 3 |
| DHC24 | Delta(24)-ste | 0 | 0.35591768 | 0 | 0 | 0 | 0 | 0 | 0.00013055 | 0 | 0 | 0 | 0.00013055 | 0 | 0 | 0 | 0 | 0 | 2 | 0 | 0 | 0 | 2 |
| TBA1C | Tubulin alpha | 0 | 0.35591768 | 0 | 0 | 0 | 0 | 0 | 0.00030006 | 0 | 0 | 0 | 0.00030006 | 0 | 0 | 0 | 0 | 0 | 167 | 0 | 0 | 0 | 167 |
| CEP78 | Centrosomal | 0 | 0.35591768 | 0 | 0 | 0 | 0 | 0 | 0 | 0.00014759 | 0 | 0 | 0.00014759 | 0 | 0 | 0 | 0 | 0 | 0 | 3 | 0 | 0 | 3 |
| TM9S3 | Transmembr | 0 | 0.35591768 | 0 | 0 | 0 | 0 | 0 | 0 | 0 | 0 | 0.00017274 | 0.00017274 | 0 | 0 | 0 | 0 | 0 | 0 | 0 | 0 | 3 | 3 |

| | | | | | | | | | | | | | | | | | | | | | | | |
|---|---|---|---|---|---|---|---|---|---|---|---|---|---|---|---|---|---|---|---|---|---|---|---|
| GEN | Flap endonuc | 0 | 0.35591768 | 0 | 0 | 0 | 0 | 0 | 0 | 0.00011199 | 0 | 0 | 0.00011199 | 0 | 0 | 0 | 0 | 0 | 0 | 3 | 0 | 0 | 3 |
| MARK2 | Serine/threo | 0 | 0.35591768 | 0 | 0 | 0 | 0 | 0 | 0 | 8.6029E-05 | 0 | 0 | 8.6029E-05 | 0 | 0 | 0 | 0 | 0 | 0 | 2 | 0 | 0 | 2 |
| SI1L1 | Signal-induce | 0 | 0.35591768 | 0 | 0 | 0 | 0 | 0 | 3.7341E-05 | 0 | 0 | 0 | 3.7341E-05 | 0 | 0 | 0 | 0 | 0 | 2 | 0 | 0 | 0 | 2 |
| UB2D2 | Ubiquitin-cor | 0 | 0.35591768 | 0 | 0 | 0 | 0 | 0 | 0 | 0 | 0.00057151 | 0 | 0.00057151 | 0 | 0 | 0 | 0 | 0 | 0 | 0 | 2 | 0 | 2 |
| UB2D3 | Ubiquitin-cor | 0 | 0.35591768 | 0 | 0 | 0 | 0 | 0 | 0 | 0 | 0.00057151 | 0 | 0.00057151 | 0 | 0 | 0 | 0 | 0 | 0 | 0 | 2 | 0 | 2 |
| UBP24 | Ubiquitin car | 0 | 0.35591768 | 0 | 0 | 0 | 0 | 0 | 0 | 0 | 4.8099E-05 | 0 | 4.8099E-05 | 0 | 0 | 0 | 0 | 0 | 0 | 0 | 3 | 0 | 3 |
| TSN | Translin OS=I | 0 | 0.35591768 | 0 | 0 | 0 | 0 | 0 | 0 | 0 | 0.00036848 | 0 | 0.00036848 | 0 | 0 | 0 | 0 | 0 | 0 | 0 | 2 | 0 | 2 |
| RENT2 | Regulator of | 0 | 0.35591768 | 0 | 0 | 0 | 0 | 0 | 5.2958E-05 | 0 | 0 | 0 | 5.2958E-05 | 0 | 0 | 0 | 0 | 0 | 2 | 0 | 0 | 0 | 2 |
| SIK2 | Serine/threo | 0 | 0.35591768 | 0 | 0 | 0 | 0 | 0 | 7.2746E-05 | 0 | 0 | 0 | 7.2746E-05 | 0 | 0 | 0 | 0 | 0 | 2 | 0 | 0 | 0 | 2 |
| PRDX4 | Peroxiredoxi | 0 | 0.35591768 | 0 | 0 | 0 | 0 | 0 | 0 | 0 | 0.00062002 | 0 | 0.00062002 | 0 | 0 | 0 | 0 | 0 | 0 | 0 | 4 | 0 | 4 |
| SYLM | Probable leu | 0 | 0.35591768 | 0 | 0 | 0 | 0 | 0 | 0 | 7.5073E-05 | 0 | 0 | 7.5073E-05 | 0 | 0 | 0 | 0 | 0 | 0 | 2 | 0 | 0 | 2 |
| NUPL2 | Nucleoporin- | 0 | 0.35591768 | 0 | 0 | 0 | 0 | 0 | 0.00015925 | 0 | 0 | 0 | 0.00015925 | 0 | 0 | 0 | 0 | 0 | 2 | 0 | 0 | 0 | 2 |
| ALBU | Serum album | 0 | 0.35591768 | 0 | 0 | 0 | 0 | 0 | 0 | 0 | 0 | 0.00016707 | 0.00016707 | 0 | 0 | 0 | 0 | 0 | 0 | 0 | 0 | 3 | 3 |
| TMED9 | Transmembra | 0 | 0.35591768 | 0 | 0 | 0 | 0 | 0 | 0.0005733 | 0 | 0 | 0 | 0.0005733 | 0 | 0 | 0 | 0 | 0 | 4 | 0 | 0 | 0 | 4 |
| RS12 | 40S ribosoma | 0 | 0.35591768 | 0 | 0 | 0 | 0 | 0 | 0 | 0.00051357 | 0 | 0 | 0.00051357 | 0 | 0 | 0 | 0 | 0 | 0 | 2 | 0 | 0 | 2 |
| HNRH2 | Heterogeneo | 0 | 0.35591768 | 0 | 0 | 0 | 0 | 0 | 0 | 0 | 0 | 0.00015107 | 0.00015107 | 0 | 0 | 0 | 0 | 0 | 0 | 0 | 0 | 9 | 9 |
| APC7 | Anaphase-pr | 0 | 0.35591768 | 0 | 0 | 0 | 0 | 0 | 0 | 0 | 0 | 0.00011324 | 0.00011324 | 0 | 0 | 0 | 0 | 0 | 0 | 0 | 0 | 2 | 2 |
| RBM8A | RNA-binding | 0 | 0.35591768 | 0 | 0 | 0 | 0 | 0 | 0 | 0.00058441 | 0 | 0 | 0.00058441 | 0 | 0 | 0 | 0 | 0 | 0 | 3 | 0 | 0 | 3 |
| CK5P2 | CDK5 regulat | 0 | 0.35591768 | 0 | 0 | 0 | 0 | 0 | 0 | 0 | 0 | 5.3747E-05 | 5.3747E-05 | 0 | 0 | 0 | 0 | 0 | 0 | 0 | 0 | 3 | 3 |
| OSBP1 | Oxysterol-bir | 0 | 0.35591768 | 0 | 0 | 0 | 0 | 0 | 0 | 8.4004E-05 | 0 | 0 | 8.4004E-05 | 0 | 0 | 0 | 0 | 0 | 0 | 2 | 0 | 0 | 2 |
| PIGT | GPI transami | 0 | 0.35591768 | 0 | 0 | 0 | 0 | 0 | 0 | 0.00011729 | 0 | 0 | 0.00011729 | 0 | 0 | 0 | 0 | 0 | 0 | 2 | 0 | 0 | 2 |
| TNPO2 | Transportin-2 | 0 | 0.35591768 | 0 | 0 | 0 | 0 | 0 | 0 | 0 | 9.3659E-05 | 0 | 9.3659E-05 | 0 | 0 | 0 | 0 | 0 | 0 | 0 | 2 | 0 | 2 |
| KRT83 | Keratin, type | 0 | 0.35591768 | 0 | 0 | 0 | 0 | 0 | 0 | 0 | 0.00076685 | 0 | 0.00076685 | 0 | 0 | 0 | 0 | 0 | 0 | 0 | 10 | 0 | 10 |
| ODPA | Pyruvate deh | 0 | 0.35591768 | 0 | 0 | 0 | 0 | 0 | 0 | 0 | 0.00021542 | 0 | 0.00021542 | 0 | 0 | 0 | 0 | 0 | 0 | 0 | 2 | 0 | 2 |
| GCP2 | Gamma-tubu | 0 | 0.35591768 | 0 | 0 | 0 | 0 | 0 | 7.4682E-05 | 0 | 0 | 0 | 7.4682E-05 | 0 | 0 | 0 | 0 | 0 | 2 | 0 | 0 | 0 | 2 |
| MYH1 | Myosin-1 OS | 0 | 0.35591768 | 0 | 0 | 0 | 0 | 0 | 0.00017371 | 0 | 0 | 0 | 0.00017371 | 0 | 0 | 0 | 0 | 0 | 10 | 0 | 0 | 0 | 10 |
| NAA40 | N-alpha-acet | 0 | 0.35591768 | 0 | 0 | 0 | 0 | 0 | 0 | 0.00028604 | 0 | 0 | 0.00028604 | 0 | 0 | 0 | 0 | 0 | 0 | 2 | 0 | 0 | 2 |
| LATS1 | Serine/threo | 0 | 0.35591768 | 0 | 0 | 0 | 0 | 0 | 0 | 5.9992E-05 | 0 | 0 | 5.9992E-05 | 0 | 0 | 0 | 0 | 0 | 0 | 2 | 0 | 0 | 2 |
| PAK4 | Serine/threo | 0 | 0.35591768 | 0 | 0 | 0 | 0 | 0 | 0 | 0.00022941 | 0 | 0 | 0.00022941 | 0 | 0 | 0 | 0 | 0 | 0 | 4 | 0 | 0 | 4 |
| ATPK | ATP synthase | 0 | 0.35591768 | 0 | 0 | 0 | 0 | 0 | 0 | 0 | 0.00089375 | 0 | 0.00089375 | 0 | 0 | 0 | 0 | 0 | 0 | 0 | 2 | 0 | 2 |
| AKP13 | A-kinase ancl | 0 | 0.35591768 | 0 | 0 | 0 | 0 | 0 | 0 | 0 | 0 | 2.4113E-05 | 2.4113E-05 | 0 | 0 | 0 | 0 | 0 | 0 | 0 | 0 | 2 | 2 |
| IQGA1 | Ras GTPase-a | 0 | 0.35591768 | 0 | 0 | 0 | 0 | 0 | 4.0654E-05 | 0 | 0 | 0 | 4.0654E-05 | 0 | 0 | 0 | 0 | 0 | 2 | 0 | 0 | 0 | 2 |
| GTPB6 | Putative GTP | 0 | 0.35591768 | 0 | 0 | 0 | 0 | 0 | 0 | 0.00013138 | 0 | 0 | 0.00013138 | 0 | 0 | 0 | 0 | 0 | 0 | 2 | 0 | 0 | 2 |
| ERD21 | ER lumen pro | 0 | 0.35591768 | 0 | 0 | 0 | 0 | 0 | 0 | 0 | 0 | 0.00031995 | 0.00031995 | 0 | 0 | 0 | 0 | 0 | 0 | 0 | 0 | 2 | 2 |
| UH1BL | UHRF1-bindi | 0 | 0.35591768 | 0 | 0 | 0 | 0 | 0 | 0 | 4.6305E-05 | 0 | 0 | 4.6305E-05 | 0 | 0 | 0 | 0 | 0 | 0 | 2 | 0 | 0 | 2 |
| K1H1 | Keratin, type | 0 | 0.35591768 | 0 | 0 | 0 | 0 | 0 | 0 | 0 | 0.00090879 | 0 | 0.00090879 | 0 | 0 | 0 | 0 | 0 | 0 | 0 | 9 | 0 | 9 |
| TF2H2 | General trans | 0 | 0.35591768 | 0 | 0 | 0 | 0 | 0 | 0 | 0 | 0 | 0.00017172 | 0.00017172 | 0 | 0 | 0 | 0 | 0 | 0 | 0 | 0 | 2 | 2 |
| SCAM3 | Secretory car | 0 | 0.35591768 | 0 | 0 | 0 | 0 | 0 | 0 | 0 | 0.00024211 | 0 | 0.00024211 | 0 | 0 | 0 | 0 | 0 | 0 | 0 | 2 | 0 | 2 |
| MRPP3 | Mitochondria | 0 | 0.35591768 | 0 | 0 | 0 | 0 | 0 | 0 | 0 | 0 | 0.00011634 | 0.00011634 | 0 | 0 | 0 | 0 | 0 | 0 | 0 | 0 | 2 | 2 |
| RM37 | 39S ribosoma | 0 | 0.35591768 | 0 | 0 | 0 | 0 | 0 | 0.00015925 | 0 | 0 | 0 | 0.00015925 | 0 | 0 | 0 | 0 | 0 | 2 | 0 | 0 | 0 | 2 |
| PDLI7 | PDZ and LIM | 0 | 0.35591768 | 0 | 0 | 0 | 0 | 0 | 0.0001474 | 0 | 0 | 0 | 0.0001474 | 0 | 0 | 0 | 0 | 0 | 2 | 0 | 0 | 0 | 2 |
| RRAGD | Ras-related G | 0 | 0.35591768 | 0 | 0 | 0 | 0 | 0 | 0 | 0 | 0 | 0.00016957 | 0.00016957 | 0 | 0 | 0 | 0 | 0 | 0 | 0 | 0 | 2 | 2 |
| WDR91 | WD repeat-c | 0 | 0.35591768 | 0 | 0 | 0 | 0 | 0 | 0 | 0 | 0 | 0.0001362 | 0.0001362 | 0 | 0 | 0 | 0 | 0 | 0 | 0 | 0 | 3 | 3 |
| TCPH | T-complex pr | 0 | 0.35591768 | 0 | 0 | 0 | 0 | 0 | 0 | 0 | 0.00015472 | 0 | 0.00015472 | 0 | 0 | 0 | 0 | 0 | 0 | 0 | 2 | 0 | 2 |
| HSPB1 | Heat shock p | 0 | 0.35591768 | 0 | 0 | 0 | 0 | 0 | 0 | 0 | 0 | 0.00049631 | 0.00049631 | 0 | 0 | 0 | 0 | 0 | 0 | 0 | 0 | 3 | 3 |
| HASP | Serine/threo | 0 | 0.35591768 | 0 | 0 | 0 | 0 | 0 | 0 | 0 | 0 | 8.4999E-05 | 8.4999E-05 | 0 | 0 | 0 | 0 | 0 | 0 | 0 | 0 | 2 | 2 |
| USP9X | Probable ubi | 0 | 0.35591768 | 0 | 0 | 0 | 0 | 0 | 0 | 0 | 0 | 3.9589E-05 | 3.9589E-05 | 0 | 0 | 0 | 0 | 0 | 0 | 0 | 0 | 3 | 3 |
| ACON | Aconitate hy | 0 | 0.35591768 | 0 | 0 | 0 | 0 | 0 | 0 | 0 | 0.00053854 | 0 | 0.00053854 | 0 | 0 | 0 | 0 | 0 | 0 | 0 | 10 | 0 | 10 |
| BI2L1 | Brain-specific | 0 | 0.35591768 | 0 | 0 | 0 | 0 | 0 | 0.00013183 | 0 | 0 | 0 | 0.00013183 | 0 | 0 | 0 | 0 | 0 | 2 | 0 | 0 | 0 | 2 |
| ELP5 | Elongator co | 0 | 0.35591768 | 0 | 0 | 0 | 0 | 0 | 0 | 0 | 0.00026586 | 0 | 0.00026586 | 0 | 0 | 0 | 0 | 0 | 0 | 0 | 2 | 0 | 2 |
| CIP2A | Protein CIP2A | 0 | 0.35591768 | 0 | 0 | 0 | 0 | 0 | 0 | 0 | 0 | 7.4949E-05 | 7.4949E-05 | 0 | 0 | 0 | 0 | 0 | 0 | 0 | 0 | 2 | 2 |
| PRKRA | Interferon-in | 0 | 0.35591768 | 0 | 0 | 0 | 0 | 0 | 0 | 0 | 0.00040262 | 0 | 0.00040262 | 0 | 0 | 0 | 0 | 0 | 0 | 0 | 3 | 0 | 3 |
| VAPA | Vesicle-assoc | 0 | 0.35591768 | 0 | 0 | 0 | 0 | 0 | 0 | 0 | 0 | 0.00027241 | 0.00027241 | 0 | 0 | 0 | 0 | 0 | 0 | 0 | 0 | 2 | 2 |
| VP26A | Vacuolar pro | 0 | 0.35591768 | 0 | 0 | 0 | 0 | 0 | 0 | 0 | 0 | 0.00020743 | 0.00020743 | 0 | 0 | 0 | 0 | 0 | 0 | 0 | 0 | 2 | 2 |
| UBP53 | Inactive ubiq | 0 | 0.35591768 | 0 | 0 | 0 | 0 | 0 | 0 | 6.3179E-05 | 0 | 0 | 6.3179E-05 | 0 | 0 | 0 | 0 | 0 | 0 | 2 | 0 | 0 | 2 |
| SI1L2 | Signal-induce | 0 | 0.35591768 | 0 | 0 | 0 | 0 | 0 | 0 | 5.9052E-05 | 0 | 0 | 5.9052E-05 | 0 | 0 | 0 | 0 | 0 | 0 | 3 | 0 | 0 | 3 |
| TTC27 | Tetratricope | 0 | 0.35591768 | 0 | 0 | 0 | 0 | 0 | 0 | 0 | 0 | 0.00012069 | 0.00012069 | 0 | 0 | 0 | 0 | 0 | 0 | 0 | 0 | 3 | 3 |
| VPS16 | Vacuolar pro | 0 | 0.35591768 | 0 | 0 | 0 | 0 | 0 | 0 | 0.0000808 | 0 | 0 | 0.0000808 | 0 | 0 | 0 | 0 | 0 | 0 | 2 | 0 | 0 | 2 |
| SELB | Selenocystei | 0 | 0.35591768 | 0 | 0 | 0 | 0 | 0 | 0 | 0 | 0 | 0.00017071 | 0.00017071 | 0 | 0 | 0 | 0 | 0 | 0 | 0 | 0 | 3 | 3 |
| WNK3 | Serine/threo | 0 | 0.35591768 | 0 | 0 | 0 | 0 | 0 | 3.7424E-05 | 0 | 0 | 0 | 3.7424E-05 | 0 | 0 | 0 | 0 | 0 | 2 | 0 | 0 | 0 | 2 |
| IKKA | Inhibitor of n | 0 | 0.35591768 | 0 | 0 | 0 | 0 | 0 | 0.00009042 | 0 | 0 | 0 | 0.00009042 | 0 | 0 | 0 | 0 | 0 | 2 | 0 | 0 | 0 | 2 |
| FA83D | Protein FAM8 | 0 | 0.35591768 | 0 | 0 | 0 | 0 | 0 | 0 | 0.00011588 | 0 | 0 | 0.00011588 | 0 | 0 | 0 | 0 | 0 | 0 | 2 | 0 | 0 | 2 |
| ZN622 | Zinc finger pr | 0 | 0.35591768 | 0 | 0 | 0 | 0 | 0 | 0 | 0 | 0 | 0.0001422 | 0.0001422 | 0 | 0 | 0 | 0 | 0 | 0 | 0 | 0 | 2 | 2 |
| AIMP1 | Aminoacyl tR | 0 | 0.35591768 | 0 | 0 | 0 | 0 | 0 | 0 | 0.00021728 | 0 | 0 | 0.00021728 | 0 | 0 | 0 | 0 | 0 | 0 | 2 | 0 | 0 | 2 |
| WDR62 | WD repeat-c | 0 | 0.35591768 | 0 | 0 | 0 | 0 | 0 | 4.4376E-05 | 0 | 0 | 0 | 4.4376E-05 | 0 | 0 | 0 | 0 | 0 | 2 | 0 | 0 | 0 | 2 |
| TNPO1 | Transportin-1 | 0 | 0.35591768 | 0 | 0 | 0 | 0 | 0 | 0 | 0 | 0 | 7.5533E-05 | 7.5533E-05 | 0 | 0 | 0 | 0 | 0 | 0 | 0 | 0 | 2 | 2 |
| TGO1 | Transport an | 0 | 0.35591768 | 0 | 0 | 0 | 0 | 0 | 0 | 0 | 0 | 3.5568E-05 | 3.5568E-05 | 0 | 0 | 0 | 0 | 0 | 0 | 0 | 0 | 2 | 2 |
| ERD22 | ER lumen pro | 0 | 0.35591768 | 0 | 0 | 0 | 0 | 0 | 0.00031775 | 0 | 0 | 0 | 0.00031775 | 0 | 0 | 0 | 0 | 0 | 3 | 0 | 0 | 0 | 3 |
| AINX | Alpha-intern | 0 | 0.35591768 | 0 | 0 | 0 | 0 | 0 | 0 | 0 | 0 | 0.00013593 | 0.00013593 | 0 | 0 | 0 | 0 | 0 | 0 | 0 | 0 | 3 | 3 |
| SPS2L | SPATS2-like p | 0 | 0.35591768 | 0 | 0 | 0 | 0 | 0 | 0 | 0.00012149 | 0 | 0 | 0.00012149 | 0 | 0 | 0 | 0 | 0 | 0 | 2 | 0 | 0 | 2 |
| CDC45 | Cell division c | 0 | 0.35591768 | 0 | 0 | 0 | 0 | 0 | 0 | 0 | 0 | 0.00017976 | 0.00017976 | 0 | 0 | 0 | 0 | 0 | 0 | 0 | 0 | 3 | 3 |
| COT1 | COUP transcr | 0 | 0.35591768 | 0 | 0 | 0 | 0 | 0 | 0 | 0 | 0 | 0.00016035 | 0.00016035 | 0 | 0 | 0 | 0 | 0 | 0 | 0 | 0 | 2 | 2 |
| HIP1R | Huntingtin-in | 0 | 0.35591768 | 0 | 0 | 0 | 0 | 0 | 0 | 0 | 0 | 0.00006351 | 0.00006351 | 0 | 0 | 0 | 0 | 0 | 0 | 0 | 0 | 2 | 2 |
| RSSA | 40S ribosoma | 0 | 0.35591768 | 0 | 0 | 0 | 0 | 0 | 0 | 0 | 0 | 0.00022993 | 0.00022993 | 0 | 0 | 0 | 0 | 0 | 0 | 0 | 0 | 2 | 2 |
| GEMI2 | Gem-associat | 0 | 0.35591768 | 0 | 0 | 0 | 0 | 0 | 0.00024058 | 0 | 0 | 0 | 0.00024058 | 0 | 0 | 0 | 0 | 0 | 2 | 0 | 0 | 0 | 2 |
| NEMO | NF-kappa-B e | 0 | 0.35591768 | 0 | 0 | 0 | 0 | 0 | 0 | 0 | 0.00020051 | 0 | 0.00020051 | 0 | 0 | 0 | 0 | 0 | 0 | 0 | 2 | 0 | 2 |
| DRS7B | Dehydrogena | 0 | 0.35591768 | 0 | 0 | 0 | 0 | 0 | 0 | 0 | 0 | 0.0002087 | 0.0002087 | 0 | 0 | 0 | 0 | 0 | 0 | 0 | 0 | 2 | 2 |
| CLAP1 | CLIP-associat | 0 | 0.35591768 | 0 | 0 | 0 | 0 | 0 | 0 | 0 | 0 | 6.6153E-05 | 6.6153E-05 | 0 | 0 | 0 | 0 | 0 | 0 | 0 | 0 | 6 | 6 |
| CSPP1 | Centrosome | 0 | 0.35591768 | 0 | 0 | 0 | 0 | 0 | 0 | 5.3974E-05 | 0 | 0 | 5.3974E-05 | 0 | 0 | 0 | 0 | 0 | 0 | 2 | 0 | 0 | 2 |
| TPPP3 | Tubulin polyr | 0 | 0.35591768 | 0 | 0 | 0 | 0 | 0 | 0.00057412 | 0 | 0 | 0 | 0.00057412 | 0 | 0 | 0 | 0 | 0 | 3 | 0 | 0 | 0 | 3 |
| PI51A | Phosphatidyl | 0 | 0.35591768 | 0 | 0 | 0 | 0 | 0 | 0.00011986 | 0 | 0 | 0 | 0.00011986 | 0 | 0 | 0 | 0 | 0 | 2 | 0 | 0 | 0 | 2 |
| TMED2 | Transmembra | 0 | 0.35591768 | 0 | 0 | 0 | 0 | 0 | 0 | 0 | 0 | 0.00033746 | 0.00033746 | 0 | 0 | 0 | 0 | 0 | 0 | 0 | 0 | 2 | 2 |
| KRT82 | Keratin, type | 0 | 0.35591768 | 0 | 0 | 0 | 0 | 0 | 0 | 0 | 0.00016377 | 0 | 0.00016377 | 0 | 0 | 0 | 0 | 0 | 0 | 0 | 4 | 0 | 4 |
| S10A7 | Protein S100 | 0 | 0.35591768 | 0 | 0 | 0 | 0 | 0 | 0 | 0 | 0 | 0.0013431 | 0.0013431 | 0 | 0 | 0 | 0 | 0 | 0 | 0 | 0 | 4 | 4 |
| LEG7 | Galectin-7 OS | 0 | 0.35591768 | 0 | 0 | 0 | 0 | 0 | 0 | 0 | 0 | 0.00049874 | 0.00049874 | 0 | 0 | 0 | 0 | 0 | 0 | 0 | 0 | 2 | 2 |
| KRT85 | Keratin, type | 0 | 0.35591768 | 0 | 0 | 0 | 0 | 0 | 0 | 0 | 0.00033141 | 0 | 0.00033141 | 0 | 0 | 0 | 0 | 0 | 0 | 0 | 10 | 0 | 10 |
| PLPL6 | Neuropathy t | 0 | 0.35591768 | 0 | 0 | 0 | 0 | 0 | 0 | 0 | 6.1502E-05 | 0 | 6.1502E-05 | 0 | 0 | 0 | 0 | 0 | 0 | 0 | 2 | 0 | 2 |
| MCCA | Methylcrotor | 5.16955061 | 3.7988E-05 | 0.0073888 | 0.0062481 | 0.0087587 | 0.0064365 | 0.00720803 | 0.034936 | 0.033428 | 0.045367 | 0.035318 | 0.03726225 | 203 | 170 | 240 | 175 | 197 | 752 | 715 | 783 | 755 | 751.25 |
| ACACA | Acetyl-CoA ca | 4.58014067 | 0.00011537 | 0.0038807 | 0.0038391 | 0.0049737 | 0.0038418 | 0.00413383 | 0.016999 | 0.016977 | 0.023832 | 0.017926 | 0.0189335 | 345 | 338 | 441 | 338 | 365.5 | 1184 | 1175 | 1331 | 1240 | 1232.5 |
| PCCA | Propionyl-Co | 7.108263 | 0.00045685 | 0.0028273 | 0.0023059 | 0.0034891 | 0.0025274 | 0.00278743 | 0.016424 | 0.017274 | 0.027062 | 0.018495 | 0.01981375 | 78 | 63 | 96 | 69 | 76.5 | 355 | 371 | 469 | 397 | 398 |
| KIF1C | Kinesin-like p | 660.708876 | 2.0065E-07 | 4.7848E-05 | 0 | 4.7976E-05 | 7.2527E-05 | 5.6117E-05 | 0.037163 | 0.037061 | 0.03359 | 0.040494 | 0.037077 | 2 | 0 | 2 | 3 | 2.33333333 | 1217 | 1206 | 882 | 1317 | 1155.5 |
| PCCB | Propionyl-Co | 8.06013126 | 0.00054371 | 0.0022521 | 0.001928 | 0.0029453 | 0.0020778 | 0.0023008 | 0.015122 | 0.016476 | 0.025718 | 0.016863 | 0.01854475 | 46 | 39 | 60 | 42 | 46.75 | 242 | 262 | 330 | 268 | 275.5 |
| 1433E | 14-3-3 protei | 5.5358756 | 2.0258E-05 | 0 | 0.00020899 | 0.00031128 | 0 | 0.00026014 | 0.0013208 | 0.0013292 | 0.0016473 | 0.001463 | 0.00144008 | 0 | 2 | 3 | 0 | 2.5 | 10 | 10 | 10 | 11 | 10.25 |
| GCN1 | eIF-2-alpha k | 4.35762498 | 4.5625E-05 | 0.00012843 | 5.9857E-05 | 0.00013868 | 7.9867E-05 | 0.00010171 | 0.00042874 | 0.0003934 | 0.00051898 | 0.00043171 | 0.00044321 | 13 | 6 | 14 | 8 | 10.25 | 34 | 31 | 33 | 34 | 33 |
| RENT1 | Regulator of | 5.01365728 | 2.2522E-05 | 0.00030385 | 0.00018881 | 0.0003984 | 0.00018895 | 0.00027 | 0.0011337 | 0.001381 | 0.0014883 | 0.0014118 | 0.0013537 | 13 | 8 | 17 | 8 | 11.5 | 38 | 46 | 40 | 47 | 42.75 |
| HOOK1 | Protein Hook | 16.3908997 | 0.00097017 | 0.00018124 | 0.00025622 | 0.00014538 | 0.00021977 | 0.00020065 | 0.0025446 | 0.0027005 | 0.0047892 | 0.0031212 | 0.00328888 | 5 | 7 | 4 | 6 | 5.5 | 55 | 58 | 83 | 67 | 65.75 |
| STAU2 | Double-stran | 5.0531629 | 3.5522E-05 | 0.00018518 | 9.3496E-05 | 0.00027851 | 0.00018713 | 0.00018608 | 0.0010045 | 0.00095146 | 0.0010317 | 0.00077349 | 0.00094029 | 4 | 2 | 6 | 4 | 4 | 17 | 16 | 14 | 13 | 15 |
| DDX3X | ATP-depende | 4.58011277 | 7.9851E-06 | 0.0033882 | 0.0036226 | 0.0029176 | 0.0026988 | 0.0031568 | 0.014704 | 0.015053 | 0.012247 | 0.01583 | 0.0144585 | 85 | 90 | 73 | 67 | 78.75 | 289 | 294 | 193 | 309 | 271.25 |
| IPO9 | Importin-9 O | 4.59486769 | 0.00121309 | 0.00020279 | 0.00012799 | 0.00020333 | 0.00015369 | 0.00017195 | 0.00087359 | 0.00058609 | 0.0010491 | 0.00065157 | 0.00079009 | 8 | 5 | 8 | 6 | 6.75 | 27 | 18 | 26 | 20 | 22.75 |
| PYR1 | CAD protein | 4.4918278 | 3.6889E-05 | 0.00004744 | 9.5808E-05 | 0.00010702 | 4.7938E-05 | 7.4552E-05 | 0.00036331 | 0.00036562 | 0.00032095 | 0.00028961 | 0.00033487 | 4 | 8 | 9 | 4 | 6.25 | 24 | 24 | 17 | 19 | 21 |
| YTHD2 | YTH domain- | 9.3255222 | 9.0727E-05 | 9.1152E-05 | 0 | 0 | 0 | 9.1152E-05 | 0.00075624 | 0.00099521 | 0.00065295 | 0.00099576 | 0.00085004 | 5 | 0 | 0 | 0 | 5 | 17 | 20 | 11 | 20 | 17 |
| IF4A3 | Eukaryotic in | 5.99246946 | 0.00048695 | 0.00012841 | 0.00032417 | 0 | 0.00025952 | 0.00023737 | 0.001721 | 0.00098965 | 0.0013287 | 0.0016503 | 0.00142241 | 3 | 6 | 0 | 4 | 4.33333333 | 25 | 17 | 15 | 22 | 19.75 |
| TOP3B | DNA topoiso | 17.1200552 | 0.00025161 | 6.1226E-05 | 0 | 0 | 0 | 6.1226E-05 | 0.0013285 | 0.001219 | 0.0007797 | 0.00086557 | 0.00104819 | 2 | 0 | 0 | 0 | 2 | 34 | 31 | 16 | 22 | 25.75 |
| ANR52 | Serine/threo | 4.50346869 | 0.000172 | 9.8098E-05 | 7.4293E-05 | 0.00014754 | 0.00014869 | 0.00011716 | 0.00062605 | 0.00040952 | 0.00050751 | 0.00056734 | 0.00052761 | 4 | 3 | 6 | 6 | 4.75 | 21 | 13 | 14 | 18 | 16.5 |
| F16A1 | Protein FAM1 | 7.18161815 | 0.00242942 | 0 | 5.1243E-05 | 0.00010176 | 0.00005128 | 6.8094E-05 | 0.00029148 | 0.00055406 | 0.00068664 | 0.00042393 | 0.00048903 | 0 | 2 | 4 | 2 | 2.66666667 | 9 | 17 | 17 | 13 | 14 |
| F120A | Constitutive | 5.50081052 | 0.00020758 | 0.00018883 | 0 | 0.00018933 | 0.00014311 | 0.00017376 | 0.0010544 | 0.0010611 | 0.00067631 | 0.0010314 | 0.0009558 | 8 | 0 | 8 | 6 | 7.33333333 | 35 | 35 | 18 | 34 | 30.5 |
| PYGL | Glycogen pho | 4.13520698 | 0.00824334 | 0.00006231 | 0.00012584 | 0 | 6.2965E-05 | 8.3705E-05 | 0.00023859 | 0.00032015 | 0.00054553 | 0.00028028 | 0.00034614 | 2 | 4 | 0 | 2 | 2.66666667 | 6 | 8 | 11 | 7 | 8 |
| YTHD1 | YTH domain- | 5.88031007 | 0.00112581 | 0.00014162 | 0.000143 | 0.00023666 | 0.00019081 | 0.00017802 | 0.0010243 | 0.0013946 | 0.00067631 | 0.0010921 | 0.00104683 | 3 | 3 | 5 | 4 | 3.75 | 17 | 23 | 9 | 18 | 16.75 |
| REN3B | Regulator of | 4.64011059 | 0.00570593 | 0.00010927 | 0 | 0 | 0.00016562 | 0.00013745 | 0.00048814 | 0.00084213 | 0.00086969 | 0.00035108 | 0.00063776 | 2 | 0 | 0 | 3 | 2.5 | 7 | 12 | 10 | 5 | 8.5 |
| PP6R2 | Serine/threo | 4.29241883 | 0.00124526 | 0.00010927 | 0.00024826 | 0.00010956 | 0.00019323 | 0.00016508 | 0.00059274 | 0.00087722 | 0.00052182 | 0.00084259 | 0.00070859 | 4 | 9 | 4 | 7 | 6 | 17 | 25 | 12 | 24 | 19.5 |
| K1C14 | Keratin, type | 8.96109681 | 0.04022865 | 0.00022363 | 0.00016936 | 0.00011211 | 0 | 0.00016837 | 0.0012131 | 0.0005745 | 0.0030259 | 0.0012215 | 0.00150875 | 9 | 11 | 7 | 0 | 9 | 26 | 19 | 46 | 29 | 30 |
| ATX2 | Ataxin-2 OS= | 5.65797338 | 0.00348517 | 6.0293E-05 | 8.1177E-05 | 8.0605E-05 | 4.0618E-05 | 6.5673E-05 | 0.00043609 | 0.00051631 | 0.00022395 | 0.00030996 | 0.00037158 | 3 | 5 | 4 | 2 | 3.5 | 17 | 20 | 7 | 13 | 14.25 |
| NFX1 | Transcription | 4.36809691 | 0.0055334 | 0 | 9.5166E-05 | 0 | 7.1426E-05 | 8.3296E-05 | 0.00054131 | 0.00036317 | 0.00018753 | 0.00036337 | 0.00036385 | 0 | 4 | 0 | 3 | 3.5 | 18 | 12 | 5 | 12 | 11.75 |
| PRC2A | Protein PRRC | 4.63473694 | 0.00511233 | 0.00014681 | 0.0001606 | 0.000184 | 0.00017307 | 0.00016612 | 0.00089006 | 0.00089571 | 0.00035054 | 0.00094338 | 0.00076992 | 13 | 14 | 15 | 15 | 14.25 | 63 | 60 | 19 | 63 | 51.25 |
| FUBP3 | Far upstream | 4.44759456 | 0.00457618 | 0.0004152 | 0.00032609 | 0.00023128 | 0.00018647 | 0.00028976 | 0.0016487 | 0.0013037 | 0.00066094 | 0.0015416 | 0.00128874 | 9 | 7 | 5 | 4 | 6.25 | 28 | 22 | 9 | 26 | 21.25 |
| SMG1 | Serine/threo | 5.52874076 | 0.00982745 | 0.00003604 | 5.8228E-05 | 6.5045E-05 | 0.00005827 | 5.4396E-05 | 0.0003588 | 0.00037034 | 0.00010327 | 0.00037055 | 0.00030074 | 5 | 8 | 9 | 8 | 7.5 | 39 | 40 | 9 | 40 | 32 |
| CE350 | Centrosome- | 6.74623181 | 0.01136747 | 0.00004233 | 2.5646E-05 | 0.00005942 | 0.00003422 | 4.0404E-05 | 0.00032417 | 0.00034798 | 8.0859E-05 | 0.00033729 | 0.00027257 | 5 | 3 | 7 | 4 | 4.75 | 30 | 32 | 6 | 31 | 24.75 |
| ECH1 | Delta(3,5)-De | 3.97403507 | 0.03082924 | 0 | 0 | 0.00032267 | 0.0001626 | 0.00024264 | 0.00051344 | 0.00062004 | 0.0017929 | 0.00093058 | 0.00096424 | 0 | 0 | 4 | 2 | 3 | 5 | 6 | 14 | 9 | 8.5 |
| SMG9 | Protein SMG9 | 5.54083565 | 0.01519032 | 0.00015224 | 0.00010249 | 0.00010176 | 0 | 0.00011883 | 0.00084204 | 0.00078221 | 0.00016156 | 0.00084786 | 0.00065842 | 3 | 2 | 2 | 0 | 2.33333333 | 13 | 12 | 2 | 13 | 10 |
| PRC2B | Protein PRRC | 6.45339976 | 0.01755679 | 0.00013023 | 9.5636E-05 | 0.00014244 | 8.3741E-05 | 0.00011301 | 0.00083108 | 0.00098843 | 0.00016961 | 0.00092812 | 0.00072931 | 12 | 9 | 12 | 8 | 10.25 | 61 | 68 | 10 | 64 | 50.75 |
| LSM12 | Protein LSM1 | 11.0024393 | 0.02801898 | 0 | 0.0002733 | 0 | 0 | 0.0002733 | 0.0031091 | 0.0027812 | 0 | 0.0031306 | 0.00300697 | 0 | 2 | 0 | 0 | 2 | 18 | 16 | 0 | 18 | 17.3333333 |
| OTUD4 | OTU domain- | 8.52391109 | 0.0249433 | 4.7376E-05 | 4.7839E-05 | 0 | 0.00007181 | 5.5675E-05 | 0.00048376 | 0.00076067 | 7.5415E-05 | 0.00057843 | 0.00047457 | 2 | 2 | 0 | 3 | 2.33333333 | 16 | 25 | 2 | 19 | 15.5 |
| AGO2 | Protein argor | 4.472447 | 0.01714128 | 0.00018432 | 0.00021714 | 0.00018481 | 0.00015521 | 0.00018537 | 0.0010587 | 0.00094702 | 0.00024451 | 0.001066 | 0.00082906 | 6 | 7 | 6 | 5 | 6 | 27 | 24 | 5 | 27 | 20.75 |
| AKAP9 | A-kinase ancl | 5.18765915 | 0.02366926 | 2.0242E-05 | 0.00002044 | 2.0296E-05 | 1.3636E-05 | 1.8654E-05 | 0.00008612 | 0.000156 | 3.2222E-05 | 0.00011273 | 9.6768E-05 | 3 | 3 | 3 | 2 | 2.75 | 10 | 18 | 3 | 13 | 11 |
| TNR6A | Trinucleotide | 4.95903104 | 0.05139382 | 6.7249E-05 | 9.5069E-05 | 0 | 0 | 8.1159E-05 | 0.00034334 | 0.00038007 | 0 | 0.000484 | 0.00040247 | 5 | 7 | 0 | 0 | 6 | 20 | 22 | 0 | 28 | 23.3333333 |
| PCM1 | Pericentriola | 6.20034052 | 0.05475431 | 2.6076E-05 | 2.6331E-05 | 3.9217E-05 | 0 | 3.0541E-05 | 0.00021633 | 0.00020096 | 0 | 0.00015081 | 0.00018937 | 2 | 2 | 3 | 0 | 2.33333333 | 13 | 12 | 0 | 9 | 11.3333333 |
| CAPR1 | Caprin-1 OS= | 4.1243037 | 0.06215537 | 0.00011166 | 0 | 0 | 0.00015044 | 0.00013105 | 0.00066508 | 0.00052588 | 0 | 0.00043051 | 0.00054049 | 3 | 0 | 0 | 4 | 3.5 | 14 | 11 | 0 | 9 | 11.3333333 |
| ZN598 | E3 ubiquitin- | 5.12535851 | 0.07186697 | 5.8381E-05 | 0 | 5.8537E-05 | 0 | 5.8459E-05 | 0.00018629 | 0.00041245 | 0 | 0.00030013 | 0.00029962 | 2 | 0 | 2 | 0 | 2 | 5 | 11 | 0 | 8 | 8 |
| ARFG2 | ADP-ribosyla | 4.25633217 | 0.07622072 | 0.0001013 | 0.00010229 | 0 | 0 | 0.0001018 | 0.00025859 | 0.00052047 | 0 | 0.00052076 | 0.00043327 | 2 | 2 | 0 | 0 | 2 | 4 | 8 | 0 | 8 | 6.66666667 |
| CE192 | Centrosomal | 4.73374099 | 0.08977164 | 5.4381E-05 | 6.8641E-05 | 2.7263E-05 | 4.1214E-05 | 4.7875E-05 | 0.00020823 | 0.00019209 | 0 | 0.00027956 | 0.00022663 | 4 | 5 | 2 | 3 | 3.5 | 12 | 11 | 0 | 16 | 13 |
| APC | Adenomatou | 4.41298036 | 0.09177458 | 5.5691E-05 | 6.5609E-05 | 0.00002792 | 8.4415E-05 | 5.8409E-05 | 0.00028433 | 0.00027422 | 0 | 0.00021472 | 0.00025776 | 6 | 7 | 3 | 9 | 6.25 | 24 | 23 | 0 | 18 | 21.6666667 |
| PUM1 | Pumilio hom | 4.25571612 | 0.09451252 | 0.000178 | 0.00008987 | 0.00013386 | 8.9935E-05 | 0.00012292 | 0.00045439 | 0.00060018 | 0 | 0.00051472 | 0.0005231 | 8 | 4 | 6 | 4 | 5.5 | 16 | 21 | 0 | 18 | 18.3333333 |
| SC16A | Protein trans | 3.86488247 | 0.09894663 | 0.00010899 | 0.00009783 | 0.00012143 | 7.3425E-05 | 0.00010042 | 0.00040189 | 0.00038889 | 0 | 0.00037354 | 0.00038811 | 9 | 8 | 10 | 6 | 8.25 | 26 | 25 | 0 | 24 | 25 |

| | | | | | | | | | | | | | | | | | | | | | |
|---|---|---|---|---|---|---|---|---|---|---|---|---|---|---|---|---|---|---|---|---|---|
| IBTK | Inhibitor of B | 3.84293617 | 0.00896895 | 5.8511E-05 | 7.8777E-05 | 5.8667E-05 | 5.9126E-05 | 6.377E-05 | 0.00029873 | 0.00017537 | 0.00015523 | 0.00035093 | 0.00024507 | 3 | 4 | 3 | 3 | 3.25 | 12 | 7 | 5 | 14 | 9.5 |
| K1C16 | Keratin, type | 3.81071999 | 0.01630542 | 0.00033474 | 0.00090136 | 0.00033563 | 0.00016913 | 0.00043522 | 0.0015666 | 0.00085993 | 0.0024866 | 0.0017208 | 0.00165848 | 9 | 19 | 7 | 6 | 10.25 | 27 | 16 | 31 | 28 | 25.5 |
| R3HD1 | R3H domain- | 3.81025049 | 0.04304071 | 0 | 4.8492E-05 | 0 | 0 | 4.8492E-05 | 0.00015324 | 0.0002159 | 0 | 0.00018516 | 0.00018477 | 0 | 2 | 0 | 0 | 2 | 5 | 7 | 0 | 6 | 6 |
| MOV10 | Putative helic | 3.79326766 | 0.00028575 | 0.00018417 | 0.0000797 | 0.00015828 | 0.00015952 | 0.00014542 | 0.00060446 | 0.00054071 | 0.00041881 | 0.00064245 | 0.00055161 | 7 | 3 | 6 | 6 | 5.5 | 18 | 16 | 10 | 19 | 15.75 |
| HNRPQ | Heterogeneo | 3.78051037 | 3.3392E-06 | 0.00038121 | 0.0002994 | 0.00038223 | 0.00034242 | 0.00035132 | 0.0012975 | 0.001469 | 0.0013485 | 0.0011976 | 0.00132815 | 9 | 7 | 9 | 8 | 8.25 | 24 | 27 | 20 | 22 | 23.25 |
| IPO8 | Importin-8 O | 3.7764593 | 0.00020472 | 0.00040715 | 0.00035974 | 0.00030618 | 0.00033428 | 0.00035184 | 0.0012018 | 0.0011113 | 0.0016608 | 0.0013409 | 0.0013287 | 16 | 14 | 12 | 13 | 13.75 | 37 | 34 | 41 | 41 | 38.25 |
| PRC2C | Protein PRRC | 3.73321493 | 0.01321235 | 0.00034626 | 0.00034964 | 0.00025582 | 0.00030386 | 0.0003139 | 0.0013956 | 0.0014513 | 0.00043515 | 0.0014053 | 0.00117184 | 38 | 38 | 28 | 33 | 34.25 | 120 | 124 | 30 | 120 | 98.5 |
| 4ET | Eukaryotic tr | 3.71742988 | 0.02392787 | 0.00010716 | 0.00013526 | 0.00010745 | 5.4143E-05 | 0.000101 | 0.00044453 | 0.00037853 | 0.00012794 | 0.00055089 | 0.00037547 | 4 | 5 | 4 | 2 | 3.75 | 13 | 11 | 3 | 16 | 10.75 |
| ACACB | Acetyl-CoA ca | 3.63373723 | 0.00069444 | 0.00053679 | 0.00047699 | 0.00048439 | 0.00049903 | 0.0004993 | 0.001658 | 0.0015445 | 0.0024267 | 0.0016281 | 0.00181433 | 79 | 84 | 84 | 82 | 82.25 | 241 | 212 | 267 | 236 | 239 |
| CAMP3 | Calmodulin-r | 3.61391254 | 0.10950553 | 0.00012677 | 0.00010667 | 0.00016947 | 0.00010675 | 0.00012742 | 0.00045844 | 0.00046135 | 0 | 0.00046161 | 0.00046047 | 6 | 5 | 8 | 5 | 6 | 17 | 17 | 0 | 17 | 17 |
| PAN3 | PAN2-PAN3 d | 3.61129451 | 0.01366102 | 0.0000595 | 6.0082E-05 | 0 | 9.0188E-05 | 6.9923E-05 | 0.00030378 | 0.00034392 | 9.4715E-05 | 0.00026764 | 0.00025251 | 2 | 2 | 0 | 3 | 2.33333333 | 8 | 9 | 2 | 7 | 6.5 |
| ECHB | Trifunctional | 3.6071662 | 0.10319722 | 0 | 0.00011243 | 0.00011164 | 0.00011251 | 0.00011219 | 0.00028423 | 0.00050057 | 0 | 0.0004293 | 0.0004047 | 0 | 2 | 2 | 2 | 2 | 4 | 7 | 0 | 6 | 5.66666667 |
| YTDC2 | Probable ATP | 3.60469541 | 9.5029E-06 | 0.00012917 | 5.5902E-05 | 9.2513E-05 | 7.4589E-05 | 8.8044E-05 | 0.0003062 | 0.00033185 | 0.00032312 | 0.00030831 | 0.00031737 | 7 | 3 | 5 | 4 | 4.75 | 13 | 14 | 11 | 13 | 12.75 |
| NAMPT | Nicotinamide | 3.58244512 | 0.00331792 | 0.00010749 | 0.00037989 | 0.00010777 | 0.00016293 | 0.00018952 | 0.00061738 | 0.00075937 | 0.00085552 | 0.00048351 | 0.00067895 | 2 | 7 | 2 | 3 | 3.5 | 9 | 11 | 10 | 7 | 9.25 |
| SR1IP | Protein SREK | 3.55580578 | 0.12018843 | 0.00051074 | 0.00034383 | 0.0005121 | 0.00051611 | 0.0004707 | 0.0015211 | 0.0015308 | 0 | 0.0019692 | 0.0016737 | 3 | 2 | 3 | 3 | 2.75 | 7 | 7 | 0 | 9 | 7.66666667 |
| RL27A | 60S ribosoma | 3.52346394 | 7.4916E-05 | 0.0003566 | 0.00054013 | 0.00053632 | 0 | 0.00047768 | 0.001593 | 0.0016032 | 0.001703 | 0.0018332 | 0.0016831 | 2 | 3 | 3 | 0 | 2.66666667 | 7 | 7 | 6 | 8 | 7 |
| WNK1 | Serine/threo | 3.51747578 | 0.15802555 | 2.2157E-05 | 0.00003356 | 5.5539E-05 | 4.4779E-05 | 3.9009E-05 | 0.00016968 | 0.00015653 | 0 | 8.5427E-05 | 0.00013721 | 2 | 3 | 5 | 4 | 3.5 | 12 | 11 | 0 | 6 | 9.66666667 |
| TT30A | Tetratricopep | 3.47949183 | 0.012969 | 0.00019841 | 0.00020035 | 0.00011936 | 0.00016039 | 0.00016963 | 0.00030389 | 0.00061165 | 0.00088434 | 0.00056099 | 0.00059022 | 5 | 5 | 3 | 4 | 4.25 | 6 | 12 | 14 | 11 | 10.75 |
| K22E | Keratin, type | 3.45969509 | 0.01749232 | 0.0011563 | 0.0010008 | 0.0010352 | 0.00075114 | 0.00098586 | 0.0041114 | 0.0018566 | 0.0051275 | 0.0025476 | 0.00341078 | 30 | 29 | 28 | 20 | 26.75 | 86 | 42 | 88 | 54 | 67.5 |
| RBNS5 | Rabenosyn-5 | 3.42967807 | 0.00040282 | 0.00037025 | 0.00023792 | 0.00050622 | 0.00017006 | 0.00032111 | 0.0012888 | 0.0010376 | 0.00091085 | 0.001168 | 0.00110131 | 11 | 7 | 15 | 5 | 9.5 | 30 | 24 | 17 | 27 | 24.5 |
| TTC28 | Tetratricopep | 3.40023359 | 0.09602492 | 0 | 3.2221E-05 | 3.1994E-05 | 0 | 3.2108E-05 | 6.7879E-05 | 0.00015028 | 0 | 0.00010936 | 0.00010917 | 0 | 3 | 3 | 0 | 3 | 5 | 11 | 0 | 8 | 8 |
| NP1L4 | Nucleosome | 3.38450792 | 0.00077001 | 0.00049258 | 0.00071057 | 0.00070556 | 0.00071108 | 0.00065495 | 0.001976 | 0.0018078 | 0.0029124 | 0.0021705 | 0.00221668 | 7 | 10 | 10 | 10 | 9.25 | 22 | 20 | 26 | 24 | 23 |
| GEMI4 | Gem-associat | 3.36218711 | 0.01784693 | 9.9767E-05 | 7.5557E-05 | 0.00010003 | 7.5611E-05 | 8.7741E-05 | 0.00022285 | 0.00019222 | 0.00047644 | 0.0002885 | 0.000295 | 4 | 3 | 4 | 3 | 3.5 | 7 | 6 | 12 | 9 | 8.5 |
| RACK1 | Receptor of a | 3.35424583 | 0.00119737 | 0.0014984 | 0.0015971 | 0.0021701 | 0.0016824 | 0.001737 | 0.0045688 | 0.007271 | 0.0067581 | 0.0047074 | 0.00582633 | 18 | 19 | 26 | 20 | 20.75 | 43 | 68 | 51 | 44 | 51.5 |
| SYQ | Glutamine--t | 3.35381547 | 0.00114544 | 6.8099E-05 | 0 | 0.00006828 | 0 | 6.819E-05 | 0.00030422 | 0.00017495 | 0.00021681 | 0.0002188 | 0.0002287 | 2 | 0 | 2 | 0 | 2 | 7 | 4 | 4 | 5 | 5 |
| PDC6I | Programmed | 3.34709758 | 0.0086631 | 9.1204E-05 | 0.00021489 | 0.00015241 | 6.1442E-05 | 0.00012999 | 0.00042684 | 0.0005467 | 0.00053234 | 0.00023443 | 0.00043508 | 3 | 7 | 5 | 2 | 4.25 | 11 | 14 | 11 | 6 | 10.5 |
| NP1L1 | Nucleosome | 3.34522017 | 0.01024548 | 0.00040494 | 0.0005452 | 0.00033835 | 0.00075018 | 0.00050967 | 0.0011198 | 0.0016471 | 0.0025784 | 0.0014745 | 0.00170495 | 9 | 10 | 8 | 14 | 10.25 | 16 | 23 | 28 | 22 | 22.25 |
| DDX6 | Probable ATP | 3.33981245 | 0.00906982 | 0.0001639 | 0 | 0 | 0.00016562 | 0.00016476 | 0.00083681 | 0.00028071 | 0.00052182 | 0.00056173 | 0.00055027 | 3 | 0 | 0 | 3 | 3 | 12 | 4 | 6 | 8 | 7.5 |
| MCCB | Methylcroton | 3.32875076 | 9.6373E-05 | 0.0037028 | 0.0024611 | 0.0041826 | 0.0024629 | 0.00320235 | 0.0087943 | 0.010476 | 0.011863 | 0.011506 | 0.01065983 | 79 | 52 | 89 | 52 | 68 | 147 | 174 | 159 | 191 | 167.75 |
| LUZP1 | Leucine zippe | 3.27168451 | 0.13246518 | 0.00017167 | 0.00019812 | 9.8359E-05 | 0.00019826 | 0.0001666 | 0.00050084 | 0.00056703 | 0 | 0.00056734 | 0.00054507 | 7 | 8 | 4 | 8 | 6.75 | 16 | 18 | 0 | 18 | 17.3333333 |
| PRPS1 | Ribose-phosp | 3.22029272 | 0.00014638 | 0.00024895 | 0.00025138 | 0.00024961 | 0.00050312 | 0.00031327 | 0.0010592 | 0.00085272 | 0.0010568 | 0.0010665 | 0.00100881 | 3 | 3 | 3 | 6 | 3.75 | 10 | 8 | 8 | 10 | 9 |
| TBB6 | Tubulin beta- | 3.17421125 | 0.07704124 | 0 | 0.00011949 | 0.00011865 | 0 | 0.00011907 | 0.00045312 | 0.000304 | 0.00037674 | 0 | 0.00037795 | 0 | 41 | 35 | 0 | 38 | 41 | 39 | 34 | 0 | 38 |
| CEP55 | Centrosomal | 3.16853126 | 0.00018987 | 0 | 0.00011486 | 0 | 0 | 0.00011486 | 0.00036295 | 0.00043831 | 0.00036212 | 0.00029237 | 0.00036394 | 0 | 2 | 0 | 0 | 2 | 5 | 6 | 4 | 4 | 4.75 |
| IF2B2 | Insulin-like gr | 3.16066001 | 8.8786E-05 | 0.00026432 | 0.00026691 | 0.00044171 | 0.00035614 | 0.00033227 | 0.0011246 | 0.00090539 | 0.00098178 | 0.001189 | 0.00105019 | 7 | 7 | 11 | 9 | 8.5 | 21 | 17 | 15 | 22 | 18.75 |
| ANR28 | Serine/threo | 3.14529317 | 2.9349E-05 | 0.00027566 | 0.00027836 | 0.00032665 | 0.00017726 | 0.00026448 | 0.00073568 | 0.00086912 | 0.00091751 | 0.00080519 | 0.00083188 | 11 | 11 | 13 | 7 | 10.5 | 23 | 27 | 23 | 25 | 24.5 |
| CSDE1 | Cold shock do | 3.07402022 | 0.00052319 | 0.0015873 | 0.0021037 | 0.0011605 | 0.0012698 | 0.00153033 | 0.0040519 | 0.0048422 | 0.005843 | 0.0040799 | 0.00470425 | 48 | 63 | 35 | 38 | 46 | 96 | 114 | 111 | 96 | 104.25 |
| SUCA | Succinate--Co | 3.01906669 | 0.00220061 | 0 | 0 | 0.00015294 | 0.00015414 | 0.00015354 | 0.00048673 | 0.00058778 | 0.00048562 | 0.00029406 | 0.00046355 | 0 | 0 | 2 | 2 | 2 | 5 | 6 | 4 | 3 | 4.5 |
| ECM29 | Proteasome- | 3.00848046 | 0.00503221 | 0 | 7.2213E-05 | 5.7363E-05 | 2.8906E-05 | 5.2827E-05 | 0.00021907 | 0.00011023 | 0.00015937 | 0.00014705 | 0.00015893 | 0 | 5 | 4 | 2 | 3.66666667 | 12 | 6 | 7 | 8 | 8.25 |
| TOM40 | Mitochondria | 2.95693466 | 0.00135431 | 0.00036549 | 0.00029525 | 0.00036646 | 0.0002216 | 0.0003122 | 0.0010263 | 0.00075115 | 0.0011636 | 0.00075157 | 0.00092316 | 5 | 4 | 5 | 3 | 4.25 | 11 | 8 | 10 | 8 | 9.25 |
| SEBP2 | Selenocystein | 2.94584949 | 0.15534918 | 0.0001236 | 0.00012481 | 9.2946E-05 | 0.0001249 | 0.00011656 | 0.00031552 | 0.00035721 | 0 | 0.00035741 | 0.00034338 | 4 | 4 | 3 | 4 | 3.75 | 8 | 9 | 0 | 9 | 8.66666667 |
| RB3GP | Rab3 GTPase | 2.94136768 | 0.01488401 | 0.00032279 | 0.00024446 | 0.00051245 | 0.00032618 | 0.00035147 | 0.00099568 | 0.0008638 | 0.0015843 | 0.00069143 | 0.0010338 | 12 | 9 | 19 | 12 | 13 | 29 | 25 | 37 | 20 | 27.75 |
| IF2B1 | Insulin-like gr | 2.9012298 | 0.00026116 | 0.00096041 | 0.00083126 | 0.00096297 | 0.00073943 | 0.00087352 | 0.0029187 | 0.0019386 | 0.0026936 | 0.0025862 | 0.00253428 | 21 | 18 | 21 | 16 | 19 | 50 | 33 | 37 | 44 | 41 |
| IFT56 | Intraflagellar | 2.89476375 | 2.2748E-05 | 0.00061922 | 0.00057718 | 0.00052535 | 0.00062573 | 0.00058687 | 0.0016415 | 0.0015908 | 0.0019714 | 0.0015917 | 0.00169885 | 13 | 12 | 11 | 13 | 12.25 | 27 | 26 | 26 | 26 | 26.25 |
| TXTP | Tricarboxylat | 2.89192705 | 0.01682476 | 0.00025455 | 0.00025704 | 0.00017015 | 0.00017148 | 0.00021331 | 0.00064981 | 0.00043596 | 0.00094548 | 0.0004362 | 0.00061686 | 3 | 3 | 2 | 2 | 2.5 | 6 | 4 | 7 | 4 | 5.25 |
| RS4X | 40S ribosoma | 2.8734702 | 0.01367555 | 0.00060202 | 0.0006079 | 0.00060362 | 0.00060834 | 0.00060547 | 0.0015368 | 0.0014177 | 0.0027152 | 0.0012895 | 0.0017398 | 6 | 6 | 6 | 6 | 6 | 12 | 11 | 17 | 10 | 12.5 |
| PAIRB | Plasminogen | 2.86936184 | 0.00074488 | 0.00077613 | 0.00097965 | 0.00058365 | 0.00065357 | 0.00074825 | 0.0024766 | 0.0016615 | 0.0019562 | 0.0024937 | 0.002147 | 12 | 15 | 9 | 10 | 11.5 | 30 | 20 | 19 | 30 | 24.75 |
| RBM14 | RNA-binding | 2.85414467 | 0.16582131 | 0.00047333 | 0.00047796 | 0.00055369 | 0.00059788 | 0.00052572 | 0.0013593 | 0.0016213 | 0 | 0.0015208 | 0.00150047 | 12 | 12 | 14 | 15 | 13.25 | 27 | 32 | 0 | 30 | 29.6666667 |
| CEP85 | Centrosomal | 2.8297832 | 0.08969653 | 6.9261E-05 | 0 | 0 | 0.00013998 | 0.00010462 | 0.00026521 | 0.00026689 | 0 | 0.00035606 | 0.00029605 | 2 | 0 | 0 | 4 | 3 | 6 | 6 | 0 | 8 | 6.66666667 |
| K1C10 | Keratin, type | 2.82944157 | 0.06688216 | 0.0024852 | 0.0022357 | 0.0023106 | 0.002146 | 0.00229438 | 0.0055367 | 0.0037146 | 0.012012 | 0.0047039 | 0.0064918 | 55 | 49 | 51 | 47 | 50.5 | 96 | 64 | 167 | 81 | 102 |
| K2C5 | Keratin, type | 2.79133684 | 0.00293978 | 0 | 0.00018065 | 0 | 0 | 0.00018065 | 0.0004567 | 0.00028725 | 0.00064077 | 0.0006323 | 0.00050426 | 0 | 16 | 0 | 0 | 16 | 41 | 12 | 46 | 37 | 34 |
| SYIC | Isoleucine--tR | 2.76913555 | 0.00231623 | 0.00016728 | 0.0001478 | 0.00023062 | 6.3389E-05 | 0.00015227 | 0.00034696 | 0.00053717 | 0.00039942 | 0.0004031 | 0.00042166 | 8 | 7 | 11 | 3 | 7.25 | 13 | 20 | 12 | 15 | 15 |
| HAUS7 | HAUS augmin | 2.76876898 | 0.01620804 | 0 | 0 | 0 | 0.00014492 | 0.00014492 | 0.00018305 | 0.00036843 | 0.00068488 | 0.00036864 | 0.00040125 | 0 | 0 | 0 | 2 | 2 | 2 | 4 | 6 | 4 | 4 |
| NKAP | NF-kappa-B-a | 2.7687145 | 0.01620893 | 0 | 0 | 0 | 0.00012851 | 0.00012851 | 0.00016232 | 0.0003267 | 0.00060732 | 0.00032689 | 0.00035581 | 0 | 0 | 0 | 2 | 2 | 2 | 4 | 6 | 4 | 4 |
| IF4E | Eukaryotic tr | 2.75277201 | 0.00054368 | 0.00060803 | 0.00036838 | 0.00036579 | 0.00036865 | 0.00042771 | 0.0012417 | 0.0010934 | 0.00096788 | 0.0014066 | 0.0011774 | 5 | 3 | 3 | 3 | 3.5 | 8 | 7 | 5 | 9 | 7.25 |
| FXR2 | Fragile X mer | 2.74249192 | 0.00043888 | 0.00011763 | 0.00023756 | 0.00015726 | 0.00011887 | 0.00015783 | 0.00050047 | 0.00045328 | 0.0003745 | 0.00040314 | 0.00043285 | 3 | 6 | 4 | 3 | 4 | 10 | 9 | 6 | 8 | 8.25 |
| DHX36 | ATP-depende | 2.74122537 | 0.01771291 | 5.2358E-05 | 0.00010574 | 0 | 5.2908E-05 | 7.0335E-05 | 0.00013366 | 0.00030264 | 0.00016669 | 0.00016823 | 0.00019281 | 2 | 4 | 0 | 2 | 2.66666667 | 4 | 9 | 4 | 5 | 5.5 |
| CMC2 | Calcium-bind | 2.73221003 | 5.033E-05 | 0.00046913 | 0.00027633 | 0.00035278 | 0.00031604 | 0.00035357 | 0.00099797 | 0.0009541 | 0.0010579 | 0.00085414 | 0.00096603 | 12 | 7 | 9 | 8 | 9 | 20 | 19 | 17 | 17 | 18.25 |
| ROA0 | Heterogeneo | 2.72152386 | 0.00050183 | 0.00060564 | 0.00052419 | 0.00026025 | 0.00034971 | 0.00043495 | 0.0013252 | 0.0012225 | 0.00096408 | 0.0012231 | 0.00118372 | 7 | 6 | 3 | 4 | 5 | 12 | 12 | 7 | 12 | 10.75 |
| DD19B | ATP-depende | 2.71958414 | 0.00454166 | 0.00027545 | 0.00038941 | 0.00016571 | 0 | 0.00027686 | 0.0008438 | 0.00063687 | 0.00096465 | 0.00056642 | 0.00075294 | 5 | 7 | 3 | 0 | 5 | 12 | 9 | 11 | 8 | 10 |
| LRC40 | Leucine-rich r | 2.69444679 | 0.01515569 | 8.7669E-05 | 0.00013279 | 8.7903E-05 | 0 | 0.00010279 | 0.00039165 | 0.00033783 | 0.00020933 | 0.00016901 | 0.00027696 | 2 | 3 | 2 | 0 | 2.33333333 | 7 | 6 | 3 | 3 | 4.75 |
| LC7L2 | Putative RNA | 2.69162013 | 0.0001952 | 0.00047122 | 0.00033988 | 0.00040498 | 0.00068025 | 0.00047408 | 0.0012029 | 0.0013835 | 0.0013931 | 0.0011247 | 0.00127605 | 7 | 5 | 6 | 10 | 7 | 14 | 16 | 13 | 13 | 14 |
| CRBG3 | Very large A- | 2.68616559 | 0.19926166 | 0.00022212 | 0.00023327 | 0.00028508 | 0.00021548 | 0.00023899 | 0.00051033 | 0.00073041 | 0 | 0.00068514 | 0.00064196 | 25 | 26 | 32 | 24 | 26.75 | 45 | 64 | 0 | 60 | 56.3333333 |
| ASCC3 | Activating sig | 2.68505546 | 8.3869E-05 | 0.00013182 | 0.00018152 | 0.00018024 | 0.00018165 | 0.00016881 | 0.00052006 | 0.00040022 | 0.00047691 | 0.00041584 | 0.00045326 | 11 | 15 | 15 | 15 | 14 | 34 | 26 | 25 | 27 | 28 |
| K2C1 | Keratin, type | 2.67628541 | 0.04660015 | 0.0031142 | 0.0027722 | 0.0030403 | 0.0023187 | 0.00281135 | 0.0057531 | 0.0039475 | 0.012654 | 0.0077413 | 0.00752398 | 76 | 67 | 74 | 56 | 68.25 | 110 | 75 | 194 | 147 | 131.5 |
| SMC3 | Structural ma | 2.67010586 | 2.0464E-05 | 8.6733E-05 | 0.00013137 | 8.6964E-05 | 0.00013147 | 0.00010913 | 0.00030443 | 0.00030637 | 0.00027613 | 0.00027867 | 0.0002914 | 4 | 6 | 4 | 6 | 5 | 11 | 11 | 8 | 10 | 10 |
| RS16 | 40S ribosoma | 2.6618669 | 0.00042917 | 0.00054223 | 0.00054753 | 0.00054367 | 0.00036528 | 0.00049968 | 0.0016149 | 0.001393 | 0.0011509 | 0.0011615 | 0.00133008 | 3 | 3 | 3 | 2 | 2.75 | 7 | 6 | 4 | 5 | 5.5 |
| RUFY1 | RUN and FYV | 2.64458746 | 0.27753676 | 7.4544E-05 | 7.5273E-05 | 0 | 7.5327E-05 | 7.5048E-05 | 0 | 0.00014363 | 0.00035598 | 9.5803E-05 | 0.00019847 | 2 | 2 | 0 | 2 | 2 | 0 | 3 | 6 | 2 | 3.66666667 |
| K1C9 | Keratin, type | 2.64384978 | 0.02125908 | 0.0018214 | 0.0015825 | 0.0017837 | 0.0015837 | 0.00169283 | 0.0035141 | 0.0027748 | 0.0068774 | 0.004736 | 0.00447558 | 43 | 37 | 42 | 37 | 39.75 | 65 | 51 | 102 | 87 | 76.25 |
| ATX2L | Ataxin-2-like | 2.63124283 | 0.01857857 | 0.00054004 | 0.00071884 | 0.00059071 | 0.0004961 | 0.00058642 | 0.0017546 | 0.0019234 | 0.00066428 | 0.0018298 | 0.00154302 | 22 | 29 | 24 | 20 | 23.75 | 56 | 61 | 17 | 58 | 48 |
| DIC | Mitochondria | 2.63055761 | 0.00350772 | 0.00027584 | 0.00055707 | 0.00046095 | 0.00037165 | 0.00041638 | 0.0011736 | 0.00070862 | 0.0013173 | 0.0011817 | 0.00109531 | 3 | 6 | 5 | 4 | 4.5 | 10 | 6 | 9 | 10 | 8.75 |
| ABCF2 | ATP-binding | 2.63020934 | 0.00117473 | 0.0002965 | 0.00034217 | 0.00021235 | 0.00038522 | 0.00030906 | 0.00097314 | 0.00087051 | 0.00080911 | 0.00059881 | 0.00081289 | 7 | 8 | 5 | 9 | 7.25 | 18 | 16 | 12 | 11 | 14.25 |
| DC1L1 | Cytoplasmic | 2.62917573 | 0.01966937 | 0 | 0 | 0.00010118 | 0 | 0.00010118 | 0.0001932 | 0.00019443 | 0.00048191 | 0.00019454 | 0.00026602 | 0 | 0 | 2 | 0 | 2 | 3 | 3 | 6 | 3 | 3.75 |
| IPO7 | Importin-7 O | 2.59991773 | 0.00210129 | 0.0002288 | 0.00033372 | 0.00015294 | 0.00025689 | 0.00024309 | 0.00071387 | 0.00048982 | 0.0007689 | 0.00055544 | 0.00063201 | 9 | 13 | 6 | 10 | 9.5 | 24 | 16 | 23 | 19 | 20.5 |
| SYMC | Methionine-- | 2.59361729 | 0.00063205 | 0.00017592 | 0.00020725 | 0.00026459 | 0.00014814 | 0.00019898 | 0.00041166 | 0.0004896 | 0.00056008 | 0.00060292 | 0.00051607 | 6 | 7 | 9 | 5 | 6.75 | 11 | 13 | 12 | 16 | 13 |
| RS11 | 40S ribosoma | 2.5923837 | 0.00093403 | 0.0016702 | 0.0023611 | 0.0015071 | 0.0011814 | 0.00167995 | 0.0053294 | 0.0045051 | 0.0037221 | 0.0038637 | 0.00435508 | 10 | 14 | 9 | 7 | 10 | 25 | 21 | 14 | 18 | 19.5 |
| ERLN2 | Erlin-2 OS=Ho | 2.58245704 | 0.00127318 | 0.00015568 | 0 | 0.0003122 | 0 | 0.00023394 | 0.00049678 | 0.00059992 | 0.00061956 | 0.0007003 | 0.00060414 | 2 | 0 | 4 | 0 | 3 | 5 | 6 | 5 | 7 | 5.75 |
| UBP10 | Ubiquitin car | 2.57116775 | 0.03711803 | 0.00056216 | 0.00060105 | 0.00056366 | 0.00056806 | 0.00057373 | 0.0017305 | 0.0019539 | 0.00047375 | 0.0017425 | 0.00147516 | 17 | 18 | 17 | 17 | 17.25 | 41 | 46 | 9 | 41 | 34.25 |
| SUCB1 | Succinate--Co | 2.5582837 | 0.01153881 | 0.00017098 | 0.0001151 | 0 | 0.00011519 | 0.00013376 | 0.00029099 | 0.00029283 | 0.00027218 | 0.00051275 | 0.00034219 | 3 | 2 | 0 | 2 | 2.33333333 | 4 | 4 | 3 | 7 | 4.5 |
| IFT46 | Intraflagellar | 2.55544967 | 0.00078525 | 0.00060763 | 0.00078888 | 0.00052221 | 0.00052629 | 0.00061125 | 0.0014403 | 0.0012265 | 0.0017963 | 0.001785 | 0.00156203 | 7 | 9 | 6 | 6 | 7 | 13 | 11 | 13 | 16 | 13.25 |
| PRS4 | 26S proteaso | 2.54949448 | 0.29152858 | 0.00011995 | 0.00024224 | 0.00012027 | 0.00012121 | 0.00015092 | 0.00022965 | 0.00053925 | 0 | 0.00038539 | 0.00038476 | 2 | 4 | 2 | 2 | 2.5 | 3 | 7 | 0 | 5 | 5 |
| GRDN | Girdin OS=Ho | 2.54782057 | 0.13983086 | 2.8208E-05 | 0 | 0 | 2.8504E-05 | 2.8356E-05 | 0.00010801 | 5.4349E-05 | 0 | 5.4379E-05 | 7.2246E-05 | 2 | 0 | 0 | 2 | 2 | 6 | 3 | 0 | 3 | 4 |
| PKHA5 | Pleckstrin ho | 2.54385842 | 0.14778358 | 4.7291E-05 | 0.00007163 | 0 | 9.5576E-05 | 7.1499E-05 | 0.00018108 | 0.00018223 | 0 | 0.00018234 | 0.00018188 | 2 | 3 | 0 | 4 | 3 | 6 | 6 | 0 | 6 | 6 |
| RS15A | 40S ribosoma | 2.53435079 | 0.00701806 | 0.0034508 | 0.0047144 | 0.0046812 | 0.0043075 | 0.00428848 | 0.012695 | 0.01408 | 0.0067856 | 0.0099134 | 0.0108685 | 17 | 23 | 23 | 21 | 21 | 49 | 54 | 21 | 38 | 40.5 |
| RPN2 | Dolichyl-diph | 2.5329989 | 0.00023192 | 0.00029274 | 0.00025337 | 0.00029352 | 0.00016904 | 0.00025217 | 0.00064054 | 0.00053717 | 0.00073228 | 0.00064497 | 0.00063874 | 7 | 6 | 7 | 4 | 6 | 12 | 10 | 11 | 12 | 11.25 |
| NMD3 | 60S ribosoma | 2.52662918 | 0.0112249 | 0.00010492 | 0.00015893 | 0.0001052 | 0 | 0.00012302 | 0.00033481 | 0.00040432 | 0.00016702 | 0.00033712 | 0.00031082 | 2 | 3 | 2 | 0 | 2.33333333 | 5 | 6 | 2 | 5 | 4.5 |
| KIF1B | Kinesin-like p | 2.52470356 | 0.00873409 | 8.7186E-05 | 0.00014673 | 0.00010199 | 0.00011747 | 0.00011334 | 0.00022257 | 0.00041063 | 0.00023131 | 0.00028013 | 0.00028616 | 7 | 11 | 9 | 10 | 9.25 | 157 | 172 | 126 | 169 | 156 |
| DYHC1 | Cytoplasmic | 2.5213958 | 0.00124777 | 0.00013632 | 0.00012618 | 0.00016515 | 0.00014349 | 0.00014279 | 0.00039148 | 0.00042315 | 0.00025316 | 0.00037228 | 0.00036002 | 24 | 22 | 29 | 25 | 25 | 54 | 58 | 28 | 51 | 47.75 |
| ZCCHV | Zinc finger CC | 2.51742806 | 5.1394E-05 | 0.00081915 | 0.00067946 | 0.00085067 | 0.00088688 | 0.00080904 | 0.0019417 | 0.0021795 | 0.0017697 | 0.0022559 | 0.0020367 | 28 | 23 | 29 | 30 | 27.5 | 52 | 58 | 38 | 60 | 52 |
| KC1E | Casein kinase | 2.51539994 | 0.00647212 | 0.00012687 | 0.00025622 | 0.00012721 | 0.0001923 | 0.00017565 | 0.00040483 | 0.00048888 | 0.00030293 | 0.00057068 | 0.00044183 | 2 | 4 | 2 | 3 | 2.75 | 5 | 6 | 3 | 7 | 5.25 |
| ANR17 | Ankyrin repe | 2.51123335 | 0.27139686 | 4.0551E-05 | 6.1421E-05 | 8.1317E-05 | 5.1221E-05 | 5.8628E-05 | 0.00018115 | 9.1152E-05 | 0 | 0.00016938 | 0.00014723 | 8 | 12 | 11 | 10 | 10.25 | 21 | 16 | 0 | 21 | 19.3333333 |
| IFT22 | Intraflagellar | 2.51044695 | 0.00850229 | 0.00085584 | 0.00028807 | 0.0007151 | 0.00072069 | 0.00064493 | 0.0014565 | 0.0012825 | 0.0022706 | 0.0014666 | 0.00161905 | 6 | 2 | 5 | 5 | 4.5 | 8 | 7 | 10 | 8 | 8.25 |
| DCTN1 | Dynactin sub | 2.50810135 | 0.0759345 | 0.00012389 | 0.00006255 | 4.1406E-05 | 0.00008346 | 7.7827E-05 | 0.00018448 | 0.00031827 | 6.5737E-05 | 0.0002123 | 0.0001952 | 6 | 3 | 2 | 4 | 3.75 | 7 | 12 | 2 | 8 | 7.25 |
| TTK | Dual specifici | 2.50163643 | 0.21955361 | 0.00015396 | 0.00012437 | 0.00015437 | 9.3345E-05 | 0.00013151 | 0.00031441 | 0.00031641 | 0 | 0.00035616 | 0.00032899 | 5 | 4 | 5 | 3 | 4.25 | 8 | 8 | 0 | 9 | 8.33333333 |
| RL28 | 60S ribosoma | 2.49956973 | 0.00337751 | 0.0015409 | 0.0013615 | 0.00038626 | 0.00077856 | 0.00101681 | 0.0027044 | 0.0019793 | 0.0027595 | 0.0027231 | 0.00254158 | 8 | 7 | 2 | 4 | 5.25 | 11 | 8 | 9 | 11 | 9.75 |
| SERPH | Serpin H1 OS | 2.49471553 | 0.17203793 | 0.00012626 | 0.0001275 | 0.0001266 | 0 | 0.00012679 | 0.00024173 | 0 | 0.00030148 | 0.00040568 | 0.0003163 | 2 | 2 | 2 | 0 | 2 | 3 | 0 | 3 | 5 | 3.66666667 |
| K2C6A | Keratin, type | 2.49250143 | 0.04023392 | 0.00028073 | 0.00033072 | 0 | 0 | 0.00030573 | 0.0010749 | 0.00042069 | 0.0011917 | 0.00036079 | 0.00076202 | 17 | 17 | 0 | 0 | 17 | 46 | 12 | 50 | 30 | 34.5 |
| IF2B3 | Insulin-like gr | 2.48157363 | 0.00241206 | 0.00022788 | 0.00027613 | 0.00027418 | 0.00013816 | 0.00022909 | 0.00069806 | 0.00040979 | 0.0005804 | 0.00058574 | 0.0005685 | 10 | 10 | 12 | 8 | 10 | 20 | 13 | 14 | 17 | 16 |
| PP6R1 | Serine/threo | 2.48063893 | 0.00201611 | 0.00026958 | 0.00024197 | 0.00024026 | 0.00027241 | 0.00025606 | 0.00076462 | 0.00065406 | 0.00042912 | 0.00069292 | 0.00063518 | 9 | 8 | 8 | 9 | 8.5 | 20 | 17 | 9 | 18 | 16 |
| PPP6 | Serine/threo | 2.4773343 | 0.0001764 | 0.00051912 | 0.00052419 | 0.00043375 | 0.000612 | 0.00052227 | 0.0012147 | 0.0011113 | 0.001515 | 0.0013343 | 0.00129383 | 6 | 6 | 5 | 7 | 6 | 11 | 10 | 11 | 12 | 11 |
| DI3L1 | DIS3-like exo | 2.45915164 | 0.24335853 | 0.00025036 | 0.00032866 | 0.00030124 | 0.00030359 | 0.00029596 | 0.00060716 | 0.00067534 | 0 | 0.00090095 | 0.00072782 | 10 | 13 | 12 | 12 | 11.75 | 19 | 21 | 0 | 28 | 22.6666667 |
| VPS45 | Vacuolar pro | 2.45316542 | 0.01269353 | 0.00074073 | 0.00070122 | 0.00097479 | 0.00084207 | 0.0008147 | 0.0014182 | 0.0019029 | 0.0029478 | 0.0017255 | 0.0019986 | 16 | 15 | 21 | 18 | 17.5 | 24 | 32 | 40 | 29 | 31.25 |
| RL15 | 60S ribosoma | 2.44472651 | 0.00023931 | 0.0012935 | 0.00091434 | 0.001297 | 0.000915 | 0.00110496 | 0.0023115 | 0.0031569 | 0.0026769 | 0.00266 | 0.00270133 | 10 | 7 | 10 | 7 | 8.5 | 14 | 19 | 13 | 16 | 15.5 |
| SCYL2 | SCY1-like pro | 2.43933288 | 0.33214332 | 0.00005681 | 5.7366E-05 | 5.6962E-05 | 5.7407E-05 | 5.7136E-05 | 0.00010877 | 0.00021892 | 9.0433E-05 | 0 | 0.00013937 | 2 | 2 | 2 | 2 | 2 | 3 | 6 | 2 | 0 | 3.66666667 |
| IFT74 | Intraflagellar | 2.43367183 | 0.01377777 | 0.00052777 | 0.00093263 | 0.00088196 | 0.00079997 | 0.00078558 | 0.0019086 | 0.0015253 | 0.0028004 | 0.0014131 | 0.00191185 | 12 | 21 | 20 | 18 | 17.75 | 34 | 27 | 40 | 25 | 31.5 |
| TIM50 | Mitochondria | 2.42933564 | 0.01072631 | 0.00029902 | 0.00030194 | 0.00022486 | 0.00022662 | 0.00026311 | 0.00076332 | 0.00057613 | 0.00083298 | 0.0003843 | 0.00063918 | 4 | 4 | 3 | 3 | 3.5 | 8 | 6 | 7 | 4 | 6.25 |
| NUFP2 | Nuclear fragi | 2.41562399 | 0.04742408 | 0.0006075 | 0.00046008 | 0.00060912 | 0.00061388 | 0.00057265 | 0.0018416 | 0.0016582 | 0.00042308 | 0.0016103 | 0.0013833 | 16 | 12 | 16 | 16 | 15 | 38 | 34 | 7 | 33 | 28 |
| ATPB | ATP synthase | 2.41477869 | 0.05123336 | 0.00079814 | 0.00075557 | 0.00075025 | 0.00055448 | 0.00071461 | 0.0014007 | 0.0010893 | 0.002938 | 0.0014745 | 0.00172563 | 16 | 15 | 15 | 11 | 14.25 | 22 | 17 | 37 | 23 | 24.75 |
| DJC13 | DnaJ homolo | 2.39555834 | 0.00396272 | 0.00002353 | 0 | 2.3592E-05 | 0 | 2.3561E-05 | 4.5049E-05 | 6.0447E-05 | 7.4911E-05 | 0.00004536 | 5.6442E-05 | 2 | 0 | 2 | 0 | 2 | 3 | 4 | 4 | 3 | 3.5 |
| GGYF2 | GRB10-intera | 2.39031101 | 0.0007716 | 0.00034534 | 0.00026667 | 0.00026479 | 0.00024633 | 0.00028078 | 0.00080379 | 0.00073062 | 0.00054973 | 0.00060049 | 0.00067116 | 17 | 13 | 13 | 12 | 13.75 | 31 | 28 | 17 | 23 | 24.75 |
| DIM1 | Probable dim | 2.38646586 | 0.00041466 | 0.00025292 | 0.0002554 | 0.00033813 | 0.00034077 | 0.00029681 | 0.00064565 | 0.00064976 | 0.00067103 | 0.00086682 | 0.00070832 | 3 | 3 | 4 | 4 | 3.5 | 6 | 6 | 5 | 8 | 6.25 |
| PCH2 | Pachytene ch | 2.36902953 | 0.00036555 | 0.00012217 | 0.00012336 | 0.00018374 | 0.00012345 | 0.00013818 | 0.00038983 | 0.00031385 | 0.00029171 | 0.00031402 | 0.00032735 | 2 | 2 | 3 | 2 | 2.25 | 5 | 4 | 3 | 4 | 4 |
| SSRD | Translocon-a | 2.36525835 | 0.00160106 | 0.00091521 | 0.0010782 | 0.0007647 | 0.00077068 | 0.0008822 | 0.002531 | 0.0017634 | 0.0016997 | 0.0023524 | 0.00208663 | 6 | 7 | 5 | 5 | 5.75 | 13 | 9 | 7 | 12 | 10.25 |
| TWF2 | Twinfilin-2 O | 2.35925697 | 0.17351807 | 0 | 0.00022905 | 0.00015163 | 0.00015281 | 0.00017783 | 0 | 0.00038849 | 0.00048145 | 0.0003887 | 0.00041955 | 0 | 3 | 2 | 2 | 2.33333333 | 0 | 4 | 4 | 4 | 4 |
| SYEP | Bifunctional g | 2.35799534 | 0.000279 | 0.00022688 | 0.00015861 | 0.00024499 | 0.000194 | 0.00020612 | 0.00042325 | 0.00051561 | 0.00044451 | 0.00056075 | 0.00048603 | 13 | 9 | 14 | 11 | 11.75 | 19 | 23 | 16 | 25 | 20.75 |
| CKAP5 | Cytoskeleton | 2.34834414 | 0.02608176 | 0.00023376 | 0.00013113 | 0.0002474 | 0.00023621 | 0.00021213 | 0.00053042 | 0.00058383 | 0.0002274 | 0.00065092 | 0.00049814 | 18 | 10 | 19 | 18 | 16.25 | 32 | 35 | 11 | 39 | 29.25 |
| ANKH1 | Ankyrin repe | 2.34676073 | 0.0606251 | 0.00012457 | 0.00015724 | 0.00011449 | 0.00011539 | 0.00012792 | 0.00033125 | 0.00040003 | 8.2624E-05 | 0.00038691 | 0.0003002 | 12 | 15 | 11 | 11 | 12.25 | 25 | 30 | 5 | 29 | 22.25 |
| DNJA3 | DnaJ homolo | 2.34614111 | 0.00891377 | 0.00016493 | 0.00033308 | 0.00022049 | 0.00027777 | 0.00024907 | 0.00063153 | 0.00049431 | 0.00078762 | 0.00042393 | 0.00058435 | 3 | 6 | 4 | 5 | 4.5 | 9 | 7 | 9 | 6 | 7.75 |
| RTCB | tRNA-splicing | 2.33340098 | 0.00147355 | 0.00041803 | 0.00047489 | 0.00031436 | 0.00036962 | 0.00039423 | 0.0010671 | 0.0010068 | 0.00066544 | 0.0009402 | 0.00091989 | 8 | 9 | 6 | 7 | 7.5 | 16 | 15 | 8 | 14 | 13.25 |
| XRN1 | 5'-3' exoribor | 2.32660112 | 0.01590406 | 0.00047951 | 0.00037486 | 0.00027917 | 0.00048455 | 0.00040452 | 0.0010464 | 0.0011921 | 0.00049245 | 0.0010337 | 0.00094116 | 31 | 24 | 18 | 31 | 26 | 53 | 60 | 20 | 52 | 46.25 |
| HAUS5 | HAUS augmin | 2.30727211 | 0.01805833 | 0.00025013 | 0.00033676 | 0.00041799 | 0.00029488 | 0.00032494 | 0.0005853 | 0.00064257 | 0.0011281 | 0.00064293 | 0.00074973 | 6 | 8 | 10 | 7 | 7.75 | 11 | 12 | 17 | 12 | 13 |
| SFXN1 | Sideroflexin-1 | 2.30235386 | 0.00591506 | 0.00049171 | 0.00082753 | 0.00065736 | 0.00074531 | 0.00068048 | 0.001046 | 0.0017895 | 0.0019568 | 0.0014745 | 0.0015667 | 6 | 10 | 8 | 9 | 8.25 | 10 | 17 | 15 | 14 | 14 |
| DHX57 | Putative ATP- | 2.29718758 | 0.01069916 | 0.00013327 | 5.7676E-05 | 0.00009545 | 7.6957E-05 | 9.0838E-05 | 0.00026731 | 0.00024456 | 0.00015154 | 0.00017128 | 0.00020867 | 7 | 4 | 5 | 4 | 5 | 11 | 11 | 5 | 7 | 8.5 |
| PSDE | 26S proteaso | 2.2969077 | 0.00325051 | 0 | 0 | 0 | 0.00017204 | 0.00017204 | 0.0004346 | 0.00032802 | 0.00027101 | 0.00054701 | 0.00039516 | 0 | 0 | 0 | 2 | 2 | 4 | 3 | 2 | 5 | 3.5 |
| IF4B | Eukaryotic tr | 2.29327617 | 0.00030849 | 0.00051827 | 0.00056695 | 0.00047634 | 0.00052371 | 0.00052132 | 0.0013781 | 0.0012759 | 0.0009625 | 0.0011656 | 0.00119553 | 12 | 13 | 11 | 12 | 12 | 25 | 23 | 14 | 21 | 20.75 |
| PSA4 | Proteasome s | 2.29033178 | 0.0004537 | 0.00020221 | 0.00030628 | 0.00030412 | 0.00020433 | 0.00025424 | 0.00051619 | 0.00051947 | 0.00064377 | 0.0006497 | 0.00058228 | 2 | 3 | 3 | 2 | 2.5 | 4 | 4 | 4 | 5 | 4.25 |
| IF4A1 | Eukaryotic in | 2.28909618 | 1.8996E-05 | 0.0023399 | 0.0025596 | 0.0028023 | 0.0023644 | 0.00251655 | 0.0051435 | 0.0060945 | 0.0062078 | 0.0055967 | 0.00576063 | 36 | 39 | 43 | 36 | 38.5 | 62 | 73 | 60 | 67 | 65.5 |
| TNR6B | Trinucleotide | 2.2774568 | 0.13401731 | 0.00021594 | 0.00018898 | 0.00018765 | 0.00011638 | 0.00017724 | 0.0005145 | 0.00064722 | 4.5833E-05 | 0.00040705 | 0.00040365 | 15 | 13 | 13 | 8 | 12.25 | 28 | 35 | 2 | 22 | 21.75 |
| DHX30 | Putative ATP- | 2.27687438 | 0.00179657 | 0.00019891 | 0.00015622 | 0.00019944 | 0.00015633 | 0.00017773 | 0.00045134 | 0.00028388 | 0.00045735 | 0.00042606 | 0.00040466 | 9 | 7 | 9 | 7 | 8 | 16 | 10 | 13 | 15 | 13.5 |
| HAUS3 | HAUS augmin | 2.27362002 | 0.01433334 | 8.7524E-05 | 0.00026514 | 0.00026327 | 0.00022111 | 0.00020926 | 0.00055857 | 0.00028106 | 0.0005573 | 0.00050619 | 0.00047578 | 2 | 6 | 6 | 5 | 4.75 | 10 | 5 | 8 | 9 | 8 |
| CDK2 | Cyclin-depen | 2.2633826 | 0.01036545 | 0.00026566 | 0.00017884 | 0.00017758 | 0.00017896 | 0.00020026 | 0.00033908 | 0.00056872 | 0.00056384 | 0.00034142 | 0.00045327 | 4 | 2 | 3 | 3 | 3 | 3 | 6 | 4 | 4 | 4.25 |
| IF4G2 | Eukaryotic tr | 2.25974266 | 0.00267147 | 0.00066917 | 0.00064633 | 0.00058343 | 0.00091139 | 0.00070258 | 0.0018568 | 0.0014201 | 0.0012041 | 0.0018696 | 0.00158765 | 23 | 22 | 20 | 31 | 24 | 50 | 38 | 26 | 50 | 41 |
| RL18A | 60S ribosoma | 2.25176692 | 1.6227E-05 | 0.0013494 | 0.001514 | 0.0015033 | 0.0013636 | 0.00143258 | 0.0036361 | 0.0030814 | 0.0031027 | 0.0030831 | 0.00322583 | 9 | 10 | 10 | 9 | 9.5 | 19 | 16 | 13 | 16 | 16 |
| QCR2 | Cytochrome | 2.25167915 | 0.00276769 | 0.00099029 | 0.00082351 | 0.00046726 | 0.00064751 | 0.00073214 | 0.0017101 | 0.0013468 | 0.00204 | 0.0014973 | 0.00164855 | 17 | 14 | 8 | 11 | 12.5 | 23 | 18 | 22 | 20 | 20.75 |

| | | | | | | | | | | | | | | | | | | | | | |
|---|---|---|---|---|---|---|---|---|---|---|---|---|---|---|---|---|---|---|---|---|---|
| HNRPD | Heterogeneo | 2.24811219 | 0.00106227 | 0.000223 | 0.00022518 | 0.00044719 | 0.00030046 | 0.00029896 | 0.00066414 | 0.00076384 | 0.00059164 | 0.00066874 | 0.00067209 | 3 | 3 | 6 | 4 | 4 | 7 | 8 | 5 | 7 | 6.75 |
| CP131 | Centrosomal | 2.24218771 | 0.35017937 | 0.0001462 | 0.00014763 | 0.00019545 | 0.00012311 | 0.0001531 | 0.0004976 | 0.00025038 | 0 | 0.00028184 | 0.00034327 | 6 | 6 | 8 | 5 | 6.25 | 16 | 8 | 0 | 9 | 11 |
| RP9 | Retinitis pigr | 2.22862107 | 0.20708879 | 0.00023881 | 0.00024114 | 0.00023945 | 0 | 0.0002398 | 0.00060962 | 0.0006135 | 0.00038015 | 0 | 0.00053442 | 2 | 2 | 2 | 0 | 2 | 4 | 4 | 2 | 0 | 3.33333333 |
| RN219 | RING finger p | 2.21269509 | 0.00113268 | 0.0017447 | 0.0014314 | 0.0017493 | 0.0014692 | 0.00159865 | 0.0029692 | 0.0034549 | 0.0044552 | 0.00327 | 0.00353733 | 48 | 39 | 48 | 40 | 43.75 | 64 | 74 | 77 | 70 | 71.25 |
| TKT | Transketolas | 2.21069625 | 0.00490394 | 8.4714E-05 | 0 | 0.00016988 | 0 | 0.0001273 | 0.00027032 | 0.00032644 | 0.00020228 | 0.00032662 | 0.00028142 | 2 | 0 | 4 | 0 | 3 | 5 | 6 | 3 | 6 | 5 |
| PELO | Protein pelot | 2.20852906 | 0.15199578 | 0.00013708 | 0 | 0 | 0.00020778 | 0.00017243 | 0.00034994 | 0.00052824 | 0 | 0.00026427 | 0.00038082 | 2 | 0 | 0 | 3 | 2.5 | 4 | 6 | 0 | 3 | 4.33333333 |
| EIF3E | Eukaryotic tr | 2.17884259 | 0.00267858 | 0.0003558 | 0.00047904 | 0.00047566 | 0.00023969 | 0.00038755 | 0.0006812 | 0.0007617 | 0.00094396 | 0.00099076 | 0.00084441 | 6 | 8 | 8 | 4 | 6.5 | 9 | 10 | 10 | 13 | 10.5 |
| C1TM | Monofunctio | 2.16532042 | 0.12695646 | 8.0946E-05 | 5.4492E-05 | 0 | 0 | 6.7719E-05 | 0.00013776 | 0.00017329 | 0.00012885 | 0 | 0.00014663 | 3 | 2 | 0 | 0 | 2.5 | 4 | 5 | 3 | 0 | 4 |
| NBN | Nibrin OS=Hc | 2.16171951 | 0.29980757 | 0.00055997 | 0.00049476 | 0.00056146 | 0.00045975 | 0.00051899 | 0.0010274 | 0.0011239 | 0 | 0.0012144 | 0.0011219 | 16 | 14 | 16 | 13 | 14.75 | 23 | 25 | 0 | 27 | 25 |
| RL7 | 60S ribosoma | 2.16106975 | 0.00031893 | 0.00074483 | 0.00064467 | 0.00053344 | 0.00064514 | 0.00064202 | 0.0016298 | 0.0013668 | 0.0011857 | 0.0013675 | 0.00138745 | 7 | 6 | 5 | 6 | 6 | 12 | 10 | 7 | 10 | 9.75 |
| IFT81 | Intraflagellar | 2.1594154 | 0.00096597 | 0.00085879 | 0.00090661 | 0.00086108 | 0.00074948 | 0.00084399 | 0.0018933 | 0.0015042 | 0.002237 | 0.0016556 | 0.00182253 | 22 | 23 | 22 | 19 | 21.5 | 38 | 30 | 36 | 33 | 34.25 |
| CLAP2 | CLIP-associat | 2.15313239 | 0.34699286 | 0.00010196 | 8.2369E-05 | 4.0894E-05 | 8.2428E-05 | 7.6913E-05 | 0.00015617 | 0.00013097 | 0 | 0.00020967 | 0.0001656 | 5 | 4 | 2 | 4 | 3.75 | 6 | 5 | 0 | 8 | 6.33333333 |
| RL13A | 60S ribosoma | 2.15054636 | 0.00929007 | 0.0019499 | 0.0014439 | 0.0015641 | 0.0015763 | 0.00163355 | 0.0031525 | 0.0023376 | 0.0045524 | 0.0040096 | 0.00351303 | 15 | 11 | 12 | 12 | 12.5 | 19 | 14 | 22 | 24 | 19.75 |
| DDX1 | ATP-depende | 2.1479663 | 0.00046858 | 0.0005349 | 0.0003961 | 0.00057208 | 0.00046845 | 0.00049288 | 0.0010924 | 0.0010993 | 0.00085148 | 0.0011916 | 0.0010587 | 15 | 11 | 16 | 13 | 13.75 | 24 | 24 | 15 | 26 | 22.25 |
| HAUS8 | HAUS augmir | 2.13318754 | 0.00060332 | 0.00038617 | 0.00019497 | 0.00032267 | 0.00039023 | 0.00032351 | 0.0006572 | 0.00074405 | 0.00061472 | 0.00074446 | 0.00069011 | 6 | 3 | 5 | 6 | 5 | 8 | 9 | 6 | 9 | 8 |
| CDK1 | Cyclin-depen | 2.12910668 | 0.001003 | 0.0007108 | 0.00098691 | 0.0011581 | 0.00089783 | 0.00093841 | 0.0017011 | 0.0023967 | 0.0018387 | 0.0020554 | 0.00199798 | 9 | 11 | 14 | 11 | 11.25 | 15 | 22 | 13 | 19 | 17.25 |
| PTGR3 | Prostaglandi | 2.12247896 | 0.04275377 | 0.00020999 | 0.00014136 | 0.00014036 | 0.00014146 | 0.00015829 | 0.00017868 | 0.00026973 | 0.00044569 | 0.00044979 | 0.00033597 | 3 | 2 | 2 | 2 | 2.25 | 2 | 3 | 4 | 5 | 3.5 |
| LC7L3 | Luc7-like pro | 2.12106275 | 0.00076047 | 0.00048867 | 0.00043177 | 0.00055122 | 0.00037036 | 0.00046051 | 0.0010136 | 0.00086308 | 0.0011668 | 0.00086356 | 0.00097676 | 8 | 7 | 9 | 6 | 7.5 | 13 | 11 | 12 | 11 | 11.75 |
| YJ005 | Uncharacteri | 2.12050714 | 0.4009058 | 0.00014867 | 0.0003753 | 0.00014906 | 0.00022534 | 0.00022459 | 0.00056927 | 0.00057288 | 0 | 0.0002866 | 0.00047625 | 2 | 5 | 2 | 3 | 3 | 6 | 6 | 0 | 3 | 5 |
| MACF1 | Microtubule- | 2.12043688 | 0.27080216 | 1.0715E-05 | 7.2135E-06 | 0 | 1.4437E-05 | 1.0789E-05 | 3.1913E-05 | 1.3764E-05 | 0 | 2.2952E-05 | 2.2876E-05 | 3 | 2 | 0 | 4 | 3 | 7 | 3 | 0 | 5 | 5 |
| VP33B | Vacuolar pro | 2.11135532 | 0.03473843 | 0 | 0 | 8.5766E-05 | 0 | 8.5766E-05 | 0.00010918 | 0.00016481 | 0.00034041 | 0.00010993 | 0.00018108 | 0 | 0 | 2 | 0 | 2 | 2 | 3 | 5 | 2 | 3 |
| XPO1 | Exportin-1 O! | 2.09356867 | 0.00728841 | 7.3917E-05 | 0 | 0.00014823 | 0 | 0.00011107 | 0.00025159 | 0.00015824 | 0.00023533 | 0.000285 | 0.00023254 | 3 | 0 | 6 | 0 | 4.5 | 8 | 5 | 6 | 9 | 7 |
| YBOX1 | Nuclease-ser | 2.09244717 | 0.00011655 | 0.00032578 | 0.00024673 | 0.00032665 | 0.0002469 | 0.00028652 | 0.00062373 | 0.0006277 | 0.00051859 | 0.00062805 | 0.00059952 | 4 | 3 | 4 | 3 | 3.5 | 6 | 6 | 4 | 6 | 5.5 |
| SURF4 | Surfeit locus | 2.07331144 | 0.01565586 | 0.00049049 | 0.00059435 | 0.00029508 | 0.00049564 | 0.00046889 | 0.0011269 | 0.00088204 | 0.0012493 | 0.00063038 | 0.00097216 | 5 | 6 | 3 | 5 | 4.75 | 9 | 7 | 8 | 5 | 7.25 |
| PFKAL | ATP-depende | 2.07233207 | 0.01399658 | 0.00023682 | 0.0002733 | 0.00016961 | 0.00017093 | 0.00021267 | 0.000475 | 0.0003911 | 0.00059239 | 0.00030436 | 0.00044071 | 7 | 8 | 5 | 5 | 6.25 | 11 | 9 | 11 | 7 | 9.5 |
| 43345 | Septin-2 OS= | 2.07224273 | 0.0104628 | 0.00051169 | 0.00044288 | 0.00073293 | 0.00059093 | 0.00056961 | 0.0010263 | 0.0011267 | 0.001629 | 0.00093946 | 0.00118037 | 7 | 6 | 10 | 8 | 7.75 | 11 | 12 | 14 | 10 | 11.75 |
| RL11 | 60S ribosoma | 2.05264869 | 0.00967285 | 0.00074125 | 0.0007485 | 0.00059458 | 0.00089884 | 0.00074579 | 0.0013246 | 0.0020947 | 0.0011799 | 0.0015242 | 0.00153085 | 5 | 5 | 4 | 6 | 5 | 7 | 11 | 5 | 8 | 7.75 |
| VPS51 | Vacuolar pro | 2.05206002 | 0.00159705 | 0.0006749 | 0.0006815 | 0.00074436 | 0.00075018 | 0.00071274 | 0.0012491 | 0.0015171 | 0.0018264 | 0.0012577 | 0.00146258 | 20 | 20 | 22 | 22 | 21 | 29 | 35 | 34 | 29 | 31.75 |
| AT1A1 | Sodium/pota | 2.04531143 | 0.01341958 | 0.00005159 | 0 | 5.1728E-05 | 7.8198E-05 | 6.0505E-05 | 9.8773E-05 | 0.00013253 | 0.00016425 | 9.9456E-05 | 0.00012375 | 2 | 0 | 2 | 3 | 2.33333333 | 3 | 4 | 4 | 3 | 3.5 |
| EIF3A | Eukaryotic tr | 2.04227411 | 5.2009E-05 | 0.0003246 | 0.00028922 | 0.00044034 | 0.00032801 | 0.00034554 | 0.00073115 | 0.00071127 | 0.00066869 | 0.00071166 | 0.00070569 | 17 | 15 | 23 | 17 | 18 | 30 | 29 | 22 | 29 | 27.5 |
| RL23 | 60S ribosoma | 2.03838846 | 0.00022286 | 0.0033928 | 0.0041873 | 0.0037798 | 0.0034284 | 0.00369708 | 0.0079392 | 0.0075055 | 0.0084012 | 0.0062984 | 0.00753608 | 18 | 22 | 20 | 18 | 19.5 | 33 | 31 | 28 | 26 | 29.5 |
| ABCD3 | ATP-binding | 2.02810447 | 0.07004892 | 0.00024026 | 0.00040435 | 0.00020075 | 0.00012139 | 0.00024169 | 0.00030666 | 0.00072009 | 0.00057368 | 0.00036024 | 0.00049017 | 6 | 10 | 5 | 3 | 6 | 6 | 14 | 9 | 7 | 9 |
| HNRPC | Heterogeneo | 2.02749288 | 0.0577825 | 0.00068989 | 0.00026124 | 0.00034587 | 0.00017429 | 0.00036782 | 0.00055035 | 0.00099693 | 0.0005491 | 0.00088665 | 0.00074576 | 8 | 3 | 4 | 2 | 4.25 | 5 | 9 | 4 | 8 | 6.5 |
| EIF3G | Eukaryotic tr | 2.02430763 | 0.00014362 | 0.00049478 | 0.00041635 | 0.00041342 | 0.00049998 | 0.00045613 | 0.00084204 | 0.00095331 | 0.0010502 | 0.00084786 | 0.00092335 | 6 | 5 | 5 | 6 | 5.5 | 8 | 9 | 8 | 8 | 8.25 |
| CAND1 | Cullin-associ | 2.02138139 | 0.01411856 | 8.5816E-05 | 0.00017331 | 0.00010756 | 0.0001084 | 0.00011877 | 0.00030122 | 0.00022046 | 0.00027321 | 0.00016544 | 0.00024008 | 4 | 8 | 5 | 5 | 5.5 | 11 | 8 | 8 | 6 | 8.25 |
| PYGB | Glycogen pho | 2.01944067 | 0.00066937 | 0 | 6.3218E-05 | 0 | 0 | 6.3218E-05 | 0.00011986 | 0.00012062 | 0.00014949 | 0.00012069 | 0.00012767 | 0 | 3 | 0 | 0 | 3 | 3 | 3 | 5 | 4 | 3.75 |
| STRAP | Serine-threo | 2.01868449 | 0.00107145 | 0.00082935 | 0.00083746 | 0.00075596 | 0.00068569 | 0.00077712 | 0.0013473 | 0.0015495 | 0.0014402 | 0.001938 | 0.00156875 | 11 | 11 | 10 | 9 | 10.25 | 14 | 16 | 12 | 20 | 15.5 |
| STML2 | Stomatin-like | 2.0151227 | 0.00558135 | 0.00022237 | 0.00022455 | 0.00029729 | 0.00014981 | 0.00022351 | 0.00056767 | 0.00038085 | 0.00047198 | 0.00038106 | 0.00045039 | 3 | 3 | 4 | 2 | 3 | 6 | 4 | 4 | 4 | 4.5 |
| PCBP2 | Poly(rC)-bind | 2.0093089 | 0.0028938 | 0.00057838 | 0.00058403 | 0.00057992 | 0.00080362 | 0.00063649 | 0.0011073 | 0.0014858 | 0.0010358 | 0.0014867 | 0.0012789 | 15 | 19 | 17 | 19 | 17.5 | 19 | 24 | 13 | 20 | 19 |
| MARE2 | Microtubule- | 2.00302696 | 0.12207933 | 0 | 0.00016298 | 0 | 0 | 0.00016298 | 0.00051501 | 0 | 0.00025692 | 0.00020743 | 0.00032645 | 0 | 2 | 0 | 0 | 2 | 5 | 0 | 2 | 2 | 3 |
| EIF3F | Eukaryotic tr | 2.0000434 | 0.03456747 | 0.00051742 | 0.00052248 | 0.00074114 | 0.00052286 | 0.00057598 | 0.00094346 | 0.00094946 | 0.001765 | 0.00094998 | 0.00115198 | 7 | 7 | 10 | 7 | 7.75 | 10 | 10 | 15 | 10 | 11.25 |
| EIF3D | Eukaryotic tr | 1.99447524 | 0.00372563 | 0.00014446 | 0.00034037 | 0.00033798 | 0.0002433 | 0.00026653 | 0.00061463 | 0.00055668 | 0.00045992 | 0.0004951 | 0.00053158 | 3 | 7 | 7 | 5 | 5.5 | 10 | 9 | 6 | 8 | 8.25 |
| PYC | Pyruvate carl | 1.98766694 | 6.8905E-06 | 0.016689 | 0.018729 | 0.017025 | 0.017045 | 0.017372 | 0.035226 | 0.032457 | 0.037299 | 0.033137 | 0.03452975 | 745 | 828 | 758 | 753 | 771 | 1232 | 1128 | 1046 | 1151 | 1139.25 |
| MTAP | S-methyl-5'-t | 1.98489885 | 0.07810427 | 0.00037298 | 0.00037663 | 0.00028048 | 0.00047112 | 0.0003753 | 0.0007141 | 0.00071863 | 0.0011875 | 0.00035952 | 0.00074494 | 4 | 4 | 3 | 5 | 4 | 6 | 6 | 8 | 3 | 5.75 |
| HNRDL | Heterogeneo | 1.98005591 | 0.00170693 | 0 | 0 | 0 | 0.00012698 | 0.00012698 | 0.00024058 | 0.00024211 | 0.00020003 | 0.00032299 | 0.00025143 | 0 | 0 | 0 | 4 | 4 | 8 | 7 | 5 | 8 | 7 |
| ADT2 | ADP/ATP trar | 1.97783841 | 5.3493E-05 | 0.010183 | 0.010015 | 0.0095891 | 0.010201 | 0.00999703 | 0.018649 | 0.018426 | 0.022554 | 0.019461 | 0.0197725 | 115 | 112 | 108 | 114 | 112.25 | 165 | 162 | 160 | 171 | 164.5 |
| PHB | Prohibitin OS | 1.97700653 | 0.00037713 | 0.00097016 | 0.00088169 | 0.0011673 | 0.00068625 | 0.00092635 | 0.0018574 | 0.00162 | 0.0018532 | 0.001995 | 0.0018314 | 10 | 9 | 12 | 7 | 9.5 | 15 | 13 | 12 | 16 | 14 |
| RL31 | 60S ribosoma | 1.96719301 | 0.0453711 | 0.00084443 | 0.00042634 | 0.00063501 | 0.00042665 | 0.00058311 | 0.0016167 | 0.00081349 | 0.0013442 | 0.00081395 | 0.00114709 | 4 | 2 | 3 | 2 | 2.75 | 6 | 3 | 4 | 3 | 4 |
| NT5D2 | 5'-nucleotida | 1.96485377 | 0.02692367 | 0 | 0.00020497 | 0.00010176 | 0.00010256 | 0.00013643 | 0.00032386 | 0.00026074 | 0.00016156 | 0.0003261 | 0.00026807 | 0 | 4 | 2 | 2 | 2.66666667 | 5 | 4 | 2 | 5 | 4 |
| IF4G3 | Eukaryotic tr | 1.95416029 | 0.02972849 | 0.0010988 | 0.00099189 | 0.0011351 | 0.0011608 | 0.00109665 | 0.0026775 | 0.0023524 | 0.0010601 | 0.0024821 | 0.00214303 | 66 | 59 | 68 | 69 | 65.5 | 126 | 110 | 40 | 116 | 98 |
| IF2G | Eukaryotic tr | 1.95241429 | 0.01820057 | 0.00055908 | 0.00045164 | 0.0003924 | 0.00028247 | 0.0004214 | 0.00057087 | 0.00093357 | 0.001068 | 0.00071853 | 0.00082274 | 10 | 8 | 7 | 5 | 7.5 | 8 | 13 | 12 | 10 | 10.75 |
| SYFA | Phenylalanin | 1.94708464 | 0.00811522 | 0.00036362 | 0.00047208 | 0.0003125 | 0.00015747 | 0.00032642 | 0.00059672 | 0.00066724 | 0.0007442 | 0.00053409 | 0.00063556 | 7 | 9 | 6 | 3 | 6.25 | 9 | 10 | 9 | 8 | 9 |
| IFT52 | Intraflagellar | 1.93965471 | 0.02301382 | 0.00048308 | 0.00091464 | 0.00066601 | 0.00036612 | 0.00060746 | 0.0013873 | 0.0013962 | 0.0011535 | 0.00077607 | 0.00117827 | 8 | 15 | 11 | 6 | 10 | 18 | 18 | 12 | 10 | 14.5 |
| CNOT1 | CCR4-NOT tr | 1.93397607 | 0.00584656 | 0.00039982 | 0.00039252 | 0.00062361 | 0.00038158 | 0.00044938 | 0.00097813 | 0.0010271 | 0.00067182 | 0.00079933 | 0.0008691 | 36 | 35 | 56 | 34 | 40.25 | 69 | 72 | 38 | 56 | 58.75 |
| EIF3M | Eukaryotic tr | 1.93321364 | 0.06300621 | 0.00035279 | 0.00042748 | 0.00035373 | 0.00057039 | 0.0004261 | 0.0006304 | 0.00045315 | 0.0011232 | 0.0010882 | 0.00082374 | 5 | 6 | 5 | 8 | 6 | 7 | 5 | 10 | 12 | 8.5 |
| FR1OP | FGFR1 oncog | 1.92994437 | 0.00580632 | 0.00066136 | 0.00060105 | 0.00033156 | 0.00046782 | 0.00051545 | 0.0010974 | 0.00084951 | 0.00084223 | 0.00119 | 0.00099479 | 10 | 9 | 5 | 7 | 7.75 | 13 | 10 | 8 | 14 | 11.25 |
| AHSA1 | Activator of ! | 1.92875925 | 0.01739391 | 0.00054651 | 0.00063069 | 0 | 0.00063114 | 0.00060278 | 0.0010961 | 0.001404 | 0.00074567 | 0.0014047 | 0.00116262 | 7 | 8 | 0 | 8 | 7.66666667 | 11 | 14 | 6 | 14 | 11.25 |
| HIG1A | HIG1 domain | 1.9229355 | 0.09492989 | 0.00056749 | 0 | 0 | 0 | 0.00056749 | 0.0010865 | 0.0014579 | 0 | 0.00072934 | 0.00109125 | 2 | 0 | 0 | 0 | 2 | 3 | 4 | 0 | 2 | 3 |
| UN45A | Protein unc-4 | 1.92072954 | 0.01644473 | 0.00011182 | 0.00022582 | 0.00014014 | 0.00014124 | 0.00015476 | 0.00024976 | 0.00028725 | 0.00040048 | 0.00025148 | 0.00029724 | 4 | 8 | 5 | 5 | 5.5 | 7 | 8 | 9 | 7 | 7.75 |
| NAV1 | Neuron navig | 1.91865009 | 0.43807775 | 0.00011247 | 0.00012777 | 8.4578E-05 | 0.00011365 | 0.00010962 | 0.00019739 | 0.00016253 | 0 | 0.00027103 | 0.00021032 | 8 | 9 | 6 | 8 | 7.75 | 11 | 9 | 0 | 15 | 11.6666667 |
| CNOT2 | CCR4-NOT tr | 1.91046214 | 0.08610382 | 0.00034207 | 0.00029607 | 0.00048998 | 0.00034567 | 0.00036845 | 0.0010603 | 0.0006277 | 0.00031116 | 0.00081646 | 0.00070391 | 7 | 6 | 10 | 7 | 7.5 | 17 | 10 | 4 | 13 | 11 |
| P5CR2 | Pyrroline-5-c | 1.90891979 | 0.04446026 | 0.00032986 | 0.00049962 | 0.00016537 | 0.00033332 | 0.00033204 | 0.00042102 | 0.00084739 | 0.00052508 | 0.00074188 | 0.00063384 | 4 | 6 | 2 | 4 | 4 | 4 | 8 | 4 | 7 | 5.75 |
| C1TC | C-1-tetrahyd | 1.90861636 | 0.02589518 | 0.00014111 | 0.00014249 | 0.00016979 | 0.00017112 | 0.00015613 | 0.00021614 | 0.00039877 | 0.00035941 | 0.00021763 | 0.00029799 | 5 | 5 | 6 | 6 | 5.5 | 6 | 11 | 8 | 6 | 7.75 |
| RAN | GTP-binding | 1.90558537 | 3.1809E-05 | 0.002932 | 0.0024673 | 0.0024499 | 0.0027159 | 0.00264128 | 0.0045221 | 0.0053354 | 0.0052508 | 0.0050244 | 0.00503318 | 24 | 20 | 20 | 22 | 21.5 | 29 | 34 | 27 | 32 | 30.5 |
| RS7 | 40S ribosoma | 1.90361726 | 0.00080492 | 0.0017683 | 0.0012362 | 0.0016366 | 0.0013745 | 0.0015039 | 0.0027779 | 0.003145 | 0.0023818 | 0.0031467 | 0.00286285 | 13 | 9 | 12 | 10 | 11 | 16 | 18 | 11 | 18 | 15.75 |
| HELZ | Probable heli | 1.9017066 | 0.09548102 | 0 | 0 | 0 | 2.7462E-05 | 2.7462E-05 | 6.9375E-05 | 3.4908E-05 | 0 | 5.2391E-05 | 5.2225E-05 | 0 | 0 | 0 | 2 | 2 | 4 | 2 | 0 | 3 | 3 |
| VDAC2 | Voltage-depe | 1.89653953 | 0.02188184 | 0.00071805 | 0.00063444 | 0.00062997 | 0.00081629 | 0.00069969 | 0.0013748 | 0.0011529 | 0.0018574 | 0.00092284 | 0.00132699 | 8 | 7 | 7 | 9 | 7.75 | 12 | 10 | 13 | 8 | 10.75 |
| LIPA1 | Liprin-alpha- | 1.89509504 | 0.5468437 | 8.7815E-05 | 0.00015518 | 4.4024E-05 | 8.8737E-05 | 9.3939E-05 | 0.00028021 | 0.000141 | 0 | 0.00011286 | 0.00017802 | 4 | 7 | 2 | 4 | 4.25 | 10 | 5 | 0 | 4 | 6.33333333 |
| RL26 | 60S ribosoma | 1.89450466 | 0.01887729 | 0.0010919 | 0.0012864 | 0.0010948 | 0.0012873 | 0.0011901 | 0.0020906 | 0.0016363 | 0.0031867 | 0.002105 | 0.00225465 | 6 | 7 | 6 | 7 | 6.5 | 9 | 7 | 11 | 9 | 9 |
| AIFM1 | Apoptosis-in | 1.88566503 | 0.00880801 | 0.00012914 | 0.00017388 | 0.00017265 | 0.000174 | 0.00016242 | 0.00038462 | 0.00022118 | 0.00034263 | 0.00027663 | 0.00030627 | 3 | 4 | 4 | 4 | 3.75 | 7 | 4 | 5 | 5 | 5.25 |
| RS6 | 40S ribosoma | 1.88447929 | 0.00858914 | 0.0029674 | 0.0035315 | 0.0032941 | 0.0033198 | 0.0032782 | 0.0047344 | 0.0059896 | 0.0082663 | 0.0057205 | 0.0061777 | 28 | 33 | 31 | 31 | 30.75 | 35 | 44 | 49 | 42 | 42.5 |
| TBA1B | Tubulin alpha | 1.88394569 | 4.7154E-05 | 0.0083085 | 0.0097487 | 0.0094453 | 0.0078637 | 0.00884155 | 0.017401 | 0.017812 | 0.0163 | 0.015115 | 0.016657 | 142 | 165 | 161 | 133 | 150.25 | 233 | 237 | 175 | 201 | 211.5 |
| RL13 | 60S ribosoma | 1.87821176 | 0.28154706 | 0.00037519 | 0.00063143 | 0.00037619 | 0 | 0.00046094 | 0.00079814 | 0 | 0.00099541 | 0.00080366 | 0.00086574 | 3 | 5 | 3 | 0 | 3.66666667 | 5 | 0 | 5 | 5 | 5 |
| NSDHL | Sterol-4-alph | 1.87806164 | 0.00220986 | 0.00035373 | 0.00021431 | 0.00049654 | 0.00035745 | 0.00035551 | 0.00072239 | 0.00063611 | 0.0006757 | 0.00063646 | 0.00066767 | 5 | 3 | 7 | 5 | 5 | 8 | 7 | 6 | 7 | 7 |
| PSMD3 | 26S proteaso | 1.8774394 | 0.00477035 | 0.00039533 | 0.0004491 | 0.00064413 | 0.00044942 | 0.0004845 | 0.00081996 | 0.00088865 | 0.00078663 | 0.0011432 | 0.00090961 | 8 | 9 | 13 | 9 | 9.75 | 13 | 14 | 10 | 18 | 13.75 |
| NDUAA | NADH dehyd | 1.87505348 | 0.03656381 | 0.00029733 | 0.00045036 | 0.00037266 | 0.00037557 | 0.00037398 | 0.00094878 | 0.00038192 | 0.00070996 | 0.00076427 | 0.00070123 | 4 | 6 | 5 | 5 | 5 | 10 | 4 | 6 | 8 | 7 |
| EXOC4 | Exocyst comp | 1.87502304 | 0.02207954 | 5.4186E-05 | 0 | 0.00005433 | 0 | 5.4258E-05 | 0.00013832 | 6.9601E-05 | 0.00012938 | 6.9639E-05 | 0.00010174 | 2 | 0 | 2 | 0 | 2 | 4 | 2 | 3 | 2 | 2.75 |
| P5CR1 | Pyrroline-5-c | 1.87089168 | 0.44711577 | 0.00024817 | 0.00016706 | 0.00024883 | 0.00016718 | 0.00020781 | 0.00042234 | 0.00031877 | 0 | 0.00042526 | 0.00038879 | 4 | 4 | 4 | 3 | 3.75 | 6 | 5 | 0 | 5 | 5.33333333 |
| ASNS | Asparagine s | 1.86797167 | 0.02285434 | 0 | 9.4996E-05 | 9.4327E-05 | 0 | 9.4662E-05 | 0.00012008 | 0.00024168 | 0.00022463 | 0.00012091 | 0.00017683 | 0 | 2 | 2 | 0 | 2 | 2 | 4 | 3 | 2 | 2.75 |
| DLDH | Dihydrolipoy | 1.85972625 | 0.02674444 | 0.00020737 | 0.00047116 | 0.00015595 | 0.00020955 | 0.00026101 | 0.0004632 | 0.00053274 | 0.00041263 | 0.00053304 | 0.0004854 | 4 | 9 | 3 | 4 | 5 | 7 | 8 | 5 | 8 | 7 |
| FAKD5 | FAST kinase c | 1.85758355 | 0.09327641 | 0 | 0 | 6.9264E-05 | 0 | 6.9264E-05 | 0.00013226 | 0 | 0.00016495 | 8.8781E-05 | 0.00012866 | 0 | 0 | 2 | 0 | 2 | 3 | 0 | 3 | 2 | 2.66666667 |
| MAGD2 | Melanoma-a | 1.85718011 | 0.02954304 | 0.00021773 | 0.0003078 | 0 | 8.8005E-05 | 0.00020451 | 0.00033348 | 0.0005034 | 0.00034659 | 0.00033579 | 0.00037982 | 6 | 9 | 0 | 4 | 6.33333333 | 7 | 11 | 5 | 6 | 7.25 |
| TMM33 | Transmembr | 1.85089245 | 0.03188658 | 0.00021367 | 0.00043152 | 0.00021424 | 0 | 0.00028648 | 0.00054545 | 0.00068615 | 0.00034013 | 0.00054922 | 0.00053024 | 2 | 4 | 2 | 0 | 2.66666667 | 4 | 5 | 2 | 4 | 3.75 |
| K1671 | Uncharacteri | 1.84972196 | 0.16440285 | 0.00032145 | 0.00028033 | 0.00023441 | 0.00028054 | 0.00027918 | 0.00046625 | 0.0007695 | 0.0001163 | 0.00071359 | 0.00051641 | 22 | 19 | 16 | 19 | 19 | 25 | 41 | 5 | 38 | 27.25 |
| RPN1 | Dolichyl-diph | 1.84896247 | 0.03304974 | 0.00021737 | 0.00021949 | 0.00026154 | 0.00021965 | 0.00022951 | 0.00038842 | 0.00027921 | 0.00041522 | 0.00061459 | 0.00042436 | 5 | 5 | 6 | 5 | 5.25 | 7 | 5 | 6 | 11 | 7.25 |
| RS3 | 40S ribosoma | 1.84265051 | 0.0062506 | 0.0041266 | 0.0029607 | 0.003702 | 0.0039505 | 0.00368495 | 0.0059601 | 0.0071139 | 0.0086432 | 0.0054431 | 0.00679008 | 38 | 27 | 34 | 36 | 33.75 | 43 | 51 | 50 | 39 | 45.75 |
| TBB5 | Tubulin beta | 1.84114268 | 2.2475E-05 | 0.0010104 | 0.0011403 | 0.0011918 | 0.0012012 | 0.00113593 | 0.0021241 | 0.0019849 | 0.0022706 | 0.001986 | 0.0020914 | 95 | 102 | 106 | 94 | 99.25 | 126 | 116 | 115 | 123 | 120 |
| FLII | Protein flight | 1.83543165 | 0.00526328 | 0 | 0 | 0.0000417 | 0 | 0.0000417 | 7.9625E-05 | 5.3421E-05 | 6.6204E-05 | 0.0001069 | 7.6538E-05 | 0 | 0 | 2 | 0 | 2 | 3 | 2 | 2 | 4 | 2.75 |
| TMEDA | Transmembr | 1.83152596 | 0.05763616 | 0.00036149 | 0.00060837 | 0.00048326 | 0.0006088 | 0.00051548 | 0.00092278 | 0.0012382 | 0.0011509 | 0.00046458 | 0.00094412 | 3 | 5 | 4 | 5 | 4.25 | 6 | 8 | 6 | 3 | 5.75 |
| COPD | Coatomer su | 1.83057717 | 0.02374756 | 0.00036149 | 0.00052146 | 0.00020711 | 0.00041747 | 0.00037688 | 0.00079096 | 0.00086231 | 0.00057543 | 0.00053095 | 0.00068991 | 7 | 10 | 4 | 8 | 7.25 | 12 | 13 | 7 | 8 | 10 |
| IF4G1 | Eukaryotic tr | 1.8285337 | 0.0003226 | 0.00057761 | 0.00063325 | 0.00061224 | 0.00060035 | 0.00060586 | 0.0012007 | 0.0012083 | 0.00091946 | 0.0011029 | 0.00110784 | 44 | 49 | 49 | 49 | 47.75 | 68 | 68 | 45 | 67 | 62 |
| LTV1 | Protein LTV1 | 1.82736739 | 0.10389897 | 0.00022222 | 0.00039269 | 0.00016711 | 0.00033683 | 0.00027971 | 0.00077999 | 0.00057087 | 0.0002653 | 0.00042839 | 0.00051114 | 4 | 7 | 3 | 6 | 5 | 11 | 8 | 3 | 6 | 7 |
| DDX20 | Probable ATF | 1.82540669 | 0.01896855 | 0.0001281 | 6.4676E-05 | 0.00009633 | 6.4722E-05 | 8.8457E-05 | 0.00012263 | 0.00020568 | 0.00015294 | 0.00016463 | 0.00016147 | 4 | 2 | 3 | 2 | 2.75 | 3 | 5 | 3 | 4 | 3.75 |
| EIF3H | Eukaryotic tr | 1.82359643 | 0.02361254 | 0 | 0.0001514 | 0 | 0 | 0.0001514 | 0.00019137 | 0.00019259 | 0.00023867 | 0.00048174 | 0.00027609 | 0 | 2 | 0 | 0 | 2 | 2 | 2 | 2 | 5 | 2.75 |
| ARL1 | ADP-ribosyla | 1.82127256 | 0.53089802 | 0.00058317 | 0.00044165 | 0.00043854 | 0.0005893 | 0.00051317 | 0.00093043 | 0.00056181 | 0 | 0.0013116 | 0.00093461 | 4 | 3 | 3 | 4 | 3.5 | 5 | 3 | 0 | 7 | 5 |
| PCNA | Proliferating | 1.81996646 | 0.02342495 | 0.00060663 | 0.00040838 | 0.00050687 | 0.000613 | 0.00053372 | 0.00090334 | 0.0010389 | 0.00064377 | 0.0012994 | 0.00097135 | 6 | 4 | 5 | 6 | 5.25 | 7 | 8 | 4 | 10 | 7.25 |
| TBCD4 | TBC1 domain | 1.8125165 | 0.02594444 | 0 | 6.1587E-05 | 0.00010192 | 4.1087E-05 | 6.8198E-05 | 0.0001038 | 0.00015668 | 0.00012945 | 0.00010451 | 0.00012361 | 0 | 3 | 5 | 2 | 3.33333333 | 4 | 6 | 4 | 4 | 4.5 |
| 43354 | Septin-11 OS | 1.81092984 | 0.04600811 | 0.00018453 | 0.00031057 | 0.00030838 | 0.00018647 | 0.00024749 | 0.00039256 | 0.00063209 | 0.00029375 | 0.00047433 | 0.00044818 | 3 | 6 | 7 | 5 | 5.25 | 6 | 11 | 6 | 8 | 7.75 |
| SMC2 | Structural ma | 1.80990604 | 0.01496956 | 0.00013227 | 0.00017809 | 0.00011052 | 0.00015594 | 0.00014421 | 0.00022511 | 0.00033981 | 0.00028074 | 0.00019833 | 0.000261 | 6 | 8 | 5 | 7 | 6.5 | 8 | 12 | 8 | 7 | 8.75 |
| EIF3I | Eukaryotic tr | 1.8082623 | 0.00413234 | 0.00097434 | 0.00098387 | 0.00097694 | 0.00098458 | 0.00097993 | 0.0018654 | 0.0016687 | 0.0021972 | 0.0013566 | 0.00177198 | 12 | 12 | 12 | 12 | 12 | 18 | 16 | 17 | 13 | 16 |
| RAD50 | DNA repair p | 1.80557082 | 0.08437977 | 0.00018102 | 0.00018279 | 0.00020167 | 0.0001626 | 0.00018202 | 0.00033374 | 0.0004392 | 0.00012807 | 0.00041359 | 0.00032865 | 9 | 9 | 10 | 8 | 9 | 13 | 17 | 4 | 16 | 12.5 |
| MYPT1 | Protein phos | 1.80493366 | 0.49942415 | 0.00010248 | 0.00010348 | 0.00010275 | 0.00010356 | 0.00010307 | 0.0002289 | 0.00013163 | 0 | 0.00019756 | 0.00018603 | 4 | 4 | 4 | 4 | 4 | 7 | 4 | 0 | 6 | 5.66666667 |
| MCM7 | DNA replicati | 1.8037951 | 0.00038154 | 0.00055052 | 0.00074121 | 0.00077278 | 0.00074174 | 0.00070156 | 0.0014054 | 0.0012729 | 0.00111 | 0.0012736 | 0.00126548 | 15 | 20 | 21 | 20 | 19 | 30 | 27 | 19 | 27 | 25.75 |
| NOC2L | Nucleolar co | 1.79994087 | 0.18066337 | 0.00021139 | 0.00024903 | 0.0001413 | 0.00014241 | 0.00018603 | 0.00026981 | 0.00022627 | 0.00061691 | 0.0002264 | 0.00033485 | 6 | 7 | 4 | 4 | 5.25 | 6 | 5 | 11 | 5 | 6.75 |
| STT3B | Dolichyl-diph | 1.79389017 | 0.07938291 | 0 | 6.4519E-05 | 0 | 0 | 6.4519E-05 | 0.00012233 | 0 | 0.00010171 | 0.00012318 | 0.00011574 | 0 | 2 | 0 | 0 | 2 | 3 | 0 | 2 | 3 | 2.66666667 |
| RL5 | 60S ribosoma | 1.79149971 | 0.05436832 | 0.00062195 | 0.00080747 | 0.001069 | 0.00035913 | 0.00071439 | 0.0011341 | 0.001826 | 0.0011315 | 0.0010277 | 0.00127983 | 7 | 9 | 12 | 4 | 8 | 10 | 16 | 8 | 9 | 10.75 |
| RFLB | Refilin-B OS= | 1.79026497 | 0.58365792 | 0.00036993 | 0.00037355 | 0.0012364 | 0.00037382 | 0.00058843 | 0.0012591 | 0.00095034 | 0 | 0.00095087 | 0.00105344 | 3 | 3 | 10 | 3 | 4.75 | 8 | 6 | 0 | 6 | 6.66666667 |
| GTPB1 | GTP-binding | 1.78809787 | 0.42551406 | 7.8889E-05 | 7.9661E-05 | 0.0001582 | 0 | 0.00010558 | 0.00015104 | 0 | 0.00031395 | 0.00010139 | 0.00018879 | 2 | 2 | 4 | 0 | 2.66666667 | 3 | 0 | 5 | 2 | 3.33333333 |
| UBAP2 | Ubiquitin-ass | 1.78673835 | 0.08241063 | 0.00044806 | 0.00033338 | 0.00030738 | 0.00042894 | 0.00037944 | 0.00078259 | 0.00081786 | 0.00026277 | 0.00084862 | 0.00067796 | 19 | 14 | 13 | 18 | 16 | 26 | 27 | 7 | 28 | 22 |
| CNOT3 | CCR4-NOT tr | 1.78551827 | 0.50589146 | 0.00021027 | 0.00021232 | 0.00017569 | 0.00010624 | 0.00017613 | 0.00031311 | 0.00036011 | 0 | 0.00027023 | 0.00031448 | 6 | 6 | 5 | 3 | 5 | 7 | 8 | 0 | 6 | 7 |
| TOM22 | Mitochondri | 1.78413368 | 0.00582758 | 0.0016725 | 0.0011259 | 0.0018633 | 0.0015023 | 0.001541 | 0.0033207 | 0.0021483 | 0.0026624 | 0.002866 | 0.00274935 | 9 | 6 | 10 | 8 | 8.25 | 14 | 9 | 9 | 12 | 11 |
| PIGU | Phosphatidyl | 1.78271568 | 0.0643612 | 0.00018199 | 0.00012251 | 0.00018247 | 0.0001226 | 0.00015239 | 0.00015486 | 0.00031168 | 0.00038626 | 0.00023389 | 0.00027167 | 3 | 2 | 3 | 2 | 2.5 | 2 | 4 | 4 | 3 | 3.25 |
| ALMS1 | Alstrom synd | 1.77972067 | 0.5163154 | 3.1664E-05 | 4.4763E-05 | 2.5398E-05 | 3.1996E-05 | 3.3455E-05 | 5.6581E-05 | 7.3209E-05 | 0 | 4.8833E-05 | 5.9541E-05 | 5 | 7 | 4 | 5 | 5.25 | 7 | 9 | 0 | 6 | 7.33333333 |
| RTRAF | RNA transcri | 1.7767503 | 0.56445908 | 0.0006489 | 0.00043683 | 0.00065062 | 0.00065571 | 0.00059802 | 0.0012424 | 0.0013892 | 0 | 0.00055597 | 0.00106252 | 6 | 4 | 6 | 6 | 5.5 | 9 | 10 | 0 | 4 | 7.66666667 |
| RAB21 | Ras-related p | 1.77412113 | 0.01921805 | 0.00058641 | 0.00047372 | 0.00058797 | 0.00059257 | 0.00056017 | 0.0011976 | 0.0010545 | 0.0011202 | 0.00060292 | 0.00099381 | 5 | 4 | 5 | 5 | 4.75 | 8 | 7 | 6 | 4 | 6.25 |
| ARGL1 | Arginine and | 1.77315839 | 0.01173427 | 0.00038664 | 0.00068324 | 0.00038767 | 0.0003907 | 0.00046206 | 0.00086363 | 0.0006208 | 0.00092321 | 0.0008696 | 0.00081931 | 4 | 7 | 4 | 4 | 4.75 | 7 | 5 | 6 | 7 | 6.25 |
| IMA1 | Importin sub | 1.76602843 | 0.01073377 | 0.00034919 | 0.00030223 | 0.0003001 | 0.00040326 | 0.0003387 | 0.00057303 | 0.0005126 | 0.00079407 | 0.00051288 | 0.00059815 | 7 | 6 | 6 | 8 | 6.75 | 9 | 8 | 10 | 8 | 8.75 |
| RL10 | 60S ribosoma | 1.76155477 | 0.00423604 | 0.00098648 | 0.00087161 | 0.00074183 | 0.00087224 | 0.00086804 | 0.0015739 | 0.0019007 | 0.001374 | 0.0012678 | 0.0015291 | 8 | 7 | 6 | 7 | 7 | 10 | 12 | 7 | 8 | 9.25 |
| ULA1 | NEDD8-activ | 1.76004951 | 0.02712048 | 0.00014825 | 0.0001996 | 0.00014864 | 0.00014981 | 0.00016158 | 0.00037844 | 0.0002539 | 0.00031465 | 0.00019053 | 0.00028438 | 3 | 4 | 3 | 3 | 3.25 | 6 | 4 | 4 | 3 | 4.25 |
| FA98A | Protein FAM! | 1.75902562 | 8.9626E-05 | 0.00040676 | 0.00046208 | 0.00035686 | 0.00041103 | 0.00040918 | 0.00077876 | 0.0007184 | 0.00072843 | 0.00065346 | 0.00071976 | 8 | 9 | 7 | 8 | 8 | 12 | 11 | 9 | 10 | 10.5 |
| HACD3 | Very-long-ch | 1.74938723 | 0.01008609 | 0.00051027 | 0.00058887 | 0.00058472 | 0.00036831 | 0.00051304 | 0.00083739 | 0.00074907 | 0.0011604 | 0.00084318 | 0.00089751 | 7 | 8 | 8 | 5 | 7 | 9 | 8 | 10 | 9 | 9 |
| G3BP2 | Ras GTPase-a | 1.74699871 | 0.02832311 | 0.00049273 | 0.00060811 | 0.00049404 | 0.00044258 | 0.00050937 | 0.00076867 | 0.00098452 | 0.00061005 | 0.0011962 | 0.00088986 | 13 | 15 | 11 | 11 | 12.5 | 14 | 16 | 9 | 20 | 14.75 |
| G3BP1 | Ras GTPase-a | 1.74322812 | 0.0180694 | 0.0021518 | 0.0021157 | 0.0018737 | 0.0023461 | 0.00212183 | 0.0036862 | 0.0048734 | 0.002524 | 0.0037117 | 0.00369883 | 38 | 37 | 33 | 41 | 37.25 | 51 | 67 | 28 | 51 | 49.25 |
| HS90B | Heat shock p | 1.73452681 | 0.00044384 | 0.00051027 | 0.00069929 | 0.00065781 | 0.00055246 | 0.00060496 | 0.00107 | 0.00103 | 0.0011604 | 0.00093686 | 0.00104932 | 15 | 20 | 19 | 16 | 17.5 | 23 | 23 | 20 | 21 | 21.75 |
| PLK1 | Serine/threo | 1.73147879 | 0.03004085 | 0.00017505 | 0.00022095 | 0.00026327 | 0.00013266 | 0.00019798 | 0.000391 | 0.00033727 | 0.00041797 | 0.00022497 | 0.0003428 | 4 | 5 | 6 | 3 | 4.5 | 7 | 6 | 6 | 4 | 5.75 |
| FXR1 | Fragile X mer | 1.72989622 | 0.02854189 | 0.00016997 | 0.00012873 | 0 | 0 | 0.00014935 | 0.00021695 | 0.00038208 | 0.00027057 | 0.00016384 | 0.00025836 | 4 | 4 | 0 | 0 | 4 | 5 | 7 | 4 | 3 | 4.75 |
| HAUS6 | HAUS augmir | 1.72258365 | 0.00316893 | 0.00049737 | 0.00033482 | 0.0004987 | 0.00036299 | 0.00042347 | 0.00084645 | 0.00074535 | 0.0006158 | 0.00071025 | 0.00072946 | 18 | 12 | 18 | 13 | 15.25 | 24 | 21 | 14 | 20 | 19.75 |
| PABP4 | Polyadenylat | 1.71728729 | 0.11924095 | 0.00028683 | 0.00016551 | 0.00032868 | 0.00037266 | 0.00028842 | 0.00067991 | 0.00052633 | 0.00019568 | 0.00057928 | 0.0004953 | 28 | 23 | 28 | 33 | 28 | 35 | 29 | 10 | 29 | 25.75 |
| RS13 | 40S ribosoma | 1.71452418 | 0.00724328 | 0 | 0 | 0.00035045 | 0 | 0.00035045 | 0.00044611 | 0.00044895 | 0.00083456 | 0.0006738 | 0.00060086 | 0 | 0 | 2 | 0 | 2 | 2 | 2 | 3 | 3 | 2.5 |
| EFTU | Elongation fa | 1.71416837 | 1.1622E-05 | 0.0012844 | 0.0013559 | 0.0011122 | 0.0012389 | 0.00124785 | 0.002161 | 0.0022497 | 0.0020445 | 0.0021009 | 0.00213903 | 22 | 23 | 19 | 21 | 21.25 | 29 | 30 | 22 | 28 | 27.25 |
| CNOT9 | CCR4-NOT tr | 1.71276724 | 0.05144242 | 0.00097081 | 0.0011585 | 0.00079642 | 0.00080264 | 0.00093209 | 0.0020277 | 0.0018138 | 0.00084293 | 0.0017014 | 0.00159646 | 11 | 13 | 9 | 9 | 10.5 | 18 | 16 | 6 | 15 | 13.75 |
| BCR | Breakpoint cl | 1.71029156 | 0.00796056 | 0.00062286 | 0.00058702 | 0.0006037 | 0.00065038 | 0.00061599 | 0.0011395 | 0.0011201 | 0.00072709 | 0.0012274 | 0.00105352 | 30 | 28 | 29 | 31 | 29.5 | 43 | 42 | 22 | 46 | 38.25 |
| RL36A | 60S ribosoma | 1.70441222 | 0.09345179 | 0 | 0 | 0.00049922 | 0 | 0.00049922 | 0.00095325 | 0.00063954 | 0 | 0.00095984 | 0.00085088 | 0 | 0 | 2 | 0 | 2 | 3 | 2 | 0 | 3 | 2.66666667 |

| | | | | | | | | | | | | | | | | | | | | | | | |
|---|---|---|---|---|---|---|---|---|---|---|---|---|---|---|---|---|---|---|---|---|---|---|---|
| NO40 | Nucleolar prc | 1.70325101 | 0.69963653 | 0.00043798 | 0.0003317 | 0.00043915 | 0.00022129 | 0.00035753 | 0.00027952 | 0.0011252 | 0 | 0.00042217 | 0.00060896 | 4 | 3 | 4 | 2 | 3.25 | 2 | 8 | 0 | 3 | 4.33333333 |
| AIMP2 | Aminoacyl tR | 1.70319372 | 0.02437402 | 0.00016493 | 0 | 0.00033073 | 0 | 0.00024783 | 0.00031576 | 0.00052962 | 0.00052508 | 0.00031795 | 0.0004221 | 2 | 0 | 4 | 0 | 3 | 3 | 5 | 4 | 3 | 3.75 |
| UBAC2 | Ubiquitin-ass | 1.70170972 | 0.00602932 | 0 | 0 | 0.00023074 | 0 | 0.00023074 | 0.00029373 | 0.00049267 | 0.00048844 | 0.00029577 | 0.00039265 | 0 | 0 | 3 | 0 | 3 | 3 | 5 | 4 | 3 | 3.75 |
| EIF3B | Eukaryotic tra | 1.69968011 | 0.00116029 | 0.00038902 | 0.00049103 | 0.00055258 | 0.00045862 | 0.00047281 | 0.00078618 | 0.00079117 | 0.00092888 | 0.00070829 | 0.00080363 | 12 | 15 | 17 | 14 | 14.5 | 19 | 19 | 18 | 17 | 18.25 |
| STAU1 | Double-stran | 1.6980273 | 0.00337404 | 0.0004116 | 0.00036945 | 0.00032099 | 0.00027729 | 0.00034483 | 0.00046699 | 0.00064619 | 0.00058241 | 0.00064655 | 0.00058554 | 9 | 8 | 7 | 6 | 7.5 | 9 | 12 | 9 | 12 | 10.5 |
| CTBP2 | C-terminal-bi | 1.69632413 | 0.02805052 | 0.0002372 | 0.00035928 | 0.00029729 | 0.00035954 | 0.00031333 | 0.00060551 | 0.00045702 | 0.00037758 | 0.00068591 | 0.00053151 | 4 | 6 | 5 | 6 | 5.25 | 8 | 6 | 4 | 9 | 6.75 |
| HSDL2 | Hydroxysterc | 1.69000471 | 0.04094552 | 0 | 0.00019124 | 0.00018989 | 0.00025517 | 0.0002121 | 0.00048347 | 0.00040545 | 0.00030148 | 0.0002434 | 0.00035845 | 0 | 3 | 3 | 4 | 3.33333333 | 6 | 5 | 3 | 3 | 4.25 |
| RAB1A | Ras-related p | 1.68952091 | 0.03573939 | 0.0005149 | 0.00064991 | 0.00064533 | 0.00065038 | 0.00061513 | 0.0014787 | 0.00082672 | 0.0010245 | 0.00082718 | 0.00103928 | 4 | 5 | 5 | 5 | 4.75 | 9 | 5 | 5 | 5 | 6 |
| CDR2 | Cerebellar de | 1.68648448 | 0.03467604 | 0.00046499 | 0.00035216 | 0.00052451 | 0.00041114 | 0.0004382 | 0.00051932 | 0.00067194 | 0.0010178 | 0.00074701 | 0.00073902 | 8 | 6 | 9 | 7 | 7.5 | 7 | 9 | 11 | 10 | 9.25 |
| DVL2 | Segment pola | 1.68063634 | 0.05698435 | 0.0019361 | 0.0020999 | 0.0020132 | 0.0017391 | 0.00194708 | 0.0038441 | 0.0037764 | 0.0015981 | 0.0038707 | 0.00327233 | 54 | 58 | 56 | 48 | 54 | 84 | 82 | 28 | 84 | 69.5 |
| RBGPR | Rab3 GTPase | 1.67816251 | 0.01170993 | 0.00045465 | 0.00036345 | 0.00068379 | 0.00038285 | 0.00047119 | 0.00079791 | 0.00080298 | 0.00090466 | 0.00065735 | 0.00079073 | 24 | 19 | 36 | 20 | 24.75 | 33 | 33 | 30 | 27 | 30.75 |
| TRAP1 | Heat shock p | 1.67785797 | 0.0034726 | 0.00063722 | 0.0006056 | 0.00041342 | 0.00053028 | 0.00054663 | 0.00090902 | 0.00077035 | 0.001074 | 0.0009153 | 0.00091717 | 17 | 16 | 11 | 14 | 14.5 | 19 | 16 | 18 | 19 | 18 |
| P53 | Cellular tumc | 1.67677778 | 0.00430586 | 0.0023501 | 0.0023053 | 0.0025583 | 0.0027819 | 0.0024989 | 0.003771 | 0.0035362 | 0.0042754 | 0.0051778 | 0.0041901 | 35 | 34 | 38 | 41 | 37 | 44 | 41 | 40 | 60 | 46.25 |
| DESP | Desmoplakin | 1.67626846 | 0.00047054 | 0.00030332 | 0.00038981 | 0.00034099 | 0.00037152 | 0.00035141 | 0.00057485 | 0.0005667 | 0.00054135 | 0.00067333 | 0.00058906 | 33 | 42 | 37 | 40 | 38 | 49 | 48 | 37 | 57 | 47.75 |
| COPG1 | Coatomer sul | 1.67557589 | 0.03977518 | 0.00021135 | 0.00021342 | 0.00039355 | 0.00021357 | 0.00025797 | 0.00034684 | 0.00034904 | 0.00052868 | 0.00050445 | 0.00043225 | 7 | 7 | 13 | 7 | 8.5 | 9 | 9 | 11 | 13 | 10.5 |
| GARS | Glycine--tRN/ | 1.66828857 | 0.05905347 | 0.00017854 | 0.00039663 | 0.00014321 | 0.00028867 | 0.00025176 | 0.00036462 | 0.0005504 | 0.00039789 | 0.00036714 | 0.00042001 | 5 | 11 | 4 | 8 | 7 | 8 | 12 | 7 | 8 | 8.75 |
| ENOA | Alpha-enolas | 1.65527219 | 0.0186256 | 0.00036482 | 0.00049118 | 0.00054868 | 0.00067586 | 0.00052014 | 0.00077607 | 0.0010934 | 0.0008711 | 0.00070329 | 0.00086097 | 6 | 8 | 9 | 11 | 8.5 | 10 | 14 | 9 | 9 | 10.5 |
| XPO5 | Exportin-5 O! | 1.65174581 | 0.60500987 | 4.3835E-05 | 6.6395E-05 | 6.5927E-05 | 4.4295E-05 | 5.5113E-05 | 8.3924E-05 | 0 | 0.00010467 | 8.4504E-05 | 9.1033E-05 | 2 | 3 | 3 | 2 | 2.5 | 3 | 0 | 3 | 3 | 3 |
| BICD2 | Protein bicau | 1.64573594 | 0.00200257 | 0.0015052 | 0.0017139 | 0.0016055 | 0.001165 | 0.0014974 | 0.0020847 | 0.0025915 | 0.0027528 | 0.0024283 | 0.00246433 | 47 | 53 | 50 | 36 | 46.5 | 51 | 63 | 54 | 59 | 56.75 |
| MA7D3 | MAP7 domai | 1.64259627 | 0.15952484 | 0.00045186 | 0.00027377 | 0.00045306 | 0.00039572 | 0.0003936 | 0.00080744 | 0.00081257 | 0.00019181 | 0.0007743 | 0.00064653 | 15 | 9 | 15 | 13 | 13 | 21 | 21 | 4 | 20 | 16.5 |
| COQ8A | Atypical kina: | 1.63881441 | 0.02946267 | 0.00012236 | 0.00020592 | 8.1789E-05 | 0.00012364 | 0.00013343 | 0.00020823 | 0.00020956 | 0.00019477 | 0.00026209 | 0.00021866 | 3 | 5 | 2 | 3 | 3.25 | 4 | 4 | 3 | 5 | 4 |
| RS30 | 40S ribosoma | 1.63501286 | 0.01073396 | 0 | 0 | 0.0017938 | 0.0013559 | 0.00157485 | 0.0028544 | 0.002298 | 0.0028479 | 0.0022993 | 0.0025749 | 0 | 0 | 4 | 3 | 3.5 | 5 | 4 | 4 | 4 | 4.25 |
| KBP | KIF1-binding | 1.63384091 | 0.01086285 | 0 | 0.00017164 | 0.00012782 | 0 | 0.00014973 | 0.00027119 | 0.00021833 | 0.00027057 | 0.00021845 | 0.00024464 | 0 | 4 | 3 | 0 | 3.5 | 5 | 4 | 4 | 4 | 4.25 |
| FAF2 | FAS-associate | 1.63267751 | 0.47401849 | 0.0001186 | 0 | 0.00011892 | 0.00017977 | 0.0001391 | 0.00015138 | 0.00015234 | 0.00037758 | 0 | 0.0002271 | 2 | 0 | 2 | 3 | 2.33333333 | 2 | 2 | 4 | 0 | 2.66666667 |
| RPTOR | Regulatory-a: | 1.62876928 | 0.01146441 | 3.9533E-05 | 0 | 5.9458E-05 | 0 | 4.9496E-05 | 7.5689E-05 | 0.00007617 | 9.4396E-05 | 7.6212E-05 | 8.0617E-05 | 2 | 0 | 3 | 0 | 2.5 | 3 | 3 | 3 | 3 | 3 |
| TCPD | T-complex pr | 1.62544477 | 0.00034284 | 0.00073437 | 0.00079099 | 0.00073633 | 0.00084103 | 0.00077568 | 0.0011873 | 0.0013206 | 0.0014028 | 0.0011326 | 0.00126083 | 20 | 20 | 20 | 22 | 20.5 | 20 | 24 | 20 | 19 | 20.75 |
| TCPG | T-complex pr | 1.62113909 | 0.00102994 | 0.0011136 | 0.0010756 | 0.0011652 | 0.0012721 | 0.00115663 | 0.0020394 | 0.001928 | 0.0015415 | 0.0019913 | 0.00187505 | 23 | 22 | 24 | 26 | 23.75 | 33 | 31 | 20 | 32 | 29 |
| ECD | Protein ecdy: | 1.61727312 | 0.03761172 | 0 | 0.00012413 | 0.00008217 | 0.00020703 | 0.00013778 | 0.0002092 | 0.00021053 | 0.00026091 | 0.00021065 | 0.00022282 | 0 | 3 | 2 | 5 | 3.33333333 | 4 | 4 | 4 | 4 | 4 |
| PP6R3 | Serine/threo | 1.61281274 | 0.03670476 | 0.00027205 | 0.00042732 | 0.00030308 | 0.00058035 | 0.0003957 | 0.00054014 | 0.00069888 | 0.00076987 | 0.00054387 | 0.00063819 | 9 | 14 | 10 | 19 | 13 | 14 | 18 | 17 | 15 | 16 |
| RL9 | 60S ribosoma | 1.60989754 | 0.00058254 | 0.0043981 | 0.005135 | 0.0055122 | 0.0038887 | 0.0047335 | 0.0071924 | 0.0079443 | 0.0072198 | 0.0081253 | 0.00762045 | 32 | 37 | 40 | 28 | 34.25 | 41 | 45 | 33 | 46 | 41.25 |
| MSH3 | DNA mismatc | 1.603913 | 0.68828465 | 4.6418E-05 | 7.0307E-05 | 4.6541E-05 | 4.6905E-05 | 5.2543E-05 | 0 | 5.9623E-05 | 7.3889E-05 | 0.00011931 | 8.4274E-05 | 2 | 3 | 2 | 2 | 2.25 | 0 | 2 | 2 | 4 | 2.66666667 |
| PUR2 | Trifunctional | 1.60347753 | 0.00210388 | 0.00054867 | 0.00047489 | 0.00034056 | 0.00052803 | 0.00047304 | 0.00080035 | 0.00077188 | 0.00079022 | 0.00067157 | 0.00075851 | 21 | 18 | 13 | 20 | 18 | 24 | 23 | 19 | 20 | 21.5 |
| RS23 | 40S ribosoma | 1.59699483 | 0.01826738 | 0.0059051 | 0.0063355 | 0.0055508 | 0.006713 | 0.0061261 | 0.0077727 | 0.010429 | 0.012631 | 0.0083007 | 0.00978335 | 32 | 34 | 30 | 36 | 33 | 33 | 44 | 43 | 35 | 38.75 |
| TECR | Very-long-cha | 1.59621343 | 0.04021442 | 0.00025703 | 0.00069212 | 0 | 0.00051946 | 0.00048954 | 0.00065613 | 0.00088041 | 0.0008183 | 0.00077078 | 0.00078141 | 3 | 8 | 0 | 6 | 5.66666667 | 6 | 8 | 6 | 7 | 6.75 |
| RS8 | 40S ribosoma | 1.59549852 | 3.539E-05 | 0.0021567 | 0.0020497 | 0.0021625 | 0.0021794 | 0.00213708 | 0.0037244 | 0.0034222 | 0.0032312 | 0.003261 | 0.0034097 | 17 | 16 | 17 | 17 | 16.75 | 23 | 21 | 16 | 20 | 20 |
| ZCHC3 | Zinc finger CC | 1.58918301 | 0.22172485 | 0.00013064 | 0.00013191 | 0 | 0 | 0.00013128 | 0.00025011 | 0.0001678 | 0.00020795 | 0 | 0.00020862 | 2 | 2 | 0 | 0 | 2 | 3 | 2 | 2 | 0 | 2.33333333 |
| DHB12 | Very-long-cha | 1.58757069 | 0.06529184 | 0.00033831 | 0.00076865 | 0.00059362 | 0.00085467 | 0.00063881 | 0.00097158 | 0.00097776 | 0.0013464 | 0.0007609 | 0.00101416 | 4 | 9 | 7 | 10 | 7.5 | 9 | 9 | 10 | 7 | 8.75 |
| TXLNA | Alpha-taxilin | 1.58723757 | 0.05781056 | 0.00024165 | 9.7606E-05 | 0.00019384 | 0 | 0.0001777 | 0.00024675 | 0.00024832 | 0.00038467 | 0.00024846 | 0.00028205 | 5 | 2 | 4 | 0 | 3.66666667 | 4 | 4 | 5 | 4 | 4.25 |
| DCP1A | mRNA-decap | 1.58367923 | 0.70331924 | 0.00031739 | 0.00032049 | 0.00031823 | 0.00036654 | 0.00033066 | 0.00028936 | 0.00064064 | 0 | 0.00064099 | 0.00052366 | 7 | 7 | 7 | 8 | 7.25 | 5 | 11 | 0 | 11 | 9 |
| TBB4B | Tubulin beta- | 1.58118082 | 0.00072464 | 0.0050405 | 0.0054491 | 0.0055296 | 0.0049137 | 0.00523323 | 0.0082501 | 0.0073885 | 0.009534 | 0.0079261 | 0.00827468 | 85 | 91 | 93 | 82 | 87.75 | 109 | 97 | 101 | 104 | 102.75 |
| SYRC | Arginine--tRN | 1.5787047 | 0.00728901 | 0.00019991 | 0.00032299 | 0.00032071 | 0.00032322 | 0.00029171 | 0.00040826 | 0.00051357 | 0.00050917 | 0.00041108 | 0.00046052 | 5 | 8 | 8 | 8 | 7.25 | 8 | 10 | 8 | 8 | 8.5 |
| LDHA | L-lactate deh | 1.57409724 | 0.12985988 | 0.00055638 | 0.0004013 | 0.00015939 | 0.00048191 | 0.00039975 | 0.0004058 | 0.00061257 | 0.00088567 | 0.00061291 | 0.00062924 | 7 | 5 | 2 | 6 | 5 | 4 | 6 | 7 | 6 | 5.75 |
| RS27A | Ubiquitin-40S | 1.57025255 | 0.00242927 | 0.0028757 | 0.0020497 | 0.0027137 | 0.0027349 | 0.0025935 | 0.0036704 | 0.0036938 | 0.0045776 | 0.004348 | 0.00407245 | 17 | 12 | 16 | 16 | 15.25 | 17 | 17 | 17 | 20 | 17.75 |
| SKIV2 | Helicase SKI2 | 1.57023644 | 0.06047334 | 4.2357E-05 | 4.2771E-05 | 6.3705E-05 | 0 | 4.9611E-05 | 0.00010813 | 8.1611E-05 | 6.7426E-05 | 5.4437E-05 | 7.7901E-05 | 2 | 2 | 3 | 0 | 2.33333333 | 4 | 3 | 2 | 2 | 2.75 |
| PDLI5 | PDZ and LIM | 1.56744209 | 0.11775991 | 0.00013283 | 0 | 0 | 0 | 0.00013283 | 0.00016954 | 0.00028436 | 0 | 0.00017071 | 0.0002082 | 3 | 0 | 0 | 0 | 3 | 3 | 5 | 0 | 3 | 3.66666667 |
| UBC9 | SUMO-conju | 1.56026944 | 0.20161044 | 0.0011691 | 0.0011805 | 0.0015071 | 0.0010126 | 0.00121733 | 0.0023449 | 0.0025743 | 0.00053172 | 0.0021465 | 0.00189936 | 7 | 7 | 9 | 6 | 7.25 | 11 | 12 | 2 | 10 | 8.75 |
| DVL3 | Segment pola | 1.55961447 | 0.0712678 | 0.0011425 | 0.0013398 | 0.0011825 | 0.0011545 | 0.00120483 | 0.0023991 | 0.002083 | 0.00099735 | 0.0020368 | 0.00187906 | 38 | 45 | 39 | 41 | 40.75 | 66 | 59 | 27 | 59 | 52.75 |
| SMD1 | Small nuclear | 1.55816595 | 0.27097128 | 0.00066525 | 0 | 0.00066703 | 0 | 0.00066614 | 0.0011322 | 0.00056967 | 0.001412 | 0 | 0.00103796 | 3 | 0 | 3 | 0 | 3 | 4 | 2 | 4 | 0 | 3.33333333 |
| ATPA | ATP synthase | 1.54697894 | 0.01360627 | 0.0032449 | 0.0025538 | 0.0035884 | 0.0022181 | 0.0029013 | 0.0043244 | 0.0051487 | 0.0048615 | 0.0036184 | 0.00448825 | 68 | 53 | 75 | 46 | 60.5 | 71 | 84 | 64 | 59 | 69.5 |
| DYST | Dystonin OS= | 1.53815231 | 0.1129518 | 6.9718E-06 | 0 | 6.9904E-06 | 2.4658E-05 | 1.2873E-05 | 2.2247E-05 | 0.00001791 | 1.6647E-05 | 2.2401E-05 | 1.9801E-05 | 2 | 0 | 2 | 7 | 3.66666667 | 5 | 4 | 3 | 5 | 4.25 |
| SRS11 | Serine/argini | 1.5360596 | 0.73726532 | 0.00016356 | 0.00027527 | 0.000328 | 0.00038566 | 0.00028812 | 0.00048713 | 0.00049023 | 0 | 0.00035036 | 0.00044257 | 3 | 5 | 6 | 7 | 5.25 | 7 | 7 | 0 | 5 | 6.33333333 |
| GAPD1 | GTPase-activ | 1.53505706 | 0.13833324 | 0.00010712 | 0.00010817 | 0.00012531 | 0.00016238 | 0.00012575 | 0.0002051 | 0.00022933 | 8.5263E-05 | 0.00025241 | 0.00019303 | 6 | 6 | 7 | 9 | 7 | 9 | 10 | 3 | 11 | 8.25 |
| LARP1 | La-related pr | 1.53488658 | 0.15294394 | 0.00072231 | 0.000778 | 0.00055525 | 0.00063258 | 0.00067204 | 0.0011678 | 0.0014226 | 0.00042159 | 0.001114 | 0.0010315 | 30 | 32 | 23 | 26 | 27.75 | 38 | 46 | 11 | 36 | 32.75 |
| P2R3A | Serine/threo | 1.53429965 | 0.50446634 | 6.8839E-05 | 4.6342E-05 | 0.00011504 | 0 | 7.674E-05 | 8.7865E-05 | 8.8423E-05 | 0 | 0.00017694 | 0.00011774 | 3 | 2 | 5 | 0 | 3.33333333 | 3 | 3 | 0 | 6 | 4 |
| BCCIP | BRCA2 and C | 1.5332589 | 0.0459475 | 0.00067232 | 0.00093348 | 0.00058984 | 0.00067938 | 0.00071876 | 0.0013945 | 0.0011874 | 0.0010702 | 0.00075605 | 0.00110204 | 8 | 11 | 7 | 8 | 8.5 | 13 | 11 | 8 | 7 | 9.75 |
| RAPH1 | Ras-associate | 1.53192604 | 0.44452843 | 8.4443E-05 | 6.3952E-05 | 6.3501E-05 | 0 | 7.0632E-05 | 0.00013473 | 8.1349E-05 | 0 | 0.00010853 | 0.0001082 | 4 | 3 | 3 | 0 | 3.33333333 | 5 | 3 | 0 | 4 | 4 |
| ROA2 | Heterogeneo | 1.53121924 | 0.78259539 | 0.00097181 | 0.0011323 | 0.00082449 | 0.00083094 | 0.00093989 | 0.00066791 | 0.0022085 | 0 | 0.0014411 | 0.00143917 | 13 | 15 | 11 | 11 | 12.5 | 7 | 23 | 0 | 15 | 15 |
| CE170 | Centrosomal | 1.52190674 | 0.24278785 | 0.00028321 | 0.00033645 | 0.00033407 | 0.00025252 | 0.00030156 | 0.00061665 | 0.00051357 | 0.00010608 | 0.0005995 | 0.00045895 | 17 | 20 | 20 | 15 | 18 | 29 | 24 | 4 | 28 | 21.25 |
| SRC8 | Src substrate | 1.52177307 | 0.05930245 | 0.0009116 | 0.0011143 | 0.00048107 | 0.00082421 | 0.0008328 | 0.0010411 | 0.0013558 | 0.0010692 | 0.0016032 | 0.00126733 | 19 | 23 | 10 | 17 | 17.25 | 17 | 22 | 14 | 26 | 19.75 |
| MCM6 | DNA replicati | 1.52117976 | 0.07455306 | 0.00028928 | 0.00045439 | 0.00029005 | 0.00029231 | 0.00033151 | 0.00041025 | 0.00070186 | 0.00040932 | 0.0004957 | 0.00050428 | 9 | 14 | 9 | 9 | 10.25 | 10 | 17 | 8 | 12 | 11.75 |
| RBP2 | E3 SUMO-prc | 1.51303749 | 0.21651546 | 0.0014487 | 0.0015456 | 0.0015675 | 0.0013895 | 0.00148783 | 0.0027789 | 0.0028702 | 0.00059934 | 0.0027561 | 0.00225114 | 177 | 187 | 191 | 168 | 180.75 | 266 | 273 | 46 | 262 | 211.75 |
| PYM1 | Partner of Y1 | 1.50991515 | 0.04117639 | 0.00025871 | 0.00026124 | 0 | 0.00026143 | 0.00026046 | 0.00033021 | 0.00033231 | 0.00041183 | 0.00049874 | 0.00039327 | 2 | 2 | 0 | 2 | 2 | 2 | 2 | 2 | 3 | 2.25 |
| S61A1 | Protein trans | 1.50922771 | 0.20860533 | 0.00027719 | 0.0002799 | 0.00027793 | 0.0002801 | 0.00027878 | 0.00049532 | 0.00014242 | 0.00061774 | 0.00042749 | 0.00042074 | 5 | 5 | 5 | 5 | 5 | 7 | 2 | 7 | 6 | 5.5 |
| RIOK1 | Serine/threo | 1.50524194 | 0.09608844 | 0.00018583 | 0.00023456 | 0.00023291 | 0.00023473 | 0.00022201 | 0.00035579 | 0.00035805 | 0.00044373 | 0.00017913 | 0.00033418 | 4 | 5 | 5 | 5 | 4.75 | 6 | 6 | 6 | 3 | 5.25 |
| RS3A | 40S ribosoma | 1.50486982 | 0.15356115 | 0.00089961 | 0.0012112 | 0.000902 | 0.00070704 | 0.00092996 | 0.00076549 | 0.0019259 | 0.0017503 | 0.0011562 | 0.00139947 | 9 | 12 | 9 | 7 | 9.25 | 6 | 15 | 11 | 9 | 10.25 |
| CBPC1 | Cytosolic carl | 1.5036565 | 0.00431416 | 0.00064572 | 0.00084765 | 0.00071218 | 0.00067425 | 0.00071995 | 0.0010714 | 0.0010506 | 0.0012677 | 0.00094053 | 0.00108256 | 30 | 39 | 33 | 31 | 33.25 | 39 | 38 | 37 | 34 | 37 |
| KCC2D | Calcium/calm | 1.49415304 | 0.02940214 | 0 | 0 | 0.00015907 | 0.00010688 | 0.00013298 | 0.00020249 | 0.00020378 | 0.00025254 | 0.00013593 | 0.00019869 | 0 | 0 | 3 | 2 | 2.5 | 3 | 3 | 3 | 2 | 2.75 |
| RL21 | 60S ribosoma | 1.4876485 | 0.0691866 | 0.00065971 | 0 | 0.00082683 | 0.00066664 | 0.00071773 | 0.0012631 | 0.0010592 | 0.0013127 | 0.0006359 | 0.00106773 | 4 | 0 | 5 | 4 | 4.33333333 | 6 | 5 | 5 | 3 | 4.75 |
| IF5 | Eukaryotic tra | 1.48606267 | 0.79758475 | 0.00018368 | 0.00030912 | 0.00018417 | 0.00030935 | 0.00024658 | 0.00031259 | 0.00031458 | 0 | 0.00047213 | 0.00036643 | 3 | 5 | 3 | 5 | 4 | 4 | 4 | 0 | 6 | 4.66666667 |
| EF1G | Elongation fa | 1.48571954 | 0.02327339 | 0.00072462 | 0.00073171 | 0.00084765 | 0.00079326 | 0.00077431 | 0.00092489 | 0.0013962 | 0.00096124 | 0.0013193 | 0.00115041 | 12 | 12 | 14 | 13 | 12.75 | 12 | 18 | 10 | 17 | 14.25 |
| WASC5 | WASH compl | 1.47978921 | 0.15110691 | 0.00091073 | 0.00085066 | 0.0012328 | 0.00075925 | 0.00093836 | 0.0010171 | 0.0011991 | 0.0021384 | 0.0011997 | 0.00138858 | 40 | 37 | 54 | 33 | 41 | 35 | 41 | 59 | 41 | 44 |
| RANG | Ran-specific ( | 1.47514292 | 0.00835536 | 0.00039386 | 0.00026514 | 0.00039491 | 0.00039799 | 0.00036298 | 0.00050271 | 0.0005059 | 0.00062696 | 0.00050619 | 0.00053544 | 3 | 2 | 3 | 3 | 2.75 | 3 | 3 | 3 | 3 | 3 |
| KPYM | Pyruvate kina | 1.47504016 | 0.00086224 | 0.0030314 | 0.0035629 | 0.0036374 | 0.0032139 | 0.0033614 | 0.0051379 | 0.004979 | 0.0043509 | 0.005365 | 0.0049582 | 61 | 71 | 73 | 64 | 67.25 | 81 | 78 | 55 | 84 | 74.5 |
| SYVC | Valine--tRNA | 1.47475967 | 0.06769444 | 0.00018789 | 0.00023189 | 0.00018839 | 0.00025315 | 0.00021533 | 0.00031976 | 0.00034861 | 0.0001994 | 0.00040247 | 0.00031756 | 9 | 11 | 9 | 12 | 10.25 | 12 | 13 | 6 | 15 | 11.5 |
| TBB4A | Tubulin beta- | 1.46203462 | 0.0411796 | 0 | 0.00018004 | 0.00011918 | 0 | 0.00014961 | 0.00022758 | 0.00030537 | 0.00018922 | 0.00015277 | 0.00021874 | 0 | 87 | 84 | 0 | 85.5 | 103 | 93 | 93 | 97 | 96.5 |
| CAMP1 | Calmodulin-r | 1.46130733 | 0.30240723 | 0.00019767 | 0.0002994 | 0.00016516 | 0.00023303 | 0.00022382 | 0.00044152 | 0.00048664 | 0.00010488 | 0.00027521 | 0.00032706 | 12 | 18 | 10 | 14 | 13.5 | 21 | 23 | 4 | 13 | 15.25 |
| KIF11 | Kinesin-like p | 1.46113577 | 0.00181709 | 0.00037484 | 0.0005299 | 0.00047606 | 0.00047978 | 0.00046515 | 0.00073359 | 0.00064196 | 0.00063646 | 0.00070655 | 0.00067964 | 15 | 21 | 19 | 19 | 18.5 | 23 | 20 | 16 | 22 | 20.25 |
| XPOT | Exportin-T O! | 1.46047753 | 0.02802595 | 0.00016458 | 0.00019389 | 0.00013752 | 0.00011088 | 0.00015172 | 0.00024508 | 0.00024664 | 0.00021833 | 0.00017627 | 0.00022158 | 6 | 7 | 5 | 4 | 5.5 | 7 | 7 | 5 | 5 | 6 |
| RS2 | 40S ribosoma | 1.45976844 | 0.00027431 | 0.0017112 | 0.001546 | 0.0014448 | 0.0014561 | 0.00153953 | 0.002414 | 0.002198 | 0.0022939 | 0.0020835 | 0.00224735 | 19 | 17 | 16 | 16 | 17 | 21 | 19 | 16 | 18 | 18.5 |
| GNL3 | Guanine nucl | 1.4574607 | 0.0327494 | 0.000721 | 0.00092219 | 0.0008193 | 0.00092285 | 0.00084634 | 0.0011657 | 0.0011731 | 0.0016068 | 0.0009884 | 0.0012335 | 15 | 19 | 17 | 19 | 17.5 | 19 | 19 | 21 | 16 | 18.75 |
| AL1B1 | Aldehyde del | 1.45645248 | 0.83076318 | 0.00025521 | 0.00030924 | 0.00020471 | 0.00030947 | 0.00026966 | 0.00032574 | 0.0005245 | 0 | 0.00032799 | 0.00039274 | 5 | 6 | 4 | 6 | 5.25 | 5 | 8 | 0 | 5 | 6 |
| CHM4B | Charged mult | 1.45480151 | 0.06765879 | 0.00035342 | 0.00035687 | 0.0005906 | 0.00035713 | 0.00041451 | 0.00045109 | 0.00060528 | 0.00075011 | 0.00060561 | 0.00060302 | 3 | 3 | 5 | 3 | 3.5 | 3 | 4 | 4 | 4 | 3.75 |
| TNPO3 | Transportin-3 | 1.45053821 | 0.60809245 | 0.00008577 | 5.7739E-05 | 0.00011466 | 0 | 8.6056E-05 | 0 | 7.3447E-05 | 0.00022755 | 7.3487E-05 | 0.00012483 | 3 | 2 | 4 | 0 | 3 | 0 | 2 | 5 | 2 | 3 |
| FHL1 | Four and a ha | 1.44933127 | 0.07310757 | 0.00040849 | 0.00065997 | 0.00065532 | 0.00049534 | 0.00055478 | 0.00052139 | 0.00083952 | 0.00091035 | 0.00094498 | 0.00080406 | 5 | 8 | 8 | 6 | 6.75 | 5 | 8 | 7 | 9 | 7.25 |
| TAGL2 | Transgelin-2 | 1.4489632 | 0.04773472 | 0.0014587 | 0.0014729 | 0.00093071 | 0.001876 | 0.00143458 | 0.0016925 | 0.002044 | 0.002533 | 0.0020451 | 0.00207865 | 11 | 11 | 7 | 14 | 10.75 | 10 | 12 | 12 | 12 | 11.5 |
| IPO11 | Importin-11 ( | 1.44507329 | 0.89127708 | 0.00005413 | 0.00010932 | 0.00010855 | 5.4699E-05 | 8.1675E-05 | 0.00006909 | 0 | 0.00021542 | 6.9568E-05 | 0.00011803 | 2 | 4 | 4 | 2 | 3 | 2 | 0 | 5 | 2 | 3 |
| DVL1 | Segment pola | 1.4402062 | 0.18492022 | 0.00041766 | 0.00065178 | 0.00060912 | 0.00057552 | 0.00056352 | 0.00092079 | 0.0011705 | 0.00042308 | 0.00073197 | 0.00081159 | 11 | 19 | 17 | 15 | 15.5 | 21 | 25 | 7 | 18 | 17.75 |
| TCPA | T-complex pr | 1.43785948 | 0.00265973 | 0.0029426 | 0.0035465 | 0.0028553 | 0.0030694 | 0.00310345 | 0.0044222 | 0.0046332 | 0.0038531 | 0.0049408 | 0.00446233 | 62 | 74 | 60 | 64 | 65 | 73 | 76 | 51 | 81 | 70.25 |
| PPM1F | Protein phos | 1.43667188 | 0.52735414 | 0.00011625 | 0.00017608 | 0.00017484 | 0 | 0.00015572 | 0.00014838 | 0.00022398 | 0 | 0.00029881 | 0.00022372 | 2 | 3 | 3 | 0 | 2.66666667 | 2 | 3 | 0 | 4 | 3 |
| RL8 | 60S ribosoma | 1.43520683 | 0.00692166 | 0.00092411 | 0.00093315 | 0.0011325 | 0.0012451 | 0.00105872 | 0.0015727 | 0.0017146 | 0.001471 | 0.0013196 | 0.00151948 | 9 | 9 | 11 | 12 | 10.25 | 12 | 13 | 9 | 10 | 11 |
| KAPCA | cAMP-depen | 1.42583286 | 0.03085209 | 0 | 0.00022775 | 0 | 0.00022791 | 0.00022783 | 0.00038384 | 0.00028971 | 0.00023935 | 0.00038649 | 0.00032485 | 0 | 3 | 0 | 3 | 3 | 4 | 3 | 2 | 4 | 3.25 |
| ECHD1 | Ethylmalonyl | 1.42500452 | 0.03090448 | 0 | 0.00034719 | 0 | 0.00026058 | 0.00030389 | 0.00054856 | 0.00033123 | 0.00041048 | 0.00044188 | 0.00043304 | 0 | 4 | 0 | 3 | 3.5 | 5 | 3 | 3 | 4 | 3.75 |
| P5CS | Delta-1-pyrrc | 1.42239013 | 0.01806072 | 0.0011286 | 0.0010055 | 0.0010983 | 0.00093917 | 0.00104289 | 0.0014828 | 0.0012791 | 0.0018493 | 0.0013224 | 0.0014834 | 34 | 30 | 33 | 28 | 31.25 | 35 | 30 | 35 | 31 | 32.75 |
| CTNB1 | Catenin beta- | 1.41870539 | 0.5600988 | 0.00010136 | 0 | 0.00010163 | 0.00010243 | 0.00010181 | 0.00012938 | 8.6801E-05 | 0 | 0.00021712 | 0.00014443 | 4 | 0 | 4 | 4 | 4 | 3 | 2 | 0 | 5 | 3.33333333 |
| RAB5C | Ras-related p | 1.41410245 | 0.02047592 | 0.00061084 | 0.00049345 | 0.00048998 | 0.00061726 | 0.00055288 | 0.00077967 | 0.0006277 | 0.00077789 | 0.00094207 | 0.00078183 | 5 | 4 | 4 | 5 | 4.5 | 5 | 4 | 4 | 6 | 4.75 |
| HUWE1 | E3 ubiquitin- | 1.41336753 | 0.16933242 | 0 | 0.00003046 | 4.2344E-05 | 1.2193E-05 | 2.8332E-05 | 6.1603E-05 | 3.8747E-05 | 2.8811E-05 | 3.1015E-05 | 4.0044E-05 | 0 | 5 | 7 | 2 | 4.66666667 | 8 | 5 | 3 | 4 | 5 |
| DBNL | Drebrin-like p | 1.40994882 | 0.16543472 | 0.00042958 | 0.00049575 | 0.00012306 | 0.00037208 | 0.00035512 | 0.00039165 | 0.00055179 | 0.00058613 | 0.00047322 | 0.0005007 | 7 | 8 | 2 | 6 | 5.75 | 5 | 7 | 6 | 6 | 6 |
| WDR75 | WD repeat-c | 1.40653149 | 0.13600479 | 0.00038152 | 0.00022473 | 0.00019127 | 0.00019276 | 0.00024757 | 0.00032464 | 0.0003267 | 0.00045549 | 0.00028603 | 0.00034822 | 12 | 7 | 6 | 6 | 7.75 | 8 | 8 | 9 | 7 | 8 |
| IRAK1 | Interleukin-1 | 1.40636803 | 0.88191759 | 0.00025944 | 0.0002994 | 0.00026013 | 0.00026216 | 0.00027028 | 0.00037844 | 0.00038085 | 0 | 0.00038106 | 0.00038012 | 7 | 8 | 7 | 7 | 7.25 | 8 | 8 | 0 | 8 | 8 |
| FLNA | Filamin-A OS | 1.40521437 | 0.00052934 | 0.0052139 | 0.0048018 | 0.0052578 | 0.0045333 | 0.0049517 | 0.0066803 | 0.0068892 | 0.0076649 | 0.0065984 | 0.0069582 | 523 | 477 | 526 | 450 | 494 | 525 | 538 | 483 | 515 | 515.25 |
| LGUL | Lactoylglutat | 1.39675272 | 0.91133226 | 0.00057366 | 0.00086891 | 0.00028759 | 0.00072461 | 0.00061369 | 0 | 0.00092108 | 0.00091318 | 0.00073727 | 0.00085718 | 4 | 6 | 2 | 5 | 4.25 | 0 | 5 | 4 | 4 | 4.33333333 |
| UBR5 | E3 ubiquitin- | 1.3943474 | 0.59772895 | 2.8283E-05 | 0 | 4.7265E-05 | 2.8581E-05 | 3.471E-05 | 2.4067E-05 | 7.2659E-05 | 0 | 4.8466E-05 | 4.8397E-05 | 3 | 0 | 5 | 3 | 3.66666667 | 2 | 6 | 0 | 4 | 4 |
| DX39A | ATP-depende | 1.39102979 | 0.15483135 | 0.00080339 | 0.00068644 | 0.00061964 | 0.00056204 | 0.00066788 | 0.001341 | 0.00063505 | 0.000787 | 0.0009531 | 0.00092904 | 13 | 11 | 10 | 9 | 10.75 | 17 | 8 | 8 | 12 | 11.25 |
| FA83H | Protein FAM8 | 1.39076414 | 0.35604703 | 0.00017906 | 0.00024861 | 0.00024686 | 0.00018094 | 0.00021387 | 0.00045709 | 0.00034499 | 7.1257E-05 | 0.00031642 | 0.00029744 | 8 | 11 | 11 | 8 | 9.5 | 16 | 12 | 2 | 11 | 10.25 |
| STT3A | Dolichyl-diph | 1.38912081 | 0.39261428 | 0.00018715 | 7.5593E-05 | 0.00007506 | 7.5647E-05 | 0.00010336 | 9.5551E-05 | 0.00014424 | 0.00023833 | 9.6211E-05 | 0.00014358 | 5 | 2 | 2 | 2 | 2.75 | 2 | 3 | 4 | 2 | 2.75 |
| TPPC5 | Trafficking pr | 1.38427889 | 0.9330341 | 0.00042109 | 0.00056695 | 0.00056295 | 0.00056735 | 0.00052959 | 0 | 0.00054089 | 0.0011172 | 0.00054119 | 0.00073309 | 3 | 4 | 4 | 4 | 3.75 | 0 | 3 | 5 | 3 | 3.66666667 |
| CDK5 | Cyclin-depen | 1.38241795 | 0.28391067 | 0.00018074 | 0 | 0.00018122 | 0 | 0.00018098 | 0.0002307 | 0.00023216 | 0.00028771 | 0 | 0.00025019 | 3 | 0 | 3 | 0 | 3 | 2 | 3 | 2 | 0 | 2.33333333 |
| MAGD1 | Melanoma-a: | 1.37849267 | 0.13996387 | 0.00016959 | 0.0002055 | 0.00017004 | 0.00017137 | 0.00017913 | 0.00034634 | 0.00026141 | 0.00016198 | 0.00021796 | 0.00024692 | 5 | 6 | 5 | 5 | 5.25 | 8 | 6 | 3 | 5 | 5.5 |
| IMB1 | Importin sub | 1.37540848 | 0.21859563 | 0.0012953 | 0.0013688 | 0.00148 | 0.0012785 | 0.00135565 | 0.0021532 | 0.0019734 | 0.00081519 | 0.0025165 | 0.00186457 | 43 | 45 | 49 | 42 | 44.75 | 56 | 51 | 17 | 65 | 47.25 |
| COPB | Coatomer sul | 1.37451532 | 0.02921128 | 0.00033228 | 0.00036349 | 0.00033316 | 0.00036375 | 0.00034817 | 0.00056548 | 0.00049794 | 0.00035262 | 0.00049822 | 0.00047857 | 12 | 13 | 12 | 13 | 12.5 | 16 | 14 | 8 | 14 | 13 |
| RIC8A | Synembryn-A | 1.37035193 | 0.28904449 | 0 | 0.00010036 | 0 | 0.00010044 | 0.0001004 | 0.00012686 | 0.00012767 | 0.00015822 | 0 | 0.00013758 | 0 | 2 | 0 | 2 | 2 | 2 | 2 | 2 | 0 | 2 |
| LAT1 | Large neutral | 1.36860421 | 0.17984618 | 0.00015614 | 0.00010511 | 0.00015656 | 0.00010519 | 0.00013075 | 0.00013287 | 0.00020057 | 0.00024856 | 0.00013378 | 0.00017895 | 3 | 2 | 3 | 2 | 2.5 | 2 | 3 | 3 | 2 | 2.5 |
| RL18 | 60S ribosoma | 1.36682921 | 0.23914511 | 0.0022458 | 0.0018426 | 0.0014074 | 0.0017021 | 0.00179948 | 0.0019707 | 0.0037862 | 0.0024578 | 0.0016236 | 0.00245958 | 16 | 13 | 10 | 12 | 12.75 | 11 | 21 | 11 | 9 | 13 |
| RP25L | Ribonuclease | 1.36560825 | 0.30727748 | 0.00048568 | 0 | 0.00032465 | 0 | 0.00040517 | 0.00061991 | 0.00062385 | 0 | 0.00041613 | 0.0005533 | 3 | 0 | 2 | 0 | 2.5 | 3 | 3 | 0 | 2 | 2.66666667 |
| AAAT | Neutral amin | 1.35823285 | 0.96463118 | 9.7554E-05 | 0.00024627 | 0.00019563 | 0.00019716 | 0.00018415 | 0.00024903 | 0.00031327 | 0 | 0.00018807 | 0.00025012 | 2 | 5 | 4 | 4 | 3.75 | 4 | 5 | 0 | 3 | 4 |
| CSK2B | Casein kinase | 1.3564726 | 0.13081894 | 0.00049095 | 0.00037181 | 0.00049225 | 0.0004961 | 0.00046278 | 0.00078329 | 0.00047296 | 0.00078151 | 0.00047322 | 0.00062775 | 4 | 3 | 4 | 4 | 3.75 | 5 | 3 | 4 | 3 | 3.75 |
| RL22 | 60S ribosoma | 1.35560369 | 0.1064768 | 0.0022678 | 0.0014572 | 0.0022738 | 0.00125 | 0.0018122 | 0.0021051 | 0.0026481 | 0.0029536 | 0.0021197 | 0.00245663 | 11 | 7 | 11 | 6 | 8.75 | 8 | 10 | 9 | 8 | 8.75 |
| F193A | Protein FAM1 | 1.35451945 | 0.35282632 | 0 | 6.3193E-05 | 0 | 4.2159E-05 | 5.2676E-05 | 5.3252E-05 | 0.00010718 | 0 | 0.00005362 | 7.1351E-05 | 0 | 3 | 0 | 2 | 2.5 | 2 | 4 | 0 | 2 | 2.66666667 |
| WDCP | WD repeat an | 1.35282063 | 0.2609323 | 0.0005856 | 0.00081307 | 0.0011376 | 0.00059175 | 0.00078201 | 0.0011679 | 0.0012223 | 0.00052435 | 0.0013171 | 0.00105791 | 16 | 22 | 31 | 16 | 21.25 | 25 | 26 | 9 | 28 | 22 |
| AT2A2 | Sarcoplasmic | 1.35215194 | 0.20481698 | 0.00022792 | 0.0002813 | 0.00027931 | 0.00023032 | 0.00025471 | 0.00025859 | 0.00022771 | 0.00040313 | 0.00048821 | 0.00034441 | 9 | 11 | 11 | 9 | 10 | 8 | 7 | 10 | 15 | 10 |
| SH3G1 | Endophilin-A | 1.3507106 | 0.04781734 | 0.00035854 | 0.00050686 | 0.00050329 | 0.00065215 | 0.00050521 | 0.00064068 | 0.00064475 | 0.00079903 | 0.00064511 | 0.00068239 | 5 | 7 | 7 | 9 | 7 | 7 | 7 | 7 | 7 | 7 |
| CHRD1 | Cysteine and | 1.34867125 | 0.13809434 | 0.00087432 | 0.00072234 | 0.00055786 | 0.0011245 | 0.00081976 | 0.0013189 | 0.001021 | 0.0012652 | 0.00081722 | 0.00110558 | 11 | 9 | 7 | 14 | 10.25 | 13 | 10 | 10 | 8 | 10.25 |
| P5CR3 | Pyrroline-5-c | 1.34618916 | 0.03293188 | 0.00028892 | 0 | 0.00038626 | 0 | 0.00033759 | 0.0004917 | 0.00037112 | 0.00045992 | 0.0004951 | 0.00045446 | 3 | 0 | 4 | 0 | 3.5 | 4 | 3 | 3 | 4 | 3.5 |
| RL3 | 60S ribosoma | 1.34585663 | 0.05108197 | 0.0015715 | 0.0019175 | 0.0011818 | 0.0017865 | 0.00161433 | 0.0025073 | 0.0019345 | 0.0023974 | 0.0018514 | 0.00217265 | 24 | 29 | 18 | 27 | 24.5 | 30 | 23 | 23 | 22 | 24.5 |
| RL24 | 60S ribosoma | 1.3381355 | 0.01392636 | 0.0013446 | 0.0016972 | 0.0015167 | 0.0011889 | 0.00143685 | 0.0019308 | 0.002159 | 0.0018729 | 0.0017281 | 0.0019227 | 8 | 10 | 9 | 7 | 8.5 | 9 | 10 | 7 | 8 | 8.5 |
| RNF10 | RING finger p | 1.33737018 | 0.0393694 | 0.00019523 | 0.00032856 | 0.00032625 | 0.00026304 | 0.00027827 | 0.00041531 | 0.00033436 | 0.00036257 | 0.00037636 | 0.00037215 | 6 | 10 | 10 | 8 | 8.5 | 10 | 8 | 7 | 9 | 8.5 |
| TFB1M | Dimethylade | 1.33199753 | 0.18126116 | 0.0002288 | 0.00015403 | 0.00022941 | 0.0002312 | 0.00021086 | 0.00019469 | 0.00029389 | 0.00024281 | 0.00039207 | 0.00028087 | 3 | 2 | 3 | 3 | 2.75 | 2 | 3 | 2 | 4 | 2.75 |
| EIF3L | Eukaryotic tra | 1.3297501 | 0.24379567 | 0.00028073 | 0.00037796 | 0.00056295 | 0.00033096 | 0.00038815 | 0.00041803 | 0.00042069 | 0.00074479 | 0.00048106 | 0.00051614 | 6 | 8 | 12 | 7 | 8.25 | 7 | 7 | 10 | 8 | 8 |
| MYO1C | Unconventio | 1.32299969 | 0.36868555 | 7.4473E-05 | 5.0134E-05 | 0.00012445 | 7.5256E-05 | 8.1078E-05 | 0.00012674 | 6.3773E-05 | 7.9033E-05 | 0.00015952 | 0.00010727 | 3 | 2 | 5 | 3 | 3.25 | 4 | 2 | 2 | 5 | 3.25 |
| RS27 | 40S ribosoma | 1.32175031 | 0.3186891 | 0.00094244 | 0.0012689 | 0.0025199 | 0.0015872 | 0.00157961 | 0.0016039 | 0.0024211 | 0.0015002 | 0.0028262 | 0.00208785 | 3 | 4 | 8 | 5 | 5 | 4 | 6 | 3 | 7 | 5 |
| DIAP1 | Protein diaph | 1.32041232 | 0.26057664 | 8.2982E-05 | 6.2845E-05 | 0.000104 | 0.00016771 | 0.00010438 | 0.0001324 | 0.00015988 | 9.9071E-05 | 0.00015997 | 0.00013783 | 4 | 3 | 5 | 8 | 5 | 5 | 6 | 3 | 6 | 5 |
| sp\|Q9C0C9\|I | E2 ubiquitin- | 1.31997513 | 0.00670587 | 0.00024509 | 0.00024749 | 0.00026623 | 0.00020639 | 0.0002413 | 0.00031283 | 0.00031482 | 0.00035764 | 0.00028875 | 0.00031851 | 12 | 12 | 13 | 10 | 11.75 | 12 | 12 | 11 | 11 | 11.5 |
| SNX9 | Sorting nexin | 1.31996684 | 0.13626684 | 0.00088701 | 0.0008509 | 0.00057809 | 0.00062743 | 0.00073586 | 0.0010189 | 0.00079754 | 0.0012708 | 0.00079799 | 0.00097131 | 20 | 19 | 13 | 14 | 16.5 | 18 | 14 | 18 | 14 | 16 |

| | | | | | | | | | | | | | | | | | | | | | | | |
|---|---|---|---|---|---|---|---|---|---|---|---|---|---|---|---|---|---|---|---|---|---|---|---|
| MCM5 | DNA replicati | 1.31543876 | 0.08349372 | 0 | 0.00018152 | 7.2095E-05 | 0 | 0.00012681 | 0.00022944 | 0.00018472 | 0.00011446 | 0.00013861 | 0.00016681 | 0 | 5 | 2 | 0 | 3.5 | 5 | 4 | 2 | 3 | 3.5 |
| RS14 | 40S ribosoma | 1.30956829 | 0.0951903 | 0.0029709 | 0.0031764 | 0.0028036 | 0.0028255 | 0.0029441 | 0.0046842 | 0.0035916 | 0.004451 | 0.0026952 | 0.0038555 | 17 | 18 | 16 | 16 | 16.75 | 21 | 16 | 16 | 12 | 16.25 |
| KC1A | Casein kinase | 1.30786838 | 0.43405923 | 0.00054813 | 0.00047442 | 0.00078512 | 0.00087039 | 0.00066952 | 0.00079956 | 0.00080464 | 0.0014958 | 0.00040255 | 0.00087564 | 7 | 6 | 10 | 11 | 8.5 | 8 | 8 | 12 | 4 | 8 |
| TM10C | tRNA methyl | 1.30650043 | 0.29950002 | 0.00058932 | 0.00052896 | 0.00032827 | 0.00046317 | 0.00047743 | 0.00075219 | 0.00084108 | 0.0003127 | 0.00058908 | 0.00062376 | 9 | 8 | 5 | 7 | 7.25 | 9 | 10 | 3 | 7 | 7.25 |
| MPCP | Phosphate ca | 1.30623672 | 0.05171361 | 0.00087475 | 0.00066248 | 0.0007309 | 0.0005893 | 0.00071436 | 0.00083739 | 0.0011236 | 0.00092831 | 0.00084318 | 0.00093312 | 12 | 9 | 10 | 8 | 9.75 | 9 | 12 | 8 | 9 | 9.5 |
| PGAM5 | Serine/threo | 1.30503409 | 0.28021173 | 0.00054786 | 0.00046101 | 0.00054932 | 0.00036907 | 0.00048182 | 0.00069927 | 0.00070371 | 0.0002907 | 0.00082146 | 0.00062879 | 6 | 5 | 6 | 4 | 5.25 | 6 | 6 | 2 | 7 | 5.25 |
| DPM1 | Dolichol-pho | 1.30219554 | 0.30039374 | 0.00050747 | 0.00051243 | 0.00040706 | 0.00061536 | 0.00051058 | 0.00077727 | 0.0003911 | 0.00096937 | 0.00052176 | 0.00066488 | 5 | 5 | 4 | 6 | 5 | 6 | 3 | 6 | 4 | 4.75 |
| C2D1A | Coiled-coil ar | 1.29967443 | 0.05404726 | 0.00027748 | 0.00025217 | 0.00019475 | 0.00025236 | 0.00024419 | 0.00035417 | 0.00024949 | 0.00030919 | 0.00035662 | 0.00031737 | 10 | 9 | 7 | 9 | 8.75 | 10 | 7 | 7 | 10 | 8.5 |
| KIF14 | Kinesin-like p | 1.29666898 | 0.3844075 | 0.00022417 | 0.00017786 | 0.00012844 | 0.00016181 | 0.00017307 | 0.00026569 | 0.00024681 | 7.6468E-05 | 0.00030869 | 0.00022441 | 14 | 11 | 8 | 10 | 10.75 | 13 | 12 | 3 | 15 | 10.75 |
| M2OM | Mitochondria | 1.29382712 | 0.15400735 | 0.0010925 | 0.00067889 | 0.0009269 | 0.0010191 | 0.00092935 | 0.0013945 | 0.0010795 | 0.0014716 | 0.00086406 | 0.00120242 | 13 | 8 | 11 | 12 | 11 | 13 | 10 | 11 | 8 | 10.5 |
| UBP2L | Ubiquitin-ass | 1.29071663 | 0.15371332 | 0.0010439 | 0.0010786 | 0.00075457 | 0.0010548 | 0.00098297 | 0.0014873 | 0.0013097 | 0.00081153 | 0.0014664 | 0.00126873 | 43 | 44 | 31 | 43 | 40.25 | 48 | 42 | 21 | 47 | 39.5 |
| RL23A | 60S ribosoma | 1.28966941 | 0.19809173 | 0.0015224 | 0.0017081 | 0.0011872 | 0.0015384 | 0.00148903 | 0.0015114 | 0.0019555 | 0.0026927 | 0.0015218 | 0.00192035 | 9 | 10 | 7 | 9 | 8.75 | 7 | 9 | 10 | 7 | 8.25 |
| TBC15 | TBC1 domain | 1.28932328 | 0.35877366 | 0.00015275 | 0.00011569 | 0.00022974 | 0.00019295 | 0.00017278 | 0.00024372 | 0.00024526 | 0.00030395 | 0.00009816 | 0.00022277 | 4 | 3 | 6 | 5 | 4.5 | 5 | 5 | 5 | 2 | 4.25 |
| PHB2 | Prohibitin-2 ( | 1.28818989 | 0.00612121 | 0.0033537 | 0.0032974 | 0.0026547 | 0.0032106 | 0.0031291 | 0.0037174 | 0.0041944 | 0.0043552 | 0.0038565 | 0.00403088 | 38 | 37 | 30 | 36 | 35.25 | 33 | 37 | 31 | 34 | 33.75 |
| FACR1 | Fatty acyl-Co | 1.27943206 | 0.19917109 | 0.00010248 | 0.00010348 | 0 | 0.00020711 | 0.00013769 | 0.0001308 | 0.00019745 | 0.0002447 | 0.00013171 | 0.00017617 | 2 | 2 | 0 | 4 | 2.66666667 | 2 | 3 | 3 | 2 | 2.5 |
| NCBP1 | Nuclear cap-l | 1.27797022 | 0.64694012 | 0.00016702 | 0.00013492 | 0.00013397 | 0 | 0.0001453 | 0.0001279 | 0.00021453 | 0 | 0.00021465 | 0.00018569 | 5 | 4 | 4 | 0 | 4.33333333 | 3 | 5 | 0 | 5 | 4.33333333 |
| TANC1 | Protein TANC | 1.2757261 | 0.90476996 | 0.00015598 | 0.00010023 | 0.00012796 | 0.00012896 | 0.00012828 | 0.00014479 | 0.00014571 | 0 | 0.00020046 | 0.00016365 | 11 | 7 | 9 | 9 | 9 | 8 | 8 | 0 | 11 | 9 |
| RS26 | 40S ribosoma | 1.27364413 | 0.03264501 | 0.0016063 | 0.0011585 | 0.0013805 | 0.0011594 | 0.00132618 | 0.0017573 | 0.0017685 | 0.0014611 | 0.0017694 | 0.00168908 | 7 | 5 | 6 | 5 | 5.75 | 6 | 6 | 4 | 6 | 5.5 |
| SKA1 | Spindle and k | 1.27203088 | 0.65140161 | 0.00020697 | 0.00031349 | 0 | 0.00031371 | 0.00027806 | 0.00039625 | 0.00026585 | 0 | 0.00039899 | 0.0003537 | 2 | 3 | 0 | 3 | 2.66666667 | 3 | 2 | 0 | 3 | 2.66666667 |
| VDAC1 | Voltage-depe | 1.27115858 | 0.92184157 | 0.00027974 | 0.00018831 | 0.00028048 | 0.0003769 | 0.00028136 | 0 | 0.00023954 | 0.00059373 | 0.00023968 | 0.00035765 | 9 | 5 | 8 | 9 | 7.75 | 0 | 3 | 7 | 5 | 5 |
| RAGP1 | Ran GTPase-a | 1.26930505 | 0.00835393 | 0.0028321 | 0.0027237 | 0.0034257 | 0.0031345 | 0.003029 | 0.0039592 | 0.0037533 | 0.0035065 | 0.0041599 | 0.00384473 | 63 | 60 | 76 | 69 | 67 | 69 | 65 | 49 | 72 | 63.75 |
| NEPRO | Nucleolus an | 1.26929818 | 0.89749119 | 0.00013962 | 0.00023498 | 0.00018666 | 0.00018812 | 0.00018735 | 0.00023761 | 0 | 0.00029634 | 0.00017944 | 0.0002378 | 3 | 5 | 4 | 4 | 4 | 4 | 0 | 4 | 3 | 3.66666667 |
| VIME | Vimentin OS= | 1.26814753 | 0.05167911 | 0.0065122 | 0.0052035 | 0.0071541 | 0.0066378 | 0.0063769 | 0.0084565 | 0.0070555 | 0.0073015 | 0.0095339 | 0.00808685 | 115 | 91 | 126 | 116 | 112 | 117 | 97 | 81 | 131 | 106.5 |
| TCPZ | T-complex pr | 1.26501438 | 0.06883009 | 0.0015903 | 0.0013549 | 0.0013454 | 0.0013057 | 0.00139908 | 0.0013955 | 0.0021065 | 0.0016613 | 0.0019161 | 0.00176985 | 32 | 27 | 27 | 26 | 28 | 22 | 33 | 21 | 30 | 26.5 |
| MSH2 | DNA mismatc | 1.26455754 | 0.27486066 | 0.00022602 | 0.00042794 | 0.00031161 | 0.0002855 | 0.00031277 | 0.00050486 | 0.00029033 | 0.00031482 | 0.00047204 | 0.00039551 | 8 | 15 | 11 | 10 | 11 | 14 | 8 | 7 | 13 | 10.5 |
| SRP68 | Signal recogr | 1.26314225 | 0.36418983 | 0.00029461 | 0.00021249 | 0.00037979 | 0.00042529 | 0.00032805 | 0.00053719 | 0.00043248 | 0.00020099 | 0.00048681 | 0.00041437 | 7 | 5 | 9 | 10 | 7.75 | 10 | 8 | 3 | 9 | 7.5 |
| PABP1 | Polyadenylat | 1.25925292 | 0.23290432 | 0.0018256 | 0.0018854 | 0.0019553 | 0.0019286 | 0.00189873 | 0.0030186 | 0.0025582 | 0.0013209 | 0.0026662 | 0.00239098 | 44 | 45 | 47 | 46 | 45.5 | 57 | 48 | 20 | 50 | 43.75 |
| VDAC3 | Voltage-depe | 1.25848072 | 0.05381111 | 0.0018649 | 0.0013182 | 0.0014024 | 0.0014134 | 0.00149973 | 0.0016662 | 0.0017966 | 0.0019296 | 0.0021571 | 0.00188738 | 20 | 14 | 15 | 15 | 16 | 14 | 15 | 13 | 18 | 15 |
| TSPO | Translocator | 1.24848185 | 0.31537082 | 0.00046843 | 0.00047301 | 0.00046968 | 0.00078892 | 0.00055001 | 0.0007972 | 0.00040113 | 0.00074567 | 0.00080271 | 0.00068668 | 3 | 3 | 3 | 5 | 3.5 | 4 | 2 | 3 | 4 | 3.25 |
| EXOS4 | Exosome con | 1.24408783 | 0.86496727 | 0.00096937 | 0.00065257 | 0.00064797 | 0.0005442 | 0.00070353 | 0.0005499 | 0.00096845 | 0 | 0.0011074 | 0.00087525 | 9 | 6 | 6 | 5 | 6.5 | 4 | 7 | 0 | 8 | 6.33333333 |
| PTBP1 | Polypyrimidir | 1.24184878 | 0.33276842 | 0.00084483 | 0.00080291 | 0.00064776 | 0.00065283 | 0.00073708 | 0.00088803 | 0.0010852 | 0.00047465 | 0.0012135 | 0.00091535 | 17 | 16 | 13 | 13 | 14.75 | 14 | 17 | 6 | 19 | 14 |
| SYWC | Tryptophan-- | 1.23893416 | 0.12661379 | 0.00067232 | 0.00062232 | 0.00078646 | 0.00050953 | 0.00064766 | 0.00092964 | 0.00079162 | 0.0006243 | 0.00086406 | 0.00080241 | 12 | 11 | 14 | 9 | 11.5 | 13 | 11 | 7 | 12 | 10.75 |
| TTC37 | Tetratricoper | 1.23857019 | 0.37560237 | 3.3745E-05 | 5.1112E-05 | 3.3835E-05 | 5.1149E-05 | 4.246E-05 | 4.3071E-05 | 4.3345E-05 | 8.0575E-05 | 4.3369E-05 | 0.00005259 | 2 | 3 | 2 | 3 | 2.5 | 2 | 2 | 3 | 2 | 2.25 |
| HBS1L | HBS1-like prc | 1.23782759 | 0.26112151 | 0.00065585 | 0.00042853 | 0.00046419 | 0.00062376 | 0.00054308 | 0.00073863 | 0.00044599 | 0.00085978 | 0.00064457 | 0.00067224 | 17 | 11 | 12 | 16 | 14 | 15 | 9 | 14 | 13 | 12.75 |
| ACOT9 | Acyl-coenzyn | 1.23458461 | 0.85641303 | 0.00030055 | 0.00030349 | 0.00036162 | 0.00036445 | 0.00033253 | 0.00061379 | 0.00038605 | 0 | 0.00023176 | 0.00041053 | 5 | 5 | 6 | 6 | 5.5 | 8 | 5 | 0 | 3 | 5.33333333 |
| ELAV1 | ELAV-like prc | 1.23246057 | 0.13429742 | 0.00064757 | 0.0006539 | 0.00064929 | 0.00065437 | 0.00065128 | 0.00061991 | 0.00083179 | 0.0010308 | 0.00072822 | 0.00080268 | 8 | 8 | 8 | 8 | 8 | 6 | 8 | 8 | 7 | 7.25 |
| CNO11 | CCR4-NOT tra | 1.23219403 | 0.0417454 | 0.00031045 | 0.00036574 | 0.0002594 | 0.00031371 | 0.00031233 | 0.00039625 | 0.00033231 | 0.00041183 | 0.00039899 | 0.00038485 | 6 | 7 | 5 | 6 | 6 | 6 | 5 | 5 | 6 | 5.5 |
| PFKAM | ATP-depende | 1.22092986 | 0.36706121 | 0.00013533 | 0.00010249 | 0.00010176 | 6.8373E-05 | 0.00010199 | 0.00012954 | 8.6912E-05 | 0.00010771 | 0.00017392 | 0.00012452 | 4 | 4 | 3 | 2 | 3.25 | 3 | 2 | 3 | 5 | 3.25 |
| PYRG1 | CTP synthase | 1.21984639 | 0.21057284 | 0.00084836 | 0.00063122 | 0.00089539 | 0.00054143 | 0.0007291 | 0.00079787 | 0.0010897 | 0.00092399 | 0.000746 | 0.00088939 | 19 | 14 | 20 | 12 | 16.25 | 14 | 19 | 13 | 13 | 14.75 |
| ZW10 | Centromere/ | 1.21803895 | 0.2671274 | 0.00010162 | 0 | 0.00010189 | 0.00013692 | 0.00011348 | 8.6474E-05 | 0.00021756 | 0.00016177 | 8.7072E-05 | 0.00013822 | 3 | 0 | 3 | 4 | 3.33333333 | 2 | 5 | 3 | 2 | 3 |
| NDUA9 | NADH dehyd | 1.21517772 | 0.79069438 | 0.00027998 | 0.00021204 | 0.00028073 | 0.00028292 | 0.00026392 | 0.00026802 | 0 | 0.00033427 | 0.00035983 | 0.00032071 | 4 | 3 | 4 | 4 | 3.75 | 3 | 0 | 3 | 4 | 3.33333333 |
| MIC60 | MICOS comp | 1.20806773 | 0.12421125 | 0.00066145 | 0.000457 | 0.00062831 | 0.00045733 | 0.00055102 | 0.00066652 | 0.00076019 | 0.00060959 | 0.00062639 | 0.00066567 | 19 | 13 | 18 | 13 | 15.75 | 15 | 17 | 11 | 14 | 14.25 |
| PCBP1 | Poly(rC)-bind | 1.20289386 | 0.25280957 | 0.0012601 | 0.0021706 | 0.0015608 | 0.0014981 | 0.0016224 | 0.0019868 | 0.0023803 | 0.0015339 | 0.0019053 | 0.00195158 | 17 | 29 | 21 | 20 | 21.75 | 21 | 25 | 13 | 20 | 19.75 |
| EF1D | Elongation fa | 1.20280063 | 0.26781652 | 0.00075127 | 0.00056896 | 0.00065911 | 0.00056937 | 0.00063718 | 0.00047945 | 0.00084437 | 0.00089693 | 0.00084484 | 0.0007664 | 8 | 6 | 7 | 6 | 6.75 | 4 | 7 | 6 | 7 | 6 |
| EIF3C | Eukaryotic tr | 1.20013531 | 0.30878739 | 0.00049135 | 0.00035023 | 0.00055062 | 0.00023365 | 0.00040646 | 0.00047958 | 0.00044551 | 0.0005061 | 0.00052005 | 0.00048781 | 17 | 12 | 19 | 8 | 14 | 13 | 12 | 11 | 14 | 12.5 |
| EIFCL | Eukaryotic tr | 1.20013531 | 0.30878739 | 0.00049135 | 0.00035023 | 0.00055062 | 0.00023365 | 0.00040646 | 0.00047958 | 0.00044551 | 0.0005061 | 0.00052005 | 0.00048781 | 17 | 12 | 19 | 8 | 14 | 13 | 12 | 11 | 14 | 12.5 |
| MOT1 | Monocarbox | 1.1991418 | 0.51709378 | 0.0011611 | 0.0007461 | 0.00074084 | 0.00058664 | 0.00080867 | 0.0010104 | 0.00061012 | 0.0015122 | 0.00074612 | 0.00096971 | 22 | 14 | 14 | 11 | 15.25 | 15 | 9 | 18 | 11 | 13.25 |
| ARHG2 | Rho guanine | 1.19879727 | 0.78686821 | 8.0289E-05 | 0.00016215 | 0.00010734 | 0.00010818 | 0.00011449 | 0.00013664 | 0.00010313 | 0 | 0.00017198 | 0.00013725 | 3 | 6 | 4 | 4 | 4.25 | 4 | 3 | 0 | 5 | 4 |
| RS9 | 40S ribosoma | 1.19818944 | 0.40105596 | 0.0017683 | 0.0012362 | 0.001773 | 0.00096216 | 0.00143492 | 0.0010417 | 0.0017472 | 0.0021653 | 0.001923 | 0.0017193 | 13 | 9 | 13 | 7 | 10.5 | 6 | 10 | 10 | 11 | 9.25 |
| UBC12 | NEDD8-conju | 1.19759812 | 0.60770716 | 0.0004326 | 0.0011649 | 0.00057833 | 0.00102 | 0.00079896 | 0.0016565 | 0.00074089 | 0.00068863 | 0.0007413 | 0.00095683 | 3 | 8 | 4 | 7 | 5.5 | 9 | 4 | 3 | 4 | 5 |
| RL6 | 60S ribosoma | 1.19592672 | 0.1129711 | 0.0025655 | 0.0017579 | 0.0022968 | 0.0021295 | 0.00218743 | 0.002339 | 0.0025892 | 0.0030629 | 0.0024729 | 0.002616 | 28 | 19 | 25 | 23 | 23.75 | 20 | 22 | 21 | 21 | 21 |
| TTF2 | Transcription | 1.19002028 | 0.44282689 | 0.00049961 | 0.00055036 | 0.00031878 | 0.00036717 | 0.00043398 | 0.00049276 | 0.00067091 | 0.0002892 | 0.00061291 | 0.00051645 | 22 | 24 | 14 | 16 | 19 | 17 | 23 | 8 | 21 | 17.25 |
| MMTA2 | Multiple mye | 1.18881269 | 0.35739661 | 0.00080269 | 0.00081054 | 0.00070422 | 0.00070973 | 0.0007568 | 0.0007684 | 0.0012888 | 0.00063888 | 0.00090267 | 0.00089969 | 8 | 8 | 7 | 7 | 7.5 | 6 | 10 | 4 | 7 | 6.75 |
| NU188 | Nucleoporin | 1.18762411 | 0.64749254 | 3.0175E-05 | 0.00012188 | 9.0767E-05 | 4.5739E-05 | 7.214E-05 | 0.00011555 | 0.00003876 | 7.2052E-05 | 0.00011634 | 8.5676E-05 | 2 | 8 | 6 | 3 | 4.75 | 6 | 2 | 3 | 6 | 4.25 |
| NEMP1 | Nuclear enve | 1.18503465 | 0.40949756 | 0 | 0.00018004 | 0 | 0.00012012 | 0.00015008 | 0.00015172 | 0.00015268 | 0 | 0.00022915 | 0.00017785 | 0 | 3 | 0 | 2 | 2.5 | 2 | 2 | 0 | 3 | 2.33333333 |
| DRG2 | Development | 1.18243819 | 0.4600303 | 0.00021749 | 0.00043923 | 0.00029075 | 0.00036629 | 0.00032844 | 0.00055519 | 0.00027936 | 0.0003462 | 0.00037269 | 0.00038836 | 3 | 6 | 4 | 5 | 4.5 | 6 | 3 | 3 | 4 | 4 |
| NU205 | Nuclear pore | 1.17825257 | 0.04294475 | 0.00024919 | 0.00027812 | 0.00027616 | 0.00026507 | 0.00026714 | 0.00028459 | 0.00028639 | 0.00033405 | 0.00035398 | 0.00031475 | 19 | 21 | 21 | 20 | 20.25 | 17 | 17 | 16 | 21 | 17.75 |
| PRKDC | DNA-depend | 1.17618683 | 0.10358148 | 0.00050501 | 0.0003873 | 0.00051277 | 0.00056845 | 0.00049338 | 0.00053851 | 0.00059941 | 0.00064108 | 0.00054224 | 0.00058031 | 79 | 60 | 80 | 88 | 76.75 | 66 | 73 | 63 | 66 | 67 |
| AKA11 | A-kinase ancl | 1.17249495 | 0.60496622 | 0.00018046 | 0.00019624 | 0.00020877 | 0.00011222 | 0.00017442 | 0.00024805 | 0.00016047 | 8.8388E-05 | 0.00032113 | 0.00020451 | 13 | 14 | 15 | 8 | 12.5 | 14 | 9 | 4 | 18 | 11.25 |
| EDC3 | Enhancer of m | 1.17189964 | 0.22098534 | 0 | 0.00015736 | 0.00026042 | 0.00020997 | 0.00020925 | 0.00019891 | 0.00033362 | 0.00024807 | 0.00020028 | 0.00024522 | 0 | 3 | 5 | 4 | 4 | 3 | 5 | 3 | 3 | 3.5 |
| VIGLN | Vigilin OS=Hc | 1.16858565 | 0.02594605 | 0.0018522 | 0.0019544 | 0.0018988 | 0.0022502 | 0.0019889 | 0.0024438 | 0.0023791 | 0.0021202 | 0.0023537 | 0.0023242 | 89 | 93 | 91 | 107 | 95 | 92 | 89 | 64 | 88 | 83.25 |
| LIMD1 | LIM domain-c | 1.16837662 | 0.57931746 | 0.00054651 | 0.00047301 | 0.00046968 | 0.0005128 | 0.0005005 | 0.0005979 | 0.0008524 | 0.00018642 | 0.00070237 | 0.00058477 | 14 | 12 | 12 | 13 | 12.75 | 12 | 17 | 3 | 14 | 11.5 |
| HNRH1 | Heterogeneo | 1.15771798 | 0.38819132 | 0.00099912 | 0.00065281 | 0.00088392 | 0.00071267 | 0.00081213 | 0.0012002 | 0.0010569 | 0.00074844 | 0.00075533 | 0.00094022 | 17 | 11 | 15 | 12 | 13.75 | 16 | 14 | 8 | 10 | 12 |
| HCD2 | 3-hydroxyacy | 1.15124401 | 0.27993473 | 0.00050553 | 0.00091884 | 0.00081099 | 0.00071517 | 0.00073763 | 0.00090334 | 0.00090908 | 0.00080472 | 0.00077964 | 0.0008492 | 5 | 9 | 8 | 7 | 7.25 | 7 | 7 | 5 | 6 | 6.25 |
| ARF4 | ADP-ribosyla | 1.14234619 | 0.11051512 | 0.0026388 | 0.0020725 | 0.0023519 | 0.0026666 | 0.00243245 | 0.0028068 | 0.002448 | 0.0030338 | 0.0028262 | 0.0027787 | 18 | 14 | 16 | 18 | 16.5 | 15 | 13 | 13 | 15 | 14 |
| ASPP2 | Apoptosis-sti | 1.14183172 | 0.49936409 | 0.00028073 | 0.00023623 | 0.00023456 | 0.00033096 | 0.00027062 | 0.00032846 | 0.00042069 | 0.0001862 | 0.00030066 | 0.000309 | 12 | 10 | 10 | 14 | 11.5 | 11 | 14 | 5 | 10 | 10 |
| ZN326 | DBIRD compl | 1.14005405 | 0.48590567 | 0.00045341 | 0.00036627 | 0.00054554 | 0.00036654 | 0.00043294 | 0.00057872 | 0.00052416 | 0.0002887 | 0.00058272 | 0.00049358 | 10 | 8 | 12 | 8 | 9.5 | 10 | 9 | 4 | 10 | 8.25 |
| DNJA1 | DnaJ homolo | 1.13857846 | 0.2557075 | 0.00053176 | 0.00040272 | 0.00046653 | 0.00033584 | 0.00043421 | 0.00050904 | 0.0004269 | 0.00052904 | 0.00051256 | 0.00049439 | 8 | 6 | 7 | 5 | 6.5 | 6 | 5 | 5 | 6 | 5.5 |
| SC23A | Protein trans | 1.13365067 | 0.43270603 | 0.00010348 | 0.0001045 | 0 | 0 | 0.00010399 | 0.00013208 | 0.00013292 | 0 | 8.8665E-05 | 0.00011789 | 3 | 3 | 0 | 0 | 3 | 3 | 3 | 0 | 2 | 2.66666667 |
| DHCR7 | 7-dehydroch | 1.133598 | 0.45433 | 0 | 0.0001122 | 0.00022281 | 0 | 0.00016751 | 0.00021273 | 0.00014272 | 0 | 0.0002142 | 0.00018988 | 0 | 2 | 4 | 0 | 3 | 3 | 2 | 0 | 3 | 2.66666667 |
| CDR2L | Cerebellar de | 1.13353564 | 0.30987874 | 0.00045399 | 0.00034383 | 0.000569 | 0.00045876 | 0.0004564 | 0.00057947 | 0.00051026 | 0.00054202 | 0.00043761 | 0.00051734 | 8 | 6 | 10 | 8 | 8 | 8 | 7 | 6 | 6 | 6.75 |
| PTN23 | Tyrosine-pro | 1.13307156 | 0.68170719 | 8.0649E-05 | 0.00006515 | 4.8518E-05 | 4.8898E-05 | 6.0804E-05 | 8.2351E-05 | 8.2874E-05 | 0 | 0.00004146 | 6.8895E-05 | 5 | 4 | 3 | 3 | 3.75 | 4 | 4 | 0 | 2 | 3.33333333 |
| MTX2 | Metaxin-2 OS | 1.12876937 | 0.69625389 | 0.00030101 | 0.00050659 | 0.00040241 | 0.00030417 | 0.00037855 | 0.00025613 | 0 | 0.00063888 | 0.00038686 | 0.00042729 | 3 | 5 | 4 | 3 | 3.75 | 2 | 0 | 4 | 3 | 3 |
| UIF | UAP56-intera | 1.12038562 | 0.03554175 | 0.00024895 | 0 | 0 | 0 | 0.00024895 | 0.00021183 | 0.00042636 | 0.00026419 | 0.0002133 | 0.00027892 | 3 | 0 | 0 | 0 | 3 | 2 | 4 | 2 | 2 | 2.5 |
| NUP93 | Nuclear pore | 1.11575365 | 0.41640954 | 0.00077329 | 0.00078085 | 0.0005492 | 0.00052094 | 0.00065607 | 0.00086363 | 0.00070357 | 0.00061548 | 0.00074537 | 0.00073201 | 24 | 24 | 17 | 16 | 20.25 | 21 | 17 | 12 | 18 | 17 |
| ILF3 | Interleukin er | 1.11573211 | 0.66746718 | 0.00029517 | 0.00017884 | 0.00044394 | 0.00020879 | 0.00028169 | 0.00026373 | 0.00041706 | 0.00023493 | 0.00034142 | 0.00031429 | 10 | 6 | 15 | 7 | 9.5 | 7 | 11 | 5 | 9 | 8 |
| VATH | V-type proto | 1.11342658 | 0.63545554 | 0.00027317 | 0.00016551 | 0.00021912 | 0.00022083 | 0.00021966 | 0.0002092 | 0.00014035 | 0.00034788 | 0.00028086 | 0.00024457 | 5 | 3 | 4 | 4 | 4 | 3 | 2 | 4 | 4 | 3.25 |
| GASP2 | G-protein cou | 1.11197427 | 0.85105323 | 0 | 0.00012719 | 6.3147E-05 | 6.3641E-05 | 8.4659E-05 | 0.00012058 | 8.0896E-05 | 0 | 8.0941E-05 | 9.4139E-05 | 0 | 4 | 2 | 2 | 2.66666667 | 3 | 2 | 0 | 2 | 2.33333333 |
| HNRPL | Heterogeneo | 1.10757593 | 0.61230507 | 0.00076164 | 0.00085956 | 0.00053906 | 0.00081491 | 0.00074379 | 0.00057184 | 0.00097831 | 0 | 0.00092127 | 0.00082381 | 17 | 19 | 12 | 18 | 16.5 | 10 | 17 | 0 | 16 | 14.3333333 |
| SERA | D-3-phospho | 1.10751773 | 0.61963703 | 0.00024755 | 0.00039995 | 0.00034749 | 0.00045026 | 0.00036131 | 0.00031596 | 0.00031797 | 0.00039405 | 0.00057266 | 0.00040016 | 5 | 8 | 7 | 9 | 7.25 | 5 | 5 | 5 | 9 | 6 |
| TBG1 | Tubulin gamr | 1.10445423 | 0.11082261 | 0.00081915 | 0.00076808 | 0.000704 | 0.00070951 | 0.00075019 | 0.00074682 | 0.00082672 | 0.00083826 | 0.00090238 | 0.00082855 | 14 | 13 | 12 | 12 | 12.75 | 10 | 11 | 9 | 12 | 10.5 |
| CBSL | Cystathionin | 1.1008846 | 0.59353106 | 0.00014368 | 0.0002418 | 0.00019208 | 0.00014518 | 0.00018069 | 0.00018338 | 0 | 0.00022871 | 0.00018465 | 0.00019891 | 3 | 5 | 4 | 3 | 3.75 | 3 | 0 | 3 | 3 | 3 |
| CBS | Cystathionin | 1.1008846 | 0.59353106 | 0.00014368 | 0.0002418 | 0.00019208 | 0.00014518 | 0.00018069 | 0.00018338 | 0 | 0.00022871 | 0.00018465 | 0.00019891 | 3 | 5 | 4 | 3 | 3.75 | 3 | 0 | 3 | 3 | 3 |
| VP13C | Vacuolar pro | 1.09674988 | 0.63972191 | 2.8125E-05 | 0.0000284 | 0.0000423 | 5.6841E-05 | 3.8917E-05 | 2.6924E-05 | 4.5158E-05 | 5.5963E-05 | 0 | 4.2682E-05 | 4 | 4 | 6 | 8 | 5.5 | 3 | 5 | 5 | 0 | 4.33333333 |
| GUAA | GMP synthas | 1.08728463 | 0.64622974 | 0.00011424 | 0.0001538 | 0.00011454 | 0.00015391 | 0.00013412 | 9.7205E-05 | 9.7823E-05 | 0.00024246 | 0 | 0.00014583 | 3 | 4 | 3 | 4 | 3.5 | 2 | 2 | 4 | 0 | 2.66666667 |
| PUR6 | Multifunction | 1.08717379 | 0.43143282 | 0.0018627 | 0.0011913 | 0.0016809 | 0.0015686 | 0.00157588 | 0.001902 | 0.0017546 | 0.0016802 | 0.0015162 | 0.00171325 | 30 | 19 | 27 | 25 | 25.25 | 24 | 22 | 17 | 19 | 20.5 |
| EXOS6 | Exosome con | 1.08260821 | 0.59574279 | 0.0011642 | 0.0012735 | 0.00087547 | 0.00098035 | 0.00107338 | 0.00061915 | 0.0013708 | 0 | 0.0014962 | 0.00116205 | 12 | 13 | 9 | 10 | 11 | 5 | 11 | 0 | 12 | 9.33333333 |
| SMC1A | Structural ma | 1.08234214 | 0.5605367 | 6.4205E-05 | 6.4833E-05 | 6.4376E-05 | 4.3253E-05 | 5.9167E-05 | 5.4634E-05 | 8.2471E-05 | 0 | 5.5011E-05 | 6.4039E-05 | 3 | 3 | 3 | 2 | 2.75 | 2 | 3 | 0 | 2 | 2.33333333 |
| SYK | Lysine--tRNA | 1.08038059 | 0.59637179 | 8.8403E-05 | 0.00017854 | 0.00013296 | 8.9332E-05 | 0.00012231 | 0.00016925 | 0.00011355 | 0 | 0.00011362 | 0.00013214 | 2 | 4 | 3 | 2 | 2.75 | 3 | 2 | 0 | 2 | 2.33333333 |
| RL4 | 60S ribosoma | 1.07149559 | 0.3024063 | 0.0011742 | 0.0013729 | 0.0012393 | 0.0014363 | 0.00130568 | 0.0012621 | 0.0015082 | 0.0014756 | 0.0013502 | 0.00139903 | 19 | 22 | 20 | 23 | 21 | 16 | 19 | 15 | 17 | 16.75 |
| DDX54 | ATP-depende | 1.06644244 | 0.67038466 | 0.00026958 | 0.0003327 | 0.00024026 | 0.00027241 | 0.00027874 | 0.00034408 | 0.00034627 | 0.00019072 | 0.00030796 | 0.00029726 | 9 | 11 | 8 | 9 | 9.25 | 9 | 9 | 4 | 8 | 7.5 |
| TRIPB | Thyroid recep | 1.06454039 | 0.87958917 | 6.6671E-05 | 5.3858E-05 | 0.00016044 | 6.7371E-05 | 8.7085E-05 | 6.8078E-05 | 0.00011989 | 0.00014858 | 3.4274E-05 | 9.2706E-05 | 5 | 4 | 12 | 5 | 6.5 | 4 | 7 | 7 | 2 | 5 |
| SYYM | Tyrosine--tRN | 1.05848881 | 0.53012949 | 0.00011064 | 0.00016759 | 0.00022188 | 0.00016771 | 0.00016696 | 0.00021183 | 0 | 0.00017613 | 0.0001422 | 0.00017672 | 2 | 3 | 4 | 3 | 3 | 3 | 0 | 2 | 2 | 2.33333333 |
| HLTF | Helicase-like | 1.0524751 | 0.84263513 | 0.00013077 | 7.9226E-05 | 5.2445E-05 | 7.9283E-05 | 8.5431E-05 | 6.6762E-05 | 0.00010078 | 0.00012489 | 6.7224E-05 | 8.9914E-05 | 5 | 3 | 2 | 3 | 3.25 | 2 | 3 | 3 | 2 | 2.5 |
| DDX47 | Probable ATF | 1.05154495 | 0.74547602 | 0.00040598 | 0.00052707 | 0.00063966 | 0.00035163 | 0.00048109 | 0.00044415 | 0.00052147 | 0.00046161 | 0.0005963 | 0.00050588 | 7 | 9 | 11 | 6 | 8.25 | 6 | 7 | 5 | 8 | 6.5 |
| PRDX6 | Peroxiredoxi | 1.05070545 | 0.7677346 | 0.0024739 | 0.0029739 | 0.0030711 | 0.0034522 | 0.00299278 | 0.0036087 | 0.0021185 | 0.0041256 | 0.0027253 | 0.00314453 | 21 | 25 | 26 | 29 | 25.25 | 24 | 14 | 22 | 18 | 19.5 |
| SREK1 | Splicing regul | 1.04520522 | 0.48125608 | 0.00015584 | 0.00015736 | 0.00015625 | 0.00020997 | 0.00016986 | 0.00019891 | 0.00020017 | 0 | 0.00013352 | 0.00017753 | 3 | 3 | 3 | 4 | 3.25 | 3 | 3 | 0 | 2 | 2.66666667 |
| TADBP | TAR DNA-bin | 1.04468008 | 0.83188037 | 0.00044618 | 0.00038618 | 0.00038346 | 0.00038646 | 0.00040057 | 0.00048814 | 0.00040937 | 0.00020293 | 0.00057343 | 0.00041847 | 7 | 6 | 6 | 6 | 6.25 | 6 | 5 | 2 | 7 | 5 |
| MCM4 | DNA replicati | 1.04387537 | 0.65776058 | 0.00064213 | 0.00083367 | 0.00085845 | 0.00083427 | 0.00079213 | 0.00066348 | 0.00090336 | 0.00087614 | 0.00086456 | 0.00082689 | 21 | 27 | 28 | 27 | 25.75 | 17 | 23 | 18 | 22 | 20 |
| FAS | Fatty acid syr | 1.0280666 | 0.83217138 | 0.0011035 | 0.0010081 | 0.00095888 | 0.0011363 | 0.0010517 | 0.00099261 | 0.00098542 | 0.0014554 | 0.00089142 | 0.00108121 | 105 | 95 | 91 | 107 | 99.5 | 74 | 73 | 87 | 66 | 75 |
| COPA | Coatomer su | 1.0199091 | 0.87031733 | 0.00038806 | 0.00037009 | 0.00036748 | 0.00034857 | 0.00036855 | 0.00027518 | 0.00036 | 0.00048046 | 0.00038791 | 0.00037589 | 18 | 17 | 17 | 16 | 17 | 10 | 13 | 14 | 14 | 12.75 |
| COPE | Coatomer su | 1.01978609 | 0.48744426 | 0.00042838 | 0.00069212 | 0.00042952 | 0.00060604 | 0.00053902 | 0.00032807 | 0.0006603 | 0 | 0.00066067 | 0.00054968 | 5 | 8 | 5 | 7 | 6.25 | 3 | 6 | 0 | 6 | 5 |
| SRRM1 | Serine/argini | 1.01825112 | 0.51714504 | 0.00023353 | 0.00020633 | 0.00017561 | 0.00041296 | 0.00025711 | 0.00033533 | 0.00022497 | 0 | 0.0002251 | 0.0002618 | 8 | 7 | 6 | 14 | 8.75 | 9 | 6 | 0 | 6 | 7 |
| GPN1 | GPN-loop GT | 1.01817005 | 0.97474805 | 0.00021167 | 0.00035624 | 0 | 0.0001426 | 0.00023684 | 0.00027017 | 0.00027189 | 0 | 0.00018136 | 0.00024114 | 3 | 5 | 0 | 2 | 3.33333333 | 3 | 3 | 0 | 2 | 2.66666667 |
| TRIP4 | Activating sig | 1.01752831 | 0.89906862 | 0.00022709 | 0.00022932 | 0.00018216 | 0.00027538 | 0.00022849 | 0.00017392 | 0.00023336 | 0.0002892 | 0.00023349 | 0.00023249 | 5 | 5 | 4 | 6 | 5 | 3 | 4 | 4 | 4 | 3.75 |
| MRE11 | Double-stran | 1.0112656 | 0.90810852 | 0.0018263 | 0.0013925 | 0.0016817 | 0.0016948 | 0.00164883 | 0.0016175 | 0.0017235 | 0.0013646 | 0.001964 | 0.0016674 | 49 | 37 | 45 | 45 | 44 | 34 | 36 | 23 | 41 | 33.5 |
| MCM3 | DNA replicati | 1.00969318 | 0.46085554 | 9.7977E-05 | 9.8935E-05 | 0.00013098 | 9.9006E-05 | 0.00010672 | 0.00008337 | 0 | 0.00015596 | 8.3947E-05 | 0.00010776 | 3 | 3 | 4 | 3 | 3.25 | 2 | 0 | 3 | 2 | 2.33333333 |
| HSP7C | Heat shock c | 1.00842796 | 0.9342085 | 0.0031454 | 0.0033824 | 0.0031128 | 0.0039214 | 0.0033905 | 0.0035976 | 0.0040402 | 0.0027311 | 0.0033074 | 0.00341908 | 85 | 92 | 83 | 107 | 91.75 | 75 | 82 | 46 | 70 | 68.25 |
| 43350 | Septin-7 OS= | 1.00742011 | 0.96635422 | 0.00036231 | 0.00054878 | 0.00066601 | 0.0006102 | 0.00054683 | 0.00038537 | 0.00054295 | 0.00057674 | 0.00069847 | 0.00055088 | 6 | 9 | 11 | 10 | 9 | 5 | 7 | 6 | 9 | 6.75 |
| SYYC | Tyrosine--tRN | 1.00655569 | 0.96823899 | 0.0004498 | 0.00055514 | 0.00030067 | 0.00050503 | 0.00045266 | 0.00051033 | 0.00032098 | 0.00047734 | 0.00051386 | 0.00045563 | 9 | 11 | 6 | 10 | 9 | 8 | 5 | 6 | 8 | 6.75 |
| SEC13 | Protein SEC1 | 1.00585234 | 0.48480655 | 0.00057366 | 0.00024826 | 0.00049302 | 0.00049687 | 0.00045295 | 0.0003138 | 0.00042106 | 0 | 0.00063195 | 0.0004556 | 7 | 3 | 6 | 6 | 5.5 | 3 | 4 | 0 | 6 | 4.33333333 |
| SPD2B | SH3 and PX d | 1.00331763 | 0.45726439 | 0.00011587 | 0.0001755 | 0.0002033 | 0.00014635 | 0.00016026 | 0.00022183 | 0.00011162 | 0 | 0.00014891 | 0.00016079 | 4 | 6 | 7 | 5 | 5.5 | 6 | 3 | 0 | 4 | 4.33333333 |
| SRRM2 | Serine/argini | 0.99933948 | 0.99840931 | 0.00048903 | 0.00040667 | 0.00031727 | 0.00035851 | 0.00039287 | 0.00046508 | 0.00046804 | 4.5792E-05 | 0.00059153 | 0.00039261 | 51 | 42 | 33 | 37 | 40.75 | 38 | 38 | 3 | 48 | 31.75 |
| EPN4 | Clathrin inter | 0.99598668 | 0.97596259 | 0.00042221 | 0.00034108 | 0.00029634 | 0.00025599 | 0.00032891 | 0.00032334 | 0.00037963 | 0.00033605 | 0.00027132 | 0.00032759 | 10 | 8 | 7 | 6 | 7.75 | 6 | 7 | 5 | 5 | 5.75 |
| DDX17 | Probable ATF | 0.99588364 | 0.98597958 | 0.00083256 | 0.00069449 | 0.00079848 | 0.0010974 | 0.00085573 | 0.00097025 | 0.0011159 | 0.00034573 | 0.00097696 | 0.00085221 | 25 | 23 | 23 | 31 | 25.5 | 25 | 28 | 8 | 25 | 21.5 |
| EF1A1 | Elongation fa | 0.99529642 | 0.94877953 | 0.0033699 | 0.0032876 | 0.0032644 | 0.0031745 | 0.0032741 | 0.0034994 | 0.0032282 | 0.0026368 | 0.0036704 | 0.0032587 | 59 | 57 | 57 | 55 | 57 | 48 | 44 | 29 | 50 | 42.75 |
| MYO1B | Unconventio | 0.99061145 | 0.96877806 | 0.00013938 | 9.3826E-05 | 0.00011646 | 0.00007042 | 0.00010502 | 5.9299E-05 | 8.9513E-05 | 0.00014791 | 0.00011942 | 0.00010404 | 6 | 4 | 5 | 3 | 4.5 | 2 | 3 | 4 | 4 | 3.25 |
| CSK22 | Casein kinase | 0.98886194 | 0.50913377 | 0.00030158 | 0.00053293 | 0.00030238 | 0.00068569 | 0.00045565 | 0.00067363 | 0.00038738 | 0 | 0.0002907 | 0.00045057 | 4 | 7 | 4 | 9 | 6 | 7 | 4 | 0 | 3 | 4.66666667 |
| EXOS9 | Exosome con | 0.9859846 | 0.42834184 | 0.00054099 | 0.00048559 | 0.00024108 | 0.00060742 | 0.00046877 | 0.00053706 | 0.00046327 | 0 | 0.00038627 | 0.0004622 | 9 | 8 | 4 | 10 | 7.75 | 7 | 6 | 0 | 5 | 6 |
| COBL | Protein cordo | 0.97959959 | 0.3957884 | 0.00014649 | 0.00016905 | 8.3929E-05 | 0.00014802 | 0.00013687 | 0.00016026 | 0.0001344 | 0 | 0.00010758 | 0.00013408 | 7 | 8 | 4 | 7 | 6.5 | 6 | 5 | 0 | 4 | 5 |
| TBCE | Tubulin-speci | 0.97899532 | 0.44034876 | 0.00015022 | 0.00015169 | 0.00025103 | 0.0001012 | 0.00016354 | 0.00012782 | 0 | 0.00015942 | 0.00019306 | 0.0001601 | 3 | 3 | 5 | 2 | 3.25 | 2 | 0 | 2 | 3 | 2.33333333 |
| KINH | Kinesin-1 hea | 0.97880378 | 0.90835463 | 0.00024662 | 0.00033204 | 0.00030223 | 0.00019383 | 0.00026868 | 0.00017488 | 0.00028158 | 0.00034896 | 0.00024652 | 0.00026299 | 9 | 12 | 11 | 7 | 9.75 | 5 | 8 | 8 | 7 | 7 |
| CNOT7 | CCR4-NOT tra | 0.97185046 | 0.44039316 | 0.00037036 | 0.00065448 | 0.00064986 | 0.00028069 | 0.00048885 | 0.00035454 | 0.00047573 | 0 | 0.00059499 | 0.00047509 | 4 | 7 | 7 | 3 | 5.25 | 3 | 4 | 0 | 5 | 4 |
| TBK1 | Serine/threo | 0.97131121 | 0.34747915 | 0.00018099 | 0.00018276 | 0.00021777 | 0.00018289 | 0.0001911 | 0.00018481 | 0.00023248 | 0 | 0.00013957 | 0.00018562 | 5 | 5 | 6 | 5 | 5.25 | 4 | 5 | 0 | 3 | 4 |
| AMOT | Angiomotin ( | 0.97119695 | 0.32150744 | 0.00075465 | 0.00076203 | 0.00095193 | 0.00078718 | 0.00081395 | 0.00083893 | 0.00081299 | 0 | 0.00071959 | 0.0007905 | 31 | 31 | 39 | 32 | 33.25 | 27 | 26 | 0 | 23 | 25.3333333 |
| REPS1 | RalBP1-assoc | 0.97062828 | 0.36411012 | 0.00019891 | 0.00020085 | 0.00013296 | 0.0001675 | 0.00017506 | 0.00021157 | 0.00012775 | 0 | 0.00017042 | 0.00016991 | 6 | 6 | 4 | 5 | 5.25 | 5 | 3 | 0 | 4 | 4 |
| SPAG5 | Sperm-associ | 0.96724997 | 0.87348278 | 0.0001106 | 0.00013401 | 0.0001996 | 0.00013411 | 0.00014458 | 0.00014116 | 0.00019888 | 0.00010563 | 0.00011371 | 0.00013985 | 5 | 6 | 9 | 6 | 6.5 | 5 | 7 | 3 | 4 | 4.75 |
| RSMN | Small nuclea | 0.96319841 | 0.77546276 | 0.00098957 | 0.00099924 | 0.00088196 | 0.00088885 | 0.00093991 | 0.00084204 | 0.00070616 | 0.0012252 | 0.00084786 | 0.00090532 | 9 | 9 | 8 | 8 | 8.5 | 6 | 5 | 7 | 6 | 6 |
| 43352 | Septin-9 OS= | 0.96233019 | 0.31939026 | 0.00031522 | 0.00036377 | 0.00031606 | 0.00036404 | 0.00033977 | 0.00040234 | 0.00028921 | 0 | 0.00028937 | 0.00032697 | 7 | 8 | 7 | 8 | 7.5 | 7 | 5 | 0 | 5 | 5.66666667 |
| TBCD5 | TBC1 domain | 0.96086067 | 0.32320228 | 0.00016596 | 0.00016759 | 0.00023297 | 0.00020125 | 0.00019194 | 0.00016947 | 0.00021318 | 0 | 0.00017064 | 0.00018443 | 5 | 5 | 7 | 6 | 5.75 | 4 | 5 | 0 | 4 | 4.33333333 |
| DDX5 | Probable ATF | 0.95881893 | 0.753873 | 0.00085956 | 0.00095476 | 0.00090494 | 0.00091202 | 0.00090782 | 0.00082284 | 0.00088327 | 0.00061573 | 0.0011599 | 0.00087044 | 30 | 31 | 32 | 30 | 30.75 | 22 | 24 | 15 | 28 | 22.25 |
| DDX52 | Probable ATF | 0.94810753 | 0.71228215 | 0.00066081 | 0.00066727 | 0.00044171 | 0.00048969 | 0.00056487 | 0.0005623 | 0.00039611 | 0.00056102 | 0.0006228 | 0.00053556 | 15 | 15 | 10 | 11 | 12.75 | 10 | 7 | 8 | 11 | 9 |
| CAMP2 | Calmodulin-r | 0.94331574 | 0.37739509 | 5.3167E-05 | 0.00010737 | 8.8847E-05 | 7.1634E-05 | 8.0255E-05 | 9.0481E-05 | 4.5528E-05 | 0 | 9.1107E-05 | 7.5705E-05 | 3 | 6 | 5 | 4 | 4.5 | 4 | 2 | 0 | 4 | 3.33333333 |

| | | | | | | | | | | | | | | | | | | | | | | | |
|---|---|---|---|---|---|---|---|---|---|---|---|---|---|---|---|---|---|---|---|---|---|---|---|
| CD2AP | CD2-associat | 0.94326901 | 0.80241611 | 0.00045426 | 0.0005421 | 0.0006625 | 0.00045903 | 0.00052947 | 0.0005271 | 0.00063654 | 0.00019721 | 0.00063689 | 0.00049944 | 11 | 13 | 16 | 11 | 12.75 | 10 | 12 | 3 | 12 | 9.25 |
| SNAPN | SNARE-assoc | 0.94178853 | 0.75808106 | 0.0013582 | 0.00097965 | 0.0015564 | 0.0011764 | 0.00126766 | 0.001486 | 0.0007477 | 0.0015443 | 0.00099748 | 0.00119387 | 7 | 5 | 8 | 6 | 6.5 | 6 | 3 | 5 | 4 | 4.5 |
| IRS4 | Insulin recep | 0.94005878 | 0.63347551 | 0.00079774 | 0.00078434 | 0.00082091 | 0.00070005 | 0.00077576 | 0.00080386 | 0.0008629 | 0.00046785 | 0.00078243 | 0.00072926 | 38 | 37 | 39 | 33 | 36.75 | 30 | 32 | 14 | 29 | 26.25 |
| MAP1B | Microtubule- | 0.93729815 | 0.30158911 | 0.0004063 | 0.00036709 | 0.00030018 | 0.00038896 | 0.00036563 | 0.00032754 | 0.00043949 | 0 | 0.00026109 | 0.00034271 | 38 | 34 | 28 | 36 | 34 | 24 | 32 | 0 | 19 | 25 |
| FERM2 | Fermitin fam | 0.93703729 | 0.76977225 | 0.00023284 | 0.00023512 | 0.00011673 | 0.00015686 | 0.00018539 | 0.0001486 | 0.00014954 | 0.0002471 | 0.00014962 | 0.00017372 | 6 | 6 | 3 | 4 | 4.75 | 3 | 3 | 4 | 3 | 3.25 |
| CYFP2 | Cytoplasmic | 0.93317153 | 0.84161517 | 0.00010324 | 0.00022935 | 0.00016563 | 0.00018779 | 0.0001715 | 7.9065E-05 | 0.00015913 | 0.00029582 | 0.00010615 | 0.00016004 | 5 | 11 | 8 | 9 | 8.25 | 3 | 6 | 9 | 4 | 5.5 |
| SYLC | Leucine--tRN | 0.92875971 | 0.56684475 | 0.00022439 | 0.00020393 | 0.00022499 | 0.00020407 | 0.00021435 | 0.00022913 | 0.00017294 | 0.00025004 | 0.00014419 | 0.00019908 | 10 | 9 | 10 | 9 | 9.5 | 8 | 6 | 7 | 5 | 6.5 |
| NKRF | NF-kappa-B-r | 0.92739259 | 0.44216904 | 0.0009561 | 0.00096545 | 0.00080526 | 0.0009275 | 0.00091358 | 0.0010251 | 0.00068774 | 0.00079142 | 0.00088472 | 0.00084725 | 25 | 25 | 21 | 24 | 23.75 | 21 | 14 | 13 | 18 | 16.5 |
| FBRL | rRNA 2'-O-m | 0.92697046 | 0.58808136 | 0.0011509 | 0.0007471 | 0.00082426 | 0.0010799 | 0.00095054 | 0.0010493 | 0.00073916 | 0.00078516 | 0.00095087 | 0.00088112 | 14 | 9 | 10 | 13 | 11.5 | 10 | 7 | 6 | 9 | 8 |
| TAF2 | Transcription | 0.92587402 | 0.31156692 | 0.00011004 | 0.00015557 | 8.8269E-05 | 0.00013344 | 0.00012183 | 0.00014046 | 0.00011308 | 0 | 8.4857E-05 | 0.0001128 | 5 | 7 | 4 | 6 | 5.5 | 5 | 4 | 0 | 3 | 4 |
| PKN2 | Serine/threo | 0.92372022 | 0.68733523 | 0.00026817 | 0.00018956 | 0.00021511 | 0.0001355 | 0.00020209 | 0.0002396 | 0.00020668 | 0.00012807 | 0.00017233 | 0.00018667 | 10 | 7 | 8 | 5 | 7.5 | 7 | 6 | 3 | 5 | 5.25 |
| CNN3 | Calponin-3 O | 0.92114099 | 0.33744996 | 0.00056146 | 0.00064794 | 0.00064337 | 0.00097261 | 0.00070635 | 0.00092138 | 0.00051513 | 0 | 0.00051542 | 0.00065064 | 7 | 8 | 8 | 12 | 8.75 | 9 | 5 | 0 | 5 | 6.33333333 |
| RS24 | 40S ribosoma | 0.91821616 | 0.65983723 | 0.0047618 | 0.0030052 | 0.0043766 | 0.0036089 | 0.00393813 | 0.0032922 | 0.0022937 | 0.0050534 | 0.0038249 | 0.00361605 | 24 | 15 | 22 | 18 | 19.75 | 13 | 9 | 16 | 15 | 13.25 |
| RBM39 | RNA-binding | 0.91261219 | 0.54683995 | 0.0010954 | 0.0011564 | 0.0011981 | 0.0012075 | 0.00116435 | 0.00095325 | 0.00076745 | 0.0015059 | 0.0010238 | 0.0010626 | 22 | 23 | 24 | 24 | 23.25 | 15 | 12 | 19 | 16 | 15.5 |
| AHNK | Neuroblast d | 0.91241159 | 0.76715156 | 0.00053762 | 0.00045693 | 0.00059745 | 0.00051611 | 0.00052703 | 0.00065762 | 0.00069057 | 5.7054E-05 | 0.00051822 | 0.00048087 | 115 | 97 | 128 | 107 | 111.75 | 112 | 118 | 8 | 88 | 81.5 |
| TAB2 | TGF-beta-act | 0.90624645 | 0.33672144 | 0.00011424 | 0.00019225 | 0.00011454 | 0.00015391 | 0.00014374 | 9.7205E-05 | 9.7823E-05 | 0 | 0.00019575 | 0.00013026 | 3 | 5 | 3 | 4 | 3.75 | 2 | 2 | 0 | 4 | 2.66666667 |
| PALLD | Palladin OS=I | 0.90570702 | 0.73534934 | 0.00049609 | 0.00069362 | 0.00053568 | 0.00061699 | 0.0005856 | 0.00065756 | 0.00075977 | 0.00009112 | 0.00061306 | 0.00053038 | 26 | 36 | 28 | 32 | 30.5 | 27 | 31 | 3 | 25 | 21.5 |
| RL7A | 60S ribosoma | 0.90467254 | 0.45470527 | 0.0049602 | 0.0032056 | 0.0038793 | 0.0051126 | 0.00428943 | 0.0037987 | 0.0033131 | 0.0039479 | 0.0044624 | 0.00388053 | 50 | 32 | 39 | 51 | 43 | 30 | 26 | 25 | 35 | 29 |
| CYBP | Calcyclin-bin | 0.90449479 | 0.72461537 | 0.00081017 | 0.00093496 | 0.00081233 | 0.00046782 | 0.00075632 | 0.00073863 | 0.00029733 | 0.0011054 | 0.00059499 | 0.00068409 | 7 | 8 | 7 | 4 | 6.5 | 5 | 2 | 6 | 4 | 4.25 |
| ARF1 | ADP-ribosyla | 0.90402581 | 0.4996134 | 0.0020411 | 0.0022083 | 0.0026313 | 0.0023572 | 0.00230948 | 0.0013026 | 0.00206 | 0.0025529 | 0.0024358 | 0.00208783 | 26 | 25 | 29 | 27 | 26.75 | 16 | 18 | 19 | 23 | 19 |
| DOCK7 | Dedicator of | 0.90025446 | 0.61098116 | 0.0001973 | 0.00016187 | 0.00017309 | 0.00018691 | 0.00017979 | 0.00022035 | 0.00014255 | 7.8516E-05 | 0.00020602 | 0.00016186 | 16 | 13 | 14 | 15 | 14.5 | 14 | 9 | 4 | 13 | 10 |
| TLN1 | Talin-1 OS=H | 0.89927253 | 0.49589123 | 0.00036348 | 0.00039849 | 0.00033321 | 0.00033581 | 0.00035775 | 0.00037115 | 0.00038684 | 0.00018184 | 0.00034702 | 0.00032171 | 35 | 38 | 32 | 32 | 34.25 | 28 | 29 | 11 | 26 | 23.5 |
| DHX37 | Probable ATP | 0.89416231 | 0.28506471 | 0.00009123 | 6.9092E-05 | 0.00013721 | 0.00013828 | 0.00010895 | 0.00011644 | 8.7888E-05 | 0 | 8.7937E-05 | 9.7422E-05 | 4 | 3 | 6 | 6 | 4.75 | 4 | 3 | 0 | 3 | 3.33333333 |
| H4 | Histone H4 O | 0.89093296 | 0.20499475 | 0.0040992 | 0.0038806 | 0.0033394 | 0.0044011 | 0.00393008 | 0 | 0.003949 | 0.0032626 | 0.0032927 | 0.00350143 | 16 | 15 | 13 | 17 | 15.25 | 0 | 12 | 8 | 10 | 10 |
| BAG2 | BAG family m | 0.89032234 | 0.34024193 | 0.00037519 | 0.00050515 | 0.00050159 | 0.00050551 | 0.00047186 | 0.00047888 | 0.00048193 | 0.00039816 | 0.00032146 | 0.00042011 | 3 | 4 | 4 | 4 | 3.75 | 3 | 3 | 2 | 2 | 2.5 |
| NUP88 | Nuclear pore | 0.88708149 | 0.28419108 | 0.0008903 | 0.0010069 | 0.00082125 | 0.0010076 | 0.00093151 | 0.00072727 | 0.0010521 | 0.00079364 | 0.0007323 | 0.00082633 | 25 | 28 | 23 | 28 | 26 | 16 | 23 | 14 | 16 | 17.25 |
| SYFB | Phenylalanin | 0.88286561 | 0.32694878 | 0.00035842 | 0.00036192 | 0.00013476 | 0.00027164 | 0.00028169 | 0.00034311 | 0.00017264 | 0 | 0.00023032 | 0.00024869 | 8 | 8 | 3 | 6 | 6.25 | 6 | 3 | 0 | 4 | 4.33333333 |
| PDLI1 | PDZ and LIM | 0.88007523 | 0.14616942 | 0.0010427 | 0.00080992 | 0.00088464 | 0.0010537 | 0.00094774 | 0.00092138 | 0.00082421 | 0.00076607 | 0.00082467 | 0.00083408 | 13 | 10 | 11 | 13 | 11.75 | 9 | 8 | 6 | 8 | 7.75 |
| CDC27 | Cell division | 0.87576798 | 0.57824837 | 0.00035227 | 0.00022637 | 0.00028899 | 0.00029125 | 0.00028972 | 0.00032701 | 0.00016454 | 0.00015294 | 0.00037042 | 0.00025373 | 11 | 7 | 9 | 9 | 9 | 8 | 4 | 3 | 9 | 6 |
| ERBIN | Erbin OS=Hor | 0.8688548 | 0.17647726 | 0.00044853 | 0.00043404 | 0.00035603 | 0.00041547 | 0.00041352 | 0.00040551 | 0.00031207 | 0 | 0.00036028 | 0.00035929 | 24 | 23 | 19 | 22 | 22 | 17 | 13 | 0 | 15 | 15 |
| RPGF6 | Rap guanine | 0.8657522 | 0.60522552 | 0.00029668 | 0.00028294 | 0.0002479 | 0.00028315 | 0.00027767 | 0.00023142 | 0.00033874 | 5.2475E-05 | 0.00033893 | 0.00024039 | 18 | 17 | 15 | 17 | 16.75 | 11 | 16 | 2 | 16 | 11.25 |
| ZO2 | Tight junctior | 0.85952639 | 0.5930294 | 0.0013083 | 0.0013435 | 0.0009783 | 0.0013445 | 0.00124365 | 0.0015567 | 0.0011678 | 0.0002118 | 0.0013395 | 0.00106895 | 60 | 62 | 44 | 60 | 56.5 | 55 | 42 | 6 | 47 | 37.5 |
| DDX21 | Nucleolar RN | 0.85786839 | 0.07104671 | 0.0018873 | 0.0018377 | 0.001453 | 0.0016347 | 0.00170318 | 0.0015916 | 0.0013853 | 0.0013948 | 0.0014727 | 0.0014611 | 56 | 54 | 43 | 48 | 50.25 | 37 | 32 | 26 | 34 | 32.25 |
| DJC11 | DnaJ homolo | 0.8537378 | 0.52022704 | 0.00033045 | 0.00023834 | 0.000142 | 0.00019081 | 0.0002254 | 0.00024101 | 0.00012127 | 0.00022544 | 0.00018201 | 0.00019243 | 7 | 5 | 3 | 4 | 4.75 | 4 | 2 | 3 | 3 | 3 |
| SRPK1 | SRSF protein | 0.85090776 | 0.24555953 | 0.00024173 | 0.00012204 | 0.00016158 | 0.00020355 | 0.00018223 | 0.00010284 | 0.000207 | 0 | 0.00015533 | 0.00015506 | 6 | 3 | 4 | 5 | 4.5 | 2 | 4 | 0 | 3 | 3 |
| SRSF1 | Serine/argini | 0.85084743 | 0.22272346 | 0.0025537 | 0.0013968 | 0.001707 | 0.0018279 | 0.00187135 | 0.0012223 | 0.0020501 | 0 | 0.0015043 | 0.00159223 | 24 | 13 | 16 | 17 | 17.5 | 9 | 15 | 0 | 11 | 11.6666667 |
| RICTR | Rapamycin-ir | 0.84996033 | 0.17234457 | 0.00010815 | 0.00014041 | 0.00010844 | 0.00010929 | 0.00011657 | 0.00007888 | 9.9226E-05 | 0 | 0.00011914 | 9.9082E-05 | 7 | 9 | 7 | 7 | 7.5 | 4 | 5 | 0 | 6 | 5 |
| IF4H | Eukaryotic tr | 0.8484683 | 0.16579766 | 0.00031921 | 0.00032234 | 0.00032006 | 0.00043009 | 0.00034793 | 0 | 0.00027335 | 0.00033876 | 0.0002735 | 0.0002952 | 3 | 3 | 3 | 4 | 3.25 | 0 | 2 | 2 | 2 | 2 |
| CND2 | Condensin cc | 0.84663497 | 0.34560176 | 0.00078346 | 0.00086304 | 0.00074984 | 0.00086366 | 0.000815 | 0.00077272 | 0.00086912 | 0.00034013 | 0.00077806 | 0.00069001 | 22 | 24 | 21 | 24 | 22.75 | 17 | 19 | 6 | 17 | 14.75 |
| CH60 | 60 kDa heat | 0.84409021 | 0.35105198 | 0.00082896 | 0.00055804 | 0.00092351 | 0.00074459 | 0.00076378 | 0.00082294 | 0.00047324 | 0.00051316 | 0.00076944 | 0.0006447 | 18 | 12 | 20 | 16 | 16.5 | 14 | 8 | 7 | 13 | 10.5 |
| H2B1L | Histone H2B | 0.83832473 | 0.27533762 | 0.0037698 | 0.0027492 | 0.0035698 | 0.0025396 | 0.0031571 | 0.0018712 | 0.0026901 | 0.0033338 | 0.0026916 | 0.00264668 | 18 | 13 | 17 | 12 | 15 | 7 | 10 | 10 | 10 | 9.25 |
| SPART | Spartin OS=H | 0.83503045 | 0.22868243 | 0.00067358 | 0.00080019 | 0.00083428 | 0.00056054 | 0.00071715 | 0.00075859 | 0.00045805 | 0.00056765 | 0.00061107 | 0.00059884 | 17 | 20 | 21 | 14 | 18 | 15 | 9 | 9 | 12 | 11.25 |
| DRG1 | Developmen | 0.83366307 | 0.52516679 | 0.00050332 | 0.00029042 | 0.00050466 | 0.00021798 | 0.0003791 | 0.00045888 | 0.00027708 | 0.00034337 | 0.00018482 | 0.00031604 | 7 | 4 | 7 | 3 | 5.25 | 5 | 3 | 3 | 2 | 3.25 |
| TRI25 | E3 ubiquitin/ | 0.83018103 | 0.39720327 | 0.00033509 | 0.00042296 | 0.00016799 | 0.00038094 | 0.00032675 | 0.00026731 | 0.00026901 | 0.00033338 | 0.00021533 | 0.00027126 | 8 | 10 | 4 | 9 | 7.75 | 5 | 5 | 5 | 4 | 4.75 |
| OAT | Ornithine am | 0.82833914 | 0.30033522 | 0 | 0 | 0.00030135 | 0 | 0.00030135 | 0.00023017 | 0.00023163 | 0.00028706 | 0 | 0.00024962 | 0 | 0 | 5 | 0 | 5 | 3 | 3 | 3 | 0 | 3 |
| AP2M1 | AP-2 comple: | 0.82814427 | 0.20966993 | 0.00072796 | 0.0009801 | 0.00066907 | 0.0008582 | 0.00080883 | 0.00069686 | 0.00054545 | 0.0005794 | 0.00085761 | 0.00066983 | 12 | 16 | 11 | 14 | 13.25 | 9 | 7 | 6 | 11 | 8.25 |
| NXF1 | Nuclear RNA | 0.81983763 | 0.30359629 | 0.00042631 | 0.00051657 | 0.00047019 | 0.00030155 | 0.00042866 | 0.0004353 | 0.00038331 | 0.00020358 | 0.00038352 | 0.00035143 | 10 | 12 | 11 | 7 | 10 | 8 | 7 | 3 | 7 | 6.25 |
| PLEC | Plectin OS=H | 0.81774656 | 0.51036622 | 0.00014648 | 5.6888E-05 | 0.00014687 | 9.1087E-05 | 0.00011033 | 0.00010786 | 0.00010131 | 3.5872E-05 | 0.00011585 | 9.0223E-05 | 26 | 10 | 26 | 16 | 19.5 | 15 | 14 | 4 | 16 | 12.25 |
| GIT1 | ARF GTPase-: | 0.80959066 | 0.194315 | 0.00017338 | 0.00021009 | 0.00020861 | 0.00014016 | 0.00018306 | 0.00013278 | 0.0002227 | 0 | 8.9131E-05 | 0.0001482 | 5 | 6 | 6 | 4 | 5.25 | 3 | 5 | 0 | 2 | 3.33333333 |
| PICAL | Phosphatidyl | 0.80847829 | 0.23568323 | 8.0946E-05 | 0.00028608 | 0.00024348 | 0.00024539 | 0.00021397 | 0.00015498 | 0.00020795 | 0 | 0.00015605 | 0.00017299 | 2 | 7 | 6 | 6 | 5.25 | 3 | 4 | 0 | 3 | 3.33333333 |
| BCAS3 | Breast carcin | 0.80782029 | 0.208021 | 0.00034123 | 0.00031585 | 0.00028512 | 0.00022988 | 0.00029302 | 0.00021777 | 0.00032873 | 0.00018106 | 0.00021927 | 0.00023671 | 12 | 11 | 10 | 8 | 10.25 | 6 | 9 | 4 | 6 | 6.25 |
| TB182 | 182 kDa tank | 0.80323584 | 0.13889349 | 0.00045787 | 0.00061646 | 0.00045909 | 0.00041641 | 0.00048746 | 0.00029221 | 0.0004705 | 0 | 0.00041192 | 0.00039154 | 30 | 40 | 30 | 27 | 31.75 | 15 | 24 | 0 | 21 | 20 |
| RRP44 | Exosome con | 0.80196241 | 0.16023302 | 0.000303 | 0.00036159 | 0.00030381 | 0.00025051 | 0.00030473 | 0.00028127 | 0.00017691 | 0.00030693 | 0.00021241 | 0.00024438 | 11 | 13 | 11 | 9 | 11 | 8 | 5 | 7 | 6 | 6.5 |
| MSH6 | DNA mismat | 0.80084495 | 0.20996857 | 0.00019403 | 0.00035267 | 0.00023346 | 0.00023528 | 0.00025386 | 0.00019813 | 0.00022431 | 0.00021621 | 0.00017456 | 0.0002033 | 10 | 18 | 12 | 12 | 13 | 8 | 9 | 7 | 7 | 7.75 |
| RTN4 | Reticulon-4 C | 0.79976798 | 0.16000353 | 8.8552E-05 | 0.00013413 | 6.6591E-05 | 0.00011185 | 0.00010028 | 8.4769E-05 | 0 | 0.00007048 | 8.5355E-05 | 8.0201E-05 | 4 | 6 | 3 | 5 | 4.5 | 3 | 0 | 2 | 3 | 2.66666667 |
| RIR2 | Ribonucleosi | 0.79973096 | 0.14639185 | 0.00027135 | 0.0002055 | 0.00034009 | 0.00034275 | 0.00028992 | 0.00025976 | 0.00017427 | 0 | 0.00026155 | 0.00023186 | 4 | 3 | 5 | 5 | 4.25 | 3 | 2 | 0 | 3 | 2.66666667 |
| ATD3B | ATPase famil | 0.79932994 | 0.11890023 | 0.00016289 | 0.00016448 | 0.00016333 | 0.00020575 | 0.00017411 | 0.00015593 | 0.00015692 | 0 | 0.00010467 | 0.00013917 | 4 | 4 | 4 | 5 | 4.25 | 3 | 3 | 0 | 2 | 2.66666667 |
| NU160 | Nuclear pore | 0.79500433 | 0.37265797 | 0.00033077 | 0.00020412 | 0.00018425 | 0.00020426 | 0.00023085 | 0.0002111 | 9.4417E-05 | 0.00026327 | 0.00016532 | 0.00018353 | 18 | 11 | 10 | 11 | 12.5 | 9 | 4 | 9 | 7 | 7.25 |
| ZYX | Zyxin OS=Hor | 0.79323659 | 0.32108673 | 0.00032294 | 0.00046585 | 0.00018503 | 0.00041956 | 0.00034835 | 0.00011777 | 0.00053332 | 0 | 0.00017787 | 0.00027632 | 7 | 10 | 4 | 9 | 7.5 | 2 | 9 | 0 | 3 | 4.66666667 |
| NU133 | Nuclear pore | 0.78640847 | 0.30670598 | 0.00029676 | 0.00036881 | 0.00043487 | 0.00041521 | 0.00037891 | 0.00034964 | 0.00032254 | 0.00010901 | 0.00041073 | 0.00029798 | 13 | 16 | 19 | 18 | 16.5 | 12 | 11 | 3 | 14 | 10 |
| LMNB1 | Lamin-B1 OS | 0.78143525 | 0.23480297 | 0.00049535 | 0.00036377 | 0.00058697 | 0.00045504 | 0.00047528 | 0.00034486 | 0.00052058 | 0.00021505 | 0.00040512 | 0.0003714 | 11 | 8 | 13 | 10 | 10.5 | 6 | 9 | 3 | 7 | 6.25 |
| HNRPU | Heterogeneo | 0.77736456 | 0.17249194 | 0.00095958 | 0.0010659 | 0.00080178 | 0.00087269 | 0.00092499 | 0.0010207 | 0.00078063 | 0.00045825 | 0.00061663 | 0.00071905 | 30 | 33 | 25 | 27 | 28.75 | 25 | 19 | 9 | 15 | 17 |
| DHX9 | ATP-depende | 0.77655657 | 0.01637659 | 0.0023479 | 0.0023499 | 0.0024375 | 0.0018267 | 0.0022405 | 0.0018565 | 0.0016014 | 0.0018192 | 0.0016824 | 0.00173988 | 113 | 112 | 117 | 87 | 107.25 | 70 | 60 | 55 | 63 | 62 |
| EMAL4 | Echinoderm | 0.77317968 | 0.19598035 | 0.0001076 | 0.00019014 | 0.0001888 | 0.00010873 | 0.00014882 | 6.8668E-05 | 0.00010366 | 0 | 0.00017286 | 0.00011506 | 4 | 7 | 7 | 4 | 5.5 | 2 | 3 | 0 | 5 | 3.33333333 |
| DHX29 | ATP-depende | 0.77158718 | 0.00131688 | 0.0030841 | 0.0032894 | 0.0029377 | 0.003136 | 0.0031118 | 0.0026079 | 0.0024512 | 0.0021172 | 0.0024278 | 0.00240103 | 160 | 169 | 152 | 161 | 160.5 | 106 | 99 | 69 | 98 | 93 |
| CCD43 | Coiled-coil dc | 0.77030224 | 0.28881055 | 0.00047122 | 0.00047583 | 0.00047248 | 0.0008333 | 0.00056321 | 0.00030073 | 0.00045396 | 0.00037506 | 0.00060561 | 0.00043384 | 4 | 4 | 4 | 7 | 4.75 | 2 | 3 | 2 | 4 | 2.75 |
| FLNB | Filamin-B OS | 0.76807554 | 0.0744104 | 0.00016227 | 0.00019457 | 0.00017287 | 0.00013323 | 0.00016574 | 0.00014239 | 0.00013027 | 0.00014529 | 9.1238E-05 | 0.0001273 | 74 | 67 | 72 | 80 | 73.25 | 49 | 38 | 37 | 42 | 41.5 |
| EXOS2 | Exosome con | 0.76506353 | 0.07318507 | 0.001441 | 0.0013642 | 0.0014448 | 0.0011831 | 0.00135828 | 0.0010346 | 0.0010412 | 0 | 0.0010417 | 0.00103917 | 16 | 15 | 16 | 13 | 15 | 9 | 9 | 0 | 9 | 9 |
| AP3D1 | AP-3 comple: | 0.7590201 | 0.13075489 | 0.00057217 | 0.00041599 | 0.0004819 | 0.00062443 | 0.00052362 | 0.00037976 | 0.00055856 | 0.00032789 | 0.00032355 | 0.00039744 | 25 | 18 | 21 | 27 | 22.75 | 13 | 19 | 9 | 11 | 13 |
| ADT3 | ADP/ATP trar | 0.75819634 | 0.05871728 | 0.0046047 | 0.0031296 | 0.0039067 | 0.0033108 | 0.00373795 | 0.0031647 | 0.0029573 | 0.0022554 | 0.002959 | 0.0028341 | 89 | 76 | 76 | 79 | 80 | 93 | 99 | 84 | 100 | 94 |
| P4HA1 | Prolyl 4-hydr | 0.75591616 | 0.10435125 | 0.00034592 | 0.0002495 | 0.00044593 | 0.00029961 | 0.00033524 | 0.0002523 | 0.0002539 | 0 | 0.00025404 | 0.00025341 | 7 | 5 | 9 | 6 | 6.75 | 4 | 4 | 0 | 4 | 4 |
| CND3 | Condensin cc | 0.75582151 | 0.13698486 | 0.00025998 | 0.00023627 | 0.00023461 | 0.00031526 | 0.00026153 | 0.00023229 | 0.00026716 | 0.00012416 | 0.00016707 | 0.00019767 | 10 | 9 | 9 | 12 | 10 | 7 | 8 | 3 | 5 | 5.75 |
| GRP78 | 78 kDa gluco | 0.74923365 | 0.12084049 | 0.00080699 | 0.00073339 | 0.00089005 | 0.00089701 | 0.00083186 | 0.00082401 | 0.00072559 | 0.00032115 | 0.00062228 | 0.00062326 | 28 | 25 | 28 | 31 | 28 | 21 | 22 | 11 | 17 | 17.75 |
| PRS8 | 26S proteaso | 0.74348168 | 0.09797707 | 0.00025998 | 0.00026253 | 0.00019551 | 0.00032839 | 0.0002616 | 0.00016592 | 0.00016697 | 0 | 0.0002506 | 0.0001945 | 4 | 4 | 3 | 5 | 4 | 2 | 2 | 0 | 3 | 2.33333333 |
| TSNAX | Translin-asso | 0.74187806 | 0.58323218 | 0.00054597 | 0 | 0.00036495 | 0.00045975 | 0.00045689 | 0.00034843 | 0 | 0.00043455 | 0.00023389 | 0.00033896 | 6 | 0 | 4 | 5 | 5 | 3 | 0 | 3 | 2 | 2.66666667 |
| STRN3 | Striatin-3 OS | 0.73788246 | 0.16629678 | 0.00023177 | 0.00023403 | 0.00026558 | 0.00020074 | 0.00023303 | 0.00016904 | 0.00017012 | 0.00026353 | 8.5105E-05 | 0.00017195 | 7 | 7 | 8 | 6 | 7 | 4 | 4 | 5 | 2 | 3.75 |
| AP2A2 | AP-2 comple: | 0.73615599 | 0.16138091 | 0.00028103 | 0.00034053 | 0.00036631 | 0.00028398 | 0.00031796 | 0.00014348 | 0.00025268 | 0.00017894 | 0.00036118 | 0.00023407 | 17 | 20 | 18 | 22 | 19.25 | 7 | 8 | 5 | 14 | 8.5 |
| PSMD2 | 26S proteaso | 0.73590262 | 0.24222326 | 0.0002325 | 0.00029346 | 0.00023312 | 0.0001762 | 0.00023382 | 0.00025966 | 0.00011199 | 9.2525E-05 | 0.0002241 | 0.00017207 | 8 | 10 | 8 | 6 | 8 | 7 | 3 | 2 | 6 | 4.5 |
| SYDC | Aspartate--tF | 0.73228949 | 0.30763287 | 0.00026336 | 0.00021275 | 0.00042249 | 0.00015967 | 0.00026457 | 0.00026891 | 0.00020297 | 0.00016769 | 0.00013539 | 0.00019374 | 5 | 4 | 8 | 3 | 5 | 4 | 3 | 2 | 2 | 2.75 |
| STMN1 | Stathmin OS | 0.73034256 | 0.30914885 | 0.0010626 | 0.0017884 | 0.0007103 | 0.0017896 | 0.00133773 | 0.00067815 | 0.0011374 | 0.0014096 | 0.00068284 | 0.000977 | 6 | 10 | 4 | 10 | 7.5 | 3 | 5 | 5 | 3 | 4 |
| F1142 | Protein FAM | 0.72943142 | 0.16904178 | 0.00041803 | 0.00042212 | 0.00073351 | 0.00058084 | 0.00053863 | 0.00026678 | 0.0003356 | 0.00049908 | 0.0004701 | 0.00039289 | 8 | 8 | 14 | 11 | 10.25 | 4 | 5 | 6 | 7 | 5.5 |
| SSFA2 | Sperm-specif | 0.72873956 | 0.16106255 | 0.00018864 | 0.00019048 | 8.4063E-05 | 0.00012708 | 0.00014757 | 8.0258E-05 | 0.00016154 | 0 | 8.0813E-05 | 0.00010754 | 9 | 9 | 4 | 6 | 7 | 3 | 6 | 0 | 3 | 4 |
| NUP62 | Nuclear pore | 0.72827343 | 0.14171342 | 0.00070774 | 0.0008678 | 0.00086168 | 0.0009195 | 0.00083918 | 0.00077429 | 0.00077921 | 0.00024141 | 0.0006497 | 0.00061115 | 14 | 17 | 17 | 18 | 16.5 | 12 | 12 | 3 | 10 | 9.25 |
| H12 | Histone H1.2 | 0.72719054 | 0.01433658 | 0.0027256 | 0.0030024 | 0.0033539 | 0.0027542 | 0.00295903 | 0.0018976 | 0.002387 | 0.0017749 | 0.0025476 | 0.00215178 | 22 | 24 | 27 | 22 | 23.75 | 12 | 15 | 9 | 16 | 13 |
| SYSC | Serine--tRNA | 0.72364875 | 0.13035126 | 0.00046205 | 0.00051841 | 0.00077214 | 0.00067442 | 0.00060676 | 0.0004587 | 0.00046161 | 0.00057207 | 0.00026393 | 0.00043908 | 9 | 10 | 15 | 13 | 11.75 | 7 | 7 | 7 | 4 | 6.25 |
| DDX18 | ATP-depende | 0.71922617 | 0.18637005 | 0.00078771 | 0.00075565 | 0.00063185 | 0.00083579 | 0.00075275 | 0.00090488 | 0.00025295 | 0.00050157 | 0.00050619 | 0.0005414 | 20 | 19 | 16 | 21 | 19 | 18 | 5 | 8 | 10 | 10.25 |
| ATPG | ATP synthase | 0.71825042 | 0.04840226 | 0.0018596 | 0.0019672 | 0.0019533 | 0.0011633 | 0.00173585 | 0.0013563 | 0.0012512 | 0.0011277 | 0.0012519 | 0.00124678 | 21 | 22 | 22 | 13 | 19.5 | 12 | 11 | 8 | 11 | 10.5 |
| EF2 | Elongation fa | 0.71135586 | 0.00577522 | 0.0011072 | 0.0011491 | 0.00095597 | 0.0010878 | 0.00107502 | 0.00078512 | 0.00067159 | 0.00093021 | 0.00067196 | 0.00076472 | 36 | 37 | 31 | 35 | 34.75 | 20 | 17 | 19 | 17 | 18.25 |
| KNL1 | Kinetochore | 0.7097843 | 0.04562432 | 0.0004507 | 0.00051199 | 0.00039541 | 0.00043266 | 0.00044769 | 0.00033078 | 0.00030393 | 0 | 0.00031858 | 0.00031776 | 40 | 45 | 35 | 38 | 39.5 | 23 | 21 | 0 | 22 | 22 |
| TAB1 | TGF-beta-act | 0.70261195 | 0.05704165 | 0.0006283 | 0.00063444 | 0.00057747 | 0.00058199 | 0.00060555 | 0.00026731 | 0.00047077 | 0 | 0.00053832 | 0.00042547 | 12 | 12 | 11 | 11 | 11.5 | 4 | 7 | 0 | 8 | 6.33333333 |
| NSUN2 | tRNA (cytosir | 0.69635704 | 0.12979202 | 0.00034405 | 0.00041689 | 0.00041396 | 0.0003129 | 0.00037195 | 0.00026348 | 0.00039773 | 0.00010953 | 0.0002653 | 0.00025901 | 10 | 12 | 12 | 9 | 10.75 | 6 | 9 | 2 | 6 | 5.75 |
| NU214 | Nuclear pore | 0.69295908 | 0.18400779 | 0.00084594 | 0.00085422 | 0.00074692 | 0.00089311 | 0.00083505 | 0.00067685 | 0.00076225 | 8.0395E-05 | 0.00079512 | 0.00057865 | 67 | 67 | 59 | 70 | 65.75 | 42 | 47 | 4 | 49 | 35.5 |
| WDR3 | WD repeat-c | 0.68837289 | 0.0778678 | 0.00025185 | 0.00014129 | 0.00022446 | 0.0002545 | 0.00021803 | 0.00010715 | 0.00017972 | 0.00013364 | 0.00017982 | 0.00015008 | 9 | 5 | 8 | 9 | 7.75 | 3 | 5 | 3 | 5 | 4 |
| NU107 | Nuclear pore | 0.68637438 | 0.01442879 | 0.00054203 | 0.00046091 | 0.00062929 | 0.00057655 | 0.0005522 | 0.00040054 | 0.00043973 | 0.00027247 | 0.00040331 | 0.00037901 | 19 | 16 | 22 | 20 | 19.25 | 11 | 12 | 6 | 11 | 10 |
| RPP30 | Ribonuclease | 0.68440707 | 0.12519535 | 0.00049232 | 0.00049714 | 0.00069109 | 0.00089549 | 0.00064401 | 0.00037703 | 0.00037943 | 0.00062696 | 0.00037964 | 0.00044077 | 5 | 5 | 7 | 9 | 6.5 | 3 | 3 | 4 | 3 | 3.25 |
| SMC4 | Structural ma | 0.68190572 | 0.02658371 | 0.00034829 | 0.0003517 | 0.00028759 | 0.00028984 | 0.00031936 | 0.00023535 | 0.00018422 | 0.00029352 | 0.00015799 | 0.00021777 | 17 | 17 | 14 | 14 | 15.5 | 9 | 7 | 9 | 6 | 7.75 |
| PTN13 | Tyrosine-pro | 0.68044216 | 0.06445841 | 0.00019114 | 9.6506E-05 | 0.00014906 | 0.00015023 | 0.00014673 | 9.4878E-05 | 0.00010912 | 0 | 9.5534E-05 | 9.9844E-05 | 18 | 9 | 14 | 14 | 13.75 | 7 | 8 | 0 | 7 | 7.33333333 |
| LIMA1 | LIM domain | 0.67995118 | 0.05421506 | 0.00031291 | 0.00031597 | 0.00020916 | 0.0002108 | 0.00026221 | 0.00017751 | 0.00017863 | 0 | 0.00017873 | 0.00017829 | 9 | 9 | 6 | 6 | 7.5 | 4 | 4 | 0 | 4 | 4 |
| PIHD1 | PIH1 domain | 0.67764637 | 0.0074137 | 0.00081895 | 0.00082696 | 0.00082113 | 0.0010115 | 0.00086964 | 0.00058072 | 0.00046753 | 0.00072424 | 0.00058473 | 0.00058931 | 9 | 9 | 9 | 11 | 9.5 | 5 | 4 | 5 | 5 | 4.75 |
| PTSS1 | Phosphatidyl | 0.67747986 | 0.79875947 | 0.00027895 | 0.00016901 | 0.00022375 | 0 | 0.0002239 | 0.00014242 | 0.00014332 | 0.00017762 | 0.0001434 | 0.00015169 | 5 | 3 | 4 | 0 | 4 | 2 | 2 | 2 | 2 | 2 |
| RBM15 | Putative RNA | 0.67408049 | 0.1610214 | 0.00035113 | 0.00024546 | 0.00013541 | 0.00019105 | 0.00023076 | 0.0001379 | 0.00013877 | 0.00017198 | 0.00017356 | 0.00015555 | 13 | 9 | 5 | 7 | 8.5 | 4 | 4 | 4 | 5 | 4.25 |
| SRSF7 | Serine/argini | 0.67309881 | 0.06134896 | 0.00088701 | 0.0005598 | 0.00055585 | 0.00089632 | 0.00072475 | 0.00042456 | 0.00042726 | 0.00052949 | 0.00056999 | 0.00048783 | 8 | 5 | 5 | 8 | 6.5 | 3 | 3 | 3 | 4 | 3.25 |
| M3K7 | Mitogen-acti | 0.6712655 | 0.04051891 | 0.00034836 | 0.00043971 | 0.00043661 | 0.00039602 | 0.00040518 | 0.0002779 | 0.0003356 | 0.00013863 | 0.00033579 | 0.00027198 | 8 | 10 | 10 | 9 | 9.25 | 5 | 6 | 2 | 6 | 4.75 |
| AP2B1 | AP-2 comple: | 0.66534102 | 0.05391233 | 0.00047877 | 0.00068251 | 0.00042356 | 0.00051225 | 0.00052427 | 0.00043135 | 0.00036175 | 0 | 0.00025336 | 0.00034882 | 17 | 24 | 15 | 18 | 18.5 | 12 | 10 | 0 | 7 | 9.66666667 |
| ABCE1 | ATP-binding | 0.66523426 | 0.05804095 | 0.00044054 | 0.00035588 | 0.00035337 | 0.00031162 | 0.00036535 | 0.00016869 | 0.00016976 | 0.00035064 | 0.00028309 | 0.00024305 | 10 | 8 | 8 | 7 | 8.25 | 3 | 3 | 5 | 5 | 4 |
| CTND1 | Catenin delta | 0.66497015 | 0.09866484 | 0.00035439 | 0.00046797 | 0.00021867 | 0.00022038 | 0.00031535 | 0.00020877 | 0.00017508 | 0 | 0.00024525 | 0.0002097 | 13 | 17 | 8 | 8 | 11.5 | 6 | 5 | 0 | 7 | 6 |
| WAC2A | WASH compl | 0.65894308 | 0.03754658 | 0.00076745 | 0.00073521 | 0.00076949 | 0.00059655 | 0.00071718 | 0.00055257 | 0.00063191 | 0.0002506 | 0.00045523 | 0.00047258 | 39 | 37 | 39 | 30 | 36.25 | 22 | 25 | 8 | 18 | 18.25 |
| EMD | Emerin OS=H | 0.65772387 | 0.07713293 | 0.0012467 | 0.00094417 | 0.0010417 | 0.0013648 | 0.00114934 | 0.00039781 | 0.00080068 | 0.0011577 | 0.00066761 | 0.00075595 | 12 | 9 | 10 | 13 | 11 | 3 | 6 | 7 | 5 | 5.25 |
| IF5A1 | Eukaryotic tr | 0.65518706 | 0.0328543 | 0.0042838 | 0.0051909 | 0.0044671 | 0.0053678 | 0.0048274 | 0.0039368 | 0.0028613 | 0.0040915 | 0.0017618 | 0.00316285 | 25 | 30 | 26 | 31 | 28 | 18 | 13 | 15 | 8 | 13.5 |
| PP1G | Serine/threo | 0.65237166 | 0.00136457 | 0.0048202 | 0.0043723 | 0.0037681 | 0.0048708 | 0.00445785 | 0.0031283 | 0.0030433 | 0.0027311 | 0.00273 | 0.00290818 | 59 | 53 | 46 | 59 | 54.25 | 30 | 29 | 21 | 26 | 26.5 |
| ESYT2 | Extended syr | 0.65223926 | 0.02377554 | 0.00025787 | 0.00026039 | 0.00022983 | 0.00037639 | 0.00028112 | 0.00018285 | 0.00018402 | 0.00018244 | 0.00018412 | 0.00018336 | 9 | 9 | 8 | 13 | 9.75 | 5 | 5 | 4 | 5 | 4.75 |
| MRCKA | Serine/threo | 0.64926874 | 0.13852498 | 0.00028948 | 0.0002 | 0.00019859 | 0.00016935 | 0.00021436 | 0.00011668 | 0.0001957 | 4.8506E-05 | 0.00019581 | 0.00013917 | 19 | 13 | 13 | 11 | 14 | 6 | 10 | 2 | 10 | 7 |
| GEMI5 | Gem-associat | 0.64650106 | 0.00075604 | 0.00082245 | 0.00091884 | 0.00082464 | 0.00076036 | 0.00083157 | 0.00044671 | 0.00060688 | 0.00055711 | 0.00053975 | 0.00053761 | 47 | 52 | 47 | 43 | 47.25 | 20 | 27 | 20 | 24 | 22.75 |
| MAP4 | Microtubule- | 0.64582213 | 0.03254721 | 0.0017867 | 0.0017579 | 0.0015159 | 0.0015972 | 0.00166443 | 0.001111 | 0.0014712 | 0.00051049 | 0.001207 | 0.00107492 | 78 | 76 | 66 | 69 | 72.25 | 38 | 50 | 14 | 41 | 35.75 |
| FUS | RNA-binding | 0.64374949 | 0.07639441 | 0.00030101 | 0.00030395 | 0.00045272 | 0.00040556 | 0.00036581 | 0.0003842 | 0.00019332 | 0 | 0.00012895 | 0.00023549 | 6 | 6 | 9 | 8 | 7.25 | 6 | 3 | 0 | 2 | 3.66666667 |
| AFAD | Afadin OS=H | 0.63838091 | 0.08935148 | 0.00091144 | 0.00092036 | 0.00071079 | 0.00086254 | 0.00085128 | 0.0007017 | 0.00072474 | 0.00011515 | 0.00063218 | 0.00054344 | 63 | 63 | 49 | 59 | 58.5 | 38 | 39 | 5 | 34 | 29 |
| SC22B | Vesicle-traffi | 0.63766442 | 0.03252476 | 0.00049095 | 0.00049575 | 0.00061532 | 0.00037208 | 0.00049353 | 0.00031332 | 0.00031531 | 0 | 0.00031548 | 0.0003147 | 4 | 4 | 5 | 3 | 4 | 2 | 2 | 0 | 2 | 2 |
| PRP4 | U4/U6 small | 0.62613517 | 0.10732506 | 0.00040442 | 0.00040838 | 0.00050687 | 0.00020433 | 0.000381 | 0.0002581 | 0.00012987 | 0.00024141 | 0.00032485 | 0.00023856 | 8 | 8 | 10 | 4 | 7.5 | 4 | 2 | 3 | 5 | 3.5 |
| RL14 | 60S ribosoma | 0.62395268 | 0.00343607 | 0.0018411 | 0.0016112 | 0.0015998 | 0.0013643 | 0.0016041 | 0.0010966 | 0.00094592 | 0.0011723 | 0.00078871 | 0.00100088 | 15 | 13 | 13 | 11 | 13 | 7 | 6 | 6 | 5 | 6 |
| CP070 | UPF0183 pro | 0.62091688 | 0.05741446 | 0.00075038 | 0.00063143 | 0.00068968 | 0.00069508 | 0.00069164 | 0.00063851 | 0.00024096 | 0.00059724 | 0.0002411 | 0.00042945 | 12 | 10 | 11 | 11 | 11 | 8 | 3 | 6 | 3 | 5 |
| STRN | Striatin OS=H | 0.6154517 | 0.05217889 | 0.00043981 | 0.00068324 | 0.00061058 | 0.00034187 | 0.00051888 | 0.00030227 | 0.0003911 | 0.00032312 | 0.00026088 | 0.00031934 | 13 | 20 | 18 | 10 | 15.25 | 7 | 9 | 6 | 6 | 7 |
| XPO2 | Exportin-2 O | 0.61195372 | 0.05382766 | 0.00013588 | 0.00016465 | 0.00027249 | 0.00016477 | 0.00018445 | 0.00010406 | 0 | 0.00012978 | 0.00010478 | 0.00011287 | 5 | 6 | 10 | 6 | 6.75 | 3 | 0 | 3 | 3 | 3 |
| PUF60 | Poly(U)-bindi | 0.60972909 | 0.02937686 | 0.00066089 | 0.00057202 | 0.00056799 | 0.00071554 | 0.00062911 | 0.00024101 | 0.00054573 | 0 | 0.00036402 | 0.00038359 | 14 | 12 | 12 | 15 | 13.25 | 4 | 9 | 0 | 6 | 6.33333333 |
| ERC6L | DNA excision | 0.60968141 | 0.06014882 | 0.0004011 | 0.0004903 | 0.00033867 | 0.00053331 | 0.00044085 | 0.00032334 | 0.00037963 | 0.00010081 | 0.00027132 | 0.00026878 | 19 | 23 | 16 | 25 | 20.75 | 12 | 14 | 3 | 10 | 9.75 |
| GOGA2 | Golgin subfa | 0.60760792 | 0.08316578 | 0.00015801 | 0.00018615 | 0.00029046 | 0.00010645 | 0.00018527 | 0.00010084 | 0.00010148 | 0 | 0.00013539 | 0.00011257 | 6 | 7 | 11 | 4 | 7 | 3 | 3 | 0 | 4 | 3.33333333 |
| HS71A | Heat shock 7 | 0.60453922 | 0.00354746 | 0.0087687 | 0.0091039 | 0.0087508 | 0.008528 | 0.00878785 | 0.0056749 | 0.0064513 | 0.0031455 | 0.0059787 | 0.0053126 | 213 | 219 | 212 | 205 | 212.25 | 108 | 122 | 48 | 113 | 97.75 |
| HS71B | Heat shock 7 | 0.60453922 | 0.00354746 | 0.0087687 | 0.0091039 | 0.0087508 | 0.008528 | 0.00878785 | 0.0056749 | 0.0064513 | 0.0031455 | 0.0059787 | 0.0053126 | 213 | 219 | 212 | 205 | 212.25 | 108 | 122 | 48 | 113 | 97.75 |
| WDR6 | WD repeat-c | 0.59582322 | 0.00332832 | 0.00021186 | 0.00016639 | 0.00018882 | 0.00016651 | 0.0001834 | 0.00012018 | 0.00012095 | 7.4944E-05 | 0.00012101 | 0.00010927 | 9 | 7 | 8 | 7 | 7.75 | 4 | 4 | 2 | 4 | 3.5 |
| DYN2 | Dynamin-2 O | 0.59042808 | 0.0401859 | 0.00027298 | 0.00027565 | 0.00021289 | 0.0003065 | 0.00026701 | 7.7429E-05 | 0.0001948 | 0.00024141 | 0.00011695 | 0.00015765 | 9 | 9 | 7 | 10 | 8.75 | 2 | 5 | 5 | 3 | 3.75 |
| TCPE | T-complex pr | 0.58563869 | 0.00041928 | 0.0033656 | 0.0033985 | 0.003179 | 0.0026616 | 0.00315118 | 0.0017432 | 0.0020049 | 0.0019411 | 0.0016926 | 0.00184545 | 69 | 69 | 65 | 54 | 64.25 | 28 | 32 | 25 | 27 | 28 |
| NOG1 | Nucleolar GT | 0.58176253 | 0.03662803 | 0.00049947 | 0.0002942 | 0.0003756 | 0.00029442 | 0.00036592 | 0.00026563 | 0.00016039 | 0.00026502 | 0.00016048 | 0.00021288 | 12 | 7 | 9 | 7 | 8.75 | 5 | 3 | 4 | 3 | 3.75 |
| GRP75 | Stress-70 pro | 0.58165501 | 0.00051542 | 0.0013602 | 0.0013343 | 0.0010911 | 0.0012174 | 0.00125075 | 0.00084328 | 0.00069888 | 0.00061865 | 0.00074921 | 0.00072751 | 35 | 34 | 28 | 31 | 32 | 17 | 14 | 10 | 15 | 14 |
| LBR | Lamin-B rece | 0.57597823 | 0.0008866 | 0.00064362 | 0.00060659 | 0.00068836 | 0.00056366 | 0.00062556 | 0.00043813 | 0.00027557 | 0.00034151 | 0.00038602 | 0.00036031 | 15 | 14 | 16 | 13 | 14.5 | 8 | 5 | 5 | 7 | 6.25 |

| | | | | | | | | | | | | | | | | | | | | | | | |
|---|---|---|---|---|---|---|---|---|---|---|---|---|---|---|---|---|---|---|---|---|---|---|---|
| SRPRB | Signal recogr | 0.57307185 | 0.01716749 | 0.00068162 | 0.00068829 | 0.00097634 | 0.00059038 | 0.00073416 | 0.00049715 | 0.0005003 | 0.00031001 | 0.00037544 | 0.00042073 | 7 | 7 | 10 | 6 | 7.5 | 4 | 4 | 2 | 3 | 3.25 |
| COF1 | Cofilin-1 OS= | 0.5715378 | 0.00119945 | 0.0022255 | 0.0024078 | 0.0022315 | 0.001767 | 0.00215795 | 0.0012174 | 0.0014293 | 0.0012652 | 0.0010215 | 0.00123335 | 14 | 15 | 14 | 11 | 13.5 | 6 | 7 | 5 | 5 | 5.75 |
| EDC4 | Enhancer of r | 0.56868786 | 0.02497901 | 0.0011678 | 0.0012553 | 0.0015297 | 0.0012372 | 0.0012975 | 0.00069719 | 0.00099195 | 0.00026985 | 0.0009925 | 0.00073787 | 62 | 66 | 81 | 65 | 68.5 | 29 | 41 | 9 | 41 | 30 |
| PER1 | Period circad | 0.56748178 | 0.0804316 | 0.00022502 | 0.00012394 | 0.00034868 | 0.00022738 | 0.00023126 | 7.8329E-05 | 0.00021021 | 0 | 0.00010516 | 0.00013123 | 11 | 6 | 17 | 11 | 11.25 | 3 | 8 | 0 | 4 | 5 |
| U2AF2 | Splicing facto | 0.56738687 | 0.04371999 | 0.00083332 | 0.0010659 | 0.00072413 | 0.00095435 | 0.00089443 | 0.00042545 | 0.00064223 | 0.00017687 | 0.00078539 | 0.00050749 | 15 | 19 | 13 | 17 | 16 | 6 | 9 | 2 | 11 | 7 |
| F135A | Protein FAM1 | 0.5663277 | 0.03000508 | 0.00017418 | 0.00026383 | 0.00017464 | 0.00017601 | 0.00019717 | 0.00011116 | 0.00006712 | 0 | 0.0001567 | 0.00011166 | 10 | 15 | 10 | 10 | 11.25 | 5 | 3 | 0 | 7 | 5 |
| EP15R | Epidermal gr | 0.56578205 | 0.00977195 | 0.0005803 | 0.00070934 | 0.00067372 | 0.00080244 | 0.00069145 | 0.0004678 | 0.00043154 | 0.00019447 | 0.00047103 | 0.00039121 | 19 | 23 | 22 | 26 | 22.5 | 12 | 11 | 4 | 12 | 9.75 |
| SENP3 | Sentrin-speci | 0.55719917 | 0.05422956 | 0.00027584 | 0.00018569 | 0.00041486 | 0.00023228 | 0.00027717 | 0.00017604 | 0.0001181 | 0.00014636 | 0.00017725 | 0.00015444 | 6 | 4 | 9 | 5 | 6 | 3 | 2 | 2 | 3 | 2.5 |
| XRN2 | 5'-3' exoribor | 0.54897078 | 0.01582076 | 0.00099998 | 0.00092561 | 0.0010026 | 0.00070173 | 0.00090748 | 0.00067363 | 0.00032112 | 0 | 0.00049979 | 0.00049818 | 36 | 33 | 36 | 25 | 32.5 | 19 | 9 | 0 | 14 | 14 |
| 41 | Protein 4.1 O | 0.5460074 | 0.01351538 | 0.0002138 | 0.00024673 | 0.00015312 | 0.0002469 | 0.00021514 | 0.00011695 | 0.00011769 | 0 | 0.00011776 | 0.00011747 | 7 | 8 | 5 | 8 | 7 | 3 | 3 | 0 | 3 | 3 |
| ARFG1 | ADP-ribosyla | 0.54415167 | 0.03566014 | 0.00058496 | 0.00045942 | 0.00032585 | 0.00045975 | 0.0004575 | 0.00016592 | 0 | 0.00041385 | 0.00016707 | 0.00024895 | 9 | 7 | 5 | 7 | 7 | 2 | 0 | 4 | 2 | 2.66666667 |
| ZO1 | Tight junctior | 0.54275012 | 0.04437804 | 0.0022192 | 0.0019665 | 0.0016347 | 0.0019069 | 0.00193183 | 0.0014259 | 0.0014155 | 7.2093E-05 | 0.0012805 | 0.0010485 | 147 | 129 | 108 | 125 | 127.25 | 74 | 73 | 3 | 66 | 54 |
| TCPB | T-complex pr | 0.54204519 | 0.00094204 | 0.00029594 | 0.00039845 | 0.00034619 | 0.00029905 | 0.00033491 | 0.00018887 | 0.00019007 | 0.00015703 | 0.00019017 | 0.00018154 | 6 | 8 | 7 | 6 | 6.75 | 3 | 3 | 2 | 3 | 2.75 |
| NOP58 | Nucleolar prc | 0.54164457 | 0.00122316 | 0.0005986 | 0.00045334 | 0.00065021 | 0.0006553 | 0.00058936 | 0.00031835 | 0.00032037 | 0.00031763 | 0.00032055 | 0.00031923 | 12 | 9 | 13 | 13 | 11.75 | 5 | 5 | 4 | 5 | 4.75 |
| RBM4 | RNA-binding | 0.53605098 | 0.0150414 | 0.00072496 | 0.00080525 | 0.00058151 | 0.00065931 | 0.00069276 | 0.00055519 | 0.00027936 | 0 | 0.00027951 | 0.00037135 | 10 | 11 | 8 | 9 | 9.5 | 6 | 3 | 0 | 3 | 4 |
| ARP3 | Actin-related | 0.52973534 | 0.0232536 | 0.0006313 | 0.00038249 | 0.00056968 | 0.00044655 | 0.00050751 | 0.00016116 | 0 | 0.00040197 | 0.0002434 | 0.00026884 | 10 | 6 | 9 | 7 | 8 | 2 | 0 | 4 | 3 | 3 |
| CING | Cingulin OS=I | 0.52835268 | 0.02060766 | 0.0015211 | 0.0012911 | 0.001282 | 0.0013589 | 0.00136328 | 0.001013 | 0.00093447 | 0.00014037 | 0.00079332 | 0.00072029 | 69 | 58 | 58 | 61 | 61.5 | 36 | 33 | 4 | 28 | 25.25 |
| ACLY | ATP-citrate sy | 0.52146524 | 0.00472388 | 0.00086284 | 0.00096808 | 0.00067288 | 0.00092034 | 0.00085604 | 0.0003671 | 0.00036943 | 0.0006486 | 0.00040044 | 0.00044639 | 36 | 40 | 28 | 38 | 35.5 | 12 | 12 | 17 | 13 | 13.5 |
| KRI1 | Protein KRI1 | 0.51378188 | 0.00536788 | 0.00041291 | 0.00041694 | 0.00033873 | 0.00045517 | 0.00040594 | 0.00019164 | 0.00024108 | 0 | 0.00019297 | 0.00020856 | 11 | 11 | 9 | 12 | 10.75 | 4 | 5 | 0 | 4 | 4.33333333 |
| CALD1 | Caldesmon O | 0.50983892 | 0.01063459 | 0.00023294 | 0.00030242 | 0.00026692 | 0.00020176 | 0.00025101 | 0.00012742 | 0.00017097 | 0 | 8.5534E-05 | 0.00012797 | 7 | 9 | 8 | 6 | 7.5 | 3 | 4 | 0 | 2 | 3 |
| ILF2 | Interleukin er | 0.50737877 | 0.02717849 | 0.0014209 | 0.0012298 | 0.00061058 | 0.0012991 | 0.0011401 | 0.00051818 | 0.00043456 | 0 | 0.00078264 | 0.00057846 | 21 | 18 | 9 | 19 | 16.75 | 6 | 5 | 0 | 9 | 6.66666667 |
| RL1D1 | Ribosomal L1 | 0.50672071 | 0.00326012 | 0.0017233 | 0.0010876 | 0.0012419 | 0.0012517 | 0.00132613 | 0.00068738 | 0.00062257 | 0.00068582 | 0.00069213 | 0.00067198 | 32 | 20 | 23 | 23 | 24.5 | 10 | 9 | 8 | 10 | 9.25 |
| DDX27 | Probable ATF | 0.50589151 | 0.01538587 | 0.00029836 | 0.0002678 | 0.00036564 | 0.0003015 | 0.00030833 | 8.4627E-05 | 0.00025549 | 0 | 0.00012782 | 0.00015598 | 9 | 8 | 11 | 9 | 9.25 | 2 | 6 | 0 | 3 | 3.66666667 |
| HNRPK | Heterogeneo | 0.50578064 | 0.0033173 | 0.0018808 | 0.0016115 | 0.0012572 | 0.0016702 | 0.00160493 | 0.0010184 | 0.00095171 | 0.00054436 | 0.00073249 | 0.00081174 | 33 | 28 | 22 | 29 | 28 | 14 | 13 | 6 | 10 | 10.75 |
| NGAP | Ras GTPase-a | 0.5038304 | 0.00113455 | 0.00013901 | 0.00016376 | 0.00013938 | 0.00011706 | 0.0001398 | 5.9142E-05 | 5.9518E-05 | 0.00007376 | 8.9327E-05 | 7.0437E-05 | 6 | 7 | 6 | 5 | 6 | 2 | 2 | 2 | 3 | 2.25 |
| CSK21 | Casein kinase | 0.50251539 | 0.00626703 | 0.0006749 | 0.00074965 | 0.0008797 | 0.00061379 | 0.00072951 | 0.00043071 | 0.00034676 | 0.0003223 | 0 | 0.00036659 | 10 | 11 | 13 | 9 | 10.75 | 5 | 4 | 3 | 0 | 4 |
| RANB9 | Ran-binding p | 0.49900076 | 0.02436934 | 0.00036198 | 0.00021931 | 0.00032665 | 0.00018289 | 0.00027271 | 9.2405E-05 | 0.00013949 | 0.00017286 | 0.00013957 | 0.00013608 | 10 | 6 | 9 | 5 | 7.5 | 2 | 3 | 3 | 3 | 2.75 |
| NACAM | Nascent poly | 0.49709585 | 0.00151139 | 0.00013969 | 0.00019235 | 0.00014006 | 0.00014116 | 0.00015332 | 8.1043E-05 | 8.1558E-05 | 6.0644E-05 | 8.1604E-05 | 7.6212E-05 | 11 | 15 | 11 | 11 | 12 | 5 | 5 | 3 | 5 | 4.5 |
| RL30 | 60S ribosoma | 0.49561872 | 0.0037794 | 0.0020652 | 0.0032439 | 0.0023008 | 0.0023187 | 0.00248215 | 0.0011715 | 0.001179 | 0.0010958 | 0.0014745 | 0.0012302 | 9 | 14 | 10 | 10 | 10.75 | 4 | 4 | 3 | 5 | 4 |
| ATG2B | Autophagy-re | 0.49192693 | 0.00098448 | 0.00033017 | 0.00039752 | 0.00039472 | 0.00042347 | 0.00038647 | 0.00016209 | 0.00019574 | 0.0001415 | 0.00026113 | 0.00019012 | 26 | 31 | 31 | 33 | 30.25 | 10 | 12 | 7 | 16 | 11.25 |
| DKC1 | H/ACA ribonu | 0.48617353 | 0.02126725 | 0.00046205 | 0.00025921 | 0.00051476 | 0.00057066 | 0.00045167 | 0.00013106 | 0.00026378 | 0 | 0.00026393 | 0.00021959 | 9 | 5 | 10 | 11 | 8.75 | 2 | 4 | 0 | 4 | 3.33333333 |
| CAPZB | F-actin-cappi | 0.48566571 | 0.00525702 | 0.00095265 | 0.0011544 | 0.0013373 | 0.0012515 | 0.00117396 | 0.00048638 | 0.00048947 | 0 | 0.00073461 | 0.00057015 | 10 | 12 | 14 | 13 | 12.25 | 4 | 4 | 0 | 6 | 4.66666667 |
| ANM7 | Protein argin | 0.48279811 | 0.07056693 | 0.00026693 | 0.00069312 | 0.00030588 | 0.00065508 | 0.00048025 | 0.00019469 | 0.00024491 | 0.00024281 | 0.00024505 | 0.00023187 | 7 | 18 | 8 | 17 | 12.5 | 4 | 5 | 4 | 5 | 4.5 |
| DNJA2 | DnaJ homolo | 0.47721117 | 0.00100814 | 0.0006405 | 0.00051741 | 0.00051376 | 0.00051778 | 0.00054736 | 0.00024525 | 0.00032908 | 0.00030587 | 0.00016463 | 0.00026121 | 10 | 8 | 8 | 8 | 8.5 | 3 | 4 | 3 | 2 | 3 |
| NCKP1 | Nck-associate | 0.47331022 | 0.02622094 | 0.00021055 | 0.00035434 | 0.00023456 | 0.00021276 | 0.00025305 | 0.00020902 | 6.0099E-05 | 0 | 9.0198E-05 | 0.00011977 | 9 | 15 | 10 | 9 | 10.75 | 7 | 2 | 0 | 3 | 4 |
| COR1C | Coronin-1C C | 0.47280845 | 0.00306215 | 0.00061239 | 0.00056216 | 0.00050238 | 0.00061882 | 0.00057394 | 0.00021317 | 0.00021453 | 0.0004431 | 0.00021465 | 0.00027136 | 11 | 10 | 9 | 11 | 10.25 | 3 | 3 | 5 | 3 | 3.5 |
| KAP2 | cAMP-depen | 0.4679591 | 0.00035369 | 0.0024821 | 0.0024404 | 0.0028161 | 0.0021781 | 0.00247918 | 0.0011672 | 0.0015102 | 0.0010398 | 0.00092341 | 0.00116015 | 38 | 37 | 43 | 33 | 37.75 | 14 | 18 | 10 | 11 | 13.25 |
| AP2A1 | AP-2 comple: | 0.46780383 | 0.00381816 | 0.00040514 | 0.0005182 | 0.00040622 | 0.00054587 | 0.00046886 | 0.00020685 | 0.00020816 | 0 | 0.00024299 | 0.00021933 | 15 | 19 | 15 | 20 | 17.25 | 6 | 6 | 0 | 7 | 6.33333333 |
| PRS7 | 26S proteaso | 0.46747447 | 0.01303683 | 0.00060943 | 0.00036924 | 0.00061106 | 0.0003695 | 0.00048981 | 0.00023336 | 0.00023484 | 0.00029104 | 0.00015665 | 0.00022897 | 10 | 6 | 10 | 6 | 8 | 3 | 3 | 3 | 2 | 2.75 |
| NUMA1 | Nuclear mito | 0.46553179 | 0.00089006 | 0.0014598 | 0.0013985 | 0.0014762 | 0.0012608 | 0.00139883 | 0.00081218 | 0.00073721 | 0.00031778 | 0.00073762 | 0.0006512 | 117 | 111 | 118 | 100 | 111.5 | 51 | 46 | 16 | 46 | 39.75 |
| PARP1 | Poly [ADP-rib | 0.46184375 | 0.00275719 | 0.0011451 | 0.0011825 | 0.0010176 | 0.0015253 | 0.00121763 | 0.0005979 | 0.00046799 | 0.00041426 | 0.00076926 | 0.00056235 | 44 | 45 | 39 | 58 | 46.5 | 18 | 14 | 10 | 23 | 16.25 |
| DDX24 | ATP-depende | 0.46084761 | 0.00455351 | 0.00043008 | 0.00037224 | 0.00036962 | 0.00031043 | 0.00037059 | 0.00011763 | 0.00015784 | 0 | 0.00023689 | 0.00017079 | 14 | 12 | 12 | 10 | 12 | 3 | 4 | 0 | 6 | 4.33333333 |
| RRP7A | Ribosomal RI | 0.46015768 | 0.00295644 | 0.00075395 | 0.00076133 | 0.00085046 | 0.00095234 | 0.00082952 | 0.00036087 | 0.00048422 | 0.00030004 | 0 | 0.00038171 | 8 | 8 | 9 | 10 | 8.75 | 3 | 4 | 2 | 0 | 3 |
| U2AF1 | Splicing facto | 0.45579698 | 0.00389324 | 0.0012095 | 0.0011103 | 0.00088196 | 0.0013333 | 0.00113377 | 0.00056136 | 0.0004237 | 0 | 0.00056524 | 0.00051677 | 11 | 10 | 8 | 12 | 10.25 | 4 | 3 | 0 | 4 | 3.66666667 |
| U2AF5 | Splicing facto | 0.45579698 | 0.00389324 | 0.0012095 | 0.0011103 | 0.00088196 | 0.0013333 | 0.00113377 | 0.00056136 | 0.0004237 | 0 | 0.00056524 | 0.00051677 | 11 | 10 | 8 | 12 | 10.25 | 4 | 3 | 0 | 4 | 3.66666667 |
| CAZA1 | F-actin-cappi | 0.45485579 | 0.00497959 | 0.0012917 | 0.0013975 | 0.0010176 | 0.0014918 | 0.00129965 | 0.00070661 | 0.0007111 | 0 | 0.00035575 | 0.00059115 | 14 | 15 | 11 | 16 | 14 | 6 | 6 | 0 | 3 | 5 |
| RAE1L | mRNA export | 0.45313014 | 0.00338574 | 0.0010039 | 0.00086891 | 0.0012223 | 0.0012318 | 0.00108173 | 0.00054916 | 0.00046054 | 0 | 0.00046079 | 0.00049016 | 14 | 12 | 17 | 17 | 15 | 6 | 5 | 0 | 5 | 5.33333333 |
| EXOS3 | Exosome con | 0.44600385 | 0.00333045 | 0.0012475 | 0.0013565 | 0.0010583 | 0.0011636 | 0.00120648 | 0.00036744 | 0.0008628 | 0.0003055 | 0.00061663 | 0.00053809 | 13 | 14 | 11 | 12 | 12.5 | 3 | 7 | 2 | 5 | 4.25 |
| APBB1 | Amyloid-beta | 0.44448422 | 0.00341509 | 0.0013752 | 0.001276 | 0.0011552 | 0.0013896 | 0.001299 | 0.00047439 | 0.0010025 | 0.00035498 | 0.00047767 | 0.00057739 | 37 | 34 | 31 | 37 | 34.75 | 10 | 21 | 6 | 10 | 11.75 |
| RPAP3 | RNA polymer | 0.43810738 | 0.00455567 | 0.0004365 | 0.00056098 | 0.00035809 | 0.00044109 | 0.00044917 | 0.00030389 | 0.00015291 | 0.00012633 | 0.000204 | 0.00019678 | 11 | 14 | 9 | 11 | 11.25 | 6 | 3 | 2 | 4 | 3.75 |
| DDX50 | ATP-depende | 0.43204659 | 0.00204189 | 0.00053708 | 0.00054233 | 0.00050261 | 0.00054272 | 0.00053119 | 0.0002285 | 0.00032194 | 0 | 0.00013805 | 0.0002295 | 17 | 19 | 16 | 17 | 17.25 | 7 | 7 | 0 | 5 | 6.33333333 |
| ROA1 | Heterogeneo | 0.42899841 | 0.00304757 | 0.0019862 | 0.0012177 | 0.0016359 | 0.0019354 | 0.0016938 | 0.00090542 | 0.00072894 | 0.00045168 | 0.00082051 | 0.00072664 | 28 | 19 | 23 | 27 | 24.25 | 11 | 9 | 4 | 10 | 8.5 |
| HNRPM | Heterogeneo | 0.42818199 | 0.00158502 | 0.0011929 | 0.0011681 | 0.001486 | 0.0012054 | 0.0012631 | 0.00046139 | 0.00060362 | 0 | 0.0005575 | 0.00054084 | 33 | 32 | 41 | 33 | 34.75 | 10 | 13 | 0 | 12 | 11.6666667 |
| IF2P | Eukaryotic tra | 0.42584383 | 0.00122894 | 0.00036771 | 0.00045867 | 0.000347 | 0.00043714 | 0.00040263 | 0.00016565 | 0.0001667 | 0.00010329 | 0.00025019 | 0.00017146 | 17 | 21 | 16 | 20 | 18.5 | 6 | 6 | 3 | 9 | 6 |
| NUP98 | Nuclear pore | 0.42468496 | 0.00418553 | 0.00031951 | 0.00035196 | 0.00026211 | 0.00029351 | 0.00030677 | 0.0001483 | 0.00018655 | 0 | 5.5995E-05 | 0.00013028 | 22 | 24 | 18 | 20 | 21 | 8 | 10 | 0 | 3 | 7 |
| IMA5 | Importin sub | 0.42349367 | 0.00734903 | 0.00093193 | 0.0006934 | 0.00078688 | 0.0012391 | 0.00091283 | 0.00031303 | 0 | 0.00046847 | 0.00037823 | 0.00038658 | 33 | 27 | 28 | 39 | 31.75 | 10 | 0 | 9 | 9 | 9.33333333 |
| BCLF1 | Bcl-2-associa | 0.41481824 | 0.02846562 | 0.00060234 | 0.0010427 | 0.00043139 | 0.00052172 | 0.00064954 | 0.0003661 | 0.0002579 | 0 | 0.00018432 | 0.00026944 | 21 | 37 | 15 | 18 | 22.75 | 10 | 7 | 0 | 5 | 7.33333333 |
| MBB1A | Myb-binding | 0.41451307 | 2.7826E-05 | 0.00097367 | 0.00088287 | 0.00093641 | 0.00090358 | 0.00092413 | 0.00035508 | 0.00028076 | 0.0004112 | 0.00048522 | 0.00038307 | 49 | 44 | 47 | 45 | 46.25 | 14 | 11 | 13 | 19 | 14.25 |
| SPTN1 | Spectrin alph | 0.41448339 | 0.00339338 | 0.00077927 | 0.00074377 | 0.00099541 | 0.00057171 | 0.00077254 | 0.00031338 | 0.00042507 | 0.00032286 | 0.00021951 | 0.00032021 | 73 | 69 | 93 | 53 | 72 | 23 | 31 | 19 | 16 | 22.25 |
| TPD54 | Tumor protei | 0.41133651 | 0.00044658 | 0.0015372 | 0.0019403 | 0.0021835 | 0.0018122 | 0.0018683 | 0.00098102 | 0.00082271 | 0.00061174 | 0.00065853 | 0.0007685 | 12 | 15 | 17 | 14 | 14.5 | 6 | 5 | 3 | 4 | 4.5 |
| KIF23 | Kinesin-like p | 0.38158174 | 0.0190247 | 0.00052227 | 0.00030532 | 0.00055122 | 0.00022221 | 0.00040026 | 0.00014034 | 0.00017654 | 0 | 0.00014131 | 0.00015273 | 19 | 11 | 20 | 8 | 14.5 | 4 | 5 | 0 | 4 | 4.33333333 |
| RBM27 | RNA-binding | 0.38117586 | 0.00078581 | 0.000946 | 0.00080442 | 0.00094852 | 0.00072953 | 0.00085712 | 0.00047663 | 0.00031977 | 0.00015851 | 0.00035194 | 0.00032671 | 38 | 32 | 38 | 29 | 34.25 | 15 | 10 | 4 | 11 | 10 |
| SPTB2 | Spectrin beta | 0.37209708 | 7.7236E-05 | 0.00070324 | 0.0005974 | 0.00070512 | 0.00063167 | 0.00065936 | 0.00024221 | 0.00022941 | 0.00033761 | 0.00017215 | 0.00024535 | 63 | 53 | 63 | 56 | 58.75 | 17 | 16 | 19 | 12 | 16 |
| LAS1L | Ribosomal bi | 0.37054499 | 0.00148003 | 0.00043142 | 0.00036303 | 0.00043257 | 0.00050861 | 0.00043391 | 0.00022944 | 9.2359E-05 | 0.00022892 | 0.00009241 | 0.00016078 | 12 | 10 | 12 | 14 | 12 | 5 | 2 | 4 | 2 | 3.25 |
| NONO | Non-POU dor | 0.36301883 | 0.00037733 | 0.0067232 | 0.0088256 | 0.0067411 | 0.0077562 | 0.00751153 | 0.0032895 | 0.0028786 | 0.001427 | 0.0033122 | 0.00272683 | 142 | 173 | 139 | 159 | 153.25 | 46 | 43 | 16 | 48 | 38.25 |
| RING2 | E3 ubiquitin-p | 0.3604367 | 0.00062067 | 0.00070683 | 0.00055514 | 0.00070871 | 0.0006349 | 0.0006514 | 0.00030073 | 0.00020176 | 0 | 0.00020187 | 0.00023479 | 9 | 7 | 9 | 8 | 8.25 | 3 | 2 | 0 | 2 | 2.33333333 |
| SCRIB | Protein scribl | 0.35838957 | 0.00122382 | 0.00055043 | 0.00039234 | 0.00043827 | 0.0004417 | 0.00045569 | 0.0002273 | 0.00018715 | 5.1541E-05 | 0.00018726 | 0.00016331 | 34 | 24 | 27 | 27 | 28 | 11 | 9 | 2 | 9 | 7.75 |
| SGT1 | Protein SGT1 | 0.3581536 | 0.00078892 | 0.0010122 | 0.0010951 | 0.0010149 | 0.0010228 | 0.00103625 | 0.00018456 | 0.00055719 | 0 | 0.00037166 | 0.00037114 | 14 | 15 | 14 | 14 | 14.25 | 2 | 6 | 0 | 4 | 4 |
| NOLC1 | Nucleolar and | 0.35366791 | 0.01147447 | 0.00067953 | 0.00083866 | 0.00037852 | 0.00049593 | 0.00059816 | 0.00024093 | 0.00019397 | 0.00012019 | 0.00029111 | 0.00021155 | 18 | 22 | 10 | 13 | 15.75 | 5 | 4 | 2 | 6 | 4.25 |
| SLK | STE20-like se | 0.35278212 | 0.00096923 | 0.00034187 | 0.00023734 | 0.00025709 | 0.00030228 | 0.00028465 | 0.00010909 | 0.00010978 | 0 | 8.2383E-05 | 0.00010042 | 16 | 11 | 12 | 14 | 13.25 | 4 | 4 | 0 | 3 | 3.66666667 |
| STAT1 | Signal transd | 0.35138214 | 0.00205555 | 0.00038703 | 0.00060399 | 0.00045862 | 0.00060442 | 0.00051352 | 0.00017964 | 0.00013558 | 0 | 0.0002261 | 0.00018044 | 11 | 17 | 13 | 17 | 14.5 | 4 | 3 | 0 | 5 | 4 |
| ABCF1 | ATP-binding c | 0.34911931 | 0.00017047 | 0.00040598 | 0.00050455 | 0.00046968 | 0.00053647 | 0.00047917 | 0.00023916 | 0.00016045 | 0.00014913 | 0.00012041 | 0.00016729 | 13 | 16 | 15 | 17 | 15.25 | 6 | 4 | 3 | 3 | 4 |
| EHBP1 | EH domain-b | 0.33964005 | 0.00391282 | 0.00019293 | 0.00030305 | 0.00017195 | 0.00019496 | 0.00021572 | 8.2083E-05 | 0.00005507 | 0 | 8.2651E-05 | 7.3268E-05 | 9 | 14 | 8 | 9 | 10 | 3 | 2 | 0 | 3 | 2.66666667 |
| RL27 | 60S ribosoma | 0.3324047 | 0.00170098 | 0.0023284 | 0.0015674 | 0.0025291 | 0.0025489 | 0.00224345 | 0.00099063 | 0.00049846 | 0 | 0.00074811 | 0.00074573 | 12 | 8 | 13 | 13 | 11.5 | 4 | 2 | 0 | 3 | 3 |
| MYO1D | Unconventio | 0.32852981 | 0.03479111 | 0.00049839 | 7.9463E-05 | 0.00047342 | 0.00045061 | 0.00037547 | 0.00010044 | 0.00013477 | 0 | 0.00013485 | 0.00012335 | 19 | 3 | 18 | 17 | 14.25 | 3 | 4 | 0 | 4 | 3.66666667 |
| RL12 | 60S ribosoma | 0.32838271 | 9.2996E-05 | 0.0027188 | 0.0027454 | 0.002245 | 0.0025858 | 0.00257375 | 0.00081652 | 0.00082171 | 0.00050917 | 0.0012333 | 0.00084518 | 17 | 17 | 14 | 16 | 16 | 4 | 4 | 2 | 6 | 4 |
| RFC1 | Replication fa | 0.32800197 | 6.0671E-05 | 0.00062063 | 0.0006267 | 0.00055314 | 0.00051101 | 0.00057787 | 0.00020538 | 0.00020668 | 0.00010977 | 0.00023634 | 0.00018954 | 27 | 27 | 24 | 22 | 25 | 7 | 7 | 3 | 8 | 6.25 |
| PCNP | PEST proteol | 0.32654704 | 3.3442E-05 | 0.0014825 | 0.001497 | 0.0011892 | 0.0013483 | 0.00137925 | 0.00056767 | 0.00038085 | 0.00047198 | 0.00038106 | 0.00045039 | 10 | 10 | 8 | 9 | 9.25 | 3 | 2 | 2 | 2 | 2.25 |
| WDR44 | WD repeat-co | 0.3259886 | 0.00024316 | 0.00034684 | 0.00032104 | 0.0002898 | 0.00023365 | 0.00029783 | 0.00011067 | 0.00011138 | 9.2018E-05 | 7.4292E-05 | 0.00009709 | 12 | 11 | 10 | 8 | 10.25 | 3 | 3 | 2 | 2 | 2.5 |
| AKAP1 | A-kinase ancl | 0.32452269 | 0.00021663 | 0.0010813 | 0.0012984 | 0.001172 | 0.0011517 | 0.00117585 | 0.000373 | 0.00045044 | 0.00013956 | 0.00056336 | 0.00038159 | 37 | 44 | 40 | 39 | 40 | 10 | 12 | 3 | 15 | 10 |
| SMCA5 | SWI/SNF-rela | 0.32427461 | 4.0448E-05 | 0.0011288 | 0.0008612 | 0.00095573 | 0.00096321 | 0.00097724 | 0.00028815 | 0.00028998 | 0.0003993 | 0.00029014 | 0.00031689 | 45 | 34 | 38 | 38 | 38.75 | 9 | 9 | 10 | 9 | 9.25 |
| TCPQ | T-complex pr | 0.32317415 | 8.4377E-05 | 0.0019262 | 0.0022367 | 0.0021727 | 0.0023357 | 0.00216783 | 0.0004917 | 0.00055668 | 0.0010732 | 0.00068076 | 0.00070059 | 40 | 46 | 45 | 48 | 44.75 | 8 | 9 | 14 | 11 | 10.5 |
| NOP56 | Nucleolar prc | 0.32038922 | 0.00027376 | 0.0008885 | 0.00076261 | 0.00075723 | 0.00067337 | 0.00077043 | 0.00022681 | 0.00028532 | 0 | 0.00022838 | 0.00024684 | 20 | 17 | 17 | 15 | 17.25 | 4 | 5 | 0 | 4 | 4.33333333 |
| RUVB1 | RuvB-like 1 C | 0.30670301 | 3.4406E-05 | 0.007523 | 0.0082394 | 0.0072529 | 0.0085377 | 0.00788825 | 0.0018466 | 0.002007 | 0.0035926 | 0.0022312 | 0.00241935 | 130 | 141 | 125 | 146 | 135.5 | 25 | 27 | 39 | 30 | 30.25 |
| ESYT1 | Extended syn | 0.30474428 | 7.3673E-06 | 0.0010517 | 0.00098959 | 0.00088675 | 0.0010869 | 0.00100374 | 0.0003356 | 0.00024562 | 0.00030439 | 0.00033792 | 0.00030588 | 44 | 41 | 37 | 45 | 41.75 | 11 | 8 | 8 | 11 | 9.5 |
| RUVB2 | RuvB-like 2 C | 0.30430746 | 4.3865E-06 | 0.0083212 | 0.0085752 | 0.0081719 | 0.009618 | 0.00867158 | 0.0024734 | 0.0021963 | 0.0031754 | 0.0027102 | 0.00263883 | 146 | 149 | 143 | 167 | 151.25 | 34 | 30 | 35 | 37 | 34 |
| MBD3 | Methyl-CpG- | 0.30242714 | 0.00153298 | 0.00090682 | 0.0010073 | 0.0013638 | 0.00082471 | 0.00102566 | 0.00023149 | 0.00034944 | 0 | 0.00034963 | 0.00031019 | 10 | 11 | 15 | 9 | 11.25 | 2 | 3 | 0 | 3 | 2.66666667 |
| MTREX | Exosome RNA | 0.29969397 | 0.00040906 | 0.00058247 | 0.00066488 | 0.00048245 | 0.00051182 | 0.00056041 | 0.0001293 | 0.00013012 | 0.00028219 | 0.00013019 | 0.00016795 | 23 | 26 | 19 | 20 | 22 | 4 | 4 | 7 | 4 | 4.75 |
| IMA7 | Importin sub | 0.295537 | 2.3156E-05 | 0.0017724 | 0.0018891 | 0.0022213 | 0.0018407 | 0.00193088 | 0.00062839 | 0.00044267 | 0.00070533 | 0.00050619 | 0.00057065 | 36 | 38 | 45 | 37 | 39 | 10 | 7 | 9 | 8 | 8.5 |
| HDAC2 | Histone deac | 0.29139899 | 2.2811E-05 | 0.00070297 | 0.00092826 | 0.00081328 | 0.00081964 | 0.00081604 | 0.00027608 | 0.00020837 | 0.00025823 | 0.00020849 | 0.00023779 | 13 | 17 | 15 | 15 | 15 | 4 | 3 | 3 | 3 | 3.25 |
| TR150 | Thyroid horm | 0.29125771 | 0.00010033 | 0.00052501 | 0.00050224 | 0.00047099 | 0.00044675 | 0.00048625 | 0.00017634 | 0.00010648 | 0 | 0.00014205 | 0.00014162 | 19 | 18 | 17 | 16 | 17.5 | 5 | 3 | 0 | 4 | 4 |
| RLA2 | 60S acidic rib | 0.28766355 | 0.00043889 | 0.003442 | 0.0046342 | 0.0034511 | 0.0034781 | 0.00375135 | 0.00087865 | 0.00088423 | 0 | 0.0014745 | 0.00107913 | 15 | 20 | 15 | 15 | 16.25 | 3 | 3 | 0 | 5 | 3.66666667 |
| PR40A | Pre-mRNA-pr | 0.28281577 | 0.00122391 | 0.00082722 | 0.00072394 | 0.00063589 | 0.00097523 | 0.00079057 | 0.00035195 | 0.00024793 | 0 | 7.0877E-05 | 0.00022359 | 30 | 26 | 23 | 35 | 28.5 | 10 | 7 | 0 | 2 | 6.33333333 |
| RLA0 | 60S acidic rib | 0.27106203 | 4.439E-07 | 0.0021643 | 0.0021855 | 0.002337 | 0.0023553 | 0.00226053 | 0.00063751 | 0.00074848 | 0.00053005 | 0.00053493 | 0.00061274 | 26 | 26 | 28 | 28 | 27 | 6 | 7 | 4 | 5 | 5.5 |
| NAT10 | RNA cytidine | 0.26958746 | 3.8715E-06 | 0.0027804 | 0.0030676 | 0.0025555 | 0.0026015 | 0.00275125 | 0.0006572 | 0.00082672 | 0.00065571 | 0.00082718 | 0.0007417 | 108 | 118 | 99 | 100 | 106.25 | 20 | 25 | 16 | 25 | 21.5 |
| DICER | Endoribonucl | 0.26453742 | 0.00024016 | 0.00017849 | 0.00012478 | 0.00015143 | 0.00016649 | 0.0001553 | 3.5048E-05 | 5.2907E-05 | 0 | 3.5291E-05 | 4.1082E-05 | 13 | 9 | 11 | 12 | 11.25 | 2 | 3 | 0 | 2 | 2.33333333 |
| DNJB1 | DnaJ homolo | 0.25712231 | 0.00054861 | 0.00085374 | 0.0011756 | 0.00070038 | 0.00094114 | 0.00091772 | 0.00019813 | 0.00019939 | 0.0002471 | 0.00029924 | 0.00023597 | 11 | 15 | 9 | 12 | 11.75 | 2 | 2 | 2 | 3 | 2.25 |
| ACTG | Actin, cytopla | 0.25542297 | 0.0001606 | 0.01534 | 0.015135 | 0.018486 | 0.019839 | 0.0172 | 0.0070058 | 0.0046098 | 0.0036965 | 0.002261 | 0.00439328 | 218 | 213 | 262 | 279 | 243 | 78 | 51 | 33 | 25 | 46.75 |
| DSRAD | Double-stran | 0.24631854 | 8.4818E-06 | 0.00066724 | 0.0006303 | 0.00075535 | 0.00071775 | 0.00069266 | 0.00016484 | 0.00019353 | 0.00010279 | 0.0002213 | 0.00017062 | 31 | 29 | 35 | 33 | 32 | 6 | 7 | 3 | 8 | 6 |
| U520 | U5 small nuc | 0.23562557 | 2.3486E-05 | 0.00064241 | 0.0006986 | 0.00073083 | 0.00061171 | 0.00067089 | 7.8843E-05 | 0.00012695 | 0.00023599 | 0.00019053 | 0.00015808 | 52 | 56 | 59 | 49 | 54 | 5 | 8 | 12 | 12 | 9.25 |
| SFPQ | Splicing facto | 0.23202763 | 3.2876E-05 | 0.0028367 | 0.0036182 | 0.0030313 | 0.0029796 | 0.00311645 | 0.00085752 | 0.00095886 | 0.00035649 | 0.00071954 | 0.0007231 | 76 | 96 | 81 | 79 | 83 | 18 | 20 | 6 | 15 | 14.75 |
| DNM1L | Dynamin-1-li | 0.22247715 | 2.5644E-05 | 0.00057366 | 0.00050686 | 0.00053924 | 0.00050723 | 0.00053175 | 9.1526E-05 | 0 | 0.00017122 | 9.2159E-05 | 0.0001183 | 16 | 14 | 15 | 14 | 14.75 | 2 | 0 | 3 | 2 | 2.33333333 |
| STAT3 | Signal transd | 0.21575043 | 0.00028359 | 0.00041125 | 0.0006229 | 0.00044671 | 0.00055409 | 0.00050874 | 0 | 8.8041E-05 | 0.00010911 | 0.00013213 | 0.00010976 | 12 | 18 | 13 | 16 | 14.75 | 0 | 2 | 2 | 3 | 2.33333333 |
| NOL6 | Nucleolar prc | 0.20885869 | 1.3546E-05 | 0.0014967 | 0.0015811 | 0.0012698 | 0.0014659 | 0.00145338 | 0.00029391 | 0.00023662 | 0.00047651 | 0.00020716 | 0.00030355 | 65 | 68 | 55 | 63 | 62.75 | 10 | 8 | 13 | 7 | 9.5 |
| XRCC5 | X-ray repair c | 0.20813492 | 0.00368335 | 0.0018385 | 0.0011285 | 0.0015181 | 0.00255 | 0.00175878 | 0.00041412 | 0.00037044 | 0.0004017 | 0.00027799 | 0.00036606 | 51 | 31 | 42 | 70 | 48.5 | 9 | 8 | 7 | 6 | 7.5 |
| LAP2B | Lamina-assoc | 0.20688852 | 2.0705E-05 | 0.0012206 | 0.0012325 | 0.001049 | 0.0013509 | 0.00121325 | 0.00037094 | 0.00029864 | 0.00018505 | 0.0001494 | 0.00025101 | 52 | 48 | 54 | 59 | 53.25 | 13 | 9 | 4 | 7 | 8.25 |
| SUGP2 | SURP and G-p | 0.20653491 | 2.5877E-05 | 0.0016828 | 0.001256 | 0.001565 | 0.0013555 | 0.00146483 | 0.00024903 | 0.00037592 | 0.00027176 | 0.00031344 | 0.00030254 | 69 | 51 | 64 | 55 | 59.75 | 8 | 12 | 7 | 10 | 9.25 |
| ML12A | Myosin regul | 0.20406734 | 0.00014378 | 0.018364 | 0.014959 | 0.020115 | 0.019804 | 0.0183105 | 0.0072878 | 0.0033697 | 0.0027022 | 0.0015866 | 0.00373658 | 119 | 96 | 130 | 127 | 118 | 37 | 17 | 11 | 8 | 18.25 |
| SON | Protein SON | 0.20219675 | 7.0515E-06 | 0.00027193 | 0.00027459 | 0.00031628 | 0.00027479 | 0.0002844 | 4.1651E-05 | 6.9859E-05 | 0.00003463 | 8.3877E-05 | 5.7504E-05 | 25 | 25 | 29 | 25 | 26 | 3 | 5 | 2 | 6 | 4 |
| MYCB2 | E3 ubiquitin-p | 0.20166662 | 0.00061295 | 0.00018199 | 0.00014931 | 0.00021669 | 0.00012643 | 0.00016861 | 4.3554E-05 | 3.6525E-05 | 0 | 2.1927E-05 | 3.4002E-05 | 32 | 26 | 38 | 22 | 29.5 | 6 | 5 | 0 | 3 | 4.66666667 |
| NPM | Nucleophosm | 0.19232407 | 5.6141E-07 | 0.016066 | 0.018308 | 0.016469 | 0.015872 | 0.01667875 | 0.0033223 | 0.002767 | 0.0038577 | 0.0028839 | 0.00320773 | 179 | 202 | 183 | 175 | 184.75 | 29 | 24 | 27 | 25 | 26.25 |
| NUCL | Nucleolin OS= | 0.19165573 | 1.9794E-06 | 0.0036423 | 0.0037155 | 0.0030931 | 0.0032675 | 0.0034296 | 0.00066414 | 0.0007161 | 0.00053247 | 0.0007165 | 0.0006573 | 98 | 99 | 83 | 87 | 91.75 | 14 | 15 | 9 | 15 | 13.25 |
| E41L2 | Band 4.1-like | 0.17892398 | 4.2497E-05 | 0.0014704 | 0.0010871 | 0.0012374 | 0.0010879 | 0.0012207 | 0.00030163 | 0.00020236 | 0.00016719 | 0.00020247 | 0.00021841 | 56 | 41 | 47 | 41 | 46.25 | 9 | 6 | 4 | 6 | 6.25 |
| WDR11 | WD repeat-co | 0.1758895 | 6.6863E-06 | 0.00049586 | 0.00047894 | 0.0003891 | 0.00047928 | 0.0004608 | 5.5035E-05 | 8.3077E-05 | 0.00010296 | 8.3124E-05 | 8.1049E-05 | 23 | 22 | 18 | 22 | 21.25 | 2 | 3 | 3 | 3 | 2.75 |
| BIRC6 | Baculoviral IA | 0.1757857 | 3.2943E-05 | 0.00016299 | 0.0001591 | 0.00017977 | 0.00013176 | 0.00015841 | 3.4673E-05 | 2.7915E-05 | 0 | 2.0948E-05 | 2.7845E-05 | 30 | 29 | 33 | 24 | 29 | 5 | 4 | 0 | 3 | 4 |
| SMHD1 | Structural ma | 0.17510574 | 1.2592E-07 | 0.0033561 | 0.0032561 | 0.0032067 | 0.003019 | 0.00320948 | 0.00047037 | 0.0006086 | 0.00071232 | 0.0004567 | 0.000562 | 255 | 245 | 243 | 227 | 242.5 | 28 | 36 | 34 | 27 | 31.25 |
| C1QBP | Complement | 0.17399344 | 0.00018253 | 0.0040238 | 0.0030237 | 0.0033777 | 0.0047279 | 0.00378828 | 0.00047775 | 0.00084138 | 0.00059583 | 0.00072158 | 0.00065914 | 43 | 32 | 36 | 50 | 40.25 | 4 | 7 | 4 | 6 | 5.25 |
| PSPC1 | Paraspeckle c | 0.17223183 | 8.4213E-05 | 0.001665 | 0.0021399 | 0.0015683 | 0.0021414 | 0.00187865 | 0.0002576 | 0.00038886 | 0 | 0.00032423 | 0.00032356 | 33 | 42 | 31 | 42 | 37 | 4 | 6 | 0 | 5 | 5 |
| RRBP1 | Ribosome-bir | 0.17113747 | 0.00026134 | 0.00069246 | 0.00090711 | 0.00061925 | 0.00058627 | 0.00070127 | 9.5551E-05 | 0.00016828 | 0 | 9.6211E-05 | 0.00012001 | 37 | 48 | 33 | 31 | 37.25 | 4 | 7 | 0 | 4 | 5 |
| EPS15 | Epidermal gr | 0.16998906 | 2.126E-06 | 0.00047122 | 0.00041635 | 0.00044295 | 0.00044641 | 0.00044423 | 7.5182E-05 | 0.00007566 | 0 | 7.5702E-05 | 7.5515E-05 | 16 | 14 | 15 | 15 | 15 | 2 | 2 | 0 | 2 | 2 |
| CPSF1 | Cleavage and | 0.16796159 | 2.233E-06 | 0.00080464 | 0.00084944 | 0.00086179 | 0.00083157 | 0.00083686 | 0.00018673 | 0.00014094 | 0 | 9.4011E-05 | 0.00014056 | 44 | 46 | 47 | 45 | 45.5 | 8 | 6 | 0 | 4 | 6 |
| E41L3 | Band 4.1-like | 0.16722929 | 4.1632E-05 | 0.00043697 | 0.00039222 | 0.00034077 | 0.00031891 | 0.00037222 | 6.1972E-05 | 6.2365E-05 | 0 | 0.0000624 | 6.2246E-05 | 19 | 16 | 14 | 14 | 15.75 | 2 | 2 | 0 | 2 | 2 |
| THOC2 | THO complex | 0.1618808 | 1.0054E-06 | 0.00094422 | 0.00097018 | 0.0010132 | 0.00083696 | 0.00094114 | 0.00012686 | 0.00014894 | 0.00018458 | 0.00014903 | 0.00015235 | 57 | 58 | 61 | 50 | 56.5 | 6 | 7 | 7 | 7 | 6.75 |
| MYH9 | Myosin-9 OS= | 0.15370067 | 9.5472E-05 | 0.011175 | 0.0097749 | 0.012905 | 0.014843 | 0.01217448 | 0.0022684 | 0.0016948 | 0.0022932 | 0.0012285 | 0.00187123 | 830 | 719 | 956 | 1091 | 899 | 132 | 98 | 107 | 71 | 102 |
| KI67 | Proliferation | 0.13981752 | 1.7259E-06 | 0.0016614 | 0.0018741 | 0.0016252 | 0.0017772 | 0.00173448 | 0.00020689 | 0.00029149 | 0 | 0.00022915 | 0.00024251 | 205 | 229 | 200 | 217 | 212.75 | 20 | 28 | 0 | 22 | 23.3333333 |
| RBM25 | RNA-binding | 0.13799544 | 5.6163E-05 | 0.0011895 | 0.0012644 | 0.00084743 | 0.001202 | 0.00112583 | 0.00015982 | 0.00020104 | 9.9659E-05 | 0.00016092 | 0.00015536 | 38 | 40 | 27 | 38 | 35.75 | 4 | 5 | 2 | 4 | 3.75 |
| PRP8 | Pre-mRNA-pr | 0.13650283 | 4.6655E-06 | 0.00055376 | 0.00051353 | 0.00063456 | 0.00052532 | 0.00055679 | 4.3274E-05 | 7.2582E-05 | 7.1959E-05 | 0.0001162 | 7.6004E-05 | 49 | 45 | 56 | 46 | 49 | 3 | 5 | 4 | 8 | 5 |
| PRP19 | Pre-mRNA-pr | 0.13219452 | 5.284E-06 | 0.0023561 | 0.0021148 | 0.0018899 | 0.0020105 | 0.00209283 | 0.00026731 | 0.00047077 | 0.00016669 | 0.00020187 | 0.00027666 | 45 | 40 | 36 | 38 | 39.75 | 4 | 7 | 2 | 3 | 4 |
| DHX15 | Pre-mRNA-sp | 0.1235395 | 0.00025733 | 0.0009626 | 0.00063683 | 0.0011316 | 0.00093917 | 0.00091755 | 0.00016947 | 8.5272E-05 | 0 | 8.5319E-05 | 0.00011335 | 29 | 19 | 34 | 28 | 27.5 | 4 | 2 | 0 | 2 | 2.66666667 |
| LAP2A | Lamina-assoc | 0.12252551 | 7.7441E-07 | 0.0020913 | 0.001843 | 0.002135 | 0.0020748 | 0.00203603 | 0.00038826 | 0.0002442 | 0.00012106 | 0.00024434 | 0.00024947 | 55 | 48 | 56 | 54 | 53.25 | 8 | 5 | 2 | 5 | 5 |
| HCFC1 | Host cell fact | 0.12024175 | 1.3002E-05 | 0.0017636 | 0.0016237 | 0.0017422 | 0.0013235 | 0.00161325 | 0.00018206 | 0.00016656 | 0 | 0.00023332 | 0.00019398 | 136 | 124 | 134 | 101 | 123.75 | 11 | 10 | 0 | 14 | 11.6666667 |
| CPSF2 | Cleavage and | 0.11709696 | 2.9671E-05 | 0.0010798 | 0.00081779 | 0.00091354 | 0.0011253 | 0.00098411 | 0.00017228 | 0.00008669 | 0 | 8.6738E-05 | 0.00011524 | 32 | 24 | 27 | 33 | 29 | 4 | 2 | 0 | 2 | 2.66666667 |
| MGAP | MAX gene-as | 0.1124412 | 9.9259E-06 | 0.00064532 | 0.00050193 | 0.00065578 | 0.0005816 | 0.00059616 | 8.9046E-05 | 4.4806E-05 | 0 | 6.7246E-05 | 6.7033E-05 | 74 | 57 | 75 | 66 | 68 | 8 | 4 | 0 | 6 | 6 |

| | | | | | | | | | | | | | | | | | | | | | | | |
|---|---|---|---|---|---|---|---|---|---|---|---|---|---|---|---|---|---|---|---|---|---|---|---|
| SF3B1 | Splicing facto | 0.10643138 | 6.0599E-08 | 0.0020844 | 0.0020639 | 0.0021711 | 0.0020858 | 0.0021013 | 0.00028412 | 0.00025994 | 6.4427E-05 | 0.00028609 | 0.00022364 | 103 | 101 | 107 | 102 | 103.25 | 11 | 10 | 2 | 11 | 8.5 |
| MYH10 | Myosin-10 O | 0.09525846 | 0.00023056 | 0.0049144 | 0.0044231 | 0.0060925 | 0.0073411 | 0.00569278 | 0.00046022 | 0.00061753 | 0.00076529 | 0.0003261 | 0.00054229 | 412 | 374 | 523 | 625 | 483.5 | 34 | 45 | 47 | 28 | 38.5 |
| RIF1 | Telomere-ass | 0.08906358 | 1.3791E-05 | 0.0014625 | 0.0011318 | 0.0015413 | 0.001316 | 0.0013629 | 0.0001635 | 9.5983E-05 | 3.3986E-05 | 0.00019207 | 0.00012138 | 137 | 105 | 144 | 122 | 127 | 12 | 7 | 2 | 14 | 8.75 |
| TASOR | Protein TASC | 0.08831837 | 2.0411E-06 | 0.00060046 | 0.00060633 | 0.0007288 | 0.00065466 | 0.00064756 | 8.0674E-05 | 4.0594E-05 | 5.0307E-05 | 0 | 5.7192E-05 | 38 | 38 | 46 | 41 | 40.75 | 4 | 2 | 2 | 0 | 2.66666667 |
| XRCC6 | X-ray repair c | 0.0816509 | 0.0009643 | 0.0016032 | 0.0010501 | 0.0018247 | 0.0023207 | 0.00169968 | 0 | 0.00011132 | 0.00013795 | 0.00016707 | 0.00013878 | 37 | 24 | 42 | 53 | 39 | 0 | 2 | 2 | 3 | 2.33333333 |
| MYL6 | Myosin light | 0.07987353 | 0.00012658 | 0.0068156 | 0.007941 | 0.0099877 | 0.010949 | 0.00892333 | 0.00066917 | 0.00067342 | 0.00083456 | 0.0006738 | 0.00071274 | 48 | 52 | 67 | 73 | 60 | 3 | 4 | 4 | 5 | 4 |
| SR140 | U2 snRNP-as | 0.06857573 | 3.7141E-06 | 0.0026927 | 0.0025119 | 0.0028027 | 0.0021509 | 0.00253955 | 0.00026186 | 0.00013176 | 0.00020411 | 9.8876E-05 | 0.00017415 | 105 | 97 | 109 | 83 | 98.5 | 8 | 4 | 5 | 3 | 5 |
| CHD4 | Chromodom | 0.06599888 | 8.6363E-09 | 0.00080049 | 0.00086406 | 0.00085797 | 0.00082284 | 0.00083634 | 7.0464E-05 | 7.0911E-05 | 0.00004394 | 3.5475E-05 | 5.5198E-05 | 58 | 62 | 62 | 59 | 60.25 | 4 | 4 | 2 | 2 | 3 |
| ELYS | Protein ELYS | 0.05993616 | 1.8809E-07 | 0.0049609 | 0.0045861 | 0.0044837 | 0.0050013 | 0.004758 | 0.0005351 | 0.000359 | 3.7075E-05 | 0.00020953 | 0.00028518 | 426 | 390 | 384 | 425 | 406.25 | 36 | 24 | 2 | 14 | 19 |
| SF3B6 | Splicing facto | 0.55877168 | 0.00589131 | 0.0014778 | 0.0019185 | 0.0016934 | 0.0017066 | 0.00169908 | 0 | 0.00081349 | 0 | 0.0010853 | 0.0009494 | 7 | 9 | 8 | 8 | 8 | 0 | 3 | 0 | 4 | 3.5 |
| IFT25 | Intraflagellar | 1.59773094 | 0.61247964 | 0.00036651 | 0 | 0 | 0.00037036 | 0.00036844 | 0 | 0.00047077 | 0 | 0.00070655 | 0.00058866 | 2 | 0 | 0 | 2 | 2 | 0 | 2 | 0 | 3 | 2.5 |
| ROA3 | Heterogeneo | 3.18739339 | 0.41652111 | 0 | 0.00014099 | 0.00020999 | 0.00028218 | 0.00021105 | 0 | 0.00071737 | 0 | 0.00062805 | 0.00067271 | 0 | 2 | 3 | 4 | 3 | 0 | 9 | 0 | 8 | 8.5 |
| RL10A | 60S ribosoma | 0.16265865 | 0.00015717 | 0.0025537 | 0.0035611 | 0.0023167 | 0.0030721 | 0.0028759 | 0.00031043 | 0 | 0 | 0.00062515 | 0.00046779 | 21 | 29 | 19 | 25 | 23.5 | 2 | 0 | 0 | 4 | 3 |
| TRA2B | Transformer- | 0.55211721 | 0.0437112 | 0.0013744 | 0.00055514 | 0.0009187 | 0.00055553 | 0.00085094 | 0.00035085 | 0 | 0 | 0.00058879 | 0.00046982 | 15 | 6 | 10 | 6 | 9.25 | 3 | 0 | 0 | 5 | 4 |
| CENPM | Centromere p | 1.42988899 | 0.94788741 | 0.0002932 | 0.00029607 | 0 | 0.00059257 | 0.00039395 | 0.00056136 | 0 | 0 | 0.00056524 | 0.0005633 | 2 | 2 | 0 | 4 | 2.66666667 | 3 | 0 | 0 | 3 | 3 |
| RL7L | 60S ribosoma | 1.03383577 | 0.57900801 | 0.00053635 | 0.00043328 | 0.00043022 | 0 | 0.00046662 | 0 | 0.00041336 | 0 | 0.00055145 | 0.00048241 | 5 | 4 | 4 | 0 | 4.33333333 | 0 | 3 | 0 | 4 | 3.5 |
| SMD3 | Small nuclea | 1.90313533 | 0.94711563 | 0.0006283 | 0.00084592 | 0.00041998 | 0.00042326 | 0.00057937 | 0 | 0 | 0.0016669 | 0.00053832 | 0.00110261 | 3 | 4 | 2 | 2 | 2.75 | 0 | 0 | 5 | 2 | 3.5 |
| RAB2A | Ras-related p | 2.1577626 | 0.26716942 | 0.00024895 | 0 | 0 | 0 | 0.00024895 | 0 | 0 | 0.00059443 | 0.00047992 | 0.00053718 | 2 | 0 | 0 | 0 | 2 | 0 | 0 | 3 | 3 | 3 |
| CAZA2 | F-actin-cappi | 1.90184609 | 0.29356747 | 0 | 0.00027951 | 0 | 0 | 0.00027951 | 0.00058884 | 0 | 0 | 0.00047433 | 0.00053159 | 0 | 4 | 0 | 0 | 4 | 6 | 0 | 0 | 5 | 5.5 |
| NDUAD | NADH dehyd | 0.85541972 | 0.85535305 | 0.00054976 | 0 | 0.00055122 | 0 | 0.00055049 | 0 | 0.00047077 | 0 | 0.00047103 | 0.0004709 | 3 | 0 | 3 | 0 | 3 | 0 | 2 | 0 | 2 | 2 |
| RBMX | RNA-binding | 1.5712493 | 0.94935523 | 0.00033745 | 0 | 0.00020301 | 0.0002046 | 0.00024835 | 0 | 0.00034676 | 0 | 0.00043369 | 0.00039023 | 5 | 0 | 3 | 3 | 3.66666667 | 0 | 4 | 0 | 5 | 4.5 |
| EXOS5 | Exosome con | 0.71018837 | 0.04539831 | 0.00067375 | 0.00034017 | 0.00045036 | 0.00056735 | 0.00050791 | 0 | 0.00028847 | 0 | 0.00043295 | 0.00036071 | 6 | 3 | 4 | 5 | 4.5 | 0 | 2 | 0 | 3 | 2.5 |
| HAP28 | 28 kDa heat- | 1.26851207 | 0.44820466 | 0.00029158 | 0.00058887 | 0.00058472 | 0.0005893 | 0.00051362 | 0 | 0 | 0.00092831 | 0.00037475 | 0.00065153 | 2 | 4 | 4 | 4 | 3.5 | 0 | 0 | 4 | 2 | 3 |
| EXOS8 | Exosome con | 0.38357298 | 0.00260026 | 0.0010517 | 0.00077236 | 0.0012462 | 0.00077292 | 0.0009608 | 0 | 0.00036843 | 0 | 0.00036864 | 0.00036854 | 11 | 8 | 13 | 8 | 10 | 0 | 3 | 0 | 3 | 3 |
| JUPI2 | Jupiter micro | 0.95605055 | 0.50865871 | 0 | 0.00056098 | 0.00041777 | 0.00042104 | 0.0004666 | 0 | 0.00053519 | 0 | 0.00035699 | 0.00044609 | 0 | 4 | 3 | 3 | 3.33333333 | 0 | 3 | 0 | 2 | 2.5 |
| RAB9A | Ras-related p | 0.85428325 | 0.63193733 | 0 | 0 | 0.00039491 | 0 | 0.00039491 | 0 | 0.00033727 | 0 | 0.00033746 | 0.00033737 | 0 | 0 | 3 | 0 | 3 | 0 | 2 | 0 | 2 | 2 |
| SPF27 | Pre-mRNA-sp | 0.23675785 | 0.00133617 | 0.00093825 | 0.0017764 | 0.0011759 | 0.0011851 | 0.00126891 | 0.00029939 | 0 | 0 | 0.00030146 | 0.00030043 | 8 | 15 | 10 | 10 | 10.75 | 2 | 0 | 0 | 2 | 2 |
| ATX10 | Ataxin-10 OS | 1.37353727 | 0.72568401 | 0 | 0.00016829 | 0 | 0.00016841 | 0.00016835 | 0 | 0 | 0.00017687 | 0.0002856 | 0.00023124 | 0 | 3 | 0 | 3 | 3 | 0 | 0 | 2 | 4 | 3 |
| NDUS3 | NADH dehyd | 1.21246697 | 0.46350023 | 0 | 0.0003028 | 0 | 0 | 0.0003028 | 0 | 0 | 0.00047734 | 0.00025693 | 0.00036714 | 0 | 3 | 0 | 0 | 3 | 0 | 0 | 3 | 2 | 2.5 |
| SUPT3 | Transcription | 0.33511368 | 0.00146323 | 0.00046295 | 0.0008014 | 0.00059681 | 0.00066831 | 0.00063237 | 0.00016883 | 0 | 0 | 0.000255 | 0.00021192 | 7 | 12 | 9 | 10 | 9.5 | 2 | 0 | 0 | 3 | 2.5 |
| RT27 | 28S ribosoma | 1.28356946 | 0.78121751 | 0.00019122 | 0 | 0.00012782 | 0 | 0.00015952 | 0 | 0.00016375 | 0 | 0.00024576 | 0.00020476 | 3 | 0 | 2 | 0 | 2.5 | 0 | 2 | 0 | 3 | 2.5 |
| ATPF2 | ATP synthase | 1.28138824 | 0.41517911 | 0 | 0 | 0.00018311 | 0 | 0.00018311 | 0 | 0.00023457 | 0 | 0.0002347 | 0.00023464 | 0 | 0 | 2 | 0 | 2 | 0 | 2 | 0 | 2 | 2 |
| TBB2B | Tubulin beta- | 1.58658985 | 0.35167747 | 0 | 0.00011976 | 0 | 0 | 0.00011976 | 0.00015138 | 0 | 0 | 0.00022864 | 0.00019001 | 0 | 72 | 0 | 0 | 72 | 89 | 0 | 0 | 88 | 88.5 |
| OST48 | Dolichyl-diph | 1.90608483 | 0.93678065 | 0.00011574 | 0.00011687 | 0.00011605 | 0.00017543 | 0.00013102 | 0 | 0 | 0.00027636 | 0.00022312 | 0.00024974 | 2 | 2 | 2 | 3 | 2.25 | 0 | 0 | 3 | 3 | 3 |
| SMAP1 | Stromal mem | 0.79975473 | 0.35131133 | 0.00022602 | 0.00022824 | 0.00022663 | 0 | 0.00022696 | 0 | 0.00014516 | 0 | 0.00021787 | 0.00018152 | 4 | 4 | 4 | 0 | 4 | 0 | 2 | 0 | 3 | 2.5 |
| PURA | Transcription | 1.60170987 | 0.34836926 | 0 | 0 | 0.00016434 | 0 | 0.00016434 | 0 | 0.0003158 | 0 | 0.00021065 | 0.00026323 | 0 | 0 | 2 | 0 | 2 | 0 | 3 | 0 | 2 | 2.5 |
| NUP54 | Nucleoporin | 0.45513778 | 0.00131998 | 0.00036434 | 0.0003679 | 0.00036531 | 0.00036816 | 0.00036643 | 0.00013287 | 0 | 0 | 0.00020068 | 0.00016678 | 7 | 7 | 7 | 7 | 7 | 2 | 0 | 0 | 3 | 2.5 |
| TXLNG | Gamma-taxil | 1.27469553 | 0.78705132 | 0 | 0.00010093 | 0.00015033 | 0 | 0.00012563 | 0.00012758 | 0 | 0 | 0.0001927 | 0.00016014 | 0 | 2 | 3 | 0 | 2.5 | 2 | 0 | 0 | 3 | 2.5 |
| LYAR | Cell growth-r | 0.50966216 | 0.00430418 | 0.00041776 | 0.00035154 | 0.00034906 | 0.00028143 | 0.00034995 | 0.00017774 | 0 | 0 | 0.00017897 | 0.00017836 | 6 | 5 | 5 | 4 | 5 | 2 | 0 | 0 | 2 | 2 |
| RPAP2 | Putative RNA | 1.91026328 | 0.93779938 | 8.6237E-05 | 0.00008708 | 8.6466E-05 | 8.7143E-05 | 8.6732E-05 | 0.00016511 | 0 | 0 | 0.00016625 | 0.00016568 | 2 | 2 | 2 | 2 | 2 | 3 | 0 | 0 | 3 | 3 |
| FMR1 | Synaptic func | 1.90727025 | 0.28874533 | 0 | 0 | 0 | 8.4385E-05 | 8.4385E-05 | 0 | 0.0001609 | 0 | 0.00016099 | 0.00016095 | 0 | 0 | 0 | 3 | 3 | 0 | 5 | 0 | 5 | 5 |
| HNRPR | Heterogeneo | 0.98318711 | 0.14582031 | 0.00016675 | 0.00012629 | 0.0001254 | 0.00012638 | 0.00013621 | 0 | 0.0001071 | 0 | 0.00016073 | 0.00013392 | 6 | 6 | 5 | 4 | 5.25 | 0 | 6 | 0 | 9 | 7.5 |
| RBBP7 | Histone-bind | 0.24933196 | 0.00020473 | 0.0006209 | 0.00050158 | 0.00074707 | 0.00069017 | 0.00063993 | 0 | 0.00015951 | 0 | 0.0001596 | 0.00015956 | 10 | 8 | 12 | 11 | 10.25 | 0 | 2 | 0 | 2 | 2 |
| ORC4 | Origin recogr | 1.28075176 | 0.41539116 | 0.00012105 | 0 | 0 | 0 | 0.00012105 | 0.0001545 | 0 | 0 | 0.00015557 | 0.00015504 | 2 | 0 | 0 | 0 | 2 | 2 | 0 | 0 | 2 | 2 |
| WIPI2 | WD repeat d | 1.46760389 | 0.68522703 | 0.00017437 | 0 | 0.00011656 | 0 | 0.00014547 | 0 | 0 | 0.00027757 | 0.0001494 | 0.00021349 | 3 | 0 | 2 | 0 | 2.5 | 0 | 0 | 3 | 2 | 2.5 |
| CHERP | Calcium hom | 0.12984077 | 1.3439E-05 | 0.0012099 | 0.0010182 | 0.0012998 | 0.0010189 | 0.0011367 | 0.00014708 | 0 | 0 | 0.0001481 | 0.00014759 | 42 | 35 | 45 | 35 | 39.25 | 4 | 0 | 0 | 4 | 4 |
| CPVL | Probable seri | 0.74882067 | 0.04688456 | 0.00027719 | 0.00016794 | 0.00022234 | 0.0002801 | 0.00023689 | 0.00021228 | 0 | 0 | 0.0001425 | 0.00017739 | 5 | 3 | 4 | 5 | 4.25 | 3 | 0 | 0 | 2 | 2.5 |
| TFCP2 | Alpha-globin | 0.60580958 | 0.0140585 | 0.00021027 | 0.00031848 | 0.00031624 | 0.00026559 | 0.00027765 | 0.00020128 | 0 | 0 | 0.00013512 | 0.0001682 | 4 | 6 | 6 | 5 | 5.25 | 3 | 0 | 0 | 2 | 2.5 |
| SRP54 | Signal recogr | 0.30092093 | 0.00152047 | 0.00041886 | 0.00068731 | 0.00041998 | 0.00047617 | 0.00050058 | 0 | 0 | 0.00016669 | 0.00013458 | 0.00015064 | 8 | 13 | 8 | 9 | 9.5 | 0 | 0 | 2 | 2 | 2 |
| SEC63 | Translocatior | 1.22069246 | 0.81928597 | 0 | 0.00014024 | 0 | 0.00010526 | 0.00012275 | 0 | 0 | 0.00016581 | 0.00013387 | 0.00014984 | 0 | 4 | 0 | 3 | 3.5 | 0 | 0 | 3 | 3 | 3 |
| GCR | Glucocorticoi | 0.23217928 | 0.00031392 | 0.00047547 | 0.0005487 | 0.00034052 | 0.00051478 | 0.00046987 | 0 | 8.7247E-05 | 0 | 0.00013094 | 0.00010909 | 14 | 16 | 10 | 15 | 13.75 | 0 | 2 | 0 | 3 | 2.5 |
| RINT1 | RAD50-intera | 0.84880023 | 0.09099813 | 9.9956E-05 | 0.00013458 | 0.00010022 | 0.00016834 | 0.00012577 | 8.5054E-05 | 0 | 0 | 0.00012846 | 0.00010676 | 3 | 4 | 3 | 5 | 3.75 | 2 | 0 | 0 | 3 | 2.5 |
| CPSF6 | Cleavage and | 0.30424248 | 0.00023994 | 0.0005747 | 0.00053196 | 0.00048019 | 0.00043555 | 0.0005056 | 0 | 0.00018455 | 0 | 0.0001231 | 0.00015383 | 12 | 11 | 10 | 9 | 10.5 | 0 | 3 | 0 | 2 | 2.5 |
| GLYR1 | Putative oxid | 0.08561373 | 0.0001331 | 0.0017179 | 0.001301 | 0.0010526 | 0.0016395 | 0.00142775 | 0.00012181 | 0 | 0 | 0.00012266 | 0.00012224 | 36 | 27 | 22 | 34 | 29.75 | 2 | 0 | 0 | 2 | 2 |
| MATR3 | Matrin-3 OS= | 0.28731977 | 0.0003497 | 0.00049848 | 0.00056628 | 0.00037486 | 0.00050372 | 0.00048584 | 0.00015906 | 0 | 0 | 0.00012012 | 0.00013959 | 16 | 18 | 12 | 16 | 15.5 | 4 | 0 | 0 | 3 | 3.5 |
| FIP1 | Pre-mRNA 3' | 0.17429047 | 5.3691E-05 | 0.00075522 | 0.00071775 | 0.00080178 | 0.00098762 | 0.00081559 | 0.00017011 | 0 | 0 | 0.00011419 | 0.00014215 | 17 | 16 | 18 | 22 | 18.25 | 3 | 0 | 0 | 2 | 2.5 |
| SF3B3 | Splicing facto | 0.10069804 | 2.9869E-06 | 0.0011709 | 0.0012261 | 0.0010218 | 0.0010079 | 0.00110668 | 0 | 0.00011141 | 0 | 0.00011147 | 0.00011144 | 54 | 56 | 47 | 46 | 50.75 | 0 | 4 | 0 | 4 | 4 |
| SYNRG | Synergin gan | 0.58800069 | 0.01633518 | 0.00010041 | 0.00014195 | 0.00012082 | 0.00016235 | 0.00013138 | 5.1266E-05 | 0 | 0 | 0.00010324 | 7.7253E-05 | 5 | 7 | 6 | 8 | 6.5 | 2 | 0 | 0 | 4 | 3 |
| FANCI | Fanconi aner | 0.57524891 | 0.00915248 | 0.00011922 | 0.00016052 | 0.00017931 | 0.00016064 | 0.00015492 | 7.6088E-05 | 0 | 0 | 0.00010215 | 8.9119E-05 | 6 | 8 | 9 | 8 | 7.75 | 3 | 0 | 0 | 4 | 3.5 |
| PEAK1 | Pseudopodiu | 1.27735164 | 0.43250982 | 9.0682E-05 | 9.1569E-05 | 3.0308E-05 | 0.00006109 | 6.8412E-05 | 0 | 7.7653E-05 | 0 | 0.00009712 | 8.7387E-05 | 6 | 6 | 2 | 4 | 4.5 | 0 | 4 | 0 | 5 | 4.5 |
| NOL9 | Polynucleotic | 0.54495799 | 0.00737642 | 0.00018795 | 0.00015183 | 0.00022614 | 0.00022791 | 0.00019846 | 0 | 0 | 0.00011968 | 9.6622E-05 | 0.00010815 | 5 | 4 | 6 | 6 | 5.25 | 0 | 0 | 2 | 2 | 2 |
| IPO4 | Importin-4 O | 2.24240072 | 0.26339963 | 0 | 0 | 4.8952E-05 | 0 | 4.8952E-05 | 0 | 0.00012542 | 0 | 0.00009412 | 0.00010977 | 0 | 0 | 2 | 0 | 2 | 0 | 4 | 0 | 3 | 3.5 |
| TRM1L | TRMT1-like p | 0.50919908 | 0.00451051 | 0.00018 | 0.00018176 | 0.00014439 | 0.00021827 | 0.00018111 | 9.1901E-05 | 0 | 0 | 9.2536E-05 | 9.2219E-05 | 5 | 5 | 4 | 6 | 5 | 2 | 0 | 0 | 2 | 2 |
| BAZ1B | Tyrosine-pro | 0.23201647 | 0.00372454 | 0.0002847 | 0.00023358 | 0.00044603 | 0.00021577 | 0.00029502 | 4.5424E-05 | 0 | 0 | 9.1475E-05 | 6.845E-05 | 16 | 13 | 25 | 12 | 16.5 | 2 | 0 | 0 | 4 | 3 |
| DNMBP | Dynamin-bin | 1.90340943 | 0.32713958 | 0 | 0 | 0 | 3.3818E-05 | 3.3818E-05 | 4.2716E-05 | 0 | 0 | 8.6023E-05 | 6.437E-05 | 0 | 0 | 0 | 2 | 2 | 2 | 0 | 0 | 4 | 3 |
| NOM1 | Nucleolar MI | 0.85208539 | 0.85182729 | 9.2053E-05 | 0 | 0 | 0.00009302 | 9.2537E-05 | 0 | 7.8827E-05 | 0 | 7.8871E-05 | 7.8849E-05 | 3 | 0 | 0 | 3 | 3 | 0 | 2 | 0 | 2 | 2 |
| GAK | Cyclin-G-assc | 1.27457326 | 0.39919781 | 6.0385E-05 | 6.0976E-05 | 4.0364E-05 | 0.00004068 | 5.0601E-05 | 5.1383E-05 | 0 | 0 | 7.7607E-05 | 6.4495E-05 | 3 | 3 | 2 | 2 | 2.5 | 2 | 0 | 0 | 3 | 2.5 |
| N4BP2 | NEDD4-bindi | 1.01885664 | 0.21318099 | 0.00013418 | 0.00010538 | 7.4742E-05 | 6.0261E-05 | 9.3641E-05 | 0.00011417 | 0 | 0 | 7.6643E-05 | 9.5407E-05 | 9 | 7 | 5 | 4 | 6.25 | 6 | 0 | 0 | 4 | 5 |
| ACINU | Apoptotic ch | 0.32517432 | 0.00118277 | 0.00025582 | 0.00019871 | 0.00029596 | 0.00033804 | 0.00027213 | 0 | 0.00010111 | 0 | 7.5871E-05 | 8.8491E-05 | 13 | 10 | 15 | 17 | 13.75 | 0 | 4 | 0 | 3 | 3.5 |
| PAXB1 | PAX3- and PA | 0.16975014 | 4.7236E-05 | 0.00043165 | 0.00040682 | 0.0003751 | 0.00052343 | 0.00043425 | 0.00007346 | 0 | 0 | 7.3968E-05 | 7.3714E-05 | 15 | 14 | 13 | 18 | 15 | 2 | 0 | 0 | 2 | 2 |
| AAK1 | AP2-associat | 0.91255335 | 0.10577764 | 0.00010984 | 0.00011091 | 8.2597E-05 | 8.3243E-05 | 9.6648E-05 | 0 | 0.00010581 | 0 | 7.0582E-05 | 8.8196E-05 | 4 | 4 | 3 | 3 | 3.5 | 0 | 3 | 0 | 2 | 2.5 |
| DPOD1 | DNA polymei | 0.49747842 | 0.01107809 | 9.5351E-05 | 0.00012035 | 0.00019121 | 0.00014453 | 0.00013786 | 0 | 0 | 7.5892E-05 | 6.1273E-05 | 6.8583E-05 | 4 | 5 | 8 | 6 | 5.75 | 0 | 0 | 2 | 2 | 2 |
| ZN638 | Zinc finger pr | 0.03371109 | 3.7489E-06 | 0.0013875 | 0.0012528 | 0.0010567 | 0.0013751 | 0.00126803 | 3.4056E-05 | 0 | 0 | 5.1437E-05 | 4.2747E-05 | 104 | 93 | 79 | 102 | 94.5 | 2 | 0 | 0 | 3 | 2.5 |
| NU153 | Nuclear pore | 0.09263491 | 6.8806E-05 | 0.00094819 | 0.00097553 | 0.00062783 | 0.000922 | 0.00086839 | 0 | 0.0001149 | 0 | 4.5986E-05 | 8.0443E-05 | 53 | 54 | 35 | 51 | 48.25 | 0 | 5 | 0 | 2 | 3.5 |
| ZFY16 | Zinc finger FY | 0.23225755 | 4.7874E-05 | 0.00017146 | 0.00017314 | 0.00020631 | 0.00020792 | 0.00018971 | 0 | 4.4049E-05 | 0 | 4.4073E-05 | 4.4061E-05 | 10 | 10 | 12 | 12 | 11 | 0 | 2 | 0 | 2 | 2 |
| BAZ1A | Bromodomai | 0.31936855 | 0.00071826 | 0.00018655 | 0.000137 | 0.00015304 | 0.00020565 | 0.00017056 | 0 | 6.5351E-05 | 0 | 4.3592E-05 | 5.4472E-05 | 11 | 8 | 9 | 12 | 10 | 0 | 3 | 0 | 2 | 2.5 |
| P3C2A | Phosphatidyl | 0.63549113 | 0.23383885 | 0 | 4.7414E-05 | 6.2773E-05 | 0.00007908 | 6.3089E-05 | 3.9954E-05 | 0 | 0 | 4.0231E-05 | 4.0093E-05 | 0 | 3 | 4 | 5 | 4 | 2 | 0 | 0 | 2 | 2 |
| MDN1 | Midasin OS=I | 0.51054014 | 0.17531894 | 0 | 1.9047E-05 | 3.3097E-05 | 0.00001906 | 2.3735E-05 | 0 | 1.2114E-05 | 0 | 1.2121E-05 | 1.2118E-05 | 0 | 4 | 7 | 4 | 5 | 0 | 2 | 0 | 2 | 2 |
| TRIO | Triple functic | 1.06769815 | 0.51125601 | 2.5562E-05 | 0 | 0 | 0 | 2.5562E-05 | 2.1751E-05 | 3.2834E-05 | 0 | 0 | 2.7293E-05 | 3 | 0 | 0 | 0 | 3 | 2 | 3 | 0 | 0 | 2.5 |
| RGPD8 | RANBP2-like | 1.26712669 | 0.41991236 | 0 | 0 | 0 | 3.0216E-05 | 3.0216E-05 | 3.8166E-05 | 3.8409E-05 | 0 | 0 | 3.8288E-05 | 0 | 0 | 0 | 46 | 46 | 64 | 53 | 0 | 0 | 58.5 |
| GSK3A | Glycogen syn | 1.28044539 | 0.41549102 | 0.0001639 | 0 | 0 | 0 | 0.0001639 | 0.0002092 | 0.00021053 | 0 | 0 | 0.00020987 | 3 | 0 | 0 | 0 | 3 | 3 | 4 | 0 | 0 | 3.5 |
| SYDM | Aspartate--tF | 1.4202118 | 0.37886948 | 0 | 8.2625E-05 | 0 | 0 | 8.2625E-05 | 0.00010444 | 0 | 0.00013025 | 0 | 0.00011735 | 0 | 2 | 0 | 0 | 2 | 2 | 0 | 2 | 0 | 2 |
| KIF22 | Kinesin-like p | 1.49057473 | 0.36503201 | 0 | 0 | 0.00011936 | 0 | 0.00011936 | 0.00015195 | 0.00020388 | 0 | 0 | 0.00017792 | 0 | 0 | 3 | 0 | 3 | 3 | 4 | 0 | 0 | 3.5 |
| SNX18 | Sorting nexin | 1.58606427 | 0.35173953 | 0 | 8.4861E-05 | 0 | 0 | 8.4861E-05 | 0.00010727 | 0.00016192 | 0 | 0 | 0.0001346 | 0 | 2 | 0 | 0 | 2 | 2 | 3 | 0 | 0 | 2.5 |
| Z3H7A | Zinc finger CC | 2.95874986 | 0.21640206 | 0 | 8.2327E-05 | 0 | 0 | 8.2327E-05 | 0.00024281 | 0.00024436 | 0 | 0 | 0.00024359 | 0 | 3 | 0 | 0 | 3 | 7 | 7 | 0 | 0 | 7 |
| LARP4 | La-related pr | 2.55150179 | 0.44469575 | 7.2896E-05 | 0 | 0 | 7.3662E-05 | 7.3279E-05 | 9.3043E-05 | 0.0002809 | 0 | 0 | 0.00018697 | 2 | 0 | 0 | 2 | 2 | 2 | 6 | 0 | 0 | 4 |
| CLH1 | Clathrin heav | 1.95658894 | 0.51553074 | 0 | 4.7725E-05 | 4.7389E-05 | 0 | 4.7557E-05 | 0 | 6.0709E-05 | 0.00012539 | 0 | 9.305E-05 | 0 | 3 | 3 | 0 | 3 | 0 | 3 | 5 | 0 | 4 |
| 2A5G | Serine/threo | 1.75076573 | 0.54546476 | 0.00010072 | 0.0001017 | 0 | 0 | 0.00010121 | 0 | 0.00019406 | 0.00016033 | 0 | 0.0001772 | 2 | 2 | 0 | 0 | 2 | 0 | 3 | 2 | 0 | 2.5 |
| PI3R4 | Phosphoinos | 1.27826901 | 0.78448545 | 5.8295E-05 | 0 | 3.8967E-05 | 0 | 4.8631E-05 | 7.4407E-05 | 0.00004992 | 0 | 0 | 6.2164E-05 | 3 | 0 | 2 | 0 | 2.5 | 3 | 2 | 0 | 0 | 2.5 |
| COX2 | Cytochrome | 1.22240282 | 0.81824371 | 0.00034875 | 0.00046954 | 0 | 0 | 0.00040915 | 0.00044513 | 0 | 0.00055515 | 0 | 0.00050014 | 3 | 4 | 0 | 0 | 3.5 | 3 | 0 | 3 | 0 | 3 |
| GCDH | Glutaryl-CoA | 2.14671512 | 0.4378277 | 0.0001205 | 0.00012167 | 0 | 0 | 0.00012109 | 0 | 0.00023216 | 0.00028771 | 0 | 0.00025994 | 2 | 2 | 0 | 0 | 2 | 0 | 3 | 3 | 0 | 3 |
| GFPT1 | Glutamine--fi | 1.27619234 | 0.78602685 | 0.00011326 | 7.6242E-05 | 0 | 0 | 9.4751E-05 | 9.6371E-05 | 0.00014547 | 0 | 0 | 0.00012092 | 3 | 2 | 0 | 0 | 2.5 | 2 | 3 | 0 | 0 | 2.5 |
| ACTZ | Alpha-centra | 2.03232753 | 0.4824794 | 0 | 0 | 0.00021111 | 0.00014184 | 0.00017648 | 0 | 0.00027044 | 0.00044687 | 0 | 0.00035866 | 0 | 0 | 3 | 2 | 2.5 | 0 | 3 | 4 | 0 | 3.5 |
| SNX4 | Sorting nexin | 1.14422359 | 0.87760558 | 0 | 0.00011843 | 0.00017639 | 0 | 0.00014741 | 0 | 0.00015065 | 0.00018669 | 0 | 0.00016867 | 0 | 2 | 3 | 0 | 2.5 | 0 | 2 | 2 | 0 | 2 |
| UBA1 | Ubiquitin-like | 0.71570095 | 0.27563369 | 0.00014965 | 0.00012593 | 0 | 0.00012602 | 0.00013387 | 9.5505E-05 | 9.6112E-05 | 0 | 0 | 9.5809E-05 | 6 | 5 | 0 | 5 | 5.33333333 | 3 | 3 | 0 | 0 | 3 |
| TIM23 | Mitochondria | 1.53288726 | 0.97713497 | 0.00050504 | 0 | 0.00050639 | 0.00025517 | 0.0004222 | 0.00048347 | 0.0008109 | 0 | 0 | 0.00064719 | 4 | 0 | 4 | 2 | 3.33333333 | 3 | 5 | 0 | 0 | 4 |
| MCM2 | DNA replicati | 2.14090229 | 0.65089569 | 0 | 5.8952E-05 | 5.8537E-05 | 5.8995E-05 | 5.8828E-05 | 0 | 0.00011249 | 0.0001394 | 0 | 0.00012595 | 0 | 2 | 2 | 2 | 2 | 0 | 3 | 3 | 0 | 3 |
| IDH3B | Isocitrate del | 2.45069111 | 0.55719281 | 0.00013708 | 0.00013842 | 0 | 0.00020778 | 0.00016109 | 0.00043742 | 0.00035216 | 0 | 0 | 0.00039479 | 2 | 2 | 0 | 3 | 2.33333333 | 5 | 4 | 0 | 0 | 4.5 |
| DDX31 | Probable ATF | 0.76525263 | 0.32232038 | 0.00012403 | 0 | 9.3274E-05 | 9.4003E-05 | 0.00010377 | 7.9158E-05 | 7.9661E-05 | 0 | 0 | 7.941E-05 | 4 | 0 | 3 | 3 | 3.33333333 | 2 | 2 | 0 | 0 | 2 |
| EMC1 | ER membran | 0.85360796 | 0.45018983 | 5.3149E-05 | 0.00013417 | 0 | 8.0561E-05 | 8.9293E-05 | 6.7838E-05 | 0 | 8.4605E-05 | 0 | 7.6222E-05 | 2 | 5 | 0 | 3 | 3.33333333 | 2 | 0 | 2 | 0 | 2 |
| LYPA2 | Acyl-protein | 1.36729639 | 0.89975772 | 0.00022847 | 0 | 0.00034362 | 0.00034631 | 0.00030613 | 0.00029162 | 0 | 0.00054553 | 0 | 0.00041858 | 2 | 0 | 3 | 3 | 2.66666667 | 2 | 0 | 3 | 0 | 2.5 |
| TCOF | Treacle prote | 0.09380099 | 1.2666E-05 | 0.0011527 | 0.0011461 | 0.00087129 | 0.0011827 | 0.0010882 | 0.00013581 | 6.8338E-05 | 0 | 0 | 0.00010207 | 65 | 64 | 49 | 66 | 61 | 6 | 3 | 0 | 0 | 4.5 |
| NUP50 | Nuclear pore | 0.06467218 | 9.8524E-09 | 0.0028757 | 0.002733 | 0.0027702 | 0.0027919 | 0.0027927 | 0.00014394 | 0.00021728 | 0 | 0 | 0.00018061 | 51 | 48 | 49 | 49 | 49.25 | 2 | 3 | 0 | 0 | 2.5 |
| GTF2I | General trans | 0.15697309 | 0.00011804 | 0.00095189 | 0.0009879 | 0.001193 | 0.00074813 | 0.00097023 | 0.00016875 | 0.00013585 | 0 | 0 | 0.0001523 | 36 | 37 | 45 | 28 | 36.5 | 5 | 4 | 0 | 0 | 4.5 |
| NPA1P | Nucleolar pre | 0.0833226 | 9.7072E-05 | 0.00044155 | 0.00030507 | 0.00048933 | 0.000364 | 0.00039999 | 2.9662E-05 | 0 | 3.6994E-05 | 0 | 3.3328E-05 | 38 | 26 | 42 | 31 | 34.25 | 2 | 0 | 2 | 0 | 2 |
| DDX46 | Probable ATF | 0.15687927 | 0.00342491 | 0.00040952 | 0.00074951 | 0.00030796 | 0.00062073 | 0.00052193 | 9.8007E-05 | 6.5753E-05 | 0 | 0 | 0.00008188 | 16 | 29 | 12 | 24 | 20.25 | 3 | 2 | 0 | 0 | 2.5 |
| TFP11 | Tuftelin-inter | 0.14980309 | 0.00010282 | 0.00059902 | 0.00041386 | 0.00050578 | 0.00063717 | 0.00053896 | 8.0482E-05 | 8.0993E-05 | 0 | 0 | 8.0738E-05 | 19 | 13 | 16 | 20 | 17 | 2 | 2 | 0 | 0 | 2 |
| SRSF6 | Serine/argini | 0.58660821 | 0.03930438 | 0.00023013 | 0.00054223 | 0.00069223 | 0.00054261 | 0.0005018 | 0.00039165 | 0.00019707 | 0 | 0 | 0.00029436 | 3 | 7 | 9 | 7 | 6.5 | 4 | 2 | 0 | 0 | 3 |
| SPDLY | Protein Spinc | 0.20513367 | 0.00010384 | 0.00052341 | 0.0007047 | 0.00074347 | 0.00074928 | 0.00068022 | 0.00016702 | 0.00011205 | 0 | 0 | 0.00013954 | 12 | 16 | 17 | 17 | 15.5 | 3 | 2 | 0 | 0 | 2.5 |
| STIM1 | Stromal inter | 0.22134159 | 0.00015436 | 0.00034671 | 0.0004279 | 0.00050214 | 0.00050606 | 0.0004457 | 0.00009834 | 9.8965E-05 | 0 | 0 | 9.8653E-05 | 9 | 11 | 13 | 13 | 11.5 | 2 | 2 | 0 | 0 | 2 |
| NUDC | Nuclear migr | 0.42223198 | 0.00381144 | 0.00087696 | 0.001288 | 0.00079936 | 0.00088617 | 0.00096262 | 0.00030527 | 0 | 0.00050763 | 0 | 0.00040645 | 11 | 16 | 10 | 11 | 12 | 3 | 0 | 4 | 0 | 3.5 |
| WASC4 | WASH compl | 0.63729106 | 0.02395335 | 0.00017997 | 0.00022717 | 0.00027068 | 0.00022733 | 0.00022629 | 8.6142E-05 | 0.00020228 | 0 | 0 | 0.00014421 | 8 | 10 | 12 | 10 | 10 | 3 | 7 | 0 | 0 | 5 |
| IST1 | IST1 homolog | 1.34084458 | 0.46889615 | 0.00028998 | 0.00036602 | 0.00050882 | 0.00021977 | 0.00034615 | 0.00046266 | 0.0004656 | 0 | 0 | 0.00046413 | 4 | 5 | 7 | 3 | 4.75 | 5 | 5 | 0 | 0 | 5 |
| PPHLN | Periphilin-1 C | 0.67074334 | 0.02530367 | 0.0003457 | 0.0002909 | 0.00023108 | 0.00023289 | 0.00027514 | 0.00014708 | 0.00022202 | 0 | 0 | 0.00018455 | 6 | 5 | 4 | 4 | 4.75 | 2 | 3 | 0 | 0 | 2.5 |
| TF65 | Transcription | 0.46479283 | 0.00305106 | 0.00028735 | 0.00038688 | 0.00043217 | 0.00033876 | 0.00036129 | 0.00018338 | 0 | 0.00015247 | 0 | 0.00016793 | 6 | 8 | 9 | 7 | 7.5 | 3 | 0 | 2 | 0 | 2.5 |
| HNRPF | Heterogeneo | 0.33558739 | 0.0014049 | 0.00050869 | 0.00064208 | 0.00063756 | 0.00038553 | 0.00054347 | 0.00016232 | 0 | 0.00020244 | 0 | 0.00018238 | 14 | 14 | 17 | 10 | 13.75 | 8 | 0 | 7 | 0 | 7.5 |
| HAT1 | Histone acety | 0.33471083 | 0.00141088 | 0.0006298 | 0.00076315 | 0.00044203 | 0.00057277 | 0.00060194 | 0.00024116 | 0.00016179 | 0 | 0 | 0.00020148 | 10 | 12 | 7 | 9 | 9.5 | 3 | 2 | 0 | 0 | 2.5 |
| STK4 | Serine/threo | 1.45491481 | 0.54785248 | 0.00016256 | 0.00021886 | 0.00021732 | 0.00016426 | 0.00019075 | 0.00027665 | 0.0002784 | 0 | 0 | 0.00027753 | 3 | 4 | 4 | 3 | 3.5 | 4 | 4 | 0 | 0 | 4 |
| PHOCN | MOB-like pro | 0.92672268 | 0.11256693 | 0.00070369 | 0.00071057 | 0.00058797 | 0.00059257 | 0.0006487 | 0.00044909 | 0.00075324 | 0 | 0 | 0.00060117 | 6 | 6 | 5 | 5 | 5.5 | 3 | 5 | 0 | 0 | 4 |
| PLAK | Junction plak | 1.5278406 | 0.63019624 | 0.00014168 | 0.00014307 | 0.00014206 | 0.00017896 | 0.00015144 | 0.00018084 | 0 | 0.00028192 | 0 | 0.00023138 | 4 | 4 | 4 | 5 | 4.25 | 4 | 0 | 5 | 0 | 4.5 |
| PRS6A | 26S proteaso | 0.9712126 | 0.19867492 | 0.00024044 | 0.00030349 | 0.00036162 | 0.00036445 | 0.0003175 | 0.00015345 | 0.00046327 | 0 | 0 | 0.00030836 | 4 | 5 | 6 | 6 | 5.25 | 2 | 6 | 0 | 0 | 4 |
| AP3B1 | AP-3 comple: | 0.8708368 | 0.14964759 | 4.8242E-05 | 7.3071E-05 | 0.00012093 | 0.00014625 | 9.7123E-05 | 9.2363E-05 | 0 | 7.6794E-05 | 0 | 8.4579E-05 | 2 | 3 | 5 | 6 | 4 | 3 | 0 | 2 | 0 | 2.5 |
| H1X | Histone H1x | 0.87472408 | 0.10296313 | 0.00061945 | 0.0003753 | 0.00037266 | 0.00062595 | 0.00049834 | 0 | 0.0004774 | 0.00039442 | 0 | 0.00043591 | 5 | 3 | 3 | 5 | 4 | 0 | 3 | 2 | 0 | 2.5 |
| ARP2 | Actin-related | 0.85733999 | 0.09804528 | 0.0002679 | 0.00020289 | 0.00033577 | 0.0003384 | 0.00028624 | 0.00017097 | 0 | 0.00031984 | 0 | 0.00024541 | 4 | 3 | 5 | 5 | 4.25 | 2 | 0 | 3 | 0 | 2.5 |
| PP1A | Serine/threo | 0.63577846 | 0.0284836 | 0.00023989 | 0.00048448 | 0.00048107 | 0.00040402 | 0.00040237 | 0.0003062 | 0.00020543 | 0 | 0 | 0.00025582 | 51 | 49 | 44 | 57 | 50.25 | 28 | 27 | 0 | 0 | 27.5 |
| SYAP1 | Synapse-assc | 0.90335779 | 0.08734465 | 0.00037484 | 0.0003785 | 0.00037583 | 0.00030302 | 0.00035805 | 0 | 0.00028888 | 0.00035801 | 0 | 0.00032345 | 5 | 5 | 5 | 4 | 4.75 | 0 | 3 | 3 | 0 | 3 |
| H32 | Histone H3.2 | 0.45785222 | 0.63957573 | 0 | 0 | 0 | 0.0049018 | 0.0049018 | 0 | 0 | 0 | 0.0022443 | 0.0022443 | 0 | 0 | 0 | 25 | 25 | 0 | 0 | 0 | 9 | 9 |
| RS20 | 40S ribosoma | 1.92270846 | 0.68513315 | 0 | 0 | 0.00044468 | 0 | 0.00044468 | 0 | 0 | 0 | 0.00085499 | 0.00085499 | 0 | 0 | 2 | 0 | 2 | 0 | 0 | 0 | 3 | 3 |
| RUXE | Small nuclea | 0.53749809 | 0.00865375 | 0.0011473 | 0.0020274 | 0.00086278 | 0.0014492 | 0.00137167 | 0 | 0 | 0 | 0.00073727 | 0.00073727 | 4 | 7 | 3 | 5 | 4.75 | 0 | 0 | 0 | 2 | 2 |
| SUMO1 | Small ubiquit | 0.85355686 | 0.23245526 | 0.00052254 | 0 | 0.0013098 | 0.00052803 | 0.00078679 | 0 | 0 | 0 | 0.00067157 | 0.00067157 | 2 | 0 | 5 | 2 | 3 | 0 | 0 | 0 | 2 | 2 |
| H2B1B | Histone H2B | 1.02211526 | 0.03787509 | 0.00041886 | 0.00042296 | 0.00062997 | 0.0006349 | 0.00052667 | 0 | 0 | 0 | 0.00053832 | 0.00053832 | 17 | 14 | 17 | 14 | 15.5 | 0 | 0 | 0 | 10 | 10 |
| H2B1J | Histone H2B | 1.02211526 | 0.03787509 | 0.00041886 | 0.00042296 | 0.00062997 | 0.0006349 | 0.00052667 | 0 | 0 | 0 | 0.00053832 | 0.00053832 | 17 | 14 | 17 | 14 | 15.5 | 0 | 0 | 0 | 10 | 10 |
| H2B2E | Histone H2B | 1.02211526 | 0.03787509 | 0.00041886 | 0.00042296 | 0.00062997 | 0.0006349 | 0.00052667 | 0 | 0 | 0 | 0.00053832 | 0.00053832 | 17 | 14 | 17 | 14 | 15.5 | 0 | 0 | 0 | 10 | 10 |

| | | | | | | | | | | | | | | | | | | | | | |
|---|---|---|---|---|---|---|---|---|---|---|---|---|---|---|---|---|---|---|---|---|---|
| CFA20 | Cilia- and flag | 1.09314671 | 0.62074719 | 0 | 0.00069032 | 0.00027418 | 0 | 0.00048225 | 0 | 0 | 0 | 0.00052717 | 0.00052717 | 0 | 5 | 2 | 0 | 3.5 | 0 | 0 | 0 | 3 | 3 |
| RT17 | 28S ribosoma | 1.27274058 | 0.87172365 | 0 | 0.00040995 | 0 | 0 | 0.00040995 | 0 | 0 | 0 | 0.00052176 | 0.00052176 | 0 | 2 | 0 | 0 | 2 | 0 | 0 | 0 | 2 | 2 |
| JUPI1 | Jupiter micro | 0.85210729 | 0.02057406 | 0.00068541 | 0.00051909 | 0.00034362 | 0.00051946 | 0.0005169 | 0 | 0 | 0 | 0.00044045 | 0.00044045 | 4 | 3 | 2 | 3 | 3 | 0 | 0 | 0 | 2 | 2 |
| CENPV | Centromere | 1.53582399 | 0.81956179 | 0.00028787 | 0 | 0 | 0.00019393 | 0.0002409 | 0 | 0 | 0 | 0.00036998 | 0.00036998 | 3 | 0 | 0 | 2 | 2.5 | 0 | 0 | 0 | 3 | 3 |
| PTBP3 | Polypyrimidi | 3.81555659 | 0.50215566 | 0 | 0 | 0 | 9.6615E-05 | 9.6615E-05 | 0 | 0 | 0 | 0.00036864 | 0.00036864 | 0 | 0 | 0 | 3 | 3 | 0 | 0 | 0 | 7 | 7 |
| MYL6B | Myosin light | 0.10761932 | 0.00010848 | 0.0030448 | 0.0021778 | 0.0031801 | 0.0037178 | 0.00303013 | 0 | 0 | 0 | 0.0003261 | 0.0003261 | 24 | 17 | 25 | 29 | 23.75 | 0 | 0 | 0 | 2 | 2 |
| NUBP2 | Cytosolic Fe- | 1.09526518 | 0.26081485 | 0.00029212 | 0.00019665 | 0 | 0.00019679 | 0.00022852 | 0 | 0 | 0 | 0.00025029 | 0.00025029 | 3 | 2 | 0 | 2 | 2.33333333 | 0 | 0 | 0 | 2 | 2 |
| PDIP3 | Polymerase c | 0.90140057 | 0.02211227 | 0.00025072 | 0.00031647 | 0.00018854 | 0.00031669 | 0.00026811 | 0 | 0 | 0 | 0.00024167 | 0.00024167 | 4 | 5 | 3 | 5 | 4.25 | 0 | 0 | 0 | 3 | 3 |
| EXOS7 | Exosome con | 0.48016892 | 0.08089254 | 0.00063477 | 0.00045784 | 0.00036369 | 0 | 0.00048543 | 0 | 0 | 0 | 0.00023309 | 0.00023309 | 7 | 5 | 4 | 0 | 5.33333333 | 0 | 0 | 0 | 2 | 2 |
| DC1L2 | Cytoplasmic | 1.53332839 | 0.17448408 | 0.00010727 | 0.00016248 | 0.00016133 | 0.0001084 | 0.00013487 | 0 | 0 | 0 | 0.0002068 | 0.0002068 | 2 | 3 | 3 | 2 | 2.5 | 0 | 0 | 0 | 4 | 4 |
| RFC5 | Replication fa | 1.27183476 | 0.87208937 | 0 | 0 | 0 | 0.00015686 | 0.00015686 | 0 | 0 | 0 | 0.0001995 | 0.0001995 | 0 | 0 | 0 | 2 | 2 | 0 | 0 | 0 | 2 | 2 |
| CNO10 | CCR4-NOT tra | 2.03581716 | 0.98839843 | 0 | 0.00010745 | 0 | 7.1682E-05 | 8.9566E-05 | 0 | 0 | 0 | 0.00018234 | 0.00018234 | 0 | 3 | 0 | 2 | 2.5 | 0 | 0 | 0 | 4 | 4 |
| ERR2 | Steroid horm | 1.27846242 | 0.68992074 | 0.00012189 | 0 | 0 | 0.00012317 | 0.00012253 | 0 | 0 | 0 | 0.00015665 | 0.00015665 | 2 | 0 | 0 | 2 | 2 | 0 | 0 | 0 | 2 | 2 |
| SRP72 | Signal recogn | 1.43496612 | 0.41458311 | 7.8654E-05 | 0.00011913 | 0 | 0.00011922 | 0.00010567 | 0 | 0 | 0 | 0.00015163 | 0.00015163 | 2 | 3 | 0 | 3 | 2.66666667 | 0 | 0 | 0 | 3 | 3 |
| NOL10 | Nucleolar pro | 0.49454886 | 0.00073655 | 0.00026849 | 0.00027111 | 0.00030766 | 0.00034882 | 0.00029902 | 0 | 0 | 0 | 0.00014788 | 0.00014788 | 7 | 7 | 8 | 9 | 7.75 | 0 | 0 | 0 | 3 | 3 |
| ACSL3 | Long-chain-fa | 1.02395212 | 0.07190342 | 0.0002199 | 7.4018E-05 | 0.00014699 | 0.00011111 | 0.000138 | 0 | 0 | 0 | 0.00014131 | 0.00014131 | 6 | 2 | 4 | 3 | 3.75 | 0 | 0 | 0 | 3 | 3 |
| TOE1 | Target of EGI | 1.28527252 | 0.866702 | 0.00010348 | 0 | 0 | 0 | 0.00010348 | 0 | 0 | 0 | 0.000133 | 0.000133 | 2 | 0 | 0 | 0 | 2 | 0 | 0 | 0 | 2 | 2 |
| SMRD1 | SWI/SNF-rela | 0.30998247 | 0.00035977 | 0.00046116 | 0.00031044 | 0.00051376 | 0.00041422 | 0.0004249 | 0 | 0 | 0 | 0.00013171 | 0.00013171 | 9 | 6 | 10 | 8 | 8.25 | 0 | 0 | 0 | 2 | 2 |
| MAVS | Mitochondria | 0.28409072 | 0.00011357 | 0.00043981 | 0.00039476 | 0.00053897 | 0.00039505 | 0.00044215 | 0 | 0 | 0 | 0.00012561 | 0.00012561 | 9 | 8 | 11 | 8 | 9 | 0 | 0 | 0 | 2 | 2 |
| SLMAP | Sarcolemmal | 0.40367274 | 0.00030318 | 0.00035057 | 0.00032182 | 0.00028759 | 0.00025764 | 0.00030441 | 0 | 0 | 0 | 0.00012288 | 0.00012288 | 11 | 10 | 9 | 8 | 9.5 | 0 | 0 | 0 | 3 | 3 |
| EIF2D | Eukaryotic tra | 0.84966667 | 0.18134926 | 0 | 0.00018251 | 9.0612E-05 | 0.00013698 | 0.0001367 | 0 | 0 | 0 | 0.00011615 | 0.00011615 | 0 | 4 | 2 | 3 | 3 | 0 | 0 | 0 | 2 | 2 |
| ILVBL | Acetolactate | 1.27271002 | 0.87173598 | 0 | 8.4324E-05 | 0 | 0 | 8.4324E-05 | 0 | 0 | 0 | 0.00010732 | 0.00010732 | 0 | 2 | 0 | 0 | 2 | 0 | 0 | 0 | 2 | 2 |
| ABCF3 | ATP-binding | 1.27894976 | 0.69017685 | 7.4438E-05 | 7.5166E-05 | 0 | 0 | 7.4802E-05 | 0 | 0 | 0 | 9.5668E-05 | 9.5668E-05 | 2 | 2 | 0 | 0 | 2 | 0 | 0 | 0 | 2 | 2 |
| TERA | Transitional | 0.18249042 | 1.5111E-05 | 0.00039288 | 0.00052896 | 0.00045958 | 0.00046317 | 0.00046115 | 0 | 0 | 0 | 8.4155E-05 | 8.4155E-05 | 12 | 16 | 14 | 14 | 14 | 0 | 0 | 0 | 2 | 2 |
| NEB2 | Neurabin-2 C | 0.93059869 | 0.03778343 | 0.00012951 | 0.00006539 | 9.7394E-05 | 6.5437E-05 | 8.9433E-05 | 0 | 0 | 0 | 8.3226E-05 | 8.3226E-05 | 4 | 2 | 3 | 2 | 2.75 | 0 | 0 | 0 | 2 | 2 |
| UBP54 | Inactive ubiq | 2.57042119 | 0.58975776 | 0.00003134 | 0 | 0 | 0 | 0.00003134 | 0 | 0 | 0 | 8.0557E-05 | 8.0557E-05 | 2 | 0 | 0 | 0 | 2 | 0 | 0 | 0 | 4 | 4 |
| COPB2 | Coatomer sul | 0.68088319 | 0.06034925 | 5.8253E-05 | 0.00020588 | 0.00011682 | 5.8864E-05 | 0.00010995 | 0 | 0 | 0 | 7.4866E-05 | 7.4866E-05 | 2 | 7 | 4 | 2 | 3.75 | 0 | 0 | 0 | 2 | 2 |
| PRP4B | Serine/threo | 0.2763846 | 0.00011908 | 0.00026205 | 0.00018523 | 0.00026275 | 0.0002648 | 0.00024371 | 0 | 0 | 0 | 6.7357E-05 | 6.7357E-05 | 10 | 7 | 10 | 10 | 9.25 | 0 | 0 | 0 | 2 | 2 |
| SYIM | Isoleucine--tI | 1.02652658 | 0.56034844 | 5.2151E-05 | 0 | 7.8435E-05 | 0 | 6.5293E-05 | 0 | 0 | 0 | 6.7025E-05 | 6.7025E-05 | 2 | 0 | 3 | 0 | 2.5 | 0 | 0 | 0 | 2 | 2 |
| MDC1 | Mediator of I | 0.0885041 | 1.3587E-07 | 0.00072003 | 0.00070156 | 0.00072195 | 0.00079142 | 0.00073374 | 0 | 0 | 0 | 6.4939E-05 | 6.4939E-05 | 57 | 55 | 57 | 62 | 57.75 | 0 | 0 | 0 | 4 | 4 |
| MKL2 | MKL/myocar | 1.27183892 | 0.87208769 | 0 | 0 | 0 | 4.9018E-05 | 4.9018E-05 | 0 | 0 | 0 | 6.2343E-05 | 6.2343E-05 | 0 | 0 | 0 | 2 | 2 | 0 | 0 | 0 | 2 | 2 |
| DEN4C | DENN domai | 0.63861055 | 0.00554714 | 5.5293E-05 | 9.7708E-05 | 0.00009702 | 0.00008381 | 8.3458E-05 | 0 | 0 | 0 | 5.3297E-05 | 5.3297E-05 | 4 | 7 | 7 | 6 | 6 | 0 | 0 | 0 | 3 | 3 |
| YETS2 | YEATS domai | 0.17332219 | 3.0663E-05 | 0.0002598 | 0.00026234 | 0.00033492 | 0.00024378 | 0.00027521 | 0 | 0 | 0 | 0.0000477 | 0.0000477 | 14 | 14 | 18 | 13 | 14.75 | 0 | 0 | 0 | 2 | 2 |
| ZMYM4 | Zinc finger M | 0.06911553 | 1.9286E-05 | 0.00076711 | 0.00058526 | 0.0006495 | 0.000534 | 0.00063397 | 0 | 0 | 0 | 4.3817E-05 | 4.3817E-05 | 45 | 34 | 38 | 31 | 37 | 0 | 0 | 0 | 2 | 2 |
| KDIS | Kinase D-inte | 0.22718173 | 2.1793E-05 | 0.000149 | 0.00016551 | 0.00019422 | 0.00016562 | 0.00016859 | 0 | 0 | 0 | 0.0000383 | 0.0000383 | 10 | 11 | 13 | 11 | 11.25 | 0 | 0 | 0 | 2 | 2 |
| RHG21 | Rho GTPase- | 0.51225018 | 0.03422016 | 9.4389E-05 | 4.0848E-05 | 0.00010816 | 2.7252E-05 | 6.7662E-05 | 0 | 0 | 0 | 0.00003466 | 0.00003466 | 7 | 3 | 8 | 2 | 5 | 0 | 0 | 0 | 2 | 2 |
| SETD2 | Histone-lysin | 0.08663997 | 4.5076E-06 | 0.00033963 | 0.0002806 | 0.0002683 | 0.0003328 | 0.00030533 | 0 | 0 | 0 | 2.6454E-05 | 2.6454E-05 | 33 | 27 | 26 | 32 | 29.5 | 0 | 0 | 0 | 2 | 2 |
| HS105 | Heat shock p | 0.42133734 | 0.61301756 | 0 | 0.00018634 | 0 | 0 | 0.00018634 | 7.8512E-05 | 0 | 0 | 0 | 7.8512E-05 | 0 | 6 | 0 | 0 | 6 | 2 | 0 | 0 | 0 | 2 |
| ELP1 | Elongator co | 0.85404081 | 0.9152436 | 0 | 0 | 5.9592E-05 | 0 | 5.9592E-05 | 0 | 5.0894E-05 | 0 | 0 | 5.0894E-05 | 0 | 0 | 3 | 0 | 3 | 0 | 2 | 0 | 0 | 2 |
| ARL2 | ADP-ribosyla | 1.01763593 | 0.99053835 | 0 | 0.00072409 | 0 | 0 | 0.00072409 | 0 | 0.00073686 | 0 | 0 | 0.00073686 | 0 | 5 | 0 | 0 | 5 | 0 | 4 | 0 | 0 | 4 |
| K1841 | Uncharacteri | 1.0584351 | 0.96929095 | 0 | 0 | 0.00011055 | 0 | 0.00011055 | 0 | 0 | 0.00011701 | 0 | 0.00011701 | 0 | 0 | 3 | 0 | 3 | 0 | 0 | 2 | 0 | 2 |
| LTN1 | E3 ubiquitin- | 1.27113481 | 0.87237222 | 0 | 0 | 0 | 3.0199E-05 | 3.0199E-05 | 0 | 3.8387E-05 | 0 | 0 | 3.8387E-05 | 0 | 0 | 0 | 2 | 2 | 0 | 2 | 0 | 0 | 2 |
| RFOX2 | RNA binding | 1.27297516 | 0.871629 | 0 | 0 | 0.00013569 | 0 | 0.00013569 | 0.00017273 | 0 | 0 | 0 | 0.00017273 | 0 | 0 | 2 | 0 | 2 | 2 | 0 | 0 | 0 | 2 |
| CEP44 | Centrosomal | 1.28100818 | 0.86840283 | 0 | 0 | 0.00013569 | 0 | 0.00013569 | 0 | 0.00017382 | 0 | 0 | 0.00017382 | 0 | 0 | 2 | 0 | 2 | 0 | 2 | 0 | 0 | 2 |
| MLXIP | MLX-interact | 1.2810823 | 0.8683732 | 0 | 0 | 5.7581E-05 | 0 | 5.7581E-05 | 0 | 7.3766E-05 | 0 | 0 | 7.3766E-05 | 0 | 0 | 2 | 0 | 2 | 0 | 2 | 0 | 0 | 2 |
| CAP1 | Adenylyl cycl | 1.28114901 | 0.86834653 | 0 | 0 | 0.0001114 | 0 | 0.0001114 | 0 | 0.00014272 | 0 | 0 | 0.00014272 | 0 | 0 | 2 | 0 | 2 | 0 | 2 | 0 | 0 | 2 |
| SPTC1 | Serine palmit | 1.28445958 | 0.86702561 | 0.00011158 | 0 | 0 | 0 | 0.00011158 | 0 | 0.00014332 | 0 | 0 | 0.00014332 | 2 | 0 | 0 | 0 | 2 | 0 | 2 | 0 | 0 | 2 |
| CNO6L | CCR4-NOT tra | 1.28449243 | 0.86701253 | 0.00014264 | 0 | 0 | 0 | 0.00014264 | 0 | 0.00018322 | 0 | 0 | 0.00018322 | 3 | 0 | 0 | 0 | 3 | 0 | 3 | 0 | 0 | 3 |
| ELP4 | Elongator co | 1.57529019 | 0.76826285 | 0 | 0 | 0 | 0.00018867 | 0.00018867 | 0 | 0 | 0.00029721 | 0 | 0.00029721 | 0 | 0 | 0 | 3 | 3 | 0 | 0 | 3 | 0 | 3 |
| NCLN | Nicalin OS=H | 1.57639527 | 0.76794566 | 0 | 9.4659E-05 | 0 | 0 | 9.4659E-05 | 0 | 0 | 0.00014922 | 0 | 0.00014922 | 0 | 2 | 0 | 0 | 2 | 0 | 0 | 2 | 0 | 2 |
| WDR61 | WD repeat-c | 1.57642121 | 0.76793822 | 0 | 0.0002621 | 0 | 0 | 0.0002621 | 0 | 0 | 0.00041318 | 0 | 0.00041318 | 0 | 3 | 0 | 0 | 3 | 0 | 0 | 3 | 0 | 3 |
| LMF2 | Lipase matur | 1.58761757 | 0.76474638 | 0 | 0 | 7.4848E-05 | 0 | 7.4848E-05 | 0 | 0 | 0.00011883 | 0 | 0.00011883 | 0 | 0 | 2 | 0 | 2 | 0 | 0 | 2 | 0 | 2 |
| RL36 | 60S ribosoma | 1.59183511 | 0.76355428 | 0.00050264 | 0 | 0 | 0 | 0.00050264 | 0 | 0 | 0.00080012 | 0 | 0.00080012 | 2 | 0 | 0 | 0 | 2 | 0 | 0 | 2 | 0 | 2 |
| ARF6 | ADP-ribosyla | 1.92675244 | 0.68433391 | 0.00030158 | 0 | 0 | 0 | 0.00030158 | 0 | 0.00058107 | 0 | 0 | 0.00058107 | 2 | 0 | 0 | 0 | 2 | 0 | 3 | 0 | 0 | 3 |
| OSB11 | Oxysterol-bir | 1.92676782 | 0.68433088 | 7.0652E-05 | 0 | 0 | 0 | 7.0652E-05 | 0 | 0.00013613 | 0 | 0 | 0.00013613 | 2 | 0 | 0 | 0 | 2 | 0 | 3 | 0 | 0 | 3 |
| PDCD5 | Programmed | 2.36282667 | 0.61438248 | 0 | 0 | 0 | 0.00042665 | 0.00042665 | 0 | 0 | 0.0010081 | 0 | 0.0010081 | 0 | 0 | 0 | 2 | 2 | 0 | 0 | 3 | 0 | 3 |
| AP3S1 | AP-3 comple: | 2.36464709 | 0.61414645 | 0 | 0.00027613 | 0 | 0 | 0.00027613 | 0 | 0 | 0.00065295 | 0 | 0.00065295 | 0 | 2 | 0 | 0 | 2 | 0 | 0 | 3 | 0 | 3 |
| MARE1 | Microtubule- | 1.0505714 | 0.5653643 | 0 | 0.00029828 | 0 | 0.0002985 | 0.00029839 | 0 | 0 | 0.00031348 | 0 | 0.00031348 | 0 | 3 | 0 | 3 | 3 | 0 | 0 | 2 | 0 | 2 |
| LDHB | L-lactate deh | 1.69547875 | 0.88685104 | 0 | 0.00023934 | 0 | 0.00023951 | 0.00023943 | 0 | 0.00040594 | 0 | 0 | 0.00040594 | 0 | 4 | 0 | 4 | 4 | 0 | 6 | 0 | 0 | 6 |
| METK2 | S-adenosylm | 2.54315774 | 0.85222579 | 0 | 0.00013492 | 0 | 0.00013502 | 0.00013497 | 0 | 0.00034325 | 0 | 0 | 0.00034325 | 0 | 2 | 0 | 2 | 2 | 0 | 4 | 0 | 0 | 4 |
| LPP | Lipoma-prefe | 0.72391375 | 0.43768316 | 8.6237E-05 | 0 | 0 | 0.00021786 | 0.00015205 | 0.00011007 | 0 | 0 | 0 | 0.00011007 | 2 | 0 | 0 | 5 | 3.5 | 2 | 0 | 0 | 0 | 2 |
| PIMT | Protein-L-iso | 0.51080133 | 0.2976764 | 0.00046499 | 0.00070431 | 0 | 0 | 0.00058465 | 0 | 0.00029864 | 0 | 0 | 0.00029864 | 4 | 6 | 0 | 0 | 5 | 0 | 2 | 0 | 0 | 2 |
| PFD5 | Prefoldin sub | 0.90061579 | 0.52336867 | 0 | 0.00086515 | 0 | 0.00034631 | 0.00060573 | 0 | 0 | 0.00054553 | 0 | 0.00054553 | 0 | 5 | 0 | 2 | 3.5 | 0 | 0 | 2 | 0 | 2 |
| BORG1 | Cdc42 effect | 1.01720444 | 0.55516567 | 0 | 0.00025378 | 0 | 0.00038094 | 0.00031736 | 0 | 0.00032282 | 0 | 0 | 0.00032282 | 0 | 2 | 0 | 3 | 2.5 | 0 | 2 | 0 | 0 | 2 |
| WAC2C | WASH compl | 1.27467485 | 0.68792146 | 4.0043E-05 | 0 | 0.00004015 | 0 | 4.0097E-05 | 0.00005111 | 0 | 0 | 0 | 0.00005111 | 33 | 0 | 32 | 0 | 32.5 | 18 | 0 | 0 | 0 | 18 |
| HAUS4 | HAUS augmi | 3.15173799 | 0.74320643 | 0 | 0.00014681 | 0 | 0.00014692 | 0.00014687 | 0 | 0 | 0.00046288 | 0 | 0.00046288 | 0 | 2 | 0 | 2 | 2 | 0 | 0 | 4 | 0 | 4 |
| ZC3HF | Zinc finger C | 1.01077056 | 0.55164164 | 0 | 0.0001251 | 0 | 0.00018779 | 0.00015645 | 0.00015813 | 0 | 0 | 0 | 0.00015813 | 0 | 2 | 0 | 3 | 2.5 | 2 | 0 | 0 | 0 | 2 |
| LRCH2 | Leucine-rich | 1.9166132 | 0.97148161 | 6.8989E-05 | 0 | 0 | 6.9714E-05 | 6.9352E-05 | 0 | 0.00013292 | 0 | 0 | 0.00013292 | 2 | 0 | 0 | 2 | 2 | 0 | 3 | 0 | 0 | 3 |
| IKBE | NF-kappa-B i | 1.05341693 | 0.58851792 | 0 | 0.00021317 | 0.00010583 | 0 | 0.0001595 | 0 | 0 | 0.00016802 | 0 | 0.00016802 | 0 | 4 | 2 | 0 | 3 | 0 | 0 | 2 | 0 | 2 |
| EMC3 | ER membran | 1.01472352 | 0.55419841 | 0.00020221 | 0 | 0 | 0.0003065 | 0.00025436 | 0.0002581 | 0 | 0 | 0 | 0.0002581 | 2 | 0 | 0 | 3 | 2.5 | 2 | 0 | 0 | 0 | 2 |
| ELP6 | Elongator co | 1.89536972 | 0.96392895 | 0 | 0.00020035 | 0 | 0.00020049 | 0.00020042 | 0.00037987 | 0 | 0 | 0 | 0.00037987 | 0 | 2 | 0 | 2 | 2 | 3 | 0 | 0 | 0 | 3 |
| RGPD3 | RanBP2-like | 1.28279166 | 0.69219496 | 3.0021E-05 | 0 | 3.0101E-05 | 0 | 3.0061E-05 | 0 | 3.8562E-05 | 0 | 0 | 3.8562E-05 | 50 | 0 | 51 | 0 | 50.5 | 0 | 67 | 0 | 0 | 67 |
| RO60 | 60 kDa SS-A/ | 1.53238098 | 0.81824762 | 9.8098E-05 | 0.00014859 | 0 | 0 | 0.00012334 | 0 | 0.00018901 | 0 | 0 | 0.00018901 | 2 | 3 | 0 | 0 | 2.5 | 0 | 3 | 0 | 0 | 3 |
| AP3M1 | AP-3 comple: | 1.27382054 | 0.39288925 | 0 | 0.00031874 | 0.0001266 | 0.00012759 | 0.00019098 | 0 | 0.00024327 | 0 | 0 | 0.00024327 | 0 | 5 | 2 | 2 | 3 | 0 | 3 | 0 | 0 | 3 |
| PHF8 | Histone lysin | 0.85161479 | 0.18237858 | 4.9789E-05 | 0 | 7.4883E-05 | 0.00010062 | 7.5097E-05 | 0 | 6.3954E-05 | 0 | 0 | 6.3954E-05 | 3 | 0 | 4 | 5 | 4 | 0 | 2 | 0 | 0 | 2 |
| PR38B | Pre-mRNA-sp | 1.92011877 | 0.63984831 | 0.00014499 | 9.7606E-05 | 0.00014538 | 0 | 0.00012933 | 0 | 0.00024832 | 0 | 0 | 0.00024832 | 3 | 2 | 3 | 0 | 2.66666667 | 0 | 4 | 0 | 0 | 4 |
| HEAT1 | HEAT repeat- | 0.87955135 | 0.18109978 | 0.00006154 | 0 | 3.7022E-05 | 6.2187E-05 | 5.3583E-05 | 4.7129E-05 | 0 | 0 | 0 | 4.7129E-05 | 5 | 0 | 3 | 5 | 4.33333333 | 3 | 0 | 0 | 0 | 3 |
| SRBD1 | S1 RNA-bindi | 1.08529841 | 0.257704 | 0 | 5.3561E-05 | 5.3183E-05 | 8.0399E-05 | 6.2381E-05 | 6.7702E-05 | 0 | 0 | 0 | 6.7702E-05 | 0 | 2 | 2 | 3 | 2.33333333 | 2 | 0 | 0 | 0 | 2 |
| ASCC2 | Activating sig | 1.04312375 | 0.29854555 | 0.00010458 | 0.0000704 | 0 | 0.00021135 | 0.00012878 | 0 | 0.00013433 | 0 | 0 | 0.00013433 | 3 | 2 | 0 | 6 | 3.66666667 | 0 | 3 | 0 | 0 | 3 |
| GHC1 | Mitochondria | 1.4379282 | 0.41586781 | 0.0001634 | 0 | 0.00024575 | 0.00024767 | 0.00021894 | 0 | 0.00031482 | 0 | 0 | 0.00031482 | 2 | 0 | 3 | 3 | 2.66666667 | 0 | 3 | 0 | 0 | 3 |
| GID8 | Glucose-indu | 1.0970912 | 0.26187954 | 0.00034722 | 0.00023374 | 0.00023209 | 0 | 0.00027102 | 0 | 0.00029733 | 0 | 0 | 0.00029733 | 3 | 2 | 2 | 0 | 2.33333333 | 0 | 2 | 0 | 0 | 2 |
| PFD3 | Prefoldin sub | 2.37747735 | 0.81898508 | 0.00040185 | 0.00040578 | 0.00026862 | 0 | 0.00035875 | 0 | 0 | 0.00085292 | 0 | 0.00085292 | 3 | 3 | 2 | 0 | 2.66666667 | 0 | 0 | 4 | 0 | 4 |
| ADT1 | ADP/ATP tra | 1.90665767 | 0.62970071 | 0.0001771 | 0.00017884 | 0.00017758 | 0 | 0.00017784 | 0.00033908 | 0 | 0 | 0 | 0.00033908 | 61 | 56 | 55 | 0 | 57.3333333 | 83 | 0 | 0 | 0 | 83 |
| GRWD1 | Glutamate-ri | 1.63844109 | 0.51371465 | 0 | 0.00017924 | 0.00011865 | 0.00011958 | 0.00013916 | 0 | 0.000228 | 0 | 0 | 0.000228 | 0 | 3 | 2 | 2 | 2.33333333 | 0 | 3 | 0 | 0 | 3 |
| CYLD | Ubiquitin car | 0.95097001 | 0.20442767 | 0.00011041 | 0.00013936 | 0 | 8.3679E-05 | 0.00011115 | 0.0001057 | 0 | 0 | 0 | 0.0001057 | 4 | 5 | 0 | 3 | 4 | 3 | 0 | 0 | 0 | 3 |
| APC5 | Anaphase-pr | 0.73186293 | 0.12123743 | 0.00017476 | 0.00014117 | 0.00014018 | 0 | 0.00015204 | 0 | 0 | 0.00011127 | 0 | 0.00011127 | 5 | 4 | 4 | 0 | 4.33333333 | 0 | 0 | 2 | 0 | 2 |
| AAR2 | Protein AAR2 | 0.69825574 | 0.13209656 | 0.0003436 | 0.00020818 | 0.00020671 | 0 | 0.00025283 | 0 | 0.00017654 | 0 | 0 | 0.00017654 | 5 | 3 | 3 | 0 | 3.66666667 | 0 | 2 | 0 | 0 | 2 |
| TRUA | tRNA pseudo | 0.95919765 | 0.20358872 | 0.0001854 | 0.00018721 | 0.00012393 | 0 | 0.00016551 | 0 | 0.00015876 | 0 | 0 | 0.00015876 | 3 | 3 | 2 | 0 | 2.66666667 | 0 | 2 | 0 | 0 | 2 |
| NEDD1 | Protein NEDI | 0.95541816 | 0.20244547 | 0 | 0.00012112 | 8.0178E-05 | 0.00012121 | 0.0001075 | 0 | 0.00010271 | 0 | 0 | 0.00010271 | 0 | 3 | 2 | 3 | 2.66666667 | 0 | 2 | 0 | 0 | 2 |
| BYST | Bystin OS=Hc | 1.05533293 | 0.25610864 | 0.00024154 | 0.00018293 | 0 | 0.00012204 | 0.00018217 | 0 | 0 | 0.00019225 | 0 | 0.00019225 | 4 | 3 | 0 | 2 | 3 | 0 | 0 | 2 | 0 | 2 |
| BORC5 | BLOC-1-relat | 1.09464078 | 0.26054961 | 0.0004039 | 0.0002719 | 0 | 0.0002721 | 0.00031597 | 0 | 0.00034587 | 0 | 0 | 0.00034587 | 3 | 2 | 0 | 2 | 2.33333333 | 0 | 2 | 0 | 0 | 2 |
| PEBP1 | Phosphatidyl | 0.85258702 | 0.1533618 | 0.00042334 | 0 | 0.00042447 | 0.00042779 | 0.0004252 | 0 | 0.00036252 | 0 | 0 | 0.00036252 | 3 | 0 | 3 | 3 | 3 | 0 | 2 | 0 | 0 | 2 |
| VIR | Protein viriliz | 0.11451924 | 1.6126E-06 | 0.00046602 | 0.00049999 | 0.00054027 | 0.00044148 | 0.00048694 | 5.5764E-05 | 0 | 0 | 0 | 5.5764E-05 | 32 | 34 | 37 | 30 | 33.25 | 3 | 0 | 0 | 0 | 3 |
| SPF45 | Splicing facto | 0.11734981 | 0.00010745 | 0.0011187 | 0.0016613 | 0.0012537 | 0.0017289 | 0.00144065 | 0 | 0.00016906 | 0 | 0 | 0.00016906 | 17 | 25 | 19 | 26 | 21.75 | 0 | 2 | 0 | 0 | 2 |
| SNUT1 | U4/U6.U5 tri | 0.0860045 | 6.6968E-05 | 0.00085762 | 0.0012657 | 0.00089298 | 0.00089996 | 0.00097907 | 8.4204E-05 | 0 | 0 | 0 | 8.4204E-05 | 26 | 38 | 27 | 27 | 29.5 | 2 | 0 | 0 | 0 | 2 |
| SET1A | Histone-lysin | 0.1075852 | 5.322E-06 | 0.00043285 | 0.00034342 | 0.000341 | 0.00035929 | 0.00036914 | 0 | 3.9714E-05 | 0 | 0 | 3.9714E-05 | 28 | 22 | 22 | 23 | 23.75 | 0 | 2 | 0 | 0 | 2 |
| SF3B2 | Splicing facto | 0.14099179 | 5.1583E-07 | 0.00050123 | 0.00056568 | 0.00053213 | 0.00053629 | 0.00053383 | 7.5266E-05 | 0 | 0 | 0 | 7.5266E-05 | 17 | 19 | 18 | 18 | 18 | 2 | 0 | 0 | 0 | 2 |
| CDC5L | Cell division | 0.19498493 | 0.0048914 | 0.00052645 | 0.00096353 | 0.00032991 | 0.00076472 | 0.00064615 | 0.00012599 | 0 | 0 | 0 | 0.00012599 | 16 | 29 | 10 | 23 | 19.5 | 3 | 0 | 0 | 0 | 3 |
| NOG2 | Nucleolar GT | 0.1857258 | 3.7184E-06 | 0.00046929 | 0.00054678 | 0.00047054 | 0.0005107 | 0.00049933 | 0 | 9.2738E-05 | 0 | 0 | 9.2738E-05 | 13 | 15 | 13 | 14 | 13.75 | 0 | 2 | 0 | 0 | 2 |
| ZCH18 | Zinc finger C | 0.32210621 | 0.00238377 | 0.00041535 | 0.00064309 | 0.0003054 | 0.00039173 | 0.00043889 | 0.00014137 | 0 | 0 | 0 | 0.00014137 | 15 | 23 | 11 | 14 | 15.75 | 4 | 0 | 0 | 0 | 4 |
| TOP1 | DNA topoiso | 0.22074982 | 9.8741E-06 | 0.00041394 | 0.00034832 | 0.00041504 | 0.00041828 | 0.0003989 | 8.8056E-05 | 0 | 0 | 0 | 8.8056E-05 | 12 | 10 | 12 | 12 | 11.5 | 2 | 0 | 0 | 0 | 2 |
| TRIPC | E3 ubiquitin- | 0.1893226 | 4.1924E-05 | 0.00030469 | 0.00024078 | 0.0003055 | 0.00022757 | 0.00026964 | 0 | 5.1048E-05 | 0 | 0 | 5.1048E-05 | 23 | 18 | 23 | 17 | 20.25 | 0 | 3 | 0 | 0 | 3 |
| SMRC1 | SWI/SNF con | 0.16646392 | 5.4059E-06 | 0.00040598 | 0.00038583 | 0.00033522 | 0.00033784 | 0.00036622 | 6.0962E-05 | 0 | 0 | 0 | 6.0962E-05 | 17 | 16 | 14 | 14 | 15.25 | 2 | 0 | 0 | 0 | 2 |
| RLA1 | 60S acidic rib | 0.17704715 | 4.3126E-06 | 0.0046295 | 0.0056098 | 0.004874 | 0.0049121 | 0.00500635 | 0.00088636 | 0 | 0 | 0 | 0.00088636 | 20 | 24 | 21 | 21 | 21.5 | 3 | 0 | 0 | 0 | 3 |
| MTA2 | Metastasis-a | 0.21140129 | 5.7088E-05 | 0.00043454 | 0.00055846 | 0.00039609 | 0.00051894 | 0.00047701 | 0.00010084 | 0 | 0 | 0 | 0.00010084 | 14 | 19 | 14 | 14 | 15.25 | 2 | 0 | 0 | 0 | 2 |
| MTA1 | Metastasis-a | 0.15725309 | 5.5663E-05 | 0.00059051 | 0.00059629 | 0.0007401 | 0.00048483 | 0.00060293 | 0 | 9.4813E-05 | 0 | 0 | 9.4813E-05 | 16 | 16 | 20 | 13 | 16.25 | 0 | 2 | 0 | 0 | 2 |
| RRP5 | Protein RRP5 | 0.34837986 | 0.0004354 | 0.00016925 | 0.00011393 | 0.00015556 | 0.00018528 | 0.00015601 | 0 | 5.4349E-05 | 0 | 0 | 5.4349E-05 | 12 | 8 | 11 | 13 | 11 | 0 | 3 | 0 | 0 | 3 |
| CEBPZ | CCAAT/enha | 0.11801245 | 1.8786E-05 | 0.00047569 | 0.00050563 | 0.00052717 | 0.00065779 | 0.00054157 | 6.3912E-05 | 0 | 0 | 0 | 6.3912E-05 | 19 | 20 | 21 | 26 | 21.5 | 2 | 0 | 0 | 0 | 2 |
| DDX55 | ATP-depende | 0.22701633 | 0.00012638 | 0.00039583 | 0.00057734 | 0.00057327 | 0.00044443 | 0.00049772 | 0 | 0.00011299 | 0 | 0 | 0.00011299 | 9 | 13 | 13 | 10 | 11.25 | 0 | 2 | 0 | 0 | 2 |
| VCIP1 | Deubiquitina | 0.49078094 | 0.00299274 | 0.00012957 | 0.00021806 | 0.00012991 | 0.00019639 | 0.00016848 | 8.2688E-05 | 0 | 0 | 0 | 8.2688E-05 | 6 | 10 | 6 | 9 | 7.75 | 3 | 0 | 0 | 0 | 3 |
| CKAP4 | Cytoskeleton | 2.53227989 | 0.60052012 | 0.00039451 | 0.0004869 | 0.00021976 | 0.00022148 | 0.00033066 | 0 | 0 | 0.00083733 | 0 | 0.00083733 | 9 | 11 | 5 | 5 | 7.5 | 0 | 0 | 12 | 0 | 12 |
| PININ | Pinin OS=Hor | 0.37569505 | 0.00127874 | 0.00018402 | 0.00026015 | 0.00022141 | 0.00033471 | 0.00025007 | 9.3951E-05 | 0 | 0 | 0 | 9.3951E-05 | 5 | 7 | 6 | 9 | 6.75 | 2 | 0 | 0 | 0 | 2 |
| ADDA | Alpha-adduci | 0.28384339 | 6.7255E-05 | 0.00035805 | 0.00036155 | 0.0002872 | 0.00028945 | 0.00032406 | 0 | 9.1983E-05 | 0 | 0 | 9.1983E-05 | 10 | 10 | 8 | 8 | 9 | 0 | 2 | 0 | 0 | 2 |
| PARD3 | Partitioning | 0.47580493 | 0.00050513 | 0.00015568 | 0.00017686 | 0.00013659 | 0.00015732 | 0.00015661 | 7.4517E-05 | 0 | 0 | 0 | 7.4517E-05 | 8 | 9 | 7 | 8 | 8 | 3 | 0 | 0 | 0 | 3 |
| F91A1 | Protein FAM | 0.46886327 | 0.00067525 | 0.00018894 | 0.00025438 | 0.00022102 | 0.00019092 | 0.00021382 | 0 | 0 | 0.00010025 | 0 | 0.00010025 | 6 | 8 | 7 | 6 | 6.75 | 0 | 0 | 2 | 0 | 2 |
| MPRIP | Myosin phos | 0.9298631 | 0.03787136 | 0.00015447 | 0.00010399 | 0.00020651 | 0.00010406 | 0.00014226 | 0 | 0.00013228 | 0 | 0 | 0.00013228 | 6 | 4 | 8 | 4 | 5.5 | 0 | 4 | 0 | 0 | 4 |
| DPY30 | Protein dpy- | 0.20030422 | 1.8597E-05 | 0.002932 | 0.0032299 | 0.0037416 | 0.0037709 | 0.0034186 | 0 | 0.00068476 | 0 | 0 | 0.00068476 | 11 | 12 | 14 | 14 | 12.75 | 0 | 2 | 0 | 0 | 2 |
| HDAC1 | Histone deac | 0.42325543 | 0.0012671 | 0.00043798 | 0.00027642 | 0.00027447 | 0.00033194 | 0.0003302 | 0.00013976 | 0 | 0 | 0 | 0.00013976 | 12 | 11 | 9 | 13 | 11.25 | 5 | 0 | 0 | 0 | 5 |
| BRX1 | Ribosome bi | 0.89559477 | 0.01549777 | 0.00029902 | 0.00030194 | 0.00029982 | 0.0003777 | 0.00031962 | 0.00028625 | 0 | 0 | 0 | 0.00028625 | 4 | 4 | 4 | 5 | 4.25 | 3 | 0 | 0 | 0 | 3 |
| RRP8 | Ribosomal R | 0.22699216 | 9.266E-05 | 0.00063656 | 0.00081809 | 0.00058023 | 0.00058477 | 0.00065491 | 0 | 0.00014866 | 0 | 0 | 0.00014866 | 11 | 14 | 10 | 10 | 11.25 | 0 | 2 | 0 | 0 | 2 |
| STIP1 | Stress-induce | 0.27417483 | 0.00055771 | 0.00034018 | 0.00058887 | 0.00038981 | 0.00049108 | 0.00045249 | 0.00012406 | 0 | 0 | 0 | 0.00012406 | 7 | 12 | 8 | 10 | 9.25 | 2 | 0 | 0 | 0 | 2 |
| SRPRA | Signal recogr | 0.26715418 | 1.4779E-05 | 0.00037225 | 0.00037589 | 0.00041471 | 0.00041796 | 0.0003952 | 0.00010558 | 0 | 0 | 0 | 0.00010558 | 9 | 9 | 10 | 10 | 9.5 | 2 | 0 | 0 | 0 | 2 |
| CC85C | Coiled-coil dc | 0.51040673 | 0.00240017 | 0.00025192 | 0.00038157 | 0.00025259 | 0.00038185 | 0.00031698 | 0 | 0.00016179 | 0 | 0 | 0.00016179 | 4 | 6 | 4 | 6 | 5 | 0 | 2 | 0 | 0 | 2 |
| LRC59 | Leucine-rich | 0.86266557 | 0.01904915 | 0.00042978 | 0.00052078 | 0.00034474 | 0.00060801 | 0.00047583 | 0 | 0 | 0.00041048 | 0 | 0.00041048 | 5 | 6 | 4 | 7 | 5.5 | 0 | 0 | 3 | 0 | 3 |
| TF3C5 | General tran | 0.760409 | 0.01318817 | 0.00040676 | 0.00020537 | 0.00035686 | 0.00030827 | 0.00031932 | 0 | 0 | 0.00024281 | 0 | 0.00024281 | 8 | 4 | 7 | 6 | 6.25 | 0 | 0 | 3 | 0 | 3 |
| ANKL2 | Ankyrin repe | 0.32778101 | 0.00101924 | 0.00028133 | 0.00022726 | 0.00022566 | 0.00014214 | 0.0002191 | 7.1816E-05 | 0 | 0 | 0 | 7.1816E-05 | 10 | 8 | 8 | 5 | 7.75 | 2 | 0 | 0 | 0 | 2 |
| PRP31 | U4/U6 small | 0.67740479 | 0.00763788 | 0.00021153 | 0.0001602 | 0.00026512 | 0.00016031 | 0.00019929 | 0.000135 | 0 | 0 | 0 | 0.000135 | 4 | 3 | 5 | 3 | 3.75 | 2 | 0 | 0 | 0 | 2 |
| RU17 | U1 small nuc | 1.17949108 | 0.07601957 | 0.00024154 | 0.0002439 | 0.00018164 | 0.00012204 | 0.00019728 | 0 | 0.00023269 | 0 | 0 | 0.00023269 | 4 | 4 | 3 | 2 | 3.25 | 0 | 3 | 0 | 0 | 3 |
| RNPS1 | RNA-binding | 0.72929808 | 0.04783312 | 0.00017304 | 0.00052419 | 0.000347 | 0.00017486 | 0.00030477 | 0 | 0.00022227 | 0 | 0 | 0.00022227 | 2 | 6 | 4 | 2 | 3.5 | 0 | 2 | 0 | 0 | 2 |
| TEX10 | Testis-expres | 0.50686171 | 0.00555738 | 0.00017043 | 0.0001721 | 0.00025633 | 0.00011481 | 0.00017842 | 0 | 0 | 9.0433E-05 | 0 | 9.0433E-05 | 6 | 6 | 9 | 4 | 6.25 | 0 | 0 | 2 | 0 | 2 |
| CEP97 | Centrosomal | 1.52310541 | 0.17054151 | 9.1521E-05 | 9.2416E-05 | 6.1176E-05 | 6.1655E-05 | 7.6692E-05 | 0.00011681 | 0 | 0 | 0 | 0.00011681 | 3 | 3 | 2 | 2 | 2.5 | 3 | 0 | 0 | 0 | 3 |
| NEK9 | Serine/threo | 1.09562888 | 0.0563944 | 0.00013477 | 8.1654E-05 | 8.1079E-05 | 8.1713E-05 | 9.4804E-05 | 0 | 0.00010387 | 0 | 0 | 0.00010387 | 5 | 3 | 3 | 3 | 3.5 | 0 | 3 | 0 | 0 | 3 |
| ORC3 | Origin recogr | 0.70317986 | 0.00614836 | 0.0002598 | 0.00029982 | 0.00018607 | 0.00026253 | 0.00025206 | 0 | 0 | 0.00017724 | 0 | 0.00017724 | 7 | 8 | 5 | 7 | 6.75 | 0 | 0 | 3 | 0 | 3 |

| | | | | | | | | | | | | | | | | | | | | | |
|---|---|---|---|---|---|---|---|---|---|---|---|---|---|---|---|---|---|---|---|---|---|
| EXOS1 | Exosome con | 0.68022619 | 0.01999908 | 0.00040598 | 0.00068324 | 0.00027137 | 0.00068373 | 0.00051108 | 0 | 0.00034765 | 0 | 0 | 0.00034765 | 3 | 5 | 2 | 5 | 3.75 | 0 | 2 | 0 | 0 | 2 |
| AFTIN | Aftiphilin OS= | 0.84630507 | 0.03288582 | 0.00016898 | 0.00017063 | 8.4713E-05 | 8.5375E-05 | 0.00012742 | 0.00010784 | 0 | 0 | 0 | 0.00010784 | 6 | 6 | 3 | 3 | 4.5 | 3 | 0 | 0 | 0 | 3 |
| RPOM | DNA-directed | 0.9009443 | 0.02861178 | 8.5816E-05 | 0.00012998 | 6.4533E-05 | 8.6717E-05 | 9.1762E-05 | 0 | 8.2672E-05 | 0 | 0 | 8.2672E-05 | 4 | 6 | 3 | 4 | 4.25 | 0 | 3 | 0 | 0 | 3 |
| DPOD3 | DNA polymer | 0.35024046 | 0.00204652 | 0.00045302 | 0.00022873 | 0.00045423 | 0.000515 | 0.00041275 | 0.00014456 | 0 | 0 | 0 | 0.00014456 | 8 | 4 | 8 | 9 | 7.25 | 2 | 0 | 0 | 0 | 2 |
| OSBL8 | Oxysterol-bir | 0.85109391 | 0.02218421 | 0.00014842 | 0.00011989 | 8.9287E-05 | 0.00017997 | 0.00013439 | 0 | 0.00011438 | 0 | 0 | 0.00011438 | 5 | 4 | 3 | 6 | 4.5 | 0 | 3 | 0 | 0 | 3 |
| CSN8 | COP9 signalo | 0.40594222 | 0.00188425 | 0.00050504 | 0.00076497 | 0.0010128 | 0.00089311 | 0.00079398 | 0.00032231 | 0 | 0 | 0 | 0.00032231 | 4 | 6 | 8 | 7 | 6.25 | 2 | 0 | 0 | 0 | 2 |
| OCRL | Inositol polyp | 1.94890766 | 0.33831373 | 0.00011715 | 8.8723E-05 | 8.8098E-05 | 8.8787E-05 | 9.569E-05 | 0 | 0 | 0.00018649 | 0 | 0.00018649 | 4 | 3 | 3 | 3 | 3.25 | 0 | 0 | 4 | 0 | 4 |
| COR1B | Coronin-1B C | 0.72480467 | 0.01232638 | 0.00016189 | 0.00027246 | 0.00016232 | 0.00016359 | 0.00019007 | 0.00013776 | 0 | 0 | 0 | 0.00013776 | 3 | 5 | 3 | 3 | 3.5 | 2 | 0 | 0 | 0 | 2 |
| ACTN4 | Alpha-actinin | 1.94615983 | 0.36526679 | 8.6899E-05 | 8.7749E-05 | 5.8087E-05 | 0.00014635 | 9.4771E-05 | 0 | 0 | 0.00018444 | 0 | 0.00018444 | 3 | 3 | 2 | 5 | 3.25 | 0 | 0 | 4 | 0 | 4 |
| COIL | Coilin OS=Ho | 0.53426832 | 0.0009476 | 0.00022907 | 0.00023131 | 0.00018374 | 0.00023147 | 0.0002189 | 0.00011695 | 0 | 0 | 0 | 0.00011695 | 5 | 5 | 4 | 5 | 4.75 | 2 | 0 | 0 | 0 | 2 |
| KIF2A | Kinesin-like p | 1.09386066 | 0.07024895 | 7.4755E-05 | 0.00018871 | 0.00014991 | 0.00011331 | 0.00013167 | 0 | 0.00014403 | 0 | 0 | 0.00014403 | 2 | 5 | 4 | 3 | 3.5 | 0 | 3 | 0 | 0 | 3 |
| HS90A | Heat shock p | 0.5077997 | 0.0036967 | 0.00018025 | 0.00010921 | 0.00021687 | 0.00021857 | 0.00018123 | 9.2026E-05 | 0 | 0 | 0 | 9.2026E-05 | 7 | 8 | 9 | 11 | 8.75 | 7 | 0 | 0 | 0 | 7 |
| SRS10 | Serine/argini | 1.27742979 | 0.10000945 | 0.0005036 | 0.00050852 | 0.00030296 | 0.00030533 | 0.0004051 | 0 | 0.00051749 | 0 | 0 | 0.00051749 | 5 | 5 | 3 | 3 | 4 | 0 | 4 | 0 | 0 | 4 |
| DCP1B | mRNA-decap | 1.91576047 | 0.31838564 | 8.5538E-05 | 8.6374E-05 | 8.5766E-05 | 8.6436E-05 | 8.6029E-05 | 0 | 0.00016481 | 0 | 0 | 0.00016481 | 2 | 2 | 2 | 2 | 2 | 0 | 3 | 0 | 0 | 3 |
| NSMA3 | Sphingomyel | 1.1798459 | 0.09212127 | 9.5726E-05 | 6.4441E-05 | 0.00015997 | 9.6731E-05 | 0.00010422 | 0 | 0.00012296 | 0 | 0 | 0.00012296 | 3 | 2 | 5 | 3 | 3.25 | 0 | 3 | 0 | 0 | 3 |
| FKB15 | FK506-bindin | 0.56774022 | 0.00528759 | 0.00012989 | 8.7437E-05 | 6.5116E-05 | 0.00010937 | 9.7953E-05 | 0 | 5.5612E-05 | 0 | 0 | 5.5612E-05 | 6 | 4 | 3 | 5 | 4.5 | 0 | 2 | 0 | 0 | 2 |
| SKA2 | Spindle and k | 0.8519445 | 0.02052395 | 0.00065426 | 0.00044044 | 0.00087467 | 0.00066113 | 0.00065763 | 0 | 0.00056026 | 0 | 0 | 0.00056026 | 3 | 2 | 4 | 3 | 3 | 0 | 2 | 0 | 0 | 2 |
| ABI2 | Abl interacto | 1.17784776 | 0.07643894 | 0.00015432 | 0.00020777 | 0.00010315 | 0.00020792 | 0.00016829 | 0 | 0.00019822 | 0 | 0 | 0.00019822 | 3 | 4 | 2 | 4 | 3.25 | 0 | 3 | 0 | 0 | 3 |
| SIN1 | Target of rap | 1.0229206 | 0.06084226 | 0.00020221 | 0.00010209 | 0.00010137 | 0.00010217 | 0.00012696 | 0 | 0.00012987 | 0 | 0 | 0.00012987 | 4 | 2 | 2 | 2 | 2.5 | 0 | 2 | 0 | 0 | 2 |
| DERL1 | Derlin-1 OS=I | 1.01537938 | 0.03670535 | 0.0003154 | 0.00021232 | 0.00021083 | 0.00031871 | 0.00026432 | 0.00026838 | 0 | 0 | 0 | 0.00026838 | 3 | 2 | 2 | 3 | 2.5 | 2 | 0 | 0 | 0 | 2 |
| 1433T | 14-3-3 protei | 2.30158317 | 0.4936184 | 0.00032312 | 0.00032628 | 0.00021599 | 0.00032652 | 0.00029798 | 0 | 0 | 0.00068582 | 0 | 0.00068582 | 4 | 4 | 3 | 4 | 3.75 | 0 | 0 | 8 | 0 | 8 |
| EIF3K | Eukaryotic tr | 1.02235592 | 0.06115077 | 0.0002421 | 0.00024446 | 0.00048548 | 0.00024464 | 0.00030417 | 0 | 0.00031097 | 0 | 0 | 0.00031097 | 2 | 2 | 4 | 2 | 2.5 | 0 | 2 | 0 | 0 | 2 |
| APC4 | Anaphase-pr | 0.79086978 | 0.02363988 | 9.7977E-05 | 9.8935E-05 | 0.00013098 | 0.00019801 | 0.00013148 | 0 | 0 | 0.00010398 | 0 | 0.00010398 | 3 | 3 | 4 | 6 | 4 | 0 | 0 | 2 | 0 | 2 |
| EFHD2 | EF-hand dom | 0 | 0.35591768 | 0 | 0 | 0.00044098 | 0 | 0.00044098 | 0 | 0 | 0 | 0 | 0 | 0 | 0 | 4 | 0 | 4 | 0 | 0 | 0 | 0 | 0 |
| MYH14 | Myosin-14 O | 0 | 0.35591768 | 0 | 0 | 0 | 4.0099E-05 | 4.0099E-05 | 0 | 0 | 0 | 0 | 0 | 0 | 0 | 0 | 100 | 100 | 0 | 0 | 0 | 0 | 0 |
| AK17A | A-kinase ancl | 0 | 0.35591768 | 7.5938E-05 | 0 | 0 | 0 | 7.5938E-05 | 0 | 0 | 0 | 0 | 0 | 2 | 0 | 0 | 0 | 2 | 0 | 0 | 0 | 0 | 0 |
| LSG1 | Large subunit | 0 | 0.35591768 | 0 | 0 | 0 | 0.00008105 | 0.00008105 | 0 | 0 | 0 | 0 | 0 | 0 | 0 | 0 | 2 | 2 | 0 | 0 | 0 | 0 | 0 |
| KPCD | Protein kinas | 0 | 0.35591768 | 7.8072E-05 | 0 | 0 | 0 | 7.8072E-05 | 0 | 0 | 0 | 0 | 0 | 2 | 0 | 0 | 0 | 2 | 0 | 0 | 0 | 0 | 0 |
| FOXK1 | Forkhead box | 0 | 0.35591768 | 0 | 7.2705E-05 | 0 | 0 | 7.2705E-05 | 0 | 0 | 0 | 0 | 0 | 0 | 2 | 0 | 0 | 2 | 0 | 0 | 0 | 0 | 0 |
| PRS6B | 26S proteaso | 0 | 0.35591768 | 0 | 0 | 0.00018989 | 0 | 0.00018989 | 0 | 0 | 0 | 0 | 0 | 0 | 0 | 3 | 0 | 3 | 0 | 0 | 0 | 0 | 0 |
| PRS10 | 26S proteaso | 0 | 0.35591768 | 0 | 0.0002055 | 0 | 0 | 0.0002055 | 0 | 0 | 0 | 0 | 0 | 0 | 3 | 0 | 0 | 3 | 0 | 0 | 0 | 0 | 0 |
| LZTL1 | Leucine zippe | 0 | 0.35591768 | 0 | 0 | 0 | 0.00026755 | 0.00026755 | 0 | 0 | 0 | 0 | 0 | 0 | 0 | 0 | 3 | 3 | 0 | 0 | 0 | 0 | 0 |
| DCTN2 | Dynactin sub | 0 | 0.35591768 | 0 | 0 | 0.00019795 | 0 | 0.00019795 | 0 | 0 | 0 | 0 | 0 | 0 | 0 | 3 | 0 | 3 | 0 | 0 | 0 | 0 | 0 |
| PSMD1 | 26S proteaso | 0 | 0.35591768 | 0.00005538 | 0 | 0 | 0 | 0.00005538 | 0 | 0 | 0 | 0 | 0 | 2 | 0 | 0 | 0 | 2 | 0 | 0 | 0 | 0 | 0 |
| CBX2 | Chromobox p | 0 | 0.35591768 | 0 | 0 | 0 | 0.00010025 | 0.00010025 | 0 | 0 | 0 | 0 | 0 | 0 | 0 | 0 | 2 | 2 | 0 | 0 | 0 | 0 | 0 |
| PKHG3 | Pleckstrin ho | 0 | 0.35591768 | 4.3295E-05 | 0 | 0 | 0 | 4.3295E-05 | 0 | 0 | 0 | 0 | 0 | 2 | 0 | 0 | 0 | 2 | 0 | 0 | 0 | 0 | 0 |
| ATP5L | ATP synthase | 0 | 0.35591768 | 0 | 0 | 0.00051376 | 0 | 0.00051376 | 0 | 0 | 0 | 0 | 0 | 0 | 0 | 2 | 0 | 2 | 0 | 0 | 0 | 0 | 0 |
| ORC5 | Origin recogr | 0 | 0.35591768 | 0 | 0 | 0.00012165 | 0 | 0.00012165 | 0 | 0 | 0 | 0 | 0 | 0 | 0 | 2 | 0 | 2 | 0 | 0 | 0 | 0 | 0 |
| ERH | Enhancer of r | 0 | 0.35591768 | 0.0007612 | 0 | 0 | 0 | 0.0007612 | 0 | 0 | 0 | 0 | 0 | 3 | 0 | 0 | 0 | 3 | 0 | 0 | 0 | 0 | 0 |
| NISCH | Nischarin OS= | 0 | 0.35591768 | 0 | 0 | 3.5184E-05 | 0 | 3.5184E-05 | 0 | 0 | 0 | 0 | 0 | 0 | 0 | 2 | 0 | 2 | 0 | 0 | 0 | 0 | 0 |
| CL16A | Protein CLEC | 0 | 0.35591768 | 7.5181E-05 | 0 | 0 | 0 | 7.5181E-05 | 0 | 0 | 0 | 0 | 0 | 3 | 0 | 0 | 0 | 3 | 0 | 0 | 0 | 0 | 0 |
| SPC24 | Kinetochore | 0 | 0.35591768 | 0 | 0.00027052 | 0 | 0 | 0.00027052 | 0 | 0 | 0 | 0 | 0 | 0 | 2 | 0 | 0 | 2 | 0 | 0 | 0 | 0 | 0 |
| SND1 | Staphylococc | 0 | 0.35591768 | 5.7997E-05 | 0 | 0 | 0 | 5.7997E-05 | 0 | 0 | 0 | 0 | 0 | 2 | 0 | 0 | 0 | 2 | 0 | 0 | 0 | 0 | 0 |
| DACH1 | Dachshund h | 0 | 0.35591768 | 6.9443E-05 | 0 | 0 | 0 | 6.9443E-05 | 0 | 0 | 0 | 0 | 0 | 2 | 0 | 0 | 0 | 2 | 0 | 0 | 0 | 0 | 0 |
| SNAG | Gamma-solul | 0 | 0.35591768 | 0 | 0 | 0 | 0.00017093 | 0.00017093 | 0 | 0 | 0 | 0 | 0 | 0 | 0 | 0 | 2 | 2 | 0 | 0 | 0 | 0 | 0 |
| PPIG | Peptidyl-prol | 0 | 0.35591768 | 0 | 0 | 0 | 0.0001061 | 0.0001061 | 0 | 0 | 0 | 0 | 0 | 0 | 0 | 0 | 3 | 3 | 0 | 0 | 0 | 0 | 0 |
| MYO1E | Unconventio | 0 | 0.35591768 | 0 | 0 | 4.7759E-05 | 0 | 4.7759E-05 | 0 | 0 | 0 | 0 | 0 | 0 | 0 | 2 | 0 | 2 | 0 | 0 | 0 | 0 | 0 |
| SBNO1 | Protein straw | 0 | 0.35591768 | 0 | 0 | 7.5976E-05 | 0 | 7.5976E-05 | 0 | 0 | 0 | 0 | 0 | 0 | 0 | 4 | 0 | 4 | 0 | 0 | 0 | 0 | 0 |
| FNBP1 | Formin-bindi | 0 | 0.35591768 | 0.00012831 | 0 | 0 | 0 | 0.00012831 | 0 | 0 | 0 | 0 | 0 | 3 | 0 | 0 | 0 | 3 | 0 | 0 | 0 | 0 | 0 |
| GNAS2 | Guanine nucl | 0 | 0.35591768 | 0 | 0 | 0.00013431 | 0 | 0.00013431 | 0 | 0 | 0 | 0 | 0 | 0 | 0 | 2 | 0 | 2 | 0 | 0 | 0 | 0 | 0 |
| SVIL | Supervillin O | 0 | 0.35591768 | 2.3838E-05 | 0 | 0 | 0 | 2.3838E-05 | 0 | 0 | 0 | 0 | 0 | 2 | 0 | 0 | 0 | 2 | 0 | 0 | 0 | 0 | 0 |
| ATG2A | Autophagy-re | 0 | 0.35591768 | 2.7233E-05 | 0 | 0 | 0 | 2.7233E-05 | 0 | 0 | 0 | 0 | 0 | 2 | 0 | 0 | 0 | 2 | 0 | 0 | 0 | 0 | 0 |
| SYTC | Threonine--tl | 0 | 0.35591768 | 0 | 0 | 7.3191E-05 | 0 | 7.3191E-05 | 0 | 0 | 0 | 0 | 0 | 0 | 0 | 2 | 0 | 2 | 0 | 0 | 0 | 0 | 0 |
| ARHGB | Rho guanine | 0 | 0.35591768 | 3.4676E-05 | 0 | 0 | 0 | 3.4676E-05 | 0 | 0 | 0 | 0 | 0 | 2 | 0 | 0 | 0 | 2 | 0 | 0 | 0 | 0 | 0 |
| BL1S2 | Biogenesis of | 0 | 0.35591768 | 0 | 0.00075061 | 0 | 0 | 0.00075061 | 0 | 0 | 0 | 0 | 0 | 0 | 4 | 0 | 0 | 4 | 0 | 0 | 0 | 0 | 0 |
| NUDT5 | ADP-sugar py | 0 | 0.35591768 | 0 | 0 | 0 | 0.00036528 | 0.00036528 | 0 | 0 | 0 | 0 | 0 | 0 | 0 | 0 | 3 | 3 | 0 | 0 | 0 | 0 | 0 |
| MORC2 | MORC family | 0 | 0.35591768 | 7.6711E-05 | 0 | 0 | 0 | 7.6711E-05 | 0 | 0 | 0 | 0 | 0 | 3 | 0 | 0 | 0 | 3 | 0 | 0 | 0 | 0 | 0 |
| RAVR1 | Ribonucleopr | 0 | 0.35591768 | 0.00021773 | 0 | 0 | 0 | 0.00021773 | 0 | 0 | 0 | 0 | 0 | 5 | 0 | 0 | 0 | 5 | 0 | 0 | 0 | 0 | 0 |
| RBM12 | RNA-binding | 0 | 0.35591768 | 0 | 5.7181E-05 | 0 | 0 | 5.7181E-05 | 0 | 0 | 0 | 0 | 0 | 0 | 2 | 0 | 0 | 2 | 0 | 0 | 0 | 0 | 0 |
| PLOD1 | Procollagen-l | 0 | 0.35591768 | 0 | 0 | 7.2789E-05 | 0 | 7.2789E-05 | 0 | 0 | 0 | 0 | 0 | 0 | 0 | 2 | 0 | 2 | 0 | 0 | 0 | 0 | 0 |
| WIPI4 | WD repeat d | 0 | 0.35591768 | 0 | 0.00022205 | 0 | 0 | 0.00022205 | 0 | 0 | 0 | 0 | 0 | 0 | 3 | 0 | 0 | 3 | 0 | 0 | 0 | 0 | 0 |
| ABI1 | Abl interacto | 0 | 0.35591768 | 0 | 0.00010491 | 0 | 0 | 0.00010491 | 0 | 0 | 0 | 0 | 0 | 0 | 3 | 0 | 0 | 3 | 0 | 0 | 0 | 0 | 0 |
| BABA2 | BRISC and BR | 0 | 0.35591768 | 0 | 0 | 0.00020725 | 0 | 0.00020725 | 0 | 0 | 0 | 0 | 0 | 0 | 0 | 3 | 0 | 3 | 0 | 0 | 0 | 0 | 0 |
| PAPD5 | Non-canonic | 0 | 0.35591768 | 0 | 0 | 9.2513E-05 | 0 | 9.2513E-05 | 0 | 0 | 0 | 0 | 0 | 0 | 0 | 2 | 0 | 2 | 0 | 0 | 0 | 0 | 0 |
| XPR1 | Xenotropic a | 0 | 0.35591768 | 0 | 0 | 0 | 0.00019156 | 0.00019156 | 0 | 0 | 0 | 0 | 0 | 0 | 0 | 0 | 5 | 5 | 0 | 0 | 0 | 0 | 0 |
| TAF1 | Transcription | 0 | 0.35591768 | 0 | 0 | 0 | 7.1222E-05 | 7.1222E-05 | 0 | 0 | 0 | 0 | 0 | 0 | 0 | 0 | 5 | 5 | 0 | 0 | 0 | 0 | 0 |
| SC24B | Protein trans | 0 | 0.35591768 | 0 | 0 | 6.2599E-05 | 0 | 6.2599E-05 | 0 | 0 | 0 | 0 | 0 | 0 | 0 | 3 | 0 | 3 | 0 | 0 | 0 | 0 | 0 |
| CEPT1 | Choline/etha | 0 | 0.35591768 | 0.00012687 | 0 | 0 | 0 | 0.00012687 | 0 | 0 | 0 | 0 | 0 | 2 | 0 | 0 | 0 | 2 | 0 | 0 | 0 | 0 | 0 |
| SSBP | Single-strand | 0 | 0.35591768 | 0 | 0 | 0.00035755 | 0 | 0.00035755 | 0 | 0 | 0 | 0 | 0 | 0 | 0 | 2 | 0 | 2 | 0 | 0 | 0 | 0 | 0 |
| KI21A | Kinesin-like p | 0 | 0.35591768 | 0 | 0 | 3.1611E-05 | 0 | 3.1611E-05 | 0 | 0 | 0 | 0 | 0 | 0 | 0 | 2 | 0 | 2 | 0 | 0 | 0 | 0 | 0 |
| SI1L3 | Signal-induce | 0 | 0.35591768 | 0 | 0 | 2.9712E-05 | 0 | 2.9712E-05 | 0 | 0 | 0 | 0 | 0 | 0 | 0 | 2 | 0 | 2 | 0 | 0 | 0 | 0 | 0 |
| LENG8 | Leukocyte re | 0 | 0.35591768 | 0 | 0 | 0.00006793 | 0 | 0.00006793 | 0 | 0 | 0 | 0 | 0 | 0 | 0 | 2 | 0 | 2 | 0 | 0 | 0 | 0 | 0 |
| RCOR3 | REST corepre | 0 | 0.35591768 | 0 | 0 | 0 | 0.00010774 | 0.00010774 | 0 | 0 | 0 | 0 | 0 | 0 | 0 | 0 | 2 | 2 | 0 | 0 | 0 | 0 | 0 |
| MTA3 | Metastasis-a | 0 | 0.35591768 | 0 | 0 | 0 | 8.9783E-05 | 8.9783E-05 | 0 | 0 | 0 | 0 | 0 | 0 | 0 | 0 | 5 | 5 | 0 | 0 | 0 | 0 | 0 |
| NDUA4 | Cytochrome | 0 | 0.35591768 | 0.00065157 | 0 | 0 | 0 | 0.00065157 | 0 | 0 | 0 | 0 | 0 | 2 | 0 | 0 | 0 | 2 | 0 | 0 | 0 | 0 | 0 |
| ODBB | 2-oxoisovaler | 0 | 0.35591768 | 0 | 0 | 0 | 0.00013605 | 0.00013605 | 0 | 0 | 0 | 0 | 0 | 0 | 0 | 0 | 2 | 2 | 0 | 0 | 0 | 0 | 0 |
| NDUS1 | NADH-ubiqui | 0 | 0.35591768 | 7.2595E-05 | 0 | 0 | 0 | 7.2595E-05 | 0 | 0 | 0 | 0 | 0 | 2 | 0 | 0 | 0 | 2 | 0 | 0 | 0 | 0 | 0 |
| CX056 | UPF0428 pro | 0 | 0.35591768 | 0 | 0 | 0.00035755 | 0 | 0.00035755 | 0 | 0 | 0 | 0 | 0 | 0 | 0 | 3 | 0 | 3 | 0 | 0 | 0 | 0 | 0 |
| MED17 | Mediator of I | 0 | 0.35591768 | 0 | 8.1863E-05 | 0 | 0 | 8.1863E-05 | 0 | 0 | 0 | 0 | 0 | 0 | 2 | 0 | 0 | 2 | 0 | 0 | 0 | 0 | 0 |
| ANLN | Anillin OS=Hc | 0 | 0.35591768 | 0 | 9.4827E-05 | 0 | 0 | 9.4827E-05 | 0 | 0 | 0 | 0 | 0 | 0 | 4 | 0 | 0 | 4 | 0 | 0 | 0 | 0 | 0 |
| RB15B | Putative RNA | 0 | 0.35591768 | 0 | 0 | 0 | 5.9923E-05 | 5.9923E-05 | 0 | 0 | 0 | 0 | 0 | 0 | 0 | 0 | 2 | 2 | 0 | 0 | 0 | 0 | 0 |
| KDM2B | Lysine-specif | 0 | 0.35591768 | 0 | 0 | 3.9609E-05 | 0 | 3.9609E-05 | 0 | 0 | 0 | 0 | 0 | 0 | 0 | 2 | 0 | 2 | 0 | 0 | 0 | 0 | 0 |
| RAB13 | Ras-related p | 0 | 0.35591768 | 0 | 0.00026253 | 0 | 0 | 0.00026253 | 0 | 0 | 0 | 0 | 0 | 0 | 2 | 0 | 0 | 2 | 0 | 0 | 0 | 0 | 0 |
| SGMR2 | Sigma intrace | 0 | 0.35591768 | 0 | 0 | 0 | 0.00045453 | 0.00045453 | 0 | 0 | 0 | 0 | 0 | 0 | 0 | 0 | 3 | 3 | 0 | 0 | 0 | 0 | 0 |
| M18BP | Mis18-bindin | 0 | 0.35591768 | 4.6623E-05 | 0 | 0 | 0 | 4.6623E-05 | 0 | 0 | 0 | 0 | 0 | 2 | 0 | 0 | 0 | 2 | 0 | 0 | 0 | 0 | 0 |
| MTMRC | Myotubularir | 0 | 0.35591768 | 0 | 7.1343E-05 | 0 | 0 | 7.1343E-05 | 0 | 0 | 0 | 0 | 0 | 0 | 2 | 0 | 0 | 2 | 0 | 0 | 0 | 0 | 0 |
| PARVB | Beta-parvin C | 0 | 0.35591768 | 0 | 0 | 0.00014538 | 0 | 0.00014538 | 0 | 0 | 0 | 0 | 0 | 0 | 0 | 2 | 0 | 2 | 0 | 0 | 0 | 0 | 0 |
| UBP19 | Ubiquitin car | 0 | 0.35591768 | 0 | 0 | 0.00004015 | 0 | 0.00004015 | 0 | 0 | 0 | 0 | 0 | 0 | 0 | 2 | 0 | 2 | 0 | 0 | 0 | 0 | 0 |
| PARN | Poly(A)-speci | 0 | 0.35591768 | 0 | 8.3401E-05 | 0 | 0 | 8.3401E-05 | 0 | 0 | 0 | 0 | 0 | 0 | 2 | 0 | 0 | 2 | 0 | 0 | 0 | 0 | 0 |
| CAN1 | Calpain-1 cat | 0 | 0.35591768 | 0 | 0 | 0 | 7.4694E-05 | 7.4694E-05 | 0 | 0 | 0 | 0 | 0 | 0 | 0 | 0 | 2 | 2 | 0 | 0 | 0 | 0 | 0 |
| MEPCE | 7SK snRNA m | 0 | 0.35591768 | 0 | 0 | 7.6803E-05 | 0 | 7.6803E-05 | 0 | 0 | 0 | 0 | 0 | 0 | 0 | 2 | 0 | 2 | 0 | 0 | 0 | 0 | 0 |
| TYY1 | Transcription | 0 | 0.35591768 | 0 | 0 | 0 | 0.00012882 | 0.00012882 | 0 | 0 | 0 | 0 | 0 | 0 | 0 | 0 | 2 | 2 | 0 | 0 | 0 | 0 | 0 |
| NDE1 | Nuclear distr | 0 | 0.35591768 | 0 | 0 | 0.00022941 | 0 | 0.00022941 | 0 | 0 | 0 | 0 | 0 | 0 | 0 | 3 | 0 | 3 | 0 | 0 | 0 | 0 | 0 |
| INTU | Protein intur | 0 | 0.35591768 | 0 | 5.6574E-05 | 0 | 0 | 5.6574E-05 | 0 | 0 | 0 | 0 | 0 | 0 | 2 | 0 | 0 | 2 | 0 | 0 | 0 | 0 | 0 |
| DIAP3 | Protein diaph | 0 | 0.35591768 | 0 | 0 | 4.4357E-05 | 0 | 4.4357E-05 | 0 | 0 | 0 | 0 | 0 | 0 | 0 | 2 | 0 | 2 | 0 | 0 | 0 | 0 | 0 |
| POP7 | Ribonuclease | 0 | 0.35591768 | 0.00037698 | 0 | 0 | 0 | 0.00037698 | 0 | 0 | 0 | 0 | 0 | 2 | 0 | 0 | 0 | 2 | 0 | 0 | 0 | 0 | 0 |
| VP13A | Vacuolar pro | 0 | 0.35591768 | 0 | 0 | 1.6672E-05 | 0 | 1.6672E-05 | 0 | 0 | 0 | 0 | 0 | 0 | 0 | 2 | 0 | 2 | 0 | 0 | 0 | 0 | 0 |
| SNRPA | U1 small nuc | 0 | 0.35591768 | 0 | 0 | 0.00018765 | 0 | 0.00018765 | 0 | 0 | 0 | 0 | 0 | 0 | 0 | 2 | 0 | 2 | 0 | 0 | 0 | 0 | 0 |
| BUB1B | Mitotic check | 0 | 0.35591768 | 0 | 0 | 7.5596E-05 | 0 | 7.5596E-05 | 0 | 0 | 0 | 0 | 0 | 0 | 0 | 3 | 0 | 3 | 0 | 0 | 0 | 0 | 0 |
| SNUT2 | U4/U6.U5 tri | 0 | 0.35591768 | 0 | 0 | 0 | 0.00014159 | 0.00014159 | 0 | 0 | 0 | 0 | 0 | 0 | 0 | 0 | 3 | 3 | 0 | 0 | 0 | 0 | 0 |
| API5 | Apoptosis inh | 0 | 0.35591768 | 0 | 0 | 0 | 0.00015267 | 0.00015267 | 0 | 0 | 0 | 0 | 0 | 0 | 0 | 0 | 3 | 3 | 0 | 0 | 0 | 0 | 0 |
| CLSPN | Claspin OS=H | 0 | 0.35591768 | 0 | 0 | 0 | 3.9829E-05 | 3.9829E-05 | 0 | 0 | 0 | 0 | 0 | 0 | 0 | 0 | 2 | 2 | 0 | 0 | 0 | 0 | 0 |
| IMA4 | Importin sub | 0 | 0.35591768 | 0 | 0 | 0 | 0.00010236 | 0.00010236 | 0 | 0 | 0 | 0 | 0 | 0 | 0 | 0 | 2 | 2 | 0 | 0 | 0 | 0 | 0 |
| RFA3 | Replication p | 0 | 0.35591768 | 0.00043617 | 0 | 0 | 0 | 0.00043617 | 0 | 0 | 0 | 0 | 0 | 2 | 0 | 0 | 0 | 2 | 0 | 0 | 0 | 0 | 0 |
| KDM1B | Lysine-specif | 0 | 0.35591768 | 0 | 0 | 6.4376E-05 | 0 | 6.4376E-05 | 0 | 0 | 0 | 0 | 0 | 0 | 0 | 2 | 0 | 2 | 0 | 0 | 0 | 0 | 0 |
| WDR18 | WD repeat-c | 0 | 0.35591768 | 0 | 0 | 0.00012249 | 0 | 0.00012249 | 0 | 0 | 0 | 0 | 0 | 0 | 0 | 2 | 0 | 2 | 0 | 0 | 0 | 0 | 0 |
| C19L2 | CWF19-like p | 0 | 0.35591768 | 8.8552E-05 | 0 | 0 | 0 | 8.8552E-05 | 0 | 0 | 0 | 0 | 0 | 3 | 0 | 0 | 0 | 3 | 0 | 0 | 0 | 0 | 0 |
| N4BP1 | NEDD4-bindi | 0 | 0.35591768 | 0 | 0 | 0 | 5.9521E-05 | 5.9521E-05 | 0 | 0 | 0 | 0 | 0 | 0 | 0 | 0 | 2 | 2 | 0 | 0 | 0 | 0 | 0 |
| ARHGI | Rho guanine | 0 | 0.35591768 | 0 | 0.00006815 | 0 | 0 | 0.00006815 | 0 | 0 | 0 | 0 | 0 | 0 | 3 | 0 | 0 | 3 | 0 | 0 | 0 | 0 | 0 |
| VRK2 | Serine/threo | 0 | 0.35591768 | 0 | 0.00010491 | 0 | 0 | 0.00010491 | 0 | 0 | 0 | 0 | 0 | 0 | 2 | 0 | 0 | 2 | 0 | 0 | 0 | 0 | 0 |
| FAKD2 | FAST kinase c | 0 | 0.35591768 | 7.4334E-05 | 0 | 0 | 0 | 7.4334E-05 | 0 | 0 | 0 | 0 | 0 | 2 | 0 | 0 | 0 | 2 | 0 | 0 | 0 | 0 | 0 |
| RPAB1 | DNA-directed | 0 | 0.35591768 | 0 | 0 | 0.00025199 | 0 | 0.00025199 | 0 | 0 | 0 | 0 | 0 | 0 | 0 | 2 | 0 | 2 | 0 | 0 | 0 | 0 | 0 |
| ARC1A | Actin-related | 0 | 0.35591768 | 0 | 0 | 0 | 0.00014414 | 0.00014414 | 0 | 0 | 0 | 0 | 0 | 0 | 0 | 0 | 2 | 2 | 0 | 0 | 0 | 0 | 0 |
| PP1R8 | Nuclear inhib | 0 | 0.35591768 | 0.00015036 | 0 | 0 | 0 | 0.00015036 | 0 | 0 | 0 | 0 | 0 | 2 | 0 | 0 | 0 | 2 | 0 | 0 | 0 | 0 | 0 |
| FA98B | Protein FAM9 | 0 | 0.35591768 | 0 | 0 | 0.00016036 | 0 | 0.00016036 | 0 | 0 | 0 | 0 | 0 | 0 | 0 | 2 | 0 | 2 | 0 | 0 | 0 | 0 | 0 |
| GOGA4 | Golgin subfar | 0 | 0.35591768 | 0.0000355 | 0 | 0 | 0 | 0.0000355 | 0 | 0 | 0 | 0 | 0 | 4 | 0 | 0 | 0 | 4 | 0 | 0 | 0 | 0 | 0 |
| TF2B | Transcription | 0 | 0.35591768 | 0 | 0.00025297 | 0 | 0 | 0.00025297 | 0 | 0 | 0 | 0 | 0 | 0 | 3 | 0 | 0 | 3 | 0 | 0 | 0 | 0 | 0 |
| AKAP2 | A-kinase ancl | 0 | 0.35591768 | 0 | 6.2041E-05 | 0 | 0 | 6.2041E-05 | 0 | 0 | 0 | 0 | 0 | 0 | 2 | 0 | 0 | 2 | 0 | 0 | 0 | 0 | 0 |
| PPGB | Lysosomal pr | 0 | 0.35591768 | 0 | 0 | 0 | 0.00011111 | 0.00011111 | 0 | 0 | 0 | 0 | 0 | 0 | 0 | 0 | 2 | 2 | 0 | 0 | 0 | 0 | 0 |
| SKP2 | S-phase kina | 0 | 0.35591768 | 0 | 0 | 0.00012481 | 0 | 0.00012481 | 0 | 0 | 0 | 0 | 0 | 0 | 0 | 2 | 0 | 2 | 0 | 0 | 0 | 0 | 0 |
| SC31A | Protein trans | 0 | 0.35591768 | 0 | 0 | 4.3375E-05 | 0 | 4.3375E-05 | 0 | 0 | 0 | 0 | 0 | 0 | 0 | 2 | 0 | 2 | 0 | 0 | 0 | 0 | 0 |
| FGOP2 | FGFR1 oncog | 0 | 0.35591768 | 0 | 0.00031597 | 0 | 0 | 0.00031597 | 0 | 0 | 0 | 0 | 0 | 0 | 3 | 0 | 0 | 3 | 0 | 0 | 0 | 0 | 0 |
| MS3L1 | Male-specific | 0 | 0.35591768 | 0.0001013 | 0 | 0 | 0 | 0.0001013 | 0 | 0 | 0 | 0 | 0 | 2 | 0 | 0 | 0 | 2 | 0 | 0 | 0 | 0 | 0 |
| NSE3 | Non-structur | 0 | 0.35591768 | 0 | 0 | 0 | 0.00017543 | 0.00017543 | 0 | 0 | 0 | 0 | 0 | 0 | 0 | 0 | 2 | 2 | 0 | 0 | 0 | 0 | 0 |
| SGO2 | Shugoshin 2 | 0 | 0.35591768 | 0 | 0 | 0 | 4.2159E-05 | 4.2159E-05 | 0 | 0 | 0 | 0 | 0 | 0 | 0 | 0 | 2 | 2 | 0 | 0 | 0 | 0 | 0 |
| MTF2 | Metal-respor | 0 | 0.35591768 | 0.000089 | 0 | 0 | 0 | 0.000089 | 0 | 0 | 0 | 0 | 0 | 2 | 0 | 0 | 0 | 2 | 0 | 0 | 0 | 0 | 0 |
| CLK2 | Dual specifici | 0 | 0.35591768 | 0 | 0 | 0 | 0.00010688 | 0.00010688 | 0 | 0 | 0 | 0 | 0 | 0 | 0 | 0 | 2 | 2 | 0 | 0 | 0 | 0 | 0 |
| ZFP91 | E3 ubiquitin- | 0 | 0.35591768 | 0 | 9.3496E-05 | 0 | 0 | 9.3496E-05 | 0 | 0 | 0 | 0 | 0 | 0 | 2 | 0 | 0 | 2 | 0 | 0 | 0 | 0 | 0 |
| MRP1 | Multidrug res | 0 | 0.35591768 | 0 | 0 | 6.9128E-05 | 0 | 6.9128E-05 | 0 | 0 | 0 | 0 | 0 | 0 | 0 | 4 | 0 | 4 | 0 | 0 | 0 | 0 | 0 |
| TELO2 | Telomere len | 0 | 0.35591768 | 0 | 0 | 6.3223E-05 | 0 | 6.3223E-05 | 0 | 0 | 0 | 0 | 0 | 0 | 0 | 2 | 0 | 2 | 0 | 0 | 0 | 0 | 0 |
| SSRP1 | FACT comple | 0 | 0.35591768 | 0 | 7.5166E-05 | 0 | 0 | 7.5166E-05 | 0 | 0 | 0 | 0 | 0 | 0 | 2 | 0 | 0 | 2 | 0 | 0 | 0 | 0 | 0 |
| KMT2B | Histone-lysin | 0 | 0.35591768 | 0 | 1.9629E-05 | 0 | 0 | 1.9629E-05 | 0 | 0 | 0 | 0 | 0 | 0 | 2 | 0 | 0 | 2 | 0 | 0 | 0 | 0 | 0 |
| PHRF1 | PHD and RIN | 0 | 0.35591768 | 0 | 0 | 3.2091E-05 | 0 | 3.2091E-05 | 0 | 0 | 0 | 0 | 0 | 0 | 0 | 2 | 0 | 2 | 0 | 0 | 0 | 0 | 0 |

| | | | | | | | | | | | | | | | | | | | | | | | |
|---|---|---|---|---|---|---|---|---|---|---|---|---|---|---|---|---|---|---|---|---|---|---|---|
| IF2A | Eukaryotic tr | 0 | 0.35591768 | 0 | 0 | 0.00016799 | 0 | 0.00016799 | 0 | 0 | 0 | 0 | 0 | 0 | 0 | 2 | 0 | 2 | 0 | 0 | 0 | 0 | 0 |
| MAP1A | Microtubule- | 0 | 0.35591768 | 0 | 2.8519E-05 | 0 | 0 | 2.8519E-05 | 0 | 0 | 0 | 0 | 0 | 0 | 4 | 0 | 0 | 4 | 0 | 0 | 0 | 0 | 0 |
| TAF7 | Transcription | 0 | 0.35591768 | 0 | 0 | 0.00015163 | 0 | 0.00015163 | 0 | 0 | 0 | 0 | 0 | 0 | 0 | 2 | 0 | 2 | 0 | 0 | 0 | 0 | 0 |
| AATF | Protein AATF | 0 | 0.35591768 | 9.4244E-05 | 0 | 0 | 0 | 9.4244E-05 | 0 | 0 | 0 | 0 | 0 | 2 | 0 | 0 | 0 | 2 | 0 | 0 | 0 | 0 | 0 |
| ZN335 | Zinc finger pr | 0 | 0.35591768 | 5.8991E-05 | 0 | 0 | 0 | 5.8991E-05 | 0 | 0 | 0 | 0 | 0 | 3 | 0 | 0 | 0 | 3 | 0 | 0 | 0 | 0 | 0 |
| MTX3 | Metaxin-3 OS | 0 | 0.35591768 | 0 | 0 | 0.00016961 | 0 | 0.00016961 | 0 | 0 | 0 | 0 | 0 | 0 | 0 | 2 | 0 | 2 | 0 | 0 | 0 | 0 | 0 |
| IMA6 | Importin sub | 0 | 0.35591768 | 0 | 0.00014914 | 0 | 0 | 0.00014914 | 0 | 0 | 0 | 0 | 0 | 0 | 25 | 0 | 0 | 25 | 0 | 0 | 0 | 0 | 0 |
| PUM3 | Pumilio hom | 0 | 0.35591768 | 0 | 8.2242E-05 | 0 | 0 | 8.2242E-05 | 0 | 0 | 0 | 0 | 0 | 0 | 2 | 0 | 0 | 2 | 0 | 0 | 0 | 0 | 0 |
| MO4L2 | Mortality fac | 0 | 0.35591768 | 0.00027488 | 0 | 0 | 0 | 0.00027488 | 0 | 0 | 0 | 0 | 0 | 3 | 0 | 0 | 0 | 3 | 0 | 0 | 0 | 0 | 0 |
| RGPA1 | Ral GTPase-a | 0 | 0.35591768 | 0 | 0 | 2.5991E-05 | 0 | 2.5991E-05 | 0 | 0 | 0 | 0 | 0 | 0 | 0 | 2 | 0 | 2 | 0 | 0 | 0 | 0 | 0 |
| PPM1A | Protein phos | 0 | 0.35591768 | 0 | 0.00013951 | 0 | 0 | 0.00013951 | 0 | 0 | 0 | 0 | 0 | 0 | 2 | 0 | 0 | 2 | 0 | 0 | 0 | 0 | 0 |
| WASF1 | Wiskott-Aldri | 0 | 0.35591768 | 0.00014162 | 0 | 0 | 0 | 0.00014162 | 0 | 0 | 0 | 0 | 0 | 3 | 0 | 0 | 0 | 3 | 0 | 0 | 0 | 0 | 0 |
| DHE3 | Glutamate de | 0 | 0.35591768 | 0 | 0 | 0 | 9.5576E-05 | 9.5576E-05 | 0 | 0 | 0 | 0 | 0 | 0 | 0 | 0 | 2 | 2 | 0 | 0 | 0 | 0 | 0 |
| MED15 | Mediator of | 0 | 0.35591768 | 0.00010046 | 0 | 0 | 0 | 0.00010046 | 0 | 0 | 0 | 0 | 0 | 3 | 0 | 0 | 0 | 3 | 0 | 0 | 0 | 0 | 0 |
| EXOC1 | Exocyst comp | 0 | 0.35591768 | 0 | 5.9612E-05 | 0 | 0 | 5.9612E-05 | 0 | 0 | 0 | 0 | 0 | 0 | 2 | 0 | 0 | 2 | 0 | 0 | 0 | 0 | 0 |
| BORC7 | BLOC-1-relat | 0 | 0.35591768 | 0 | 0 | 0.00050397 | 0 | 0.00050397 | 0 | 0 | 0 | 0 | 0 | 0 | 0 | 2 | 0 | 2 | 0 | 0 | 0 | 0 | 0 |
| ABCBA | ATP-binding | 0 | 0.35591768 | 7.1513E-05 | 0 | 0 | 0 | 7.1513E-05 | 0 | 0 | 0 | 0 | 0 | 2 | 0 | 0 | 0 | 2 | 0 | 0 | 0 | 0 | 0 |
| AASD1 | Alanyl-tRNA | 0 | 0.35591768 | 0 | 0 | 0 | 0.00012944 | 0.00012944 | 0 | 0 | 0 | 0 | 0 | 0 | 0 | 0 | 2 | 2 | 0 | 0 | 0 | 0 | 0 |
| TTF1 | Transcription | 0 | 0.35591768 | 0 | 5.8887E-05 | 0 | 0 | 5.8887E-05 | 0 | 0 | 0 | 0 | 0 | 0 | 2 | 0 | 0 | 2 | 0 | 0 | 0 | 0 | 0 |
| SALL1 | Sal-like prote | 0 | 0.35591768 | 7.9723E-05 | 0 | 0 | 0 | 7.9723E-05 | 0 | 0 | 0 | 0 | 0 | 5 | 0 | 0 | 0 | 5 | 0 | 0 | 0 | 0 | 0 |
| CL043 | Uncharacteri | 0 | 0.35591768 | 0 | 0 | 0.00020197 | 0 | 0.00020197 | 0 | 0 | 0 | 0 | 0 | 0 | 0 | 2 | 0 | 2 | 0 | 0 | 0 | 0 | 0 |
| NECT2 | Nectin-2 OS= | 0 | 0.35591768 | 0 | 0 | 9.8359E-05 | 0 | 9.8359E-05 | 0 | 0 | 0 | 0 | 0 | 0 | 0 | 2 | 0 | 2 | 0 | 0 | 0 | 0 | 0 |
| I2BP2 | Interferon re | 0 | 0.35591768 | 0 | 0 | 0 | 0.00013628 | 0.00013628 | 0 | 0 | 0 | 0 | 0 | 0 | 0 | 0 | 3 | 3 | 0 | 0 | 0 | 0 | 0 |
| TMOD1 | Tropomoduli | 0 | 0.35591768 | 0 | 0 | 0 | 0.00022283 | 0.00022283 | 0 | 0 | 0 | 0 | 0 | 0 | 0 | 0 | 3 | 3 | 0 | 0 | 0 | 0 | 0 |
| SMAL1 | SWI/SNF-rela | 0 | 0.35591768 | 0 | 0 | 0 | 8.3854E-05 | 8.3854E-05 | 0 | 0 | 0 | 0 | 0 | 0 | 0 | 0 | 3 | 3 | 0 | 0 | 0 | 0 | 0 |
| PK3C3 | Phosphatidyl | 0 | 0.35591768 | 0 | 0 | 0 | 6.0125E-05 | 6.0125E-05 | 0 | 0 | 0 | 0 | 0 | 0 | 0 | 0 | 2 | 2 | 0 | 0 | 0 | 0 | 0 |
| PCGF1 | Polycomb gro | 0 | 0.35591768 | 0 | 0 | 0.00020431 | 0 | 0.00020431 | 0 | 0 | 0 | 0 | 0 | 0 | 0 | 2 | 0 | 2 | 0 | 0 | 0 | 0 | 0 |
| TET2 | Methylcytosi | 0 | 0.35591768 | 0 | 0.00003993 | 0 | 0 | 0.00003993 | 0 | 0 | 0 | 0 | 0 | 0 | 3 | 0 | 0 | 3 | 0 | 0 | 0 | 0 | 0 |
| T2EB | Transcription | 0 | 0.35591768 | 0 | 0.00018314 | 0 | 0 | 0.00018314 | 0 | 0 | 0 | 0 | 0 | 0 | 2 | 0 | 0 | 2 | 0 | 0 | 0 | 0 | 0 |
| KCRB | Creatine kina | 0 | 0.35591768 | 0.00013852 | 0 | 0 | 0 | 0.00013852 | 0 | 0 | 0 | 0 | 0 | 2 | 0 | 0 | 0 | 2 | 0 | 0 | 0 | 0 | 0 |
| MLH1 | DNA mismatc | 0 | 0.35591768 | 0 | 0 | 6.9997E-05 | 0 | 6.9997E-05 | 0 | 0 | 0 | 0 | 0 | 0 | 0 | 2 | 0 | 2 | 0 | 0 | 0 | 0 | 0 |
| PP4C | Serine/threo | 0 | 0.35591768 | 0 | 0 | 0.00017237 | 0 | 0.00017237 | 0 | 0 | 0 | 0 | 0 | 0 | 0 | 2 | 0 | 2 | 0 | 0 | 0 | 0 | 0 |
| LRRF1 | Leucine-rich | 0 | 0.35591768 | 0 | 0 | 9.8238E-05 | 0 | 9.8238E-05 | 0 | 0 | 0 | 0 | 0 | 0 | 0 | 3 | 0 | 3 | 0 | 0 | 0 | 0 | 0 |
| DHB11 | Estradiol 17- | 0 | 0.35591768 | 0 | 0 | 0 | 0.00026666 | 0.00026666 | 0 | 0 | 0 | 0 | 0 | 0 | 0 | 0 | 3 | 3 | 0 | 0 | 0 | 0 | 0 |
| PQBP1 | Polyglutamin | 0 | 0.35591768 | 0 | 0.00030166 | 0 | 0 | 0.00030166 | 0 | 0 | 0 | 0 | 0 | 0 | 3 | 0 | 0 | 3 | 0 | 0 | 0 | 0 | 0 |
| SMC6 | Structural ma | 0 | 0.35591768 | 0 | 0 | 4.8504E-05 | 0 | 4.8504E-05 | 0 | 0 | 0 | 0 | 0 | 0 | 0 | 2 | 0 | 2 | 0 | 0 | 0 | 0 | 0 |
| ACTN1 | Alpha-actinin | 0 | 0.35591768 | 0 | 0 | 0 | 5.9788E-05 | 5.9788E-05 | 0 | 0 | 0 | 0 | 0 | 0 | 0 | 0 | 5 | 5 | 0 | 0 | 0 | 0 | 0 |
| ABRX1 | BRCA1-A com | 0 | 0.35591768 | 0 | 0 | 0 | 0.00026079 | 0.00026079 | 0 | 0 | 0 | 0 | 0 | 0 | 0 | 0 | 4 | 4 | 0 | 0 | 0 | 0 | 0 |
| HTR5B | HEAT repeat- | 0 | 0.35591768 | 0 | 0 | 0 | 2.5751E-05 | 2.5751E-05 | 0 | 0 | 0 | 0 | 0 | 0 | 0 | 0 | 2 | 2 | 0 | 0 | 0 | 0 | 0 |
| ZNT7 | Zinc transpor | 0 | 0.35591768 | 0 | 0 | 0 | 0.00014184 | 0.00014184 | 0 | 0 | 0 | 0 | 0 | 0 | 0 | 0 | 2 | 2 | 0 | 0 | 0 | 0 | 0 |
| GLMN | Glomulin OS= | 0 | 0.35591768 | 0 | 0 | 0 | 8.9783E-05 | 8.9783E-05 | 0 | 0 | 0 | 0 | 0 | 0 | 0 | 0 | 2 | 2 | 0 | 0 | 0 | 0 | 0 |
| UBXN4 | UBX domain- | 0 | 0.35591768 | 0 | 0.00010491 | 0 | 0 | 0.00010491 | 0 | 0 | 0 | 0 | 0 | 0 | 2 | 0 | 0 | 2 | 0 | 0 | 0 | 0 | 0 |
| RPP29 | Ribonuclease | 0 | 0.35591768 | 0 | 0 | 0.00024053 | 0 | 0.00024053 | 0 | 0 | 0 | 0 | 0 | 0 | 0 | 2 | 0 | 2 | 0 | 0 | 0 | 0 | 0 |
| S38A1 | Sodium-coup | 0 | 0.35591768 | 0 | 0 | 0 | 0.00016426 | 0.00016426 | 0 | 0 | 0 | 0 | 0 | 0 | 0 | 0 | 3 | 3 | 0 | 0 | 0 | 0 | 0 |
| C19L1 | CWF19-like p | 0 | 0.35591768 | 0 | 9.9058E-05 | 0 | 0 | 9.9058E-05 | 0 | 0 | 0 | 0 | 0 | 0 | 2 | 0 | 0 | 2 | 0 | 0 | 0 | 0 | 0 |
| F169A | Soluble lamir | 0 | 0.35591768 | 0 | 0 | 0 | 7.9599E-05 | 7.9599E-05 | 0 | 0 | 0 | 0 | 0 | 0 | 0 | 0 | 2 | 2 | 0 | 0 | 0 | 0 | 0 |
| CTNL1 | Alpha-catulin | 0 | 0.35591768 | 0 | 7.2606E-05 | 0 | 0 | 7.2606E-05 | 0 | 0 | 0 | 0 | 0 | 0 | 2 | 0 | 0 | 2 | 0 | 0 | 0 | 0 | 0 |
| AF9 | Protein AF-9 | 0 | 0.35591768 | 0 | 0 | 0 | 9.3893E-05 | 9.3893E-05 | 0 | 0 | 0 | 0 | 0 | 0 | 0 | 0 | 2 | 2 | 0 | 0 | 0 | 0 | 0 |
| ILK | Integrin-linke | 0 | 0.35591768 | 0 | 0.00017686 | 0 | 0 | 0.00017686 | 0 | 0 | 0 | 0 | 0 | 0 | 3 | 0 | 0 | 3 | 0 | 0 | 0 | 0 | 0 |
| TDIF1 | Deoxynucleo | 0 | 0.35591768 | 0 | 0.00016198 | 0 | 0 | 0.00016198 | 0 | 0 | 0 | 0 | 0 | 0 | 2 | 0 | 0 | 2 | 0 | 0 | 0 | 0 | 0 |
| TTC4 | Tetratricope | 0 | 0.35591768 | 0 | 0.00020656 | 0 | 0 | 0.00020656 | 0 | 0 | 0 | 0 | 0 | 0 | 3 | 0 | 0 | 3 | 0 | 0 | 0 | 0 | 0 |
| CELF2 | CUGBP Elav-l | 0 | 0.35591768 | 0 | 0.00010491 | 0 | 0 | 0.00010491 | 0 | 0 | 0 | 0 | 0 | 0 | 2 | 0 | 0 | 2 | 0 | 0 | 0 | 0 | 0 |
| IPYR2 | Inorganic pyr | 0 | 0.35591768 | 0 | 0.00023934 | 0 | 0 | 0.00023934 | 0 | 0 | 0 | 0 | 0 | 0 | 3 | 0 | 0 | 3 | 0 | 0 | 0 | 0 | 0 |
| RT02 | 28S riboson | 0 | 0.35591768 | 0 | 0 | 0 | 0.00018017 | 0.00018017 | 0 | 0 | 0 | 0 | 0 | 0 | 0 | 0 | 2 | 2 | 0 | 0 | 0 | 0 | 0 |
| PEX3 | Peroxisomal | 0 | 0.35591768 | 0 | 0.00021431 | 0 | 0 | 0.00021431 | 0 | 0 | 0 | 0 | 0 | 0 | 3 | 0 | 0 | 3 | 0 | 0 | 0 | 0 | 0 |
| RMD5A | Protein RMD | 0 | 0.35591768 | 0 | 0 | 0.00013534 | 0 | 0.00013534 | 0 | 0 | 0 | 0 | 0 | 0 | 0 | 2 | 0 | 2 | 0 | 0 | 0 | 0 | 0 |
| SRGP2 | SLIT-ROBO R | 0 | 0.35591768 | 0 | 0 | 0 | 4.9796E-05 | 4.9796E-05 | 0 | 0 | 0 | 0 | 0 | 0 | 0 | 0 | 2 | 2 | 0 | 0 | 0 | 0 | 0 |
| VPS72 | Vacuolar pro | 0 | 0.35591768 | 0 | 0 | 0.00014538 | 0 | 0.00014538 | 0 | 0 | 0 | 0 | 0 | 0 | 0 | 2 | 0 | 2 | 0 | 0 | 0 | 0 | 0 |
| CAPON | Carboxyl-terr | 0 | 0.35591768 | 0 | 0 | 0 | 0.0001054 | 0.0001054 | 0 | 0 | 0 | 0 | 0 | 0 | 0 | 0 | 2 | 2 | 0 | 0 | 0 | 0 | 0 |
| BEND3 | BEN domain- | 0 | 0.35591768 | 0 | 0 | 0.00006391 | 0 | 0.00006391 | 0 | 0 | 0 | 0 | 0 | 0 | 0 | 2 | 0 | 2 | 0 | 0 | 0 | 0 | 0 |
| ZN609 | Zinc finger pr | 0 | 0.35591768 | 3.7404E-05 | 0 | 0 | 0 | 3.7404E-05 | 0 | 0 | 0 | 0 | 0 | 2 | 0 | 0 | 0 | 2 | 0 | 0 | 0 | 0 | 0 |
| ZKSC3 | Zinc finger pr | 0 | 0.35591768 | 9.8098E-05 | 0 | 0 | 0 | 9.8098E-05 | 0 | 0 | 0 | 0 | 0 | 3 | 0 | 0 | 0 | 3 | 0 | 0 | 0 | 0 | 0 |
| PPIH | Peptidyl-prol | 0 | 0.35591768 | 0 | 0.00045164 | 0 | 0 | 0.00045164 | 0 | 0 | 0 | 0 | 0 | 0 | 3 | 0 | 0 | 3 | 0 | 0 | 0 | 0 | 0 |
| F207A | Protein FAM2 | 0 | 0.35591768 | 0 | 0.00034756 | 0 | 0 | 0.00034756 | 0 | 0 | 0 | 0 | 0 | 0 | 3 | 0 | 0 | 3 | 0 | 0 | 0 | 0 | 0 |
| TBPL1 | TATA box-bir | 0 | 0.35591768 | 0 | 0 | 0.0002845 | 0 | 0.0002845 | 0 | 0 | 0 | 0 | 0 | 0 | 0 | 2 | 0 | 2 | 0 | 0 | 0 | 0 | 0 |
| CASP | Protein CASP | 0 | 0.35591768 | 0 | 0 | 7.8049E-05 | 0 | 7.8049E-05 | 0 | 0 | 0 | 0 | 0 | 0 | 0 | 16 | 0 | 16 | 0 | 0 | 0 | 0 | 0 |
| KAT6A | Histone acety | 0 | 0.35591768 | 2.6336E-05 | 0 | 0 | 0 | 2.6336E-05 | 0 | 0 | 0 | 0 | 0 | 2 | 0 | 0 | 0 | 2 | 0 | 0 | 0 | 0 | 0 |
| ALG12 | Dol-P-Man:M | 0 | 0.35591768 | 0 | 0 | 0.00010844 | 0 | 0.00010844 | 0 | 0 | 0 | 0 | 0 | 0 | 0 | 2 | 0 | 2 | 0 | 0 | 0 | 0 | 0 |
| TCAL1 | Transcription | 0 | 0.35591768 | 0 | 0 | 0.00033705 | 0 | 0.00033705 | 0 | 0 | 0 | 0 | 0 | 0 | 0 | 2 | 0 | 2 | 0 | 0 | 0 | 0 | 0 |
| PARVA | Alpha-parvin | 0 | 0.35591768 | 0.00014187 | 0 | 0 | 0 | 0.00014187 | 0 | 0 | 0 | 0 | 0 | 2 | 0 | 0 | 0 | 2 | 0 | 0 | 0 | 0 | 0 |
| CELF1 | CUGBP Elav-l | 0 | 0.35591768 | 0 | 0 | 0 | 0.00010974 | 0.00010974 | 0 | 0 | 0 | 0 | 0 | 0 | 0 | 0 | 2 | 2 | 0 | 0 | 0 | 0 | 0 |
| AT7L1 | Ataxin-7-like | 0 | 0.35591768 | 0 | 0 | 9.2191E-05 | 0 | 9.2191E-05 | 0 | 0 | 0 | 0 | 0 | 0 | 0 | 3 | 0 | 3 | 0 | 0 | 0 | 0 | 0 |
| BRAP | BRCA1-assoc | 0 | 0.35591768 | 0 | 0 | 0 | 9.0087E-05 | 9.0087E-05 | 0 | 0 | 0 | 0 | 0 | 0 | 0 | 0 | 2 | 2 | 0 | 0 | 0 | 0 | 0 |
| CHK1 | Serine/threo | 0 | 0.35591768 | 0 | 0.00011196 | 0 | 0 | 0.00011196 | 0 | 0 | 0 | 0 | 0 | 0 | 2 | 0 | 0 | 2 | 0 | 0 | 0 | 0 | 0 |
| PSF3 | DNA replicati | 0 | 0.35591768 | 0 | 0.00024673 | 0 | 0 | 0.00024673 | 0 | 0 | 0 | 0 | 0 | 0 | 2 | 0 | 0 | 2 | 0 | 0 | 0 | 0 | 0 |
| DPOLQ | DNA polymer | 0 | 0.35591768 | 2.0377E-05 | 0 | 0 | 0 | 2.0377E-05 | 0 | 0 | 0 | 0 | 0 | 2 | 0 | 0 | 0 | 2 | 0 | 0 | 0 | 0 | 0 |
| CUL3 | Cullin-3 OS=H | 0 | 0.35591768 | 0 | 6.9392E-05 | 0 | 0 | 6.9392E-05 | 0 | 0 | 0 | 0 | 0 | 0 | 2 | 0 | 0 | 2 | 0 | 0 | 0 | 0 | 0 |
| CDKAL | Threonylcarb | 0 | 0.35591768 | 0 | 0 | 0 | 9.2109E-05 | 9.2109E-05 | 0 | 0 | 0 | 0 | 0 | 0 | 0 | 0 | 2 | 2 | 0 | 0 | 0 | 0 | 0 |
| TFG | Protein TFG | 0 | 0.35591768 | 0 | 0 | 0.00013229 | 0 | 0.00013229 | 0 | 0 | 0 | 0 | 0 | 0 | 0 | 2 | 0 | 2 | 0 | 0 | 0 | 0 | 0 |
| NR2C1 | Nuclear rece | 0 | 0.35591768 | 0 | 0.00008838 | 0 | 0 | 0.00008838 | 0 | 0 | 0 | 0 | 0 | 0 | 2 | 0 | 0 | 2 | 0 | 0 | 0 | 0 | 0 |
| WWP1 | NEDD4-like E | 0 | 0.35591768 | 0 | 0 | 5.7394E-05 | 0 | 5.7394E-05 | 0 | 0 | 0 | 0 | 0 | 0 | 0 | 2 | 0 | 2 | 0 | 0 | 0 | 0 | 0 |
| CT027 | UPF0687 pro | 0 | 0.35591768 | 0 | 0.00030628 | 0 | 0 | 0.00030628 | 0 | 0 | 0 | 0 | 0 | 0 | 2 | 0 | 0 | 2 | 0 | 0 | 0 | 0 | 0 |
| TRPM7 | Transient rec | 0 | 0.35591768 | 0 | 2.8575E-05 | 0 | 0 | 2.8575E-05 | 0 | 0 | 0 | 0 | 0 | 0 | 2 | 0 | 0 | 2 | 0 | 0 | 0 | 0 | 0 |
| PRDX1 | Peroxiredoxi | 0 | 0.13794297 | 0 | 0.00053561 | 0 | 0.00066999 | 0.0006028 | 0 | 0 | 0 | 0 | 0 | 0 | 4 | 0 | 5 | 4.5 | 0 | 0 | 0 | 0 | 0 |
| M4K4 | Mitogen-acti | 0 | 0.13398221 | 4.2596E-05 | 4.3013E-05 | 0 | 0 | 4.2805E-05 | 0 | 0 | 0 | 0 | 0 | 2 | 2 | 0 | 0 | 2 | 0 | 0 | 0 | 0 | 0 |
| STRN4 | Striatin-4 OS | 0 | 0.13397947 | 0 | 0 | 7.0275E-05 | 7.0825E-05 | 0.00007055 | 0 | 0 | 0 | 0 | 0 | 0 | 0 | 3 | 2 | 2.5 | 0 | 0 | 0 | 0 | 0 |
| KHDR1 | KH domain-c | 0 | 0.13397517 | 0.0001787 | 0 | 0.00017918 | 0 | 0.00017894 | 0 | 0 | 0 | 0 | 0 | 3 | 0 | 3 | 0 | 3 | 0 | 0 | 0 | 0 | 0 |
| STX5 | Syntaxin-5 O | 0 | 0.13659744 | 0 | 0.00045036 | 0 | 0.00037557 | 0.00041297 | 0 | 0 | 0 | 0 | 0 | 0 | 6 | 0 | 5 | 5.5 | 0 | 0 | 0 | 0 | 0 |
| RU2B | U2 small nuc | 0 | 0.16908107 | 0.00035185 | 0.00071057 | 0 | 0 | 0.00053121 | 0 | 0 | 0 | 0 | 0 | 3 | 6 | 0 | 0 | 4.5 | 0 | 0 | 0 | 0 | 0 |
| RB12B | RNA-binding | 0 | 0.13397862 | 0 | 0.00005324 | 5.2864E-05 | 0 | 5.3052E-05 | 0 | 0 | 0 | 0 | 0 | 0 | 2 | 2 | 0 | 2 | 0 | 0 | 0 | 0 | 0 |
| P4R3A | Serine/threo | 0 | 0.13397862 | 0 | 9.5966E-05 | 9.5289E-05 | 0 | 9.5628E-05 | 0 | 0 | 0 | 0 | 0 | 0 | 3 | 3 | 0 | 3 | 0 | 0 | 0 | 0 | 0 |
| DNMT1 | DNA (cytosin | 0 | 0.13397859 | 0 | 3.2978E-05 | 3.2746E-05 | 0 | 3.2862E-05 | 0 | 0 | 0 | 0 | 0 | 0 | 2 | 2 | 0 | 2 | 0 | 0 | 0 | 0 | 0 |
| SRSF9 | Serine/argini | 0 | 0.15378186 | 0.00035821 | 0 | 0.00059861 | 0 | 0.00047841 | 0 | 0 | 0 | 0 | 0 | 3 | 0 | 5 | 0 | 4 | 0 | 0 | 0 | 0 | 0 |
| PRAF3 | PRA1 family | 0 | 0.16890947 | 0 | 0 | 0.00028148 | 0.00056735 | 0.00042442 | 0 | 0 | 0 | 0 | 0 | 0 | 0 | 2 | 4 | 3 | 0 | 0 | 0 | 0 | 0 |
| IFT27 | Intraflagellar | 0 | 0.13397947 | 0 | 0 | 0.00042675 | 0.00043009 | 0.00042842 | 0 | 0 | 0 | 0 | 0 | 0 | 0 | 3 | 3 | 3 | 0 | 0 | 0 | 0 | 0 |
| EFTS | Elongation fa | 0 | 0.14647286 | 0.00024359 | 0 | 0.00016282 | 0 | 0.00020321 | 0 | 0 | 0 | 0 | 0 | 3 | 0 | 2 | 0 | 2.5 | 0 | 0 | 0 | 0 | 0 |
| GBF1 | Golgi-specific | 0 | 0.16760863 | 0 | 2.8668E-05 | 5.6931E-05 | 0 | 4.28E-05 | 0 | 0 | 0 | 0 | 0 | 0 | 2 | 4 | 0 | 3 | 0 | 0 | 0 | 0 | 0 |
| CSN1 | COP9 signalo | 0 | 0.14678765 | 0.00021498 | 0 | 0.00032332 | 0 | 0.00026915 | 0 | 0 | 0 | 0 | 0 | 4 | 0 | 6 | 0 | 5 | 0 | 0 | 0 | 0 | 0 |
| HERC2 | E3 ubiquitin- | 0 | 0.13397515 | 1.6377E-05 | 0 | 0.00001642 | 0 | 1.6399E-05 | 0 | 0 | 0 | 0 | 0 | 3 | 0 | 3 | 0 | 3 | 0 | 0 | 0 | 0 | 0 |
| RL19 | 60S ribosoma | 0 | 0.13397946 | 0 | 0 | 0.00026999 | 0.0002721 | 0.00027105 | 0 | 0 | 0 | 0 | 0 | 0 | 0 | 2 | 2 | 2 | 0 | 0 | 0 | 0 | 0 |
| BRD2 | Bromodomai | 0 | 0.16908471 | 6.5889E-05 | 0.00013307 | 0 | 0 | 9.948E-05 | 0 | 0 | 0 | 0 | 0 | 2 | 4 | 0 | 0 | 3 | 0 | 0 | 0 | 0 | 0 |
| ZHX2 | Zinc fingers a | 0 | 0.13397463 | 0 | 0.00015918 | 0 | 0.00015929 | 0.00015924 | 0 | 0 | 0 | 0 | 0 | 0 | 5 | 0 | 6 | 5.5 | 0 | 0 | 0 | 0 | 0 |
| DYL1 | Dynein light | 0 | 0.13397516 | 0.000593 | 0 | 0.00059458 | 0 | 0.00059379 | 0 | 0 | 0 | 0 | 0 | 2 | 0 | 2 | 0 | 2 | 0 | 0 | 0 | 0 | 0 |
| JIP4 | C-Jun-amino- | 0 | 0.16846404 | 5.9928E-05 | 0 | 0.00012018 | 0 | 9.0054E-05 | 0 | 0 | 0 | 0 | 0 | 3 | 0 | 6 | 0 | 4.5 | 0 | 0 | 0 | 0 | 0 |
| HTR5A | HEAT repeat- | 0 | 0.14658606 | 0 | 3.9186E-05 | 0 | 2.6143E-05 | 3.2665E-05 | 0 | 0 | 0 | 0 | 0 | 0 | 3 | 0 | 2 | 2.5 | 0 | 0 | 0 | 0 | 0 |
| EDF1 | Endothelial d | 0 | 0.14658582 | 0 | 0.00054013 | 0 | 0.00036035 | 0.00045024 | 0 | 0 | 0 | 0 | 0 | 0 | 3 | 0 | 2 | 2.5 | 0 | 0 | 0 | 0 | 0 |
| TEX2 | Testis-expres | 0 | 0.14658698 | 0 | 7.0931E-05 | 0 | 4.7321E-05 | 5.9126E-05 | 0 | 0 | 0 | 0 | 0 | 0 | 3 | 0 | 2 | 2.5 | 0 | 0 | 0 | 0 | 0 |
| DHB4 | Peroxisomal | 0 | 0.1528455 | 0.00017927 | 0 | 0 | 0.00010869 | 0.00014398 | 0 | 0 | 0 | 0 | 0 | 5 | 0 | 0 | 3 | 4 | 0 | 0 | 0 | 0 | 0 |
| SURF6 | Surfeit locus | 0 | 0.18803577 | 0.00036549 | 0 | 0 | 0.00014773 | 0.00025661 | 0 | 0 | 0 | 0 | 0 | 5 | 0 | 0 | 2 | 3.5 | 0 | 0 | 0 | 0 | 0 |
| CRNL1 | Crooked neck | 0 | 0.13397859 | 0 | 6.2845E-05 | 6.2403E-05 | 0 | 6.2624E-05 | 0 | 0 | 0 | 0 | 0 | 0 | 2 | 2 | 0 | 2 | 0 | 0 | 0 | 0 | 0 |
| LRBA | Lipopolysacc | 0 | 0.16816403 | 0 | 3.7229E-05 | 0 | 1.8628E-05 | 2.7929E-05 | 0 | 0 | 0 | 0 | 0 | 0 | 4 | 0 | 2 | 3 | 0 | 0 | 0 | 0 | 0 |
| AKP8L | A-kinase ancl | 0 | 0.14601211 | 0.00012255 | 0 | 0 | 8.2556E-05 | 0.00010255 | 0 | 0 | 0 | 0 | 0 | 3 | 0 | 0 | 2 | 2.5 | 0 | 0 | 0 | 0 | 0 |
| SNP47 | Synaptosoma | 0 | 0.1466713 | 0 | 0.00011486 | 0 | 0.00017241 | 0.00014364 | 0 | 0 | 0 | 0 | 0 | 0 | 2 | 0 | 3 | 2.5 | 0 | 0 | 0 | 0 | 0 |
| TAM41 | Phosphatidat | 0 | 0.14647121 | 0.00017514 | 0 | 0.00011707 | 0 | 0.00014611 | 0 | 0 | 0 | 0 | 0 | 3 | 0 | 2 | 0 | 2.5 | 0 | 0 | 0 | 0 | 0 |
| AURKB | Aurora kinas | 0 | 0.14601114 | 0.00023013 | 0 | 0 | 0.00015503 | 0.00019258 | 0 | 0 | 0 | 0 | 0 | 3 | 0 | 0 | 2 | 2.5 | 0 | 0 | 0 | 0 | 0 |
| PSMD8 | 26S proteaso | 0 | 0.13397464 | 0 | 0.00030453 | 0 | 0.00030475 | 0.00030464 | 0 | 0 | 0 | 0 | 0 | 0 | 4 | 0 | 4 | 4 | 0 | 0 | 0 | 0 | 0 |
| RFC4 | Replication fa | 0 | 0.13826482 | 0.00029078 | 0.00036703 | 0 | 0 | 0.00032891 | 0 | 0 | 0 | 0 | 0 | 4 | 5 | 0 | 0 | 4.5 | 0 | 0 | 0 | 0 | 0 |
| RHOG | Rho-related | 0 | 0.18803554 | 0.0006908 | 0 | 0 | 0.00027922 | 0.00048501 | 0 | 0 | 0 | 0 | 0 | 5 | 0 | 0 | 2 | 3.5 | 0 | 0 | 0 | 0 | 0 |
| SMN | Survival moto | 0 | 0.13397517 | 0.00026927 | 0 | 0.00026999 | 0 | 0.00026963 | 0 | 0 | 0 | 0 | 0 | 3 | 0 | 3 | 0 | 3 | 0 | 0 | 0 | 0 | 0 |
| SCML2 | Sex comb on | 0 | 0.1460508 | 0.00011309 | 7.6133E-05 | 0 | 0 | 9.4612E-05 | 0 | 0 | 0 | 0 | 0 | 3 | 2 | 0 | 0 | 2.5 | 0 | 0 | 0 | 0 | 0 |
| RAD21 | Double-stran | 0 | 0.16846289 | 0.00008364 | 0 | 0.00016773 | 0 | 0.00012569 | 0 | 0 | 0 | 0 | 0 | 2 | 0 | 4 | 0 | 3 | 0 | 0 | 0 | 0 | 0 |
| RPAC1 | DNA-directed | 0 | 0.15284656 | 0.00038134 | 0 | 0 | 0.0002312 | 0.00030627 | 0 | 0 | 0 | 0 | 0 | 5 | 0 | 0 | 3 | 4 | 0 | 0 | 0 | 0 | 0 |
| TAF4B | Transcription | 0 | 0.14678956 | 6.1226E-05 | 0 | 9.2084E-05 | 0 | 7.6655E-05 | 0 | 0 | 0 | 0 | 0 | 4 | 0 | 5 | 0 | 4.5 | 0 | 0 | 0 | 0 | 0 |
| CARF | CDKN2A-inte | 0 | 0.2067668 | 0.00027298 | 0 | 9.1237E-05 | 0 | 0.00018211 | 0 | 0 | 0 | 0 | 0 | 6 | 0 | 2 | 0 | 4 | 0 | 0 | 0 | 0 | 0 |
| SLTM | SAFB-like tra | 0 | 0.1466718 | 0 | 5.1541E-05 | 0 | 7.7366E-05 | 6.4454E-05 | 0 | 0 | 0 | 0 | 0 | 0 | 2 | 0 | 3 | 2.5 | 0 | 0 | 0 | 0 | 0 |
| IPP2 | Protein phos | 0 | 0.14658738 | 0 | 0.0007799 | 0 | 0.0005203 | 0.0006501 | 0 | 0 | 0 | 0 | 0 | 0 | 6 | 0 | 4 | 5 | 0 | 0 | 0 | 0 | 0 |
| ASXL2 | Putative Poly | 0 | 0.14678937 | 3.6778E-05 | 0 | 5.5314E-05 | 0 | 4.6046E-05 | 0 | 0 | 0 | 0 | 0 | 2 | 0 | 3 | 0 | 2.5 | 0 | 0 | 0 | 0 | 0 |
| INT13 | Integrator co | 0 | 0.13398219 | 7.4755E-05 | 7.5486E-05 | 0 | 0 | 7.5121E-05 | 0 | 0 | 0 | 0 | 0 | 2 | 2 | 0 | 0 | 2 | 0 | 0 | 0 | 0 | 0 |
| MKL1 | MKL/myocar | 0 | 0.13398218 | 8.5033E-05 | 8.5864E-05 | 0 | 0 | 8.5449E-05 | 0 | 0 | 0 | 0 | 0 | 3 | 3 | 0 | 0 | 3 | 0 | 0 | 0 | 0 | 0 |
| SNX1 | Sorting nexin | 0 | 0.14658465 | 0 | 0.00015314 | 0 | 0.00010217 | 0.00012766 | 0 | 0 | 0 | 0 | 0 | 0 | 3 | 0 | 2 | 2.5 | 0 | 0 | 0 | 0 | 0 |
| SCAFB | Protein SCAF | 0 | 0.14605336 | 5.4112E-05 | 3.6427E-05 | 0 | 0 | 4.527E-05 | 0 | 0 | 0 | 0 | 0 | 3 | 2 | 0 | 0 | 2.5 | 0 | 0 | 0 | 0 | 0 |
| SMRCD | SWI/SNF-rela | 0 | 0.14679007 | 5.1439E-05 | 0 | 7.7365E-05 | 0 | 6.4402E-05 | 0 | 0 | 0 | 0 | 0 | 2 | 0 | 3 | 0 | 2.5 | 0 | 0 | 0 | 0 | 0 |
| CSN4 | COP9 signalo | 0 | 0.18974088 | 0 | 0.00032816 | 0.00013034 | 0 | 0.00022925 | 0 | 0 | 0 | 0 | 0 | 0 | 5 | 2 | 0 | 3.5 | 0 | 0 | 0 | 0 | 0 |

| | | | | | | | | | | | | | | | | | | | | | | | |
|---|---|---|---|---|---|---|---|---|---|---|---|---|---|---|---|---|---|---|---|---|---|---|---|
| YETS4 | YEATS domai | 0 | 0.13398334 | 0.0002325 | 0 | 0 | 0.00023494 | 0.00023372 | 0 | 0 | 0 | 0 | 0 | 2 | 0 | 0 | 2 | 2 | 0 | 0 | 0 | 0 | 0 |
| SKP1 | S-phase kina | 0 | 0.13397516 | 0.00048568 | 0 | 0.00048697 | 0 | 0.00048633 | 0 | 0 | 0 | 0 | 0 | 3 | 0 | 3 | 0 | 3 | 0 | 0 | 0 | 0 | 0 |
| TP53B | TP53-binding | 0 | 0.16760991 | 0 | 2.7025E-05 | 5.3669E-05 | 0 | 4.0347E-05 | 0 | 0 | 0 | 0 | 0 | 0 | 2 | 4 | 0 | 3 | 0 | 0 | 0 | 0 | 0 |
| ZN830 | Zinc finger pr | 0 | 0.13397464 | 0 | 0.00021489 | 0 | 0.00021505 | 0.00021497 | 0 | 0 | 0 | 0 | 0 | 0 | 3 | 0 | 3 | 3 | 0 | 0 | 0 | 0 | 0 |
| ARPC3 | Actin-related | 0 | 0.13397516 | 0.0002965 | 0 | 0.00029729 | 0 | 0.0002969 | 0 | 0 | 0 | 0 | 0 | 2 | 0 | 2 | 0 | 2 | 0 | 0 | 0 | 0 | 0 |
| TPD52 | Tumor protei | 0 | 0.14667154 | 0 | 0.00023792 | 0 | 0.00035713 | 0.00029753 | 0 | 0 | 0 | 0 | 0 | 0 | 2 | 0 | 3 | 2.5 | 0 | 0 | 0 | 0 | 0 |
| RU1C | U1 small nuc | 0 | 0.1380112 | 0.00066386 | 0 | 0.00083203 | 0 | 0.00074795 | 0 | 0 | 0 | 0 | 0 | 4 | 0 | 5 | 0 | 4.5 | 0 | 0 | 0 | 0 | 0 |
| MYBB | Myb-related | 0 | 0.1339795 | 0 | 0 | 0.00011339 | 0.00011428 | 0.00011384 | 0 | 0 | 0 | 0 | 0 | 0 | 0 | 3 | 3 | 3 | 0 | 0 | 0 | 0 | 0 |
| ZDBF2 | DBF4-type zi | 0 | 0.14658553 | 0 | 3.3959E-05 | 0 | 2.2656E-05 | 2.8308E-05 | 0 | 0 | 0 | 0 | 0 | 0 | 3 | 0 | 2 | 2.5 | 0 | 0 | 0 | 0 | 0 |
| PLOD2 | Procollagen-l | 0 | 0.1472602 | 0.00007161 | 0 | 0 | 0.00010854 | 9.0075E-05 | 0 | 0 | 0 | 0 | 0 | 2 | 0 | 0 | 3 | 2.5 | 0 | 0 | 0 | 0 | 0 |
| SAMH1 | Deoxynucleo | 0 | 0.14678986 | 8.4308E-05 | 0 | 0.0001268 | 0 | 0.00010555 | 0 | 0 | 0 | 0 | 0 | 2 | 0 | 3 | 0 | 2.5 | 0 | 0 | 0 | 0 | 0 |
| AUP1 | Ancient ubiq | 0 | 0.13397514 | 0.00011088 | 0 | 0.00011117 | 0 | 0.00011103 | 0 | 0 | 0 | 0 | 0 | 2 | 0 | 2 | 0 | 2 | 0 | 0 | 0 | 0 | 0 |
| SF3B5 | Splicing facto | 0 | 0.13398221 | 0.00061368 | 0.00061969 | 0 | 0 | 0.00061669 | 0 | 0 | 0 | 0 | 0 | 2 | 2 | 0 | 0 | 2 | 0 | 0 | 0 | 0 | 0 |
| SYAC | Alanine--tRN. | 0 | 0.1400196 | 0.00010904 | 0 | 0 | 8.2641E-05 | 9.5841E-05 | 0 | 0 | 0 | 0 | 0 | 4 | 0 | 0 | 3 | 3.5 | 0 | 0 | 0 | 0 | 0 |
| OGA | Protein O-Glc | 0 | 0.14616659 | 0 | 0 | 8.6655E-05 | 5.8222E-05 | 7.2439E-05 | 0 | 0 | 0 | 0 | 0 | 0 | 0 | 3 | 2 | 2.5 | 0 | 0 | 0 | 0 | 0 |
| RPB1 | DNA-directed | 0 | 0.13398339 | 0.00002679 | 0 | 0 | 2.7072E-05 | 2.6931E-05 | 0 | 0 | 0 | 0 | 0 | 2 | 0 | 0 | 2 | 2 | 0 | 0 | 0 | 0 | 0 |
| PEX1 | Peroxisome b | 0 | 0.14647056 | 6.1703E-05 | 0 | 4.1245E-05 | 0 | 5.1474E-05 | 0 | 0 | 0 | 0 | 0 | 3 | 0 | 2 | 0 | 2.5 | 0 | 0 | 0 | 0 | 0 |
| REQU | Zinc finger pr | 0 | 0.14059382 | 0.00020247 | 0 | 0.00027068 | 0 | 0.00023658 | 0 | 0 | 0 | 0 | 0 | 3 | 0 | 4 | 0 | 3.5 | 0 | 0 | 0 | 0 | 0 |
| RHG05 | Rho GTPase-a | 0 | 0.13398216 | 3.5138E-05 | 3.5481E-05 | 0 | 0 | 3.531E-05 | 0 | 0 | 0 | 0 | 0 | 2 | 2 | 0 | 0 | 2 | 0 | 0 | 0 | 0 | 0 |
| ANFY1 | Rabankyrin-5 | 0 | 0.14616725 | 0 | 0 | 6.7901E-05 | 4.5621E-05 | 5.6761E-05 | 0 | 0 | 0 | 0 | 0 | 0 | 0 | 3 | 2 | 2.5 | 0 | 0 | 0 | 0 | 0 |
| PEX14 | Peroxisomal | 0 | 0.14679094 | 0.00013999 | 0 | 0.00021055 | 0 | 0.00017527 | 0 | 0 | 0 | 0 | 0 | 2 | 0 | 3 | 0 | 2.5 | 0 | 0 | 0 | 0 | 0 |
| RANB3 | Ran-binding p | 0 | 0.14001958 | 0.00018616 | 0 | 0 | 0.00014109 | 0.00016363 | 0 | 0 | 0 | 0 | 0 | 4 | 0 | 0 | 3 | 3.5 | 0 | 0 | 0 | 0 | 0 |
| PAK2 | Serine/threo | 0 | 0.20695674 | 0 | 0.00030511 | 0 | 0.00010178 | 0.00020345 | 0 | 0 | 0 | 0 | 0 | 0 | 6 | 0 | 2 | 4 | 0 | 0 | 0 | 0 | 0 |
| RABL3 | Rab-like prot | 0 | 0.13397464 | 0 | 0.00022582 | 0 | 0.00022598 | 0.0002259 | 0 | 0 | 0 | 0 | 0 | 0 | 2 | 0 | 2 | 2 | 0 | 0 | 0 | 0 | 0 |
| SHCBP | SHC SH2 dom | 0 | 0.14658783 | 0 | 0.00011896 | 0 | 7.9362E-05 | 9.9161E-05 | 0 | 0 | 0 | 0 | 0 | 0 | 3 | 0 | 2 | 2.5 | 0 | 0 | 0 | 0 | 0 |
| DHX16 | Pre-mRNA-sp | 0 | 0.14705645 | 0 | 7.6791E-05 | 5.0833E-05 | 0 | 6.3812E-05 | 0 | 0 | 0 | 0 | 0 | 0 | 3 | 4 | 0 | 3.5 | 0 | 0 | 0 | 0 | 0 |
| KDM5A | Lysine-specif | 0 | 0.13398335 | 3.1229E-05 | 0 | 0 | 3.1557E-05 | 3.1393E-05 | 0 | 0 | 0 | 0 | 0 | 2 | 0 | 0 | 2 | 2 | 0 | 0 | 0 | 0 | 0 |
| AKA12 | A-kinase ancl | 0 | 0.14678819 | 2.9617E-05 | 0 | 4.4543E-05 | 0 | 0.00003708 | 0 | 0 | 0 | 0 | 0 | 2 | 0 | 3 | 0 | 2.5 | 0 | 0 | 0 | 0 | 0 |
| PNO1 | RNA-binding | 0 | 0.1339786 | 0 | 0.00021148 | 0.00020999 | 0 | 0.00021074 | 0 | 0 | 0 | 0 | 0 | 0 | 2 | 2 | 0 | 2 | 0 | 0 | 0 | 0 | 0 |
| PAR3L | Partitioning c | 0 | 0.1339822 | 6.5697E-05 | 0.00006634 | 0 | 0 | 6.6019E-05 | 0 | 0 | 0 | 0 | 0 | 3 | 3 | 0 | 0 | 3 | 0 | 0 | 0 | 0 | 0 |
| NF1 | Neurofibrom | 0 | 0.14678872 | 0.00001859 | 0 | 2.7959E-05 | 0 | 2.3275E-05 | 0 | 0 | 0 | 0 | 0 | 2 | 0 | 3 | 0 | 2.5 | 0 | 0 | 0 | 0 | 0 |
| CALX | Calnexin OS= | 0 | 0.14667261 | 0 | 9.0022E-05 | 0 | 0.00013513 | 0.00011258 | 0 | 0 | 0 | 0 | 0 | 0 | 2 | 0 | 3 | 2.5 | 0 | 0 | 0 | 0 | 0 |
| TCP4 | Activated RN | 0 | 0.13397464 | 0 | 0.00062944 | 0 | 0.0006299 | 0.00062967 | 0 | 0 | 0 | 0 | 0 | 0 | 3 | 0 | 3 | 3 | 0 | 0 | 0 | 0 | 0 |
| BTAF1 | TATA-binding | 0 | 0.14647146 | 4.2815E-05 | 0 | 2.8619E-05 | 0 | 3.5717E-05 | 0 | 0 | 0 | 0 | 0 | 3 | 0 | 2 | 0 | 2.5 | 0 | 0 | 0 | 0 | 0 |
| UFD1 | Ubiquitin rec | 0 | 0.13398218 | 0.00025787 | 0.00026039 | 0 | 0 | 0.00025913 | 0 | 0 | 0 | 0 | 0 | 3 | 3 | 0 | 0 | 3 | 0 | 0 | 0 | 0 | 0 |
| KI18B | Kinesin-like p | 0 | 0.13397516 | 9.1626E-05 | 0 | 0.00009187 | 0 | 9.1748E-05 | 0 | 0 | 0 | 0 | 0 | 3 | 0 | 3 | 0 | 3 | 0 | 0 | 0 | 0 | 0 |
| RIPK2 | Receptor-inte | 0 | 0.14705793 | 0 | 0.00014804 | 9.7995E-05 | 0 | 0.00012302 | 0 | 0 | 0 | 0 | 0 | 0 | 3 | 2 | 0 | 2.5 | 0 | 0 | 0 | 0 | 0 |
| BOP1 | Ribosome bic | 0 | 0.13397464 | 0 | 7.1438E-05 | 0 | 0.00007149 | 7.1464E-05 | 0 | 0 | 0 | 0 | 0 | 0 | 2 | 0 | 2 | 2 | 0 | 0 | 0 | 0 | 0 |
| CIZ1 | Cip1-interact | 0 | 0.18803524 | 0.00014693 | 0 | 0 | 5.9389E-05 | 0.00010316 | 0 | 0 | 0 | 0 | 0 | 5 | 0 | 0 | 2 | 3.5 | 0 | 0 | 0 | 0 | 0 |
| SIAS | Sialic acid syr | 0 | 0.13397864 | 0 | 0.00014845 | 0.0001474 | 0 | 0.00014793 | 0 | 0 | 0 | 0 | 0 | 0 | 2 | 2 | 0 | 2 | 0 | 0 | 0 | 0 | 0 |
| PFD2 | Prefoldin sub | 0 | 0.14658635 | 0 | 0.00051909 | 0 | 0.00034631 | 0.0004327 | 0 | 0 | 0 | 0 | 0 | 0 | 3 | 0 | 2 | 2.5 | 0 | 0 | 0 | 0 | 0 |
| MEN1 | Menin OS=Hc | 0 | 0.14646925 | 0.00012872 | 0 | 8.6044E-05 | 0 | 0.00010738 | 0 | 0 | 0 | 0 | 0 | 3 | 0 | 2 | 0 | 2.5 | 0 | 0 | 0 | 0 | 0 |
| DPOA2 | DNA polymer | 0 | 0.13397463 | 0 | 0.00013368 | 0 | 0.00013377 | 0.00013373 | 0 | 0 | 0 | 0 | 0 | 0 | 3 | 0 | 3 | 3 | 0 | 0 | 0 | 0 | 0 |
| DDX51 | ATP-depende | 0 | 0.14658696 | 0 | 0.00012003 | 0 | 8.0077E-05 | 0.00010005 | 0 | 0 | 0 | 0 | 0 | 0 | 3 | 0 | 2 | 2.5 | 0 | 0 | 0 | 0 | 0 |
| CARL2 | Capping prot | 0 | 0.1339822 | 5.5167E-05 | 5.5707E-05 | 0 | 0 | 5.5437E-05 | 0 | 0 | 0 | 0 | 0 | 3 | 3 | 0 | 0 | 3 | 0 | 0 | 0 | 0 | 0 |
| RADI | Radixin OS=H | 0 | 0.13397516 | 9.0526E-05 | 0 | 9.0767E-05 | 0 | 9.0647E-05 | 0 | 0 | 0 | 0 | 0 | 3 | 0 | 3 | 0 | 3 | 0 | 0 | 0 | 0 | 0 |
| SP130 | Histone deac | 0 | 0.16846207 | 0.00005036 | 0 | 0.00010099 | 0 | 7.5675E-05 | 0 | 0 | 0 | 0 | 0 | 2 | 0 | 4 | 0 | 3 | 0 | 0 | 0 | 0 | 0 |
| REV3L | DNA polymer | 0 | 0.14678843 | 1.6862E-05 | 0 | 0.00002536 | 0 | 2.1111E-05 | 0 | 0 | 0 | 0 | 0 | 2 | 0 | 3 | 0 | 2.5 | 0 | 0 | 0 | 0 | 0 |
| TSH3 | Teashirt hom | 0 | 0.14647076 | 7.3233E-05 | 0 | 4.8952E-05 | 0 | 6.1093E-05 | 0 | 0 | 0 | 0 | 0 | 3 | 0 | 2 | 0 | 2.5 | 0 | 0 | 0 | 0 | 0 |
| ELOV1 | Elongation of | 0 | 0.13398219 | 0.00018916 | 0.00019101 | 0 | 0 | 0.00019009 | 0 | 0 | 0 | 0 | 0 | 2 | 2 | 0 | 0 | 2 | 0 | 0 | 0 | 0 | 0 |
| TRA2A | Transformer- | 0 | 0.14647129 | 0.00028073 | 0 | 0.00018765 | 0 | 0.00023419 | 0 | 0 | 0 | 0 | 0 | 4 | 0 | 3 | 0 | 3.5 | 0 | 0 | 0 | 0 | 0 |
| ICAL | Calpastatin C | 0 | 0.14036027 | 0.00014909 | 0 | 0.00011211 | 0 | 0.0001306 | 0 | 0 | 0 | 0 | 0 | 4 | 0 | 3 | 0 | 3.5 | 0 | 0 | 0 | 0 | 0 |
| UCHL5 | Ubiquitin car | 0 | 0.13398211 | 0.00016042 | 0.00016198 | 0 | 0 | 0.0001612 | 0 | 0 | 0 | 0 | 0 | 2 | 2 | 0 | 0 | 2 | 0 | 0 | 0 | 0 | 0 |
| ARBK1 | Beta-adrener | 0 | 0.14705506 | 0 | 0.00011602 | 7.6803E-05 | 0 | 9.6412E-05 | 0 | 0 | 0 | 0 | 0 | 0 | 3 | 2 | 0 | 2.5 | 0 | 0 | 0 | 0 | 0 |
| RPA49 | DNA-directed | 0 | 0.13397464 | 0 | 0.0001108 | 0 | 0.00011088 | 0.00011084 | 0 | 0 | 0 | 0 | 0 | 0 | 2 | 0 | 2 | 2 | 0 | 0 | 0 | 0 | 0 |
| PWP2 | Periodic tryp | 0 | 0.13397516 | 5.7429E-05 | 0 | 5.7581E-05 | 0 | 5.7505E-05 | 0 | 0 | 0 | 0 | 0 | 2 | 0 | 2 | 0 | 2 | 0 | 0 | 0 | 0 | 0 |
| RBM22 | Pre-mRNA-sp | 0 | 0.1400206 | 0.00025132 | 0 | 0 | 0.00019047 | 0.0002209 | 0 | 0 | 0 | 0 | 0 | 4 | 0 | 0 | 3 | 3.5 | 0 | 0 | 0 | 0 | 0 |
| TSC1 | Hamartin OS: | 0 | 0.13397945 | 0 | 0 | 4.5462E-05 | 4.5817E-05 | 4.564E-05 | 0 | 0 | 0 | 0 | 0 | 0 | 0 | 2 | 2 | 2 | 0 | 0 | 0 | 0 | 0 |
| BMP2K | BMP-2-induc | 0 | 0.13397948 | 0 | 0 | 4.5579E-05 | 4.5936E-05 | 4.5758E-05 | 0 | 0 | 0 | 0 | 0 | 0 | 0 | 2 | 2 | 2 | 0 | 0 | 0 | 0 | 0 |
| FWCH2 | FLYWCH fam | 0 | 0.13397948 | 0 | 0 | 0.00056697 | 0.00057141 | 0.00056919 | 0 | 0 | 0 | 0 | 0 | 0 | 0 | 3 | 3 | 3 | 0 | 0 | 0 | 0 | 0 |
| PRR12 | Proline-rich p | 0 | 0.13819602 | 0 | 0 | 8.7107E-05 | 0.00010974 | 9.8424E-05 | 0 | 0 | 0 | 0 | 0 | 0 | 0 | 4 | 5 | 4.5 | 0 | 0 | 0 | 0 | 0 |
| ASPC1 | Tether contai | 0 | 0.14016403 | 0 | 0.00014456 | 0.00019138 | 0 | 0.00016797 | 0 | 0 | 0 | 0 | 0 | 0 | 3 | 4 | 0 | 3.5 | 0 | 0 | 0 | 0 | 0 |
| NHLC2 | NHL repeat-c | 0 | 0.13397517 | 7.2695E-05 | 0 | 7.2889E-05 | 0 | 7.2792E-05 | 0 | 0 | 0 | 0 | 0 | 2 | 0 | 2 | 0 | 2 | 0 | 0 | 0 | 0 | 0 |
| SAE1 | SUMO-activa | 0 | 0.13397864 | 0 | 0.00015403 | 0.00015294 | 0 | 0.00015349 | 0 | 0 | 0 | 0 | 0 | 0 | 2 | 2 | 0 | 2 | 0 | 0 | 0 | 0 | 0 |
| HGS | Hepatocyte g | 0 | 0.14658567 | 0 | 0.00010288 | 0 | 6.8637E-05 | 8.5759E-05 | 0 | 0 | 0 | 0 | 0 | 0 | 3 | 0 | 2 | 2.5 | 0 | 0 | 0 | 0 | 0 |
| ZGRF1 | Protein ZGRF | 0 | 0.13398217 | 2.5084E-05 | 2.5329E-05 | 0 | 0 | 2.5207E-05 | 0 | 0 | 0 | 0 | 0 | 2 | 2 | 0 | 0 | 2 | 0 | 0 | 0 | 0 | 0 |
| E2F6 | Transcription | 0 | 0.14658646 | 0 | 0.00028448 | 0 | 0.00018979 | 0.00023714 | 0 | 0 | 0 | 0 | 0 | 0 | 3 | 0 | 2 | 2.5 | 0 | 0 | 0 | 0 | 0 |
| NECP2 | Adaptin ear-b | 0 | 0.13397518 | 0.00020067 | 0 | 0.00020121 | 0 | 0.00020094 | 0 | 0 | 0 | 0 | 0 | 2 | 0 | 2 | 0 | 2 | 0 | 0 | 0 | 0 | 0 |
| KAT7 | Histone acety | 0 | 0.1339822 | 0.00017276 | 0.00017445 | 0 | 0 | 0.00017361 | 0 | 0 | 0 | 0 | 0 | 4 | 4 | 0 | 0 | 4 | 0 | 0 | 0 | 0 | 0 |
| IWS1 | Protein IWS1 | 0 | 0.13397861 | 0 | 6.5071E-05 | 6.4612E-05 | 0 | 6.4842E-05 | 0 | 0 | 0 | 0 | 0 | 0 | 2 | 2 | 0 | 2 | 0 | 0 | 0 | 0 | 0 |
| TRM7 | Putative tRN/ | 0 | 0.13397516 | 0.00024062 | 0 | 0.00024126 | 0 | 0.00024094 | 0 | 0 | 0 | 0 | 0 | 3 | 0 | 3 | 0 | 3 | 0 | 0 | 0 | 0 | 0 |
| CUL1 | Cullin-1 OS=H | 0 | 0.16828865 | 0 | 6.8677E-05 | 0 | 0.00013745 | 0.00010306 | 0 | 0 | 0 | 0 | 0 | 0 | 2 | 0 | 4 | 3 | 0 | 0 | 0 | 0 | 0 |
| MPC1 | Mitochondria | 0 | 0.13398221 | 0.00048419 | 0.00048893 | 0 | 0 | 0.00048656 | 0 | 0 | 0 | 0 | 0 | 2 | 2 | 0 | 0 | 2 | 0 | 0 | 0 | 0 | 0 |
| BAIP2 | Brain-specific | 0 | 0.14667197 | 0 | 9.6545E-05 | 0 | 0.00014492 | 0.00012073 | 0 | 0 | 0 | 0 | 0 | 0 | 2 | 0 | 3 | 2.5 | 0 | 0 | 0 | 0 | 0 |
| NCOA7 | Nuclear recep | 0 | 0.13397464 | 0 | 5.6574E-05 | 0 | 5.6615E-05 | 5.6595E-05 | 0 | 0 | 0 | 0 | 0 | 0 | 2 | 0 | 2 | 2 | 0 | 0 | 0 | 0 | 0 |
| NSD2 | Histone-lysin | 0 | 0.1460538 | 5.7997E-05 | 3.9042E-05 | 0 | 0 | 4.852E-05 | 0 | 0 | 0 | 0 | 0 | 3 | 2 | 0 | 0 | 2.5 | 0 | 0 | 0 | 0 | 0 |
| AT7L3 | Ataxin-7-like | 0 | 0.14679124 | 0.00015209 | 0 | 0.00022875 | 0 | 0.00019042 | 0 | 0 | 0 | 0 | 0 | 2 | 0 | 3 | 0 | 2.5 | 0 | 0 | 0 | 0 | 0 |
| UBP11 | Ubiquitin car | 0 | 0.14616724 | 0 | 0 | 8.2426E-05 | 0.00005538 | 6.8903E-05 | 0 | 0 | 0 | 0 | 0 | 0 | 0 | 3 | 2 | 2.5 | 0 | 0 | 0 | 0 | 0 |
| RING1 | E3 ubiquitin-p | 0 | 0.13397518 | 0.00012999 | 0 | 0.00013034 | 0 | 0.00013017 | 0 | 0 | 0 | 0 | 0 | 5 | 0 | 3 | 0 | 4 | 0 | 0 | 0 | 0 | 0 |
| PRDM2 | PR domain zi | 0 | 0.13801117 | 0.00006144 | 0 | 7.7004E-05 | 0 | 6.9222E-05 | 0 | 0 | 0 | 0 | 0 | 4 | 0 | 5 | 0 | 4.5 | 0 | 0 | 0 | 0 | 0 |
| PAXI1 | PAX-interacti | 0 | 0.14705619 | 0 | 0.00007478 | 4.9502E-05 | 0 | 6.2141E-05 | 0 | 0 | 0 | 0 | 0 | 0 | 3 | 2 | 0 | 2.5 | 0 | 0 | 0 | 0 | 0 |
| 4F2 | 4F2 cell-surfa | 0 | 0.13397464 | 0 | 8.4592E-05 | 0 | 8.4653E-05 | 8.4623E-05 | 0 | 0 | 0 | 0 | 0 | 0 | 2 | 0 | 2 | 2 | 0 | 0 | 0 | 0 | 0 |
| PDLI3 | PDZ and LIM | 0 | 0.13397517 | 0.00014499 | 0 | 0.00014538 | 0 | 0.00014519 | 0 | 0 | 0 | 0 | 0 | 2 | 0 | 2 | 0 | 2 | 0 | 0 | 0 | 0 | 0 |
| MIRO2 | Mitochondria | 0 | 0.13398216 | 0.0001281 | 0.00012935 | 0 | 0 | 0.00012873 | 0 | 0 | 0 | 0 | 0 | 3 | 3 | 0 | 0 | 3 | 0 | 0 | 0 | 0 | 0 |
| HSP74 | Heat shock 7 | 0 | 0.13397464 | 0 | 6.3444E-05 | 0 | 0.00006349 | 6.3467E-05 | 0 | 0 | 0 | 0 | 0 | 0 | 2 | 0 | 2 | 2 | 0 | 0 | 0 | 0 | 0 |
| PRI2 | DNA primase | 0 | 0.13397464 | 0 | 0.0001047 | 0 | 0.00010478 | 0.00010474 | 0 | 0 | 0 | 0 | 0 | 0 | 2 | 0 | 2 | 2 | 0 | 0 | 0 | 0 | 0 |
| MSD2 | Myb/SANT-lil | 0 | 0.13398338 | 0.00014162 | 0 | 0 | 0.00014311 | 0.00014237 | 0 | 0 | 0 | 0 | 0 | 3 | 0 | 0 | 3 | 3 | 0 | 0 | 0 | 0 | 0 |
| MBD2 | Methyl-CpG- | 0 | 0.13397866 | 0 | 0.00012967 | 0.00012875 | 0 | 0.00012921 | 0 | 0 | 0 | 0 | 0 | 0 | 2 | 2 | 0 | 2 | 0 | 0 | 0 | 0 | 0 |
| NEP1 | Ribosomal RI | 0 | 0.13397949 | 0 | 0 | 0.00021687 | 0.00021857 | 0.00021772 | 0 | 0 | 0 | 0 | 0 | 0 | 0 | 2 | 2 | 2 | 0 | 0 | 0 | 0 | 0 |
| P4HA2 | Prolyl 4-hydr | 0 | 0.14621052 | 0 | 9.9613E-05 | 0.00014837 | 0 | 0.00012399 | 0 | 0 | 0 | 0 | 0 | 0 | 2 | 3 | 0 | 2.5 | 0 | 0 | 0 | 0 | 0 |
| TM131 | Transmembr. | 0 | 0.14705521 | 0 | 4.2453E-05 | 2.8103E-05 | 0 | 3.5278E-05 | 0 | 0 | 0 | 0 | 0 | 0 | 3 | 2 | 0 | 2.5 | 0 | 0 | 0 | 0 | 0 |
| SNP23 | Synaptosoma | 0 | 0.14647139 | 0.00037519 | 0 | 0.00025079 | 0 | 0.00031299 | 0 | 0 | 0 | 0 | 0 | 3 | 0 | 2 | 0 | 2.5 | 0 | 0 | 0 | 0 | 0 |
| SYUA | Alpha-synucl | 0 | 0.13398335 | 0.00037698 | 0 | 0 | 0.00038094 | 0.00037896 | 0 | 0 | 0 | 0 | 0 | 2 | 0 | 0 | 2 | 2 | 0 | 0 | 0 | 0 | 0 |
| RT09 | 28S ribosoma | 0 | 0.14679007 | 0.00013327 | 0 | 0.00020044 | 0 | 0.00016686 | 0 | 0 | 0 | 0 | 0 | 2 | 0 | 3 | 0 | 2.5 | 0 | 0 | 0 | 0 | 0 |
| BAG1 | BAG family m | 0 | 0.13397862 | 0 | 0.00015447 | 0.00015338 | 0 | 0.00015393 | 0 | 0 | 0 | 0 | 0 | 0 | 2 | 2 | 0 | 2 | 0 | 0 | 0 | 0 | 0 |
| ARHGG | Rho guanine | 0 | 0.13397464 | 0 | 7.5166E-05 | 0 | 0.00007522 | 7.5193E-05 | 0 | 0 | 0 | 0 | 0 | 0 | 2 | 0 | 2 | 2 | 0 | 0 | 0 | 0 | 0 |
| PTN2 | Tyrosine-prot | 0 | 0.14667 | 0 | 0.00012842 | 0 | 0.00019276 | 0.00016059 | 0 | 0 | 0 | 0 | 0 | 0 | 2 | 0 | 3 | 2.5 | 0 | 0 | 0 | 0 | 0 |
| PIAS3 | E3 SUMO-pro | 0 | 0.13398217 | 0.00008404 | 8.4861E-05 | 0 | 0 | 8.4451E-05 | 0 | 0 | 0 | 0 | 0 | 5 | 4 | 0 | 0 | 4.5 | 0 | 0 | 0 | 0 | 0 |
| MKLN1 | Muskelin OS= | 0 | 0.1464723 | 0.00010771 | 0 | 7.1996E-05 | 0 | 8.9853E-05 | 0 | 0 | 0 | 0 | 0 | 3 | 0 | 2 | 0 | 2.5 | 0 | 0 | 0 | 0 | 0 |
| CWC15 | Spliceosome- | 0 | 0.13397464 | 0 | 0.00023272 | 0 | 0.00023289 | 0.00023281 | 0 | 0 | 0 | 0 | 0 | 0 | 2 | 0 | 2 | 2 | 0 | 0 | 0 | 0 | 0 |
| ARIP4 | Helicase ARIF | 0 | 0.13397517 | 3.5976E-05 | 0 | 3.6072E-05 | 0 | 3.6024E-05 | 0 | 0 | 0 | 0 | 0 | 2 | 0 | 2 | 0 | 2 | 0 | 0 | 0 | 0 | 0 |
| SYCC | Cysteine--tRI | 0 | 0.13397464 | 0 | 7.1247E-05 | 0 | 7.1298E-05 | 7.1273E-05 | 0 | 0 | 0 | 0 | 0 | 0 | 2 | 0 | 2 | 2 | 0 | 0 | 0 | 0 | 0 |
| UBXN1 | UBX domain- | 0 | 0.13397464 | 0 | 0.00017944 | 0 | 0.00017957 | 0.00017951 | 0 | 0 | 0 | 0 | 0 | 0 | 2 | 0 | 2 | 2 | 0 | 0 | 0 | 0 | 0 |
| CSN7A | COP9 signalo | 0 | 0.1339833 | 0.00019192 | 0 | 0 | 0.00019393 | 0.00019293 | 0 | 0 | 0 | 0 | 0 | 2 | 0 | 0 | 2 | 2 | 0 | 0 | 0 | 0 | 0 |
| KAP3 | cAMP-depen | 0 | 0.1467879 | 0.00012626 | 0 | 0.00018989 | 0 | 0.00015808 | 0 | 0 | 0 | 0 | 0 | 6 | 0 | 8 | 0 | 7 | 0 | 0 | 0 | 0 | 0 |
| CAN2 | Calpain-2 cat | 0 | 0.1339822 | 7.5395E-05 | 7.6133E-05 | 0 | 0 | 7.5764E-05 | 0 | 0 | 0 | 0 | 0 | 2 | 2 | 0 | 0 | 2 | 0 | 0 | 0 | 0 | 0 |
| CCNL2 | Cyclin-L2 OS= | 0 | 0.13397464 | 0 | 0.00015373 | 0 | 0.00015384 | 0.00015379 | 0 | 0 | 0 | 0 | 0 | 0 | 3 | 0 | 3 | 3 | 0 | 0 | 0 | 0 | 0 |
| NACC1 | Nucleus accu | 0 | 0.13397464 | 0 | 0.00015169 | 0 | 0.0001518 | 0.00015175 | 0 | 0 | 0 | 0 | 0 | 0 | 3 | 0 | 3 | 3 | 0 | 0 | 0 | 0 | 0 |
| SAMD1 | Atherin OS=H | 0 | 0.14721985 | 9.8098E-05 | 0.00014859 | 0 | 0 | 0.00012334 | 0 | 0 | 0 | 0 | 0 | 2 | 3 | 0 | 0 | 2.5 | 0 | 0 | 0 | 0 | 0 |
| ATX3 | Ataxin-3 OS= | 0 | 0.13398329 | 0.0001462 | 0 | 0 | 0.00014773 | 0.00014697 | 0 | 0 | 0 | 0 | 0 | 2 | 0 | 0 | 2 | 2 | 0 | 0 | 0 | 0 | 0 |
| SKA3 | Spindle and k | 0 | 0.04517299 | 0.00057645 | 0.00038806 | 0.00025688 | 0 | 0.00040713 | 0 | 0 | 0 | 0 | 0 | 9 | 6 | 4 | 0 | 6.33333333 | 0 | 0 | 0 | 0 | 0 |
| SMD2 | Small nuclear | 0 | 0.02619463 | 0.00089452 | 0.00090327 | 0.0011211 | 0 | 0.00097296 | 0 | 0 | 0 | 0 | 0 | 4 | 4 | 5 | 0 | 4.33333333 | 0 | 0 | 0 | 0 | 0 |
| EF1B | Elongation fa | 0 | 0.03182677 | 0.00023456 | 0 | 0.00035278 | 0.00023703 | 0.00027479 | 0 | 0 | 0 | 0 | 0 | 2 | 0 | 3 | 2 | 2.33333333 | 0 | 0 | 0 | 0 | 0 |
| ANM5 | Protein argin | 0 | 0.10442789 | 0.00012428 | 0 | 0.00033229 | 8.3722E-05 | 0.0001801 | 0 | 0 | 0 | 0 | 0 | 3 | 0 | 8 | 2 | 4.33333333 | 0 | 0 | 0 | 0 | 0 |
| BTF3 | Transcription | 0 | 0.0250909 | 0 | 0.00090546 | 0.00077064 | 0.00077667 | 0.00081759 | 0 | 0 | 0 | 0 | 0 | 0 | 7 | 6 | 6 | 6.33333333 | 0 | 0 | 0 | 0 | 0 |
| SHIP2 | Phosphatidyl | 0 | 0.02797134 | 6.2929E-05 | 6.3545E-05 | 0 | 8.4787E-05 | 7.042E-05 | 0 | 0 | 0 | 0 | 0 | 3 | 3 | 0 | 4 | 3.33333333 | 0 | 0 | 0 | 0 | 0 |
| CDV3 | Protein CDV3 | 0 | 0.03184217 | 0.00020456 | 0.00020656 | 0.00030766 | 0 | 0.00023959 | 0 | 0 | 0 | 0 | 0 | 2 | 2 | 3 | 0 | 2.33333333 | 0 | 0 | 0 | 0 | 0 |
| CC124 | Coiled-coil dc | 0 | 0.02790175 | 0 | 0.00047796 | 0.00035595 | 0.00035873 | 0.00039755 | 0 | 0 | 0 | 0 | 0 | 0 | 4 | 3 | 3 | 3.33333333 | 0 | 0 | 0 | 0 | 0 |
| TBL3 | Transducin b | 0 | 0.05052652 | 9.7977E-05 | 0.00019787 | 9.8238E-05 | 0 | 0.00013136 | 0 | 0 | 0 | 0 | 0 | 3 | 6 | 3 | 0 | 4 | 0 | 0 | 0 | 0 | 0 |
| H2AV | Histone H2A. | 0 | 0.02705052 | 0.00082464 | 0.00062453 | 0 | 0.0008333 | 0.00076082 | 0 | 0 | 0 | 0 | 0 | 4 | 3 | 0 | 4 | 3.66666667 | 0 | 0 | 0 | 0 | 0 |
| H2AZ | Histone H2A. | 0 | 0.02705052 | 0.00082464 | 0.00062453 | 0 | 0.0008333 | 0.00076082 | 0 | 0 | 0 | 0 | 0 | 4 | 3 | 0 | 4 | 3.66666667 | 0 | 0 | 0 | 0 | 0 |
| ARFP2 | Arfaptin-2 OS | 0 | 0.0496318 | 0 | 0.00039071 | 0.00038796 | 0.0001564 | 0.00031169 | 0 | 0 | 0 | 0 | 0 | 0 | 5 | 5 | 2 | 4 | 0 | 0 | 0 | 0 | 0 |
| AAAS | Aladin OS=Hc | 0 | 0.03009596 | 0.00014499 | 0 | 9.6918E-05 | 0.00014651 | 0.00012947 | 0 | 0 | 0 | 0 | 0 | 3 | 0 | 2 | 3 | 2.66666667 | 0 | 0 | 0 | 0 | 0 |
| MED14 | Mediator of I | 0 | 0.03925212 | 3.6298E-05 | 7.3305E-05 | 5.4592E-05 | 0 | 5.4732E-05 | 0 | 0 | 0 | 0 | 0 | 2 | 4 | 3 | 0 | 3 | 0 | 0 | 0 | 0 | 0 |
| PAF1 | RNA polymer | 0 | 0.03326213 | 0 | 0.00020073 | 0.0003488 | 0.00030131 | 0.00028361 | 0 | 0 | 0 | 0 | 0 | 0 | 4 | 7 | 6 | 5.66666667 | 0 | 0 | 0 | 0 | 0 |
| URFB1 | UHRF1-bindi | 0 | 0.03971643 | 7.3301E-05 | 3.7009E-05 | 7.3496E-05 | 0 | 6.1269E-05 | 0 | 0 | 0 | 0 | 0 | 4 | 2 | 4 | 0 | 3.33333333 | 0 | 0 | 0 | 0 | 0 |
| BICRA | BRD4-interac | 0 | 0.02764162 | 6.7663E-05 | 5.1243E-05 | 5.0882E-05 | 0 | 5.6596E-05 | 0 | 0 | 0 | 0 | 0 | 4 | 3 | 3 | 0 | 3.33333333 | 0 | 0 | 0 | 0 | 0 |
| ARID2 | AT-rich intera | 0 | 0.03197558 | 7.1903E-05 | 5.8085E-05 | 4.3257E-05 | 0 | 5.7748E-05 | 0 | 0 | 0 | 0 | 0 | 5 | 4 | 3 | 0 | 4 | 0 | 0 | 0 | 0 | 0 |
| PLST | Plastin-3 OS= | 0 | 0.05961232 | 8.3773E-05 | 0.00025378 | 0 | 0.00016931 | 0.00016895 | 0 | 0 | 0 | 0 | 0 | 2 | 6 | 0 | 4 | 4 | 0 | 0 | 0 | 0 | 0 |
| ITSN2 | Intersectin-2 | 0 | 0.02978773 | 0.00004665 | 3.1404E-05 | 4.6774E-05 | 0 | 4.1609E-05 | 0 | 0 | 0 | 0 | 0 | 4 | 3 | 4 | 0 | 3.66666667 | 0 | 0 | 0 | 0 | 0 |
| EWS | RNA-binding | 0 | 0.0316598 | 0.00012068 | 0 | 8.0667E-05 | 8.1298E-05 | 9.4215E-05 | 0 | 0 | 0 | 0 | 0 | 3 | 0 | 2 | 2 | 2.33333333 | 0 | 0 | 0 | 0 | 0 |
| WDR41 | WD repeat-co | 0 | 0.03232576 | 0.00011498 | 0 | 0.00011529 | 0.00017429 | 0.00013485 | 0 | 0 | 0 | 0 | 0 | 2 | 0 | 2 | 3 | 2.33333333 | 0 | 0 | 0 | 0 | 0 |
| SNP29 | Synaptosoma | 0 | 0.03238962 | 0.00030684 | 0.0005164 | 0 | 0.00041342 | 0.00041222 | 0 | 0 | 0 | 0 | 0 | 3 | 5 | 0 | 4 | 4 | 0 | 0 | 0 | 0 | 0 |

| | | | | | | | | | | | | | | | | | | | | | | | |
|---|---|---|---|---|---|---|---|---|---|---|---|---|---|---|---|---|---|---|---|---|---|---|---|
| HELLS | Lymphoid-sp | 0 | 0.02539418 | 0.00015745 | 0.00019079 | 0.00018944 | 0 | 0.00017923 | 0 | 0 | 0 | 0 | 0 | 5 | 6 | 6 | 0 | 5.66666667 | 0 | 0 | 0 | 0 | 0 |
| TRRAP | Transformati | 0 | 0.0272805 | 2.0514E-05 | 0 | 2.7425E-05 | 0.00002764 | 2.5193E-05 | 0 | 0 | 0 | 0 | 0 | 3 | 0 | 4 | 4 | 3.66666667 | 0 | 0 | 0 | 0 | 0 |
| UBF1 | Nucleolar tra | 0 | 0.03906611 | 0 | 0.00013951 | 6.9264E-05 | 0.00010471 | 0.00010449 | 0 | 0 | 0 | 0 | 0 | 0 | 4 | 2 | 3 | 3 | 0 | 0 | 0 | 0 | 0 |
| EM55 | 55 kDa erythi | 0 | 0.03827644 | 0 | 0.00017154 | 0.00028389 | 0.00034333 | 0.00026625 | 0 | 0 | 0 | 0 | 0 | 0 | 3 | 5 | 6 | 4.66666667 | 0 | 0 | 0 | 0 | 0 |
| EED | Polycomb pr | 0 | 0.03211539 | 0 | 0.00012085 | 0.00011999 | 0.0001814 | 0.00014075 | 0 | 0 | 0 | 0 | 0 | 0 | 2 | 2 | 3 | 2.33333333 | 0 | 0 | 0 | 0 | 0 |
| UIMC1 | BRCA1-A com | 0 | 0.07241724 | 7.3403E-05 | 7.4121E-05 | 0 | 0.00018544 | 0.00011099 | 0 | 0 | 0 | 0 | 0 | 2 | 2 | 0 | 5 | 3 | 0 | 0 | 0 | 0 | 0 |
| CTR9 | RNA polymer | 0 | 0.05289928 | 0.00006749 | 4.5433E-05 | 0.00011278 | 0 | 7.5234E-05 | 0 | 0 | 0 | 0 | 0 | 3 | 2 | 5 | 0 | 3.33333333 | 0 | 0 | 0 | 0 | 0 |
| EFL1 | Elongation fa | 0 | 0.03925298 | 4.7122E-05 | 9.5166E-05 | 7.0871E-05 | 0 | 7.1053E-05 | 0 | 0 | 0 | 0 | 0 | 2 | 4 | 3 | 0 | 3 | 0 | 0 | 0 | 0 | 0 |
| NH2L1 | NHP2-like pr | 0 | 0.04961382 | 0.00041232 | 0 | 0.00082683 | 0.00041665 | 0.00055193 | 0 | 0 | 0 | 0 | 0 | 2 | 0 | 4 | 2 | 2.66666667 | 0 | 0 | 0 | 0 | 0 |
| GRIN1 | G protein-reg | 0 | 0.03229336 | 0.00010472 | 0.00015861 | 0.00010499 | 0 | 0.00012277 | 0 | 0 | 0 | 0 | 0 | 4 | 6 | 4 | 0 | 4.66666667 | 0 | 0 | 0 | 0 | 0 |
| NADAP | Kanadaptin C | 0 | 0.04031371 | 6.6303E-05 | 0.0001339 | 0.00013296 | 0 | 0.00011105 | 0 | 0 | 0 | 0 | 0 | 2 | 4 | 4 | 0 | 3.33333333 | 0 | 0 | 0 | 0 | 0 |
| URB2 | Unhealthy rib | 0 | 0.05017879 | 0 | 3.4969E-05 | 3.4723E-05 | 6.9988E-05 | 0.00004656 | 0 | 0 | 0 | 0 | 0 | 0 | 2 | 2 | 4 | 2.66666667 | 0 | 0 | 0 | 0 | 0 |
| ANR11 | Ankyrin repe | 0 | 0.08688071 | 6.9365E-05 | 0 | 2.9807E-05 | 2.0027E-05 | 3.9733E-05 | 0 | 0 | 0 | 0 | 0 | 7 | 0 | 3 | 2 | 4 | 0 | 0 | 0 | 0 | 0 |
| HAKAI | E3 ubiquitin- | 0 | 0.04963203 | 0 | 0.00027135 | 0.00026944 | 0.00010862 | 0.00021647 | 0 | 0 | 0 | 0 | 0 | 0 | 5 | 5 | 2 | 4 | 0 | 0 | 0 | 0 | 0 |
| S4A7 | Sodium bicar | 0 | 0.03983588 | 8.6947E-05 | 8.7797E-05 | 0 | 0.00004393 | 7.2891E-05 | 0 | 0 | 0 | 0 | 0 | 4 | 4 | 0 | 2 | 3.33333333 | 0 | 0 | 0 | 0 | 0 |
| KDM2A | Lysine-specif | 0 | 0.04957543 | 0.00011355 | 0.00011466 | 0 | 4.5896E-05 | 9.1369E-05 | 0 | 0 | 0 | 0 | 0 | 5 | 5 | 0 | 2 | 4 | 0 | 0 | 0 | 0 | 0 |
| KI20B | Kinesin-like p | 0 | 0.04961454 | 4.3497E-05 | 0 | 8.7226E-05 | 4.3954E-05 | 5.8226E-05 | 0 | 0 | 0 | 0 | 0 | 3 | 0 | 6 | 3 | 4 | 0 | 0 | 0 | 0 | 0 |
| CFDP1 | Craniofacial c | 0 | 0.04059659 | 0.00035302 | 0.00062383 | 0.00035396 | 0 | 0.0004436 | 0 | 0 | 0 | 0 | 0 | 4 | 7 | 4 | 0 | 5 | 0 | 0 | 0 | 0 | 0 |
| PAPOA | Poly(A) polyn | 0 | 0.03894417 | 0.00010626 | 7.1534E-05 | 0 | 0.00014317 | 0.00010699 | 0 | 0 | 0 | 0 | 0 | 3 | 2 | 0 | 4 | 3 | 0 | 0 | 0 | 0 | 0 |
| KMT2D | Histone-lysin | 0 | 0.03295181 | 2.3829E-05 | 0 | 2.3893E-05 | 1.4448E-05 | 2.0723E-05 | 0 | 0 | 0 | 0 | 0 | 5 | 0 | 5 | 3 | 4.33333333 | 0 | 0 | 0 | 0 | 0 |
| ANM1 | Protein argin | 0 | 0.03229221 | 0.0001462 | 0.00022144 | 0.00014659 | 0 | 0.00017141 | 0 | 0 | 0 | 0 | 0 | 2 | 3 | 2 | 0 | 2.33333333 | 0 | 0 | 0 | 0 | 0 |
| STXB3 | Syntaxin-binc | 0 | 0.03845373 | 0 | 9.0022E-05 | 0.00017877 | 0.00013513 | 0.00013464 | 0 | 0 | 0 | 0 | 0 | 0 | 2 | 4 | 3 | 3 | 0 | 0 | 0 | 0 | 0 |
| SNX2 | Sorting nexin | 0 | 0.03212764 | 0.00010169 | 0.00015403 | 0 | 0.00010276 | 0.00011949 | 0 | 0 | 0 | 0 | 0 | 2 | 3 | 0 | 2 | 2.33333333 | 0 | 0 | 0 | 0 | 0 |
| UBIP1 | Upstream-bii | 0 | 0.04732576 | 0.00024434 | 9.8691E-05 | 0.00019599 | 0 | 0.00017967 | 0 | 0 | 0 | 0 | 0 | 7 | 5 | 6 | 0 | 6 | 0 | 0 | 0 | 0 | 0 |
| ARPC2 | Actin-related | 0 | 0.03189198 | 0.00035185 | 0 | 0.00044098 | 0.00026666 | 0.00035316 | 0 | 0 | 0 | 0 | 0 | 4 | 0 | 5 | 3 | 4 | 0 | 0 | 0 | 0 | 0 |
| NCOA6 | Nuclear rece | 0 | 0.0240124 | 3.8374E-05 | 3.8749E-05 | 0 | 3.8777E-05 | 3.8633E-05 | 0 | 0 | 0 | 0 | 0 | 3 | 3 | 0 | 3 | 3 | 0 | 0 | 0 | 0 | 0 |
| SENP1 | Sentrin-speci | 0 | 0.03971624 | 0.0001639 | 8.2753E-05 | 0.00016434 | 0 | 0.000137 | 0 | 0 | 0 | 0 | 0 | 4 | 2 | 4 | 0 | 3.33333333 | 0 | 0 | 0 | 0 | 0 |
| CALM1 | Calmodulin-1 | 0 | 0.03229334 | 0.00035421 | 0.00053651 | 0.00035515 | 0 | 0.00041529 | 0 | 0 | 0 | 0 | 0 | 2 | 3 | 2 | 0 | 2.33333333 | 0 | 0 | 0 | 0 | 0 |
| CALM2 | Calmodulin-2 | 0 | 0.03229334 | 0.00035421 | 0.00053651 | 0.00035515 | 0 | 0.00041529 | 0 | 0 | 0 | 0 | 0 | 2 | 3 | 2 | 0 | 2.33333333 | 0 | 0 | 0 | 0 | 0 |
| CALM3 | Calmodulin-3 | 0 | 0.03229334 | 0.00035421 | 0.00053651 | 0.00035515 | 0 | 0.00041529 | 0 | 0 | 0 | 0 | 0 | 2 | 3 | 2 | 0 | 2.33333333 | 0 | 0 | 0 | 0 | 0 |
| CDC23 | Cell division c | 0 | 0.05009176 | 0 | 0.00017854 | 8.8639E-05 | 8.9332E-05 | 0.00011884 | 0 | 0 | 0 | 0 | 0 | 0 | 4 | 2 | 2 | 2.66666667 | 0 | 0 | 0 | 0 | 0 |
| NCOR2 | Nuclear rece | 0 | 0.03656721 | 5.2254E-05 | 3.1659E-05 | 0 | 3.1682E-05 | 3.8532E-05 | 0 | 0 | 0 | 0 | 0 | 7 | 5 | 0 | 4 | 5.33333333 | 0 | 0 | 0 | 0 | 0 |
| SHB | SH2 domain- | 0 | 0.04762304 | 0.00025922 | 0 | 0.00010396 | 0.00020955 | 0.00019091 | 0 | 0 | 0 | 0 | 0 | 5 | 0 | 2 | 4 | 3.66666667 | 0 | 0 | 0 | 0 | 0 |
| SAS10 | Something al | 0 | 0.07098593 | 0.00016527 | 0.00033378 | 0.00011047 | 0 | 0.00020317 | 0 | 0 | 0 | 0 | 0 | 3 | 6 | 2 | 0 | 3.66666667 | 0 | 0 | 0 | 0 | 0 |
| CENPC | Centromere | 0 | 0.04928292 | 0.00011193 | 0 | 5.6116E-05 | 5.6555E-05 | 7.4867E-05 | 0 | 0 | 0 | 0 | 0 | 4 | 0 | 2 | 2 | 2.66666667 | 0 | 0 | 0 | 0 | 0 |
| HECD1 | E3 ubiquitin- | 0 | 0.03700749 | 3.0332E-05 | 3.0628E-05 | 5.0687E-05 | 0 | 3.7216E-05 | 0 | 0 | 0 | 0 | 0 | 3 | 3 | 5 | 0 | 3.66666667 | 0 | 0 | 0 | 0 | 0 |
| TOIP1 | Torsin-1A-int | 0 | 0.03167438 | 0.00013579 | 9.1412E-05 | 9.0767E-05 | 0 | 0.00010599 | 0 | 0 | 0 | 0 | 0 | 3 | 2 | 2 | 0 | 2.33333333 | 0 | 0 | 0 | 0 | 0 |
| ELOA1 | Elongin-A OS | 0 | 0.03837736 | 0.00013227 | 0 | 9.9469E-05 | 6.6831E-05 | 9.9523E-05 | 0 | 0 | 0 | 0 | 0 | 4 | 0 | 3 | 2 | 3 | 0 | 0 | 0 | 0 | 0 |
| ZBT10 | Zinc finger ar | 0 | 0.03439538 | 0.00021208 | 0 | 0.00015189 | 0.00012246 | 0.00016214 | 0 | 0 | 0 | 0 | 0 | 7 | 0 | 5 | 4 | 5.33333333 | 0 | 0 | 0 | 0 | 0 |
| T2FA | General trans | 0 | 0.05947983 | 0 | 0.00030924 | 0.00010235 | 0.00020631 | 0.00020597 | 0 | 0 | 0 | 0 | 0 | 0 | 6 | 2 | 4 | 4 | 0 | 0 | 0 | 0 | 0 |
| CLP1 | Polyribonucle | 0 | 0.03151068 | 0.00018627 | 0.0001254 | 0 | 0.00012549 | 0.00014572 | 0 | 0 | 0 | 0 | 0 | 3 | 2 | 0 | 2 | 2.33333333 | 0 | 0 | 0 | 0 | 0 |
| MIC19 | MICOS comp | 0 | 0.03211494 | 0 | 0.00023477 | 0.00023312 | 0.00035241 | 0.00027343 | 0 | 0 | 0 | 0 | 0 | 0 | 2 | 2 | 3 | 2.33333333 | 0 | 0 | 0 | 0 | 0 |
| RABL6 | Rab-like prot | 0 | 0.03883364 | 7.2396E-05 | 0.00010966 | 0.00014518 | 0 | 0.00010908 | 0 | 0 | 0 | 0 | 0 | 2 | 3 | 4 | 0 | 3 | 0 | 0 | 0 | 0 | 0 |
| PRD16 | PR domain zi | 0 | 0.02705068 | 8.2722E-05 | 6.2648E-05 | 0 | 8.3591E-05 | 7.632E-05 | 0 | 0 | 0 | 0 | 0 | 4 | 3 | 0 | 4 | 3.66666667 | 0 | 0 | 0 | 0 | 0 |
| THOC7 | THO complex | 0 | 0.02978757 | 0.00038806 | 0.00026124 | 0.0003891 | 0 | 0.00034613 | 0 | 0 | 0 | 0 | 0 | 3 | 2 | 3 | 0 | 2.66666667 | 0 | 0 | 0 | 0 | 0 |
| FA50A | Protein FAM5 | 0 | 0.03182779 | 0.00015568 | 0 | 0.00023415 | 0.00015732 | 0.00018238 | 0 | 0 | 0 | 0 | 0 | 2 | 0 | 3 | 2 | 2.33333333 | 0 | 0 | 0 | 0 | 0 |
| DPYL2 | Dihydropyrin | 0 | 0.03030321 | 9.2267E-05 | 0.00013975 | 0 | 0.00013985 | 0.00012396 | 0 | 0 | 0 | 0 | 0 | 2 | 3 | 0 | 3 | 2.66666667 | 0 | 0 | 0 | 0 | 0 |
| ZC3H4 | Zinc finger C( | 0 | 0.0316592 | 6.0756E-05 | 0 | 4.0612E-05 | 0.00004093 | 4.7433E-05 | 0 | 0 | 0 | 0 | 0 | 3 | 0 | 2 | 2 | 2.33333333 | 0 | 0 | 0 | 0 | 0 |
| CBPD | Carboxypepti | 0 | 0.03971666 | 7.6488E-05 | 3.8618E-05 | 7.6692E-05 | 0 | 6.3933E-05 | 0 | 0 | 0 | 0 | 0 | 4 | 2 | 4 | 0 | 3.33333333 | 0 | 0 | 0 | 0 | 0 |
| PPIL4 | Peptidyl-prol | 0 | 0.02753441 | 0.00021454 | 0.00016248 | 0 | 0.0001626 | 0.00017987 | 0 | 0 | 0 | 0 | 0 | 4 | 3 | 0 | 3 | 3.33333333 | 0 | 0 | 0 | 0 | 0 |
| GRSF1 | G-rich seque | 0 | 0.03151194 | 0.00016493 | 0.00011103 | 0 | 0.00011111 | 0.00012902 | 0 | 0 | 0 | 0 | 0 | 3 | 2 | 0 | 2 | 2.33333333 | 0 | 0 | 0 | 0 | 0 |
| TCF20 | Transcription | 0 | 0.04032931 | 2.6927E-05 | 0 | 5.3997E-05 | 0.00005442 | 4.5115E-05 | 0 | 0 | 0 | 0 | 0 | 2 | 0 | 4 | 4 | 3.33333333 | 0 | 0 | 0 | 0 | 0 |
| H1BP3 | HCLS1-bindir | 0 | 0.06984157 | 0.00020195 | 0.00013595 | 0.00040498 | 0 | 0.00024763 | 0 | 0 | 0 | 0 | 0 | 3 | 2 | 6 | 0 | 3.66666667 | 0 | 0 | 0 | 0 | 0 |
| DHX33 | Putative ATP- | 0 | 0.03212769 | 7.4649E-05 | 0.00011307 | 0 | 7.5433E-05 | 8.7717E-05 | 0 | 0 | 0 | 0 | 0 | 2 | 3 | 0 | 2 | 2.33333333 | 0 | 0 | 0 | 0 | 0 |
| MSL1 | Male-specific | 0 | 0.02577884 | 0.00021489 | 0 | 0.00021546 | 0.00017372 | 0.00020136 | 0 | 0 | 0 | 0 | 0 | 5 | 0 | 5 | 4 | 4.66666667 | 0 | 0 | 0 | 0 | 0 |
| ZRAB2 | Zinc finger Ra | 0 | 0.04931514 | 0.00031986 | 0.00016149 | 0.00016036 | 0 | 0.0002139 | 0 | 0 | 0 | 0 | 0 | 4 | 2 | 2 | 0 | 2.66666667 | 0 | 0 | 0 | 0 | 0 |
| KAT5 | Histone acety | 0 | 0.03894614 | 0.00015432 | 0.00010388 | 0 | 0.00020792 | 0.00015537 | 0 | 0 | 0 | 0 | 0 | 3 | 2 | 0 | 4 | 3 | 0 | 0 | 0 | 0 | 0 |
| RPA34 | DNA-directec | 0 | 0.030208 | 0.00010348 | 0 | 0.00015564 | 0.00015686 | 0.00013866 | 0 | 0 | 0 | 0 | 0 | 2 | 0 | 3 | 3 | 2.66666667 | 0 | 0 | 0 | 0 | 0 |
| TRIR | Telomerase F | 0 | 0.02797166 | 0.0004498 | 0.0004542 | 0 | 0.00060604 | 0.00050335 | 0 | 0 | 0 | 0 | 0 | 3 | 3 | 0 | 4 | 3.33333333 | 0 | 0 | 0 | 0 | 0 |
| C8AP2 | CASP8-associ | 0 | 0.02401187 | 2.6628E-05 | 0 | 2.6699E-05 | 2.6908E-05 | 2.6745E-05 | 0 | 0 | 0 | 0 | 0 | 2 | 0 | 2 | 2 | 2 | 0 | 0 | 0 | 0 | 0 |
| CC137 | Coiled-coil dc | 0 | 0.02401186 | 0.00018262 | 0 | 0.00018311 | 0.00018454 | 0.00018342 | 0 | 0 | 0 | 0 | 0 | 2 | 0 | 2 | 2 | 2 | 0 | 0 | 0 | 0 | 0 |
| S30BP | SAP30-bindir | 0 | 0.03212733 | 0.00017135 | 0.00025954 | 0 | 0.00017315 | 0.00020135 | 0 | 0 | 0 | 0 | 0 | 2 | 3 | 0 | 2 | 2.33333333 | 0 | 0 | 0 | 0 | 0 |
| UBP15 | Ubiquitin car | 0 | 0.04036242 | 0 | 0.00010865 | 5.3942E-05 | 0.00010873 | 9.0441E-05 | 0 | 0 | 0 | 0 | 0 | 0 | 4 | 2 | 4 | 3.33333333 | 0 | 0 | 0 | 0 | 0 |
| CNDG2 | Condensin-2 | 0 | 0.031674 | 6.9261E-05 | 4.6626E-05 | 4.6297E-05 | 0 | 5.4061E-05 | 0 | 0 | 0 | 0 | 0 | 3 | 2 | 2 | 0 | 2.33333333 | 0 | 0 | 0 | 0 | 0 |
| RBM7 | RNA-binding | 0 | 0.02794123 | 0.00029761 | 0.0004007 | 0 | 0.00030074 | 0.00033302 | 0 | 0 | 0 | 0 | 0 | 3 | 4 | 0 | 3 | 3.33333333 | 0 | 0 | 0 | 0 | 0 |
| PR38A | Pre-mRNA-sp | 0 | 0.03971633 | 0.00033831 | 0.00017081 | 0.00033921 | 0 | 0.00028278 | 0 | 0 | 0 | 0 | 0 | 4 | 2 | 4 | 0 | 3.33333333 | 0 | 0 | 0 | 0 | 0 |
| AMOL1 | Angiomotin-l | 0 | 0.02401186 | 5.5206E-05 | 0 | 5.5353E-05 | 5.5786E-05 | 5.5448E-05 | 0 | 0 | 0 | 0 | 0 | 4 | 0 | 4 | 5 | 4.33333333 | 0 | 0 | 0 | 0 | 0 |
| CSTF3 | Cleavage stin | 0 | 0.04047146 | 7.3608E-05 | 0.00014866 | 0 | 0.00014876 | 0.00012368 | 0 | 0 | 0 | 0 | 0 | 2 | 4 | 0 | 4 | 3.33333333 | 0 | 0 | 0 | 0 | 0 |
| TM209 | Transmembr | 0 | 0.03022841 | 0 | 0.00014249 | 9.4327E-05 | 0.0001426 | 0.00012647 | 0 | 0 | 0 | 0 | 0 | 0 | 3 | 2 | 3 | 2.66666667 | 0 | 0 | 0 | 0 | 0 |
| TFDP1 | Transcription | 0 | 0.05381321 | 0.00019309 | 0.00032496 | 0.00012907 | 0 | 0.00021571 | 0 | 0 | 0 | 0 | 0 | 3 | 5 | 2 | 0 | 3.33333333 | 0 | 0 | 0 | 0 | 0 |
| BORG5 | Cdc42 effect | 0 | 0.0391798 | 0.00020247 | 0.0002726 | 0.00013534 | 0 | 0.00020347 | 0 | 0 | 0 | 0 | 0 | 3 | 4 | 2 | 0 | 3 | 0 | 0 | 0 | 0 | 0 |
| TMX1 | Thioredoxin- | 0 | 0.02976787 | 0.00028273 | 0 | 0.00028349 | 0.00019047 | 0.00025223 | 0 | 0 | 0 | 0 | 0 | 3 | 0 | 3 | 2 | 2.66666667 | 0 | 0 | 0 | 0 | 0 |
| MBOA7 | Lysophospho | 0 | 0.03882572 | 0.00011182 | 0 | 0.00022423 | 0.00016948 | 0.00016851 | 0 | 0 | 0 | 0 | 0 | 2 | 0 | 4 | 3 | 3 | 0 | 0 | 0 | 0 | 0 |
| ARP8 | Actin-related | 0 | 0.02990447 | 0 | 0.00012811 | 0.00012721 | 8.5467E-05 | 0.0001136 | 0 | 0 | 0 | 0 | 0 | 0 | 3 | 3 | 2 | 2.66666667 | 0 | 0 | 0 | 0 | 0 |
| ATX7 | Ataxin-7 OS= | 0 | 0.03211563 | 0 | 5.9745E-05 | 5.9324E-05 | 8.9683E-05 | 6.9584E-05 | 0 | 0 | 0 | 0 | 0 | 0 | 2 | 2 | 3 | 2.33333333 | 0 | 0 | 0 | 0 | 0 |
| MRT4 | mRNA turnov | 0 | 0.02401048 | 0 | 0.00022298 | 0.00022141 | 0.00022314 | 0.00022251 | 0 | 0 | 0 | 0 | 0 | 0 | 2 | 2 | 2 | 2 | 0 | 0 | 0 | 0 | 0 |
| PSMD4 | 26S proteaso | 0 | 0.02705042 | 0.00027998 | 0.00021204 | 0 | 0.00028292 | 0.00025831 | 0 | 0 | 0 | 0 | 0 | 4 | 3 | 0 | 4 | 3.66666667 | 0 | 0 | 0 | 0 | 0 |
| MPP5 | MAGUK p55 | 0 | 0.04778812 | 0.00015638 | 0.00019738 | 0 | 7.9009E-05 | 0.00014426 | 0 | 0 | 0 | 0 | 0 | 4 | 5 | 0 | 2 | 3.66666667 | 0 | 0 | 0 | 0 | 0 |
| NMT1 | Glycylpeptide | 0 | 0.04961856 | 0.0001064 | 0 | 0.00021338 | 0.00010752 | 0.00014243 | 0 | 0 | 0 | 0 | 0 | 2 | 0 | 4 | 2 | 2.66666667 | 0 | 0 | 0 | 0 | 0 |
| RFC2 | Replication fa | 0 | 0.02986984 | 0.00022363 | 0.00022582 | 0 | 0.00015065 | 0.00020003 | 0 | 0 | 0 | 0 | 0 | 3 | 3 | 0 | 2 | 2.66666667 | 0 | 0 | 0 | 0 | 0 |
| PTN11 | Tyrosine-pro | 0 | 0.03217379 | 8.8403E-05 | 8.9268E-05 | 0 | 0.000134 | 0.00010389 | 0 | 0 | 0 | 0 | 0 | 2 | 2 | 0 | 3 | 2.33333333 | 0 | 0 | 0 | 0 | 0 |
| ZN516 | Zinc finger pr | 0 | 0.05017891 | 0 | 4.5824E-05 | 4.5501E-05 | 9.1713E-05 | 6.1013E-05 | 0 | 0 | 0 | 0 | 0 | 0 | 2 | 2 | 4 | 2.66666667 | 0 | 0 | 0 | 0 | 0 |
| SLU7 | Pre-mRNA-sp | 0 | 0.03217189 | 9.0063E-05 | 9.0944E-05 | 0 | 0.00013651 | 0.00010584 | 0 | 0 | 0 | 0 | 0 | 2 | 2 | 0 | 3 | 2.33333333 | 0 | 0 | 0 | 0 | 0 |
| RSBN1 | Lysine-specif | 0 | 0.0240124 | 6.5807E-05 | 0.00006645 | 0 | 6.6498E-05 | 6.6252E-05 | 0 | 0 | 0 | 0 | 0 | 2 | 2 | 0 | 2 | 2 | 0 | 0 | 0 | 0 | 0 |
| KCNQ5 | Potassium vc | 0 | 0.03888036 | 8.4941E-05 | 0.00011436 | 0 | 5.7222E-05 | 8.5508E-05 | 0 | 0 | 0 | 0 | 0 | 3 | 4 | 0 | 2 | 3 | 0 | 0 | 0 | 0 | 0 |
| LIN9 | Protein lin-9 | 0 | 0.03009663 | 0.00014606 | 0 | 9.7634E-05 | 0.0001476 | 0.00013043 | 0 | 0 | 0 | 0 | 0 | 3 | 0 | 2 | 3 | 2.66666667 | 0 | 0 | 0 | 0 | 0 |
| SMRD2 | SWI/SNF-rela | 0 | 0.03211493 | 0 | 0.00010036 | 9.9656E-05 | 0.00015065 | 0.00011689 | 0 | 0 | 0 | 0 | 0 | 0 | 3 | 6 | 4 | 4.33333333 | 0 | 0 | 0 | 0 | 0 |
| TACC1 | Transforming | 0 | 0.03853374 | 9.8342E-05 | 0 | 0.00013147 | 0.00006625 | 9.8687E-05 | 0 | 0 | 0 | 0 | 0 | 3 | 0 | 4 | 2 | 3 | 0 | 0 | 0 | 0 | 0 |
| PPIL1 | Peptidyl-prol | 0 | 0.03167438 | 0.0004769 | 0.00032104 | 0.00031878 | 0 | 0.00037224 | 0 | 0 | 0 | 0 | 0 | 3 | 2 | 2 | 0 | 2.33333333 | 0 | 0 | 0 | 0 | 0 |
| TAD2B | Transcription | 0 | 0.02401135 | 0.00012566 | 0.00012689 | 0.00012599 | 0 | 0.00012618 | 0 | 0 | 0 | 0 | 0 | 2 | 2 | 2 | 0 | 2 | 0 | 0 | 0 | 0 | 0 |
| CAF1B | Chromatin as | 0 | 0.05008885 | 0 | 0.00019067 | 9.4664E-05 | 9.5405E-05 | 0.00012691 | 0 | 0 | 0 | 0 | 0 | 0 | 4 | 2 | 2 | 2.66666667 | 0 | 0 | 0 | 0 | 0 |
| KDM4B | Lysine-specif | 0 | 0.02989953 | 7.2231E-05 | 4.8625E-05 | 0 | 0.00007299 | 6.4615E-05 | 0 | 0 | 0 | 0 | 0 | 3 | 2 | 0 | 3 | 2.66666667 | 0 | 0 | 0 | 0 | 0 |
| TAF9 | Transcription | 0 | 0.03859773 | 0.00039982 | 0 | 0.00020044 | 0.00030302 | 0.00030109 | 0 | 0 | 0 | 0 | 0 | 14 | 0 | 13 | 17 | 14.6666667 | 0 | 0 | 0 | 0 | 0 |
| ING3 | Inhibitor of g | 0 | 0.02978623 | 0.00018939 | 0.0001275 | 0.00018989 | 0 | 0.00016893 | 0 | 0 | 0 | 0 | 0 | 3 | 2 | 3 | 0 | 2.66666667 | 0 | 0 | 0 | 0 | 0 |
| RPC3 | DNA-directec | 0 | 0.02976771 | 0.00014825 | 0 | 0.00014864 | 9.9871E-05 | 0.00013225 | 0 | 0 | 0 | 0 | 0 | 3 | 0 | 3 | 2 | 2.66666667 | 0 | 0 | 0 | 0 | 0 |
| ZN536 | Zinc finger pr | 0 | 0.05017879 | 0 | 4.0995E-05 | 4.0706E-05 | 8.2048E-05 | 5.4583E-05 | 0 | 0 | 0 | 0 | 0 | 0 | 2 | 2 | 4 | 2.66666667 | 0 | 0 | 0 | 0 | 0 |
| DPCD | Protein DPCD | 0 | 0.02401243 | 0.00025998 | 0.00026253 | 0 | 0.00026272 | 0.00026174 | 0 | 0 | 0 | 0 | 0 | 2 | 2 | 0 | 2 | 2 | 0 | 0 | 0 | 0 | 0 |
| MOC2B | Molybdopter | 0 | 0.03921139 | 0.00028073 | 0.00042521 | 0 | 0.00056735 | 0.00042443 | 0 | 0 | 0 | 0 | 0 | 2 | 3 | 0 | 4 | 3 | 0 | 0 | 0 | 0 | 0 |
| PLAP | Phospholipas | 0 | 0.02702874 | 0.00013277 | 0.00013407 | 0 | 0.00010062 | 0.00012249 | 0 | 0 | 0 | 0 | 0 | 4 | 4 | 0 | 3 | 3.66666667 | 0 | 0 | 0 | 0 | 0 |
| REXO4 | RNA exonucl | 0 | 0.03211487 | 0 | 0.00012629 | 0.0001254 | 0.00018957 | 0.00014709 | 0 | 0 | 0 | 0 | 0 | 0 | 2 | 2 | 3 | 2.33333333 | 0 | 0 | 0 | 0 | 0 |
| NT5D1 | 5'-nucleotida | 0 | 0.03030494 | 0.00011599 | 0.00017569 | 0 | 0.00017582 | 0.00015583 | 0 | 0 | 0 | 0 | 0 | 2 | 3 | 0 | 3 | 2.66666667 | 0 | 0 | 0 | 0 | 0 |
| HCFC2 | Host cell fact | 0 | 0.05235585 | 0.00016659 | 6.7289E-05 | 0 | 0.00010101 | 0.00011163 | 0 | 0 | 0 | 0 | 0 | 6 | 3 | 0 | 4 | 4.33333333 | 0 | 0 | 0 | 0 | 0 |
| INT4 | Integrator co | 0 | 0.03151249 | 8.2207E-05 | 5.5341E-05 | 0 | 0.00005538 | 6.4309E-05 | 0 | 0 | 0 | 0 | 0 | 3 | 2 | 0 | 2 | 2.33333333 | 0 | 0 | 0 | 0 | 0 |
| EMC2 | ER membran | 0 | 0.03921142 | 0.0001777 | 0.00026916 | 0 | 0.00035913 | 0.00026866 | 0 | 0 | 0 | 0 | 0 | 2 | 3 | 0 | 4 | 3 | 0 | 0 | 0 | 0 | 0 |
| F192A | Protein FAM1 | 0 | 0.03217411 | 0.00020778 | 0.00020981 | 0 | 0.00031495 | 0.00024418 | 0 | 0 | 0 | 0 | 0 | 2 | 2 | 0 | 3 | 2.33333333 | 0 | 0 | 0 | 0 | 0 |
| CSN2 | COP9 signalo | 0 | 0.03162182 | 0 | 0.0001203 | 0.00017918 | 0.00012039 | 0.00013996 | 0 | 0 | 0 | 0 | 0 | 0 | 2 | 3 | 2 | 2.33333333 | 0 | 0 | 0 | 0 | 0 |
| MEF2D | Myocyte-spe | 0 | 0.02401047 | 0 | 0.00010229 | 0.00010157 | 0.00010236 | 0.00010207 | 0 | 0 | 0 | 0 | 0 | 0 | 2 | 2 | 2 | 2 | 0 | 0 | 0 | 0 | 0 |
| SENP6 | Sentrin-speci | 0 | 0.02401186 | 4.7461E-05 | 0 | 4.7588E-05 | 0.00004796 | 4.767E-05 | 0 | 0 | 0 | 0 | 0 | 2 | 0 | 2 | 2 | 2 | 0 | 0 | 0 | 0 | 0 |
| CCNL1 | Cyclin-L1 OS= | 0 | 0.03019565 | 0.00010034 | 0.00015198 | 0.00015091 | 0 | 0.00013441 | 0 | 0 | 0 | 0 | 0 | 2 | 3 | 3 | 0 | 2.66666667 | 0 | 0 | 0 | 0 | 0 |
| TOM1 | Target of My | 0 | 0.0321153 | 0 | 0.00010832 | 0.00010756 | 0.0001626 | 0.00012616 | 0 | 0 | 0 | 0 | 0 | 0 | 2 | 2 | 3 | 2.33333333 | 0 | 0 | 0 | 0 | 0 |
| RAB18 | Ras-related p | 0 | 0.0318418 | 0.0002562 | 0.0002587 | 0.00038532 | 0 | 0.00030007 | 0 | 0 | 0 | 0 | 0 | 2 | 2 | 3 | 0 | 2.33333333 | 0 | 0 | 0 | 0 | 0 |
| UTP4 | U3 small nuc | 0 | 0.02401241 | 7.6934E-05 | 7.7687E-05 | 0 | 7.7742E-05 | 7.7454E-05 | 0 | 0 | 0 | 0 | 0 | 2 | 2 | 0 | 2 | 2 | 0 | 0 | 0 | 0 | 0 |
| T2AG | Transcription | 0 | 0.0240124 | 0.00072629 | 0.00073339 | 0 | 0.00073392 | 0.0007312 | 0 | 0 | 0 | 0 | 0 | 3 | 3 | 0 | 3 | 3 | 0 | 0 | 0 | 0 | 0 |
| RASN | GTPase NRas | 0 | 0.02774256 | 0.00041886 | 0 | 0.00055997 | 0.00042326 | 0.00046736 | 0 | 0 | 0 | 0 | 0 | 3 | 0 | 4 | 3 | 3.33333333 | 0 | 0 | 0 | 0 | 0 |
| 6PGD | 6-phosphoglu | 0 | 0.02401242 | 0.00010927 | 0.00011034 | 0 | 0.00011042 | 0.00011001 | 0 | 0 | 0 | 0 | 0 | 2 | 2 | 0 | 2 | 2 | 0 | 0 | 0 | 0 | 0 |
| MED4 | Mediator of I | 0 | 0.03182788 | 0.00019547 | 0 | 0.00029399 | 0.00019752 | 0.00022899 | 0 | 0 | 0 | 0 | 0 | 2 | 0 | 3 | 2 | 2.33333333 | 0 | 0 | 0 | 0 | 0 |
| KMT2A | Histone-lysin | 0 | 5.7403E-09 | 0.000492 | 0.00053709 | 0.00053997 | 0.00052404 | 0.00052328 | 0 | 0 | 0 | 0 | 0 | 74 | 80 | 81 | 78 | 78.25 | 0 | 0 | 0 | 0 | 0 |
| INT1 | Integrator co | 0 | 1.4804E-06 | 0.0010242 | 0.0010707 | 0.00086987 | 0.00087668 | 0.00096036 | 0 | 0 | 0 | 0 | 0 | 85 | 88 | 72 | 72 | 79.25 | 0 | 0 | 0 | 0 | 0 |
| BPTF | Nucleosome- | 0 | 4.0072E-10 | 0.00032921 | 0.00034992 | 0.00033877 | 0.00033266 | 0.00033764 | 0 | 0 | 0 | 0 | 0 | 38 | 40 | 39 | 38 | 38.75 | 0 | 0 | 0 | 0 | 0 |
| NSD1 | Histone-lysin | 0 | 1.8801E-06 | 0.00038173 | 0.00029651 | 0.00032386 | 0.00031651 | 0.00032965 | 0 | 0 | 0 | 0 | 0 | 39 | 30 | 33 | 32 | 33.5 | 0 | 0 | 0 | 0 | 0 |
| PHF3 | PHD finger pr | 0 | 2.0035E-06 | 0.00050473 | 0.00041819 | 0.00040227 | 0.00040541 | 0.00043265 | 0 | 0 | 0 | 0 | 0 | 39 | 32 | 31 | 31 | 33.25 | 0 | 0 | 0 | 0 | 0 |
| ZN318 | Zinc finger pr | 0 | 4.6875E-05 | 0.00035895 | 0.000304 | 0.00042956 | 0.00028081 | 0.00034333 | 0 | 0 | 0 | 0 | 0 | 31 | 26 | 37 | 24 | 29.5 | 0 | 0 | 0 | 0 | 0 |
| MED1 | Mediator of I | 0 | 1.0966E-05 | 0.00070102 | 0.00082586 | 0.000569 | 0.00072525 | 0.00070528 | 0 | 0 | 0 | 0 | 0 | 42 | 49 | 34 | 43 | 42 | 0 | 0 | 0 | 0 | 0 |
| DIDO1 | Death-induce | 0 | 5.1125E-07 | 0.00035342 | 0.00040446 | 0.00036617 | 0.00042855 | 0.00038815 | 0 | 0 | 0 | 0 | 0 | 30 | 34 | 31 | 36 | 32.75 | 0 | 0 | 0 | 0 | 0 |
| TAF6 | Transcription | 0 | 4.3677E-06 | 0.0017151 | 0.0016137 | 0.0016024 | 0.0012604 | 0.0015479 | 0 | 0 | 0 | 0 | 0 | 44 | 41 | 41 | 32 | 39.5 | 0 | 0 | 0 | 0 | 0 |
| TPR | Nucleoprotei | 0 | 8.6316E-06 | 0.00030152 | 0.00034957 | 0.00042549 | 0.00038368 | 0.00036507 | 0 | 0 | 0 | 0 | 0 | 27 | 31 | 38 | 34 | 32.5 | 0 | 0 | 0 | 0 | 0 |
| TIF1B | Transcription | 0 | 4.6141E-06 | 0.0011693 | 0.0013084 | 0.0012358 | 0.00095805 | 0.00116789 | 0 | 0 | 0 | 0 | 0 | 37 | 41 | 39 | 30 | 36.75 | 0 | 0 | 0 | 0 | 0 |
| WDR33 | pre-mRNA 3' | 0 | 7.2605E-06 | 0.00065181 | 0.00063824 | 0.00049511 | 0.00051894 | 0.00057603 | 0 | 0 | 0 | 0 | 0 | 33 | 32 | 25 | 26 | 29 | 0 | 0 | 0 | 0 | 0 |
| PP1RA | Serine/threo | 0 | 1.9551E-05 | 0.00087026 | 0.00076538 | 0.00067554 | 0.00099287 | 0.00082601 | 0 | 0 | 0 | 0 | 0 | 31 | 27 | 24 | 35 | 29.25 | 0 | 0 | 0 | 0 | 0 |
| RBBP6 | E3 ubiquitin- | 0 | 3.1692E-06 | 0.00042704 | 0.00055018 | 0.00047248 | 0.00043153 | 0.00047031 | 0 | 0 | 0 | 0 | 0 | 29 | 37 | 32 | 29 | 31.75 | 0 | 0 | 0 | 0 | 0 |
| CDK13 | Cyclin-depen | 0 | 6.5335E-06 | 0.00050613 | 0.00038771 | 0.00047248 | 0.00038799 | 0.00043858 | 0 | 0 | 0 | 0 | 0 | 29 | 22 | 27 | 22 | 25 | 0 | 0 | 0 | 0 | 0 |
| LMBL3 | Lethal(3)mali | 0 | 1.7626E-06 | 0.0011164 | 0.0014006 | 0.0011194 | 0.0012307 | 0.00121678 | 0 | 0 | 0 | 0 | 0 | 33 | 41 | 33 | 36 | 35.75 | 0 | 0 | 0 | 0 | 0 |
| NOL8 | Nucleolar prc | 0 | 1.3593E-06 | 0.00067837 | 0.00059367 | 0.00058948 | 0.00052554 | 0.00059677 | 0 | 0 | 0 | 0 | 0 | 30 | 26 | 26 | 23 | 26.25 | 0 | 0 | 0 | 0 | 0 |

| | | | | | | | | | | | | | | | | | | | | | | | |
|---|---|---|---|---|---|---|---|---|---|---|---|---|---|---|---|---|---|---|---|---|---|---|---|
| BD1L1 | Biorientation | 0 | 4.4933E-06 | 0.00027677 | 0.00023581 | 0.00020813 | 0.0002185 | 0.0002348 | 0 | 0 | 0 | 0 | 0 | 32 | 27 | 24 | 25 | 27 | 0 | 0 | 0 | 0 | 0 |
| NIPBL | Nipped-B-like | 0 | 1.1599E-06 | 0.00018822 | 0.00023758 | 0.00020759 | 0.00022824 | 0.00021541 | 0 | 0 | 0 | 0 | 0 | 20 | 25 | 22 | 24 | 22.75 | 0 | 0 | 0 | 0 | 0 |
| TAF5 | Transcription | 0 | 1.3087E-06 | 0.0010885 | 0.00099924 | 0.0010914 | 0.00086663 | 0.00101144 | 0 | 0 | 0 | 0 | 0 | 33 | 30 | 33 | 26 | 30.5 | 0 | 0 | 0 | 0 | 0 |
| ZC11A | Zinc finger C( | 0 | 5.3235E-06 | 0.00094477 | 0.00098691 | 0.00078396 | 0.0010864 | 0.00095051 | 0 | 0 | 0 | 0 | 0 | 29 | 30 | 24 | 33 | 29 | 0 | 0 | 0 | 0 | 0 |
| CHAP1 | Chromosome | 0 | 1.6497E-05 | 0.00087745 | 0.00098448 | 0.00068428 | 0.00075531 | 0.00082538 | 0 | 0 | 0 | 0 | 0 | 27 | 30 | 21 | 23 | 25.25 | 0 | 0 | 0 | 0 | 0 |
| ADNP | Activity-depe | 0 | 5.4093E-09 | 0.00055076 | 0.00053196 | 0.0005042 | 0.00050815 | 0.00052377 | 0 | 0 | 0 | 0 | 0 | 23 | 22 | 21 | 21 | 21.75 | 0 | 0 | 0 | 0 | 0 |
| EP400 | E1A-binding | 0 | 0.00027492 | 0.0002506 | 0.00021088 | 0.00016751 | 0.00013506 | 0.00019101 | 0 | 0 | 0 | 0 | 0 | 30 | 25 | 20 | 16 | 22.75 | 0 | 0 | 0 | 0 | 0 |
| TAF6L | TAF6-like RN. | 0 | 8.0285E-05 | 0.00093335 | 0.0008568 | 0.0012336 | 0.00077167 | 0.00094886 | 0 | 0 | 0 | 0 | 0 | 22 | 20 | 29 | 18 | 22.25 | 0 | 0 | 0 | 0 | 0 |
| MINT | Msx2-interac | 0 | 4.0047E-05 | 0.00011523 | 0.00010182 | 7.2213E-05 | 9.4611E-05 | 9.5969E-05 | 0 | 0 | 0 | 0 | 0 | 16 | 14 | 10 | 13 | 13.25 | 0 | 0 | 0 | 0 | 0 |
| TAF4 | Transcription | 0 | 2.0584E-06 | 0.00063235 | 0.00066309 | 0.0006828 | 0.00081103 | 0.00069732 | 0 | 0 | 0 | 0 | 0 | 26 | 27 | 28 | 33 | 28.5 | 0 | 0 | 0 | 0 | 0 |
| PKCB1 | Protein kinas | 0 | 5.8971E-09 | 0.00051175 | 0.00053922 | 0.00055773 | 0.00056209 | 0.0005427 | 0 | 0 | 0 | 0 | 0 | 23 | 24 | 25 | 25 | 24.25 | 0 | 0 | 0 | 0 | 0 |
| KNOP1 | Lysine-rich nu | 0 | 6.5636E-07 | 0.0017285 | 0.0015709 | 0.0017909 | 0.0014555 | 0.00163645 | 0 | 0 | 0 | 0 | 0 | 30 | 27 | 31 | 25 | 28.25 | 0 | 0 | 0 | 0 | 0 |
| GANP | Germinal-cer | 0 | 3.7381E-06 | 0.00030653 | 0.00024224 | 0.00029399 | 0.00024241 | 0.00027129 | 0 | 0 | 0 | 0 | 0 | 23 | 18 | 22 | 18 | 20.25 | 0 | 0 | 0 | 0 | 0 |
| CHD8 | Chromodoma | 0 | 1.8175E-05 | 0.00014314 | 0.00015486 | 0.00017427 | 0.00020663 | 0.00016973 | 0 | 0 | 0 | 0 | 0 | 14 | 15 | 17 | 20 | 16.5 | 0 | 0 | 0 | 0 | 0 |
| CDCA2 | Cell division | 0 | 4.9065E-05 | 0.00069647 | 0.00075537 | 0.000569 | 0.00091231 | 0.00073329 | 0 | 0 | 0 | 0 | 0 | 27 | 29 | 22 | 35 | 28.25 | 0 | 0 | 0 | 0 | 0 |
| YLPM1 | YLP motif-co | 0 | 1.8443E-05 | 0.00021641 | 0.00021853 | 0.00018986 | 0.00015034 | 0.00019379 | 0 | 0 | 0 | 0 | 0 | 16 | 16 | 14 | 11 | 14.25 | 0 | 0 | 0 | 0 | 0 |
| BCOR | BCL-6 corepr | 0 | 5.5085E-05 | 0.0003759 | 0.00030366 | 0.0003769 | 0.00024311 | 0.00032489 | 0 | 0 | 0 | 0 | 0 | 25 | 20 | 25 | 16 | 21.5 | 0 | 0 | 0 | 0 | 0 |
| NCOR1 | Nuclear rece | 0 | 0.00010584 | 0.00023793 | 0.00028394 | 0.00016266 | 0.00021857 | 0.00022578 | 0 | 0 | 0 | 0 | 0 | 22 | 26 | 15 | 20 | 20.75 | 0 | 0 | 0 | 0 | 0 |
| SLX4 | Structure-spe | 0 | 2.0398E-05 | 0.00031655 | 0.00030511 | 0.00038952 | 0.00026171 | 0.00031822 | 0 | 0 | 0 | 0 | 0 | 22 | 21 | 27 | 18 | 22 | 0 | 0 | 0 | 0 | 0 |
| SAFB2 | Scaffold atta | 0 | 2.5516E-09 | 0.00058149 | 0.00055921 | 0.00055527 | 0.00053163 | 0.0005569 | 0 | 0 | 0 | 0 | 0 | 21 | 20 | 20 | 19 | 20 | 0 | 0 | 0 | 0 | 0 |
| U5S1 | 116 kDa U5 s | 0 | 8.1342E-05 | 0.00051582 | 0.00068535 | 0.00078941 | 0.00052124 | 0.00062796 | 0 | 0 | 0 | 0 | 0 | 19 | 25 | 29 | 19 | 23 | 0 | 0 | 0 | 0 | 0 |
| WAPL | Wings apart- | 0 | 3.775E-06 | 0.00048785 | 0.00038066 | 0.00037798 | 0.00040335 | 0.00041246 | 0 | 0 | 0 | 0 | 0 | 22 | 17 | 17 | 18 | 18.5 | 0 | 0 | 0 | 0 | 0 |
| XPF | DNA repair e | 0 | 2.2801E-08 | 0.00080663 | 0.00090179 | 0.00080878 | 0.00084422 | 0.00084036 | 0 | 0 | 0 | 0 | 0 | 28 | 31 | 28 | 29 | 29 | 0 | 0 | 0 | 0 | 0 |
| TRPS1 | Zinc finger tr | 0 | 1.596E-05 | 0.0004944 | 0.00037442 | 0.00053702 | 0.00054122 | 0.00048677 | 0 | 0 | 0 | 0 | 0 | 24 | 18 | 26 | 26 | 23.5 | 0 | 0 | 0 | 0 | 0 |
| RRP1B | Ribosomal R | 0 | 3.6031E-05 | 0.00066145 | 0.00066792 | 0.00062831 | 0.00091465 | 0.00071808 | 0 | 0 | 0 | 0 | 0 | 19 | 19 | 18 | 26 | 20.5 | 0 | 0 | 0 | 0 | 0 |
| TPM3 | Tropomyosin | 0 | 0.0011903 | 0.0017592 | 0.001122 | 0.0022281 | 0.0027133 | 0.00195565 | 0 | 0 | 0 | 0 | 0 | 31 | 19 | 42 | 48 | 35 | 0 | 0 | 0 | 0 | 0 |
| ATAD5 | ATPase famil | 0 | 0.00012835 | 0.00040069 | 0.00028901 | 0.00030132 | 0.00023137 | 0.0003056 | 0 | 0 | 0 | 0 | 0 | 28 | 20 | 21 | 16 | 21.25 | 0 | 0 | 0 | 0 | 0 |
| ZC3H1 | Zinc finger C3 | 0 | 1.9975E-06 | 0.00026534 | 0.00024114 | 0.00023945 | 0.0002011 | 0.00023676 | 0 | 0 | 0 | 0 | 0 | 20 | 18 | 18 | 15 | 17.75 | 0 | 0 | 0 | 0 | 0 |
| CCAR1 | Cell division | 0 | 0.00022856 | 0.00041304 | 0.00025488 | 0.00027609 | 0.00041737 | 0.00034035 | 0 | 0 | 0 | 0 | 0 | 18 | 11 | 12 | 18 | 14.75 | 0 | 0 | 0 | 0 | 0 |
| CUX1 | Homeobox p | 0 | 0.00017185 | 0.00042081 | 0.0003187 | 0.00045709 | 0.00026577 | 0.00036559 | 0 | 0 | 0 | 0 | 0 | 24 | 18 | 26 | 15 | 20.75 | 0 | 0 | 0 | 0 | 0 |
| KDM3B | Lysine-specif | 0 | 3.5503E-05 | 0.00028471 | 0.00025723 | 0.0001803 | 0.00025742 | 0.00024492 | 0 | 0 | 0 | 0 | 0 | 19 | 17 | 12 | 17 | 16.25 | 0 | 0 | 0 | 0 | 0 |
| SIN3A | Paired amphi | 0 | 7.7988E-05 | 0.00043532 | 0.00035584 | 0.00047804 | 0.00029326 | 0.00039062 | 0 | 0 | 0 | 0 | 0 | 21 | 17 | 23 | 14 | 18.75 | 0 | 0 | 0 | 0 | 0 |
| TPM4 | Tropomyosin | 0 | 0.00106454 | 0.0015961 | 0.0012893 | 0.0022405 | 0.0027956 | 0.00198038 | 0 | 0 | 0 | 0 | 0 | 15 | 12 | 21 | 26 | 18.5 | 0 | 0 | 0 | 0 | 0 |
| HIRA | Protein HIRA | 0 | 5.2374E-06 | 0.00041516 | 0.00041922 | 0.00054634 | 0.00047196 | 0.00046317 | 0 | 0 | 0 | 0 | 0 | 16 | 16 | 21 | 18 | 17.75 | 0 | 0 | 0 | 0 | 0 |
| WDHD1 | WD repeat a | 0 | 3.2627E-07 | 0.00037397 | 0.00040123 | 0.00035153 | 0.00042514 | 0.00038797 | 0 | 0 | 0 | 0 | 0 | 16 | 17 | 15 | 18 | 16.5 | 0 | 0 | 0 | 0 | 0 |
| SPB1 | pre-rRNA pro | 0 | 3.7459E-06 | 0.00059195 | 0.00050336 | 0.00043733 | 0.0005352 | 0.00051696 | 0 | 0 | 0 | 0 | 0 | 19 | 16 | 14 | 17 | 16.5 | 0 | 0 | 0 | 0 | 0 |
| MY18A | Unconventio | 0 | 0.00085115 | 0.00016702 | 0.00020757 | 0.00032204 | 0.00033754 | 0.00025854 | 0 | 0 | 0 | 0 | 0 | 13 | 16 | 25 | 26 | 20 | 0 | 0 | 0 | 0 | 0 |
| SF3A1 | Splicing facto | 0 | 4.6973E-05 | 0.00036604 | 0.00047043 | 0.00030029 | 0.00043714 | 0.00039348 | 0 | 0 | 0 | 0 | 0 | 11 | 14 | 9 | 13 | 11.75 | 0 | 0 | 0 | 0 | 0 |
| BLM | Bloom syndr | 0 | 0.00019947 | 0.00035383 | 0.00020685 | 0.00037345 | 0.00028228 | 0.0003041 | 0 | 0 | 0 | 0 | 0 | 19 | 11 | 20 | 15 | 16.25 | 0 | 0 | 0 | 0 | 0 |
| NASP | Nuclear auto | 0 | 0.00019956 | 0.00063627 | 0.00074394 | 0.00040292 | 0.00054143 | 0.00058114 | 0 | 0 | 0 | 0 | 0 | 19 | 22 | 12 | 16 | 17.25 | 0 | 0 | 0 | 0 | 0 |
| SMCA4 | Transcription | 0 | 0.00013511 | 0.00024033 | 0.00025886 | 0.00035343 | 0.00021048 | 0.00026578 | 0 | 0 | 0 | 0 | 0 | 15 | 16 | 22 | 13 | 16.5 | 0 | 0 | 0 | 0 | 0 |
| PDS5B | Sister chrom | 0 | 0.00012905 | 0.0002006 | 0.00014732 | 0.00023771 | 0.00025799 | 0.00021091 | 0 | 0 | 0 | 0 | 0 | 11 | 8 | 13 | 14 | 11.5 | 0 | 0 | 0 | 0 | 0 |
| NVL | Nuclear valos | 0 | 1.6272E-06 | 0.00064738 | 0.00052919 | 0.0006491 | 0.00068533 | 0.00062775 | 0 | 0 | 0 | 0 | 0 | 21 | 17 | 21 | 22 | 20.25 | 0 | 0 | 0 | 0 | 0 |
| EMSY | BRCA2-intera | 0 | 0.0003428 | 0.00031938 | 0.00046359 | 0.00032023 | 0.00024205 | 0.00033631 | 0 | 0 | 0 | 0 | 0 | 16 | 23 | 16 | 12 | 16.75 | 0 | 0 | 0 | 0 | 0 |
| TF3C1 | General tran | 0 | 0.00593771 | 0.00012512 | 8.8443E-05 | 0.00025091 | 0.00012644 | 0.00014773 | 0 | 0 | 0 | 0 | 0 | 10 | 7 | 20 | 10 | 11.75 | 0 | 0 | 0 | 0 | 0 |
| TPX2 | Targeting pro | 0 | 0.00053569 | 0.00031793 | 0.00039238 | 0.0001771 | 0.00032127 | 0.00030217 | 0 | 0 | 0 | 0 | 0 | 9 | 11 | 5 | 9 | 8.5 | 0 | 0 | 0 | 0 | 0 |
| DPOLA | DNA polyme | 0 | 6.083E-08 | 0.00028879 | 0.00027339 | 0.00025337 | 0.00025535 | 0.00026773 | 0 | 0 | 0 | 0 | 0 | 16 | 15 | 14 | 14 | 14.75 | 0 | 0 | 0 | 0 | 0 |
| RPRD2 | Regulation o | 0 | 6.1566E-06 | 0.00030705 | 0.00027358 | 0.00027165 | 0.00021902 | 0.00026783 | 0 | 0 | 0 | 0 | 0 | 17 | 15 | 15 | 12 | 14.75 | 0 | 0 | 0 | 0 | 0 |
| ZN106 | Zinc finger pr | 0 | 1.8591E-06 | 0.00029429 | 0.00025472 | 0.00022482 | 0.0002549 | 0.00025718 | 0 | 0 | 0 | 0 | 0 | 21 | 18 | 16 | 18 | 18.25 | 0 | 0 | 0 | 0 | 0 |
| ZMYM2 | Zinc finger M | 0 | 0.00062551 | 0.00030662 | 0.00017416 | 0.00034587 | 0.00021302 | 0.00025992 | 0 | 0 | 0 | 0 | 0 | 16 | 9 | 18 | 11 | 13.5 | 0 | 0 | 0 | 0 | 0 |
| TBL1R | F-box-like/W | 0 | 4.0522E-08 | 0.0011295 | 0.00098499 | 0.001081 | 0.0010376 | 0.00105827 | 0 | 0 | 0 | 0 | 0 | 22 | 19 | 21 | 20 | 20.5 | 0 | 0 | 0 | 0 | 0 |
| SART3 | Squamous ce | 0 | 5.0526E-09 | 0.00038363 | 0.00038738 | 0.00041213 | 0.00041535 | 0.00039962 | 0 | 0 | 0 | 0 | 0 | 14 | 14 | 15 | 15 | 14.5 | 0 | 0 | 0 | 0 | 0 |
| MPRI | Cation-indep | 0 | 6.357E-05 | 0.00014831 | 0.00011767 | 0.0001487 | 0.00019269 | 0.00015184 | 0 | 0 | 0 | 0 | 0 | 14 | 11 | 14 | 18 | 14.25 | 0 | 0 | 0 | 0 | 0 |
| EHMT1 | Histone-lysin | 0 | 0.00021713 | 0.00032528 | 0.00018476 | 0.00034653 | 0.0003287 | 0.00029632 | 0 | 0 | 0 | 0 | 0 | 16 | 9 | 17 | 16 | 14.5 | 0 | 0 | 0 | 0 | 0 |
| DDX42 | ATP-depende | 0 | 4.8124E-06 | 0.00053452 | 0.00039771 | 0.00045132 | 0.00051171 | 0.00047382 | 0 | 0 | 0 | 0 | 0 | 19 | 14 | 16 | 18 | 16.75 | 0 | 0 | 0 | 0 | 0 |
| NOP2 | Probable 28S | 0 | 5.3127E-09 | 0.00042247 | 0.00042661 | 0.00039102 | 0.00042691 | 0.00041675 | 0 | 0 | 0 | 0 | 0 | 13 | 13 | 12 | 13 | 12.75 | 0 | 0 | 0 | 0 | 0 |
| CABIN | Calcineurin-b | 0 | 0.00412925 | 0.00020207 | 0.00008402 | 0.0001311 | 8.4081E-05 | 0.00012532 | 0 | 0 | 0 | 0 | 0 | 17 | 7 | 11 | 7 | 10.5 | 0 | 0 | 0 | 0 | 0 |
| TMOD3 | Tropomoduli | 0 | 8.413E-05 | 0.0011245 | 0.0008327 | 0.0006765 | 0.00098481 | 0.00090463 | 0 | 0 | 0 | 0 | 0 | 15 | 11 | 9 | 13 | 12 | 0 | 0 | 0 | 0 | 0 |
| CPSF5 | Cleavage and | 0 | 6.879E-08 | 0.00186 | 0.0017608 | 0.0016318 | 0.0018795 | 0.00178303 | 0 | 0 | 0 | 0 | 0 | 16 | 15 | 14 | 16 | 15.25 | 0 | 0 | 0 | 0 | 0 |
| WIZ | Protein Wiz ( | 0 | 9.298E-07 | 0.00020778 | 0.00020981 | 0.00019231 | 0.00024227 | 0.00021304 | 0 | 0 | 0 | 0 | 0 | 13 | 13 | 12 | 15 | 13.25 | 0 | 0 | 0 | 0 | 0 |
| KMT2C | Histone-lysin | 0 | 0.00140871 | 0.00006448 | 6.5111E-05 | 0.0001293 | 8.1447E-05 | 8.5085E-05 | 0 | 0 | 0 | 0 | 0 | 12 | 12 | 24 | 15 | 15.75 | 0 | 0 | 0 | 0 | 0 |
| GPTC4 | G patch dom | 0 | 4.0548E-07 | 0.0008875 | 0.0010157 | 0.00083054 | 0.00089683 | 0.00090764 | 0 | 0 | 0 | 0 | 0 | 15 | 17 | 14 | 15 | 15.25 | 0 | 0 | 0 | 0 | 0 |
| ZMYM1 | Zinc finger M | 0 | 9.4879E-10 | 0.00043904 | 0.00042 | 0.00041704 | 0.00044365 | 0.00042993 | 0 | 0 | 0 | 0 | 0 | 19 | 18 | 18 | 19 | 18.5 | 0 | 0 | 0 | 0 | 0 |
| EHMT2 | Histone-lysin | 0 | 1.7776E-05 | 0.00037075 | 0.00041842 | 0.00028427 | 0.00033057 | 0.000351 | 0 | 0 | 0 | 0 | 0 | 17 | 19 | 13 | 15 | 16 | 0 | 0 | 0 | 0 | 0 |
| SRCAP | Helicase SRC | 0 | 9.417E-05 | 0.00012255 | 9.0747E-05 | 0.00013926 | 9.0812E-05 | 0.00011084 | 0 | 0 | 0 | 0 | 0 | 15 | 11 | 17 | 11 | 13.5 | 0 | 0 | 0 | 0 | 0 |
| P66B | Transcription | 0 | 1.2672E-06 | 0.0006675 | 0.00053922 | 0.00058004 | 0.00053961 | 0.00058159 | 0 | 0 | 0 | 0 | 0 | 15 | 12 | 13 | 12 | 13 | 0 | 0 | 0 | 0 | 0 |
| TCRG1 | Transcription | 0 | 0.00011801 | 0.0003605 | 0.00021841 | 0.00024097 | 0.00029143 | 0.00027783 | 0 | 0 | 0 | 0 | 0 | 15 | 9 | 10 | 12 | 11.5 | 0 | 0 | 0 | 0 | 0 |
| BBX | HMG box tra | 0 | 1.9681E-06 | 0.00042064 | 0.00048139 | 0.00036553 | 0.00042506 | 0.00042316 | 0 | 0 | 0 | 0 | 0 | 15 | 17 | 13 | 15 | 15 | 0 | 0 | 0 | 0 | 0 |
| FUBP2 | Far upstream | 0 | 6.0564E-06 | 0.00044537 | 0.00056216 | 0.00048377 | 0.00060007 | 0.00052284 | 0 | 0 | 0 | 0 | 0 | 12 | 15 | 13 | 16 | 14 | 0 | 0 | 0 | 0 | 0 |
| NFM | Neurofilame | 0 | 5.6077E-05 | 0.00040332 | 0.00031999 | 0.00051993 | 0.00040755 | 0.0004127 | 0 | 0 | 0 | 0 | 0 | 15 | 11 | 19 | 14 | 14.75 | 0 | 0 | 0 | 0 | 0 |
| CDC73 | Parafibromin | 0 | 7.3969E-06 | 0.00089452 | 0.00075273 | 0.00069759 | 0.00065283 | 0.00074942 | 0 | 0 | 0 | 0 | 0 | 18 | 15 | 14 | 13 | 15 | 0 | 0 | 0 | 0 | 0 |
| POGZ | Pogo transpo | 0 | 6.2343E-06 | 0.00026201 | 0.00020788 | 0.00028148 | 0.00022694 | 0.00024458 | 0 | 0 | 0 | 0 | 0 | 14 | 11 | 15 | 12 | 13 | 0 | 0 | 0 | 0 | 0 |
| CPIN1 | Anamorsin O | 0 | 0.00011413 | 0.00050747 | 0.00076865 | 0.00050882 | 0.0005128 | 0.00057444 | 0 | 0 | 0 | 0 | 0 | 6 | 9 | 6 | 6 | 6.75 | 0 | 0 | 0 | 0 | 0 |
| APC1 | Anaphase-pr | 0 | 0.00010822 | 0.00013574 | 8.2242E-05 | 0.00012249 | 9.6018E-05 | 0.00010912 | 0 | 0 | 0 | 0 | 0 | 10 | 6 | 9 | 7 | 8 | 0 | 0 | 0 | 0 | 0 |
| INT12 | Integrator co | 0 | 0.00029735 | 0.00074253 | 0.00051909 | 0.00040089 | 0.00069261 | 0.00058878 | 0 | 0 | 0 | 0 | 0 | 13 | 9 | 7 | 12 | 10.25 | 0 | 0 | 0 | 0 | 0 |
| CUL4B | Cullin-4B OS= | 0 | 9.4838E-06 | 0.00052025 | 0.00046697 | 0.00037674 | 0.00052572 | 0.00047242 | 0 | 0 | 0 | 0 | 0 | 18 | 16 | 13 | 18 | 16.25 | 0 | 0 | 0 | 0 | 0 |
| OGT1 | UDP-N-acety | 0 | 1.5111E-07 | 0.00032796 | 0.00038212 | 0.00037943 | 0.00038239 | 0.00036798 | 0 | 0 | 0 | 0 | 0 | 13 | 15 | 15 | 15 | 14.5 | 0 | 0 | 0 | 0 | 0 |
| UBN2 | Ubinuclein-2 | 0 | 0.00075515 | 0.00029386 | 0.00019782 | 0.00015714 | 0.00015837 | 0.0002018 | 0 | 0 | 0 | 0 | 0 | 15 | 10 | 8 | 8 | 10.25 | 0 | 0 | 0 | 0 | 0 |
| ZN250 | Zinc finger pr | 0 | 3.8174E-06 | 0.00070683 | 0.00052341 | 0.00066147 | 0.00066664 | 0.00063959 | 0 | 0 | 0 | 0 | 0 | 15 | 11 | 14 | 14 | 13.5 | 0 | 0 | 0 | 0 | 0 |
| TPM1 | Tropomyosin | 0 | 0.00353481 | 0.001115 | 0.00056295 | 0.0015838 | 0.001784 | 0.00126144 | 0 | 0 | 0 | 0 | 0 | 17 | 11 | 25 | 28 | 20.25 | 0 | 0 | 0 | 0 | 0 |
| RED | Protein Red ( | 0 | 2.0037E-07 | 0.00066326 | 0.00066975 | 0.00057003 | 0.00067023 | 0.00064332 | 0 | 0 | 0 | 0 | 0 | 14 | 14 | 12 | 14 | 13.5 | 0 | 0 | 0 | 0 | 0 |
| LMBL2 | Lethal(3)mali | 0 | 5.9077E-05 | 0.00056146 | 0.00049135 | 0.00045036 | 0.00034041 | 0.0004609 | 0 | 0 | 0 | 0 | 0 | 15 | 13 | 12 | 9 | 12.25 | 0 | 0 | 0 | 0 | 0 |
| PCF11 | Pre-mRNA cl | 0 | 0.00244027 | 0.00025455 | 0.0001885 | 8.5076E-05 | 0.00024008 | 0.00019205 | 0 | 0 | 0 | 0 | 0 | 15 | 11 | 5 | 14 | 11.25 | 0 | 0 | 0 | 0 | 0 |
| UBP36 | Ubiquitin car | 0 | 0.00014657 | 0.0002354 | 0.00019016 | 0.00028323 | 0.00016651 | 0.00021883 | 0 | 0 | 0 | 0 | 0 | 10 | 8 | 12 | 7 | 9.25 | 0 | 0 | 0 | 0 | 0 |
| F208B | Protein FAM2 | 0 | 1.9446E-05 | 0.00010859 | 0.00016448 | 0.00014155 | 0.00014266 | 0.00013932 | 0 | 0 | 0 | 0 | 0 | 10 | 15 | 13 | 13 | 12.75 | 0 | 0 | 0 | 0 | 0 |
| CPSF4 | Cleavage and | 0 | 2.7208E-06 | 0.0011772 | 0.00089152 | 0.001082 | 0.00099129 | 0.0010355 | 0 | 0 | 0 | 0 | 0 | 12 | 9 | 11 | 10 | 10.5 | 0 | 0 | 0 | 0 | 0 |
| METH | Methionine s | 0 | 0.00136572 | 0.00012516 | 0.00025277 | 0.00016733 | 0.00012648 | 0.00016794 | 0 | 0 | 0 | 0 | 0 | 6 | 12 | 8 | 6 | 8 | 0 | 0 | 0 | 0 | 0 |
| TAF9B | Transcription | 0 | 3.5534E-05 | 0.0013667 | 0.0011678 | 0.0014758 | 0.001806 | 0.00145408 | 0 | 0 | 0 | 0 | 0 | 13 | 11 | 14 | 17 | 13.75 | 0 | 0 | 0 | 0 | 0 |
| ZC3HE | Zinc finger C( | 0 | 2.4671E-06 | 0.00039439 | 0.00043445 | 0.00039544 | 0.00032607 | 0.00038759 | 0 | 0 | 0 | 0 | 0 | 11 | 12 | 11 | 9 | 10.75 | 0 | 0 | 0 | 0 | 0 |
| RBM10 | RNA-binding | 0 | 9.1322E-05 | 0.0003405 | 0.00045843 | 0.00031295 | 0.00028673 | 0.00034965 | 0 | 0 | 0 | 0 | 0 | 12 | 16 | 11 | 10 | 12.25 | 0 | 0 | 0 | 0 | 0 |
| RBM28 | RNA-binding | 0 | 1.4156E-05 | 0.00038244 | 0.0004915 | 0.0003486 | 0.00045672 | 0.00041982 | 0 | 0 | 0 | 0 | 0 | 11 | 14 | 10 | 13 | 12 | 0 | 0 | 0 | 0 | 0 |
| SAP18 | Histone deac | 0 | 1.1024E-06 | 0.002932 | 0.0024382 | 0.0031128 | 0.0027886 | 0.0028179 | 0 | 0 | 0 | 0 | 0 | 17 | 14 | 18 | 16 | 16.25 | 0 | 0 | 0 | 0 | 0 |
| UBP7 | Ubiquitin car | 0 | 4.055E-05 | 0.00023946 | 0.00031434 | 0.00021609 | 0.00031457 | 0.00027112 | 0 | 0 | 0 | 0 | 0 | 10 | 13 | 9 | 13 | 11.25 | 0 | 0 | 0 | 0 | 0 |
| TAF5L | TAF5-like RN. | 0 | 0.00010569 | 0.00044802 | 0.00058812 | 0.00058398 | 0.00036218 | 0.00049558 | 0 | 0 | 0 | 0 | 0 | 10 | 13 | 13 | 8 | 11 | 0 | 0 | 0 | 0 | 0 |
| PRPF3 | U4/U6 small | 0 | 5.0405E-06 | 0.000425 | 0.00042915 | 0.00042613 | 0.00054659 | 0.00045672 | 0 | 0 | 0 | 0 | 0 | 11 | 11 | 11 | 14 | 11.75 | 0 | 0 | 0 | 0 | 0 |
| ACL6A | Actin-like pro | 0 | 3.1577E-06 | 0.00073814 | 0.00074536 | 0.0007401 | 0.00093236 | 0.00078899 | 0 | 0 | 0 | 0 | 0 | 12 | 12 | 12 | 15 | 12.75 | 0 | 0 | 0 | 0 | 0 |
| EXOSX | Exosome con | 0 | 0.00041133 | 0.00026836 | 0.00018065 | 0.00014948 | 0.00027118 | 0.00021742 | 0 | 0 | 0 | 0 | 0 | 9 | 6 | 5 | 9 | 7.25 | 0 | 0 | 0 | 0 | 0 |
| WDR82 | WD repeat-c | 0 | 0.00058848 | 0.001096 | 0.0005108 | 0.00076079 | 0.00093713 | 0.00082618 | 0 | 0 | 0 | 0 | 0 | 13 | 6 | 9 | 11 | 9.75 | 0 | 0 | 0 | 0 | 0 |
| MAN1 | Inner nuclear | 0 | 4.799E-09 | 0.00037656 | 0.0004095 | 0.00037757 | 0.00038052 | 0.00038604 | 0 | 0 | 0 | 0 | 0 | 13 | 14 | 13 | 13 | 13.25 | 0 | 0 | 0 | 0 | 0 |
| RBBP5 | Retinoblasto | 0 | 2.0331E-05 | 0.00049049 | 0.00064387 | 0.00059016 | 0.00044608 | 0.00054265 | 0 | 0 | 0 | 0 | 0 | 10 | 13 | 12 | 9 | 11 | 0 | 0 | 0 | 0 | 0 |
| ZMYM3 | Zinc finger M | 0 | 5.168E-06 | 0.00019262 | 0.00017505 | 0.00021244 | 0.00015571 | 0.00018396 | 0 | 0 | 0 | 0 | 0 | 12 | 12 | 14 | 8 | 11.5 | 0 | 0 | 0 | 0 | 0 |
| CDK12 | Cyclin-depen | 0 | 3.1604E-06 | 0.00021252 | 0.0002146 | 0.00021309 | 0.00026845 | 0.00022717 | 0 | 0 | 0 | 0 | 0 | 20 | 19 | 20 | 19 | 19.5 | 0 | 0 | 0 | 0 | 0 |
| UT14A | U3 small nuc | 0 | 6.126E-05 | 0.00047917 | 0.00034561 | 0.00044613 | 0.00031127 | 0.00039555 | 0 | 0 | 0 | 0 | 0 | 14 | 10 | 13 | 9 | 11.5 | 0 | 0 | 0 | 0 | 0 |
| LIN54 | Protein lin-54 | 0 | 3.8712E-05 | 0.0005637 | 0.00042691 | 0.00038858 | 0.00039162 | 0.0004427 | 0 | 0 | 0 | 0 | 0 | 16 | 12 | 11 | 11 | 12.5 | 0 | 0 | 0 | 0 | 0 |
| PRCC | Proline-rich p | 0 | 0.00854027 | 0.00042995 | 0.0010311 | 0.00048499 | 0.00038016 | 0.00058155 | 0 | 0 | 0 | 0 | 0 | 8 | 19 | 9 | 7 | 10.75 | 0 | 0 | 0 | 0 | 0 |
| STAG2 | Cohesin subu | 0 | 0.00550073 | 0.00019293 | 8.6585E-05 | 0.0003224 | 0.00021662 | 0.00020463 | 0 | 0 | 0 | 0 | 0 | 9 | 4 | 15 | 10 | 9.5 | 0 | 0 | 0 | 0 | 0 |
| KAISO | Transcription | 0 | 0.00014306 | 0.00047122 | 0.00027757 | 0.00039373 | 0.00031745 | 0.00036499 | 0 | 0 | 0 | 0 | 0 | 12 | 7 | 10 | 8 | 9.25 | 0 | 0 | 0 | 0 | 0 |
| ICE2 | Little elongat | 0 | 5.774E-06 | 0.00016123 | 0.00018994 | 0.00021555 | 0.00021724 | 0.00019599 | 0 | 0 | 0 | 0 | 0 | 6 | 7 | 8 | 8 | 7.25 | 0 | 0 | 0 | 0 | 0 |
| RBBP4 | Histone-bind | 0 | 0.0001266 | 0.00099345 | 0.00075237 | 0.00087158 | 0.00056468 | 0.00079552 | 0 | 0 | 0 | 0 | 0 | 20 | 15 | 18 | 13 | 16.5 | 0 | 0 | 0 | 0 | 0 |
| NOC3L | Nucleolar co | 0 | 1.1256E-07 | 0.00026388 | 0.00026646 | 0.00029766 | 0.00029999 | 0.000282 | 0 | 0 | 0 | 0 | 0 | 8 | 8 | 9 | 9 | 8.5 | 0 | 0 | 0 | 0 | 0 |
| SF01 | Splicing facto | 0 | 0.00323108 | 0.00033037 | 0.0004587 | 0.00028984 | 0.00070941 | 0.00044708 | 0 | 0 | 0 | 0 | 0 | 8 | 11 | 7 | 17 | 10.75 | 0 | 0 | 0 | 0 | 0 |
| KDM1A | Lysine-specif | 0 | 0.00024584 | 0.00024778 | 0.00028148 | 0.00037266 | 0.00043817 | 0.00033502 | 0 | 0 | 0 | 0 | 0 | 8 | 9 | 12 | 14 | 10.75 | 0 | 0 | 0 | 0 | 0 |
| LRIF1 | Ligand-depe | 0 | 1.6577E-05 | 0.0004461 | 0.00034651 | 0.00041288 | 0.00031208 | 0.00037939 | 0 | 0 | 0 | 0 | 0 | 13 | 10 | 12 | 9 | 11 | 0 | 0 | 0 | 0 | 0 |
| KIF4A | Chromosome | 0 | 2.8535E-05 | 0.00021419 | 0.00017303 | 0.00019329 | 0.00025973 | 0.00021006 | 0 | 0 | 0 | 0 | 0 | 10 | 8 | 9 | 12 | 9.75 | 0 | 0 | 0 | 0 | 0 |
| ASH2L | Set1/Ash2 hi | 0 | 4.3137E-06 | 0.00037818 | 0.00046674 | 0.00042132 | 0.00050953 | 0.00044394 | 0 | 0 | 0 | 0 | 0 | 9 | 11 | 10 | 12 | 10.5 | 0 | 0 | 0 | 0 | 0 |
| RFA1 | Replication p | 0 | 0.00055103 | 0.00034271 | 0.00064886 | 0.00038657 | 0.00043288 | 0.00045276 | 0 | 0 | 0 | 0 | 0 | 8 | 15 | 9 | 10 | 10.5 | 0 | 0 | 0 | 0 | 0 |
| ZN687 | Zinc finger pr | 0 | 2.8334E-06 | 0.00023466 | 0.00019387 | 0.00025667 | 0.00021557 | 0.00022519 | 0 | 0 | 0 | 0 | 0 | 11 | 9 | 12 | 10 | 10.5 | 0 | 0 | 0 | 0 | 0 |
| TF3C3 | General tran | 0 | 0.000289 | 0.00026805 | 0.00021053 | 0.00035836 | 0.00021068 | 0.00026191 | 0 | 0 | 0 | 0 | 0 | 9 | 7 | 12 | 7 | 8.75 | 0 | 0 | 0 | 0 | 0 |
| LRWD1 | Leucine-rich | 0 | 3.1277E-07 | 0.0002855 | 0.00032948 | 0.00028626 | 0.00032971 | 0.00030774 | 0 | 0 | 0 | 0 | 0 | 7 | 8 | 7 | 8 | 7.5 | 0 | 0 | 0 | 0 | 0 |
| ZCHC8 | Zinc finger C( | 0 | 8.4639E-05 | 0.00044789 | 0.00056534 | 0.00041166 | 0.00033945 | 0.00044109 | 0 | 0 | 0 | 0 | 0 | 12 | 15 | 11 | 9 | 11.75 | 0 | 0 | 0 | 0 | 0 |
| CBX3 | Chromobox p | 0 | 1.5726E-05 | 0.0010094 | 0.0014561 | 0.0014458 | 0.0013114 | 0.00130568 | 0 | 0 | 0 | 0 | 0 | 7 | 10 | 10 | 9 | 9 | 0 | 0 | 0 | 0 | 0 |
| RBM6 | RNA-binding | 0 | 2.4081E-05 | 0.00018799 | 0.0001661 | 0.00023561 | 0.00023745 | 0.00020679 | 0 | 0 | 0 | 0 | 0 | 8 | 7 | 10 | 10 | 8.75 | 0 | 0 | 0 | 0 | 0 |
| TOPB1 | DNA topoiso | 0 | 4.9781E-06 | 0.00019072 | 0.00015757 | 0.00017384 | 0.00014016 | 0.00016557 | 0 | 0 | 0 | 0 | 0 | 11 | 9 | 10 | 8 | 9.5 | 0 | 0 | 0 | 0 | 0 |
| WDR5 | WD repeat-c | 0 | 0.00027794 | 0.0011061 | 0.00095736 | 0.00087139 | 0.00055886 | 0.00087343 | 0 | 0 | 0 | 0 | 0 | 14 | 12 | 11 | 7 | 11 | 0 | 0 | 0 | 0 | 0 |
| BRCA1 | Breast cance | 0 | 1.5257E-05 | 0.00012748 | 0.00011442 | 0.00011362 | 0.00008588 | 0.00011035 | 0 | 0 | 0 | 0 | 0 | 9 | 8 | 8 | 6 | 7.75 | 0 | 0 | 0 | 0 | 0 |
| GPKOW | G-patch dom | 0 | 1.5336E-05 | 0.0004435 | 0.00061578 | 0.00050027 | 0.00061622 | 0.00054394 | 0 | 0 | 0 | 0 | 0 | 8 | 11 | 9 | 11 | 9.75 | 0 | 0 | 0 | 0 | 0 |
| ZN644 | Zinc finger pr | 0 | 3.1229E-05 | 0.00017897 | 0.00018072 | 0.00023926 | 0.00016076 | 0.00018993 | 0 | 0 | 0 | 0 | 0 | 9 | 9 | 12 | 8 | 9.5 | 0 | 0 | 0 | 0 | 0 |
| NSF1C | NSFL1 cofact | 0 | 0.00128619 | 0.00078452 | 0.00036009 | 0.00050057 | 0.00079276 | 0.00060949 | 0 | 0 | 0 | 0 | 0 | 11 | 5 | 7 | 11 | 8.5 | 0 | 0 | 0 | 0 | 0 |
| DIP2B | Disco-interac | 0 | 0.0002275 | 0.00013395 | 0.00020289 | 0.00011752 | 0.00018612 | 0.00016012 | 0 | 0 | 0 | 0 | 0 | 8 | 12 | 7 | 11 | 9.5 | 0 | 0 | 0 | 0 | 0 |
| ATRX | Transcription | 0 | 3.2745E-05 | 0.00010589 | 0.00007485 | 0.00011679 | 9.6304E-05 | 9.8459E-05 | 0 | 0 | 0 | 0 | 0 | 10 | 7 | 11 | 9 | 9.25 | 0 | 0 | 0 | 0 | 0 |
| ZNF24 | Zinc finger pr | 0 | 2.2238E-06 | 0.00064537 | 0.00065168 | 0.00057519 | 0.00050723 | 0.00059487 | 0 | 0 | 0 | 0 | 0 | 9 | 9 | 8 | 7 | 8.25 | 0 | 0 | 0 | 0 | 0 |
| BUB3 | Mitotic check | 0 | 0.00050156 | 0.00032181 | 0.00024372 | 0.00016133 | 0.00032519 | 0.00026301 | 0 | 0 | 0 | 0 | 0 | 4 | 3 | 2 | 4 | 3.25 | 0 | 0 | 0 | 0 | 0 |
| CND1 | Condensin co | 0 | 5.4038E-07 | 0.00011301 | 0.00011412 | 0.0001322 | 0.00013323 | 0.00012314 | 0 | 0 | 0 | 0 | 0 | 6 | 6 | 7 | 7 | 6.5 | 0 | 0 | 0 | 0 | 0 |

| | | | | | | | | | | | | | | | | | | | | | | |
|---|---|---|---|---|---|---|---|---|---|---|---|---|---|---|---|---|---|---|---|---|---|---|
| EBP2 | Probable rRN | 0 | 5.4608E-06 | 0.00068989 | 0.0008708 | 0.00086466 | 0.00095857 | 0.00084598 | 0 | 0 | 0 | 0 | 0 | 8 | 10 | 10 | 11 | 9.75 | 0 | 0 | 0 | 0 |
| RBM26 | RNA-binding | 0 | 0.00028773 | 0.00031446 | 0.00026461 | 0.00028902 | 0.00015888 | 0.00025674 | 0 | 0 | 0 | 0 | 0 | 16 | 12 | 13 | 8 | 12.25 | 0 | 0 | 0 | 0 |
| TOX4 | TOX high mo | 0 | 0.00017357 | 0.0006374 | 0.00051491 | 0.00034085 | 0.00051528 | 0.00050211 | 0 | 0 | 0 | 0 | 0 | 15 | 12 | 8 | 12 | 11.75 | 0 | 0 | 0 | 0 |
| LA | Lupus La prot | 0 | 3.0497E-05 | 0.00051742 | 0.0006531 | 0.00045395 | 0.0004575 | 0.00052049 | 0 | 0 | 0 | 0 | 0 | 8 | 10 | 7 | 7 | 8 | 0 | 0 | 0 | 0 |
| PRP6 | Pre-mRNA-pr | 0 | 0.0028443 | 0.00011217 | 0.00028317 | 0.00014059 | 0.0002267 | 0.00019066 | 0 | 0 | 0 | 0 | 0 | 4 | 10 | 5 | 8 | 6.75 | 0 | 0 | 0 | 0 |
| ST65G | STAGA comp | 0 | 0.00010244 | 0.0009561 | 0.00057927 | 0.00095865 | 0.00077292 | 0.00081674 | 0 | 0 | 0 | 0 | 0 | 15 | 9 | 15 | 12 | 12.75 | 0 | 0 | 0 | 0 |
| K319L | Dyslexia-asso | 0 | 0.00120672 | 0.00015093 | 7.6205E-05 | 0.00012611 | 0.00007626 | 0.00010738 | 0 | 0 | 0 | 0 | 0 | 6 | 3 | 5 | 3 | 4.25 | 0 | 0 | 0 | 0 |
| SNW1 | SNW domain | 0 | 0.00022091 | 0.00064002 | 0.000348 | 0.00059236 | 0.00064674 | 0.00055678 | 0 | 0 | 0 | 0 | 0 | 13 | 7 | 12 | 13 | 11.25 | 0 | 0 | 0 | 0 |
| ARI4A | AT-rich intera | 0 | 0.00068637 | 0.00020993 | 0.00012719 | 0.00018944 | 0.00010607 | 0.00015816 | 0 | 0 | 0 | 0 | 0 | 10 | 6 | 9 | 5 | 7.5 | 0 | 0 | 0 | 0 |
| P66A | Transcription | 0 | 0.00263106 | 0.00016675 | 0.00029467 | 0.00041799 | 0.00021063 | 0.00027251 | 0 | 0 | 0 | 0 | 0 | 8 | 10 | 13 | 12 | 10.75 | 0 | 0 | 0 | 0 |
| SLIK5 | SLIT and NTR | 0 | 0.01038388 | 0.00024791 | 0.00013907 | 5.5237E-05 | 0.00025051 | 0.00017318 | 0 | 0 | 0 | 0 | 0 | 9 | 5 | 2 | 9 | 6.25 | 0 | 0 | 0 | 0 |
| RBL1 | Retinoblastor | 0 | 1.0547E-06 | 0.00022237 | 0.0001996 | 0.00017342 | 0.00019974 | 0.00019878 | 0 | 0 | 0 | 0 | 0 | 9 | 8 | 7 | 8 | 8 | 0 | 0 | 0 | 0 |
| NOC4L | Nucleolar cor | 0 | 3.8622E-07 | 0.0005114 | 0.0005164 | 0.00046149 | 0.00056845 | 0.00051444 | 0 | 0 | 0 | 0 | 0 | 10 | 10 | 9 | 11 | 10 | 0 | 0 | 0 | 0 |
| LYRIC | Protein LYRIC | 0 | 5.0349E-06 | 0.00031739 | 0.00036627 | 0.00036369 | 0.0002749 | 0.00033056 | 0 | 0 | 0 | 0 | 0 | 7 | 8 | 8 | 6 | 7.25 | 0 | 0 | 0 | 0 |
| PHF2 | Lysine-specif | 0 | 0.00074142 | 0.00014446 | 0.00014587 | 7.2423E-05 | 0.00017031 | 0.00013327 | 0 | 0 | 0 | 0 | 0 | 6 | 6 | 3 | 7 | 5.5 | 0 | 0 | 0 | 0 |
| SFSWA | Splicing facto | 0 | 0.00428927 | 8.3244E-05 | 0.00019614 | 0.00011129 | 8.4119E-05 | 0.0001187 | 0 | 0 | 0 | 0 | 0 | 3 | 7 | 4 | 3 | 4.25 | 0 | 0 | 0 | 0 |
| PB1 | Protein polyb | 0 | 0.00397361 | 9.3742E-05 | 4.7329E-05 | 0.00015665 | 0.0001263 | 0.00010601 | 0 | 0 | 0 | 0 | 0 | 6 | 3 | 10 | 8 | 6.75 | 0 | 0 | 0 | 0 |
| RGAP1 | Rac GTPase-a | 0 | 2.0256E-06 | 0.00033403 | 0.00029514 | 0.00029306 | 0.00025315 | 0.00029385 | 0 | 0 | 0 | 0 | 0 | 8 | 7 | 7 | 6 | 7 | 0 | 0 | 0 | 0 |
| CAF1A | Chromatin as | 0 | 0.00015709 | 0.00022082 | 0.00019511 | 0.00033212 | 0.00027893 | 0.00025675 | 0 | 0 | 0 | 0 | 0 | 8 | 7 | 12 | 10 | 9.25 | 0 | 0 | 0 | 0 |
| DSG2 | Desmoglein-2 | 0 | 9.4938E-05 | 0.00016522 | 0.000143 | 9.4664E-05 | 0.00014311 | 0.0001365 | 0 | 0 | 0 | 0 | 0 | 7 | 6 | 4 | 6 | 5.75 | 0 | 0 | 0 | 0 |
| P121C | Nuclear enve | 0 | 0.00080459 | 0.00012883 | 6.5044E-05 | 0.0001507 | 0.00010848 | 0.00011326 | 0 | 0 | 0 | 0 | 0 | 6 | 3 | 7 | 5 | 5.25 | 0 | 0 | 0 | 0 |
| TADA1 | Transcription | 0 | 0.00299298 | 0.00094526 | 0.00047725 | 0.0014217 | 0.00087559 | 0.00092995 | 0 | 0 | 0 | 0 | 0 | 12 | 6 | 18 | 11 | 11.75 | 0 | 0 | 0 | 0 |
| NELFA | Negative elor | 0 | 7.3646E-06 | 0.0004498 | 0.00055514 | 0.00040089 | 0.00050503 | 0.00047772 | 0 | 0 | 0 | 0 | 0 | 9 | 11 | 8 | 10 | 9.5 | 0 | 0 | 0 | 0 |
| PRP16 | Pre-mRNA-sp | 0 | 2.4124E-05 | 0.00021506 | 0.00015202 | 0.00017251 | 0.00015213 | 0.00017293 | 0 | 0 | 0 | 0 | 0 | 10 | 7 | 8 | 7 | 8 | 0 | 0 | 0 | 0 |
| BMS1 | Ribosome bio | 0 | 0.00051908 | 0.00014409 | 0.00010393 | 0.00010319 | 0.0001872 | 0.0001346 | 0 | 0 | 0 | 0 | 0 | 7 | 5 | 5 | 9 | 6.5 | 0 | 0 | 0 | 0 |
| RFIP1 | Rab11 family | 0 | 8.494E-06 | 0.00020568 | 0.00018692 | 0.00014436 | 0.00018705 | 0.000181 | 0 | 0 | 0 | 0 | 0 | 10 | 9 | 7 | 9 | 8.75 | 0 | 0 | 0 | 0 |
| RREB1 | Ras-responsiv | 0 | 0.00349561 | 6.2569E-05 | 7.8976E-05 | 0.00012547 | 0.00004742 | 7.8609E-05 | 0 | 0 | 0 | 0 | 0 | 4 | 5 | 8 | 3 | 5 | 0 | 0 | 0 | 0 |
| UBP16 | Ubiquitin car | 0 | 0.0009071 | 0.00016032 | 0.00035615 | 0.00035364 | 0.0002592 | 0.00028233 | 0 | 0 | 0 | 0 | 0 | 5 | 11 | 11 | 8 | 8.75 | 0 | 0 | 0 | 0 |
| PHF12 | PHD finger pr | 0 | 6.3537E-05 | 0.00023655 | 0.00021232 | 0.00026353 | 0.00015936 | 0.00021794 | 0 | 0 | 0 | 0 | 0 | 9 | 8 | 10 | 6 | 8.25 | 0 | 0 | 0 | 0 |
| TAF8 | Transcription | 0 | 0.00062631 | 0.00076612 | 0.0011174 | 0.0005121 | 0.00086018 | 0.00081395 | 0 | 0 | 0 | 0 | 0 | 9 | 13 | 6 | 10 | 9.5 | 0 | 0 | 0 | 0 |
| CHD7 | Chromodoma | 0 | 0.01409056 | 0.00007044 | 2.6673E-05 | 1.7657E-05 | 4.4487E-05 | 3.9814E-05 | 0 | 0 | 0 | 0 | 0 | 12 | 8 | 5 | 11 | 9 | 0 | 0 | 0 | 0 |
| CCAR2 | Cell cycle and | 0 | 1.8317E-05 | 0.00011436 | 0.00014435 | 0.000172 | 0.00014445 | 0.00014379 | 0 | 0 | 0 | 0 | 0 | 4 | 5 | 6 | 5 | 5 | 0 | 0 | 0 | 0 |
| GNPI1 | Glucosamine | 0 | 3.1217E-06 | 0.00082178 | 0.00064542 | 0.00064087 | 0.00073815 | 0.00071156 | 0 | 0 | 0 | 0 | 0 | 9 | 7 | 7 | 8 | 7.75 | 0 | 0 | 0 | 0 |
| FNBP4 | Formin-bindi | 0 | 3.2606E-05 | 0.00033732 | 0.00023581 | 0.00023415 | 0.0002622 | 0.00026737 | 0 | 0 | 0 | 0 | 0 | 13 | 9 | 9 | 10 | 10.25 | 0 | 0 | 0 | 0 |
| PPM1G | Protein phos | 0 | 2.3728E-05 | 0.0004833 | 0.00048803 | 0.00033921 | 0.0003907 | 0.00042531 | 0 | 0 | 0 | 0 | 0 | 10 | 10 | 7 | 8 | 8.75 | 0 | 0 | 0 | 0 |
| BRCA2 | Breast cancer | 0 | 0.00128999 | 6.1763E-05 | 2.3388E-05 | 5.4187E-05 | 5.4611E-05 | 4.8487E-05 | 0 | 0 | 0 | 0 | 0 | 8 | 3 | 7 | 7 | 6.25 | 0 | 0 | 0 | 0 |
| NFRKB | Nuclear facto | 0 | 0.00027641 | 0.0001422 | 0.00012308 | 0.00012221 | 0.00020528 | 0.00014819 | 0 | 0 | 0 | 0 | 0 | 7 | 6 | 6 | 10 | 7.25 | 0 | 0 | 0 | 0 |
| TAF3 | Transcription | 0 | 8.0672E-05 | 0.00036927 | 0.00028683 | 0.00025633 | 0.00022963 | 0.00028552 | 0 | 0 | 0 | 0 | 0 | 13 | 10 | 9 | 8 | 10 | 0 | 0 | 0 | 0 |
| THOC1 | THO complex | 0 | 1.462E-05 | 0.00024099 | 0.00024335 | 0.00024163 | 0.0003247 | 0.00026267 | 0 | 0 | 0 | 0 | 0 | 6 | 6 | 6 | 8 | 6.5 | 0 | 0 | 0 | 0 |
| RAI14 | Ankycorbin O | 0 | 0.0004557 | 8.0781E-05 | 8.1571E-05 | 8.0996E-05 | 0.00013605 | 9.485E-05 | 0 | 0 | 0 | 0 | 0 | 3 | 3 | 3 | 5 | 3.5 | 0 | 0 | 0 | 0 |
| ZFP1 | Zinc finger pr | 0 | 0.00023832 | 0.00032418 | 0.00058923 | 0.00039005 | 0.00045862 | 0.00044052 | 0 | 0 | 0 | 0 | 0 | 5 | 9 | 6 | 7 | 6.75 | 0 | 0 | 0 | 0 |
| RHG35 | Rho GTPase-a | 0 | 0.00327028 | 0.00010562 | 0.00010666 | 3.5302E-05 | 0.00012452 | 9.3026E-05 | 0 | 0 | 0 | 0 | 0 | 6 | 6 | 2 | 7 | 5.25 | 0 | 0 | 0 | 0 |
| NOP14 | Nucleolar pro | 0 | 0.00446058 | 0.00024633 | 0.00012437 | 0.00037048 | 0.00018669 | 0.00023197 | 0 | 0 | 0 | 0 | 0 | 8 | 4 | 12 | 6 | 7.5 | 0 | 0 | 0 | 0 |
| DDX41 | Probable ATP | 0 | 0.00014685 | 0.00008485 | 0.00012852 | 8.5076E-05 | 0.00012861 | 0.00010676 | 0 | 0 | 0 | 0 | 0 | 2 | 3 | 2 | 3 | 2.5 | 0 | 0 | 0 | 0 |
| PDS5A | Sister chroma | 0 | 1.4286E-06 | 0.00013816 | 0.00017937 | 0.00015832 | 0.00015955 | 0.00015885 | 0 | 0 | 0 | 0 | 0 | 7 | 9 | 8 | 8 | 8 | 0 | 0 | 0 | 0 |
| ZN281 | Zinc finger pr | 0 | 0.00062436 | 0.00014742 | 0.00026795 | 0.0002365 | 0.00032773 | 0.0002449 | 0 | 0 | 0 | 0 | 0 | 5 | 9 | 8 | 11 | 8.25 | 0 | 0 | 0 | 0 |
| ICE1 | Little elongat | 0 | 0.00199232 | 9.3163E-05 | 3.5278E-05 | 5.8382E-05 | 8.2374E-05 | 6.7299E-05 | 0 | 0 | 0 | 0 | 0 | 8 | 3 | 5 | 7 | 5.75 | 0 | 0 | 0 | 0 |
| SF3A3 | Splicing facto | 0 | 0.00022354 | 0.00052671 | 0.00058505 | 0.00031687 | 0.0004258 | 0.00046361 | 0 | 0 | 0 | 0 | 0 | 10 | 11 | 6 | 8 | 8.75 | 0 | 0 | 0 | 0 |
| PP1B | Serine/threo | 0 | 0.00217845 | 0.0012105 | 0.00040744 | 0.00097096 | 0.0011416 | 0.00093263 | 0 | 0 | 0 | 0 | 0 | 45 | 30 | 33 | 42 | 37.5 | 0 | 0 | 0 | 0 |
| Z512B | Zinc finger pr | 0 | 0.00353499 | 0.00020708 | 0.00017924 | 5.9324E-05 | 0.00014947 | 0.00014878 | 0 | 0 | 0 | 0 | 0 | 7 | 6 | 2 | 5 | 5 | 0 | 0 | 0 | 0 |
| CD11B | Cyclin-depen | 0 | 0.00090217 | 9.9579E-05 | 0.00023462 | 0.00016641 | 0.00020125 | 0.00017546 | 0 | 0 | 0 | 0 | 0 | 3 | 7 | 5 | 6 | 5.25 | 0 | 0 | 0 | 0 |
| KANL3 | KAT8 regulat | 0 | 3.5031E-05 | 0.00026272 | 0.00020633 | 0.00032195 | 0.00029497 | 0.00027149 | 0 | 0 | 0 | 0 | 0 | 9 | 7 | 11 | 10 | 9.25 | 0 | 0 | 0 | 0 |
| CCDC8 | Coiled-coil dc | 0 | 1.796E-05 | 0.00049049 | 0.00039623 | 0.00039344 | 0.00054521 | 0.00045634 | 0 | 0 | 0 | 0 | 0 | 10 | 8 | 8 | 11 | 9.25 | 0 | 0 | 0 | 0 |
| ILKAP | Integrin-linke | 0 | 6.6003E-05 | 0.0004039 | 0.00061178 | 0.00047248 | 0.00040815 | 0.00047408 | 0 | 0 | 0 | 0 | 0 | 6 | 9 | 7 | 6 | 7 | 0 | 0 | 0 | 0 |
| CTCF | Transcription | 0 | 0.0008557 | 0.00010889 | 0.00018326 | 0.00025476 | 0.00022007 | 0.00019175 | 0 | 0 | 0 | 0 | 0 | 3 | 5 | 7 | 6 | 5.25 | 0 | 0 | 0 | 0 |
| PIAS2 | E3 SUMO-pro | 0 | 8.2388E-05 | 0.00046743 | 0.00030036 | 0.00051128 | 0.0004294 | 0.00042712 | 0 | 0 | 0 | 0 | 0 | 11 | 7 | 12 | 10 | 10 | 0 | 0 | 0 | 0 |
| SET | Protein SET O | 0 | 0.01428712 | 0.00054597 | 0.00036754 | 0.00036495 | 0.0011034 | 0.00059547 | 0 | 0 | 0 | 0 | 0 | 6 | 4 | 4 | 12 | 6.5 | 0 | 0 | 0 | 0 |
| ITSN1 | Intersectin-1 | 0 | 2.1423E-06 | 7.6666E-05 | 6.1933E-05 | 6.1496E-05 | 6.1977E-05 | 6.5518E-05 | 0 | 0 | 0 | 0 | 0 | 5 | 4 | 4 | 4 | 4.25 | 0 | 0 | 0 | 0 |
| THOC5 | THO complex | 0 | 0.0004367 | 0.00019318 | 0.00023408 | 0.00015496 | 0.00011713 | 0.00017484 | 0 | 0 | 0 | 0 | 0 | 5 | 6 | 4 | 3 | 4.5 | 0 | 0 | 0 | 0 |
| DDHD1 | Phospholipas | 0 | 0.00189823 | 0.00020524 | 0.00011843 | 0.00011759 | 8.8885E-05 | 0.00013254 | 0 | 0 | 0 | 0 | 0 | 7 | 4 | 4 | 3 | 4.5 | 0 | 0 | 0 | 0 |
| KTN1 | Kinectin OS=H | 0 | 0.00136614 | 5.8338E-05 | 0.00011782 | 7.7992E-05 | 5.8951E-05 | 7.8275E-05 | 0 | 0 | 0 | 0 | 0 | 3 | 6 | 4 | 3 | 4 | 0 | 0 | 0 | 0 |
| ZN148 | Zinc finger pr | 0 | 5.2701E-05 | 0.00023264 | 0.0003356 | 0.00023326 | 0.00023509 | 0.00025915 | 0 | 0 | 0 | 0 | 0 | 7 | 10 | 7 | 7 | 7.75 | 0 | 0 | 0 | 0 |
| ORC2 | Origin recogr | 0 | 0.00025717 | 0.00022867 | 0.00032327 | 0.00032099 | 0.00018486 | 0.00026445 | 0 | 0 | 0 | 0 | 0 | 5 | 7 | 7 | 4 | 5.75 | 0 | 0 | 0 | 0 |
| SP20H | Transcription | 0 | 0.00032932 | 0.00023712 | 0.00020524 | 0.00027172 | 0.00037654 | 0.00027266 | 0 | 0 | 0 | 0 | 0 | 7 | 6 | 8 | 11 | 8 | 0 | 0 | 0 | 0 |
| DMAP1 | DNA methylt | 0 | 0.01849176 | 0.00033904 | 0.00011412 | 0.00022663 | 0.000571 | 0.0003127 | 0 | 0 | 0 | 0 | 0 | 6 | 2 | 4 | 10 | 5.5 | 0 | 0 | 0 | 0 |
| DDX23 | Probable ATP | 0 | 0.00358613 | 0.00019309 | 0.00022747 | 0.00022587 | 6.5038E-05 | 0.00017787 | 0 | 0 | 0 | 0 | 0 | 6 | 7 | 7 | 2 | 5.5 | 0 | 0 | 0 | 0 |
| GCFC2 | GC-rich sequ | 0 | 0.00015838 | 0.00016894 | 0.00020471 | 0.00013551 | 0.000239 | 0.00018704 | 0 | 0 | 0 | 0 | 0 | 5 | 6 | 4 | 7 | 5.5 | 0 | 0 | 0 | 0 |
| CRKL | Crk-like prote | 0 | 0.00366098 | 0.00043545 | 0.00079148 | 0.00026197 | 0.00061604 | 0.00052624 | 0 | 0 | 0 | 0 | 0 | 5 | 9 | 3 | 7 | 6 | 0 | 0 | 0 | 0 |
| RSBNL | Round sperm | 0 | 0.00129593 | 6.2384E-05 | 9.4491E-05 | 0.00006255 | 0.00012608 | 8.6376E-05 | 0 | 0 | 0 | 0 | 0 | 2 | 3 | 2 | 4 | 2.75 | 0 | 0 | 0 | 0 |
| KIF15 | Kinesin-like p | 0 | 0.00080376 | 0.00011407 | 0.00023037 | 0.0001525 | 0.00013448 | 0.00015786 | 0 | 0 | 0 | 0 | 0 | 6 | 12 | 8 | 7 | 8.25 | 0 | 0 | 0 | 0 |
| CLMN | Calmin OS=H | 0 | 0.0009782 | 0.00026336 | 0.00029253 | 0.00013203 | 0.00018629 | 0.00021855 | 0 | 0 | 0 | 0 | 0 | 10 | 11 | 5 | 7 | 8.25 | 0 | 0 | 0 | 0 |
| ESF1 | ESF1 homolo | 0 | 0.00031097 | 0.00012403 | 9.3936E-05 | 6.2183E-05 | 9.4003E-05 | 9.3538E-05 | 0 | 0 | 0 | 0 | 0 | 4 | 3 | 2 | 3 | 3 | 0 | 0 | 0 | 0 |
| IMA3 | Importin sub | 0 | 0.00373742 | 0.00015195 | 0.00040916 | 0.00020314 | 0.00025591 | 0.00025504 | 0 | 0 | 0 | 0 | 0 | 3 | 8 | 4 | 5 | 5 | 0 | 0 | 0 | 0 |
| PCGF6 | Polycomb gro | 0 | 0.00094211 | 0.00045237 | 0.00053293 | 0.00022679 | 0.00053331 | 0.00043635 | 0 | 0 | 0 | 0 | 0 | 6 | 7 | 3 | 7 | 5.75 | 0 | 0 | 0 | 0 |
| WDR70 | WD repeat-co | 0 | 2.9155E-05 | 0.00028244 | 0.00020372 | 0.00020228 | 0.00020387 | 0.00022308 | 0 | 0 | 0 | 0 | 0 | 7 | 5 | 5 | 5 | 5.5 | 0 | 0 | 0 | 0 |
| HAX1 | HCLS1-associ | 0 | 2.0274E-14 | 0.00037833 | 0.00038203 | 0.00037934 | 0.0003823 | 0.0003805 | 0 | 0 | 0 | 0 | 0 | 4 | 4 | 4 | 4 | 4 | 0 | 0 | 0 | 0 |
| SYNC | Asparagine--t | 0 | 2.7013E-05 | 0.00038523 | 0.000389 | 0.00038626 | 0.00053526 | 0.00042394 | 0 | 0 | 0 | 0 | 0 | 8 | 8 | 8 | 11 | 8.75 | 0 | 0 | 0 | 0 |
| CLCC1 | Chloride char | 0 | 0.00142215 | 0.00023946 | 0.00033852 | 0.00019208 | 0.00043555 | 0.0003014 | 0 | 0 | 0 | 0 | 0 | 5 | 7 | 4 | 9 | 6.25 | 0 | 0 | 0 | 0 |
| WRIP1 | ATPase WRN | 0 | 1.4839E-05 | 0.00039682 | 0.0004007 | 0.00027851 | 0.00036089 | 0.00035923 | 0 | 0 | 0 | 0 | 0 | 10 | 10 | 7 | 9 | 9 | 0 | 0 | 0 | 0 |
| RADX | RPA-related p | 0 | 2.2699E-05 | 9.2591E-05 | 0.00012466 | 9.2837E-05 | 0.00012475 | 0.00010871 | 0 | 0 | 0 | 0 | 0 | 3 | 4 | 3 | 4 | 3.5 | 0 | 0 | 0 | 0 |
| ZFR | Zinc finger RN | 0 | 0.00014442 | 0.00017199 | 0.00012405 | 0.00012318 | 9.9313E-05 | 0.00012963 | 0 | 0 | 0 | 0 | 0 | 7 | 5 | 5 | 4 | 5.25 | 0 | 0 | 0 | 0 |
| ARI1A | AT-rich intera | 0 | 2.7627E-05 | 0.00008084 | 6.9969E-05 | 8.1055E-05 | 0.00010503 | 8.4224E-05 | 0 | 0 | 0 | 0 | 0 | 7 | 6 | 7 | 9 | 7.25 | 0 | 0 | 0 | 0 |
| BAP1 | Ubiquitin car | 0 | 2.0308E-05 | 0.00036198 | 0.00029242 | 0.00036294 | 0.00025605 | 0.00031835 | 0 | 0 | 0 | 0 | 0 | 10 | 8 | 10 | 7 | 8.75 | 0 | 0 | 0 | 0 |
| CHD6 | Chromodoma | 0 | 0.00043697 | 3.8878E-05 | 4.9073E-05 | 3.8981E-05 | 6.8751E-05 | 4.8921E-05 | 0 | 0 | 0 | 0 | 0 | 7 | 8 | 5 | 10 | 7.5 | 0 | 0 | 0 | 0 |
| Z518B | Zinc finger pr | 0 | 0.00026545 | 0.00012285 | 0.00012405 | 0.00014781 | 7.4485E-05 | 0.0001173 | 0 | 0 | 0 | 0 | 0 | 5 | 5 | 6 | 3 | 4.75 | 0 | 0 | 0 | 0 |
| INT3 | Integrator co | 0 | 0.00128498 | 0.0001771 | 0.00025548 | 0.00012684 | 0.00012783 | 0.00017181 | 0 | 0 | 0 | 0 | 0 | 7 | 10 | 5 | 5 | 6.75 | 0 | 0 | 0 | 0 |
| CPSF7 | Cleavage and | 0 | 5.0345E-06 | 0.00039218 | 0.00045259 | 0.0004494 | 0.00033969 | 0.00040847 | 0 | 0 | 0 | 0 | 0 | 7 | 8 | 8 | 6 | 7.25 | 0 | 0 | 0 | 0 |
| RN169 | E3 ubiquitin-p | 0 | 0.00013485 | 0.00022363 | 0.00018818 | 0.00029897 | 0.00018832 | 0.00022478 | 0 | 0 | 0 | 0 | 0 | 6 | 5 | 8 | 5 | 6 | 0 | 0 | 0 | 0 |
| MACOI | Macoilin OS= | 0 | 8.3629E-06 | 0.00027819 | 0.00032104 | 0.00023908 | 0.00024095 | 0.00026982 | 0 | 0 | 0 | 0 | 0 | 7 | 8 | 6 | 6 | 6.75 | 0 | 0 | 0 | 0 |
| ERCC1 | DNA excision | 0 | 0.00033022 | 0.0008885 | 0.00062803 | 0.00071269 | 0.00044892 | 0.00066954 | 0 | 0 | 0 | 0 | 0 | 10 | 7 | 8 | 5 | 7.5 | 0 | 0 | 0 | 0 |
| NPAT | Protein NPAT | 0 | 0.00089455 | 0.00016643 | 9.3365E-05 | 9.2707E-05 | 9.3432E-05 | 0.00011148 | 0 | 0 | 0 | 0 | 0 | 9 | 5 | 5 | 5 | 6 | 0 | 0 | 0 | 0 |
| CDYL | Chromodoma | 0 | 0.0004364 | 0.00039715 | 0.00031192 | 0.00022123 | 0.00044591 | 0.00034405 | 0 | 0 | 0 | 0 | 0 | 9 | 7 | 5 | 10 | 7.75 | 0 | 0 | 0 | 0 |
| ERCC5 | DNA repair p | 0 | 0.00060687 | 0.000178 | 0.00015727 | 0.00020078 | 8.9935E-05 | 0.0001565 | 0 | 0 | 0 | 0 | 0 | 8 | 7 | 9 | 4 | 7 | 0 | 0 | 0 | 0 |
| RPB2 | DNA-directed | 0 | 0.00049087 | 8.9909E-05 | 4.5394E-05 | 6.7612E-05 | 9.0854E-05 | 7.3442E-05 | 0 | 0 | 0 | 0 | 0 | 4 | 2 | 3 | 4 | 3.25 | 0 | 0 | 0 | 0 |
| CCNT1 | Cyclin-T1 OS= | 0 | 0.00248202 | 0.00018174 | 0.00022022 | 7.2889E-05 | 0.00022038 | 0.00017381 | 0 | 0 | 0 | 0 | 0 | 5 | 6 | 2 | 6 | 4.75 | 0 | 0 | 0 | 0 |
| SAFB1 | Scaffold attac | 0 | 0.00040744 | 0.00017304 | 0.00017473 | 0.0001735 | 8.7428E-05 | 0.00015217 | 0 | 0 | 0 | 0 | 0 | 16 | 17 | 15 | 12 | 15 | 0 | 0 | 0 | 0 |
| CMTR1 | Cap-specific r | 0 | 0.00011574 | 0.00012641 | 0.00012765 | 0.00012675 | 0.00019161 | 0.00014311 | 0 | 0 | 0 | 0 | 0 | 4 | 4 | 4 | 6 | 4.5 | 0 | 0 | 0 | 0 |
| NSD3 | Histone-lysin | 0 | 0.00214286 | 0.00014691 | 9.2716E-05 | 5.5237E-05 | 9.2782E-05 | 9.6911E-05 | 0 | 0 | 0 | 0 | 0 | 11 | 7 | 4 | 6 | 7 | 0 | 0 | 0 | 0 |
| UBP47 | Ubiquitin car | 0 | 3.4353E-05 | 0.00011515 | 0.00011628 | 7.6971E-05 | 9.6966E-05 | 0.00010134 | 0 | 0 | 0 | 0 | 0 | 6 | 6 | 4 | 5 | 5.25 | 0 | 0 | 0 | 0 |
| PI4KA | Phosphatidyl | 0 | 0.00040347 | 0.00006277 | 0.00003803 | 7.5524E-05 | 5.0743E-05 | 5.6767E-05 | 0 | 0 | 0 | 0 | 0 | 5 | 3 | 6 | 4 | 4.5 | 0 | 0 | 0 | 0 |
| SMCE1 | SWI/SNF-rela | 0 | 0.00093732 | 0.00044944 | 0.00045383 | 0.00045063 | 0.00019464 | 0.00038714 | 0 | 0 | 0 | 0 | 0 | 7 | 7 | 7 | 3 | 6 | 0 | 0 | 0 | 0 |
| NUP58 | Nucleoporin | 0 | 0.0043173 | 0.00013216 | 0.00013345 | 0.00017669 | 0.00031162 | 0.00018848 | 0 | 0 | 0 | 0 | 0 | 3 | 3 | 4 | 7 | 4.25 | 0 | 0 | 0 | 0 |
| BACH1 | Transcription | 0 | 0.00138499 | 0.00010756 | 0.00021723 | 0.00010785 | 0.00018115 | 0.00015345 | 0 | 0 | 0 | 0 | 0 | 3 | 6 | 3 | 5 | 4.25 | 0 | 0 | 0 | 0 |
| HMGX4 | HMG domain | 0 | 0.00031887 | 0.00021954 | 0.0003547 | 0.00044024 | 0.00039932 | 0.00035345 | 0 | 0 | 0 | 0 | 0 | 5 | 8 | 10 | 9 | 8 | 0 | 0 | 0 | 0 |
| MK67I | MKI67 FHA d | 0 | 0.0003469 | 0.00090063 | 0.00072755 | 0.00045151 | 0.00063706 | 0.00067919 | 0 | 0 | 0 | 0 | 0 | 10 | 8 | 5 | 7 | 7.5 | 0 | 0 | 0 | 0 |
| POP1 | Ribonuclease | 0 | 0.01203228 | 0.00007731 | 0.00013011 | 0.00025839 | 0.00010416 | 0.00014249 | 0 | 0 | 0 | 0 | 0 | 3 | 5 | 10 | 4 | 5.5 | 0 | 0 | 0 | 0 |
| NUP85 | Nuclear pore | 0 | 0.00640301 | 0.0001609 | 0.00016248 | 8.0667E-05 | 0.00028454 | 0.00017215 | 0 | 0 | 0 | 0 | 0 | 4 | 4 | 2 | 7 | 4.25 | 0 | 0 | 0 | 0 |
| PALD | Paladin OS=H | 0 | 0.00135349 | 0.00021579 | 0.00015565 | 0.00012364 | 9.3454E-05 | 0.00014713 | 0 | 0 | 0 | 0 | 0 | 7 | 5 | 4 | 3 | 4.75 | 0 | 0 | 0 | 0 |
| NPM3 | Nucleoplasm | 0 | 0.0003798 | 0.0014825 | 0.0007485 | 0.0010405 | 0.0010486 | 0.00108003 | 0 | 0 | 0 | 0 | 0 | 10 | 5 | 7 | 7 | 7.25 | 0 | 0 | 0 | 0 |
| DREB | Drebrin OS=H | 0 | 0.00136103 | 0.0002033 | 0.00016423 | 0.00028538 | 0.00012326 | 0.00019404 | 0 | 0 | 0 | 0 | 0 | 5 | 4 | 7 | 3 | 4.75 | 0 | 0 | 0 | 0 |
| FHOD1 | FH1/FH2 don | 0 | 3.5832E-05 | 0.00015869 | 0.00011446 | 0.00018185 | 0.00016036 | 0.00015384 | 0 | 0 | 0 | 0 | 0 | 7 | 5 | 8 | 7 | 6.75 | 0 | 0 | 0 | 0 |
| HDAC3 | Histone deac | 0 | 0.00484201 | 0.00030828 | 0.00056032 | 0.0003091 | 0.00018691 | 0.00034115 | 0 | 0 | 0 | 0 | 0 | 6 | 10 | 7 | 5 | 7 | 0 | 0 | 0 | 0 |
| CDK9 | Cyclin-depen | 0 | 0.00243478 | 0.00035468 | 0.00035815 | 0.00014225 | 0.00021505 | 0.00026753 | 0 | 0 | 0 | 0 | 0 | 8 | 6 | 6 | 4 | 6 | 0 | 0 | 0 | 0 |
| GWL | Serine/threo | 0 | 0.01752816 | 6.0042E-05 | 0.0002122 | 6.0202E-05 | 0.00015168 | 0.00012103 | 0 | 0 | 0 | 0 | 0 | 2 | 7 | 2 | 5 | 4 | 0 | 0 | 0 | 0 |
| ZN512 | Zinc finger pr | 0 | 6.629E-05 | 0.00018616 | 0.00023498 | 0.00018666 | 0.00014109 | 0.00018722 | 0 | 0 | 0 | 0 | 0 | 4 | 5 | 4 | 3 | 4 | 0 | 0 | 0 | 0 |
| PSIP1 | PC4 and SFRS | 0 | 0.00156368 | 0.00029874 | 0.00035193 | 0.00014977 | 0.00020125 | 0.00025042 | 0 | 0 | 0 | 0 | 0 | 6 | 7 | 3 | 4 | 5 | 0 | 0 | 0 | 0 |
| ADDG | Gamma-addu | 0 | 0.00040989 | 0.00022426 | 0.00011323 | 0.00022486 | 0.00022662 | 0.00019724 | 0 | 0 | 0 | 0 | 0 | 6 | 3 | 6 | 6 | 5.25 | 0 | 0 | 0 | 0 |
| ZBT21 | Zinc finger ar | 0 | 0.00060568 | 0.00012377 | 0.00019997 | 0.00017374 | 0.00010006 | 0.00014939 | 0 | 0 | 0 | 0 | 0 | 5 | 8 | 7 | 4 | 6 | 0 | 0 | 0 | 0 |
| GELS | Gelsolin OS=H | 0 | 0.00376984 | 0.00013498 | 0.0001363 | 0.00016917 | 0.00030689 | 0.00018684 | 0 | 0 | 0 | 0 | 0 | 4 | 4 | 5 | 9 | 5.5 | 0 | 0 | 0 | 0 |
| AP1B1 | AP-1 complex | 0 | 5.4779E-06 | 0.00016684 | 0.00016847 | 0.00019516 | 0.00014049 | 0.00016774 | 0 | 0 | 0 | 0 | 0 | 9 | 9 | 9 | 8 | 8.75 | 0 | 0 | 0 | 0 |
| RU2A | U2 small nuc | 0 | 0.00011471 | 0.00051742 | 0.00041798 | 0.00031128 | 0.00052286 | 0.00044239 | 0 | 0 | 0 | 0 | 0 | 5 | 4 | 3 | 5 | 4.25 | 0 | 0 | 0 | 0 |
| PLIN3 | Perilipin-3 OS | 0 | 6.8437E-05 | 0.00048642 | 0.00067537 | 0.00048772 | 0.00043009 | 0.0005199 | 0 | 0 | 0 | 0 | 0 | 8 | 11 | 8 | 7 | 8.5 | 0 | 0 | 0 | 0 |
| E41L5 | Band 4.1-like | 0 | 0.00276887 | 0.00018 | 0.00014541 | 0.00025268 | 0.00036379 | 0.00023547 | 0 | 0 | 0 | 0 | 0 | 5 | 4 | 7 | 10 | 6.5 | 0 | 0 | 0 | 0 |
| RRP12 | RRP12-like pr | 0 | 4.6318E-06 | 8.1383E-05 | 8.2179E-05 | 0.000102 | 0.0001028 | 9.2091E-05 | 0 | 0 | 0 | 0 | 0 | 4 | 4 | 5 | 5 | 4.5 | 0 | 0 | 0 | 0 |
| RBM5 | RNA-binding | 0 | 0.00086067 | 9.7135E-05 | 0.00016348 | 0.00022725 | 0.00016359 | 0.00016286 | 0 | 0 | 0 | 0 | 0 | 3 | 5 | 7 | 5 | 5 | 0 | 0 | 0 | 0 |
| TAF12 | Transcription | 0 | 0.00149914 | 0.00049171 | 0.00066202 | 0.0008217 | 0.0011594 | 0.00078371 | 0 | 0 | 0 | 0 | 0 | 3 | 4 | 5 | 7 | 4.75 | 0 | 0 | 0 | 0 |
| ENAH | Protein enab | 0 | 0.00041904 | 0.0001786 | 0.00027052 | 0.00031339 | 0.00018048 | 0.00023575 | 0 | 0 | 0 | 0 | 0 | 4 | 6 | 7 | 4 | 5.25 | 0 | 0 | 0 | 0 |
| SF3A2 | Splicing facto | 0 | 1.0258E-05 | 0.00056872 | 0.00040199 | 0.00045618 | 0.00045975 | 0.00047166 | 0 | 0 | 0 | 0 | 0 | 10 | 7 | 8 | 8 | 8.25 | 0 | 0 | 0 | 0 |
| RPR1B | Regulation of | 0 | 4.8468E-05 | 0.00064757 | 0.00040869 | 0.00056813 | 0.00049078 | 0.00052879 | 0 | 0 | 0 | 0 | 0 | 8 | 5 | 7 | 6 | 6.5 | 0 | 0 | 0 | 0 |
| ARI3B | AT-rich intera | 0 | 5.2182E-05 | 0.00028223 | 0.00018999 | 0.00023582 | 0.00019013 | 0.00022454 | 0 | 0 | 0 | 0 | 0 | 6 | 4 | 5 | 4 | 4.75 | 0 | 0 | 0 | 0 |
| CBX8 | Chromobox p | 0 | 0.00013485 | 0.00040702 | 0.0003425 | 0.00054414 | 0.00034275 | 0.0004091 | 0 | 0 | 0 | 0 | 0 | 6 | 5 | 8 | 5 | 6 | 0 | 0 | 0 | 0 |
| DNLI3 | DNA ligase 3 | 0 | 0.00055015 | 0.00010461 | 0.00021127 | 0.00015734 | 0.00013214 | 0.00015134 | 0 | 0 | 0 | 0 | 0 | 4 | 8 | 6 | 5 | 5.75 | 0 | 0 | 0 | 0 |
| KDM6A | Lysine-specif | 0 | 9.139E-07 | 9.4177E-05 | 0.00011412 | 9.4428E-05 | 9.5166E-05 | 9.9473E-05 | 0 | 0 | 0 | 0 | 0 | 5 | 6 | 5 | 5 | 5.25 | 0 | 0 | 0 | 0 |

| | | | | | | | | | | | | | | | | | | | | | | | |
|---|---|---|---|---|---|---|---|---|---|---|---|---|---|---|---|---|---|---|---|---|---|---|---|
| QSER1 | Glutamine ar | 0 | 2.334E-07 | 0.00010647 | 0.00010751 | 0.000122 | 0.00012295 | 0.00011473 | 0 | 0 | 0 | 0 | 0 | 7 | 7 | 8 | 8 | 7.5 | 0 | 0 | 0 | 0 | 0 |
| SMRC2 | SWI/SNF com | 0 | 0.00013484 | 0.00015216 | 8.7797E-05 | 0.00015256 | 0.00013179 | 0.00013108 | 0 | 0 | 0 | 0 | 0 | 14 | 11 | 13 | 14 | 13 | 0 | 0 | 0 | 0 | 0 |
| ARPC4 | Actin-related | 0 | 9.3821E-05 | 0.00047122 | 0.00031722 | 0.00031498 | 0.00031745 | 0.00035522 | 0 | 0 | 0 | 0 | 0 | 3 | 2 | 2 | 2 | 2.25 | 0 | 0 | 0 | 0 | 0 |
| VATA | V-type proto | 0 | 0.00028559 | 0.00012831 | 0.00021594 | 0.0002573 | 0.00021609 | 0.00020441 | 0 | 0 | 0 | 0 | 0 | 3 | 5 | 6 | 5 | 4.75 | 0 | 0 | 0 | 0 | 0 |
| NHP2 | H/ACA ribonu | 0 | 5.9906E-06 | 0.00068989 | 0.00069664 | 0.0005188 | 0.00069714 | 0.00065062 | 0 | 0 | 0 | 0 | 0 | 4 | 4 | 3 | 4 | 3.75 | 0 | 0 | 0 | 0 | 0 |
| SETX | Probable heli | 0 | 0.00243454 | 4.9287E-05 | 1.9908E-05 | 2.9651E-05 | 4.9805E-05 | 3.7163E-05 | 0 | 0 | 0 | 0 | 0 | 5 | 2 | 3 | 5 | 3.75 | 0 | 0 | 0 | 0 | 0 |
| FUBP1 | Far upstream | 0 | 3.5497E-05 | 0.00028683 | 0.00028964 | 0.00032868 | 0.00020703 | 0.00027805 | 0 | 0 | 0 | 0 | 0 | 8 | 7 | 9 | 5 | 7.25 | 0 | 0 | 0 | 0 | 0 |
| NFL | Neurofilamer | 0 | 0.00150306 | 0.00034018 | 0.00014722 | 0.00019491 | 0.00029465 | 0.00024424 | 0 | 0 | 0 | 0 | 0 | 8 | 3 | 6 | 7 | 6 | 0 | 0 | 0 | 0 | 0 |
| FANCJ | Fanconi aner | 0 | 0.00129196 | 0.00010564 | 4.2669E-05 | 8.4736E-05 | 0.00010675 | 8.4949E-05 | 0 | 0 | 0 | 0 | 0 | 5 | 2 | 4 | 5 | 4 | 0 | 0 | 0 | 0 | 0 |
| RFX5 | DNA-binding | 0 | 0.00054426 | 0.00017135 | 0.00025954 | 0.00012886 | 0.00017315 | 0.00018323 | 0 | 0 | 0 | 0 | 0 | 4 | 6 | 3 | 4 | 4.25 | 0 | 0 | 0 | 0 | 0 |
| PA24A | Cytosolic phc | 0 | 0.00195272 | 0.00010569 | 0.00028461 | 0.00017663 | 0.00021361 | 0.00019514 | 0 | 0 | 0 | 0 | 0 | 3 | 8 | 5 | 6 | 5.5 | 0 | 0 | 0 | 0 | 0 |
| BAP18 | Chromatin cc | 0 | 0.00083007 | 0.00076711 | 0.00077461 | 0.0010768 | 0.0004651 | 0.00077091 | 0 | 0 | 0 | 0 | 0 | 5 | 5 | 7 | 3 | 5 | 0 | 0 | 0 | 0 | 0 |
| SPAT5 | Spermatoger | 0 | 0.01433424 | 5.9101E-05 | 8.9518E-05 | 5.9258E-05 | 0.00017916 | 9.6759E-05 | 0 | 0 | 0 | 0 | 0 | 2 | 3 | 2 | 6 | 3.25 | 0 | 0 | 0 | 0 | 0 |
| SFR19 | Splicing facto | 0 | 3.4745E-07 | 0.00012068 | 0.00012186 | 0.000121 | 0.00014227 | 0.00012645 | 0 | 0 | 0 | 0 | 0 | 6 | 6 | 6 | 7 | 6.25 | 0 | 0 | 0 | 0 | 0 |
| RPA1 | DNA-directec | 0 | 1.0822E-05 | 6.1368E-05 | 4.6476E-05 | 4.6149E-05 | 0.00004651 | 5.0126E-05 | 0 | 0 | 0 | 0 | 0 | 4 | 3 | 3 | 3 | 3.25 | 0 | 0 | 0 | 0 | 0 |
| PINX1 | PIN2/TERF1-i | 0 | 2.9167E-05 | 0.00056317 | 0.0004062 | 0.00040333 | 0.00040649 | 0.0004448 | 0 | 0 | 0 | 0 | 0 | 7 | 5 | 5 | 5 | 5.5 | 0 | 0 | 0 | 0 | 0 |
| INO80 | DNA helicase | 0 | 0.00023348 | 0.00011871 | 8.5625E-05 | 0.00011903 | 6.8549E-05 | 9.7979E-05 | 0 | 0 | 0 | 0 | 0 | 7 | 5 | 7 | 4 | 5.75 | 0 | 0 | 0 | 0 | 0 |
| UBP5 | Ubiquitin car | 0 | 9.4115E-07 | 0.00015378 | 0.00015528 | 0.00015419 | 0.00018647 | 0.00016243 | 0 | 0 | 0 | 0 | 0 | 5 | 5 | 5 | 6 | 5.25 | 0 | 0 | 0 | 0 | 0 |
| CD2B2 | CD2 antigen | 0 | 0.000116 | 0.00046431 | 0.00031257 | 0.00046555 | 0.00031279 | 0.00038881 | 0 | 0 | 0 | 0 | 0 | 6 | 4 | 6 | 4 | 5 | 0 | 0 | 0 | 0 | 0 |
| MAML1 | Mastermind- | 0 | 0.00054436 | 0.00010389 | 0.00015736 | 7.8126E-05 | 0.00010498 | 0.00011109 | 0 | 0 | 0 | 0 | 0 | 4 | 6 | 3 | 4 | 4.25 | 0 | 0 | 0 | 0 | 0 |
| SNF5 | SWI/SNF-rela | 0 | 0.00026782 | 0.00061687 | 0.00041527 | 0.00034362 | 0.00041557 | 0.00044783 | 0 | 0 | 0 | 0 | 0 | 9 | 6 | 5 | 6 | 6.5 | 0 | 0 | 0 | 0 | 0 |
| BRD4 | Bromodomai | 0 | 1.4329E-05 | 5.8124E-05 | 7.8257E-05 | 5.8279E-05 | 5.8735E-05 | 6.3349E-05 | 0 | 0 | 0 | 0 | 0 | 3 | 4 | 3 | 3 | 3.25 | 0 | 0 | 0 | 0 | 0 |
| CSN3 | COP9 signalo | 0 | 0.00525394 | 0.00018715 | 0.00012599 | 0.00031275 | 0.00012608 | 0.00018799 | 0 | 0 | 0 | 0 | 0 | 3 | 2 | 5 | 2 | 3 | 0 | 0 | 0 | 0 | 0 |
| MEP50 | Methylosom | 0 | 0.00030852 | 0.00023148 | 0.00015583 | 0.00030946 | 0.00023391 | 0.00023267 | 0 | 0 | 0 | 0 | 0 | 3 | 2 | 4 | 3 | 3 | 0 | 0 | 0 | 0 | 0 |
| TF3C4 | General trans | 0 | 0.00042583 | 0.00016051 | 0.0001945 | 9.6565E-05 | 0.00019464 | 0.00016155 | 0 | 0 | 0 | 0 | 0 | 5 | 6 | 3 | 6 | 5 | 0 | 0 | 0 | 0 | 0 |
| FEN1 | Flap endonuc | 0 | 0.00010608 | 0.00027777 | 0.00042073 | 0.00041777 | 0.00049121 | 0.00040187 | 0 | 0 | 0 | 0 | 0 | 4 | 6 | 6 | 7 | 5.75 | 0 | 0 | 0 | 0 | 0 |
| TDIF2 | Deoxynucleo | 0 | 0.00010294 | 0.00020943 | 0.00024673 | 0.00013999 | 0.00021163 | 0.00020195 | 0 | 0 | 0 | 0 | 0 | 6 | 7 | 4 | 6 | 5.75 | 0 | 0 | 0 | 0 | 0 |
| ZHX3 | Zinc fingers a | 0 | 0.0015211 | 0.00013801 | 0.00011149 | 0.00011071 | 0.00022314 | 0.00014584 | 0 | 0 | 0 | 0 | 0 | 5 | 4 | 4 | 8 | 5.25 | 0 | 0 | 0 | 0 | 0 |
| BCORL | BCL-6 corepr | 0 | 0.00023594 | 6.1691E-05 | 7.7868E-05 | 4.6392E-05 | 4.6754E-05 | 5.8176E-05 | 0 | 0 | 0 | 0 | 0 | 4 | 5 | 3 | 3 | 3.75 | 0 | 0 | 0 | 0 | 0 |
| RL34 | 60S ribosoma | 0 | 0.00012922 | 0.00067663 | 0.0004555 | 0.00045229 | 0.00068373 | 0.00056704 | 0 | 0 | 0 | 0 | 0 | 3 | 2 | 2 | 3 | 2.5 | 0 | 0 | 0 | 0 | 0 |
| PIAS1 | E3 SUMO-pro | 0 | 1.5305E-06 | 0.00020268 | 0.00020466 | 0.00016257 | 0.00020481 | 0.00019368 | 0 | 0 | 0 | 0 | 0 | 6 | 6 | 5 | 7 | 6 | 0 | 0 | 0 | 0 | 0 |
| SALL2 | Sal-like prote | 0 | 2.0554E-14 | 0.00010482 | 0.00010585 | 0.0001051 | 0.00010592 | 0.00010542 | 0 | 0 | 0 | 0 | 0 | 4 | 4 | 4 | 4 | 4 | 0 | 0 | 0 | 0 | 0 |
| TADA3 | Transcription | 0 | 2.9682E-05 | 0.00036651 | 0.00030841 | 0.00036748 | 0.0002469 | 0.00032233 | 0 | 0 | 0 | 0 | 0 | 6 | 5 | 6 | 4 | 5.25 | 0 | 0 | 0 | 0 | 0 |
| TAF10 | Transcription | 0 | 0.00079319 | 0.00084733 | 0.00061116 | 0.00072822 | 0.00036696 | 0.00063842 | 0 | 0 | 0 | 0 | 0 | 7 | 5 | 6 | 3 | 5.25 | 0 | 0 | 0 | 0 | 0 |
| SUGP1 | SURP and G-p | 0 | 3.1587E-06 | 0.00016365 | 0.00016525 | 0.00016408 | 0.00020671 | 0.00017492 | 0 | 0 | 0 | 0 | 0 | 4 | 4 | 4 | 5 | 4.25 | 0 | 0 | 0 | 0 | 0 |
| TET1 | Methylcytosi | 0 | 0.00455878 | 7.4125E-05 | 6.2375E-05 | 2.4774E-05 | 3.7452E-05 | 4.9682E-05 | 0 | 0 | 0 | 0 | 0 | 6 | 5 | 2 | 3 | 4 | 0 | 0 | 0 | 0 | 0 |
| RPA2 | DNA-directec | 0 | 9.1242E-07 | 0.00011625 | 0.00014086 | 0.00011656 | 0.00011747 | 0.00012279 | 0 | 0 | 0 | 0 | 0 | 5 | 6 | 5 | 5 | 5.25 | 0 | 0 | 0 | 0 | 0 |
| AFF4 | AF4/FMR2 fa | 0 | 0.0007159 | 0.00013614 | 6.8736E-05 | 9.1001E-05 | 0.00013757 | 0.00010836 | 0 | 0 | 0 | 0 | 0 | 6 | 3 | 4 | 6 | 4.75 | 0 | 0 | 0 | 0 | 0 |
| MYO6 | Unconventio | 0 | 4.9221E-06 | 8.1572E-05 | 8.2369E-05 | 8.1789E-05 | 6.1821E-05 | 7.6888E-05 | 0 | 0 | 0 | 0 | 0 | 4 | 4 | 4 | 3 | 3.75 | 0 | 0 | 0 | 0 | 0 |
| PAXI | Paxillin OS=H | 0 | 0.00705428 | 8.9301E-05 | 0.00022544 | 8.9539E-05 | 0.0002256 | 0.00015747 | 0 | 0 | 0 | 0 | 0 | 2 | 5 | 2 | 5 | 3.5 | 0 | 0 | 0 | 0 | 0 |
| PGAM1 | Phosphoglyc | 0 | 0.00129577 | 0.00020778 | 0.00031472 | 0.00020834 | 0.00041993 | 0.00028769 | 0 | 0 | 0 | 0 | 0 | 2 | 3 | 2 | 4 | 2.75 | 0 | 0 | 0 | 0 | 0 |
| PSME3 | Proteasome a | 0 | 1.7813E-06 | 0.00051946 | 0.00062944 | 0.00052084 | 0.0006299 | 0.00057491 | 0 | 0 | 0 | 0 | 0 | 5 | 6 | 5 | 6 | 5.5 | 0 | 0 | 0 | 0 | 0 |
| PTN1 | Tyrosine-prot | 0 | 9.1342E-07 | 0.00030332 | 0.00036754 | 0.00030412 | 0.0003065 | 0.00032037 | 0 | 0 | 0 | 0 | 0 | 5 | 6 | 5 | 5 | 5.25 | 0 | 0 | 0 | 0 | 0 |
| UBP38 | Ubiquitin car | 0 | 0.00043471 | 0.0001013 | 0.00010229 | 5.0784E-05 | 0.00010236 | 8.9184E-05 | 0 | 0 | 0 | 0 | 0 | 4 | 4 | 2 | 4 | 3.5 | 0 | 0 | 0 | 0 | 0 |
| S23IP | SEC23-intera | 0 | 0.00013387 | 5.2777E-05 | 5.3293E-05 | 7.9376E-05 | 7.9997E-05 | 6.6361E-05 | 0 | 0 | 0 | 0 | 0 | 2 | 2 | 3 | 3 | 2.5 | 0 | 0 | 0 | 0 | 0 |
| CNDD3 | Condensin-2 | 0 | 7.2051E-05 | 5.2847E-05 | 0.00008894 | 7.0651E-05 | 7.1203E-05 | 7.091E-05 | 0 | 0 | 0 | 0 | 0 | 3 | 5 | 4 | 4 | 4 | 0 | 0 | 0 | 0 | 0 |
| SRRT | Serrate RNA | 0 | 0.00232852 | 0.00015062 | 6.0837E-05 | 0.00015102 | 9.1321E-05 | 0.00011345 | 0 | 0 | 0 | 0 | 0 | 5 | 2 | 5 | 3 | 3.75 | 0 | 0 | 0 | 0 | 0 |
| THOC3 | THO complex | 0 | 0.00115217 | 0.00030072 | 0.00022775 | 0.00015076 | 0.00015194 | 0.00020779 | 0 | 0 | 0 | 0 | 0 | 4 | 3 | 2 | 2 | 2.75 | 0 | 0 | 0 | 0 | 0 |
| CDK7 | Cyclin-depen | 0 | 2.0242E-14 | 0.00030507 | 0.00030805 | 0.00030588 | 0.00030827 | 0.00030682 | 0 | 0 | 0 | 0 | 0 | 4 | 4 | 4 | 4 | 4 | 0 | 0 | 0 | 0 | 0 |
| CSR2B | Cysteine-rich | 0 | 0.00662717 | 0.00010123 | 0.00010222 | 0.00020301 | 6.8198E-05 | 0.00011866 | 0 | 0 | 0 | 0 | 0 | 3 | 3 | 6 | 2 | 3.5 | 0 | 0 | 0 | 0 | 0 |
| ZKSC4 | Zinc finger pr | 0 | 0.00911544 | 0.00019368 | 0.00034225 | 9.7096E-05 | 0.00039142 | 0.00025611 | 0 | 0 | 0 | 0 | 0 | 4 | 7 | 2 | 8 | 5.25 | 0 | 0 | 0 | 0 | 0 |
| SRSF2 | Serine/argini | 0 | 0.00214784 | 0.00035821 | 0.00060286 | 0.00035917 | 0.00024132 | 0.00039039 | 0 | 0 | 0 | 0 | 0 | 3 | 5 | 3 | 2 | 3.25 | 0 | 0 | 0 | 0 | 0 |
| PESC | Pescadillo ho | 0 | 0.00157903 | 0.00022439 | 0.00018127 | 8.9996E-05 | 0.00013605 | 0.00015793 | 0 | 0 | 0 | 0 | 0 | 5 | 4 | 2 | 3 | 3.5 | 0 | 0 | 0 | 0 | 0 |
| GOGA5 | Golgin subfar | 0 | 2.0043E-07 | 0.00025269 | 0.00025516 | 0.00021717 | 0.00025535 | 0.00024509 | 0 | 0 | 0 | 0 | 0 | 7 | 7 | 6 | 7 | 6.75 | 0 | 0 | 0 | 0 | 0 |
| ANK3 | Ankyrin-3 OS | 0 | 0.00027501 | 4.2202E-05 | 4.2615E-05 | 4.8359E-05 | 2.4369E-05 | 3.9386E-05 | 0 | 0 | 0 | 0 | 0 | 7 | 7 | 8 | 4 | 6.5 | 0 | 0 | 0 | 0 | 0 |
| SMCA1 | Probable glol | 0 | 0.00044876 | 0.00012518 | 0.00010113 | 7.5309E-05 | 0.0001518 | 0.00011335 | 0 | 0 | 0 | 0 | 0 | 23 | 11 | 15 | 19 | 17 | 0 | 0 | 0 | 0 | 0 |
| ZBTB9 | Zinc finger ar | 0 | 0.00012961 | 0.00044632 | 0.00028168 | 0.00033563 | 0.00028188 | 0.00033638 | 0 | 0 | 0 | 0 | 0 | 8 | 5 | 6 | 5 | 6 | 0 | 0 | 0 | 0 | 0 |
| ARI1B | AT-rich intera | 0 | 0.02642378 | 3.5405E-05 | 8.3419E-05 | 2.3666E-05 | 2.3851E-05 | 4.1585E-05 | 0 | 0 | 0 | 0 | 0 | 6 | 8 | 4 | 5 | 5.75 | 0 | 0 | 0 | 0 | 0 |
| SF3B4 | Splicing facto | 0 | 0.00255893 | 0.00024895 | 0.00050276 | 0.00024961 | 0.00025156 | 0.00031322 | 0 | 0 | 0 | 0 | 0 | 4 | 8 | 4 | 4 | 5 | 0 | 0 | 0 | 0 | 0 |
| JCAD | Junctional pr | 0 | 0.00923354 | 0.00011651 | 5.8822E-05 | 3.8938E-05 | 0.00013735 | 8.7905E-05 | 0 | 0 | 0 | 0 | 0 | 6 | 3 | 2 | 7 | 4.5 | 0 | 0 | 0 | 0 | 0 |
| SSF1 | Suppressor o | 0 | 0.0004644 | 0.00022316 | 0.00011267 | 0.00022375 | 0.00016913 | 0.00018218 | 0 | 0 | 0 | 0 | 0 | 4 | 2 | 4 | 3 | 3.25 | 0 | 0 | 0 | 0 | 0 |
| WBP11 | WW domain- | 0 | 0.02643977 | 8.2335E-05 | 0.00029099 | 8.2554E-05 | 0.0001248 | 0.00014517 | 0 | 0 | 0 | 0 | 0 | 2 | 7 | 2 | 3 | 3.5 | 0 | 0 | 0 | 0 | 0 |
| SYF1 | Pre-mRNA-sp | 0 | 0.0061511 | 0.00018518 | 0.00018699 | 9.2837E-05 | 6.2376E-05 | 0.00013185 | 0 | 0 | 0 | 0 | 0 | 6 | 6 | 3 | 2 | 4.25 | 0 | 0 | 0 | 0 | 0 |
| CHD3 | Chromodoma | 0 | 2.027E-14 | 3.9583E-05 | 0.00003997 | 3.9688E-05 | 3.9998E-05 | 3.981E-05 | 0 | 0 | 0 | 0 | 0 | 18 | 16 | 17 | 20 | 17.75 | 0 | 0 | 0 | 0 | 0 |
| TBL1X | F-box-like/W | 0 | 0.00025123 | 0.0001372 | 0.00023091 | 0.00013757 | 0.00018486 | 0.00017264 | 0 | 0 | 0 | 0 | 0 | 15 | 13 | 12 | 15 | 13.75 | 0 | 0 | 0 | 0 | 0 |
| RBM34 | RNA-binding | 0 | 0.00125205 | 0.00012274 | 0.00024787 | 0.0001846 | 0.00012403 | 0.00016981 | 0 | 0 | 0 | 0 | 0 | 2 | 4 | 3 | 2 | 2.75 | 0 | 0 | 0 | 0 | 0 |
| TBCB | Tubulin-foldi | 0 | 9.3852E-05 | 0.00032445 | 0.00021841 | 0.00021687 | 0.00021857 | 0.00024458 | 0 | 0 | 0 | 0 | 0 | 3 | 2 | 2 | 2 | 2.25 | 0 | 0 | 0 | 0 | 0 |
| TMF1 | TATA elemen | 0 | 0.04016569 | 4.8286E-05 | 4.8758E-05 | 0.00016945 | 4.8793E-05 | 7.8822E-05 | 0 | 0 | 0 | 0 | 0 | 2 | 2 | 7 | 2 | 3.25 | 0 | 0 | 0 | 0 | 0 |
| DIAP2 | Protein diapł | 0 | 0.00136445 | 0.00014381 | 7.2606E-05 | 0.00019225 | 0.00014532 | 0.0001385 | 0 | 0 | 0 | 0 | 0 | 6 | 3 | 8 | 6 | 5.75 | 0 | 0 | 0 | 0 | 0 |
| TF2H4 | General trans | 0 | 0.00395023 | 0.00039982 | 0.00023071 | 0.00017181 | 0.00017315 | 0.00024387 | 0 | 0 | 0 | 0 | 0 | 7 | 4 | 3 | 3 | 4.25 | 0 | 0 | 0 | 0 | 0 |
| DHX8 | ATP-depende | 0 | 0.00049633 | 0.00006489 | 8.7366E-05 | 0.00008675 | 4.3714E-05 | 0.00007068 | 0 | 0 | 0 | 0 | 0 | 4 | 4 | 6 | 2 | 4 | 0 | 0 | 0 | 0 | 0 |
| AGFG1 | Arf-GAP dom | 0 | 0.00136914 | 0.00018782 | 0.00023707 | 9.4159E-05 | 0.00023724 | 0.00018907 | 0 | 0 | 0 | 0 | 0 | 4 | 5 | 2 | 5 | 4 | 0 | 0 | 0 | 0 | 0 |
| RRS1 | Ribosome bic | 0 | 0.0005148 | 0.00014459 | 0.00021901 | 0.00028996 | 0.00029223 | 0.00023645 | 0 | 0 | 0 | 0 | 0 | 2 | 3 | 4 | 4 | 3.25 | 0 | 0 | 0 | 0 | 0 |
| EPC2 | Enhancer of p | 0 | 0.01049554 | 0.0001962 | 6.6038E-05 | 0.00016393 | 6.6086E-05 | 0.00012306 | 0 | 0 | 0 | 0 | 0 | 6 | 2 | 5 | 2 | 3.75 | 0 | 0 | 0 | 0 | 0 |
| PLRG1 | Pleiotropic re | 0 | 0.00011407 | 0.00010268 | 0.00015552 | 0.00010295 | 0.00010376 | 0.00011623 | 0 | 0 | 0 | 0 | 0 | 2 | 3 | 2 | 2 | 2.25 | 0 | 0 | 0 | 0 | 0 |
| VIP2 | Inositol hexal | 0 | 0.00257907 | 6.3689E-05 | 6.4312E-05 | 6.3858E-05 | 0.00012872 | 8.0145E-05 | 0 | 0 | 0 | 0 | 0 | 3 | 3 | 3 | 6 | 3.75 | 0 | 0 | 0 | 0 | 0 |
| SNR40 | U5 small nuc | 0 | 0.00020985 | 0.00029567 | 0.00022392 | 0.00037057 | 0.00022408 | 0.00027856 | 0 | 0 | 0 | 0 | 0 | 4 | 3 | 5 | 3 | 3.75 | 0 | 0 | 0 | 0 | 0 |
| CCNK | Cyclin-K OS=I | 0 | 0.00200077 | 0.00018199 | 0.00018377 | 9.1237E-05 | 0.00009195 | 0.00013724 | 0 | 0 | 0 | 0 | 0 | 4 | 4 | 2 | 2 | 3 | 0 | 0 | 0 | 0 | 0 |
| BRD8 | Bromodomai | 0 | 6.4321E-05 | 8.5469E-05 | 0.00012946 | 0.00010712 | 8.6366E-05 | 0.0001021 | 0 | 0 | 0 | 0 | 0 | 4 | 6 | 5 | 4 | 4.75 | 0 | 0 | 0 | 0 | 0 |
| STRBP | Spermatid pe | 0 | 2.0121E-14 | 0.00019634 | 0.00019826 | 0.00019687 | 0.0001984 | 0.00019747 | 0 | 0 | 0 | 0 | 0 | 6 | 5 | 7 | 7 | 6.25 | 0 | 0 | 0 | 0 | 0 |
| THOC6 | THO complex | 0 | 0.00010244 | 0.00038693 | 0.00023443 | 0.00038796 | 0.00031279 | 0.00033053 | 0 | 0 | 0 | 0 | 0 | 5 | 3 | 5 | 4 | 4.25 | 0 | 0 | 0 | 0 | 0 |
| CATIN | Cactin OS=Hc | 0 | 0.00261028 | 0.00013925 | 0.00021092 | 6.9812E-05 | 0.00017589 | 0.00014897 | 0 | 0 | 0 | 0 | 0 | 4 | 6 | 2 | 5 | 4.25 | 0 | 0 | 0 | 0 | 0 |
| ESS2 | Splicing facto | 0 | 0.01367065 | 0.00011088 | 0.00011196 | 0.00033351 | 0.00016806 | 0.0001811 | 0 | 0 | 0 | 0 | 0 | 2 | 2 | 6 | 3 | 3.25 | 0 | 0 | 0 | 0 | 0 |
| BRD7 | Bromodomai | 0 | 0.00190566 | 0.00016214 | 8.1863E-05 | 0.00016257 | 8.1922E-05 | 0.00012212 | 0 | 0 | 0 | 0 | 0 | 4 | 2 | 4 | 2 | 3 | 0 | 0 | 0 | 0 | 0 |
| EZRI | Ezrin OS=Hor | 0 | 0.00012797 | 0.00013509 | 0.00013642 | 9.0303E-05 | 9.1009E-05 | 0.00011321 | 0 | 0 | 0 | 0 | 0 | 3 | 3 | 2 | 2 | 2.5 | 0 | 0 | 0 | 0 | 0 |
| SGF29 | SAGA-associa | 0 | 0.00011409 | 0.00018013 | 0.00027283 | 0.00018061 | 0.00018202 | 0.0002039 | 0 | 0 | 0 | 0 | 0 | 2 | 3 | 2 | 2 | 2.25 | 0 | 0 | 0 | 0 | 0 |
| RPF2 | Ribosome pro | 0 | 0.00049904 | 0.00034495 | 0.00034832 | 0.00017293 | 0.00026143 | 0.00028191 | 0 | 0 | 0 | 0 | 0 | 4 | 4 | 2 | 3 | 3.25 | 0 | 0 | 0 | 0 | 0 |
| PPIL2 | Peptidyl-prol | 0 | 3.6948E-05 | 0.00010149 | 0.00015373 | 0.00015265 | 0.00015384 | 0.00014043 | 0 | 0 | 0 | 0 | 0 | 2 | 3 | 3 | 3 | 2.75 | 0 | 0 | 0 | 0 | 0 |
| RBMX2 | RNA-binding | 0 | 0.00011416 | 0.0001639 | 0.00024826 | 0.00016434 | 0.00016562 | 0.00018553 | 0 | 0 | 0 | 0 | 0 | 2 | 3 | 2 | 2 | 2.25 | 0 | 0 | 0 | 0 | 0 |
| GPTC1 | G patch dom | 0 | 1.1713E-05 | 8.5033E-05 | 8.5864E-05 | 0.00011368 | 8.5926E-05 | 9.2626E-05 | 0 | 0 | 0 | 0 | 0 | 3 | 3 | 4 | 3 | 3.25 | 0 | 0 | 0 | 0 | 0 |
| DDX10 | Probable ATF | 0 | 6.0377E-05 | 0.00012063 | 9.1359E-05 | 0.00015119 | 0.0001219 | 0.00012127 | 0 | 0 | 0 | 0 | 0 | 4 | 3 | 5 | 4 | 4 | 0 | 0 | 0 | 0 | 0 |
| MAT1 | CDK-activatir | 0 | 0.00153237 | 0.0003416 | 0.00068988 | 0.00034251 | 0.00043148 | 0.00045137 | 0 | 0 | 0 | 0 | 0 | 4 | 8 | 4 | 5 | 5.25 | 0 | 0 | 0 | 0 | 0 |
| RFA2 | Replication p | 0 | 0.0012889 | 0.00019547 | 0.00039476 | 0.00019599 | 0.00029628 | 0.00027063 | 0 | 0 | 0 | 0 | 0 | 2 | 4 | 2 | 3 | 2.75 | 0 | 0 | 0 | 0 | 0 |
| WDR26 | WD repeat-c | 0 | 0.00024249 | 0.00011977 | 0.00012094 | 0.00016011 | 0.00020171 | 0.00015063 | 0 | 0 | 0 | 0 | 0 | 3 | 3 | 4 | 5 | 3.75 | 0 | 0 | 0 | 0 | 0 |
| CARL1 | F-actin-uncap | 0 | 0.00531532 | 3.8495E-05 | 5.8307E-05 | 9.6494E-05 | 0.0000389 | 5.8049E-05 | 0 | 0 | 0 | 0 | 0 | 2 | 3 | 5 | 2 | 3 | 0 | 0 | 0 | 0 | 0 |
| EGLN1 | Egl nine hom | 0 | 0.00104987 | 0.00012389 | 0.0002502 | 0.00024844 | 0.00031298 | 0.00023388 | 0 | 0 | 0 | 0 | 0 | 2 | 4 | 4 | 5 | 3.75 | 0 | 0 | 0 | 0 | 0 |
| WRN | Werner synd | 0 | 0.00011045 | 9.2138E-05 | 5.5824E-05 | 7.3907E-05 | 9.3106E-05 | 7.8744E-05 | 0 | 0 | 0 | 0 | 0 | 5 | 3 | 4 | 5 | 4.25 | 0 | 0 | 0 | 0 | 0 |
| NSRP1 | Nuclear spec | 0 | 2.0405E-14 | 0.00014187 | 0.00014326 | 0.00014225 | 0.00014336 | 0.00014269 | 0 | 0 | 0 | 0 | 0 | 3 | 3 | 3 | 3 | 3 | 0 | 0 | 0 | 0 | 0 |
| INADL | InaD-like pro | 0 | 0.00473403 | 5.8608E-05 | 2.9591E-05 | 4.4073E-05 | 8.8836E-05 | 5.5277E-05 | 0 | 0 | 0 | 0 | 0 | 4 | 2 | 3 | 6 | 3.75 | 0 | 0 | 0 | 0 | 0 |
| SUZ12 | Polycomb pro | 0 | 3.1021E-05 | 0.00010712 | 0.00010817 | 0.00010741 | 7.2167E-05 | 9.8717E-05 | 0 | 0 | 0 | 0 | 0 | 3 | 3 | 3 | 2 | 2.75 | 0 | 0 | 0 | 0 | 0 |
| BRCC3 | Lys-63-specif | 0 | 0.00481835 | 0.00041754 | 0.0003373 | 0.00058611 | 0.00016877 | 0.00037743 | 0 | 0 | 0 | 0 | 0 | 5 | 4 | 7 | 2 | 4.5 | 0 | 0 | 0 | 0 | 0 |
| EMSA1 | ELM2 and SA | 0 | 0.00219502 | 5.0504E-05 | 7.6497E-05 | 7.5958E-05 | 0.00012759 | 8.2637E-05 | 0 | 0 | 0 | 0 | 0 | 2 | 3 | 3 | 5 | 3.25 | 0 | 0 | 0 | 0 | 0 |
| NCOA2 | Nuclear rece | 0 | 0.00029882 | 0.0000721 | 5.4603E-05 | 5.4219E-05 | 3.6428E-05 | 5.4338E-05 | 0 | 0 | 0 | 0 | 0 | 4 | 3 | 3 | 2 | 3 | 0 | 0 | 0 | 0 | 0 |
| XPO6 | Exportin-6 OS | 0 | 0.00124881 | 0.00011728 | 9.4743E-05 | 0.00011759 | 4.7406E-05 | 9.4255E-05 | 0 | 0 | 0 | 0 | 0 | 5 | 4 | 5 | 2 | 4 | 0 | 0 | 0 | 0 | 0 |
| NELFE | Negative elor | 0 | 4.3255E-06 | 0.00034722 | 0.00028049 | 0.00027851 | 0.00035086 | 0.00031427 | 0 | 0 | 0 | 0 | 0 | 5 | 4 | 4 | 5 | 4.5 | 0 | 0 | 0 | 0 | 0 |
| HTSF1 | HIV Tat-speci | 0 | 0.00172274 | 6.9903E-05 | 0.00017647 | 0.00010513 | 0.00014127 | 0.00012319 | 0 | 0 | 0 | 0 | 0 | 2 | 5 | 3 | 4 | 3.5 | 0 | 0 | 0 | 0 | 0 |
| EZH2 | Histone-lysin | 0 | 2.2714E-05 | 0.00010612 | 0.00014288 | 0.0001064 | 0.00014298 | 0.0001246 | 0 | 0 | 0 | 0 | 0 | 3 | 4 | 3 | 4 | 3.5 | 0 | 0 | 0 | 0 | 0 |
| CIC | Protein capic | 0 | 0.00255923 | 3.2821E-05 | 6.6285E-05 | 3.2909E-05 | 3.3166E-05 | 4.1295E-05 | 0 | 0 | 0 | 0 | 0 | 2 | 4 | 2 | 2 | 2.5 | 0 | 0 | 0 | 0 | 0 |
| CCD47 | Coiled-coil dc | 0 | 0.00011567 | 0.00010927 | 0.00011034 | 0.00010956 | 0.00016562 | 0.0001237 | 0 | 0 | 0 | 0 | 0 | 2 | 2 | 2 | 3 | 2.25 | 0 | 0 | 0 | 0 | 0 |
| CDC37 | Hsp90 co-cha | 0 | 9.9069E-05 | 0.00013962 | 0.00014099 | 0.00020999 | 0.00014109 | 0.00015792 | 0 | 0 | 0 | 0 | 0 | 2 | 2 | 3 | 2 | 2.25 | 0 | 0 | 0 | 0 | 0 |
| SFR15 | Splicing facto | 0 | 6.5276E-05 | 9.2026E-05 | 9.2926E-05 | 0.00011534 | 0.00013949 | 0.00010995 | 0 | 0 | 0 | 0 | 0 | 4 | 4 | 5 | 6 | 4.75 | 0 | 0 | 0 | 0 | 0 |
| AF17 | Protein AF-17 | 0 | 1.4617E-05 | 7.2429E-05 | 7.3138E-05 | 7.2622E-05 | 9.7587E-05 | 7.8944E-05 | 0 | 0 | 0 | 0 | 0 | 3 | 3 | 3 | 4 | 3.25 | 0 | 0 | 0 | 0 | 0 |
| SCAM1 | Secretory car | 0 | 1.9826E-05 | 0.00023422 | 0.00023651 | 0.00031312 | 0.00031557 | 0.00027486 | 0 | 0 | 0 | 0 | 0 | 3 | 3 | 4 | 4 | 3.5 | 0 | 0 | 0 | 0 | 0 |
| HP1B3 | Heterochrom | 0 | 0.00031102 | 0.00019087 | 0.00014456 | 9.5691E-05 | 0.00014466 | 0.00014395 | 0 | 0 | 0 | 0 | 0 | 4 | 3 | 2 | 3 | 3 | 0 | 0 | 0 | 0 | 0 |
| ZC3HD | Zinc finger CC | 0 | 0.0047202 | 6.3282E-05 | 9.5851E-05 | 3.1725E-05 | 0.00004796 | 5.9705E-05 | 0 | 0 | 0 | 0 | 0 | 4 | 6 | 2 | 3 | 3.75 | 0 | 0 | 0 | 0 | 0 |
| NC2B | Protein Dr1 C | 0 | 0.0026385 | 0.00059974 | 0.0003028 | 0.000902 | 0.00060604 | 0.00060265 | 0 | 0 | 0 | 0 | 0 | 4 | 2 | 6 | 4 | 4 | 0 | 0 | 0 | 0 | 0 |
| ALG5 | Dolichyl-phos | 0 | 2.0049E-14 | 0.00016289 | 0.00016448 | 0.00016333 | 0.0001646 | 0.00016383 | 0 | 0 | 0 | 0 | 0 | 2 | 2 | 2 | 2 | 2 | 0 | 0 | 0 | 0 | 0 |
| SCFD1 | Sec1 family d | 0 | 3.6938E-05 | 8.2207E-05 | 0.00012452 | 0.00012364 | 0.00012461 | 0.00011374 | 0 | 0 | 0 | 0 | 0 | 2 | 3 | 3 | 3 | 2.75 | 0 | 0 | 0 | 0 | 0 |
| ATG3 | Ubiquitin-like | 0 | 1.1708E-05 | 0.00025212 | 0.00025458 | 0.00033705 | 0.00025477 | 0.00027463 | 0 | 0 | 0 | 0 | 0 | 3 | 3 | 4 | 3 | 3.25 | 0 | 0 | 0 | 0 | 0 |
| ZN592 | Zinc finger pr | 0 | 3.1426E-05 | 6.2482E-05 | 4.2062E-05 | 6.2649E-05 | 6.3139E-05 | 5.7583E-05 | 0 | 0 | 0 | 0 | 0 | 3 | 2 | 3 | 3 | 2.75 | 0 | 0 | 0 | 0 | 0 |
| MO4L1 | Mortality fac | 0 | 7.2745E-05 | 0.00021869 | 0.00029444 | 0.00029236 | 0.00036831 | 0.00029345 | 0 | 0 | 0 | 0 | 0 | 3 | 4 | 4 | 5 | 4 | 0 | 0 | 0 | 0 | 0 |
| HNRL1 | Heterogeneo | 0 | 0.00013387 | 6.1655E-05 | 6.2258E-05 | 9.2729E-05 | 9.3454E-05 | 7.7524E-05 | 0 | 0 | 0 | 0 | 0 | 2 | 2 | 3 | 3 | 2.5 | 0 | 0 | 0 | 0 | 0 |
| GMEB2 | Glucocorticoi | 0 | 0.00172223 | 9.9579E-05 | 0.00025138 | 0.00014977 | 0.00020125 | 0.00017549 | 0 | 0 | 0 | 0 | 0 | 2 | 5 | 3 | 4 | 3.5 | 0 | 0 | 0 | 0 | 0 |
| BUD13 | BUD13 homc | 0 | 0.00214378 | 8.5261E-05 | 0.00017219 | 8.5488E-05 | 0.00017231 | 0.00012881 | 0 | 0 | 0 | 0 | 0 | 2 | 4 | 2 | 4 | 3 | 0 | 0 | 0 | 0 | 0 |
| PEX19 | Peroxisomal | 0 | 0.00560092 | 0.00026477 | 0.00017824 | 0.00017698 | 0.00044591 | 0.00026648 | 0 | 0 | 0 | 0 | 0 | 3 | 2 | 2 | 5 | 3 | 0 | 0 | 0 | 0 | 0 |
| UBP22 | Ubiquitin car | 0 | 3.5407E-05 | 0.00015079 | 0.00015227 | 0.00010079 | 0.00015237 | 0.00013906 | 0 | 0 | 0 | 0 | 0 | 3 | 3 | 2 | 3 | 2.75 | 0 | 0 | 0 | 0 | 0 |
| SPAG7 | Sperm-associ | 0 | 0.00040741 | 0.00046499 | 0.00046954 | 0.00046623 | 0.00023494 | 0.00040893 | 0 | 0 | 0 | 0 | 0 | 4 | 4 | 4 | 2 | 3.5 | 0 | 0 | 0 | 0 | 0 |
| PUR4 | Phosphoribos | 0 | 1.6032E-05 | 7.8889E-05 | 5.9745E-05 | 7.9099E-05 | 5.9788E-05 | 6.938E-05 | 0 | 0 | 0 | 0 | 0 | 4 | 3 | 4 | 3 | 3.5 | 0 | 0 | 0 | 0 | 0 |
| VAPB | Vesicle-assoc | 0 | 0.00129572 | 0.00021719 | 0.00032897 | 0.00021777 | 0.00043894 | 0.00030072 | 0 | 0 | 0 | 0 | 0 | 2 | 3 | 2 | 4 | 2.75 | 0 | 0 | 0 | 0 | 0 |
| NDC1 | Nucleoporin | 0 | 1.0829E-05 | 0.00015661 | 0.0001186 | 0.00011777 | 0.00011869 | 0.00012792 | 0 | 0 | 0 | 0 | 0 | 4 | 3 | 3 | 3 | 3.25 | 0 | 0 | 0 | 0 | 0 |
| P4R3B | Serine/threo | 0 | 0.00218027 | 6.2164E-05 | 0.00015693 | 9.3494E-05 | 9.4225E-05 | 0.0001017 | 0 | 0 | 0 | 0 | 0 | 2 | 5 | 3 | 3 | 3.25 | 0 | 0 | 0 | 0 | 0 |
| RPAB3 | DNA-directec | 0 | 2.0336E-14 | 0.00035185 | 0.00035529 | 0.00035278 | 0.00035554 | 0.00035387 | 0 | 0 | 0 | 0 | 0 | 2 | 2 | 2 | 2 | 2 | 0 | 0 | 0 | 0 | 0 |
| NIPA | Nuclear-inter | 0 | 3.5401E-05 | 0.0001577 | 0.00015924 | 0.00010541 | 0.00015936 | 0.00014543 | 0 | 0 | 0 | 0 | 0 | 3 | 3 | 2 | 3 | 2.75 | 0 | 0 | 0 | 0 | 0 |

| | | | | | | | | | | | | | | | | | | | | | | | |
|---|---|---|---|---|---|---|---|---|---|---|---|---|---|---|---|---|---|---|---|---|---|---|---|
| RCL1 | RNA 3'-termi | 0 | 0.00030739 | 0.00021224 | 0.00021431 | 0.00028374 | 0.00014298 | 0.00021332 | 0 | 0 | 0 | 0 | 0 | 3 | 3 | 4 | 2 | 3 | 0 | 0 | 0 | 0 | 0 |
| LMO7 | LIM domain o | 0 | 0.00012797 | 4.7038E-05 | 4.7498E-05 | 3.1442E-05 | 3.1688E-05 | 3.9417E-05 | 0 | 0 | 0 | 0 | 0 | 3 | 3 | 2 | 2 | 2.5 | 0 | 0 | 0 | 0 | 0 |
| KCY | UMP-CMP ki | 0 | 0.00045259 | 0.0004039 | 0.00067976 | 0.00067497 | 0.00040815 | 0.0005417 | 0 | 0 | 0 | 0 | 0 | 3 | 5 | 5 | 3 | 4 | 0 | 0 | 0 | 0 | 0 |
| ANS1A | Ankyrin repe | 0 | 0.00196108 | 0.00011635 | 7.0493E-05 | 6.9997E-05 | 4.7029E-05 | 7.5967E-05 | 0 | 0 | 0 | 0 | 0 | 5 | 3 | 3 | 2 | 3.25 | 0 | 0 | 0 | 0 | 0 |
| TF3C2 | General trans | 0 | 0.00133753 | 5.7933E-05 | 0.00014625 | 0.00014522 | 0.00011708 | 0.00011662 | 0 | 0 | 0 | 0 | 0 | 2 | 5 | 5 | 4 | 4 | 0 | 0 | 0 | 0 | 0 |
| PF21A | PHD finger pr | 0 | 0.0001326 | 7.7613E-05 | 0.00011756 | 0.00011673 | 7.8428E-05 | 9.7583E-05 | 0 | 0 | 0 | 0 | 0 | 2 | 3 | 3 | 2 | 2.5 | 0 | 0 | 0 | 0 | 0 |
| CWC27 | Peptidyl-prol | 0 | 0.0011875 | 0.00011182 | 0.00016936 | 0.00022423 | 0.00011299 | 0.0001546 | 0 | 0 | 0 | 0 | 0 | 2 | 3 | 4 | 2 | 2.75 | 0 | 0 | 0 | 0 | 0 |
| KIN17 | DNA/RNA-bir | 0 | 2.0567E-14 | 0.00013429 | 0.00013561 | 0.00013465 | 0.0001357 | 0.00013506 | 0 | 0 | 0 | 0 | 0 | 2 | 2 | 2 | 2 | 2 | 0 | 0 | 0 | 0 | 0 |
| SIX5 | Homeobox p | 0 | 0.00046433 | 0.00014283 | 7.2115E-05 | 0.00014321 | 0.00010825 | 0.0001166 | 0 | 0 | 0 | 0 | 0 | 4 | 2 | 4 | 3 | 3.25 | 0 | 0 | 0 | 0 | 0 |
| BRD9 | Bromodomai | 0 | 0.00014686 | 8.8403E-05 | 0.0001339 | 8.8639E-05 | 0.000134 | 0.00011124 | 0 | 0 | 0 | 0 | 0 | 2 | 3 | 2 | 3 | 2.5 | 0 | 0 | 0 | 0 | 0 |
| ARI3A | AT-rich intera | 0 | 0.000128 | 0.0001335 | 0.00013481 | 8.9237E-05 | 8.9935E-05 | 0.00011187 | 0 | 0 | 0 | 0 | 0 | 4 | 4 | 3 | 3 | 3.5 | 0 | 0 | 0 | 0 | 0 |
| GAB1 | GRB2-associa | 0 | 0.00012923 | 0.00011407 | 7.6791E-05 | 0.00007625 | 0.00011527 | 9.5595E-05 | 0 | 0 | 0 | 0 | 0 | 3 | 2 | 2 | 3 | 2.5 | 0 | 0 | 0 | 0 | 0 |
| STXB4 | Syntaxin-binc | 0 | 3.1041E-05 | 0.00014316 | 0.00014456 | 0.00014354 | 0.00009644 | 0.00013193 | 0 | 0 | 0 | 0 | 0 | 3 | 3 | 3 | 2 | 2.75 | 0 | 0 | 0 | 0 | 0 |
| 3MG | DNA-3-methy | 0 | 0.00012797 | 0.00026566 | 0.00026825 | 0.00017758 | 0.00017896 | 0.00022261 | 0 | 0 | 0 | 0 | 0 | 3 | 3 | 2 | 2 | 2.5 | 0 | 0 | 0 | 0 | 0 |
| MRGBP | MRG/MORF4 | 0 | 9.9083E-05 | 0.00025871 | 0.00026124 | 0.0003891 | 0.00026143 | 0.00029262 | 0 | 0 | 0 | 0 | 0 | 2 | 2 | 3 | 2 | 2.25 | 0 | 0 | 0 | 0 | 0 |
| PUS7 | Pseudouridyl | 0 | 2.0396E-14 | 7.9844E-05 | 8.0625E-05 | 8.0057E-05 | 8.0683E-05 | 8.0302E-05 | 0 | 0 | 0 | 0 | 0 | 2 | 2 | 2 | 2 | 2 | 0 | 0 | 0 | 0 | 0 |
| INT14 | Integrator co | 0 | 0.00011403 | 0.00010189 | 0.00015432 | 0.00010216 | 0.00010296 | 0.00011533 | 0 | 0 | 0 | 0 | 0 | 2 | 3 | 2 | 2 | 2.25 | 0 | 0 | 0 | 0 | 0 |
| RBM42 | RNA-binding | 0 | 0.00115209 | 0.0002199 | 0.00016654 | 0.00011024 | 0.00011111 | 0.00015195 | 0 | 0 | 0 | 0 | 0 | 4 | 3 | 2 | 2 | 2.75 | 0 | 0 | 0 | 0 | 0 |
| NR2C2 | Nuclear rece | 0 | 2.0356E-14 | 8.8552E-05 | 8.9418E-05 | 8.8788E-05 | 8.9482E-05 | 0.00008906 | 0 | 0 | 0 | 0 | 0 | 2 | 2 | 2 | 2 | 2 | 0 | 0 | 0 | 0 | 0 |
| RT29 | 28S ribosoma | 0 | 3.1434E-05 | 0.00019891 | 0.0001339 | 0.00019944 | 0.000201 | 0.00018331 | 0 | 0 | 0 | 0 | 0 | 3 | 2 | 3 | 3 | 2.75 | 0 | 0 | 0 | 0 | 0 |
